# Genome-wide association study in European patients with congenital heart disease identifies risk loci for transposition of the great arteries and anomalies of the thoracic arteries and veins and expression of discovered candidate genes in the developing heart

**DOI:** 10.1101/2020.06.19.161067

**Authors:** Harald Lahm, Meiwen Jia, Martina Dreßen, Felix Wirth, Nazan Puluca, Ralf Gilsbach, Bernard D. Keavney, Julie Cleuziou, Nicole Beck, Olga Bondareva, Elda Dzilic, Melchior Burri, Karl C. König, Johannes A. Ziegelmüller, Claudia Abou-Ajram, Irina Neb, Zhong Zhang, Stefanie A. Doppler, Elisa Mastantuono, Peter Lichtner, Gertrud Eckstein, Jürgen Hörer, Peter Ewert, James R. Priest, Lutz Hein, Rüdiger Lange, Thomas Meitinger, Heather J. Cordell, Bertram Müller-Myhsok, Markus Krane

## Abstract

**Rationale:** Genetic factors undoubtedly contribute to the development of congenital heart disease (CHD), but still remain mostly ill-defined.

**Objective:** Identification of genetic risk factors associated with CHD and functional analysis of SNP-carrying genes.

**Methods and Results:** Genetic association study of 1,440 Caucasian CHD patients from the German Heart Center Munich collected from March 2009 to June 2016, 2,594 patients of previous studies provided by the Newcastle University and 8,486 controls underwent meta-analysis to detect single nucleotide polymorphisms (SNPs) associated with CHD.

**Results:** 4,034 Caucasian CHD patients strictly classified according to the Society of Thoracic Surgeons nomenclature and 8,486 controls were included. One SNP on chromosome 5 reached genome-wide significance across all CHD phenotypes (rs185531658,OR:2.16, *p*=5.28×10^−9^) and was also indicative for septal defects (OR:2.16, *p*=6.15×10^−8^). One region on chromosome 20 pointing to the *MACROD2* locus, identified four SNPs (rs150246290,OR:3.78, *p*=1.27×10^−10^; rs149890280,OR:3.74, *p*=1.8×10^−10^; rs149467721,OR:3.53; *p*=1.39×10^−9^, rs77094733,OR:3.53, *p*=1.73×10^−9^) in patients with transposition of the great arteries (TGA). A second region was detected on chromosome 8 located at *ZBTB10* (rs148563140,OR:3.42, *p*=3.28×10^−8^; rs143638934,OR:3.42, *p*=3.51×10^−8^) in the same subgroup. Three highly significant risk variants on chromosome 17 (rs76774446,OR:1.60, *p*=9.95×10^−8^; rs11874,OR:1.60, *p*=6.64×10^−8^; rs17677363,OR:1.60, *p*=9.81×10^−8^) within the *GOSR2* locus were identified in patients with anomalies of thoracic arteries and veins (ATAV). Genetic variants associated with ATAV are suggested to influence expression of *WNT3*, and variant rs870142 related to septal defects is proposed to influence expression of *MSX1*. Cardiac differentiation of human and murine induced pluripotent stem cells and single cell RNAseq analyses of developing murine and human hearts show essential functional roles for *MACROD2, GOSR2, WNT3* and *MSX1* at all developmental stages.

**Conclusions:** For the first time genetic risk factors in CHD patients with TGA and ATAV were identified. Several candidate genes play an essential functional role in heart development at the embryonic, newborn and adult stage.

## Introduction

Congenital heart disease (CHD) accounts for approximately 28% of all congenital anomalies worldwide ^1^ with a frequency of CHD of 9.1 per 1,000 live births. ^2^ Currently, CHD represents a major global health challenge, causing more than 200,000 deaths world-wide per year. ^3^

While major progress has been made in the field of genetics during the last few decades, the exact etiologic origins of CHD still remain only partially understood. Causal genes have been identified in uncommon syndromic forms, such as *TBX5* for Holt-Oram syndrome. ^4^ CHD may also be associated with major chromosomal syndromes, ^5^ *de novo* mutations, ^6^ aneuploidy, and copy number variants. ^7-9^ Each of these genetic abnormalities are associated with roughly 10% of CHDs, while the majority of cases seem to represent a complex multifactorial disease with unknown etiology. ^9^ An increasing number of candidate genes have been implicated, which harbor low- and intermediate-effect variants. ^10^ Genetic studies in mice and humans strongly support the idea that certain variants are inherited and cause a pronounced pathology. ^11^, ^12^

Several genome-wide association studies (GWAS) have previously been conducted to determine potential genetic risk factors for CHD. ^13-17^ For atrial septal defects (ASDs) 4p16 was identified as a risk locus. ^17^, ^18^ For tetralogy of Fallot (TOF), regions of interest have been reported on chromosomes 1, 12 and 13. ^19^, ^20^ Agopian and colleagues have shown an association of a single intra-genetic single nucleotide polymorphism (SNP) with left ventricular obstructive defects. ^13^ For other major clinical subcategories, no risk loci have been identified to date.

We sought to identify genetic risk loci in CHD and clinical subpopulations thereof by GWAS due to the proven success of this approach. ^21^ We conducted a GWAS in more than 4,000 unrelated Caucasian patients diagnosed with CHD who were classified according to the standards and categories defined by the Society of Thoracic Surgeons (STS). ^22^, ^23^ We identified one risk variant for CHD in general and detected an association of single or clustered SNPs in five major subpopulations. We determined risk loci in patients with transposition of the great arteries (TGA) and anomalies of the thoracic arteries and veins (ATAV). In addition, we demonstrate differential expression of candidate genes during differentiation of murine and human pluripotent stem cells and determine their expression in pediatric and adult aortic and atrial tissue. Finally, we document the functional role of candidate genes by single cell RNA sequencing (scRNAseq) analyses in the developing murine and human heart *in vivo*.

## Methods

The complete cohort of CHD patients comprised 4,034 subjects. The first cohort of 1,440 patients (769 males, 671 females, mean age 17 y) was collected at the German Heart Center Munich between March 2009 and June 2016. The German ethnicity of the participants was confirmed by analysis of the genotype data using multidimensional scaling. In addition, two previously analyzed patient collectives with mixed CHD history (mean age 20 y) ^17^ and TOF (mean age 15 y) ^19^, comprising 2,594 patients (1,320 males, 1,274 females), were included. Patients in whom neurodevelopmental or genetic abnormalities were apparent were excluded, but since some probands were recruited as babies/young children, this would not have been evident in all cases. Genotypes were compared to 3,554 (1,726 males, 1,828 females) and 4,932 (2,498 males, 2,434 females) controls for the German and British cohorts, respectively. The German controls were recruited from the well-established KORA (Cooperative Health Research in the Region of Augsburg) F4 and S3 cohorts used in numerous studies as a control group. ^24^ Ethics approval for the study was obtained from the local ethical review boards, and informed written consent was obtained from participants, parents, or legal guardians. Genotyping was performed at the Helmholtz Zentrum (Munich, Germany) and the Centre National de Genotypage (Evry Cedex, France) using the Affymetrix Axiom Genome-Wide Human array or the Illumina 660wQUAD array, respectively. The German samples were genotyped on the Affymetrix Axiom CEU array according to the Axiom GT best practice protocol according to the manufacturer’s recommendation. The KORA controls were genotyped on the same chip type by Affymetrix. A detailed description of Methods is available in the Data Supplement.

## Results

### Association analysis in overall population of CHD patients and subgroups defined by STS classification

We performed a GWAS analysis in 4,034 CHD cases (2,089 males, 1,945 females) and 8,486 controls (4,224 males, 4,262 females) to detect possible candidate SNPs. The first group consisted of 1,440 patients collected in the German Heart Center Munich. Two further groups implicating 2,594 patients have previously been published. ^17^, ^19^ To obtain clearly defined clinical subgroups of patients, we classified all CHD patients according to the STS Congenital Heart Surgery Database (CHSD) recommendations. This classification was established under the leadership of the International Society for Nomenclature of Pediatric and Congenital Heart Disease as a clinical data registry but also reflects common developmental etiologies and is therefore a well accepted tool for research on CHD. ^22^, ^23^ The distribution of the subgroups is shown in Table 1.

**Table 1.**
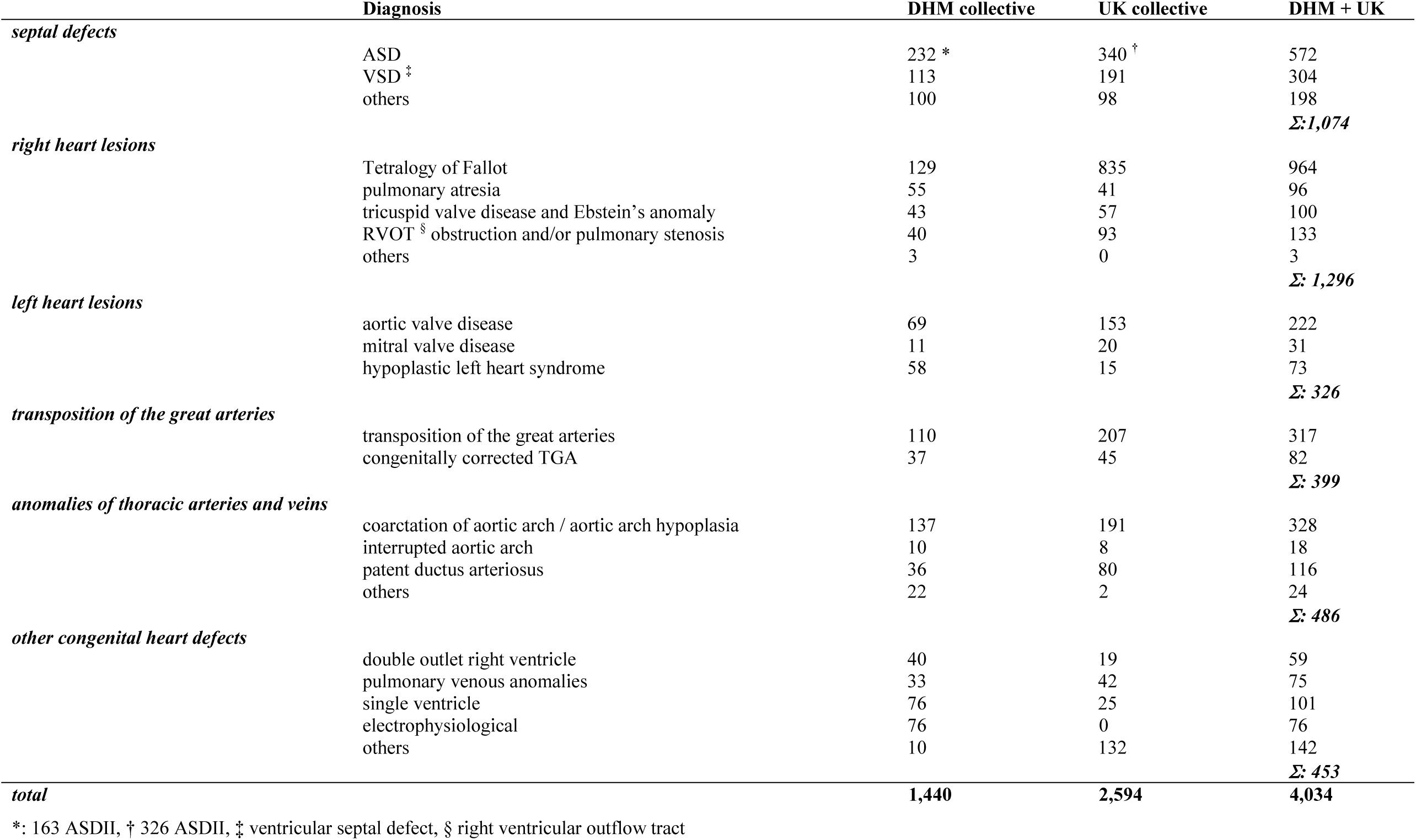
Patient collective

|  | Diagnosis | DHM collective | UK collective | DHM + UK |
| --- | --- | --- | --- | --- |
| *septal defects* |  |  |  |  |
|  | ASD | 232 * | 340 † | 572 |
|  | VSD ‡ | 113 | 191 | 304 |
|  | others | 100 | 98 | 198 |
|  |  |  |  | **Σ: 1,074** |
| *right heart lesions* |  |  |  |  |
|  | Tetralogy of Fallot | 129 | 835 | 964 |
|  | pulmonary atresia | 55 | 41 | 96 |
|  | tricuspid valve disease and Ebstein's anomaly | 43 | 57 | 100 |
|  | RVOT § obstruction and/or pulmonary stenosis | 40 | 93 | 133 |
|  | others | 3 | 0 | 3 |
|  |  |  |  | **Σ: 1,296** |
| *left heart lesions* |  |  |  |  |
|  | aortic valve disease | 69 | 153 | 222 |
|  | mitral valve disease | 11 | 20 | 31 |
|  | hypoplastic left heart syndrome | 58 | 15 | 73 |
|  |  |  |  | **Σ: 326** |
| *transposition of the great arteries* |  |  |  |  |
|  | transposition of the great arteries | 110 | 207 | 317 |
|  | congenitally corrected TGA | 37 | 45 | 82 |
|  |  |  |  | **Σ: 399** |
| *anomalies of thoracic arteries and veins* |  |  |  |  |
|  | coarctation of aortic arch / aortic arch hypoplasia | 137 | 191 | 328 |
|  | interrupted aortic arch | 10 | 8 | 18 |
|  | patent ductus arteriosus | 36 | 80 | 116 |
|  | others | 22 | 2 | 24 |
|  |  |  |  | **Σ: 486** |
| *other congenital heart defects* |  |  |  |  |
|  | double outlet right ventricle | 40 | 19 | 59 |
|  | pulmonary venous anomalies | 33 | 42 | 75 |
|  | single ventricle | 76 | 25 | 101 |
|  | electrophysiological | 76 | 0 | 76 |
|  | others | 10 | 132 | 142 |
|  |  |  |  | **Σ: 453** |
| **total** |  | **1,440** | **2,594** | **4,034** |

\*: 163 ASDII, † 326 ASDII, ‡ ventricular septal defect, § right ventricular outflow tract

We first performed an analysis across all 4,034 CHD patients and identified one SNP on chromosome 5 with genome-wide significance (rs185531658) (Figure 1).

**Figure 1.**
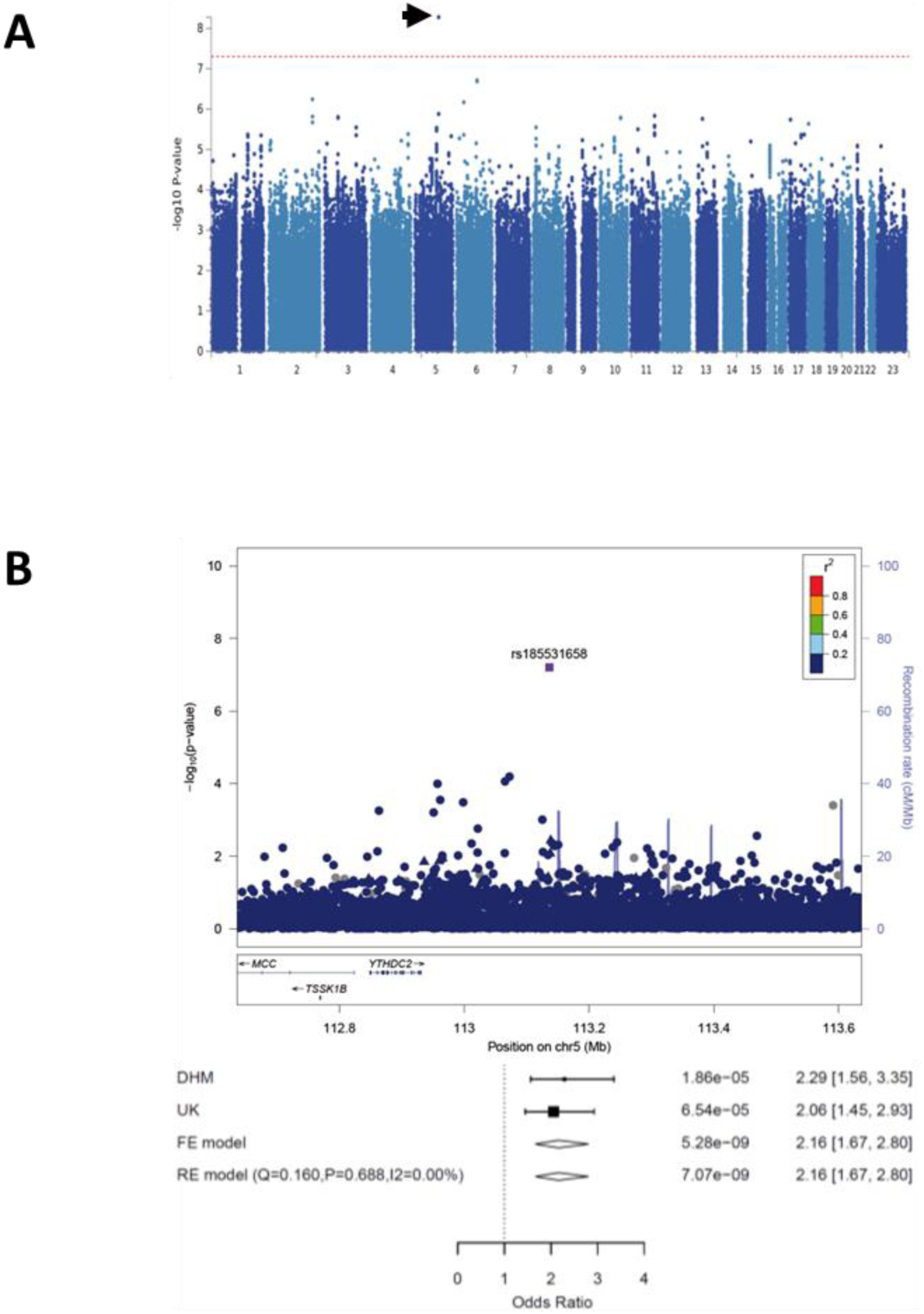
Identification of SNPs with genome-wide significance across the entire CHD collective. **A:** Manhattan plot. **B:** LocusZoom plot of the genomic region of rs185531658 on chromosome 5. The index SNP is indicated as a purple diamond. The forest plot shows the significance of the SNP and the odds ratios of both collectives separately and together. Circles represent imputed SNPs, triangles genotyped SNPs. FE: fixed effects, RE: random effects

Subsequently, we examined five diagnostic subgroups in our cohort: TGA (n=399), right heart lesions (n=1,296), left heart lesions (n=326), septal defects (n=1,074) and ATAV (n=486). In the TGA subgroup, SNPs on chromosomes 20 and 8 were identified. The lead SNP (rs150246290) and three variants on chromosome 20, all with genome-wide significance, mapped to the *MACROD2* gene (Figure 2A) implicated in chromosomal instability ^25^ and transcriptional regulation. ^26^ Two SNPs (rs149890280, rs150246290) are suggested to be possible causal variants (Table I in the Data Supplement). The identified risk locus on chromosome 8 close to *ZBTB10* included two SNPs (rs148563140, rs143638934), both with genome-wide significance (Figure 2B). Given the high levels of linkage disequilibrium between these SNPs, they are indicative of the same association signal in both loci. Unexpectedly, two risk variants at 12q24 and 13q32, previously shown to be associated with TOF ^19^ could not be substantiated in the German cohort (Figure IA and IB and Table II in the Data Supplement). A single SNP (rs146300195) on chromosome 5 at the *SLC27A6* locus with genome-wide significance was evident in this subgroup (Figure IC in the Data Supplement). In left heart lesions, three variants (rs3547121, chromosome 2 and rs114503684, rs2046060, chromosome 3), reached genome-wide significance (Figure II in the Data Supplement). The same SNP on chromosome 5 (rs185531658), indicative for the whole CHD population, also appeared in the subpopulation of septal defects with almost genome-wide significance (Figure IIIA in the Data Supplement). A second SNP (rs138741144) was evident on chromosome 17 within the *ASIC2* locus (Figure IIIB in the Data Supplement). Restricting the analysis to ASD, we confirmed the previously reported significance of the lead SNP (rs870142) and multiple variants on chromosome 4p16 ^17^ (Figure IV in the Data Supplement). Limiting ASD patients to those diagnosed with ASD type II (ASDII) (n=489) we identified two SNPs (rs145619574 and rs72917381) on chromosome 18, in the vicinity of *WDR7*, and another variant (rs187369228) on chromosome 3, located close to *LEPREL1* (= *P3H2*) (Figure VA and VB in the Data Supplement). In patients with ATAV three SNPs were apparent on chromosome 17 with sub-genome-wide significance (rs17677363, rs11874, and rs76774446), all located within the *GOSR2* locus (Figure 2C). All three variants are predicted to be possibly causal (Table I in the Data Supplement). In addition, GeneHancer analyses suggest that rs11874 may affect expression of *GOSR2* and *WNT3* may be a topolocigally associated region (Table III in the Data Supplement). One additional SNP mapped to chromosome 6 (rs117527287) without a nearby gene (the closest was *TBX18*, approximately 0.3 Mb apart) (Figure VI in the Data Supplement). Table 2 summarizes all detected SNPs and their significances. Genes located within the LD region of each locus are provided in Table IV in the Data Supplement.

**Table 2.** List of highly significant SNPs in CHD

|  | chromosomal location | gene | *p* value | OR <sup>†</sup> | MAF DHM * |  | MAF UK * |  |
| --- | --- | --- | --- | --- | --- | --- | --- | --- |
|  |  |  |  |  | cases | control | cases | control |
| *all CHD* |  |  |  |  |  |  |  |  |
| rs185531658 | NC_000005.9:g.113136521T>C | none | 5.28x10<sup>-9</sup> | 2.16 (1.67-2.80) | 0.020 | 0.011 | 0.015 | 0.008 |
| *TGA* |  |  |  |  |  |  |  |  |
| **rs150246290** | NC_000020.11:g.15132234G>C | MACROD2, intron | 1.27x10<sup>-10</sup> | 3.78 (2.51-5.64) | 0.054 | 0.014 | 0.027 | 0.012 |
| rs149890280 | NC_000020.11:g.15126433A>G | MACROD2, intron | 1.8x10<sup>-10</sup> | 3.74 (2.48-5.64) | 0.054 | 0.014 | 0.027 | 0.012 |
| rs149467721 | NC_000020.11:g.15178710G>T | MACROD2, intron | 1.39x10<sup>-9</sup> | 3.53 (2.34-5.31) | 0.053 | 0.016 | 0.026 | 0.010 |
| rs77094733 | NC_000020.11:g.15184689G>C | MACROD2, intron | 1.73x10<sup>-9</sup> | 3.53 (2.34-5.31) | 0.053 | 0.016 | 0.026 | 0.010 |
| **rs148563140** | NC_000008.10:g.81475406C>T | none | 3.28x10<sup>-8</sup> | 3.42 (2.20-5.26) | 0.020 | 0.010 | 0.037 | 0.010 |
| rs143638934 | NC_000008.10:g.81467030A>G | none | 3.51x10<sup>-8</sup> | 3.42 (1.08-5.26) | 0.020 | 0.010 | 0.037 | 0.010 |
| *right heart lesions* |  |  |  |  |  |  |  |  |
| rs146300195 | NC_000005.10:g.128991152G>A | SLC27A6, intron | 1.01x10<sup>-8</sup> | 3.60 (2.32-5.53) | 0.011 | 0.005 | 0.014 | 0.005 |
| *left heart lesions* |  |  |  |  |  |  |  |  |
| **rs35437121** | NC_000002.12:g.131011875C>T | ARHGEF4, intron | 4.31x10<sup>-8</sup> | 2.27 (1.68-3.03) | 0.075 | 0.049 | 0.100 | 0.047 |
| **rs114503684** | NC_000003.12:g.142116127C>G | TFDP2, intron | 5.1x10<sup>-8</sup> | 3.53 (2.25-5.58) | 0.028 | 0.013 | 0.042 | 0.014 |
| **rs2046060** | NC_000003.11:g.187852486A>G | none | 7.14x10<sup>-8</sup> | 1.57 (1.34-1.86) | 0.404 | 0.300 | 0.393 | 0.297 |
| *anomalies of thoracic arteries and veins* |  |  |  |  |  |  |  |  |
| rs76774446 | NC_000017.11:g.46969002C>A | GOSR2, intron | 9.95x10<sup>-8</sup> | 1.60 (1.35-1.92) | 0.156 | 0.115 | 0.203 | 0.135 |
| rs17677363 | NC_000017.11:g.46958746A>T | GOSR2, intron | 9.81x10<sup>-8</sup> | 1.60 (1.35-1.92) | 0.156 | 0.115 | 0.203 | 0.135 |
| **rs11874** | NC_000017.11:g.46939827G>A | GOSR2, intron variant,<br>utr variant 3' | 6.64x10<sup>-8</sup> | 1.60 (1.35-1.92) | 0.160 | 0.115 | 0.203 | 0.136 |
| rs117527287 | NC_000006.11:g.85729959G>A | none | 6.22x10<sup>-9</sup> | 3.63 (2.34-5.58) | 0.020 | 0.010 | 0.032 | 0.009 |
| *septal defects* |  |  |  |  |  |  |  |  |
| rs185531658 | NC_000005.9:g.113136521T>C | none | 6.15x10<sup>-8</sup> | 2.16 (1.67-3.90) | 0.023 | 0.011 | 0.019 | 0.008 |
| rs138741144 | NC_000017.11:g.33959545G>A | ASIC2, LOC107985038,<br>intron | 7.34x10<sup>-8</sup> | 2.46 (1.77-3.42) | 0.034 | 0.014 | 0.019 | 0.010 |
| *ASD* |  |  |  |  |  |  |  |  |
| **rs870142** | NC_000004.12:g.4646320C>T | STX18-AS1, intron | 4.30x10<sup>-7</sup> | 1.40 (1.23-1.60) | 0.283 | 0.238 | 0.312 | 0.227 |
| *ASDII* |  |  |  |  |  |  |  |  |
| rs187369228 | NC_000003.12:g.190084650A>G | P3H2 (=LEPREL1),<br>intron | 1.74x10<sup>-8</sup> | 2.97 (2.03-4.35) | 0.024 | 0.015 | 0.041 | 0.016 |
| rs145619574 | NC_000018.10:g.56833471A>T | WDR7, intron | 2.56x10<sup>-8</sup> | 6.11 (3.25-11.59) | 0.040 | 0.008 | 0.005 | 0.008 |
| **rs72917381** | NC_000018.10:g.56878992C>T | WDR7, intron | 2.35x10<sup>-8</sup> | 5.93 (3.19-11.13) | 0.042 | 0.009 | 0.007 | 0.009 |

\* minor allele frequency of the German (DHM) or English (UK) collective. Lead SNPs are indicated in **bold**, † odds ratio (95% confidence interval in parenthesis).

**Figure 2.**
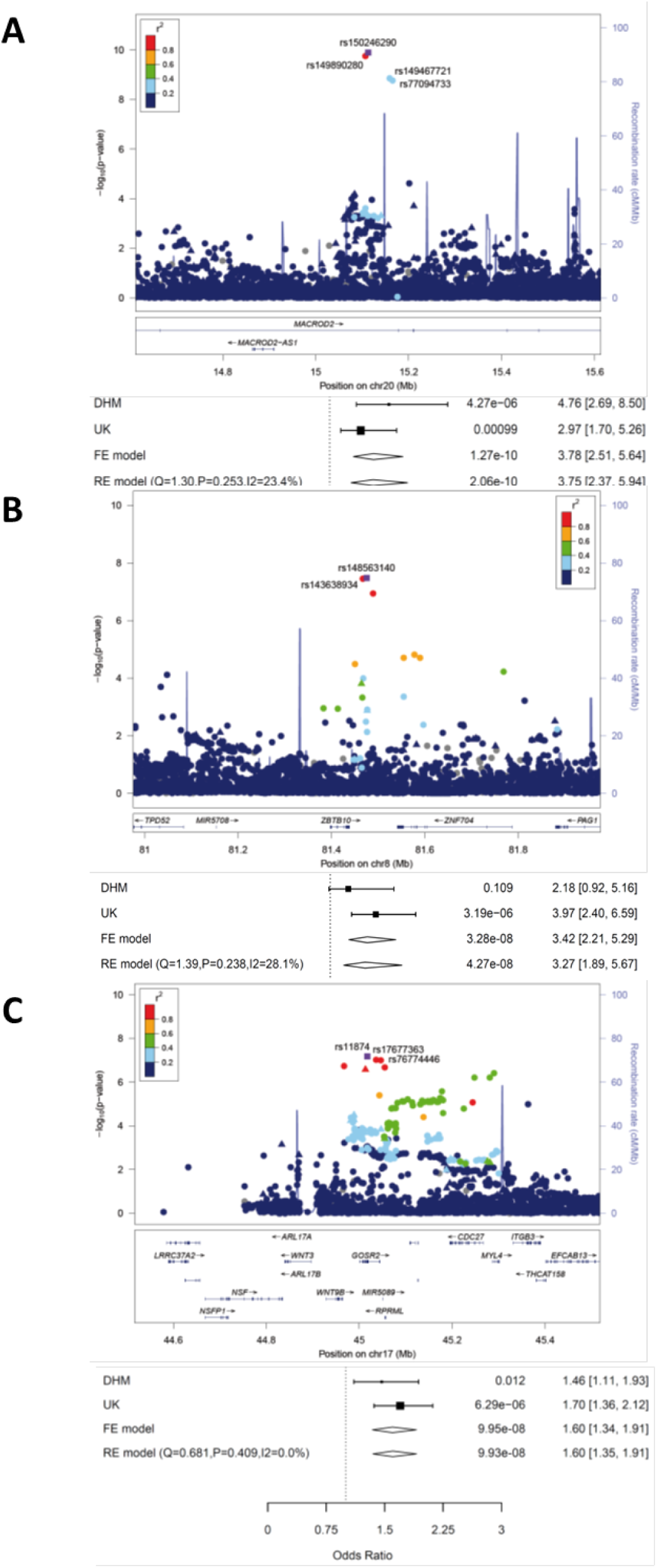
SNPs associated with transposition of the great arteries **(A, B)** or anomalies of thoracic arteries and veins **(C). A:** LocusZoom plot of the *MACROD2* region on chromosome 20. **B:** LocusZoom plot of the *ZBTB10* region on chromosome 8. **C:** LocusZoom plot of the *GOSR2* region on chromosome 17. The index SNPs are indicated as purple diamonds and the other SNPs are color coded depending on their degree of correlation (r^2^). Circles represent imputed SNPs, triangles genotyped SNPs. FE: fixed effects, RE: random effects.

Genes where SNPs with genome-wide significance, listed in Table 2 and further variants significantly enriched with *p* values <0.0005 (Table V in the Data Supplement), fall into the gene region, underwent a gene set enrichment analysis (GSEA). Terms related to cell-cell signaling, embryonic development and morphogenesis showed the highest significance (Table VI in the Data Supplement) and well-known cardiac transcription factors *GATA3, GATA4*, and *WNT9B* were involved in all signaling cascades (Figure VII in the Data Supplement).

### Expression of SNP-carrying candidate genes during cardiac differentiation of murine embryonic stem cells

We addressed the question whether SNP-carrying might be expressed by multipotent GFP-positive cardiac progenitor cells (CPCs) during differentiation of embryonic stem cells (ESCs) (Figure 3A) derived from the Nkx2.5 cardiac enhancer (CE) eGFP transgenic mouse line. ^27^ Interestingly, *Macrod2* and *Gosr2* were significantly enriched in beating GFP-positive CPCs compared to GFP-negative stage-matched counterparts, in contrast to *Wnt3* and *Msx1* (Figure 3B).

**Figure 3.**
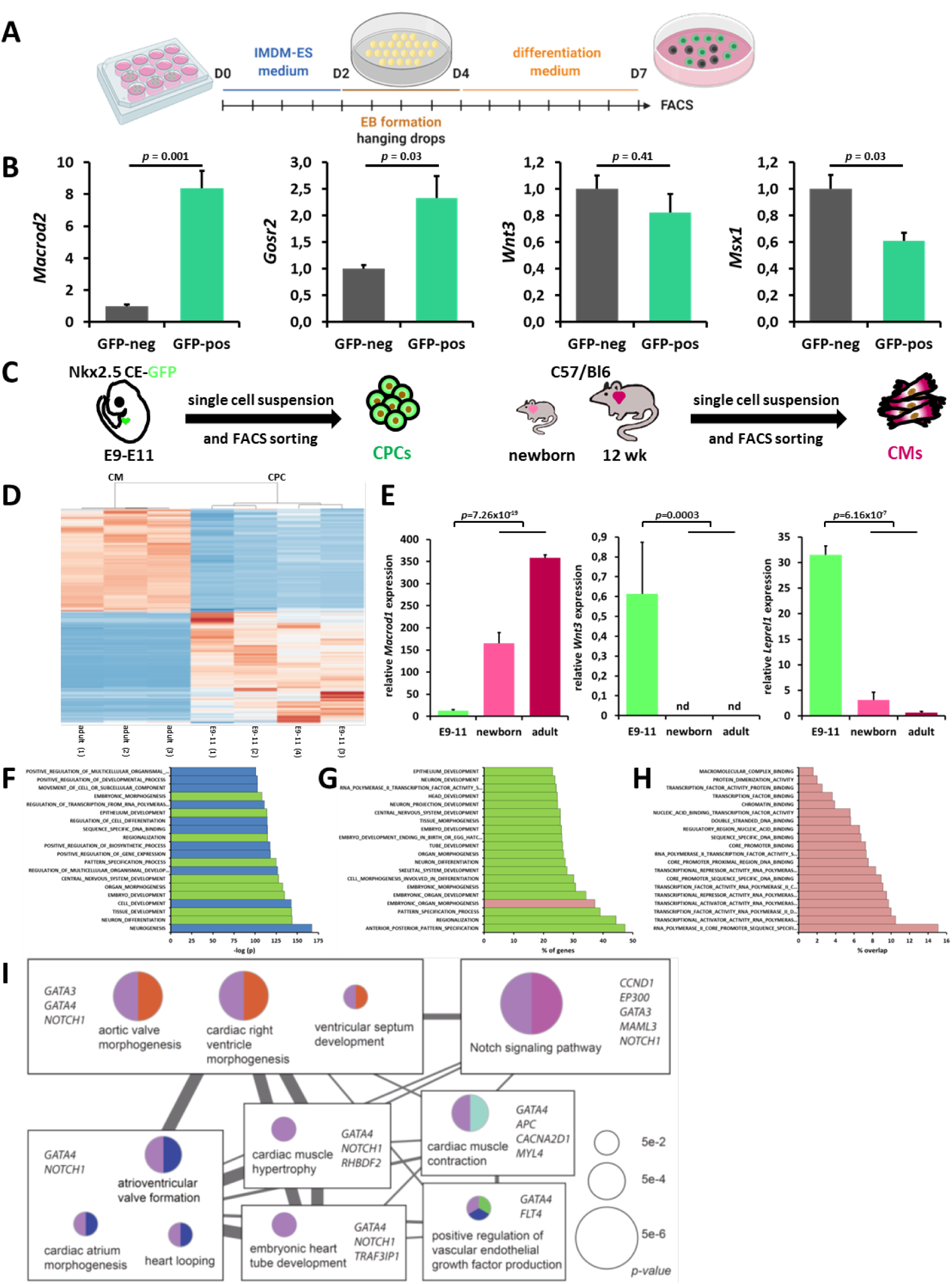
Role of SNP-carrying candidate genes in murine cardiac development. **A:** Schedule of differentiation of murine ESCs. **B:** Relative gene expression of *Macrod2, Gosr2, Wnt3* and *Msx1* in GFP neg cells and GFP pos CPCs. **C:** Schematic representation of the enrichment of murine CPCs and postnatal CMs. **D:** Heatmap of genes differentially expressed in embryonic CPCs and adult CMs. **E:** Expression of *Macrod1, Wnt3* and *Leprel1* in CPCs (E9-11), newborn and adult CMs. **F - H:** Results of GSEA of 1,649 genes overlapping between CHD-associated SNP-carrying genes and genes up-regulated in CPCs according to **(F)** significance of GO terms, **(G)** coverage of GO terms and **(H)** second level GO terms showing molecular functions. **I:** Significantly enriched gene sets by ClueGO (Bonferroni-corrected *p* values < 0.1). Each circle represents a pathway/GO-term with each type of pie chart (identified by color and pattern) represents a functional group. The size of the circles represents raw *p* values of enrichment tests for GO terms. The width of the edges represents the degree of similarity between GO terms. Rectangles encompass the GO terms that share the same significant genes.

### Role of SNP-carrying genes in murine prenatal cardiac progenitors and cardiomyocytes in vivo

We then reanalyzed our RNAseq data from purified murine CPCs and postnatal cardiomyocytes (CMs) ^28^ (Figure 3C), clearly separated by their global expression patterns (Figure 3D). Both newborn and adult CMs expressed *Macrod1*, a paralog of *Macrod2*, at a much higher level than embryonic CPCs (Figure 3E). Furthermore, *Wnt3* and *Leprel1* were both abundantly expressed in CPCs but barely expressed or undetectable in CMs of newborn or adult mice (Figure 3E).

The global RNAseq analysis (Figure 3D, Supplementary Datafile1) identified 1,915 and 1,155 significantly up-regulated genes (>2-fold, *p*<0.05) specific for CPCs and CMs, respectively. We speculated that the gene loci of the SNPs identified in our CHD cohort might be associated with either of these two gene pools. Therefore, we compared the genes of the whole CHD cohort carrying SNPs with the gene lists up-regulated in CPCs or CMs. Applying MAGMA, we detected a clear enrichment of GWAS association signals in 1,649 genes up-regulated in CPCs (*p*=0.0078) (Supplementary Datafile1), but not in those up-regulated in CMs (*p*=0.471). After GSEA of these 1,649 genes, gene ontology (GO) terms related to neural development showed the highest significance, followed by pathways regulating tissue, cell, embryo and organ morphogenesis (Figure 3F). Investigation of the deposited GO gene set revealed a high coverage for embryonic and neural development (Figure 3G). Since “embryonic” gene sets comprise many genes in common, we selected embryonic organ morphogenesis to have a closer look on the molecular function in a second-level GO analysis. The top 20 categories all referred to DNA binding or transcription factor activity (Figure 3H). A network-based functional enrichment analysis highlights several pathways directly involved in cardiac development, such as ventricular septum development, aortic valve, right ventricle and atrium morphogenesis (Figure 3I).

### Location, timepoint, and cell-type specificity of candidate genes during mouse cardiac development at single-cell resolution

Using a contemporary computational approach ^29^ we re-analyzed a dataset of 1,901 cells derived from micro-dissection of embryonic mouse hearts spanning the critical period of E8.5 to E10.5 in cardiac morphogenesis ^30^ to examine the expression of *Macrod2, Gosr2, Zbtb10*, and *Wnt3*. Consistent with our survey of CPC differentiation, *Wnt3* was minimally detected within single cells during later stages of cardiac morphogenesis. *Gosr2* was moderately (*p*=1.2×10^−4^) and *Zbtb10* was strongly (*p*=9.2×10^−10^) expressed within mesenchymal cells throughout the developing heart with *Zbtb10* expressing very strong spatial localization to mesenchymal cells within the AV canal (*p*=3.6×10^−11^). By contrast, *Macrod2* expression was scattered at a low-level throughout the developing heart and displayed moderate cell-type specificity to ventricular cardiomyocytes from the interventricular septum (*p*=4.4×10^−5^).

### Expression of SNP-carrying candidate genes during cardiac differentiation of human induced pluripotent stem cells

We then investigated the role of all candidate genes during cardiac differentiation of human induced pluripotent stem (iPS) cells (Figure 4A). Expression of *MACROD2* gradually increased and peaked around day 10 while the expression of *GOSR2* did not substantially change at any time point (Figure 4B). ATACseq analyses suggest a potential interaction of *GOSR2* variants with *WNT3* and *STX18-AS1* variants with *MSX1*, respectively, early during cardiac differentiation of human ESCs. ^31^ In line with these results, both genes are most strongly upregulated on day 2 during differentiation of human iPS cells (Figure 4B). *STX18* and *LEPREL1* also peak early while expression of all other candidate genes was not substantially changed (Figure VIII in the Data Supplement).

**Figure 4.**
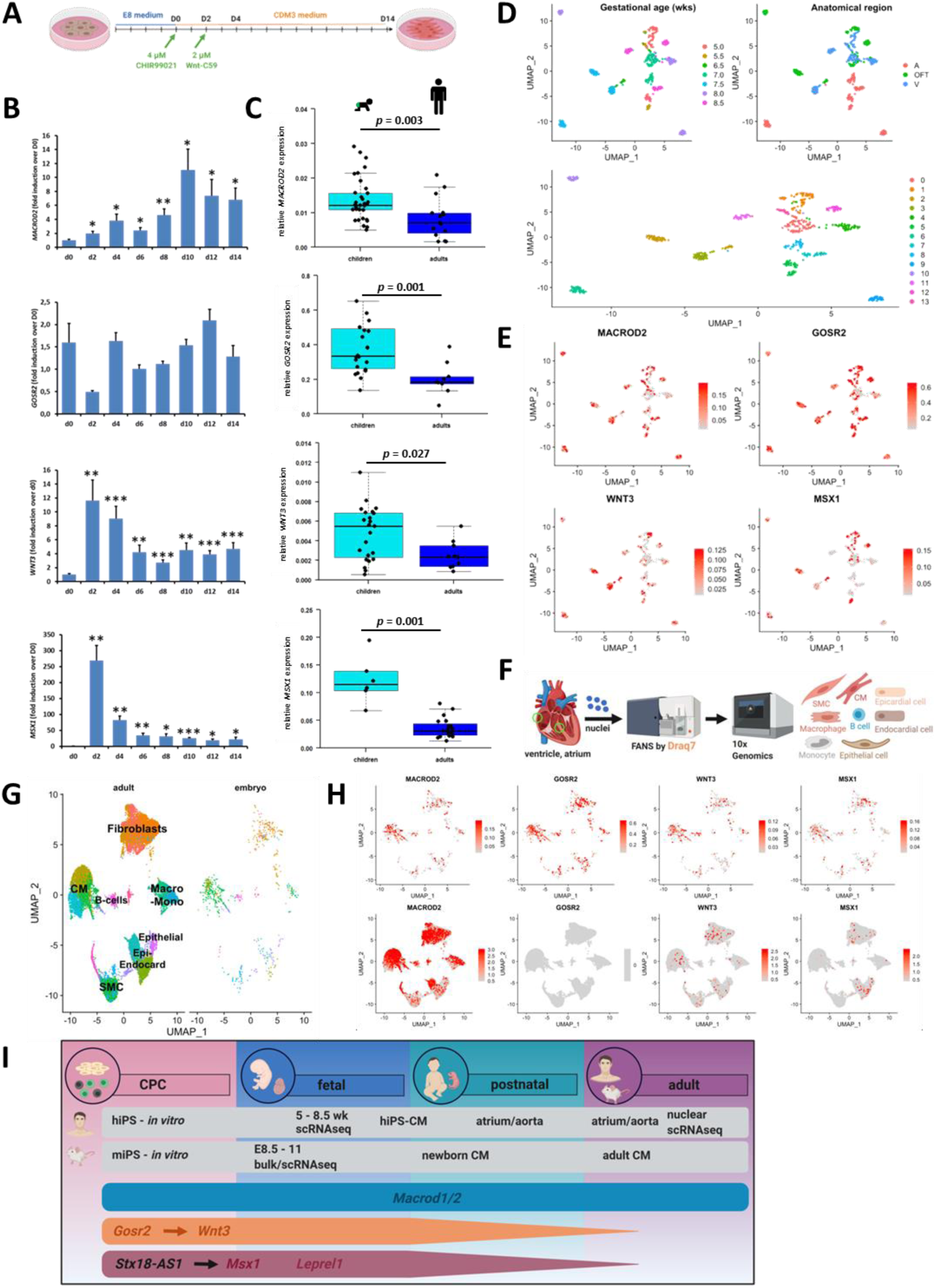
Role of SNP-carrying candidate genes in human cardiac development. **A:** Schedule of directed cardiac differentiation of human iPS cells. **B:** Expression of *MACROD2, GOSR2, WNT3* and *MSX1* during directed cardiac differentiation of human iPS cells. *: *p*<0.05, **: *p*<0.01, ***: *p*<0.001 vs. D0. **C:** Expression of *MACROD2, GOSR2, WNT3* and *MSX1* in aortic tissue of pediatric and adult surgical patients. **D:** Unbiased clustering of embryonic cells into biological entities. Cells are labeled based on age as well as anatomical localization for purposes of visualization. A: atria, OFT: outflow tract, V: ventricle. **E:** Relative expression of *MACROD2, GOSR2, WNT3* and *MSX1* in cells of embryonic heart. **F:** Schedule of single-cell RNAseq analysis of cell from atria and ventricles. **G:** Clustering of embryonic and adult cells and identification of cell types. **H:** Expression of candidate genes in the integrated dataset split by embryonic cells (upper panel) and adult cells (lower panel). **I:** Expression of candidate genes associated with TGA (turquoise), ATAV (orange) or septal defects (red) *in vitro* and *in vivo* during different stages of the developing murine and human heart.

### Expression of SNP-carrying candidate genes in tissue of CHD patients

We first analyzed whether the presence of the risk variant might influence expression of the affected gene. However, the genotype did not alter expression of *MACROD2, GOSR2* and *WNT3* (Figure IX in the Data Supplement). Therefore, we compared the expression of all candidate genes in aortic and atrial tissue of CHD patients (Table VII in the Data Supplement) with the expression in tissue of adult surgical patients (Table VIII in the Data Supplement). *MACROD2, GOSR2, WNT3* and *MSX1* were clearly expressed to a higher extent in tissues of CHD patients (Figure 4C). In addition, *ARHGEF4, STX18-AS1, STX18* and *WDR7* also showed a similar significantly higher expression in pediatric aortic tissue (Figure X in the Data Supplement). In atrial tissue expression of *SLC27A6, MSX1, LEPREL1* and *WDR7* was significantly higher in CHD samples (Figure XI in the Data Supplement). Thus, these data suggest that the majority of our candidate genes may rather have a role in early cardiac development.

### Expression of SNP-carrying candidate genes in human fetal and adult heart tissue

We extended our analysis and revisited a published scRNAseq dataset of 669 human embryonic cardiac cells. ^32^ Using principal component analysis and unsupervised clustering, we could classify cells into distinct biological entities, defined by their gestational age and anatomical region (Figure 4D). High expression among all 14 clusters was detected for *MACROD2* and especially for *GOSR2* with higher relative gene expression (Figure 4E). Expression of *WNT3* and *MSX1* appeared broad throughout all developmental stages, (Figure 4E), albeit more concentrated on fibroblasts and myocytes (Figure 4G). To pursue age-dependent differences in the expression of our candidate genes, we conducted additional scRNAseq experiments with 17,782 cells from samples of adult human atria and ventricles (Figure 4F). Integrating the data from adult and embryonic hearts, we could identify different cell types based on their expression of defined marker genes (Figure XII in the Data Supplement). Of note, cells from both adult and embryonic hearts yielded perfectly superimposable clusters (Figure 4G).

*MACROD2* shows robust expression in all adult cardiac cell types. By stark contrast, *GOSR2*, widely expressed throughout the embryonic heart, could not be detected in any adult cell (Figure 4H). *WNT3* and especially *MSX1* are expressed in cells of the adult heart, though at a much lower extent compared to embryonic cells given the much higher number of adult cells analyzed. While *WNT3* and *MSX1* show similar expression patterns in fetal and adult cell types, the expression of *MSX1* appears virtually absent in adult myocytes (Figure 4H). Thus, the four candidate genes analyzed may play a role in the developing human heart while *MACROD2* may still be important later on. Figure 4I summarizes the expression of candidate genes *in vitro* and at different stages of the developing murine and human heart *in vivo*.

## Discussion

We performed a GWAS on more than 4,000 Caucasian CHD patients which represents the largest genetic study of European individuals to date. We detected roughly 20 SNPs across five major clinical subgroups, associated with genome-wide significance (*p*<5×10^−8^).

A careful evaluation of the genes related to the identified SNPs showed no cardiac phenotype in monogenic knockout mouse models (Table IX in the Data Supplement) which is probably due to the multigenic etiology of almost all congenital heart malformations. Nevertheless, our downstream analyses of these SNPs within the subgroups of TGA, ATAV and ASD showed a clear functional association of the closely related genes during murine and human heart development using different *in vitro* and *in vivo* experimental strategies.

### TGA and MACROD2

In the TGA subgroup, four SNPs with genome-wide significance mapped to *MACROD2* which has been linked to adipogenesis and hypertension. ^26^, ^33^ Microdeletions in this gene have been implicated as a cause of chromosomal instability in cancers ^34^ and *de novo* deletion of exon 5 causes Kabuki syndrome. ^35^ Chromosomal imbalance is also frequently seen in CHD patients with different morphologies ^36-39^ including TGA ^40^ but so far the *MACROD2* locus was not associated with CHD.

Expression of *Macrod2* was significantly enhanced in murine CPCs. *Macrod1* was abundantly expressed in newborn and adult CMs, but negligibly in embryonic CPCs. *Macrod1* and *Macrod2* are paralogs with substantial structural similarity ^41^ and common biological activities, ^42^ potentially suggesting similar functions during cardiac development. scRNAseq data suggest *MARCOD2* expression during human embryonic development within ventricular and outflow tract cells (Figure 4E). We also found *MACROD2* expression in CMs which is in line with the later expression during directed cardiac differentiation of human iPS cells. Even more important for structural developmental defects is a high expression level of *MACROD2* during embryonic development in fibroblasts and endothelial cells (Figure 4 G and H upper panel). The *MACROD2* expression is not limited to the embryonic stage but shows high expression levels in different adult cardiac cell types (Figure 4 G and H lower panel).

Besides *MACROD2* two other highly significant SNPs, closely located to *ZBTB10*, have been associated with the TGA group. This gene interacts with telomeric regions of chromosomes. ^43^ Also this gene was previously not described to play a role in cardiogenesis, while our experimental data presented here suggests strong cell-type specificity in murine cardiac development for *Zbtb10*.

### ATAV and GOSR2

One risk region comprises three highly significant SNPs mapping to *GOSR2* which is involved in directed movement of macromolecules between Golgi compartments. ^44^ Genetic variants of *GOSR2* have been implicated in coronary artery disease ^45^ and myocardial infarction, with contradictory results. ^46^, ^47^ The ATAV subgroup includes patients diagnosed with coarctation of the aorta, an interrupted/hypoplastic aortic arch as well as patients with a patent ductus arteriosus. These CHD malformations all share a common origin within the aortic sac and the stepwise emerging aortic arches during embryonic development. ^48^ The proximal aorta and portions of the outflow tract derive from the bulbus cordis.

Applying ATACseq analysis Zhang et al. described a potential interaction between *GOSR2* and *WNT3* during cardiac differentiation of human ESC. ^31^ Our expression analysis showed significantly enhanced *Gosr2* expression in isolated murine CPCs, while *Wnt3* displayed similar expression in CPCs and developmentally stage-matched cells (Figure 3B) suggesting a specific role of *Gosr2* during embryonic cardiac development. Nevertheless, *Wnt3* was clearly detectable in embryonic CPCs but absent in newborn or adult CMs indicating a more distinct role for *Wnt3* during embryonic development. Furthermore, we could clearly show expression of *GOSR2* in human embryonic cells of the outflow tract (Figure 4E) by scRNAseq analysis, suggesting a potential association of this gene with the development of ATAV. In contrast, we could not detect *GOSR2* expression in the adult human heart, supporting our hypothesis that *GOSR2* exerts its biological role during embryonic cardiac development.

### ASD and STX18/MSX1

We identified SNP rs185531658 in patients with septal defects with high significance. The same SNP was also strongly associated with CHD risk in general, with *YTHDC2*, an RNA helicase involved in meiosis as the closest gene. ^49^ The second SNP for septal defects is related to *ASIC2*, whose loss leads to hypertension in null mice. ^50^ Restricting the patient cohort to ASD, we confirmed SNP rs870142, which we had previously identified. ^17^ As this SNP appeared with a much lower significance in the German cohort (Figure IV in the Data Supplement), its significance was lower compared to the original study (*p=*4.3×10^−7^ vs 2.6×10^−10^). Narrowing the cohort to ASDII patients, two risk loci were identified. The genes in the affected loci, *WDR7* and *LEPREL1*, are associated with growth regulation and tumor suppression of breast cancer ^51^, ^52^ but without cardiovascular importance. Lin and colleagues have published several risk loci for septal defects in a Chinese cohort. ^15^ We could validate one variant, rs490514, in our CHD population (Table X in the Data Supplement) supporting the validity of our GWAS results.

Zhang et al. also described a functional association between *STX18* (SNP rs870142) and *MSX1*. ^31^ This interaction is also supported by our findings of significantly higher expression levels of *STX18* and *MSX1* during cardiac differentiation of human iPS cells at early stages. Furthermore, expression of *Msx1* displayed comparable expression in CPCs and developmentally stage-matched cells suggesting a role of *Msx1* during embryonic development. The similar expression in GFP positive CPCs and GFP negative developmentally stage-matched cells could be explained either by an expression not exclusively restricted to embryonic cardiac development or a predominant expression of *Msx1* in second heart field (SHF) progenitors and cells of the outflow tract ^53^ which are not necessarily captured by our Nkx2.5 CE transgenic mouse model. ^27^

Even more important, extensive scRNAseq analyses showed a predominant expression of *MSX1* overlapping with cells of the outflow tract during embryonic human heart development with CMs and fibroblast as the main cell types at this stage. The role of *MSX1* in CMs seems to be restricted to embryonic development whereas we could still find expression of *MSX1* in fibroblasts end endothelial cells of the adult heart. This is in line with our comparative expression analysis of pediatric and adult aortic tissues (Figure 4C).

A second SNP, closely related to *LEPREL1* was associated with the subgroup of ASDII. *Leprel1* was clearly detectable in embryonic CPCs but barely evident in newborn or adult CMs. Furthermore, we could show a significantly elevated expression early during cardiac differentiation of human iPS cells suggesting a role during early cardiac development. Comparing the expression of *LEPREL1* in adult and pediatric atrial tissue we could show a significantly enhanced expression in pediatric samples, again suggesting a potential role during early cardiac development.

In summary, our GWAS identified multiple risk loci for all major clinical CHD subgroups. We detected genetic variants in the *MACROD2* and *GOSR2* loci, strongly associated with the phenotype of TGA and ATAV, respectively. The use of murine and human pluripotent stem cells and *ex vivo* results in tissue of CHD patients underline the functional role of several candidate genes during cardiac differentiation. Finally, scRNAseq analyses provide strong *in vivo* evidence that *MACROD2, GOSR2, WNT3* and *MSX1* play important roles during the embryonic development of the human heart.

## Acknowledgments

We gratefully acknowledge the support of Dr. Stefan Eichhorn for his help with biobank issues and Mrs. Elisabeth Zierler for her support with the genotyping of samples. The authors acknowledge the support of the Freiburg Galaxy Team: Dr. Mehmet Tekman and Prof. Rolf Backofen, Bioinformatics, University of Freiburg, Germany funded by Collaborative Research Centre 992 Medical Epigenetics (<u>DFG</u> grant SFB 992/1 2012) and German Federal Ministry of Education and Research (BMBF grant 031 A538A de.NBI-RBC). Parts of figure 3 and 4 were created with BioRender.com and exported under a paid subscription. Bertram Müller-Myhsok and Markus Krane had full access to all the data in the study and take responsibility for the integrity of the data and the accuracy of the data analysis.

## Sources of Funding

M. K. is supported by the Deutsche Stiftung für Herzforschung (grant no. F/37/11), the Deutsches Zentrum für Herz Kreislauf Forschung (grant no. DZHK_B 19 SE), and the Deutsche Forschungsgemeinschaft (grant no. KR3770/11-1 and KR3770/14-1). B. M.-M. is supported by the European Union’s Horizon 2020 research and innovation programme (Marie Skłodowska-Curie grant, agreement No 813533). BD. K. is supported by a British Heart Foundation personal chair (grant no. CH/13/2/30154).

## Disclosures

All authors declare no conflict of interest.

## Supplemental Materials

Expanded Materials and Methods

Online Tables I-XIV

Online Figures I-XIV

Data set (Supplementary datafile)

References ^17^, ^19^, ^27^-^30^, ^32^, ^54^-^71^

## Author contributions

Acquisition of data and material: H. L., M. J., M. D., N. B., C. A.-A., I. N., E. D., SA. D., HJ. C., BD. K. Molecular and cellular experimental work: H. L., M. D., N. B., O. B., I. N., Z. Z., SA. D., P. L., G. E. Provision and analysis of clinical and bioinformatic data: N. P., J. C., M. B., KC. K., J. Z., E. M., T. M., J. H., P. E., JR. P., HJ. C., BD. K., M. K. Bioinformatic analyses: M. J., F. W., R. G., L. H., JR. P., B. M-M. Editing and reviewing the manuscript: M. J., M. D., SA. D., R. G., L. H., J. H., P. E., JR. P., R. L., T. M., HJ. C., BD. K. Writing the manuscript: H. L., M. J., B. M-M., M. K. All authors commented on, edited and approved the manuscript. Supervision of the study: B. M.-M, M.K.

## Non-standard Abbreviations and Acronyms

ASD: atrial septal defect
ASDII: atrial septal defect type II
ATAV: anomalies of thoracic arteries and veins
CHD: congenital heart disease
CHSD: Congenital Heart Surgery Database
CE: cardiac enhancer
CM: cardiomyocyte
CPC: cardiac progenitor cells
ESC: embryonic stem cell
GO: gene ontology
GSEA: gene set enrichment analysis
GWAS: genome-wide association study
iPS: induced pluripotent stem cell
KORA: Cooperative Health Research in the Region of Augsburg
RVOT: right ventricular outflow tract
scRNAseq: single cell RNA sequencing
SHF: second heart field
SNP: single nucleotide polymorphism
STS: Society of Thoracic Surgeons
TGA: transposition of the great arteries
TOF: Tetralogy of Fallot
VSD: ventricular septal defect

## Notes

### Competing Interest Statement

The authors have declared no competing interest.

