## Supplemental Materials, Tables and Figures for "Genome-wide association study in European patients with congenital heart disease identifies risk loci for transposition of the great arteries and anomalies of the thoracic arteries and veins and expression of discovered candidate genes in the developing heart"

**Online Table I.** Possible causal variants in *MACROD2* and *GOSR2* gene loci

**Online Table II.** Analysis of loci 12q24 and 13q32 in patients with right heart lesions

**Online Table III.** Possible influence of rs11784 on expression of *GOSR2* and *WNT3*

**Online Table IV.** List of candidate genes within the LD region of associated loci

**Online Table V.** Gene list for GSEA analysis

**Online Table VI.** GSEA analysis

**Online Table VII.** Patients for candidate gene expression analyses

**Online Table VIII.** Adult control patients for candidate gene expression analyses

**Online Table IX.** Candidate gene knockout mouse models

**Online Table X.** Cross ethnical validation

**Online Table XI.** Imputation score for all lead SNPs

**Online Table XII.** Genomic inflation

**Online Table XIII.** Power analysis

**Online Table XIV.** Primers used for qRT-PCR analyses and genotyping

**Online Figure I.** SNPs associated with right heart lesions

**Online Figure II.** SNPs associated with left heart lesions

**Online Figure III.** SNPs associated with septal defects

**Online Figure IV.** SNPs associated with ASD

**Online Figure V.** SNPs associated with ASDII

**Online Figure VI.** SNPs associated with anomalies of thoracic arteries and veins

**Online Figure VII.** Participation of CHD-associated SNP-carrying genes in signaling cascades

**Online Figure VIII.** Expression of candidate genes during cardiac differentiation of human iPS cells

**Online Figure IX.** Expression of *MACROD2*, *GOSR2* and *WNT3* in patient tissue with or without risk variant

**Online Figure X.** Expression of candidate genes in pediatric and adult aortic tissue

**Online Figure XI.** Expression of candidate genes in pediatric and adult atrial tissue

**Online Figure XII.** Identification of different cell types after single-cell RNAseq by expression of defined marker genes

**Online Figure XIII.** Workflow for general GWAS quality control

**Online Figure XIV.** Quality control steps and PCA plots to analyze population stratification

**SUPPLEMENTARY METHODS**

#### Genotype calling

Genotype calling was done following the Axiom™ Genotyping Solution Data Analysis Guide (http://tools.thermofisher.com/content/sfs/manuals/axiom_genotyping_solution_analysis_guide.pdf). It provides a standard workflow to perform quality control analysis for samples and plates, SNP filtering prior to downstream analysis, and advanced genotyping methods. The workflow utilizes three software systems, including Axiom™, Analysis Suite, Power Tools (APT) and SNPolisher R package. Initially we had 20 plates and 1,921 individual samples in total. Of those, 1,803 arrays passed all quality control steps (sample DQC > 82%, sample call rate > 97%). In order to obtain a high quality of genotype calling only "PolyHighRes" and “MonoHighRes" samples were kept for the next steps.

**Quality control, imputation and association analysis**

All statistical analyses and quality control procedures for the two British cohorts are described in detail in the two respective publications. [17, 19] For the German cohort a standardized eight step GWAS quality control procedure was developed and applied to the genetic data (Online Figure XIII and XIV). Prior to imputation samples were excluded from further analysis for the following reasons: call rate < 98%, incorrect or ambiguous sex call or potential sample contamination. In addition, the thresholds for relatedness and population outliers were set at pihat ≥ 0.09 in an identical by descent (IBD) analysis and a deviation ≥ 2 SD was applied in multidimensional scaling (MDS) analysis. SNPs were excluded if their missing rate was > 3%, if the minor allele content (MAC) was < 5, if the $p$ value for the Hardy-Weinberg equilibrium was $\leq 1x10^{-5}$ in controls or if they failed the cluster quality check. The population structures were evaluated using a set of pruned autosomal variants with MAF > 0.05 $p < 1x10^{-5}$, $r^2 \leq 0.2$ between pairs of variants (indep-pairwise 50 5 0.2). For the principle component analysis (PCA) in PLINK (v1.90b3.36) [54] 119,381 independent SNPs were pruned (Online Figure XIVB and XIVC) except the quality cluster check for which Affymetrix SNPolisher (v1.5.2) [55] was used.

Genome-wide imputation was conducted based on the Haplotype Reference Consortium using the Sanger Imputation Service. All individuals were imputed on the Sanger imputation server (https://imputation.sanger.ac.uk/) with the Haplotype Reference Consortium panel and EAGLE v2.4.1 (https://data.broadinstitute.org/alkesgroup/Eagle/) and positional BWT (PBWT) pipelines. Imputed variants with an AF < 0.005 and/or an info score of < 0.7 were excluded from the statistical analysis. The application of these filters resulted in a total of 20,441,516 high-quality SNPs available for the meta-analysis in up to 1,495 cases and 3,554 controls. Due to gender mismatch and inappropriate diagnoses the number of samples for the final analysis had to be reduced to 1,440. For the British cohort 11,356,134 high-quality SNPs were available. The shared set used for the meta-analysis included 9,216,527 SNPs. The information on the imputation score of all lead SNPs is shown in Online Table XI. The analysis of single SNP genetic association was performed using SNPTEST 2.5.2. (https://mathgen.stats.ox.ac.uk/genetics_software/snptest/snptest.html) via logistic regression using probabilistic imputed allele dosages with adjustment for age, sex and the first 10 ancestry principal components. We have estimated the effective number of independent markers ($M_{eff}$) by calculating the reciprocal of the variance of the off-diagonal elements of the genetic relatedness matrix. [56, 57] The genome-wide significance cutoff is $9.5x10^{-8}$ and $1.9x10^{-7}$ with *q* of 0.05 and 0.1, respectively. In accordance with the majority of published GWAS analyses we have used $5x10^{-8}$ and $1x10^{-5}$ as genome-wide and suggestive significance cutoff levels. The value of the inflation factor λ for all CHD cases and sub-groups is given in Online Table XII. The GWAS.PC package (v1.0) in R was used to confirm that data from each subgroup can be obtained with sufficient power (Online Table XIII).

**Meta-analysis**

The quality of summary statistics of each GWAS dataset was controlled with the EasyQC pipeline, version 8.5 (https://omictools.com/easyqc-tool). For the meta-analysis we used the fixed-effect inverse-variance method with METAL, release 2011-03-25 (http://csg.sph.umich.edu/abecasis/metal/). Genomic control was done in each study separately prior to meta-analysis by calculating the inflation factor λ and adjusting for it. Lead SNPs of independent genome-wide significant signals in meta-analysis results were defined by LD-based independent "clumps" in PLINK (v1.90b3.36) with a *p* value < $1x10^{-5}$, $r^2 > 0.05$, and a clumping distance of < 500 kb. The heterogeneity of lead SNPs was estimated with random-effects meta-analysis using METASOFT v2.0.1 (http://genetics.cs.ucla.edu/meta/).

##### Identification of potentially causal variants by CAVIARBF

To prioritize the possible causal variants identified by our GWAS the fine-mapping tool CAVIARBF (https://bitbucket.org/Wenan/caviarbf/src/default/) was applied. This tool uses an approximate Bayesian method allowing to deal with multiple causal variants. [58] We used the 74 baseline annotations in stratified LD Score regression. [59] SNPs within a 50-kb radius of a lead SNP and with MAF > 0.01 were considered. 1000 genome was used as the reference panel and 0.2 was added to the main diagonal of the LD as suggested correction. Exact Bayes factor is averaged over prior variances of 0.01, 0.1, and 0.5. The elastic net parameters were selected via 10-fold cross-validation.

##### GeneHancer annotation

To detect the putative regulatory implication of the association signals, we annotated the significant SNPs to GeneHancer database. [60] The records of regulatory elements and linked genes were downloaded from UCSC table browser. A SNP is linked to a regulatory element by the colocalization for both the SNP and its proxy SNPs, which is defined with $R^2 > 0.6$ in 1000 genome EUR reference panel.

##### Gene-set enrichment analysis

For the analysis of genome-wide and highly significant SNPs the GO tool of the Broad Institute was used (http://software.broadinstitute.org/gsea/msigdb/annotate.jsp). The functional analysis was performed by ClueGO, a network based functional enrichment method, which can generate new functional groups by measuring the similarity between different pathways/terms. The method will produce both term- and group-based enrichment score for a better visualization and interpretation. Gene-level enrichment was performed by ClueGO (v2.5.4) in Cytoscape 3.7.1, with GO (Biological Processes, version from Apr 24th, 2019) (https://cytoscape.org/), GO terms level: 3–8, GO term with 2 genes and 2% total genes associated; GO terms were grouped by Kappa score with default settings.

Bonferroni-corrected *p* value < 0.1 was considered as the cutoff for significant enrichment. For the GSEA analysis in Online Table V a cutoff *p* value < 0.0005 was chosen to control the FDR at 0.05 for the gene selection by the Benjamini-Hochberg correction. There the lowest *p* value is assigned to the gene for *p* value adjustment which is equal to snp-wise=top, 1 in MAGMA. [61]

**Genotyping of patients for gene expression in cardiac tissue**

To measure gene expression in cardiac tissue a number of patients were analyzed who had not been genotyped by GWAS. In these cases genomic DNA from peripheral blood was amplified by PCR using the following conditions: 95°C for 2 min, 40 cycles of 95°C for 30 sec, 60°C for 30 sec and 72°C for 90 sec with Fast Start High Fidelity Enzyme Blend (Roche, Mannheim, Germany) and a final primer concentration of 0.4 µM. Sequences were verified by conventional Sanger sequencing. The exact sequences of all primers are indicated in Online Table XIV.

**qRT-PCR analysis of gene expression in cardiac tissue**

Tissue samples were obtained during the operation and were snap-frozen immediately in liquid nitrogen. They were kept at -196°C until further use. RNA was extracted using the RNeasy Plus Universal kit (Qiagen, Hilden, Germany) according to the manufacturer's recommendation. cDNA was synthesized from 100 ng total RNA using M-MLV reverse transcriptase (100 U), 250 ng random hexamer primers, 10 mM DTT, dNTPs (0.5 mM each), 15 mM $MgCl_2$, 375 mM KCl and 250 mM Tris-HCl pH 8.3 in a final volume of 30 µL. qRT-PCR analyses were performed on a Quant Studio 3 (ThermoFisher, Germering, Germany) using the following conditions: 95°C for 10 min, 40 cycles of 95° C for 15 sec and 60° C for 1 min using 0.3 µM of each primer. The expression of *ACTB* (β-actin) was used to normalize the expression levels in individual samples. The exact sequences of all primers are indicated in Online Table XIV.

**Spontaneous differentiation of murine embryonic cells**

Murine ESCs were differentiated according to a standard "hanging drop" protocol. [62] Cells were grown for two days on gelatine-coated 6-well plates in IMDM-ES medium (Biochrom, AG, Billierica,

MA) supplemented with 20% FCS (ThermoFisher Scientific, Waltham, MA), 0.1 mM 1-thioglycerol (Sigma-Aldrich, St. Louis, MO) and $10^3$ U/mL LIF (Millipore, Billerica, MA). Hanging drops (1,000 cells per droplet) were prepared on 15 cm cell culture dishes in differentiation medium (IMDM supplemented with 20% FCS, 0.1 mM 1-thioglycerol, 0.05 mg/mL L-ascorbic acid (Sigma-Aldrich) and antibiotics). Culture dishes were cultured upside-down for two days to allow embryoid body (EB) formation. Then, EBs were flooded with differentiation medium and cultured with medium change every other day. On day seven GFP-positive cardiac progenitors and their GFP-negative counterparts were sorted by FACS. RNA purification and cDNA production was performed as described above.

**Directed cardiac differentiation of human induced pluripotent stem cells**

The human induced pluripotent stem cell (iPSC) line S was established from peripheral blood mononuclear cells of a healthy 34 year old male proband using Sendai virus according to the manufacturer's protocol (Invitrogen, Carlsbad, CA) and meets all criteria of fully reprogrammed iPSCs. Differentiation into human cardiomyocytes was done according to a previously published protocol. [63] Human iPSCs were seeded into 24-well plates and grown to confluence in normal mTeSR E8 medium (Stem Cell Technologies, Cologne, Germany). On day 0 the medium was switched to RPMI1640 supplemented with *O. sativa*–derived recombinant human albumin (500 µg/mL, Sigma-Aldrich) and L-ascorbic acid 2-phosphate (213 µg/mL, Sigma-Aldrich) referred to as CDM3. From day 0 to 2 CDM3 was supplemented with 4 µM CHIR99021 (LC Laboratories, Woburn, MA), from day 2 to 4 the cells received CDM3 and 2 µM WNT-C59 (Selleckchem, Munich Germany). Thereafter, CDM3 was replaced every other day. Every second day cells in duplicate wells were lysed with RNA lysis buffer (Peqlab, Erlangen, Germany), purified and cDNA was produced as described above.

**RNAseq analysis in murine cardiac progenitor cells and cardiomyocytes**

Previously published RNAseq data were reanalyzed. [28] Original sequencing data were deposited at the Sequence Read Archive (PRJNA229481). For this study cardiac progenitor cells (CPCs) and cardiomyocytes (CMs) were isolated. CPCs were obtained from embryonic hearts (E9-11) of the

*Nkx2.5* cardiac enhancer-eGFP transgenic mouse line. [27] Embryos were cut into small pieces and digested with a collagenase II (10,000 U/mL, Worthington Biochemical Corporation, Lakewood, NJ) /DNase I (10,000 U/µL, Roche, Molecular Systems Inc., Rotkreuz, Switzerland) solution for 1 h at 37 °C to obtain single cell suspension. Cells were washed and resuspended in PBS/0.5% BSA/2mM EDTA for flow cytometry. GFP-positive CPCs were isolated with a FACS ARIATM Illu flow cytometer (BD Biosciences, San Jose, CA). Dead cells were excluded by propidium iodide staining (2 µg/mL, Sigma-Aldrich, Munich, Germany). FSC pulse width was used to exclude doublets from sorting. For RNAseq cells were sorted into RLTplus Buffer (Qiagen) containing β-mercaptoethanol (10 µL/mL) to extract DNA and total RNA.

CMs were obtained from C57/Bl6 mice at 12 weeks of age. Hearts were retrograde perfused with digestion buffer for 12 min. The enzymatic digest was stopped by addition of 5% FCS and gentle dissociation. Cells were passed through a 100 µm filter. CMs were identified by a high FSC signal and viable cells were discriminated by Draq5 (Cell Signaling Technology, Frankfurt, Germany). Polyadenylated RNA was isolated from total RNA using magnetic beads (NEBNext Poly(A) mRNA Magnetic Isolation Module, NEB, Frankfurt, Germany). Libraries were constructed using the NEBNext Ultra RNA Library Prep Kit for Illumina (NEB) according to the manufacturer´s instruction. A heatmap of differentially regulated genes was generated with the ClustVis software (https://biit.cs.ut.ee/clustvis_large/).

**Mouse single-cell RNAseq data**

We re-analyzed a previously published single-cell RNA sequencing dataset obtained from embryonic mouse hearts at E8.5, E9.5, and E10.5 stages of development. Technical details on dissection, library preparation, sequencing, and transcript assignment are as previously described. [30] After excluding cells with low library diversity (less than 1,000 unique genes) and extremes in total read counts (less than 100,000 or greater than 3,000,000 total reads), viable data from 1,901 cells remained and were converted to transcripts per million (TPM) for subsequent analysis using the Monocle3 set of tools. [29] After standard preprocessing, cell type assignments were made for five major classes of cells (endothelial, epicardial, mesenchymal, atrial cardiomyocytes, and ventricular cardiomyocytes) using previously described transcriptional definitions incorporating three to five cell-type specific markers.[30] We performed regression analysis as implemented in Monocle3 for four of the mouse orthologs of genes identified in the GWAS *Gosr2, Macrod2, Zbtb10,* and *Wnt3* for spatial localization (obtained at the time of dissection), assigned cell type, and embryonic timepoint. We report raw $p$ values from regression coefficients where the $q$ value adjusted for multiple comparisons by the default Benjamini-Hochberg method was less than 0.05.

**Single-cell RNAseq analysis of human embryonic cells and cells from adult atria and ventricles**

Samples from right atrium and interventricular septum were collected from two patients with no history of coronary artery disease at the German Heart Center Munich and directly snap-frozen in liquid nitrogen in the operating room. Tissue samples were minced and nuclei extracted in lysis buffer containing 5 mM $CaCl_2$, 3 mM magnesium actetate, 2 mM EDTA, 0.5 mM EGTA, 10 mM Tris, 0.2%, Triton X-100, protease inhibitors and DTT. Nuclei were centrifuged in 1 M sucrose and resuspended in PBS. After staining with Draq7, samples were purified by fluorescence-activated nuclei sorting (FANS). Nuclei were counted under the microscope and diluted for subsequent 10x Genomics Chromium™ Next GEM Single Cell 3' Solution v3. Barcoding, cDNA amplification and gene expression library construction were done according to the manufacturer's recommendations. Library sequencing was done at the EMBL Heidelberg Genomics Core Facility. The sequencing parameters were 28 bp for read1, 8 bp for the index, and 56 bp for informative read2.

Single-Cell RNA sequencing data from 676 individual cells previously published by Sahara et al.[32] was uploaded to the Galaxy web platform[64] and we used the public server at usegalaxy.eu for data pre-processing and alignment. Datasets were trimmed using trim galore[65] and aligned with RNA STAR[66] against Genome Reference Consortium Human Build 38 (hg38). Aligned reads were processed with MarkDuplicates[67] and count matrices were generated with FeatureCounts[68]. Samples from adult patients were subjected to the Cellranger pipeline from 10x Genomics with default settings using a premRNA-reference as detailed by the manufacturer.

Seurat[69] objects for Count matrices for all samples were created for downstream analyses. After quality filtering, the data was normalized, scaled and variable features were detected using

SCTransform.[70] Data from embryonic and adult cardiac tissue was integrated as described by Stuart et al.[71] Principal component analysis and Uniform manifold approximation and projection (UMAP) for dimension reduction was used to cluster cells into distinct biological identities. Cell types were identified based on the expression of known markers. For expression analysis of *MACROD2*, *GOSR2*, *WNT3* and *MSX1*, the Seurat object was split into adult and embryonic cardiac populations, retaining the clustering information of the integrated dataset. The Seurat command FeaturePlot was used for visualization of gene expression with min.cutoff = 'q10' and max.cutoff = 'q90' settings.

**Statistical analyses**

To compare the expression levels during directed cardiac differentiation a 2-sided *t*-test was performed. $P$ values < 0.05 were considered to be statistically significant. The significance of gene expression in tissue samples was estimated by the Mann-Whitney-Wilcoxon test if equal variance test or normality test failed. $P$ values < 0.05 were considered to be statistically significant.

#### Online Table I. Possible causal variants in *MACROD2* and *GOSR2* gene loci

| rs ID | Z score | PIP CAVIARBF | gene | cyto band | CHD subgroup |
|---|---|---|---|---|---|
| rs17677363 | -5.3322 | 0.9999244 | *GOSR2* | 17q21.32 | anomalies of thoracic arteries and veins |
| rs11874 | 5.40319 | 0.991485 | | | |
| rs76774446 | 5.32994 | 0.9874922 | | | |
| rs149890280 | -6.37711 | 1 | *MACROD2* | 20q12.1 | transposition of the great arteries |
| rs150246290 | 6.42989 | 1 | | | |
| rs149467721 | 6.05662 | 0.5554932 | | | |
| rs77094733 | 6.02248 | 0.4444939 | | | |

PIP: posterior inclusion probability, CAVIARBF: CAVIAR Bayes factor, CHD: congenital heart disease

### Online Table II. Analysis of loci 12q24 and 13q32 in patients with right heart lesions

| | | | | | | | | DHM collective | | | | | | meta-analysis (DHM + UK collective) | | | | |
|---|---|---|---|---|---|---|---|---|---|---|---|---|---|---|---|---|---|---|
| | ChrPosID hg19 | Strand | Chr | Pos | N [1] | Effect_allele | Other_allele | EAF [2] | Info [3] | BETA [4] | OR | SE [5] | *P* | meta_Effect | meta_StdErr | meta_*P*-value | Direction | Subgroup |
| rs7982677 (13q32) | 13:92988323 | + | 13 | 92988323 | 3999 | A | C | 0,270024 | 0,989605 | 0,00695663 | 1.006981 | 0.0811229 | 0.931689 | -0.0477 | 0.0522 | 0.3607 | +- | septal defects |
| | 13:92988323 | + | 13 | 92988323 | 3717 | A | C | 0,268711 | 0,989537 | -0,133944 | 0.874639 | 0.133799 | 0.311697 | -0.1339 | 0.1338 | 0.3168 | -- | ASDII |
| | 13:92988323 | + | 13 | 92988323 | 5049 | A | C | 0,27035 | 0,989636 | -0,00429724 | 0.995712 | 0.0507463 | 0.932507 | 0.0402 | 0.0306 | 0.1896 | -+ | all CHD |
| | 13:92988323 | + | 13 | 92988323 | 3786 | A | C | 0,268119 | 0,989515 | -0,146722 | 0.863534 | 0.112598 | 0.187516 | -0.097 | 0.0702 | 0.1671 | -- | ASD |
| | 13:92988323 | + | 13 | 92988323 | 3759 | A | C | 0,269706 | 0,989828 | -0,0193519 | 0.980834 | 0.115665 | 0.866948 | -0.0528 | 0.075 | 0.4814 | -- | anomalies of thoracic arteries and veins |
| | 13:92988323 | + | 13 | 92988323 | 3700 | A | C | 0,27129 | 0,989499 | 0,127057 | 1.135482 | 0.132544 | 0.341485 | 0.0944 | 0.0785 | 0.2295 | ++ | TGA |
| | 13:92988323 | + | 13 | 92988323 | 3692 | A | C | 0,269495 | 0,989387 | -0,0969594 | 0.907593 | 0.143157 | 0.495098 | 0.0123 | 0.0898 | 0.8908 | -+ | left heart lesions |
| | 13:92988323 | + | 13 | 92988323 | 3824 | A | C | 0,272319 | 0,989156 | 0,174637 | | 0.0997423 | 0.0828563 | 0.2123 | 0.047 | $6.29 \times 10^{-6}$ | ++ | right heart lesions |
| rs11065987 (12q24) | 12:112072424 | + | 12 | 112072424 | 3999 | G | A | 0,466492 | 1 | 0,055953 | 1.057548 | 0.0729835 | 0.443403 | 0.0084 | 0.0473 | 0.859 | -+ | septal defects |
| | 12:112072424 | + | 12 | 112072424 | 3717 | G | A | 0,465967 | 1 | 0,114226 | 1.121005 | 0.115896 | 0.324546 | -0.1142 | 0.1159 | 0.3243 | -- | ASDII |
| | 12:112072424 | + | 12 | 112072424 | 5049 | G | A | 0,46534 | 1 | 0,00368067 | 1.003687 | 0.0453026 | 0.935247 | -0.0281 | 0.0281 | 0.3163 | -- | all CHD |
| | 12:112072424 | + | 12 | 112072424 | 3786 | G | A | 0,465927 | 1 | 0,0776794 | 1.080776 | 0.0975685 | 0.42613 | 0.0363 | 0.0629 | 0.564 | -+ | ASD |
| | 12:112072424 | + | 12 | 112072424 | 3759 | G | A | 0,462623 | 1 | -0,166374 | 0.846729 | 0.104174 | 0.108993 | 0.1428 | 0.0683 | 0.03646 | ++ | anomalies of thoracic arteries and veins |
| | 12:112072424 | + | 12 | 112072424 | 3700 | G | A | 0,464595 | 1 | -0,0213772 | 0.978850 | 0.121644 | 0.860464 | 0.0905 | 0.0736 | 0.219 | ++ | TGA |
| | 12:112072424 | + | 12 | 112072424 | 3692 | G | A | 0,462893 | 1 | -0,220078 | 0.802456 | 0.127457 | 0.0825705 | 0.0746 | 0.0825 | 0.3658 | +- | left heart lesions |
| | 12:112072424 | + | 12 | 112072424 | 3824 | G | A | 0,46705 | 1 | 0,119852 | 1.127330 | 0.0909645 | 0.187722 | -0.2114 | 0.0443 | $1.79 \times 10^{-6}$ | -- | right heart lesions |

[1]: sample size, [2]: effector allele frequency, [3]: info score, [4]: $\beta = \ln OR$, [5]: standard error of $\beta$

#### Online Table III. Possible influence of rs11784 on expression of *GOSR2* and *WNT3*

| rsid | chr | pos_hg19 | cyto band | gene | type | gene/protein consequences | Ensembl Regulatory Build | roadmap enhancer(tissue) | GeneHancer Interactions | geneAssociationMethods | GeneHancer (GH) Identifier | Gene_By_topologically association domains | SV | SV_PMID |
|---|---|---|---|---|---|---|---|---|---|---|---|---|---|---|
| rs35437121 | 2 | 131769448 | 2q21.1 | ARHGEF4 | protein_coding | intronic(ARHGEF4),coding(ARHGEF4) | | | ARHGEF4 | eQTLs,Distance | GH02J131039,GH02J131036 | GPR148,AMER3,ARHGEF4,FAM168B,PLEKHB2, POTEE,CYP4F31P,MZT2A,TUBA3D | gain | 25217958 |
| rs114503684 | 3 | 141834969 | 3q23 | TFDP2 | protein_coding | intronic(TFDP2),non-coding intronic(TFDP2) | Promoter Flanking Region | | | | | GK5,TFDP2,XRN1 | gain | 19592680;21841781;21293372; 25217958;25503493;21179565 |
| rs2046060 | 3 | 187852486 | 3q27.3 | RP11-430L16.1 | lincRNA | non-coding intronic(RP11-430L16.1) | | | LPP | C-HiC,eRNA_co-expression,eQTLs | GH03J187735,GH03J188081 | LPP,AC022498.1,TPRG1 | gain | 17911159 |
| rs187369228 | 3 | 189802439 | 3q28 | LEPREL1 | protein_coding | intronic(LEPREL1) | | | P3H2 | eQTLs,C-HiC,Distance,eRNA_co-expression | GH03J189961,GH03J189983, GH03J189995,GH03J189999, GH03J190002,GH03J190010, GH03J190016,GH03J190024, GH03J190031,GH03J190033, GH03J190036,GH03J190040, GH03J190048,GH03J190053, GH03J190055,GH03J190062, GH03J190072,GH03J190075 | CLDN1,CLDN16,TMEM207,LEPREL1 | | |
| rs870142 | 4 | 4648047 | 4p16.2 | STX18-AS1 | antisense | non-coding intronic(STX18-AS1) | | | | | | TMEM128,LYAR,ZBTB49,NSG1,STX18,MSX1 | | |
| rs185531658 | 5 | 113136521 | 5q22.3 | YTHDC2 | protein_coding | upstream | | | | | | KCNN2,TRIM36,PGGT1B,CCDC112 | loss | 25217958 |
| rs146300195 | 5 | 128326845 | 5q23.3 | SLC27A6 | protein_coding | intronic(SLC27A6) | | | SLC27A6 | eQTLs,Distance | GH05J128965 | FBN2,SLC27A6,ISOC1,ADAMTS19,KIAA1024L | | |
| rs117527287 | 6 | 85729959 | 6q14.3 | RP3-435K13.1 | pseudogene | upstream | | | | | | TBX18,NT5E,SNX14 | | |
| rs148563140 | 8 | 81475406 | 8q21.13 | RPSAP47 | pseudogene | upstream | | | | | | ZBTB10 | | |
| rs11065987 | 12 | 112072424 | 12q24.12 | BRAP | protein_coding | downstream | | | ALDH2,ADAM1A,ATXN2 | eQTLs,eRNA_co-expression,TF_co-expression,Distance | GH12J111396,GH12J111402, GH12J111424,GH12J111466, GH12J112417 | | | |
| rs7982677 | 13 | 92988323 | 13q31.3 | GPC5 | protein_coding | intronic(GPC5) | | | | | | GPC6,DCT,GPC5,TGDS | gain | 25217958 |
| rs138741144 | 17 | 32286564 | 17q12 | ASIC2 | protein_coding | intronic(ASIC2) | | H3K9me3(Right Atrium) | ASIC2 | C-HiC,Distance | GH17J033905,GH17J033906, GH17J033908,GH17J033928, GH17J033930 | AC005549.3,CCL2,CCL7,CCL11,CCL8,CCL13, CCL1,ASIC2 | gain | 25217958 |
| rs11874 | 17 | 45017193 | 17q21.32 | GOSR2,RP11-156P1.2 | protein_coding | intronic(GOSR2),intronic(RP11-156P1.2),3downstream(GOSR2),3utr(GOSR2) | | H3K36me3(Left Ventricle) | KANSL1,CDC27,**GOSR2** | eQTLs,TF_co-expression,C-HiC,Distance | GH17J047098,GH17J046922, GH17J047274,GH17J046942, GH17J046940 | **WNT3**,WNT9B,GOSR2,RP11-156P1.2,RPRML, CDC27,MYL4,ITGB3,EFCAB13,NPEPPS,NSF | | |
| rs72917381 | 18 | 54546223 | 18q21.31 | WDR7 | protein_coding | intronic(WDR7) | | | WDR7 | eRNA_co-expression,Distance | GH18J056889 | BOD1L2,ST8SIA3,ONECUT2,FECH,WDR7,NARS | | |
| rs150246290 | 20 | 15112880 | 20p12.1 | MACROD2 | protein_coding | intronic(MACROD2) | | | | | | FLRT3,KIF16B,SNRPB2,OTOR,MACROD2 | loss | 21841781;25118596;25217958; 21293372;19592680;25503493 |

#### Online Table IV. List of candidate genes within the LD region of associated loci

| gene | chromosome | expression pattern, potential cardiac relevance | reference(s) |
|---|---|---|---|
| **Figure 1** | **rs185531658** | **all CHD patients** | |
| ***MCC*** (MCC regulator of WNT signaling pathway) | 5q22.2 | expressed in primitive streak, cardiac mesoderm formation | Teo et al., Stem Cells, 30;631-642 (2012) |
| ***TSSK1B*** (testis specific serine kinase 1B) | 5q22.2 | expressed in various tissue, overexpressed in heart | |
| ***YTHDC2*** (YTH domain containing 2) | 5q22.2 | mRNA widely expressed | |
| **Figure 2A** | **rs150246290** | **transposition of the great arteries** | |
| ***MACROD2*** (MACRO Domain Containing 2) | 20p12.1 | mRNA widely expressed | see manuscript |
| ***MACROD2-AS1*** (MACROD2 anti-sense RNA1) | 20p12.1 | RNA widely expressed (non-protein coding) | |
| **Figure 2B** | **rs148563140** | **transposition of the great arteries** | |
| ***ZBTB10*** (zinc finger and BTB containing domain 10) | 8q21.13 | mRNA widely expressed | |
| ***ZNF704*** (zinc finger protein 704) | 8q21.13 | mRNA widely expressed, protein in fetal and adult heart | |
| ***PAG1*** (phosphoprotein membrane anchor with glycosphingolipid microdomains 1) | 8q21.13 | mRNA widely expressed | |
| **Figure 2C** | **rs11874** | **thoracic arteries and veins** | |
| ***ARL17B*** (ADP ribosylation factor like GTPase 17b) | 17q21.31 | mRNA widely expressed | |
| ***WNT3*** (Wnt family member 3) | 17q21.31-32 | mRNA widely expressed, primitive streak and mesoderm formation | Liu et al., Nature Genet. 22;361-365 (1999); see manuscript |
| ***WNT9B*** (Wnt family member 9B) | 17q21.32 | mRNA widely expressed including atrioventricular node cells | Alfieri et al., Dev. Biol., 338;127 (2010) |
| ***GOSR2*** (Golgi SNAP receptor complex member 2) | 17q21.32 | mRNA widely expressed, coronary artery disease and myocardial infaction | see manuscript |
| ***MIR5089*** (micro RNA 5089) | 17q21.32 | RNA weakly expressed in different tissues, but not in heart | |
| ***RPRML*** (reprimo like) | 17q21.32 | mRNA widely expressed | |
| ***CDC27*** (cell division cycle 27) | 17q21.32 | mRNA widely expressed, protein in fetal heart | |
| ***MYL4*** (myosin light chain 4) | 17q21.32 | mRNA expressed in fetal heart and cardiomyocytes, atrial fibrillation | Orr et al., Nat. Commun., 12;11303 (2016) |
| **Figure IA** | **rs11065987** | **right heart lesions (TOF)** | |
| ***CUX2*** (Cut Like Homebox 2) | 12q24.11-12 | mRNA widely expressed | Liu et al., Can J Cardiol., 33; 443-449 (2017); Lin et al., Sci Rep. 7: 40377 (2017); Sinner et al., Circulation, 130;1225-35 (2014) |
| ***MIR6760*** (micro RNA 6760) | 12q24.12 | no data on RNA expression | |
| ***FAM109A*** (PH Domain Containing Endocytic Trafficking Adaptor 1) | 12q24.12 | no data on RNA expression | |
| ***SH2B3*** (SH2B Adaptor Protein 3) | 12q24.12 | mRNA widely expressed, protein in heart | Wang et al., Curr Mol Med., Epub ahead of print; Keefe et al., Hypertension, 73;497-503; Hong et al., Medicine (Baltimore), 97;e13436 (2018) |
| ***ATXN2*** (Ataxin 2) | 12q24.12 | mRNA widely expressed, protein in fetal heart | Liu et al., Oncotarget., 8(63); 106976-106988 (2017); Lv et al., J Mol Cell Cardiol., 112;1-7 (2017) |
| ***BRAP*** (BRCA1 Associated Protein) | 12q24.12 | mRNA widely expressed | Volland et al., Cardiovasc Res. Epub ahead of print |

| gene | chromosome | expression pattern, potential cardiac relevance | reference(s) |
|---|---|---|---|
| | | | (2019); Nozynski et al., Transplant Proc., 48;1746-50 (2016) |
| ***ACAD10*** (Acyl-CoA Dehydrogenase Family Member 10) | 12q24.12 | mRNA widely expressed, protein in heart and fetal heart | |
| ***ALDH2*** (Aldehyde Dehydrogenase 2 Family Member) | 12q24.12 | mRNA widely expressed, protein widely expressed and in heart and fetal heart | Chen et al., Adv Exp Med Biol., 1193; 53-67 (2019) |
| ***ADAM1A*** (ADAM Metallopeptidase Domain 1A (Pseudogene)) | 12q24.12-13 | mRNA widely expressed | |
| ***MAPKAPK5*** (MAPK Activated Protein Kinase 5) | 12q24.12-13 | mRNA widely expressed, protein in heart | Nawaito et al., Am J Physiol Heart Circ Physiol., 313;H46-H58 (2017); Dingar et al., Cell Signal, 22;1063-75 (2010) |
| ***MIR6761*** (micro RNA 6761) | 12q24.12 | no data on RNA expression | |
| ***MAPKAPK5-AS1*** (MAPKAPK5 Antisense RNA1) | 12q24.12 | RNA widely expressed | |
| ***TMEM116*** (Transmembrane Protein 116) | 12q24.13 | mRNA widely expressed | |
| ***ERP29*** (Endoplasmic Reticulum Protein 29) | 12q24.13 | mRNA widely expressed, protein widely expressed and in heart and fetal heart | Su et al., Pulm. Circ., 5;481-497 (2015) |
| ***NAA25*** (N(Alpha)-Acetyltransferase 25, NatB Auxiliary Subunit) | 12q24.13 | mRNA widely expressed, protein in fetal heart | |
| ***TRAFD1*** (TRAF-Type Zinc Finger Domain Containg 1) | 12q24.13 | mRNA widely expressed, protein in heart and fetal heart | |
| ***MIR3657*** (micro RNA 3657) | 12q24.13 | no data on RNA expression | |
| **Figure IB** | **rs7982677** | **right heart lesions (TOF)** | |
| ***GPC5-AS2*** (GPC5 Antisense RNA 2) | 13q31.3 | mRNA expressed in heart | |
| **Figure IC** | **rs146300195** | **right heart lesions** | |
| ***SLC27A6*** (solute carrier family 27 member 6) | 5q23.3 | mRNA widely expressed, heart failure | Auinger et al., Br. J. Nutr., 107;1422-1428 (2012) |
| ***ISOC1*** (isochorismatase domain containing 1) | 5q23.3 | mRNA widely expressed | |
| ***MIR4633*** (micro RNA 4633) | 5q23.3 | RNA weakly expressed in different tissues, including heart | |
| ***MIR4460*** (micro RNA 4460) | 5q23.3 | RNA only detectable in blood and esophagus | |
| **Figure IIA** | **rs35437121** | **left heart lesions** | |
| ***AMER3*** (APC membrane recruitment protein 3) | 2q21.1 | RNA weakly expressed in different tissues, protein in fetal heart and adipocytes | |
| ***ARHGEF4*** (Rho guanine exchange factor 4) | 2q21.1 | mRNA widely expressed | |
| ***FAM168B*** (family with sequence similarity 168 member B) | 2q21.1 | mRNA widely expressed | |
| ***PLEKHB2*** (pleckstrin homolgy domain containing B2) | 2q21.1 | mRNA widely expressed | |
| ***POTEE*** (POTE ankyrin domain family member E) | 2q21.1 | mRNA widely expressed, protein in heart | |
| ***LOC440910*** (uncharacterized LOC440910) | 2q21.1 | no data on RNA expression | |
| ***WTH3DI*** (= RAB6D, member ras oncogene family) | 2q21.1 | no data on mRNA expression | |
| ***MZT2A*** (mitotic spindle organizing protein 2A) | 2q21.1 | mRNA widely expressed | |

| gene | chromosome | expression pattern, potential cardiac relevance | reference(s) |
|---|---|---|---|
| ***NOC2LP2*** (NOC2 like nucleolar associated transcriptional repressor pseudogene 2) | 2q21.1 | pseudogene | |
| ***LINC01120*** (long intergenic non-protein coding RNA 1120) | 2q21.1 | ncRNA widely exprressed | |
| ***TUBA3D*** (tubulin alpha 3d) | 2q21.1 | mRNA widely expressed, cardiac arrrhthmia | Friedman et al., NPJ Genom. Med., 3;9 (2018) |
| ***MIR4784*** (micro RNA 4784) | 2q21.1 | RNA weakly expressed in different tissues | |
| ***LOC150776*** (sphingomyelin phosphodiesterase 4, neutral membrane (neutral sphingomyelinase-3)) | 2q21.1 | pseudogene | |
| **Figure IIB** | **rs114503684** | **left heart lesions** | |
| ***LOC646730*** (LINC02618) | 3q23 | no data on mRNA expression | |
| ***RNF7*** (ring finger protein 7) | 3q23 | mRNA widely expressed | |
| ***GRK7*** (G protein coupled receptor kinase 7) | 3q23 | mRNA widely expressed | |
| ***ATP1B3*** (ATPase Na+/K+ transporting subunit beta 3) | 3q23 | mRNA widely expressed | |
| ***LOC646730* (LINC02618)** | 3q23 | no data on mRNA expression | |
| ***TFDP2*** (transcription factor Dp-2) | 3q23 | mRNA widely expressed | |
| **GK5** (glycerol kinase 5) | 3q23 | mRNA widely expressed | |
| **Figure IIC** | **rs2046060** | **left heart lesions** | |
| ***LPP-AS2*** (LPP antsense RNA 2) | 3q27.3 | RNA widely expressed | |
| ***FLJ42393*** (uncharacterized LOC401105) | 3q27.3 | ncRNA widely exprressed | |
| ***LPP*** (LIM domain containing preferred translocation partner in lipoma) | 3q27.3-q28 | mRNA widely expressed | |
| **Figure IIIA** | **rs185531658** | **septal defects** | |
| **see above (Figure 1)** | | | |
| **Figure IIIB** | **rs138741144** | **septal defects** | |
| ***ASIC2*** (acid sensing ion channel subunit 2) | 17q11.2-q12 | mRNA widely expressed, protein in heart | |
| ***AA06*** (uncharacterized LOC100506677) | 17q12 | pseudogene | |
| **Figure IV** | **rs870142** | **ASD** | |
| ***STX18-AS1*** (STX18 anti-sense RNA 1 (head to head)) | 4p16.2 | ncRNA widely exprressed | |
| ***SNORD162*** (small nucleolar RNA, C/D box 162) | 4p16.2 | no data on RNA expression | |
| **Figure VA** | **rs72917381** | **ASDII** | |
| ***TXNL1*** (thioredoxin like 1) | 18q21.31 | mRNA widely expressed, lower expression in RV | Su et al., Pulm. Circ., 5;481-497 (2015) |
| ***WDR7*** (WD repeat domain 7) | 18q21.31 | mRNA widely expressed | |
| **Figure VB** | **rs187369228** | **ASDII** | |
| ***TP63*** (tumor protein 63) | 3q28 | mRNA widely expressed, arrhthmogenic cardiomyopathy | Poloni et al., Heart Rhythm, 16;773-780 (2019) |
| ***MIR944*** (micro RNA 944) | 3q28 | RNA weakly expressed in skeletal muscle, esophagus and skin | |
| ***LEPREL1*** (=*P3H2*, proloy 3 hydroxylase 2) | 3q28 | protein expressed in heart | |
| **Figure VI** | **rs117527287** | **ASDII** | |
| ***TBX18-AS1*** (TBX18 antisense RNA 1) | 6q14.3 | no data on RNA expression | |
| ***TBX18*** (T-box 18) | 6q14.3 | mRNA widely expressed, protein in heart and lung, cardiac development | Kapoor et al., Nat. Biotechnol., 31;54-62 (2013) |
| ***LOC101928820*** (long intergenic non-protein coding | 6q14.3 | no data on RNA expression | |

| gene | chromosome | expression pattern, potential cardiac relevance | reference(s) |
|---|---|---|---|
| RNA 2535) | | | |
| ***NT5E*** (5' nucleotidase ecto) | 6q14.3 | mRNA widely expressed, calcification of arteries | Hellman et al., Medicine (Baltimore), 98;e15065 (2019) |
| ***SNX14*** (sorting nexin 14) | 6q14.3 | mRNA widely expressed | |

#### Online Table V. Gene list for GSEA analysis

| CHD population | gene |
|---|---|
| all | *GRM4* |
| ASD | *IGSF21* |
| | *LEPREL1* |
| | *MAML3* |
| | *ADAMTSL1* |
| | *RHBDF2* |
| ASDII | *LEPREL1* |
| | *TMTC2* |
| | *WDR7* |
| left heart lesions | *ARHGEF4* |
| | *TRAF3IP1* |
| | *TFDP2* |
| | *LPP* |
| | *ZFC3H1* |
| | *THAP2* |
| | *RP11-293I14.2* |
| | *SMCHD1* |
| right heart lesions | *FBN2* |
| | *SLC27A6* |
| | *ISOC1* |
| | *FLT4* |
| septal defects | *CACNA2D1* |
| | *AKR1C1* |
| thoracic arteries and veins | *INHBA* |
| | *GATA4* |
| | *C8orf49* |
| | *NEIL2* |
| | *FDFT1* |
| | *RP11-297N6.4* |
| | *CTSB* |
| | *WNT9B* |
| | *GOSR2* |
| | *RP11-156P1.2* |
| | *RPRML* |
| | *CDC27* |
| | *MYL4* |
| TGA | *HS6ST1* |
| | *GABRA6* |
| | *GABRA1* |
| | *ZNF704* |
| | *TMEM74* |
| | *GATA3* |
| | *DLG2* |
| | *MACROD2* |

red: genes with genome-wide significance. Cutoff value for all genes $p < 0.0005$

#### Online Table VI. GSEA analysis

| Gene set name | # genes | *p* value | FDR *q* value |
|---|---|---|---|
| cell-cell signaling | 8 | $3.41 \times 10^{-7}$ | $2.02 \times 10^{-3}$ |
| embryonic organ development | 6 | $1.59 \times 10^{-6}$ | $2.64 \times 10^{-3}$ |
| anatomical structure formation involved in morphogenesis | 8 | $1.79 \times 10^{-6}$ | $2.64 \times 10^{-3}$ |
| embryonic morphogenesis | 6 | $8.03 \times 10^{-6}$ | $7.83 \times 10^{-3}$ |

### Online Table VII. Patients for candidate gene expression analyses

| ID | sex | age [1] | genotype [2] | genotyped by | tissue | diagnosis | STS code | gene expression |
|---|---|---|---|---|---|---|---|---|
| 604 | m | 5 d | wt | GWAS | aorta | TGA, IVS | 83 | |
| 687 | m | 6 d | wt | GWAS | aorta | TGA, IVS | 83 | |
| 690 | f | 9 d | wt | GWAS | aorta | TGA, IVS | 83 | |
| 707 | m | 9 d | wt | GWAS | aorta | TGA, IVS | 83 | |
| 726 | m | 6 d | wt | GWAS | aorta | TGA, IVS | 83 | |
| 745 | m | 4 d | wt | GWAS | aorta | TGA, IVS | 83 | |
| 841 | m | 7 d | wt | GWAS | aorta | TGA, IVS | 83 | |
| 937 | m | 6 d | wt | GWAS | aorta | TGA, IVS | 83 | |
| 1048 | m | 6 d | wt | GWAS | aorta | TGA, IVS | 83 | |
| 1638 | m | 7 d | wt | GWAS | aorta | TGA, IVS | 83 | |
| 2594 | m | 6 d | wt | GWAS | aorta | TGA, VSD | 85 | |
| 2771 | f | 8 d | wt | GWAS | aorta | TGA, IVS | 83 | |
| 2773 | m | 9 d | wt | GWAS | aorta | TGA, IVS | 83 | |
| 3234 | m | 10 d | wt | GWAS | aorta | TGA, VSD | 85 | |
| 3755 | f | 9 d | wt | GWAS | aorta | TGA, IVS | 83 | |
| 3883 | m | 8 d | wt | GWAS | aorta | TGA, IVS | 83 | |
| 703 | m | 7 d | het | GWAS | aorta | TGA, IVS | 83 | |
| 859 | m | 9 d | het | GWAS | aorta | TGA, IVS | 83 | *MACROD2* |
| 1755 | m | 5 d | het | GWAS | aorta | TGA, IVS | 83 | |
| 2812 | m | 6 d | het | GWAS | aorta | TGA, IVS | 83 | |
| 3898 | m | 8 d | het [3] | GWAS | aorta | TGA, VSD | 85 | |
| 605 | m | 10 d | wt | PCR | aorta | TGA, IVS | 83 | |
| 1026 | f | 6 d | wt | PCR | aorta | TGA, IVS | 83 | |
| 1663 | f | 18 d | wt | PCR | aorta | TGA, IVS | 83 | |
| 1665 | f | 13 d | wt | PCR | aorta | TGA, VSD | 85 | |
| 1789 | m | 12 d | wt | PCR | aorta | TGA, IVS | 83 | |
| 2855 | f | 9 d | wt | PCR | aorta | TGA, IVS | 83 | |
| 2982 | m | 10 d | wt | PCR | aorta | TGA, IVS | 83 | |
| 3156 | m | 14 d | wt | PCR | aorta | TGA, IVS | 83 | |
| 3589 | m | 6 d | wt | PCR | aorta | TGA, IVS | 83 | |
| 3624 | m | 13 d | het | PCR | aorta | TGA, IVS | 83 | |
| 3789 | m | 8 d | wt | PCR | aorta | ccTGA | 82 | |
| 4063 | m | 30 d | wt | PCR | aorta | TGA, IVS | 83 | |
| 4114 | m | 7 d | wt | PCR | aorta | TGA, VSD | 85 | |
| 4639 | m | 6 d | wt | PCR | aorta | TGA, VSD | 85 | |
| | 28m/7f | 8 (4-30) | | | | | | |
| 300 | m | 12378 d | wt | GWAS | aorta | aortic aneurysm (including pseudoaneurysm) | 102 | |
| 801 | m | 78 d | wt | GWAS | aorta | coronary artery anomaly, anomalous pulmonary origin (includes ALCAPA) | 159 | |
| 821 | m | 3 d | wt | GWAS | ductus | coarctation of aorta | 95 | |
| 1949 | m | 6 d | wt | GWAS | ductus | VSD + aortic arch hypoplasia | 168 | |
| 2061 | f | 8 d | wt | GWAS | ductus | interrupted aortic arch | 98 | |
| 2330 | f | 32 d | wt | GWAS | ductus | VSD + aortic arch hypoplasia | 168 | *GOSR2, WNT3* |
| 2405 | f | 87 d | wt | GWAS | ductus | VSD + aortic arch hypoplasia | 168 | |
| 2529 | f | 9673 d | wt | GWAS | aorta | coarctation of aorta | 95 | |
| 3466 | m | 7 d | wt | GWAS | ductus | coarctation of aorta | 95 | |
| 3815 | f | 8 d | wt | GWAS | ductus | coarctation of aorta | 95 | |
| 3992 | m | 120 d | wt | GWAS | ductus | coarctation of aorta | 95 | |
| 771 | m | 3 d | het | GWAS | ductus | aortic arch hypoplasia | 96 | |
| 1308 | m | 5455 d | het | GWAS | aorta | aortic aneurysm (including pseudoaneurysm) | 102 | |
| 1461 | m | 4 d | het | GWAS | ductus | coarctation of aorta | 95 | |

| ID | sex | age [1] | genotype [2] | genotyped by | tissue | diagnosis | STS code | gene expression |
|---|---|---|---|---|---|---|---|---|
| 2329 | m | 5 d | het | GWAS | ductus | aortic arch hypoplasia | 96 | *GOSR2, WNT3* |
| 3961 | f | 4 d | het | GWAS | ductus | coarctation of aorta | 95 | |
| 867 | m | 11 d | het | PCR | aorta | coarctation of aorta | 95 | |
| 2754 | m | 8 d | wt | PCR | ductus | aortic arch hypoplasia | 96 | |
| 2993 | f | 3 d | hom [4] | PCR | ductus | coarctation of aorta | 95 | |
| 3754 | f | 7 d | het | PCR | ductus | interrupted aortic arch | 98 | |
| 4038 | f | 6 d | het [5] | PCR | aorta | aortic arch hypoplasia | 96 | |
| 4038 | f | 6 d | het [5] | PCR | ductus | aortic arch hypoplasia | 96 | |
| 4180 | m | 60 d | hom | PCR | aorta | coarctation of aorta | 95 | |
| | 13m/10f | 8 (3-12378) | | | | | | |
| 27 | f | 1465 d | | | right atrium | HLHS | 69 | *ARHGEF4, TFDP2* |
| 266 | f | 9 d | | | right atrium | HLHS | 69 | |
| 474 | f | 540 d | | | right atrium | HLHS | 69 | |
| 2677 | m | 19 d | | | right atrium | HLHS | 69 | |
| 2684 | m | 8 d | | | right atrium | HLHS | 69 | |
| 3007 | f | 5 d | | | right atrium | HLHS | 69 | |
| 3582 | f | 10 d | | | right atrium | HLHS | 69 | |
| 5347 | f | 6 d | | | right atrium | HLHS | 69 | |
| | 2m/6f | 9.5 (5-1465) | | | | | | |
| 1531 | m | 6 d | | | aorta | HLHS | 69 | |
| 3258 | f | 6 d | | | aorta | HLHS | 69 | |
| 4221 | m | 7 d | | | aorta | HLHS | 69 | |
| 5942 | m | 8 d | | | aorta | HLHS | 69 | |
| 6179 | m | 7 d | | | aorta | HLHS | 69 | |
| | 4m/1f | 7 (6-8) | | | | | | |
| 3526 | m | 140 d | | | right atrium | TOF | 28 | *SLC27A6* |
| 3951 | f | 1004 d | | | right atrium | TOF | 28 | |
| 4499 | f | 342 d | | | right atrium | TOF | 28 | |
| 4618 | m | 106 d | | | right atrium | TOF | 28 | |
| 4758 | m | 113 d | | | right atrium | TOF | 28 | |
| 6122 | m | 130 d | | | right atrium | TOF | 28 | |
| | 4m/2f | 135 (106–1004) | | | | | | |
| 28 | f | 2272 d | | | right atrium | ASDII | 2 | *ASIC2, STX18-AS1, STX18, MSC1, P3H2, WDR7* |
| 92 | f | 4262 d | | | right atrium | ASDII | 2 | |
| 98 | f | 824 d | | | right atrium | ASDII | 2 | |
| 332 | m | 2980 d | | | right atrium | ASDII | 2 | |
| 614 | f | 5183 d | | | right atrium | ASDII | 2 | |
| 739 | f | 1887 d | | | right atrium | ASDII | 2 | |
| 2930 | f | 2600 d | | | right atrium | ASDII | 2 | |
| 3586 | f | 1069 d | | | right atrium | ASDII | 2 | |
| 5321 | m | 692 d | | | right atrium | ASDII | 2 | |
| 6166 | f | 6 d | | | right atrium | ASDII | 2 | |
| | 2m/8f | 2079.5 (6–5183) | | | | | | |
| 5352 | m | 11 d | | | aorta | ASDII | 2 | |
| 5387 | f | 9 d | | | aorta | ASDII | 2 | |
| 5632 | m | 8 d | | | aorta | ASDII | 2 | |
| 5817 | m | 7 d | | | aorta | ASDII | 2 | |
| 5920 | m | 4 d | | | aorta | ASDII | 2 | |
| 5973 | m | 7 d | | | aorta | ASDII | 2 | |
| | 5m/1f | 7.5 (4–11) | | | | | | |

[1] at the time of operation, [2] *MACROD2* locus for all four SNPs wt: wild-type, het: heterozygous, [3] only rs149890280 and rs150246290 are heterozygous, [2] *GOSR2* locus for all three SNPs wt: wild-type, het: heterozygous, hom: homozygous, [4]only rs76774446 is homozygous, [5] only rs17677363 is heterozygous. HLHS: hypoplastic left heart syndrome.

#### Online Table VIII. Adult control patients for candidate gene expression analyses

| ID | sex | age [1] | tissue | type of OP | diagnosis | tested for expression of |
|---|---|---|---|---|---|---|
| A463 | m | 77 y | aorta | CABG | atherosclerotic heart disease | *GOSR2, MACROD2, WNT3* |
| A3677 | m | 76 y | aorta | CABG | atherosclerotic heart disease | |
| A4613 | m | 57 y | aorta | CABG | atherosclerotic heart disease | |
| A4861 | f | 79 y | aorta | AVR + aorta | aortic insufficiency | |
| A4872 | f | 82 y | aorta | AVR + aorta | aortic insufficiency | |
| A4877 | f | 67 y | aorta | aorta | aortic aneurysm with dissection | |
| A4902 | m | 78 y | aorta | CABG | atherosclerotic heart disease | |
| A5199 | m | 71 y | aorta | CABG | atherosclerotic heart disease | |
| A5356 | m | 57 y | aorta | CABG | atherosclerotic heart disease | |
| A5644 | m | 53 y | aorta | CABG | atherosclerotic heart disease | |
| | 7m/3f | 73.5 (53 – 82) | | | | |
| A3100 | m | 59 | aorta | AVR + aorta | aortic insufficiency | *ARHGEF4, TFDP2, ASIC2, STX18-AS1, STX18, MSX1, LEPREL1 (=P3H2), WDR7* |
| A3111 | m | 41 | aorta | Aorta | aortic aneurysm | |
| A345 | m | 58 | aorta | Aorta | aortic insufficiency | |
| A3562 | m | 48 | aorta | Aorta | aortic insufficiency | |
| A4472 | m | 53 | aorta | Aorta | aortic insufficiency | |
| A4989 | m | 42 | aorta | AVR + aorta | aortic insufficiency | |
| A5072 | m | 60 | aorta | AVR + aorta | aortic insufficiency | |
| A5283 | f | 57 | aorta | AVR + aorta | aortic insufficiency | |
| A5851 | f | 38 | aorta | AVR + aorta | combined aortic vitium | |
| A5892 | m | 51 | aorta | AVR + aorta | aortic stenosis | |
| A6382 | m | 52 | aorta | AVR + aorta | aortic insufficiency | |
| A6478 | m | 64 | aorta | AVR + aorta | aortic insufficiency | |
| A6641 | m | 49 | aorta | AVR + aorta | combined aortic vitium | |
| A7338 | m | 53 | aorta | AVR + aorta | aortic insufficiency | |
| A7711 | f | 64 | aorta | AVR + aorta | coronary heart disease (unspecified) | |
| A7741 | f | 66 | aorta | AVR + aorta | aortic insufficiency | |
| A7857 | m | 55 | aorta | AVR + aorta | aortic insufficiency | |
| A8307 | m | 58 | aorta | AVR + aorta | aortic stenosis | |
| A8454 | m | 68 | aorta | AVR + aorta | aortic insufficiency | |
| A8684 | m | 56 | aorta | AVR + aorta | aortic insufficiency | |
| | 16m/4f | 55.5 (38 – 68) | | | | |
| A4419 | m | 51 | right atrium | AVR | aortic stenosis | *ARHGEF4, TFDP2, ASIC2, STX18-AS1, STX18, MSX1, LEPREL1 (=P3H2), WDR7* |
| A4438 | f | 64 | right atrium | AVR | aortic stenosis | |
| A4885 | m | 54 | right atrium | AVR | aortic stenosis | |
| A4968 | f | 55 | right atrium | AVR | aortic insufficiency | |
| A4990 | f | 54 | right atrium | AVR | aortic stenosis | |
| A5569 | f | 52 | right atrium | AVR | aortic stenosis | |
| A5629 | m | 71 | right atrium | AVR | aortic stenosis | |
| A5666 | f | 59 | right atrium | AVR | aortic stenosis | |
| A5945 | m | 52 | right atrium | AVR | combined aortic vitium | |
| A6115 | m | 58 | right atrium | AVR | aortic stenosis | |
| A6587 | m | 25 | right atrium | AVR | aortic insufficiency | |
| A6603 | m | 55 | right atrium | AVR | aortic stenosis | |
| | 7m/5f | 54.5 (25 – 71) | | | | |

[1] at the time of operation, CABG: coronary artery bypass graft, AVR: aortic valve replacement

#### Online Table IX. Candidate gene knockout mouse models

| gene | phenotype | reference |
|---|---|---|
| ***Macrod2*** | increase in intestinal tumorigenicity, no cardiac phenotype | Sakthianandeswaren et al., Cancer Disc., 8:988-1005 (2018); Lo Re et al., Front. Genet., 9:654 (2018) |
| ***Slc27a6*** | -- | |
| ***Arhgef4*** | atypical peripheral blood lymphocyte parameters (B cells ↓, granulocytes ↑) | Gerdin. Acta. Ophthalm., 88:925-927 (2010) |
| ***Gosr2*** | homozygous: (male/female) preweaning lethality, complete penetrance, heterozygous (male) abnormal gait | International Mouse Phenotyping Consortium |
| ***Asic2* (*LOC107985038*)** | abnormal blood vessel physiology, decreased vasoconstriction, hypertension | Gannon et al., Am. J. Physiol. Heart Circ. Physiol., 294:H1793-1803 (2008); Lu et al., Neuron, 64:885-897 (2009) |
| ***Stx18-AS1*** | -- | |
| ***P3h2* (= *Leprel1*)** | abnormal thrombosis, embryonic lethality during organogenesis (complete penetrance), embryonic lethality (incomplete penetrance) | Pokidysheva et al., Proc. Natl. Acad. Sci. USA, 111:161-166 (2014) |
| ***Wdr7*** | -- | |

#### Online Table X. Cross ethnical validation

| rsid | Lin et. al (2015) | | | | EU allele (+) | | EU all | | | EU septal defects | | | gene | cytoband |
|---|---|---|---|---|---|---|---|---|---|---|---|---|---|---|
| | ea | oa | or | *p* | ea | oa | beta | se | *p* | beta | se | *p* | | |
| rs1400558 | A | G | 1.15 | $1.63 \times 10^{-9}$ | t | c | 0.0096 | 0.0306 | 0.7547 | 0.0042 | 0.0515 | 0.9346 | EDNRA | 4q31.22 |
| rs7863990 | T | C | 1.34 | $3.71 \times 10^{-14}$ | t | c | 0.0315 | 0.0333 | 0.3437 | 0.0028 | 0.0564 | 0.9608 | SMARCA2 | 9p24.2 |
| rs2433752 | G | A | 0.83 | $1.04 \times 10\text{-}^{10}$ | a | g | -0.0146 | 0.0448 | 0.7452 | 0.0188 | 0.0763 | 0.8055 | TBX3–TBX5 | 12q24.13 |
| rs490514 | G | A | 1.19 | $1.20 \times 10^{-13}$ | t | c | -0.0909 | 0.0422 | 0.03117 | -0.1478 | 0.0691 | 0.03248 | PTPRT | 20q12 |

EU: DHM+UK meta-analysis, ea: effective allele, oa: other allele, or: odds ratio, *p*: *p* value, EU allele(+): eu allele in positive strand, beta: effective size, se: standard error

#### Online Table XI. Imputation score for all lead SNPs

| rsid | chr | pos_hg19 | A1 | A2 | DHM | UK |
|---|---|---|---|---|---|---|
| rs35437121 | 2 | 131769448 | T | C | 0.854281 | 0.871797 |
| rs114503684 | 3 | 141834969 | G | C | 0.816128 | 0.859228 |
| rs2046060 | 3 | 187852486 | G | A | 0.938114 | 0.979392 |
| rs187369228 | 3 | 189802439 | G | A | 0.761438 | 0.789581 |
| rs870142 | 4 | 4648047 | T | C | 0.990797 | genotyped |
| rs185531658 | 5 | 113136521 | C | T | 0.826139 | 0.806271 |
| rs185531658 | 5 | 113136521 | C | T | 0.826139 | 0.806271 |
| rs146300195 | 5 | 128326845 | A | G | 0.794733 | 0.880439 |
| rs117527287 | 6 | 85729959 | A | G | 0.737518 | 0.824389 |
| rs148563140 | 8 | 81475406 | T | C | 0.862684 | 0.911681 |
| rs11065987 | 12 | 112072424 | G | A | genotyped | genotyped |
| rs7982677 | 13 | 92988323 | A | C | 0.989636 | genotyped |
| rs138741144 | 17 | 32286564 | A | G | 0.785646 | 0.812021 |
| rs11874 | 17 | 45017193 | A | G | 0.990608 | 0.992829 |
| rs72917381 | 18 | 54546223 | T | C | 0.936111 | 0.949522 |
| rs150246290 | 20 | 15112880 | C | G | 0.823259 | 0.873741 |

#### Online Table XII. Genomic inflation

| | cases before imputation | controls after imputation | genomic inflation λ | |
|---|---|---|---|---|
| all CHD | 1,495 | 3,554 | 1.032 | 1.041 |
| septal defects | 445 | 3,554 | 1.023 | 1.036 |
| ASD | 232 | 3,554 | 1.014 | 1.039 |
| right heart lesions | 270 | 3,554 | 1.017 | 1.034 |
| left heart lesions | 138 | 3,554 | 1.011 | 1.022 |
| transposition of the great arteries | 146 | 3,554 | 1.024 | 1.032 |
| anomalies of thoracic arteries and veins | 205 | 3,554 | 1.007 | 1.028 |

### Online Table XIII. Power analysis

| | OR | alpha | Power with n cases and 8486 controls, MAF=0.05, prevalance=9/1000, genotyping error=0.001 | | | | | | | | | | | | | |
|---|---|---|---|---|---|---|---|---|---|---|---|---|---|---|---|---|
| | | | n=100 | n=200 | n=300 | n=326 | n=399 | n=453 | n=486 | n=500 | n=1074 | n=1296 | n=1500 | n=3000 | n=4034 | n=5000 |
| suggestive significance | 1.2 | $1x10^{-6}$ | $2.39x10^{-5}$ | $2.07x10^{-4}$ | $4.43x10^{-4}$ | $7.53x10^{-4}$ | 0.001243 | 0.001012 | 0.001751 | 0.001714 | 0.019214 | 0.031534 | 0.051429 | 0.265726 | 0.428973 | 0.553271 |
| | 1.5 | $1x10^{-6}$ | $3.70x10^{-3}$ | $2.67x10^{-2}$ | $1.09x10^{-1}$ | $1.55x10^{-1}$ | 0.231282 | 0.309943 | 0.376382 | 0.399426 | 0.957304 | 0.986148 | 0.996780 | 1 | 1 | 1 |
| | 2.0 | $1x10^{-6}$ | $1.59x10^{-1}$ | $7.95x10^{-1}$ | $9.69x10^{-1}$ | $9.86x10^{-1}$ | 0.997805 | 0.999608 | 0.999859 | 0.999934 | 1 | 1 | 1 | 1 | 1 | 1 |
| | 2.5 | $1x10^{-6}$ | $8.03x10^{-1}$ | $9.98x10^{-1}$ | $9.99x10^{0}$ | $9.99x10^{0}$ | 1 | 1 | 1 | 1 | 1 | 1 | 1 | 1 | 1 | 1 |
| | 3.0 | $1x10^{-6}$ | $9.90x10^{-1}$ | $9.99x10^{0}$ | $9.99x10^{0}$ | $9.99x10^{0}$ | 1 | 1 | 1 | 1 | 1 | 1 | 1 | 1 | 1 | 1 |
| genome-wide significance | 1.2 | $5x10^{-8}$ | $1.86x10^{-6}$ | $2.15x10^{-5}$ | $5.14x10^{-5}$ | $9.46x10^{-5}$ | 0.000169 | 0.000133 | 0.000251 | 0.000245 | 0.004270 | 0.007795 | 0.014232 | 0.117917 | 0.230057 | 0.335148 |
| | 1.5 | $5x10^{-8}$ | $6.02x10^{-4}$ | $6.37x10^{-3}$ | $3.68x10^{-2}$ | $5.78x10^{-2}$ | 0.097780 | 0.145557 | 0.190877 | 0.207678 | 0.877087 | 0.949684 | 0.984798 | 0.999996 | 1 | 1 |
| | 2.0 | $5x10^{-8}$ | $5.96x10^{-2}$ | $6.04x10^{-1}$ | $9.05x10^{-1}$ | $9.50x10^{-1}$ | 0.988961 | 0.997435 | 0.998934 | 0.999451 | 1 | 1 | 1 | 1 | 1 | 1 |
| | 2.5 | $5x10^{-8}$ | $6.15x10^{-1}$ | $9.91x10^{-1}$ | $9.99x10^{0}$ | $9.99x10^{0}$ | 1 | 1 | 1 | 1 | 1 | 1 | 1 | 1 | 1 | 1 |
| | 3.0 | $5x10^{-8}$ | $9.60x10^{-1}$ | $9.99x10^{0}$ | $9.99x10^{0}$ | $9.99x10^{0}$ | 1 | 1 | 1 | 1 | 1 | 1 | 1 | 1 | 1 | 1 |

#### Online Table XIV. Primers used for qRT-PCR analyses and genotyping

| primer | sequence | length of amplicon |
|---|---|---|
| hu MACROD2_F1100 | 5’ CAG ATG GTG TCA ACA CTG TCA CT 3’ | |
| hu MACROD2_R1190 | 5’ TTT TCA TCC TTT GCA AAA TCT TC 3’ | 91 bp |
| hu GOSR2_F71 | 5’ GGC GAC ATG GAT CCC CTG T 3’ | |
| hu GOSR2_R171 | 5’ ATG TGC ACA GAC TGC TTG TCT 3’ | 101 bp |
| hu WNT3_F1221 | 5’ CCT GCA AGT AGG GCA CCA 3’ | |
| hu WNT3_R1333 | 5’ TCT GAC GCT GAG GGC TGT 3’ | 108 bp |
| hu ARHGEF4_F4853 | 5’ CTG TAA GAA GGA CCT GCT CCG 3’ | |
| hu ARHGEF4_R4936 | 5’ GTC TCT GTC CTT CCC GTC CT 3’ | 84 bp |
| hu ASIC2_F1032 | 5’ AGC CAC CTT TCA TCC AAG AG 3’ | |
| hu ASIC2_R1126 | 5’ TGG GGG CAG GTA TGT GAG 3’ | 95 bp |
| hu MSX1_F661 | 5´ GAC CCC GTG GAT GCA GAG 3´ | |
| hu MSX1_R752 | 5´ GGT TCG TCT TGT GTT TGC GG 3´ | 92 bp |
| hu P3H2_F1278 | 5’ GCC TCT CTC CCA TCG AGA AT 3’ | |
| hu P3H2_R1366 | 5’ CAG GGC TTT CAC ATA CTC ACC 3’ | 89 bp |
| hu SLC27A6_F1446 | 5’ TTA CAT TTT TAC CTC TGG AAC AAC A 3’ | |
| hu SLC27A6_R1555 | 5’ CAT GAG CAG TAC AAC CAA AAG C 3’ | 110 bp |
| hu STX18_F986 | 5’ TCA AGG AAG GCA ACG AAG AC 3’ | |
| hu STX18_R1080 | 5’ GGA GAA GGA GCA CAT CAC G 3’ | 94 bp |
| hu STX18-AS1_F455 | 5’ GCC ATC CCT AAG ACA GCA AG 3’ | |
| hu STX18-AS1_R540 | 5’ GCT TAG ACT CTC TGA ATC TCT GCA T 3’ | 86 bp |
| hu TFDP2_F960 | 5’ GCT GGT GTC AGA GTT CAC CA 3’ | |
| hu TFDP2_R1041 | 5’ TCT TCG CCT AAT GTT CTT CTG A 3’ | 82 bp |
| hu WDR7_F1913 | 5’ TGG AGG CCT TCT GAT GAT TAC 3’ | |
| hu WDR7_R2006 | 5’ TCA CAC AAC GAT CCA ATG C 3’ | 94 bp |
| hu OCT4_F268 | 5’ GGG ATG GCG TAC TGT GGG 3’ | |
| hu OCT4_R416 | 5’ GCA CCA GGG GTG ACG GTG 3’ | 149 bp |
| hu KLF4_F1441 | 5’ TCT TCG TGC ACC CAC TTG GG 3’ | |
| hu KLF4_R1574 | 5’ CTG CTC AGC ACT TCC TCA AG 3’ | 134 bp |
| hu SOX2_F840 | 5’ ACA GCT ACG CGC ACA TGA 3’ | |
| hu SOX2_R908 | 5’ GGT AGC CCA GCT GCT CCT 3’ | 69 bp |
| hu CMYC_F1311 | 5’ CAC CAG CAG CGA CTC TGA 3’ | |
| hu CMYC_R1412 | 5’ GAT CCA GAC TCT GAC CTT TTG C 3’ | 102 bp |
| hu NANOG_F253 | 5’ TGC TTT GAA GCA TCC GAC TGT 3’ | |
| hu NANOG_R446 | 5’ GGT TGT TTG CCT TTG GGA CTG 3’ | 194 bp |
| hu REX1_F292 | 5‘ AGT AGT GCT CAC AGT CCA GCA G 3’ | |
| hu REX1_R397 | 5‘ TGT GCC CTT CTT GAA GGT TT 3’ | 106 bp |
| mu Gosr2_F217 | 5’ TGG AGA TTT TGT CAA GCA AGG A | |
| mu Gosr2_R306 | 5’ GCA AGT GCT GGA CAT CAT ACT | 90 bp |
| mu Macrod2_F264 | 5’ AAA AGG TGT GGA GAG AGG AGA A | |
| mu Macrod2_R371 | 5’ TCC ATG ATA GAA TGG TGC TCA | 108 bp |
| mu Msx1_F659 | 5’ GCC CCG AGA AAC TAG ATC G 3’ | |
| mu Msx1_R767 | 5’ TTG GTC TTG TGC TTG CGT AG 3’ | 109 bp |
| mu Wnt3_F107 | 5’ CTG CTC GGG CTG CTA CTC | |
| mu Wnt3_R211 | 5’ GGC CAG AGA TGT GTA CTG CTG | 105 bp |
| SeV_F | 5’ GGA TCA CTA GGT GAT ATC GAG C 3’ | |
| SeV_R | 5’ ACC AGA CAA GAG TTT AAG AGA TAT GTA TC 3’ | 181 bp |
| hu β-actin_F382 | 5’ CCA ACC GCG AGA AGA TGA 3’ | |
| hu β-actin_R478 | 5’ CCA GAG GCG TAC AGG GAT AG 3’ | 97 bp |
| MACROD2_SNP280_F | 5’ ACA AGG AAC CAC TAA ACA GCT T 3’ | |
| MACROD2_SNP280_R2 | 5’ GAG CTA TTT CAA TTA TGT TAC TC 3’ | 340 bp |
| MACROD2_SNP290_F | 5’ AGT GAA CAA TAA GGT AAA CAG TTA 3’ | |
| MACROD2-SNP290_R2 | 5’ ATC CAT CCA ACC AAC TGG CTA 3’ | 409 bp |
| MACROD2_SNP721_F | 5’ TGT GTG GTT ATG ACA TTT GTC C 3’ | |
| MACROD2_SNP721_R | 5’ CTG TTC AAT TTG CCC CTG CTA 3’ | 500 bp |
| MACROD2_SNP733_F | 5’ TGA GTG AGT CTC CTG TAA GCA 3’ | |
| MACROD2_SNP733_R | 5’ CAA CAA TAT GAC TTG CAG GTG T 3’ | 393 bp |
| GOSR2_SNP363_F | 5’ GGT GTC TCA CAG CCT GAC TA 3’ | |
| GOSR2_SNP363_R | 5’ ATG CTA TTC ACT CAG CCT TGT A 3’ | 467 bp |

| primer | sequence | length of amplicon |
|---|---|---|
| GOSR2_SNP446_F | 5’ CTA CCT GAG ACT GGA ACA TCA 3’ | |
| GOSR2_SNP446_R | 5’ CGA GAT CGG CTT CCG CTT C 3’ | 365 bp |
| GOSR2_SNP874_F | 5’ CCA GTG TGG GCA GAT CCC A 3’ | |
| GOSR2_SNP874_R | 5’ ACA AGA AAC TAA CAG AGC AGG 3’ | 371 bp |

#### Online Figure I. SNPs associated with right heart lesions

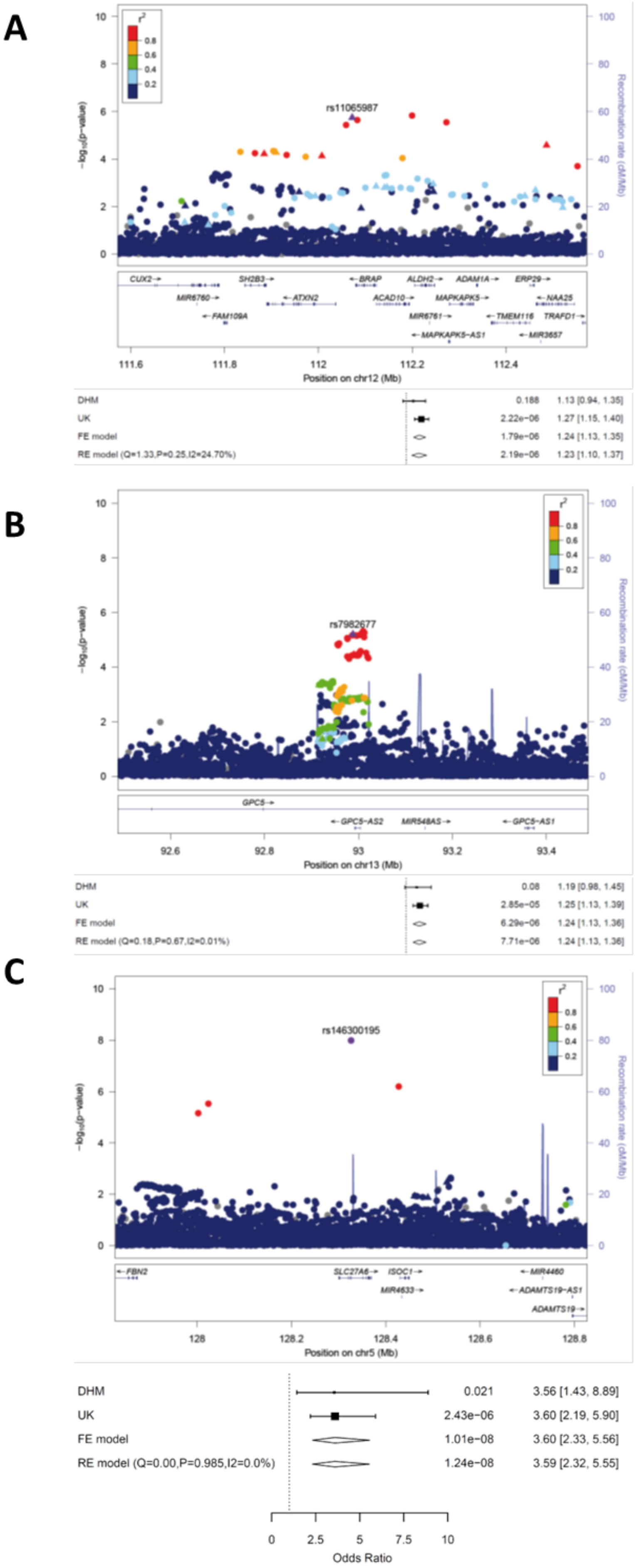

**A:** LocusZoom plot of the genomic region of rs11065987 on chromosome 12. **B:** LocusZoom plot of the genomic region of rs7982677 on chromosome 13. **C:** LocusZoom plot of the *SLC27A6* region on chromosome 5. The index SNP is indicated as purple diamonds and the other SNPs are color coded depending on their degree of correlation ($r^2$). Circles represent imputed SNPs, triangles genotyped SNPs. FE: fixed effects, RE: random effects.

### Online Figure II. SNPs associated with left heart lesions

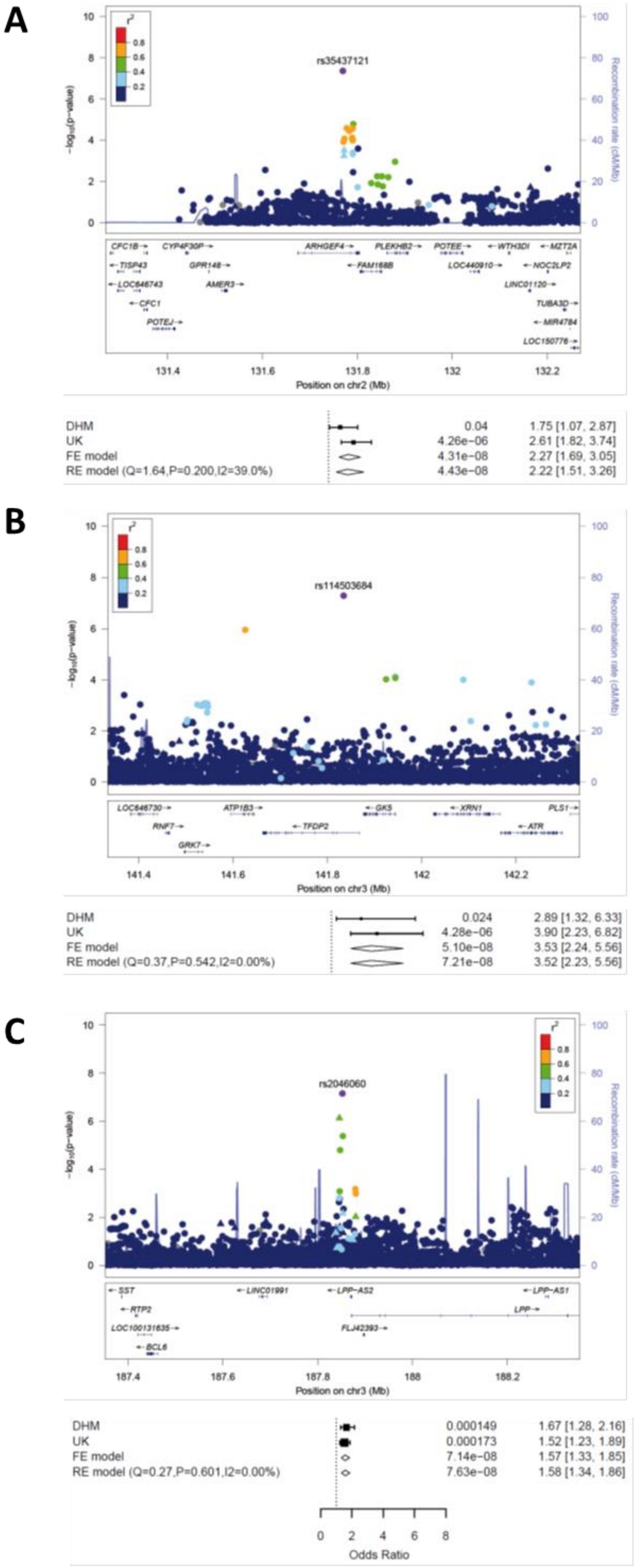

**A:** LocusZoom plot of the *ARHGEF4* region on chromosome 2. **B:** LocusZoom plot of the *TFDP2* region on chromosome 3. **C:** LocusZoom plot of the *FLJ42993* region on chromosome 3.The index SNPs are indicated as purple diamonds and the other SNPs are color coded depending on their degree of correlation ($r^2$). Circles represent imputed SNPs, triangles genotyped SNPs. FE: fixed effects, RE: random effects.

**Online Figure III. SNPs associated with septal defects**

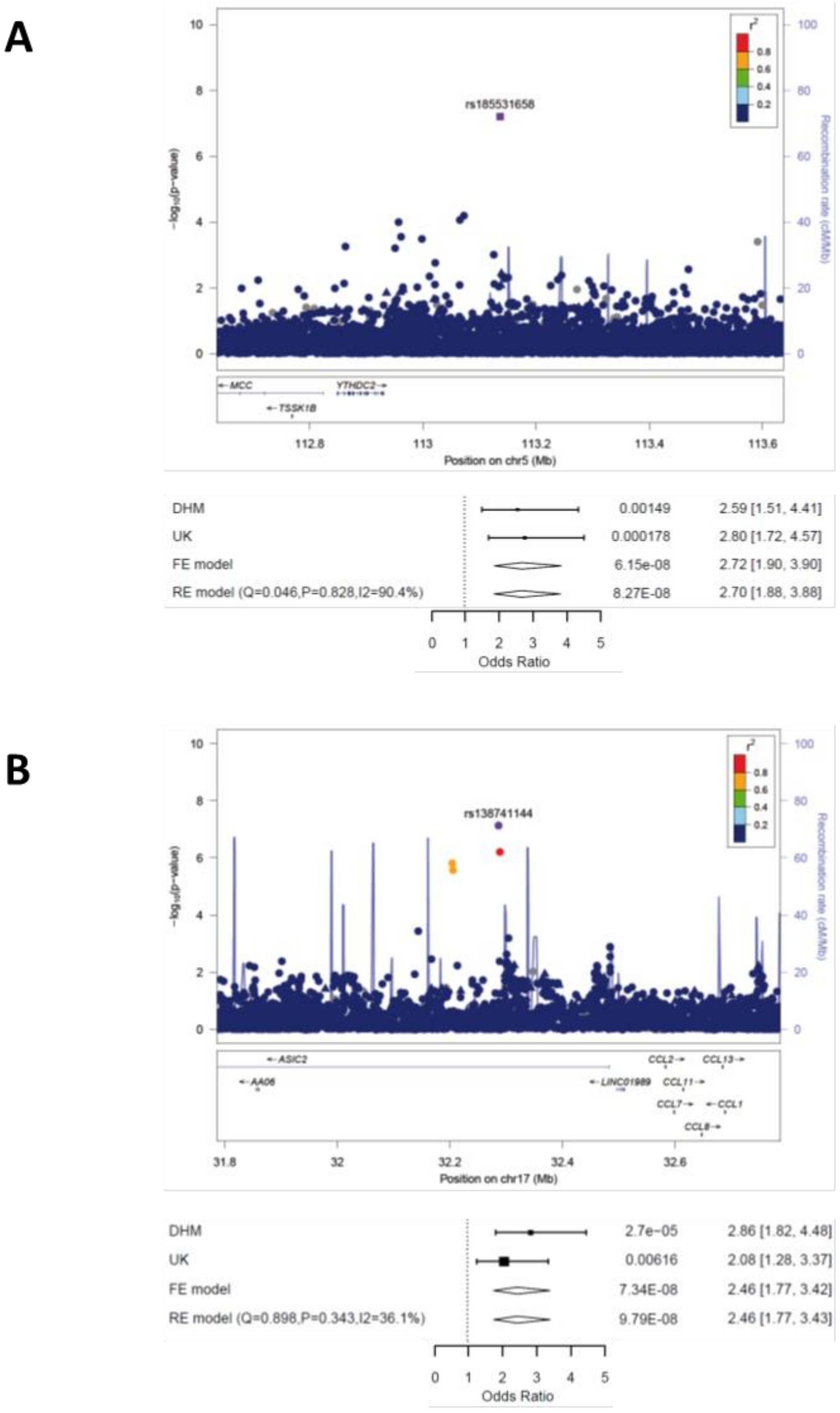

**A:** LocusZoom plot of the region on chromosome 5. **B:** LocusZoom plot of the region on chromosome 17. The index SNP is indicated as purple diamonds and the other SNPs are color coded depending on their degree of correlation ($r^2$). Circles represent imputed SNPs, triangles genotyped SNPs. FE: fixed effects, RE: random effects.

#### Online Figure IV. SNPs associated with ASD

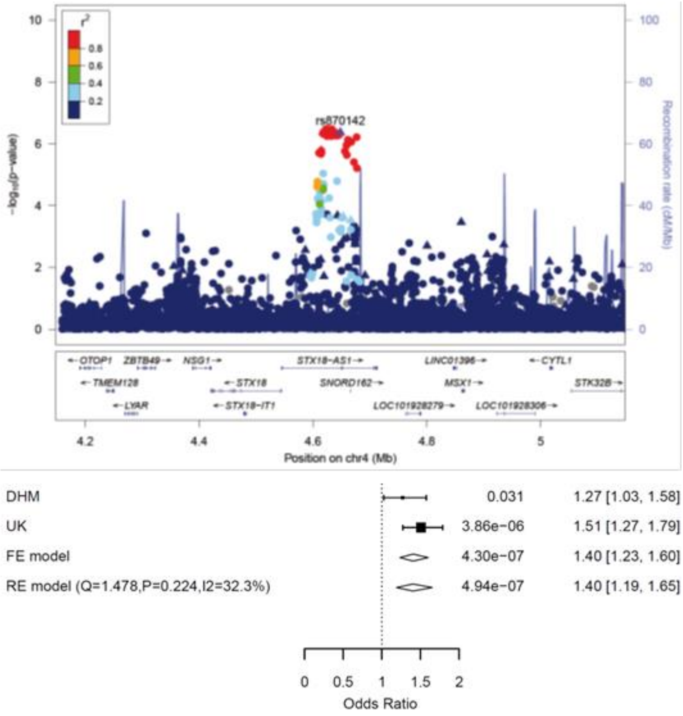

LocusZoom plot of the *SNORD162* region on chromosome 4. The index SNP is indicated as purple diamonds and the other SNPs are color coded depending on their degree of correlation ($r^2$). Circles represent imputed SNPs, triangles genotyped SNPs. FE: fixed effects, RE: random effects.

**Online Figure V. SNPs associated with ASDII**

A

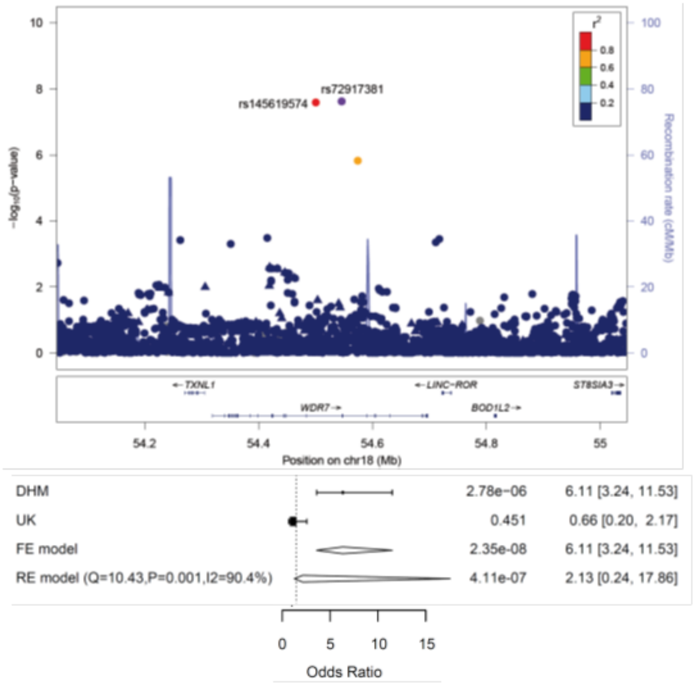

B

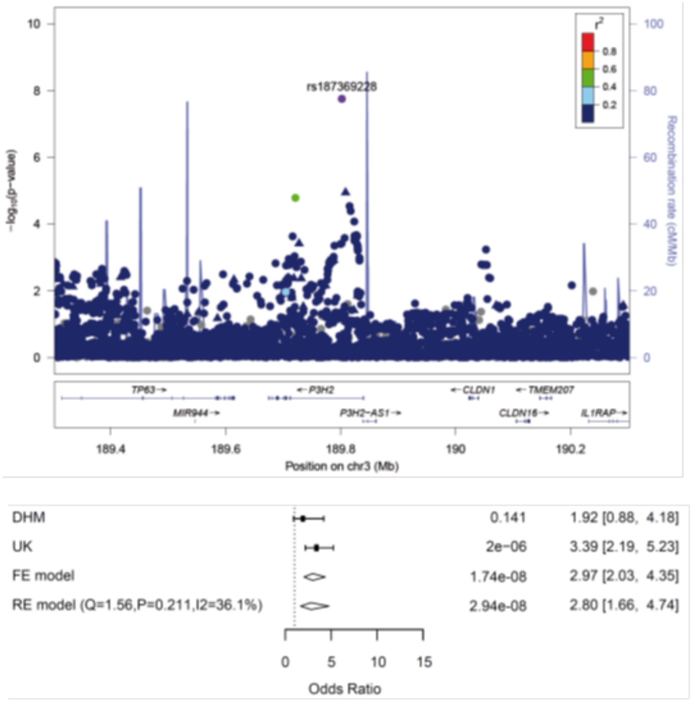

**A:** LocusZoom plot of the *WDR7* region on chromosome 18. **B:** LocusZoom plot of the *P3H2* (= *LEPREL1*) region on chromosome 3.The index SNPs are indicated as purple diamonds and the other SNPs are color coded depending on their degree of correlation ($r^2$). Circles represent imputed SNPs, triangles genotyped SNPs. FE: fixed effects, RE: random effects.

**Online Figure VI. SNPs associated with anomalies of thoracic arteries and veins**

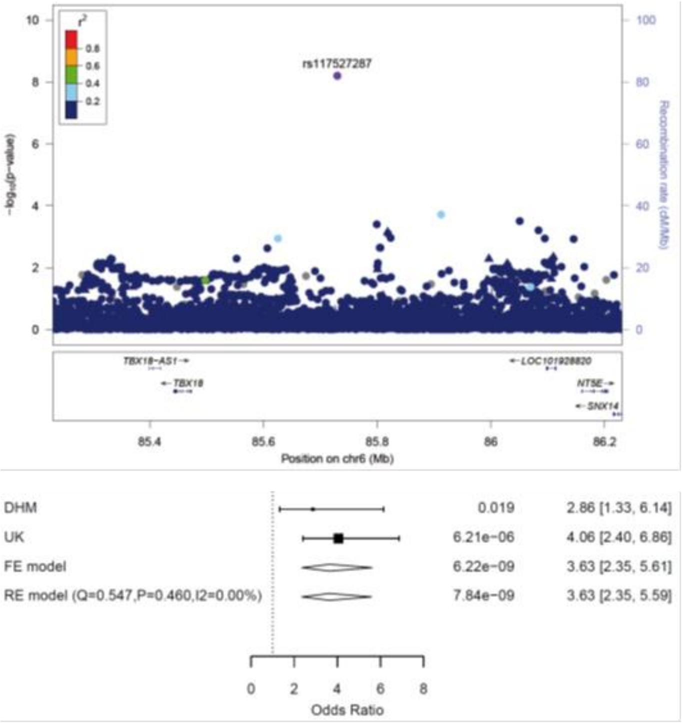

LocusZoom plot of the region on chromosome 6. The index SNP is indicated as purple diamonds and the other SNPs are color coded depending on their degree of correlation ($r^2$). Circles represent imputed SNPs, triangles genotyped SNPs. FE: fixed effects, RE: random effects.

#### Online Figure VII. Participation of CHD-associated SNP-carrying genes in signaling cascades

| Entrez Gene Id | Gene Symbol | GO_CELL_CELL_SIGNALING | GO_EMBRYONIC_ORGAN_DEVELOPMENT | GO_REPRODUCTIVE_SYSTEM_DEVELOPMENT | GO_ANATOMICAL_STRUCTURE_FORMATION_INVOLVED_IN_MORPHOGENESIS | GO_EMBRYONIC_MORPHOGENESIS | GO_MALE_SEX_DIFFERENTIATION | GO_REGULATION_OF_CYTOKINE_PRODUCTION | GO_EYE_DEVELOPMENT | GO_EMBRYO_DEVELOPMENT | GO_DEVELOPMENTAL_PROCESS_INVOLVED_IN_REPRODUCTION | Entrez | Source | Gene Description |
|---|---|---|---|---|---|---|---|---|---|---|---|---|---|---|
| 2625 | GATA3 | ■ | ■ | ■ | ■ | ■ | ■ | ■ | ■ | ■ | ■ | | S | GATA binding protein 3 |
| 2626 | GATA4 | ■ | ■ | ■ | ■ | ■ | ■ | ■ | | ■ | ■ | | S | GATA binding protein 4 |
| 7484 | WNT9B | ■ | ■ | ■ | ■ | ■ | ■ | | ■ | ■ | ■ | | S | wingless-type MMTV integration site family, member 9B |
| 3624 | INHBA | ■ | | ■ | ■ | ■ | ■ | ■ | ■ | ■ | ■ | | S | inhibin, beta A |
| 2554 | GABRA1 | ■ | | | | | | | | | | | S | gamma-aminobutyric acid (GABA) A receptor, alpha 1 |
| 1740 | DLG2 | ■ | | | | | | | | | | | S | discs, large homolog 2 (Drosophila) |
| 2914 | GRM4 | ■ | | | | | | | | | | | S | glutamate receptor, metabotropic 4 |
| 2559 | GABRA6 | ■ | | | | | | | | | | | S | gamma-aminobutyric acid (GABA) A receptor, alpha 6 |
| 9394 | HS6ST1 | | ■ | ■ | ■ | | | | | ■ | ■ | | S | heparan sulfate 6-O-sulfotransferase 1 |
| 26146 | TRAF3IP1 | | ■ | | ■ | ■ | | ■ | ■ | ■ | | | S | TNF receptor-associated factor 3 interacting protein 1 |
| 2201 | FBN2 | | ■ | | ■ | ■ | | | ■ | ■ | | | S | fibrillin 2 |
| 1508 | CTSB | | | ■ | | | | | | | ■ | | S | cathepsin B |
| 2324 | FLT4 | | | | ■ | | | ■ | | | | | S | fms-related tyrosine kinase 4 |
| 9698 | PUM1 | | | | | | | ■ | | | | | S | pumilio homolog 1 (Drosophila) |

**Online Figure VIII. Expression of candidate genes during cardiac differentiation of human iPS cells**

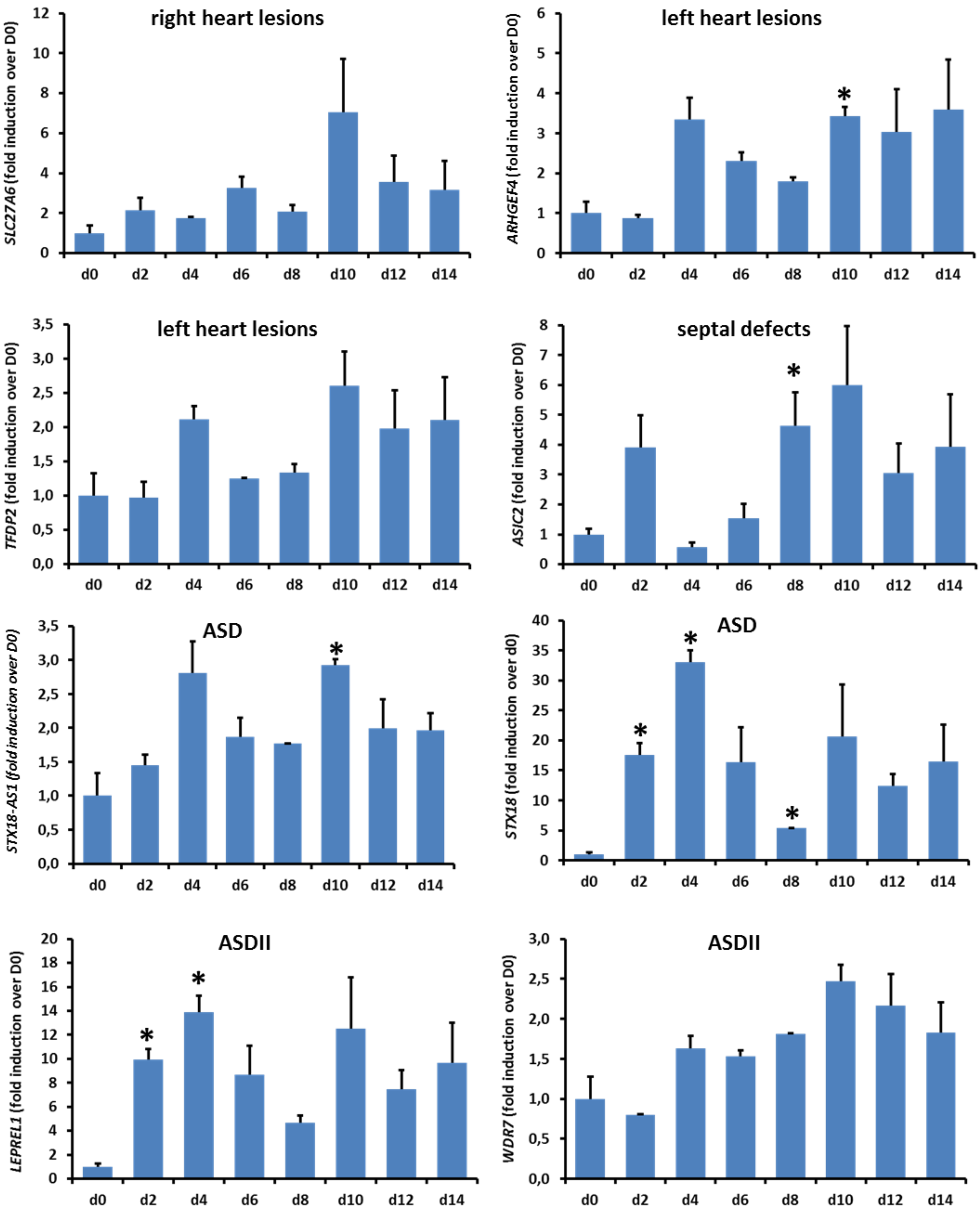

Expression of candidate genes during directed cardiac differentiation of human iPS cells. Significances were calculated *vs.* gene expression on D0. * $p < 0.05$.

**Online Figure IX. Expression of *MACROD2*, *GOSR2* and *WNT3* in patient tissue with or without risk variant**

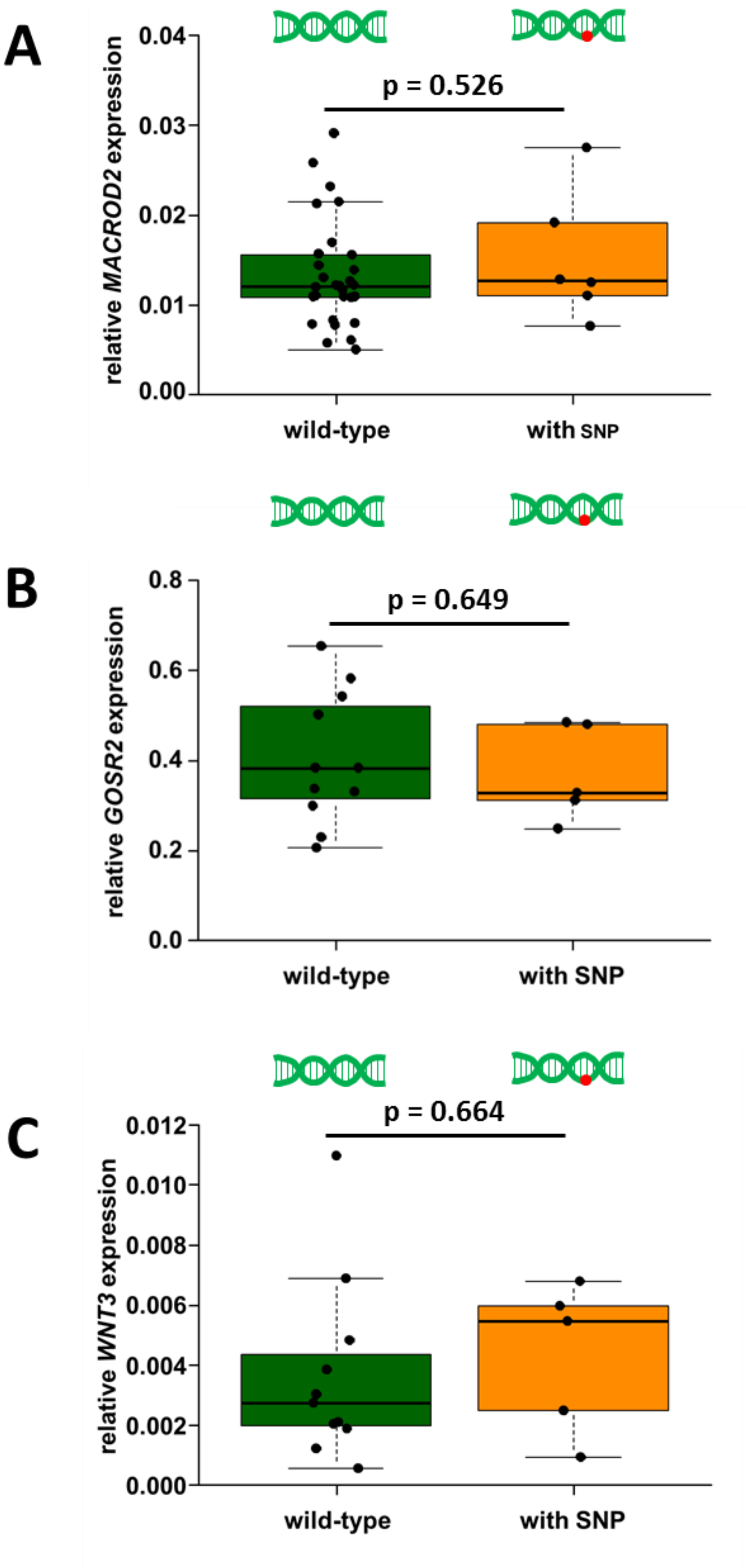

**A:** Expression of *MACROD2* in aortic tissue of pediatric TGA patients with wild-type (n=29) or heterozygous SNP genotype (n=6). **B:** Expression of *GOSR2* in tissue of pediatric patients with anomalies of thoracic arteries and veins with wild-type (n=11) or heterozygous SNP genotype (n=5). **C:** Expression of *WNT3* in tissue of pediatric patients with anomalies of thoracic arteries and veins with wild-type (n=11) or heterozygous SNP genotype (n=5).

**Online Figure X. Expression of candidate genes in pediatric and adult aortic tissue**

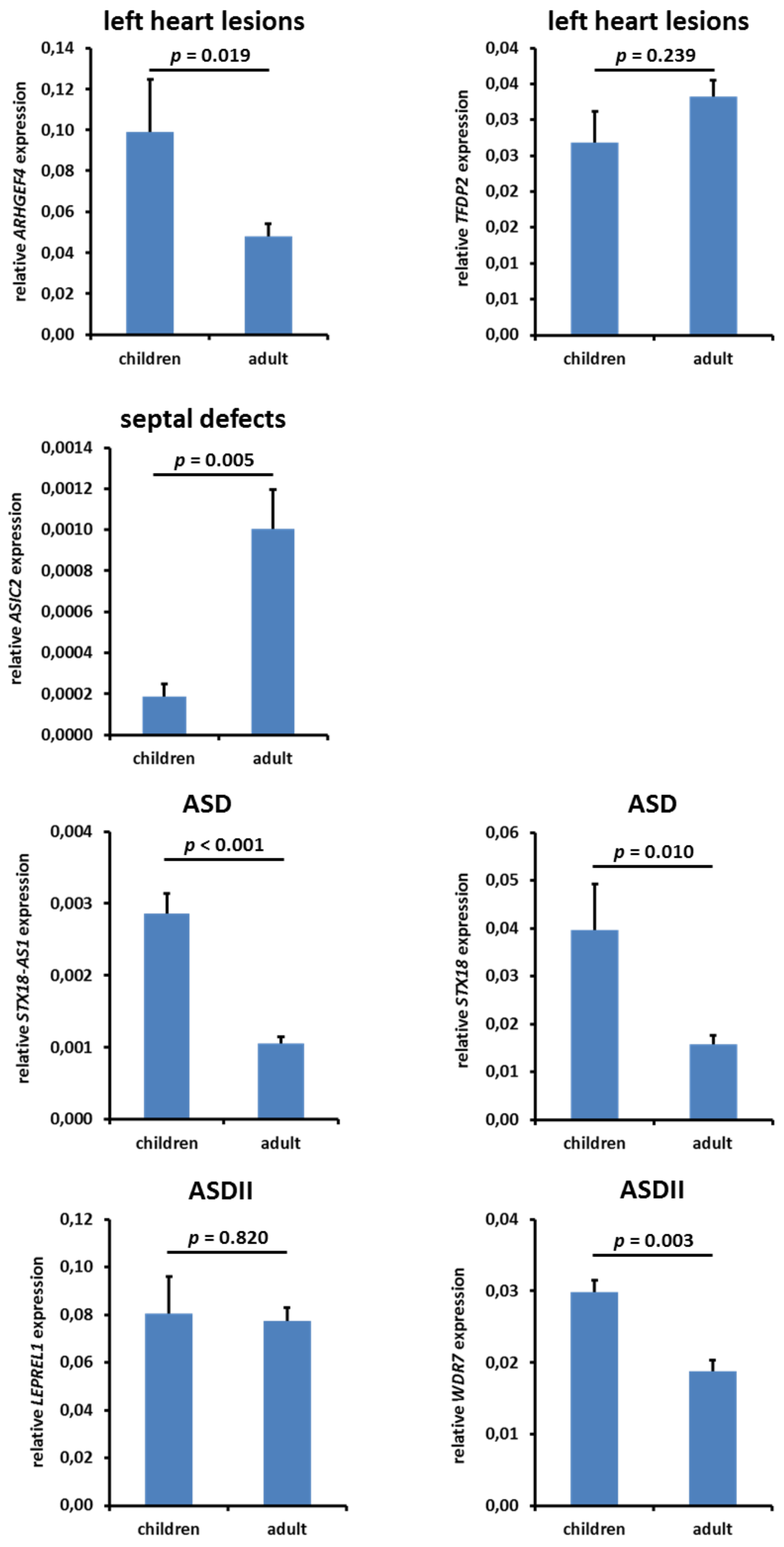

Expression of candidate genes in aortic tissue of CHD patients and adult surgical patients. The number of samples used for each gene are listed in Table VII (CHD patients) and Table VIII (adult surgical patients).

**Online Figure XI. Expression of candidate genes in pediatric and adult atrial tissue**

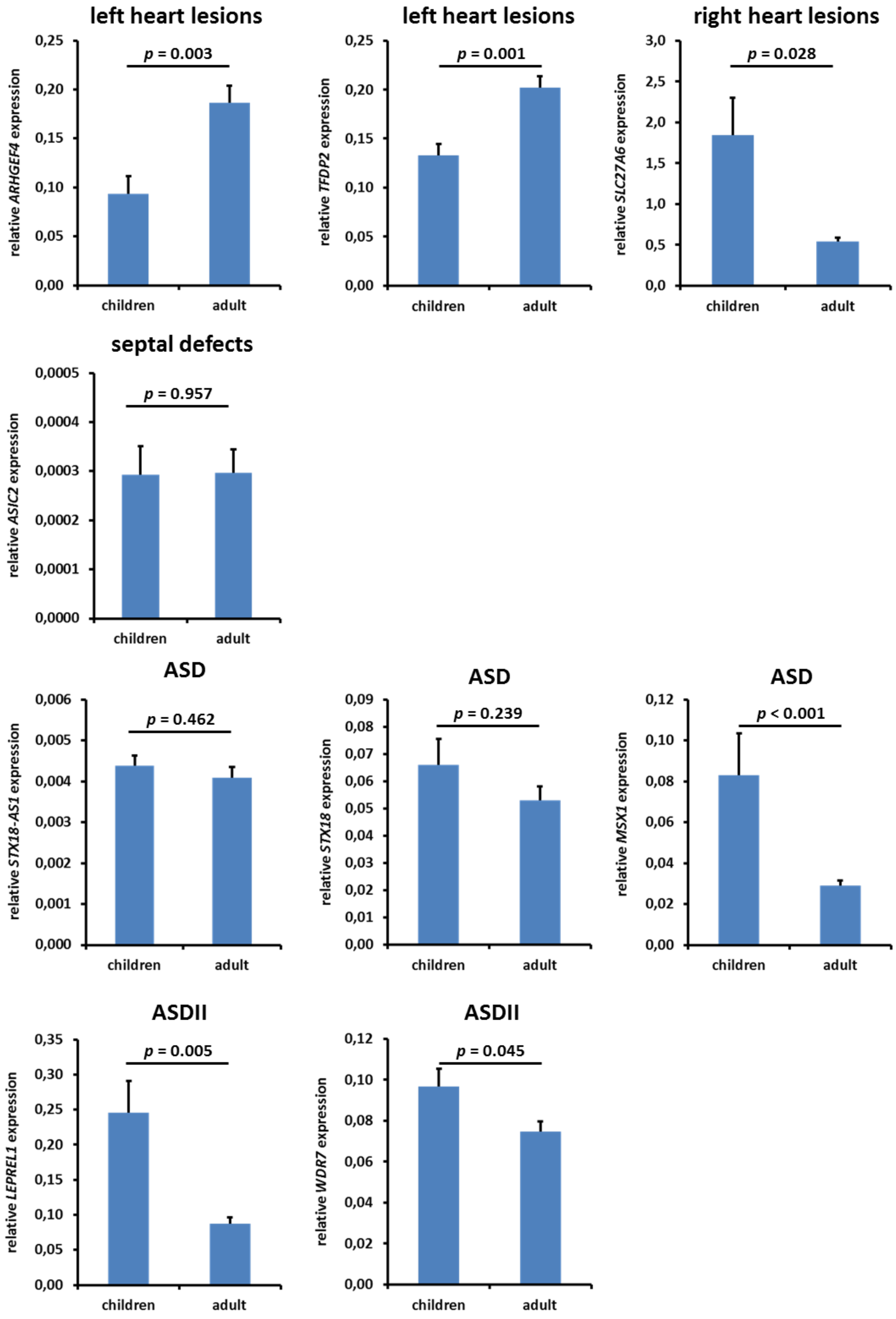

Expression of candidate genes in aortic tissue of CHD patients and adult surgical patients. The number of samples used for each gene are listed in Table VII (CHD patients) and Table VIII (adult surgical patients).

**Online Figure XII. Identification of different cell types after single-cell RNAseq by expression of defined marker genes**

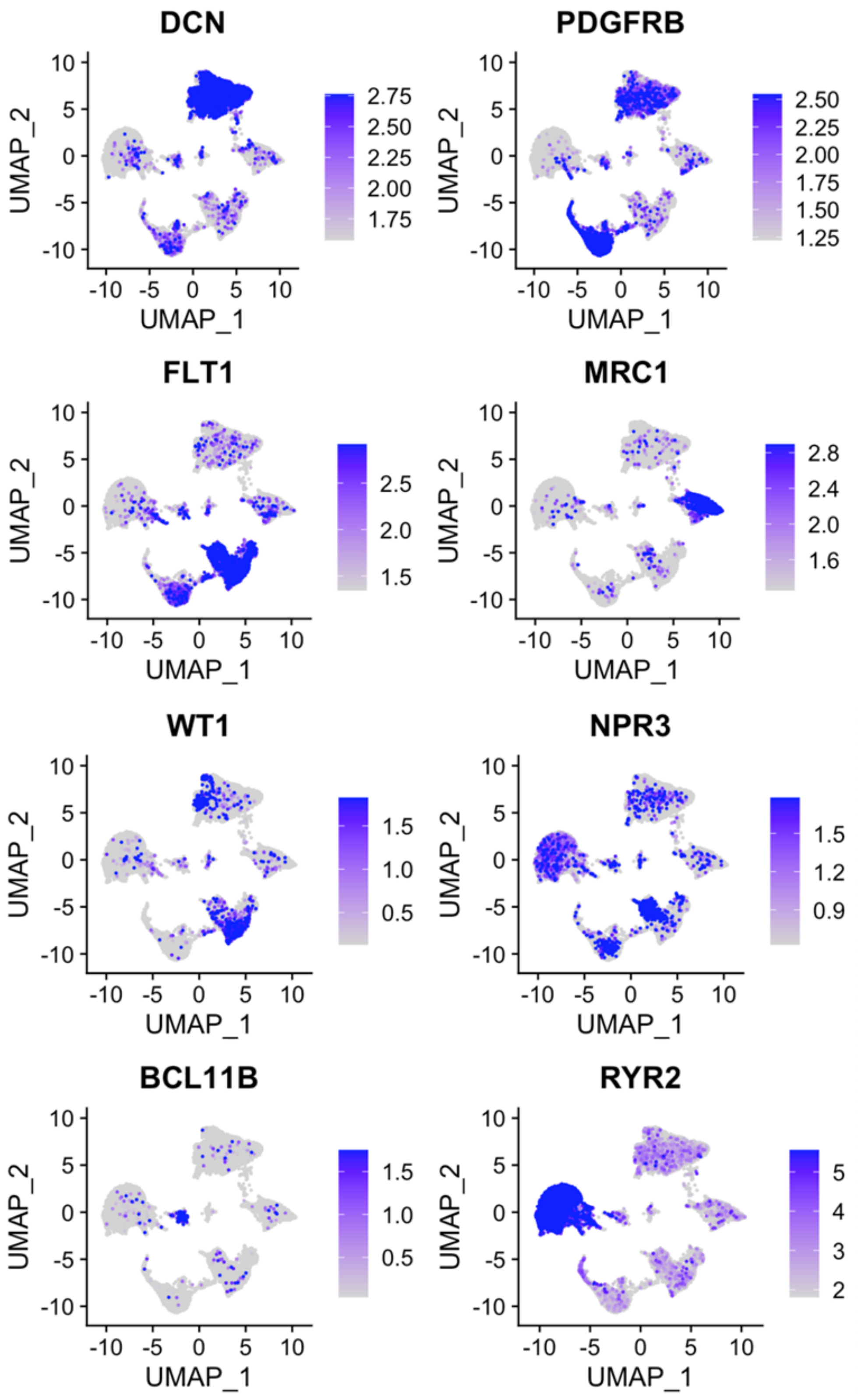

*DCN* (decorin), *PDGFRB* (platelet-derived growth factor receptor β), *FLT1* (fms related receptor kinase 1), *MRC1* (mannose receptor C-type 1), *WT1* (WT1 transcription factor), *NPR3* (natriuretic peptide receptor 3), *BCL11B* (BAF chromatin remodeling complex subunit BCL11B), *RYR2* (ryanodine receptor 2)

**Online Figure XIII. Workflow for general GWAS quality control**

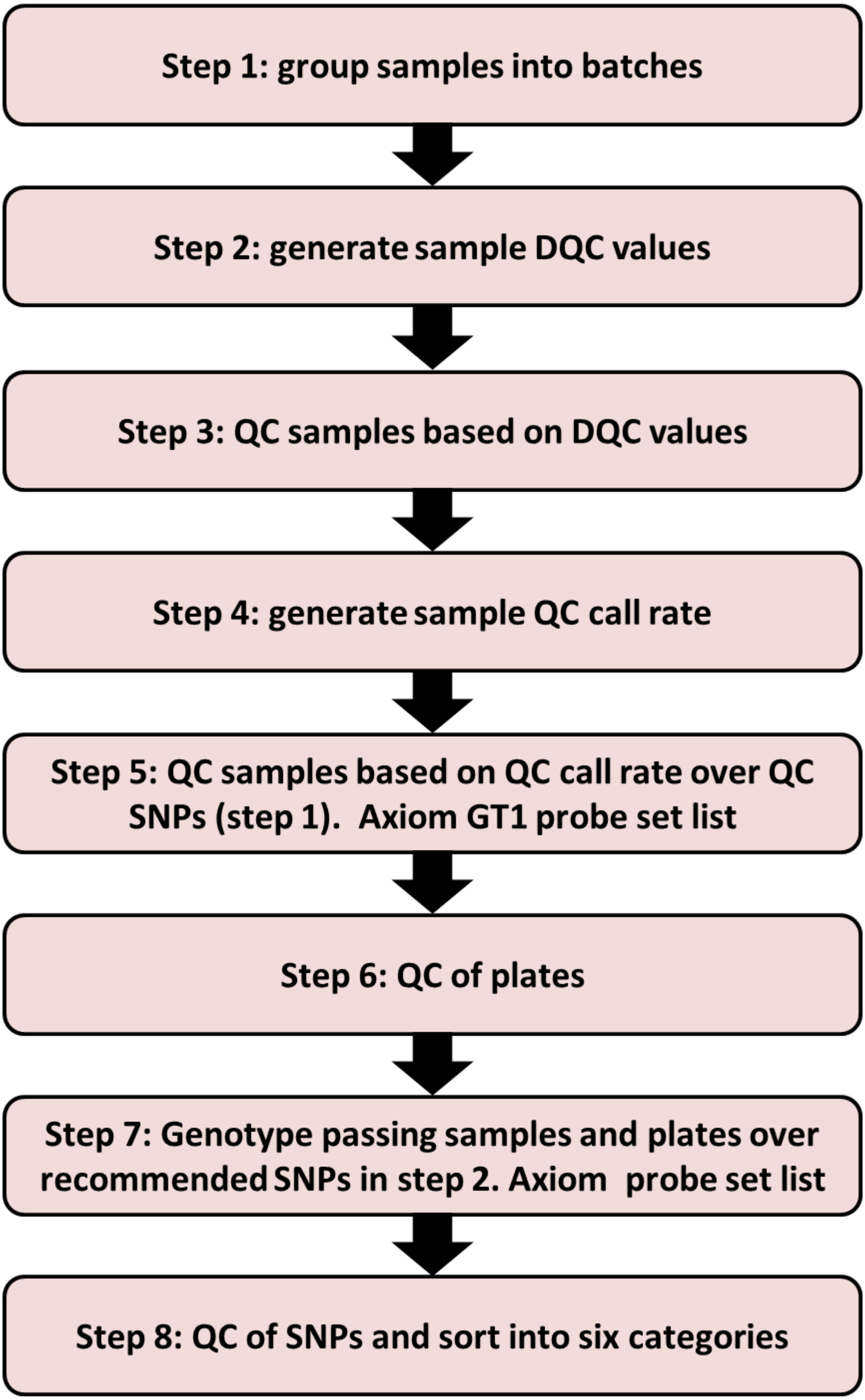

Application of these filters resulted in 1,495 cases, 3,554 controls and 432,097 variants. The genomic inflation λ is 1.03.

**Online Figure XIV. Quality control steps and PCA plots to analyze population stratification**

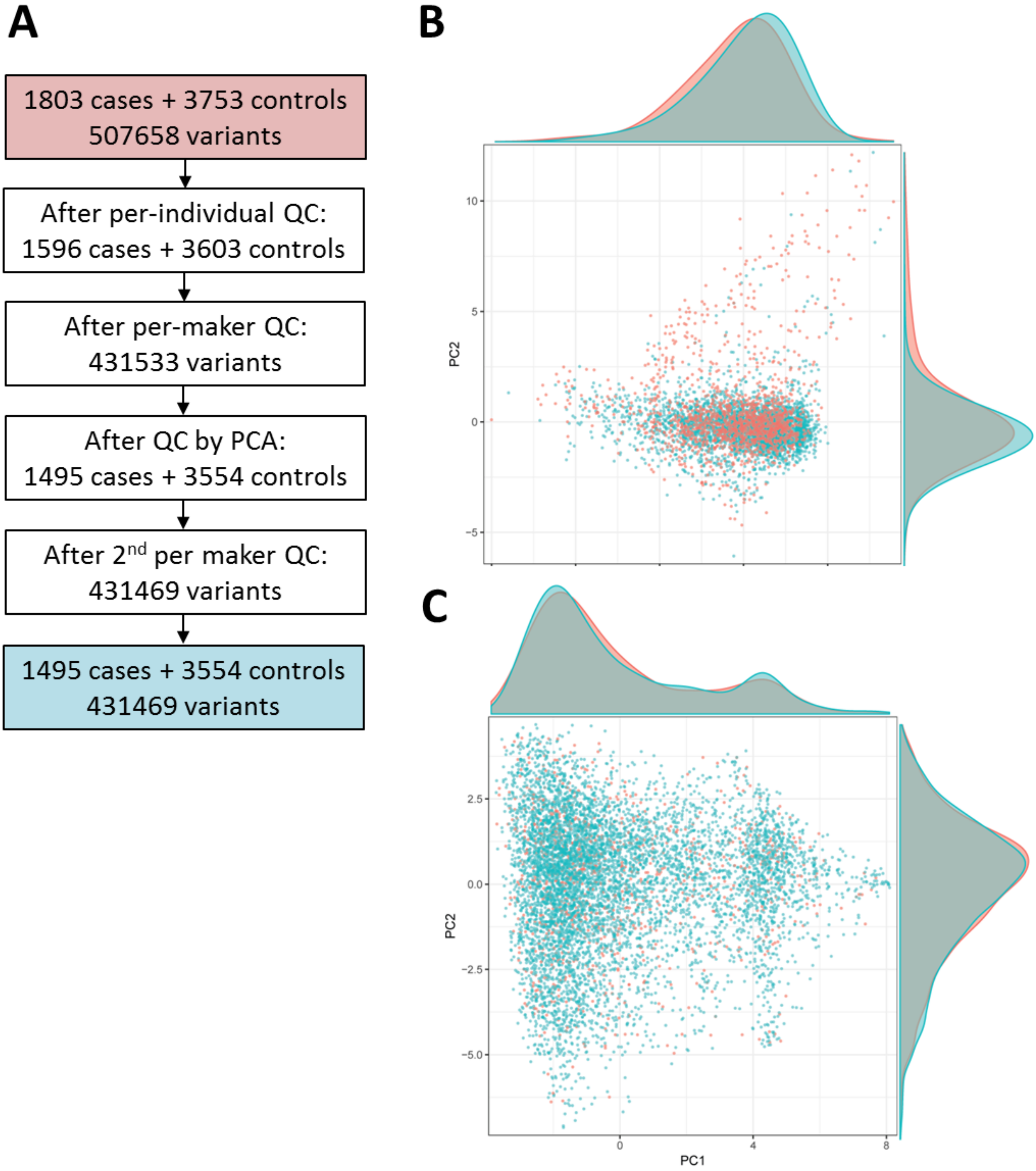

**A:** Quality control steps. **B, C:** Scatter plots with density plots in the margins for PC1 and PC2. The red dots are cases and the blue ones are the controls. **B:** DHM cohort. **C:** British cohort.

**Genes up-regulated in cardiomyocytes**

| Gene | d9-11 (1) | d9-11 (2) | d9-11(3) | d9-11(4) | newborn(1) | newborn(2) | newborn(3) | adult(1) | adult(2) | adult(3) | Gene | log2(fold) | FDR(*p*-value) | | |
|---|---|---|---|---|---|---|---|---|---|---|---|---|---|---|---|
| Ckm | 0,762222 | 1,00266 | 2,75663 | 3,32636 | 1585,26 | 2007,83 | 2208,66 | 2493,34 | 2055,72 | 2211,69 | Ckm | 9,492061172 | 6,60E-136 | | |
| Rbm20 | 17,3695 | 15,6287 | 19,9759 | 18,0303 | 136,358 | 123,718 | 114,601 | 55,8275 | 77,4142 | 72,2856 | Rbm20 | 3,143447154 | 9,12E-74 | | **Fold Change > 2.0** |
| Ehd4 | 38,8705 | 48,287 | 55,9182 | 42,1587 | 276,001 | 284,594 | 292,549 | 268,136 | 256,38 | 249,035 | Ehd4 | 2,895623802 | 5,11E-72 | | ***p* < 0.05** |
| Slc20a2 | 15,3726 | 17,4928 | 17,9115 | 16,4927 | 107,928 | 92,2464 | 90,0097 | 124,158 | 126,82 | 118,088 | Slc20a2 | 2,87918691 | 1,10E-68 | | **n=1155** |
| Pdzd2 | 1,63664 | 1,34168 | 2,0589 | 2,06727 | 31,3618 | 26,5093 | 24,5457 | 5,73279 | 8,87292 | 8,34767 | Pdzd2 | 4,166179498 | 3,16E-65 | | |
| Pnpla2 | 14,9745 | 13,5308 | 19,5362 | 15,4656 | 147,598 | 132,457 | 133,209 | 326,724 | 273,007 | 304,133 | Pnpla2 | 3,359873053 | 1,54E-63 | | |
| Myom2 | 0 | 0,0633683 | 0,589808 | 0,388324 | 149,535 | 414,439 | 417,193 | 439,363 | 403,071 | 390,186 | Myom2 | 9,980393409 | 8,86E-58 | | |
| Insr | 11,2183 | 11,9591 | 12,9768 | 12,4108 | 50,9129 | 56,9942 | 52,3837 | 39,2974 | 39,8533 | 39,3948 | Insr | 2,444134694 | 6,69E-55 | | |
| Mb | 4,46004 | 10,3492 | 17,9056 | 25,751 | 2010,31 | 4203,43 | 4592,79 | 11445,5 | 8461,92 | 8444,28 | Mb | 7,8852074 | 1,23E-49 | | |
| Sgcg | 0 | 0 | 0,154004 | 0,371765 | 58,3294 | 96,794 | 88,5408 | 101,55 | 102,889 | 105,939 | Sgcg | 8,809537838 | 2,21E-48 | | |
| Perm1 | 0,0839271 | 0,993526 | 0,560439 | 0,507342 | 42,208 | 76,7784 | 74,6222 | 154,832 | 181,74 | 159,883 | Perm1 | 6,779548881 | 1,52E-47 | | |
| Fhl2 | 48,0563 | 51,4537 | 98,5059 | 68,2492 | 919,831 | 1174,95 | 1143,32 | 1831,59 | 2009,92 | 1966,37 | Fhl2 | 4,142158266 | 7,44E-47 | | |
| Slk | 13,3541 | 18,8901 | 18,4401 | 12,8627 | 79,4129 | 73,4604 | 71,4391 | 40,1455 | 36,9003 | 41,7197 | Slk | 2,534629582 | 7,44E-47 | | |
| Acsl1 | 18,3005 | 13,4787 | 17,4875 | 14,3498 | 234,897 | 452,003 | 462,164 | 509,336 | 535,45 | 514,217 | Acsl1 | 4,816840443 | 2,43E-45 | | |
| Dhrs11 | 8,34263 | 12,968 | 11,3935 | 10,1839 | 91,8293 | 139,257 | 129,331 | 96,4384 | 105,927 | 112,785 | Dhrs11 | 3,917429528 | 5,91E-45 | | |
| Lpl | 6,53664 | 16,6627 | 21,3307 | 15,8645 | 414,213 | 479,297 | 514,193 | 1985,74 | 1809,49 | 1732,55 | Lpl | 5,278614619 | 7,03E-45 | | |
| Kcnj12 | 0,722626 | 0,784158 | 0,904785 | 1,23772 | 16,4172 | 21,059 | 19,9102 | 23,9016 | 22,1366 | 25,2423 | Kcnj12 | 4,630740015 | 2,58E-43 | | |
| Calcoco1 | 19,1988 | 25,7899 | 32,5372 | 26,102 | 136,245 | 128,38 | 135,266 | 102,587 | 110,573 | 105,586 | Calcoco1 | 2,642150659 | 3,28E-42 | | |
| Plcd3 | 2,70306 | 3,09319 | 2,16598 | 2,06962 | 31,382 | 22,9078 | 25,0649 | 7,56922 | 7,53549 | 7,94742 | Plcd3 | 3,684468335 | 3,76E-42 | | |
| Myzap | 25,7748 | 15,9823 | 16,9648 | 11,721 | 296,985 | 360,907 | 369,53 | 428,198 | 422,379 | 402,928 | Myzap | 4,433552231 | 1,21E-41 | | |
| Kif1c | 24,1384 | 23,996 | 28,7079 | 27,5522 | 104,129 | 111,079 | 114,72 | 186,397 | 194,755 | 202,218 | Kif1c | 2,369455332 | 1,22E-41 | | |
| Neat1 | 1,13067 | 2,53499 | 4,80462 | 3,72796 | 83,2882 | 68,2086 | 60,7618 | 147,111 | 160,302 | 146,883 | Neat1 | 4,731945368 | 6,19E-41 | | |
| Lynx1 | 0 | 0,562485 | 0,407948 | 0,410333 | 29,3087 | 76,1813 | 75,1415 | 141,684 | 163,971 | 164,671 | Lynx1 | 7,391041068 | 2,33E-40 | | |
| Hopx | 54,63 | 63,7348 | 86,5034 | 75,7862 | 497,413 | 731,625 | 727,974 | 475,705 | 431,431 | 426,471 | Hopx | 3,508669927 | 4,43E-40 | | |
| Art3 | 5,42687 | 8,6993 | 11,3233 | 9,79269 | 151,163 | 90,3907 | 88,1976 | 283,707 | 260,566 | 336,521 | Art3 | 4,11778722 | 1,49E-39 | | |
| Fhod3 | 29,5953 | 27,2684 | 32,4094 | 34,8267 | 152,424 | 190,15 | 191,219 | 142,6 | 142,491 | 123,497 | Fhod3 | 2,843889895 | 1,89E-39 | | |
| Gja1 | 32,9749 | 44,793 | 40,8024 | 66,3351 | 454,449 | 569,281 | 554,854 | 314,233 | 394,899 | 287,715 | Gja1 | 3,766362764 | 2,27E-39 | | |
| Fam65b | 1,28798 | 0,960742 | 2,62005 | 2,92469 | 26,0968 | 30,7262 | 28,0609 | 4,1296 | 5,11616 | 2,80652 | Fam65b | 4,290316615 | 6,09E-39 | | |
| Ryr2 | 24,8534 | 26,6006 | 37,2525 | 33,0411 | 181,964 | 265,907 | 250,978 | 249,21 | 265,162 | 307,862 | Ryr2 | 3,215638256 | 7,94E-39 | | |
| Cpeb3 | 1,22086 | 0,943716 | 2,38307 | 2,04919 | 24,5691 | 21,0263 | 21,1888 | 27,2735 | 33,6608 | 33,1868 | Cpeb3 | 4,084931981 | 8,51E-39 | | |
| Nfix | 0 | 0,737529 | 1,64757 | 0,69528 | 55,678 | 41,96 | 44,8477 | 78,7993 | 90,53 | 92,1672 | Nfix | 5,90232308 | 2,52E-37 | | |
| Per3 | 0,631529 | 0,934529 | 1,27428 | 1,00753 | 9,12313 | 11,5829 | 10,5556 | 4,07236 | 3,8316 | 4,70913 | Per3 | 3,631374476 | 2,72E-37 | | |
| Fam160a1 | 0,729384 | 1,22321 | 1,33943 | 0,514415 | 14,1122 | 15,8943 | 15,8015 | 31,6576 | 24,6774 | 30,1197 | Fam160a1 | 4,252220221 | 1,01E-36 | | |
| Tmtc1 | 0 | 0 | 0 | 0 | 14,7528 | 14,1104 | 12,8579 | 30,566 | 24,8082 | 40,09 | Tmtc1 | 4,987559822 | 4,03E-36 | | |
| Lrrc10 | 28,4019 | 26,2818 | 42,164 | 33,4127 | 392,069 | 240,797 | 233,448 | 294,712 | 286,462 | 347,057 | Lrrc10 | 3,402385537 | 4,39E-36 | | |
| Dhrs7c | 0 | 0 | 0,280127 | 0,337977 | 78,8326 | 168,009 | 164,355 | 146,925 | 187,274 | 188,585 | Dhrs7c | 8,522113277 | 4,54E-36 | | |
| Adhfe1 | 0 | 0,104181 | 0,0881499 | 0,319182 | 27,9769 | 53,6273 | 54,3655 | 56,8056 | 70,0129 | 66,4783 | Adhfe1 | 8,040141268 | 7,62E-36 | | |
| Carns1 | 0,567751 | 0,160024 | 0,744714 | 0,4903 | 19,2795 | 42,5913 | 40,3931 | 18,2991 | 21,9616 | 21,6972 | Carns1 | 6,236319336 | 1,38E-35 | | |
| Syngr1 | 3,03241 | 6,48147 | 4,77852 | 3,64245 | 54,6351 | 93,3352 | 84,0939 | 98,1638 | 88,7945 | 96,6728 | Syngr1 | 4,452352936 | 1,52E-34 | | |
| Atp2a2 | 332,802 | 358,455 | 373,496 | 404,284 | 1816,26 | 2579,26 | 2705,8 | 5209,38 | 4460,62 | 4433,08 | Atp2a2 | 2,869736863 | 3,11E-34 | | |
| Adck3 | 0 | 0 | 0 | 0 | 69,1121 | 102,005 | 101,537 | 300,365 | 323,376 | 286,771 | Adck3 | 6,209155708 | 9,80E-34 | | |
| Hlf | 0,167517 | 0,605941 | 1,07205 | 0,450097 | 18,3756 | 14,3778 | 13,1876 | 9,04764 | 9,9207 | 11,3323 | Hlf | 4,916161379 | 1,41E-33 | | |
| Plin4 | 0 | 0 | 0 | 0 | 23,4872 | 27,2882 | 25,5513 | 76,2685 | 58,6455 | 88,568 | Plin4 | 9,587908977 | 1,54E-33 | | |
| Hspb6 | 1,09891 | 2,16805 | 6,19062 | 0,829979 | 151,4 | 294,078 | 293,512 | 1158,19 | 1060,39 | 1121,1 | Hspb6 | 6,425531026 | 9,65E-32 | | |
| Cdc37l1 | 5,95324 | 10,31 | 10,3798 | 5,73067 | 48,2976 | 51,2182 | 50,5706 | 63,9518 | 55,8942 | 61,6145 | Cdc37l1 | 2,976383814 | 4,04E-31 | | |
| Cish | 14,2444 | 11,4267 | 7,05515 | 10,0914 | 138,182 | 96,9476 | 95,3943 | 44,9204 | 93,3662 | 94,1962 | Cish | 3,598678564 | 1,49E-30 | | |
| Rorc | 0,133105 | 0,131306 | 0 | 0,402263 | 19,7207 | 24,08 | 24,7464 | 16,3275 | 20,0568 | 14,1935 | Rorc | 7,074448237 | 1,62E-30 | | |
| Pde1c | 1,68922 | 1,66061 | 1,18281 | 0,290596 | 20,8175 | 35,5197 | 33,1712 | 17,1638 | 21,1482 | 20,1198 | Pde1c | 5,232188382 | 4,04E-30 | | |
| Casq2 | 0,132467 | 2,48286 | 6,41291 | 7,3395 | 499,271 | 745,582 | 729,324 | 637,703 | 680,728 | 675,988 | Casq2 | 7,181173509 | 5,98E-29 | | |
| Osbpl8 | 8,9523 | 10,6033 | 9,72081 | 9,44685 | 34,6695 | 48,281 | 49,4547 | 24,2469 | 31,3814 | 23,483 | Osbpl8 | 2,50124879 | 1,14E-28 | | |
| Naca | 141,502 | 189,843 | 147,307 | 151,161 | 253,378 | 205,572 | 190,48 | 113,463 | 133,507 | 133,592 | Naca | 2,089421608 | 1,47E-28 | | |
| Adcy9 | 0,59858 | 0,826704 | 0,436077 | 0,52272 | 6,25926 | 8,86732 | 8,16287 | 9,66608 | 12,8493 | 11,5155 | Adcy9 | 3,84279242 | 2,60E-28 | | |
| Tmem35 | 0 | 0,70594 | 0,597288 | 0,901026 | 21,1194 | 34,6564 | 31,3142 | 4,54681 | 8,30981 | 8,1322 | Tmem35 | 5,70553085 | 3,74E-28 | | |
| Hhatl | 0,198521 | 0 | 1,49123 | 0,39991 | 25,8624 | 53,318 | 56,5958 | 130,993 | 134,634 | 117,242 | Hhatl | 5,872715948 | 4,10E-28 | | |
| Popdc2 | 139,256 | 127,098 | 157,557 | 170,799 | 542,661 | 593,565 | 597,456 | 419,162 | 492,517 | 442,863 | Popdc2 | 2,226222067 | 4,91E-28 | | |
| Cacna1c | 11,8447 | 9,94705 | 15,1891 | 14,3438 | 59,9082 | 88,1205 | 82,2104 | 45,5298 | 48,1447 | 45,3988 | Cacna1c | 2,852300559 | 4,99E-28 | | |
| Acadl | 17,4044 | 42,1423 | 67,3505 | 53,4787 | 669,897 | 1328,84 | 1365,56 | 1208,27 | 1094,46 | 1129,81 | Acadl | 4,816622677 | 7,27E-28 | | |
| Lmo7 | 15,8641 | 21,5294 | 32,3221 | 29,4176 | 153 | 200,647 | 203,594 | 136,503 | 139,611 | 140,004 | Lmo7 | 3,248520816 | 7,82E-28 | | |
| Prdx5 | 75,0423 | 86,1445 | 71,6063 | 76,8044 | 331,666 | 491,445 | 477,415 | 901,713 | 805,705 | 801,187 | Prdx5 | 2,748103071 | 1,15E-27 | | |
| Rilp | 0 | 1,65491 | 0 | 0 | 23,7585 | 34,9867 | 35,8103 | 44,5309 | 48,3447 | 45,9388 | Rilp | 5,527343303 | 1,72E-27 | | |
| Scd4 | 0,570568 | 0,855931 | 0,990024 | 0,636254 | 20,4637 | 45,2077 | 46,9062 | 9,99498 | 10,8994 | 12,1017 | Scd4 | 8,560849707 | 2,03E-27 | | |
| Cd36 | 3,27044 | 9,98343 | 12,5877 | 9,97631 | 129,272 | 317,136 | 324,652 | 603,073 | 668,161 | 588,641 | Cd36 | 4,999632262 | 2,81E-27 | | |
| Sptb | 7,7116 | 7,9614 | 10,2297 | 7,2585 | 28,5526 | 34,7896 | 33,6028 | 37,3252 | 42,2691 | 44,1042 | Sptb | 2,220325645 | 4,78E-27 | | |
| Nceh1 | 2,25784 | 2,15551 | 4,55967 | 5,06338 | 38,4174 | 37,8725 | 38,7439 | 81,847 | 80,9868 | 76,2859 | Nceh1 | 3,7039388 | 4,88E-27 | | |
| Slc25a34 | 0,123436 | 0 | 0 | 0,12435 | 18,7927 | 53,9767 | 53,5369 | 186,899 | 146,596 | 174,288 | Slc25a34 | 8,58301114 | 5,95E-27 | | |
| Mlf1 | 10,1597 | 17,3714 | 20,0661 | 18,4132 | 145,612 | 381,98 | 390,623 | 184,82 | 221,276 | 243,873 | Mlf1 | 4,45532367 | 1,96E-26 | | |
| C130080G10R | 4,68458 | 6,57636 | 9,32376 | 9,61874 | 51,1638 | 78,0588 | 76,2666 | 105,644 | 146,952 | 127,507 | C130080G10R | 3,521420678 | 2,12E-26 | | |
| Ptgds | 0 | 0 | 0 | 0 | 165,363 | 152,624 | 145,592 | 247,485 | 243,182 | 193,326 | Ptgds | 9,706041362 | 2,13E-26 | | |
| Acadvl | 19,7615 | 28,7846 | 53,606 | 33,4364 | 356,714 | 970,465 | 1039,56 | 1173,99 | 1113,13 | 1088,96 | Acadvl | 4,656315718 | 2,70E-26 | | |
| Acadm | 12,8649 | 24,5684 | 34,0029 | 22,0966 | 272,351 | 577,025 | 594,435 | 959,605 | 908,843 | 838,462 | Acadm | 4,223253576 | 2,83E-26 | | |
| Ablim1 | 30,1078 | 23,4982 | 27,3494 | 24,814 | 134,898 | 96,9149 | 97,2601 | 152,893 | 143,566 | 145,06 | Ablim1 | 2,324438071 | 8,69E-26 | | |
| Slc27a1 | 4,83897 | 7,68432 | 8,66913 | 5,94497 | 50,3743 | 135,455 | 135,225 | 142,842 | 151,694 | 137,864 | Slc27a1 | 4,270924561 | 4,99E-25 | | |
| Ckmt2 | 0,719547 | 0,236602 | 1,00089 | 5,31424 | 176,911 | 272,026 | 283,686 | 2392,1 | 2254,09 | 2381,96 | Ckmt2 | 6,867566413 | 5,76E-25 | | |
| Fabp3 | 11,1658 | 39,0808 | 83,24 | 74,4617 | 863,382 | 1930,64 | 2027,68 | 2665,61 | 1930,95 | 1994,73 | Fabp3 | 5,021597428 | 9,05E-25 | | |
| Plin5 | 0 | 0,174433 | 0,147586 | 0 | 20,4763 | 61,4374 | 60,0087 | 173,944 | 155,144 | 166,642 | Plin5 | 8,312707883 | 1,28E-24 | | |
| Fgf1 | 0,741633 | 0,650328 | 1,51323 | 0,581158 | 15,8759 | 46,9177 | 43,9989 | 100,902 | 106,905 | 107,959 | Fgf1 | 5,419894993 | 1,93E-24 | | |
| Zfp651 | 2,08225 | 3,91266 | 5,21437 | 4,7456 | 20,9625 | 23,0134 | 24,1435 | 31,4146 | 35,2022 | 37,9459 | Zfp651 | 2,776796693 | 2,24E-24 | | |
| Sdha | 95,4131 | 112,131 | 179,563 | 128,781 | 601,944 | 921,835 | 963,973 | 1196,35 | 1148,82 | 1149,45 | Sdha | 2,978543987 | 2,90E-24 | | |
| Popdc3 | 5,02859 | 3,12333 | 3,73073 | 5,44025 | 61,0762 | 35,8363 | 33,1185 | 23,8317 | 19,3792 | 21,5334 | Popdc3 | 3,620561969 | 4,05E-24 | | |
| Prob1 | 2,69279 | 1,61978 | 1,9188 | 1,522 | 14,0825 | 14,8025 | 15,6753 | 32,217 | 37,2885 | 33,5899 | Prob1 | 3,121343915 | 4,15E-24 | | |
| Mef2a | 17,7246 | 33,6709 | 31,2395 | 24,3401 | 102,164 | 125,225 | 125,909 | 55,0274 | 77,3457 | 53,3717 | Mef2a | 2,437572891 | 6,05E-24 | | |
| Stbd1 | 9,46668 | 15,621 | 23,9914 | 23,5791 | 138,455 | 125,669 | 131,219 | 4,0626 | 5,55916 | 5,41942 | Stbd1 | 3,139664269 | 7,31E-24 | | |
| Fuca2 | 5,29173 | 8,11151 | 4,75685 | 9,26857 | 51,0206 | 105,291 | 103,324 | 95,393 | 81,2059 | 98,7153 | Fuca2 | 3,909312851 | 8,24E-24 | | |
| Gramd4 | 5,63354 | 9,54326 | 10,6788 | 6,77938 | 30,6358 | 34,8385 | 34,1012 | 19,2504 | 26,0017 | 21,4239 | Gramd4 | 2,362840835 | 1,14E-23 | | |
| Sorbs1 | 25,7356 | 23,8204 | 21,7844 | 17,3552 | 81,8142 | 92,6643 | 87,8001 | 237,042 | 160,971 | 225,982 | Sorbs1 | 2,382961418 | 1,29E-23 | | |
| Kifc3 | 6,48845 | 9,33786 | 7,17487 | 7,62355 | 25,0924 | 25,8484 | 26,5669 | 51,0726 | 50,8073 | 53,8943 | Kifc3 | 2,00115109 | 1,32E-23 | | |
| Asb11 | 0 | 0 | 0 | 0 | 45,9349 | 73,1377 | 72,9788 | 147,944 | 131,247 | 126,067 | Asb11 | 9,365255646 | 1,60E-23 | | |
| Scarb2 | 10,6095 | 22,518 | 20,1258 | 20,7266 | 155,924 | 95,8322 | 95,8767 | 94,5887 | 85,4692 | 81,9382 | Scarb2 | 3,001002505 | 1,63E-23 | | |
| Tmem245 | 7,15182 | 6,37375 | 7,57011 | 8,26187 | 23,5258 | 27,7493 | 27,4415 | 16,4605 | 18,0357 | 16,3118 | Tmem245 | 2,162688598 | 2,05E-23 | | |
| Pdha1 | 44,6113 | 67,3894 | 98,7815 | 79,8237 | 340,845 | 558,944 | 571,393 | 740 | 816,093 | 811,28 | Pdha1 | 3,061054206 | 2,12E-23 | | |
| Alas1 | 9,91781 | 16,0648 | 21,3599 | 15,6037 | 87,4345 | 172,07 | 175,332 | 178,332 | 224,947 | 252,811 | Alas1 | 3,487562262 | 2,89E-23 | | |
| Dhrs7 | 8,96126 | 6,6299 | 7,47885 | 10,1522 | 59,8181 | 117,461 | 110,942 | 41,8414 | 47,0549 | 46,4599 | Dhrs7 | 3,706378479 | 5,09E-23 | | |
| Rcan2 | 4,35191 | 8,01384 | 15,7814 | 7,50225 | 91,9256 | 118,304 | 128,821 | 253,366 | 239,446 | 253,85 | Rcan2 | 3,861084076 | 5,19E-23 | | |
| Smyd2 | 62,7121 | 60,854 | 79,243 | 60,2907 | 191,13 | 251,877 | 256,546 | 120,099 | 112,693 | 122,5 | Smyd2 | 2,143159724 | 5,61E-23 | | |
| C030006K11R | 4,34197 | 3,8936 | 4,23592 | 5,44336 | 26,634 | 33,5392 | 36,0734 | 74,1317 | 69,1671 | 75,7319 | C030006K11R | 2,944636437 | 5,66E-23 | | |
| Npas2 | 0 | 0 | 0,256316 | 0,232035 | 5,95173 | 8,08966 | 7,99857 | 0,907997 | 1,20654 | 0,868983 | Npas2 | 5,553486188 | 5,68E-23 | | |
| Yipf7 | 0,303815 | 0,299698 | 0,253552 | 0 | 25,8306 | 82,0913 | 80,9024 | 135,969 | 161,064 | 147,288 | Yipf7 | 7,63012096 | 6,39E-23 | | |
| Ddo | 0 | 0 | 0,365019 | 0,330422 | 10,5051 | 32,4089 | 31,0384 | 29,7685 | 36,5249 | 45,5629 | Ddo | 6,879905096 | 7,26E-23 | | |
| Rnf207 | 0 | 3,71571 | 9,50856 | 4,87837 | 51,3473 | 119,222 | 119,625 | 29,4562 | 34,1773 | 38,9889 | Rnf207 | 4,634071384 | 7,36E-23 | | |
| Xirp2 | 0,0776977 | 0,0255499 | 0,0216194 | 0,0260974 | 2,10895 | 3,31166 | 3,3756 | 58,2615 | 66,0797 | 79,5533 | Xirp2 | 6,190931313 | 1,21E-22 | | |
| Pptc7 | 3,53104 | 4,23469 | 6,24162 | 6,62703 | 25,4172 | 33,3103 | 33,2211 | 76,8656 | 79,607 | 73,5972 | Pptc7 | 2,843071604 | 1,93E-22 | | |
| Bckdha | 26,404 | 26,4327 | 36,5663 | 33,2929 | 98,9859 | 104,017 | 106,161 | 388,75 | 372,207 | 355,654 | Bckdha | 2,032252405 | 2,14E-22 | | |
| Rit1 | 8,73027 | 7,46485 | 9,75116 | 9,62903 | 32,8632 | 34,7759 | 34,4359 | 37,2742 | 33,6814 | 27,925 | Rit1 | 2,216250158 | 2,49E-22 | | |
| Cox8b | 0 | 4,00051 | 30,4499 | 10,1843 | 618,393 | 1223,39 | 1124,88 | 2644,04 | 2041,36 | 2089,9 | Cox8b | 6,274424507 | 3,27E-22 | | |
| Ndufa8 | 128,984 | 107,079 | 158,265 | 121,571 | 481,261 | 659,598 | 639,813 | 924,775 | 827,864 | 828,038 | Ndufa8 | 2,467277171 | 3,84E-22 | | |
| Phyh | 12,8334 | 26,9914 | 34,4565 | 26,0933 | 174,159 | 377,65 | 384,162 | 700,331 | 772,724 | 732,216 | Phyh | 3,867003499 | 4,42E-22 | | |
| Ogn | 0 | 0,129214 | 0,765307 | 0,131952 | 27,4838 | 12,4225 | 12,939 | 35,4689 | 65,7088 | 46,2709 | Ogn | 5,870035832 | 5,99E-22 | | |
| Ankrd9 | 0,519906 | 0,6411 | 0,86791 | 1,83312 | 11,1913 | 12,968 | 11,6829 | 35,315 | 33,0575 | 46,3977 | Ankrd9 | 3,983425454 | 7,44E-22 | | |
| Slc25a42 | 0,61777 | 1,23336 | 2,57826 | 2,8007 | 21,9687 | 58,2176 | 55,805 | 58,9188 | 61,5418 | 63,4592 | Slc25a42 | 4,841653872 | 8,47E-22 | | |
| Echs1 | 39,6586 | 77,0651 | 70,3857 | 57,746 | 225,571 | 285,232 | 287,561 | 509,531 | 528,602 | 524,677 | Echs1 | 2,411126075 | 1,05E-21 | | |
| Ehbp1 | 4,73343 | 8,51875 | 10,363 | 9,02279 | 39,2827 | 36,0398 | 34,9118 | 21,8246 | 33,7374 | 28,2066 | Ehbp1 | 2,57133464 | 1,29E-21 | | |
| Murc | 1,49904 | 1,80739 | 0,973143 | 0,838883 | 16,3504 | 41,4988 | 42,339 | 94,0621 | 93,9407 | 85,1887 | Murc | 4,810202899 | 1,41E-21 | | |

E:\Daten\Publications\2020\GWAS CHD\Manuscript\BIORXIV preprint server\2020-06-15_Lahm_Supplementary_Datafile2.xlsx

| Gene | d9-11 (1) | d9-11 (2) | d9-11(3) | d9-11(4) | newborn(1) | newborn(2) | newborn(3) | adult(1) | adult(2) | adult(3) | Gene | log2(fold) | FDR(*p* -value) |
|---|---|---|---|---|---|---|---|---|---|---|---|---|---|
| Gpr157 | 1,19926 | 0,902435 | 2,5869 | 2,09815 | 17,6925 | 18,4977 | 19,2553 | 26,9952 | 37,4936 | 31,8829 | Gpr157 | 3,907625765 | 1,77E-21 |
| Mul1 | 6,40501 | 7,0016 | 9,17544 | 9,15634 | 26,5928 | 27,7394 | 28,0682 | 27,5563 | 29,1356 | 29,0399 | Mul1 | 2,090864294 | 1,79E-21 |
| Ech1 | 29,7504 | 49,89 | 55,7249 | 48,2616 | 291,041 | 938,509 | 931,098 | 1921,8 | 1802,74 | 1509,22 | Ech1 | 4,135679811 | 1,92E-21 |
| Myh6 | 555,998 | 589,973 | 1160,99 | 957,841 | 4240,92 | 4328,16 | 4721,81 | 6633,53 | 5590,35 | 5863,09 | Myh6 | 2,791435582 | 1,94E-21 |
| Dip2c | 4,98748 | 4,80487 | 8,13137 | 4,98621 | 22,9986 | 29,2275 | 28,8566 | 14,8673 | 20,7719 | 15,7298 | Dip2c | 2,511460624 | 1,96E-21 |
| Gas6 | 30,3505 | 22,1079 | 17,3162 | 12,5138 | 183,782 | 158,936 | 163,03 | 60,6314 | 64,0158 | 59,0524 | Gas6 | 3,283246488 | 2,01E-21 |
| Pacsin3 | 7,9646 | 14,4484 | 13,0029 | 13,2141 | 48,1014 | 60,6485 | 61,8816 | 137,159 | 157,166 | 143,052 | Pacsin3 | 2,533714661 | 2,03E-21 |
| Hsdl2 | 6,97581 | 10,8854 | 18,3147 | 9,71063 | 87,0553 | 279,637 | 288,914 | 216,808 | 227,633 | 203,986 | Hsdl2 | 4,440346458 | 2,71E-21 |
| Mylk3 | 31,0334 | 39,2409 | 70,411 | 55,1159 | 255,451 | 377,157 | 375,449 | 244,147 | 289,189 | 261,772 | Mylk3 | 3,073976973 | 2,74E-21 |
| Vps13d | 2,85689 | 4,99202 | 6,2877 | 4,17359 | 19,7895 | 19,5775 | 17,9735 | 16,118 | 18,2562 | 18,1275 | Vps13d | 2,369050685 | 3,19E-21 |
| D10Jhu81e | 28,1548 | 37,992 | 53,4238 | 43,6001 | 170,282 | 274,643 | 263,417 | 573,27 | 558,893 | 531,53 | D10Jhu81e | 2,784983393 | 3,27E-21 |
| Mpc2 | 21,9942 | 27,8265 | 43,8872 | 31,2816 | 176,841 | 327,881 | 309,749 | 545,618 | 488,751 | 529,219 | Mpc2 | 3,269788666 | 3,30E-21 |
| Tnnt2 | 1328,63 | 1314,08 | 2171,26 | 1609,97 | 5693,43 | 6647,86 | 7550,29 | 8717,72 | 7496,87 | 7008,06 | Tnnt2 | 2,387649373 | 3,45E-21 |
| Ppp1r14c | 33,0391 | 41,5703 | 56,1999 | 42,1631 | 177,388 | 337,027 | 342,41 | 140,001 | 192,964 | 173,151 | Ppp1r14c | 3,000660526 | 4,10E-21 |
| Bves | 39,4123 | 32,3991 | 41,3006 | 36,6076 | 108,712 | 138,227 | 146,144 | 54,2055 | 57,9759 | 58,1353 | Bves | 2,132289035 | 6,46E-21 |
| Pik3ip1 | 2,85368 | 4,88939 | 8,27385 | 6,05185 | 66,6414 | 41,1735 | 40,5168 | 63,4341 | 47,4576 | 43,1099 | Pik3ip1 | 3,364166897 | 1,04E-20 |
| Fam174b | 8,28365 | 6,58616 | 10,2165 | 7,72225 | 41,9932 | 86,2648 | 89,6896 | 201,641 | 242,834 | 189,775 | Fam174b | 3,399483637 | 1,21E-20 |
| Pfkm | 34,6193 | 44,5211 | 68,3786 | 43,6245 | 208,541 | 388,167 | 409,676 | 737,631 | 734,748 | 711,054 | Pfkm | 3,09442266 | 1,87E-20 |
| Tbc1d16 | 11,142 | 12,6315 | 11,3633 | 10,3666 | 31,326 | 43,5987 | 42,1028 | 34,8936 | 36,4491 | 35,8078 | Tbc1d16 | 2,080190576 | 2,54E-20 |
| Ppp1r3a | 0,132204 | 0,304314 | 1,06677 | 0,13321 | 13,2264 | 15,352 | 15,5605 | 43,2953 | 45,3971 | 56,7041 | Ppp1r3a | 5,241224174 | 3,36E-20 |
| Kcnq1 | 1,08199 | 2,02802 | 4,86434 | 5,44003 | 42,8676 | 72,5615 | 74,7269 | 37,709 | 47,1164 | 41,6574 | Kcnq1 | 4,077487919 | 3,58E-20 |
| 3425401B19R | 1,42689 | 1,40763 | 3,62084 | 3,59424 | 23,659 | 22,6631 | 21,0242 | 40,87 | 46,1785 | 62,0937 | 3425401B19R | 3,428673565 | 4,22E-20 |
| Mfn1 | 15,585 | 19,34 | 26,8423 | 18,2603 | 69,9068 | 112,874 | 113,712 | 145,412 | 164,278 | 151,361 | Mfn1 | 2,595278178 | 5,62E-20 |
| Pde4dip | 8,03036 | 17,73 | 24,32 | 24,3827 | 143,287 | 160,888 | 154,66 | 448,435 | 455,863 | 513,902 | Pde4dip | 3,358743601 | 6,48E-20 |
| Acaa2 | 29,4165 | 54,5264 | 91,869 | 82,9054 | 491,29 | 1162,98 | 1197,45 | 1246,98 | 1199,78 | 1098,74 | Acaa2 | 4,058332379 | 1,02E-19 |
| Crip2 | 342,486 | 351,117 | 464,616 | 487,648 | 1228,13 | 1434,38 | 1454,22 | 1457,01 | 1451,08 | 1379,82 | Crip2 | 2,03721137 | 1,06E-19 |
| Lrrc2 | 1,96897 | 1,04758 | 1,03412 | 0,98072 | 9,02691 | 11,5978 | 11,4829 | 137,313 | 120,409 | 111,173 | Lrrc2 | 3,366032203 | 1,38E-19 |
| Lrrfip2 | 14,5626 | 22,5256 | 20,5843 | 16,8529 | 39,2697 | 51,6402 | 51,5422 | 26,8076 | 29,1803 | 26,3565 | Lrrfip2 | 2,217249693 | 1,59E-19 |
| Fabp4 | 0 | 0 | 1,79878 | 0 | 288,76 | 84,7379 | 72,9428 | 128,847 | 269,726 | 218,877 | Fabp4 | 7,454689552 | 1,77E-19 |
| Acot2 | 5,2592 | 5,45509 | 14,0433 | 13,3109 | 92,1452 | 217,482 | 224,049 | 142,765 | 102,408 | 110,696 | Acot2 | 4,443640946 | 2,14E-19 |
| Hadh | 63,1517 | 90,9937 | 94,7462 | 86,8276 | 288,888 | 556,995 | 548,473 | 800,368 | 801,194 | 757,954 | Hadh | 2,760962688 | 2,42E-19 |
| Gnptab | 4,57514 | 5,95767 | 8,75822 | 6,14632 | 22,4979 | 25,0344 | 24,9391 | 12,6591 | 12,7571 | 12,7374 | Gnptab | 2,245485854 | 3,35E-19 |
| Idh3b | 69,8424 | 72,5838 | 113,333 | 107,275 | 383,447 | 619,443 | 617,294 | 942,176 | 856,749 | 894,346 | Idh3b | 2,826068987 | 3,35E-19 |
| Uqcrc2 | 87,0008 | 124,229 | 149,043 | 158,143 | 472,218 | 761,463 | 709,82 | 902,491 | 902,671 | 921,833 | Uqcrc2 | 2,609779576 | 3,39E-19 |
| Cmya5 | 0,876128 | 1,03201 | 3,54759 | 1,54166 | 25,5663 | 44,7787 | 42,7263 | 128,848 | 148,409 | 137,269 | Cmya5 | 4,570664671 | 4,13E-19 |
| Got1 | 41,9522 | 60,0381 | 88,8964 | 81,1299 | 284,667 | 405,249 | 409,877 | 834,892 | 815,678 | 827,633 | Got1 | 2,711554153 | 4,47E-19 |
| Micall1 | 16,7672 | 12,1856 | 17,9559 | 10,3166 | 72,0415 | 74,7218 | 69,9508 | 23,0273 | 31,1642 | 35,5801 | Micall1 | 2,423124452 | 4,84E-19 |
| Mylk4 | 0 | 0 | 0,140451 | 0,226045 | 6,18673 | 3,75256 | 3,99158 | 85,3624 | 69,9453 | 86,7861 | Mylk4 | 5,635141439 | 5,40E-19 |
| Nt5e | 0,361869 | 1,07095 | 0,302054 | 0,18229 | 10,0489 | 7,73096 | 7,87562 | 8,39297 | 12,4143 | 9,77756 | Nt5e | 4,342249599 | 5,93E-19 |
| Ndufb10 | 104,974 | 163,945 | 190,974 | 185,536 | 621,307 | 1099,81 | 1129,9 | 1474,68 | 1404,12 | 1298,88 | Ndufb10 | 2,854100197 | 6,84E-19 |
| Macrod1 | 4,60354 | 11,9206 | 18,7297 | 14,197 | 107,325 | 193,896 | 194,387 | 370,944 | 362,609 | 340,837 | Macrod1 | 3,661106229 | 7,26E-19 |
| Fem1a | 5,23217 | 6,57133 | 11,1977 | 11,1013 | 45,3094 | 47,5 | 45,7037 | 88,3934 | 85,12 | 97,1164 | Fem1a | 2,74610609 | 8,43E-19 |
| Ndufa9 | 58,125 | 83,6967 | 115,642 | 88,7715 | 310,361 | 477,499 | 475,098 | 631,176 | 637,203 | 634,471 | Ndufa9 | 2,55059721 | 1,17E-18 |
| Adrbk2 | 4,84311 | 3,50053 | 4,47968 | 3,28548 | 16,2396 | 14,4384 | 14,0576 | 2,27019 | 3,40124 | 3,33067 | Adrbk2 | 2,171808742 | 1,17E-18 |
| Hadha | 32,3745 | 54,9985 | 79,245 | 69,132 | 315,758 | 859,402 | 882,027 | 833,798 | 839,302 | 769,594 | Hadha | 3,796334312 | 1,56E-18 |
| Agpat3 | 10,387 | 14,2523 | 17,419 | 13,6041 | 51,5106 | 116,818 | 113,883 | 87,4237 | 105,336 | 90,0302 | Agpat3 | 3,119273367 | 1,86E-18 |
| Tmem71 | 0 | 1,21652 | 1,68435 | 1,24235 | 19,1463 | 22,2649 | 21,9684 | 2,75171 | 3,38158 | 3,55805 | Tmem71 | 4,505458072 | 2,29E-18 |
| Nnt | 25,2846 | 39,8183 | 70,9134 | 65,0278 | 299,273 | 482,249 | 478,17 | 757,176 | 767,451 | 689,729 | Nnt | 3,32851037 | 3,00E-18 |
| Kcng2 | 14,0538 | 9,58919 | 13,2946 | 18,7595 | 74,9605 | 99,0871 | 104,2 | 99,2159 | 84,7312 | 88,4149 | Kcng2 | 2,97011851 | 3,69E-18 |
| Asb10 | 0,14483 | 0,142872 | 1,81328 | 0,87537 | 14,9515 | 21,969 | 23,0216 | 84,8576 | 84,0765 | 78,8031 | Asb10 | 4,650492962 | 3,69E-18 |
| Ears2 | 5,04403 | 9,03017 | 9,51138 | 9,22036 | 25,476 | 26,6668 | 23,0938 | 13,9019 | 15,7277 | 13,6517 | Ears2 | 2,131121831 | 3,83E-18 |
| Herc3 | 4,98945 | 8,58041 | 8,61102 | 6,04611 | 26,8142 | 53,8399 | 53,753 | 19,9417 | 27,7164 | 25,3084 | Herc3 | 2,965358794 | 3,93E-18 |
| S100a1 | 16,0777 | 27,1865 | 51,0994 | 49,2889 | 286,554 | 459,195 | 435,715 | 1131,57 | 1047,28 | 1043,21 | S100a1 | 3,647571799 | 4,75E-18 |
| Cidea | 0 | 0 | 0,536575 | 0 | 19,4764 | 50,9364 | 57,0245 | 37,8074 | 33,6327 | 36,5225 | Cidea | 7,306316551 | 5,87E-18 |
| A430105I19Ri | 0,20286 | 0,266828 | 0,338663 | 0,408781 | 3,77328 | 6,90929 | 6,36047 | 11,2611 | 13,4916 | 19,4353 | A430105I19Ri | 4,19842388 | 6,25E-18 |
| Perp | 4,70129 | 15,8752 | 22,9398 | 20,9453 | 158,684 | 203,088 | 203,507 | 98,2236 | 126,575 | 108,971 | Perp | 3,758154916 | 7,06E-18 |
| Agtr1a | 1,91377 | 2,75924 | 4,05484 | 4,00396 | 17,3276 | 25,8919 | 22,9498 | 28,1703 | 37,0622 | 30,4285 | Agtr1a | 3,068221636 | 8,02E-18 |
| Ndufb8 | 155,233 | 227,552 | 208,658 | 239,307 | 650,172 | 851,72 | 784,748 | 1196,02 | 1099,07 | 1108,52 | Ndufb8 | 2,0297262 | 8,91E-18 |
| Flot1 | 31,32 | 42,4823 | 40,9306 | 43,1676 | 103,073 | 146,11 | 146,115 | 185,04 | 173,511 | 180,186 | Flot1 | 2,048775756 | 1,11E-17 |
| Map1lc3a | 33,8308 | 39,1757 | 73,3428 | 66,641 | 257,285 | 353,515 | 348,587 | 646,286 | 633,605 | 602,875 | Map1lc3a | 2,811871705 | 1,27E-17 |
| Tcap | 29,5771 | 23,0117 | 69,1779 | 52,8396 | 328,131 | 400,774 | 416,259 | 3286,88 | 2691,99 | 3509,29 | Tcap | 3,354194373 | 1,41E-17 |
| Pink1 | 13,7346 | 13,8283 | 32,1461 | 23,9629 | 116,03 | 170,968 | 180,307 | 489,831 | 477,609 | 421,293 | Pink1 | 3,136342107 | 1,46E-17 |
| Mlxipl | 0,265944 | 0,437254 | 0,0739951 | 0,446561 | 4,21993 | 7,88609 | 7,71696 | 10,7242 | 11,5699 | 16,3782 | Mlxipl | 4,717500474 | 1,57E-17 |
| Eno3 | 120,015 | 155,205 | 178,491 | 142,729 | 489,52 | 1023,93 | 1045,46 | 1289,58 | 1273,47 | 1243,69 | Eno3 | 2,81715925 | 1,61E-17 |
| Idh2 | 159,347 | 227,594 | 430,139 | 366,64 | 1583,41 | 2105,03 | 2189,22 | 2383,06 | 1955,54 | 1771,26 | Idh2 | 3,000710483 | 1,74E-17 |
| Isca1 | 34,4552 | 50,7407 | 52,0559 | 41,6709 | 130,883 | 202,109 | 207,683 | 252,955 | 264,479 | 255,398 | Isca1 | 2,293356208 | 1,80E-17 |
| Ddc | 5,77478 | 11,7386 | 18,7684 | 20,7995 | 93,5744 | 223,951 | 222,647 | 5,55976 | 7,30172 | 6,76024 | Ddc | 3,875066947 | 2,35E-17 |
| Txnip | 44,7711 | 82,8687 | 68,1464 | 60,3152 | 223,228 | 439,209 | 412,446 | 635,212 | 481,357 | 481,642 | Txnip | 2,750354076 | 3,36E-17 |
| Cyc1 | 140,53 | 280,017 | 303,472 | 268,578 | 862,833 | 1177,62 | 1190,62 | 1348,58 | 1279,93 | 1329,22 | Cyc1 | 2,434816386 | 3,52E-17 |
| Ucp3 | 0 | 0,134544 | 0 | 0,274787 | 4,56375 | 38,4689 | 35,7361 | 95,0447 | 25,6375 | 63,5308 | Ucp3 | 7,403283772 | 4,70E-17 |
| Sh3kbp1 | 18,3722 | 18,3667 | 22,9228 | 19,9975 | 72,7193 | 128,09 | 132,449 | 40,6678 | 60,166 | 44,4229 | Sh3kbp1 | 2,396012893 | 4,75E-17 |
| Ndufa1 | 88,1363 | 90,2325 | 109,418 | 145,17 | 391,367 | 600,354 | 594,184 | 733,844 | 648,088 | 678,96 | Ndufa1 | 2,526167483 | 5,01E-17 |
| Ndufb9 | 94,0827 | 146,127 | 165,775 | 204,132 | 560,186 | 849,837 | 778,955 | 1089,98 | 1135,16 | 1075,63 | Ndufb9 | 2,516228481 | 5,27E-17 |
| Ndufv2 | 45,7806 | 77,1593 | 77,9539 | 82,9473 | 222,788 | 343,563 | 338,872 | 414,936 | 452,763 | 424,72 | Ndufv2 | 2,397997627 | 5,81E-17 |
| Wdr13 | 6,261 | 10,6537 | 15,9235 | 10,218 | 44,6918 | 62,3557 | 60,0087 | 21,3669 | 21,835 | 25,2099 | Wdr13 | 2,649311527 | 6,52E-17 |
| 9030617O03R | 0 | 0 | 0,16863 | 0,101767 | 5,18167 | 12,3318 | 13,2242 | 39,8642 | 48,9983 | 48,0167 | 9030617O03R | 6,985453129 | 6,93E-17 |
| Ndufs7 | 139,652 | 131,086 | 212,372 | 200,035 | 561,977 | 766,278 | 761,259 | 1450,58 | 1073,52 | 1203,36 | Ndufs7 | 2,28917029 | 7,46E-17 |
| Etfdh | 14,5953 | 40,2853 | 48,4828 | 43,5136 | 214,004 | 579,811 | 572,841 | 660,014 | 651,74 | 618,607 | Etfdh | 3,859693041 | 7,51E-17 |
| Mrps23 | 32,7826 | 54,3676 | 37,5961 | 38,8581 | 115,404 | 169,9 | 161,307 | 230,621 | 230,332 | 221,807 | Mrps23 | 2,059022957 | 7,54E-17 |
| Nfatc2 | 1,07474 | 2,84859 | 2,7141 | 3,18115 | 10,7308 | 12,5015 | 11,9977 | 5,97156 | 7,24753 | 6,98644 | Nfatc2 | 2,830384648 | 9,13E-17 |
| Bcl2l2 | 5,90856 | 4,81105 | 8,53298 | 6,33072 | 23,0688 | 24,8924 | 24,5529 | 22,9448 | 24,6775 | 27,0437 | Bcl2l2 | 2,199896337 | 1,37E-16 |
| Ndufs2 | 115,255 | 150,224 | 214,881 | 199,721 | 599,866 | 936,191 | 927,531 | 1288,68 | 1125,1 | 1188,86 | Ndufs2 | 2,456821982 | 1,37E-16 |
| Slc25a4 | 560,118 | 908,781 | 1269,54 | 1045,58 | 3570,94 | 4166,85 | 4446,82 | 6619,49 | 5382,96 | 5408,62 | Slc25a4 | 2,336480519 | 1,38E-16 |
| Idh3g | 81,4276 | 92,9933 | 166,246 | 122,313 | 433,518 | 639,278 | 656,434 | 719,403 | 754,791 | 759,596 | Idh3g | 2,566233621 | 1,52E-16 |
| Mfn2 | 21,6589 | 34,5992 | 63,7391 | 48,2743 | 220,273 | 321,917 | 326,434 | 386,066 | 408,308 | 411,94 | Mfn2 | 3,070551988 | 1,70E-16 |
| Gbe1 | 11,1997 | 9,53194 | 17,1385 | 12,2786 | 44,8172 | 48,0066 | 47,1161 | 35,0381 | 37,5795 | 41,684 | Gbe1 | 2,118052336 | 1,70E-16 |
| Cox4i1 | 435,29 | 621,708 | 664,172 | 834,758 | 2028,21 | 2547,83 | 2620,87 | 3710,13 | 3052,53 | 2971,96 | Cox4i1 | 2,18416847 | 1,70E-16 |
| Akap1 | 11,8138 | 21,923 | 20,3571 | 28,5225 | 82,1535 | 102,95 | 100,812 | 87,3774 | 104,341 | 91,5692 | Akap1 | 2,496404177 | 1,77E-16 |
| Ppip5k2 | 10,4038 | 16,4888 | 14,3553 | 10,8263 | 40,3913 | 87,1148 | 83,1453 | 47,3769 | 54,8288 | 61,4392 | Ppip5k2 | 2,790168833 | 2,91E-16 |
| Mdh1 | 71,8396 | 114,32 | 198,702 | 180,994 | 727,612 | 1169,45 | 1195,69 | 1544,61 | 1484,11 | 1398,44 | Mdh1 | 3,14074966 | 3,37E-16 |
| Tnnc1 | 666,425 | 884,157 | 1314,23 | 1039,56 | 3010,02 | 3435,95 | 3544,41 | 4202,68 | 3206,25 | 3594,3 | Tnnc1 | 2,055993438 | 3,66E-16 |
| Fcgrt | 5,40239 | 5,72407 | 6,34601 | 6,44905 | 28,0113 | 20,3226 | 20,7763 | 52,8236 | 52,4213 | 43,9737 | Fcgrt | 2,226232377 | 3,70E-16 |
| Strip2 | 0,36016 | 0,874107 | 4,83325 | 2,66116 | 40,8609 | 111,245 | 107,04 | 48,2203 | 50,7979 | 52,1511 | Strip2 | 5,316201246 | 4,24E-16 |
| Nr0b2 | 0 | 0,333323 | 1,12797 | 0,680456 | 15,7664 | 15,8649 | 17,6961 | 12,2229 | 18,154 | 14,5705 | Nr0b2 | 5,018555006 | 6,09E-16 |
| Nudt8 | 10,1719 | 21,6512 | 21,8886 | 17,241 | 60,5377 | 92,4515 | 93,3876 | 151,323 | 151,936 | 141,038 | Nudt8 | 2,563065979 | 6,57E-16 |
| Arhgef6 | 0,699043 | 1,9309 | 4,2596 | 2,25381 | 25,5935 | 22,1912 | 22,1259 | 15,4895 | 16,7183 | 18,1498 | Arhgef6 | 3,564473373 | 9,28E-16 |
| Mettl7a1 | 2,00931 | 2,88311 | 3,65904 | 1,37993 | 14,7168 | 16,654 | 14,9777 | 33,4885 | 34,4585 | 35,1392 | Mettl7a1 | 2,936541895 | 9,76E-16 |
| Crat | 4,22873 | 8,62417 | 21,7095 | 15,414 | 103,799 | 244,324 | 255,118 | 245,953 | 269,509 | 247,245 | Crat | 4,153329885 | 1,02E-15 |
| Klhl40 | 0 | 0 | 0,453932 | 0,410891 | 6,51573 | 11,4123 | 11,388 | 12,0443 | 13,4857 | 11,1501 | Klhl40 | 5,123754423 | 1,07E-15 |
| Aldh6a1 | 3,59883 | 7,96402 | 11,7722 | 12,2491 | 59,0016 | 75,6145 | 74,0407 | 153,555 | 180,004 | 170,287 | Aldh6a1 | 3,173621788 | 1,07E-15 |
| Cacna1g | 1,57555 | 1,44991 | 4,49086 | 1,9486 | 15,4175 | 28,9832 | 25,6915 | 5,53706 | 8,07383 | 7,39901 | Cacna1g | 3,534983551 | 1,35E-15 |
| Dst | 5,97045 | 9,95987 | 13,315 | 11,2413 | 34,4228 | 33,7269 | 31,3688 | 46,675 | 51,4961 | 57,1785 | Dst | 2,09946318 | 1,44E-15 |
| Nrap | 0,239163 | 4,01089 | 8,87955 | 3,85462 | 168,75 | 167,299 | 173,595 | 217,486 | 196,345 | 237,146 | Nrap | 5,342984843 | 1,50E-15 |
| Trp53inp2 | 20,9261 | 42,0677 | 74,2646 | 45,7676 | 245,065 | 425,943 | 419,849 | 299,158 | 351,14 | 396,801 | Trp53inp2 | 3,256224999 | 1,53E-15 |
| Mdh2 | 217,23 | 336,374 | 443,455 | 413,325 | 1147,42 | 1625,14 | 1693,99 | 2026,34 | 1869,96 | 1940,89 | Mdh2 | 2,342396967 | 1,59E-15 |
| Dnajc28 | 0,0916945 | 0,381369 | 0,968064 | 0,68159 | 6,14142 | 14,7659 | 14,7543 | 29,9714 | 34,4309 | 31,4065 | Dnajc28 | 4,497217349 | 2,03E-15 |
| Coq9 | 20,0007 | 41,0873 | 52,3165 | 43,1969 | 154,304 | 300,404 | 282,199 | 637,727 | 565,849 | 625,178 | Coq9 | 2,940992644 | 2,18E-15 |
| Cluh | 22,1908 | 39,3233 | 55,4965 | 44,1852 | 139,662 | 185,055 | 185,047 | 145,478 | 161,714 | 147,959 | Cluh | 2,382396328 | 2,67E-15 |
| Nmrk2 | 0,187507 | 0,647398 | 0,782504 | 0,755465 | 4,36375 | 13,8778 | 13,6006 | 4,08384 | 4,04135 | 3,47131 | Nmrk2 | 4,10219014 | 3,15E-15 |
| Ndufa10 | 57,6936 | 73,2586 | 131,128 | 110,318 | 371,747 | 566,764 | 586,994 | 817,875 | 784,305 | 815,247 | Ndufa10 | 2,775248211 | 3,86E-15 |
| Ndufb5 | 90,3364 | 113,18 | 126,076 | 113,682 | 272,237 | 425,479 | 420,929 | 594,727 | 566,135 | 566,569 | Ndufb5 | 2,022951608 | 4,01E-15 |
| Nrbp2 | 3,60487 | 5,40009 | 9,36104 | 9,01147 | 32,0541 | 36,5162 | 34,5715 | 10,5544 | 12,4568 | 11,8572 | Nrbp2 | 2,668304748 | 4,01E-15 |
| Coq10a | 4,38478 | 4,19445 | 4,43622 | 3,88175 | 15,9562 | 40,8409 | 38,6852 | 156,41 | 146,535 | 149,459 | Coq10a | 3,187910152 | 4,17E-15 |

| Gene | d9-11 (1) | d9-11 (2) | d9-11(3) | d9-11(4) | newborn(1) | newborn(2) | newborn(3) | adult(1) | adult(2) | adult(3) | Gene | log2(fold) | FDR(*p* -value) |
|---|---|---|---|---|---|---|---|---|---|---|---|---|---|
| Ndufs1 | 35,4038 | 67,01 | 95,5278 | 72,1819 | 276,34 | 491,784 | 512,137 | 551,542 | 624,314 | 614,234 | Ndufs1 | 2,860426988 | 4,40E-15 |
| Cul9 | 2,35983 | 1,45581 | 2,06418 | 1,50036 | 6,30115 | 8,22059 | 7,65376 | 10,2179 | 8,85532 | 9,36255 | Cul9 | 2,330404328 | 4,45E-15 |
| Vamp1 | 0,414372 | 0,613162 | 0,432344 | 0,364186 | 4,60965 | 7,65516 | 8,22417 | 4,19973 | 4,96788 | 5,48245 | Vamp1 | 3,973121225 | 4,51E-15 |
| Msrb3 | 13,9439 | 23,6425 | 33,3169 | 21,4239 | 92,9378 | 77,9612 | 75,2039 | 56,9969 | 71,3061 | 65,3887 | Msrb3 | 2,101558657 | 4,81E-15 |
| Sesn1 | 5,47268 | 6,64597 | 7,84197 | 7,22798 | 91,2681 | 33,3979 | 31,0545 | 132,21 | 123,065 | 127,677 | Sesn1 | 3,089533471 | 5,02E-15 |
| Gm12338 | 159,801 | 253,351 | 241,864 | 280,306 | 580,255 | 844,499 | 889,771 | 1222,67 | 949,698 | 914,707 | Gm12338 | 2,142096181 | 5,29E-15 |
| Esrra | 10,5187 | 18,9759 | 24,5858 | 28,1559 | 86,2662 | 146,065 | 122,095 | 146,721 | 163,168 | 160,463 | Esrra | 2,763272377 | 5,47E-15 |
| Sod2 | 19,8874 | 16,0174 | 38,1118 | 36,8526 | 134,219 | 210,12 | 220,162 | 317,428 | 317,598 | 311,638 | Sod2 | 3,007972722 | 6,06E-15 |
| Ndufv1 | 64,546 | 104,034 | 187,362 | 134,866 | 540,08 | 790,82 | 827,704 | 1030,85 | 971,483 | 964,566 | Ndufv1 | 2,829935356 | 6,81E-15 |
| Pdk4 | 4,29352 | 11,9028 | 17,4155 | 10,9996 | 66,4551 | 180,281 | 176,524 | 810,78 | 213,579 | 470,5 | Pdk4 | 3,833095993 | 6,96E-15 |
| Nlrp10 | 0,142628 | 0,140702 | 1,3096 | 0,574811 | 9,10068 | 16,0263 | 15,8606 | 12,6864 | 14,971 | 14,2969 | Nlrp10 | 4,69018295 | 7,26E-15 |
| 4430402I18Ri | 0,15726 | 0 | 0,616245 | 0 | 7,44987 | 6,82673 | 6,73192 | 5,80688 | 9,44585 | 7,15177 | 4430402I18Ri | 4,877976865 | 7,28E-15 |
| Tcp11l2 | 7,66631 | 11,9628 | 6,86424 | 4,49327 | 38,7429 | 53,2701 | 52,3274 | 109,159 | 91,7304 | 86,2947 | Tcp11l2 | 2,829583067 | 7,86E-15 |
| Gbas | 25,0754 | 73,5708 | 100,056 | 76,8355 | 378,332 | 648,065 | 650,091 | 837,614 | 830,161 | 772,992 | Gbas | 3,281134875 | 8,56E-15 |
| Ldb3 | 72,5186 | 57,8071 | 165,936 | 87,7034 | 450,584 | 738,389 | 740,969 | 970,236 | 909,664 | 1063,33 | Ldb3 | 2,666189388 | 9,19E-15 |
| Mbnl2 | 11,0856 | 18,1344 | 27,039 | 18,2384 | 68,678 | 65,7061 | 62,7877 | 82,5582 | 76,0109 | 77,3121 | Mbnl2 | 2,07584621 | 9,72E-15 |
| Myom1 | 73,9671 | 100,332 | 138,827 | 137,893 | 340,027 | 409,27 | 422,579 | 467,501 | 471,379 | 538,105 | Myom1 | 2,053083293 | 1,07E-14 |
| Ankrd63 | 0 | 0,0646239 | 0,328087 | 0,528023 | 10,9289 | 5,594 | 4,87708 | 0,775027 | 1,57611 | 1,80009 | Ankrd63 | 4,96421815 | 1,22E-14 |
| Cox7a1 | 116,867 | 102,103 | 181,006 | 117,973 | 455,982 | 829,043 | 791,471 | 2989,63 | 2595,68 | 2462,51 | Cox7a1 | 2,794740111 | 1,22E-14 |
| Mtmr1 | 3,44377 | 3,26139 | 3,16205 | 3,53914 | 9,48175 | 12,1437 | 12,3771 | 8,58675 | 8,62118 | 9,06347 | Mtmr1 | 2,074441237 | 1,29E-14 |
| Pkdrej | 0 | 0,0438674 | 0,0371187 | 0,0896112 | 2,19082 | 2,03657 | 2,00187 | 1,53857 | 2,30014 | 1,83859 | Pkdrej | 5,452100413 | 1,35E-14 |
| Sdhd | 74,9617 | 128,492 | 144,628 | 155,423 | 396,32 | 644,796 | 591,217 | 583,237 | 666,51 | 645,606 | Sdhd | 2,39976564 | 1,87E-14 |
| Mgll | 1,49525 | 1,89571 | 3,72022 | 3,01291 | 11,9413 | 18,4511 | 18,1997 | 15,9609 | 22,5754 | 20,8318 | Mgll | 2,967645108 | 2,07E-14 |
| Oxsm | 1,03277 | 1,27353 | 1,36494 | 0,867099 | 5,12097 | 7,40049 | 6,79262 | 12,7609 | 14,7009 | 13,1566 | Oxsm | 2,862529932 | 2,22E-14 |
| Klf9 | 2,19747 | 3,74436 | 2,41784 | 3,22018 | 35,951 | 13,8212 | 14,1175 | 56,2237 | 50,8116 | 66,671 | Klf9 | 3,145057693 | 2,39E-14 |
| Dgat2 | 11,0706 | 12,9871 | 12,6117 | 24,8657 | 69,7166 | 105,719 | 103,073 | 383,188 | 346,095 | 449,852 | Dgat2 | 2,818095942 | 2,51E-14 |
| Gm13375 | 3,38786 | 6,42689 | 8,04742 | 4,72404 | 29,0772 | 24,9737 | 25,7677 | 27,1506 | 30,6848 | 32,3388 | Gm13375 | 2,54308647 | 2,85E-14 |
| Fh1 | 52,4811 | 95,267 | 109,505 | 97,2673 | 285,926 | 509,638 | 495,587 | 531,74 | 576,213 | 584,977 | Fh1 | 2,529357173 | 2,88E-14 |
| Cyth1 | 7,71169 | 8,82473 | 13,3316 | 15,1241 | 37,3986 | 48,1901 | 45,8705 | 31,6947 | 31,4101 | 32,7885 | Cyth1 | 2,274892261 | 3,38E-14 |
| Dld | 30,7244 | 55,9553 | 65,4679 | 69,7123 | 193,501 | 388,796 | 410,32 | 435,122 | 455,154 | 440,658 | Dld | 2,860720628 | 3,57E-14 |
| Tec | 1,04159 | 1,79815 | 1,41277 | 0,918129 | 7,48037 | 11,181 | 9,7862 | 10,206 | 11,7859 | 9,32802 | Tec | 2,876750508 | 3,83E-14 |
| Hagh | 12,3689 | 16,9576 | 19,5339 | 24,5486 | 56,0589 | 92,2404 | 85,5385 | 97,8543 | 77,157 | 95,6617 | Hagh | 2,465014922 | 3,92E-14 |
| Uqcrfs1 | 70,5454 | 143,289 | 171,759 | 155,507 | 482,015 | 741,725 | 780,129 | 1170,8 | 1119,4 | 1122,7 | Uqcrfs1 | 2,57549477 | 4,26E-14 |
| Adrb1 | 0,669037 | 2,24399 | 2,23372 | 3,36988 | 14,1147 | 18,9363 | 18,1039 | 26,2614 | 19,5349 | 19,5166 | Adrb1 | 3,28631164 | 4,27E-14 |
| Decr1 | 13,8703 | 17,7331 | 13,8929 | 14,6442 | 50,136 | 184,53 | 190,452 | 253,756 | 273,582 | 251,234 | Decr1 | 3,447370063 | 4,44E-14 |
| Pm20d2 | 0,454472 | 1,40106 | 1,47003 | 1,14479 | 5,3897 | 7,21088 | 7,58729 | 34,2845 | 39,4164 | 38,3799 | Pm20d2 | 2,832961012 | 4,50E-14 |
| Trak1 | 10,6881 | 14,4656 | 25,8944 | 25,8058 | 80,6769 | 106,856 | 106,657 | 107,176 | 97,5952 | 108,6 | Trak1 | 2,706304027 | 4,70E-14 |
| Pank1 | 2,61914 | 5,0009 | 6,87626 | 3,36268 | 19,3528 | 17,0204 | 16,5589 | 21,7912 | 18,2398 | 17,6243 | Pank1 | 2,27390202 | 4,89E-14 |
| Nr3c1 | 0,99544 | 1,42391 | 3,94691 | 2,45734 | 15,8559 | 18,0272 | 17,6859 | 13,0955 | 18,8751 | 14,2475 | Nr3c1 | 3,220624031 | 6,49E-14 |
| Klhl24 | 8,46841 | 19,5394 | 18,654 | 18,5824 | 63,9136 | 54,8784 | 51,5354 | 73,9969 | 72,3877 | 78,7339 | Klhl24 | 2,113106716 | 6,93E-14 |
| Hist1h1c | 19,3918 | 24,5506 | 23,7692 | 34,7004 | 207,179 | 96,3651 | 88,4396 | 264,867 | 271,195 | 396,954 | Hist1h1c | 2,830243049 | 7,38E-14 |
| Pdk2 | 4,75919 | 9,24301 | 29,296 | 23,2217 | 168,584 | 264,458 | 262,807 | 384,393 | 404,316 | 396,199 | Pdk2 | 3,963930958 | 8,15E-14 |
| D830015G02F | 0 | 0 | 0 | 0 | 0 | 0 | 0 | 0 | 0 | 0 | D830015G02F | 4,557477721 | 8,21E-14 |
| Cd1d1 | 1,87161 | 2,21555 | 6,32659 | 2,82777 | 22,0174 | 40,874 | 34,0294 | 2,54771 | 2,04842 | 1,94518 | Cd1d1 | 3,679449044 | 8,77E-14 |
| Vldlr | 2,88734 | 5,54685 | 6,00379 | 8,65146 | 24,6107 | 44,2429 | 43,707 | 95,2341 | 87,8579 | 95,7535 | Vldlr | 2,978414268 | 9,54E-14 |
| Immt | 63,0081 | 94,4918 | 109,378 | 96,9546 | 239,777 | 426,627 | 442,706 | 458,656 | 471,826 | 463,997 | Immt | 2,2652273 | 1,02E-13 |
| Reep5 | 25,7713 | 38,8628 | 32,7879 | 27,5641 | 80,4393 | 150,75 | 153,112 | 189,588 | 215,192 | 199,805 | Reep5 | 2,334965727 | 1,05E-13 |
| Plscr2 | 0 | 0 | 0 | 0,36481 | 5,6098 | 7,0633 | 7,29583 | 10,0408 | 12,6376 | 11,3656 | Plscr2 | 6,13619349 | 1,07E-13 |
| Slc25a20 | 8,35644 | 21,4712 | 27,7361 | 24,0762 | 92,1458 | 215,473 | 206,393 | 225,965 | 234,136 | 209,796 | Slc25a20 | 3,305335244 | 1,38E-13 |
| Ptp4a3 | 16,6652 | 22,2156 | 41,2644 | 28,9255 | 103,191 | 202,67 | 212,285 | 212,338 | 167,702 | 210,682 | Ptp4a3 | 2,913526095 | 1,46E-13 |
| Lbh | 56,4542 | 115,638 | 172,499 | 133,317 | 481,507 | 652,233 | 650,758 | 391,123 | 404,959 | 358,374 | Lbh | 2,607193869 | 1,47E-13 |
| Fundc2 | 9,2954 | 13,9027 | 14,6 | 14,9031 | 34,1193 | 53,605 | 51,6655 | 57,9874 | 69,3556 | 68,2356 | Fundc2 | 2,083845245 | 1,57E-13 |
| Ghitm | 38,369 | 63,3534 | 75,0023 | 72,9951 | 179,652 | 295,022 | 296,307 | 380,116 | 405,292 | 405,101 | Ghitm | 2,326905862 | 1,69E-13 |
| Cers4 | 7,6699 | 6,17482 | 7,35877 | 6,57272 | 19,3596 | 34,2863 | 31,9271 | 14,3539 | 20,9858 | 18,6268 | Cers4 | 2,287035683 | 1,82E-13 |
| Slc25a11 | 66,2696 | 76,1471 | 79,4908 | 85,5822 | 186,663 | 328,702 | 326,869 | 554,551 | 578,418 | 552,269 | Slc25a11 | 2,126882228 | 1,89E-13 |
| Clasp1 | 27,6713 | 28,3752 | 45,0702 | 39,7503 | 90,8177 | 139,408 | 137,581 | 88,394 | 124,405 | 90,0578 | Clasp1 | 2,151597394 | 1,97E-13 |
| Dmpk | 17,349 | 20,1745 | 35,6072 | 19,7679 | 78,4624 | 197,27 | 204,454 | 497,43 | 444,239 | 494,37 | Dmpk | 3,032867668 | 1,98E-13 |
| Cycs | 23,1064 | 45,395 | 40,763 | 37,0299 | 80,9805 | 171,085 | 174,572 | 232,583 | 239,167 | 234,362 | Cycs | 2,404823041 | 2,50E-13 |
| Cox6c | 91,4748 | 210,951 | 188,033 | 189,9 | 659,818 | 953,524 | 1037,76 | 1315,08 | 1103,24 | 1074,07 | Cox6c | 2,208278293 | 2,53E-13 |
| Gpd1 | 1,77725 | 1,40259 | 0,593379 | 0,9549 | 7,62469 | 17,3545 | 18,5875 | 54,1708 | 55,1398 | 61,7163 | Gpd1 | 3,669001021 | 2,65E-13 |
| Uqcrq | 232,431 | 245,842 | 361,846 | 330,342 | 766,566 | 1193,2 | 1190,19 | 1946,56 | 1558,95 | 1681,59 | Uqcrq | 2,122798561 | 2,72E-13 |
| Atp5a1 | 393,751 | 577,968 | 856,451 | 717,91 | 1876,11 | 2501,94 | 2649,58 | 3496,94 | 2980,19 | 2963,71 | Atp5a1 | 2,180492461 | 3,37E-13 |
| Myadml2 | 0,58237 | 1,29262 | 1,7013 | 2,7866 | 11,3141 | 23,3886 | 23,0985 | 47,8754 | 53,9364 | 54,3873 | Myadml2 | 3,768819954 | 3,45E-13 |
| Pdhx | 12,9199 | 19,7682 | 18,4868 | 14,8747 | 44,323 | 76,196 | 73,1251 | 118,896 | 108,655 | 121,924 | Pdhx | 2,194846415 | 3,73E-13 |
| Slc6a8 | 10,3365 | 14,6932 | 20,1288 | 27,1998 | 80,7201 | 75,5095 | 78,8196 | 83,0987 | 78,2178 | 78,0613 | Slc6a8 | 2,332652753 | 3,93E-13 |
| Acot1 | 2,3669 | 5,44802 | 8,24636 | 12,7684 | 67,5734 | 163,953 | 161,67 | 56,6228 | 38,7536 | 33,5763 | Acot1 | 4,189255731 | 4,16E-13 |
| Paqr9 | 0,72866 | 1,58138 | 2,31112 | 1,90844 | 9,01039 | 30,8003 | 32,0202 | 42,4936 | 42,9266 | 39,3648 | Paqr9 | 4,031531755 | 5,17E-13 |
| Cpe | 13,4633 | 25,7624 | 44,0012 | 33,8226 | 152,179 | 148,52 | 149,48 | 276,976 | 273,372 | 286,952 | Cpe | 2,574514889 | 5,33E-13 |
| Uqcrc1 | 126,264 | 165,761 | 260,897 | 235,526 | 605,85 | 881,031 | 901,385 | 1337,78 | 1263,05 | 1287,64 | Uqcrc1 | 2,255522455 | 5,55E-13 |
| Gm15816 | 0 | 0 | 0 | 0 | 10,9465 | 32,1816 | 33,8649 | 116,94 | 119,222 | 118,125 | Gm15816 | 5,436039119 | 5,87E-13 |
| Fbxo9 | 11,6192 | 19,5166 | 13,2366 | 15,5223 | 39,9085 | 65,8995 | 65,8341 | 28,3758 | 32,9845 | 35,3509 | Fbxo9 | 2,24848654 | 5,93E-13 |
| Suclg1 | 67,0942 | 99,1507 | 114,154 | 112,18 | 258,067 | 442,618 | 461,087 | 516,325 | 539,513 | 533,71 | Suclg1 | 2,260660808 | 5,99E-13 |
| 1110007C09R | 21,2757 | 20,9864 | 51,5262 | 31,8576 | 88,5249 | 94,8365 | 98,9665 | 112,152 | 106,258 | 107,941 | 1110007C09R | 2,38520278 | 6,01E-13 |
| Smim5 | 0,842388 | 5,40117 | 3,86625 | 5,93604 | 24,7194 | 47,2325 | 43,3668 | 22,8093 | 24,6949 | 32,8518 | Smim5 | 3,671606222 | 6,06E-13 |
| Nr3c2 | 0 | 0,0775513 | 0,0656192 | 0,237611 | 2,27169 | 4,84469 | 4,72194 | 4,72043 | 9,6226 | 5,89898 | Nr3c2 | 5,249153608 | 7,51E-13 |
| Pcdhga2 | 0 | 0 | 0 | 0 | 0 | 38,9542 | 0 | 0 | 12,7859 | 14,5146 | Pcdhga2 | 2,78872766 | 7,51E-13 |
| Rnasel | 0,133989 | 0,33045 | 1,84542 | 0,57375 | 12,1781 | 24,6799 | 23,2323 | 0,365807 | 0,475769 | 0,303229 | Rnasel | 4,752508054 | 7,91E-13 |
| Eci1 | 13,2374 | 54,4652 | 58,4814 | 32,6188 | 196,006 | 454,714 | 463,181 | 546,165 | 524,74 | 492,514 | Eci1 | 3,4550766 | 8,25E-13 |
| Mlycd | 2,58568 | 3,1884 | 8,49774 | 11,3951 | 46,1437 | 102,533 | 99,3146 | 122,384 | 99,0766 | 105,081 | Mlycd | 3,86615683 | 8,97E-13 |
| Fn3k | 0 | 0,380686 | 2,94948 | 0,725457 | 19,658 | 42,4356 | 37,5709 | 33,1175 | 36,726 | 34,5646 | Fn3k | 4,695225032 | 9,14E-13 |
| Sgcb | 5,89407 | 8,55071 | 11,2867 | 12,0512 | 28,268 | 39,9201 | 38,0445 | 75,6904 | 77,3373 | 73,671 | Sgcb | 2,225884665 | 9,14E-13 |
| Fgf16 | 0 | 0 | 0 | 0,10858 | 3,55891 | 6,05317 | 6,3123 | 11,3667 | 11,4836 | 6,63073 | Fgf16 | 6,605059298 | 9,21E-13 |
| Xirp1 | 18,073 | 19,3237 | 31,7756 | 36,6107 | 105,867 | 96,6914 | 90,4805 | 97,0383 | 113,679 | 98,9933 | Xirp1 | 2,155348923 | 9,51E-13 |
| Krcc1 | 11,1245 | 14,1094 | 14,5905 | 12,8055 | 32,5367 | 47,9924 | 51,3526 | 29,7373 | 38,7622 | 32,6932 | Krcc1 | 2,01721145 | 1,01E-12 |
| Cacna2d1 | 1,5246 | 3,96897 | 5,37339 | 3,67693 | 17,0551 | 23,0352 | 23,0226 | 54,4739 | 70,3091 | 70,4961 | Cacna2d1 | 2,722410815 | 1,02E-12 |
| Rftn1 | 5,17897 | 7,54739 | 8,35091 | 6,40318 | 20,154 | 35,8378 | 35,5943 | 12,8387 | 25,6245 | 18,4662 | Rftn1 | 2,341968074 | 1,03E-12 |
| Gpr146 | 2,68812 | 6,0056 | 4,28964 | 5,49631 | 19,8455 | 14,5183 | 15,2404 | 35,3895 | 37,8126 | 38,7679 | Gpr146 | 2,124779908 | 1,24E-12 |
| BC002163 | 32,00496 | 44,5421 | 56,733 | 69,1227 | 106,834 | 176,785 | 187,4895 | 268,249 | 271,05945 | 247,226 | BC002163 | 2,358621674 | 1,24E-12 |
| Mccc1 | 9,25332 | 12,8231 | 18,0219 | 12,2076 | 39,6242 | 71,4305 | 70,6348 | 87,5295 | 102,046 | 96,8516 | Mccc1 | 2,427188501 | 1,33E-12 |
| Deb1 | 18,2614 | 28,026 | 55,873 | 29,5914 | 101,162 | 164,306 | 174,829 | 187,075 | 203,088 | 199,173 | Deb1 | 2,480139818 | 1,38E-12 |
| Irx4 | 12,7631 | 9,19119 | 17,0983 | 23,5293 | 67,4056 | 86,1113 | 88,7906 | 21,8053 | 26,2476 | 26,9658 | Irx4 | 2,659282744 | 1,40E-12 |
| Nckap5 | 0,443637 | 0,962832 | 0,444385 | 0,40231 | 5,37016 | 4,1064 | 2,76975 | 3,00241 | 2,12322 | 4,35302 | Nckap5 | 2,737425791 | 1,57E-12 |
| Acat1 | 5,10933 | 20,1613 | 36,8541 | 23,0182 | 144,652 | 329,221 | 335,962 | 366,959 | 386,72 | 362,736 | Acat1 | 3,84153813 | 1,57E-12 |
| Mtus1 | 11,5472 | 21,8713 | 30,1124 | 19,502 | 64,2198 | 91,5431 | 90,9351 | 67,6368 | 70,9214 | 70,7976 | Mtus1 | 2,407121133 | 1,60E-12 |
| Pam | 67,4851 | 101,174 | 163,906 | 156,375 | 469,93 | 490,077 | 486,15 | 252,609 | 246,316 | 223,755 | Pam | 2,272574318 | 1,69E-12 |
| Pla2g16 | 2,48474 | 6,4911 | 8,91063 | 7,78817 | 29,0505 | 39,8076 | 36,2689 | 61,0173 | 63,8873 | 57,0004 | Pla2g16 | 2,762721699 | 1,73E-12 |
| Dlst | 58,3474 | 93,6954 | 124,991 | 100,491 | 248,862 | 386,954 | 388,588 | 453,337 | 494,288 | 501,545 | Dlst | 2,156616751 | 1,92E-12 |
| Slc35f5 | 13,296 | 15,6387 | 23,1562 | 13,5231 | 42,8822 | 68,1243 | 66,8474 | 86,7865 | 97,6194 | 93,7111 | Slc35f5 | 2,127543057 | 1,96E-12 |
| Bcam | 45,5729 | 64,8313 | 92,5449 | 105,954 | 253,827 | 336,308 | 350,814 | 162,585 | 154,446 | 151,705 | Bcam | 2,31482141 | 1,97E-12 |
| Psmb10 | 6,93813 | 18,5362 | 17,1298 | 12,2267 | 47,0085 | 50,5706 | 50,7123 | 33,7624 | 31,9452 | 34,4046 | Psmb10 | 2,110222218 | 2,24E-12 |
| Nfia | 0,589012 | 2,55547 | 4,28851 | 1,97836 | 28,743 | 21,044 | 21,459 | 13,322 | 17,8849 | 14,8234 | Nfia | 3,490606317 | 2,56E-12 |
| Ak1 | 85,0407 | 65,0682 | 129,603 | 86,8219 | 278,873 | 559,168 | 569,884 | 666,506 | 714,216 | 722,437 | Ak1 | 2,61039143 | 2,56E-12 |
| Adamtsl5 | 0 | 1,16912 | 0,836621 | 1,41145 | 12,6168 | 14,1563 | 13,3913 | 45,9517 | 40,4284 | 53,6592 | Adamtsl5 | 4,098149212 | 2,88E-12 |
| Srl | 9,36106 | 25,6555 | 66,4316 | 54,6379 | 428,057 | 441,02 | 448,196 | 323,949 | 374,076 | 392,466 | Srl | 3,671006905 | 3,17E-12 |
| Prkaa2 | 8,00116 | 9,77254 | 19,6233 | 12,17 | 43,6337 | 46,6609 | 47,7855 | 35,0612 | 44,2788 | 40,3626 | Prkaa2 | 2,182138169 | 4,20E-12 |
| Etfb | 85,1122 | 128,995 | 199,012 | 149,931 | 439,974 | 841,738 | 892,539 | 1257,27 | 1131,87 | 1033,93 | Etfb | 2,652025083 | 4,31E-12 |
| Ppapdc3 | 0,946875 | 2,98901 | 2,84507 | 4,19643 | 15,5332 | 21,0609 | 23,3531 | 28,6986 | 35,0167 | 33,6159 | Ppapdc3 | 3,00371832 | 4,36E-12 |
| Tmem182 | 0,69274 | 8,88349 | 5,78113 | 1,04623 | 50,8565 | 113,792 | 102,825 | 151,259 | 144,827 | 131,974 | Tmem182 | 4,614661154 | 4,81E-12 |
| Hmgcs2 | 0 | 0,487714 | 0,825338 | 1,29502 | 32,1134 | 301,898 | 311,905 | 6,38408 | 7,64875 | 4,7369 | Hmgcs2 | 7,137355159 | 5,13E-12 |
| Mr1 | 0 | 0,654811 | 0,775661 | 0,668683 | 4,30082 | 6,86191 | 6,27219 | 4,70451 | 4,45521 | 4,86213 | Mr1 | 3,714280214 | 6,17E-12 |
| Ccser2 | 3,21794 | 7,66832 | 10,4876 | 6,51607 | 25,3653 | 38,6834 | 38,5656 | 46,4724 | 52,5137 | 57,2239 | Ccser2 | 2,428731518 | 6,45E-12 |
| Pdhb | 51,4482 | 72,931 | 98,4322 | 78,0843 | 290,656 | 480,125 | 483,814 | 452,752 | 482,748 | 512,095 | Pdhb | 2,253162891 | 7,64E-12 |
| Sntb1 | 1,25743 | 1,86065 | 9,18337 | 5,06656 | 51,4384 | 51,8982 | 48,3895 | 1,71286 | 1,7434 | 3,00846 | Sntb1 | 3,634131737 | 7,66E-12 |

| Gene | d9-11 (1) | d9-11 (2) | d9-11(3) | d9-11(4) | newborn(1) | newborn(2) | newborn(3) | adult(1) | adult(2) | adult(3) | Gene | log2(fold) | FDR(*p*-value) |
|---|---|---|---|---|---|---|---|---|---|---|---|---|---|
| Cisd1 | 26,258 | 48,1889 | 39,9172 | 36,4748 | 105,635 | 196,159 | 203,951 | 342,561 | 332,256 | 328,731 | Cisd1 | 2,20985393 | 8,82E-12 |
| Tom1l2 | 7,18726 | 9,08392 | 14,1225 | 10,5768 | 26,7524 | 42,4242 | 40,6417 | 58,0801 | 63,1253 | 56,8497 | Tom1l2 | 2,141664439 | 9,05E-12 |
| Map7 | 1,16565 | 1,31419 | 1,80697 | 3,52321 | 13,3189 | 11,5153 | 10,4674 | 15,6787 | 13,2585 | 18,3426 | Map7 | 2,775867622 | 1,03E-11 |
| Hibadh | 18,8579 | 32,456 | 34,4928 | 39,6046 | 111,241 | 192,182 | 188,427 | 274,882 | 315,819 | 309,688 | Hibadh | 2,347185027 | 1,03E-11 |
| Trabd2b | 1,09324 | 2,25075 | 4,64217 | 2,49044 | 20,9629 | 15,0376 | 12,403 | 36,8711 | 41,6782 | 48,0223 | Trabd2b | 2,878747878 | 1,15E-11 |
| Ky | 0,11629 | 0 | 0,630957 | 0,058585 | 3,55694 | 11,7189 | 10,9364 | 5,55716 | 10,429 | 11,1999 | Ky | 5,283795288 | 1,17E-11 |
| Il10rb | 3,16809 | 7,3535 | 13,0655 | 8,25922 | 34,6425 | 62,7561 | 64,5034 | 84,5046 | 108,44 | 83,7457 | Il10rb | 2,997095865 | 1,19E-11 |
| Cyp2d22 | 0,715362 | 0,94129 | 1,5904 | 0,600558 | 11,318 | 5,81565 | 5,68689 | 1,73541 | 2,6683 | 1,66273 | Cyp2d22 | 3,248572613 | 1,21E-11 |
| Clic5 | 1,02246 | 2,54821 | 5,07598 | 7,04884 | 49,7474 | 36,4205 | 36,5001 | 80,6591 | 62,015 | 83,1467 | Clic5 | 3,556878644 | 1,29E-11 |
| Sgcd | 1,03006 | 2,28626 | 4,29841 | 2,33404 | 22,9508 | 19,3901 | 19,51 | 38,7296 | 38,8811 | 43,6366 | Sgcd | 2,930140223 | 1,32E-11 |
| Mtfp1 | 7,2288 | 7,13091 | 7,75683 | 9,09992 | 25,1612 | 71,2579 | 62,4119 | 208,172 | 211,881 | 207,001 | Mtfp1 | 2,97382666 | 1,35E-11 |
| Iqsec2 | 3,23145 | 2,52321 | 1,85051 | 2,16498 | 6,56494 | 6,58759 | 7,21572 | 8,02557 | 9,12824 | 9,9139 | Iqsec2 | 2,164268179 | 1,49E-11 |
| Adamts7 | 2,49113 | 2,92561 | 6,9314 | 3,34664 | 14,9338 | 20,2858 | 19,3055 | 8,63791 | 10,2628 | 10,6394 | Adamts7 | 2,541382306 | 1,52E-11 |
| Wfdc1 | 0 | 0,630903 | 1,60126 | 0,643999 | 9,09551 | 19,9276 | 19,8924 | 31,2453 | 29,4336 | 32,1525 | Wfdc1 | 4,397645166 | 1,55E-11 |
| Klf15 | 1,61154 | 3,90214 | 1,95652 | 3,54194 | 47,3569 | 15,9969 | 15,3309 | 52,1665 | 56,1965 | 37,1516 | Klf15 | 3,408115363 | 1,59E-11 |
| Il15ra | 1,00704 | 4,15593 | 0,515582 | 1,8562 | 11,3008 | 16,349 | 14,3301 | 14,0767 | 11,7406 | 13,4831 | Il15ra | 3,424278063 | 1,66E-11 |
| Ntn4 | 0,138688 | 2,46265 | 4,16734 | 3,07363 | 29,4974 | 31,8811 | 31,1917 | 2,01602 | 6,2391 | 3,50692 | Ntn4 | 3,911889753 | 1,68E-11 |
| Ggnbp1 | 1,96652 | 2,741 | 2,71891 | 2,10918 | 13,0243 | 30,5545 | 35,0601 | 5,53201 | 9,0362 | 11,2371 | Ggnbp1 | 3,631019253 | 1,73E-11 |
| 1110001J03Ri | 48,6453 | 61,387 | 77,8804 | 67,9964 | 168,331 | 275,217 | 224,388 | 281,206 | 243,813 | 278,765 | 1110001J03Ri | 2,086607215 | 1,82E-11 |
| Pou6f1 | 0,418848 | 1,10185 | 2,21427 | 0,844014 | 10,1046 | 9,59853 | 8,7911 | 3,04917 | 2,66003 | 3,0697 | Pou6f1 | 3,130462733 | 1,86E-11 |
| Usp20 | 4,01522 | 4,03718 | 10,377 | 9,0244 | 32,7472 | 36,1426 | 39,718 | 18,9683 | 22,8867 | 22,4886 | Usp20 | 2,639976607 | 1,87E-11 |
| Nfic | 4,18846 | 6,9512 | 13,8674 | 7,91757 | 38,6168 | 38,0948 | 36,7911 | 43,9493 | 42,962 | 56,665 | Nfic | 2,448902835 | 1,93E-11 |
| Dnaja4 | 3,60132 | 3,01439 | 10,02 | 4,61762 | 24,6845 | 46,9558 | 43,6546 | 59,7282 | 67,7799 | 66,737 | Dnaja4 | 3,100448893 | 1,97E-11 |
| Fam129a | 0,326957 | 2,09653 | 0,818746 | 0,98824 | 16,2804 | 7,49531 | 8,04147 | 1,41286 | 1,47649 | 0,925438 | Fam129a | 3,48415098 | 1,98E-11 |
| Kcnj11 | 1,3683 | 3,06474 | 8,06221 | 8,67314 | 47,8781 | 70,5019 | 68,9538 | 93,4538 | 117,195 | 98,5414 | Kcnj11 | 3,747672924 | 2,02E-11 |
| Rbfox1 | 0,157307 | 0,54314 | 0,590878 | 0,237734 | 4,76857 | 15,8486 | 14,2776 | 23,5428 | 31,5221 | 25,6985 | Rbfox1 | 4,143978152 | 2,28E-11 |
| Rnf128 | 3,36339 | 8,41336 | 10,0263 | 9,55993 | 29,2184 | 46,2779 | 52,501 | 65,3987 | 68,193 | 60,3552 | Rnf128 | 2,68872969 | 2,31E-11 |
| Ror1 | 1,73853 | 3,1593 | 3,20782 | 5,34682 | 18,8759 | 14,5155 | 12,9304 | 0,749004 | 2,20513 | 1,45089 | Ror1 | 2,401839327 | 2,32E-11 |
| Asb15 | 0 | 0 | 0,125374 | 0 | 1,3616 | 5,98216 | 6,19057 | 32,0814 | 42,1483 | 35,4632 | Asb15 | 6,245934014 | 2,41E-11 |
| Ogdh | 34,9781 | 48,2932 | 81,6752 | 59,7548 | 159,17 | 396,175 | 404,487 | 745,411 | 719,593 | 698,786 | Ogdh | 2,78359652 | 2,41E-11 |
| Lclat1 | 9,2904 | 14,9863 | 19,9996 | 21,0568 | 46,0662 | 90,4883 | 90,6578 | 45,5902 | 61,6387 | 50,8706 | Lclat1 | 2,545431916 | 2,41E-11 |
| Dlat | 24,7816 | 41,8979 | 54,2747 | 47,2061 | 108,418 | 203,926 | 198,647 | 250,685 | 266,352 | 282,188 | Dlat | 2,302684643 | 2,41E-11 |
| Nfib | 2,52152 | 7,39326 | 12,3366 | 5,65758 | 47,0353 | 41,6121 | 42,3494 | 25,2287 | 33,1912 | 27,7568 | Nfib | 2,890310304 | 2,42E-11 |
| Trdn | 0,778792 | 2,65405 | 7,15081 | 4,13731 | 28,4585 | 51,3161 | 53,0157 | 77,342 | 92,7506 | 87,5976 | Trdn | 3,823319692 | 2,42E-11 |
| Ehhadh | 0,108936 | 0,967176 | 0,727423 | 0,877972 | 4,84776 | 6,41888 | 7,11517 | 16,5395 | 7,77507 | 10,9353 | Ehhadh | 3,410165612 | 2,51E-11 |
| Gpr22 | 0 | 0,343383 | 0,581102 | 0,280565 | 2,7312 | 8,02907 | 7,77504 | 11,3397 | 16,684 | 16,7018 | Gpr22 | 4,414127528 | 2,74E-11 |
| 2300009A05R | 4,79838 | 8,67729 | 12,677 | 6,43541 | 39,6491 | 55,2254 | 59,163 | 27,9088 | 35,2902 | 42,2558 | 2300009A05R | 2,716740875 | 2,74E-11 |
| Ndufa6 | 96,8011 | 194,481 | 183,755 | 181,325 | 402,934 | 678,09 | 712,289 | 1022,38 | 793,757 | 833,678 | Ndufa6 | 2,162517198 | 3,42E-11 |
| Trim63 | 9,29916 | 22,6637 | 55,3957 | 33,7935 | 249,262 | 206,382 | 201,61 | 163,998 | 213,189 | 238,347 | Trim63 | 3,058464046 | 3,65E-11 |
| Palmd | 1,18748 | 4,13648 | 4,19989 | 1,54882 | 14,3092 | 14,1363 | 13,9811 | 27,3012 | 29,0715 | 34,8462 | Palmd | 2,958046017 | 3,68E-11 |
| Pdp1 | 2,63068 | 6,10647 | 4,4959 | 4,8796 | 11,9195 | 16,6937 | 17,0139 | 14,2538 | 18,8081 | 15,3571 | Pdp1 | 2,083570474 | 3,84E-11 |
| Fyco1 | 3,85936 | 7,71476 | 16,0824 | 12,3348 | 42,8983 | 95,368 | 89,4643 | 130,093 | 144,148 | 151,238 | Fyco1 | 3,191580217 | 3,95E-11 |
| Zfp30 | 0,892295 | 0,75449 | 1,59596 | 0,513655 | 5,2406 | 5,27656 | 5,74811 | 2,66433 | 3,87014 | 3,84148 | Zfp30 | 2,827387053 | 4,20E-11 |
| Sh3rf2 | 0 | 0,13425 | 0,0567974 | 0 | 4,08496 | 1,96846 | 1,81437 | 10,9049 | 11,2196 | 12,1461 | Sh3rf2 | 5,167847898 | 4,35E-11 |
| Frmd5 | 0,992965 | 3,08938 | 6,18445 | 5,23305 | 21,8335 | 28,2733 | 26,2781 | 24,393 | 36,218 | 29,236 | Frmd5 | 3,221068024 | 4,45E-11 |
| Gm16793 | 0 | 0 | 0 | 0 | 0 | 14,8578 | 14,77 | 0 | 10,7281 | 0 | Gm16793 | 4,926848795 | 4,64E-11 |
| Insig2 | 4,87934 | 5,68847 | 7,47568 | 6,82241 | 15,9967 | 24,7433 | 26,4947 | 32,3471 | 35,7552 | 33,9461 | Insig2 | 2,087472649 | 5,23E-11 |
| 4930429F24R | 1,40152 | 0,230425 | 1,94953 | 0,705759 | 10,5487 | 7,8181 | 7,89883 | 11,6462 | 12,2289 | 16,1881 | 4930429F24R | 3,16773546 | 5,32E-11 |
| Aco2 | 80,0323 | 120,191 | 200,505 | 166,419 | 625,591 | 968,516 | 1021,35 | 1223,81 | 1209,64 | 1200,97 | Aco2 | 2,331719901 | 5,39E-11 |
| Hadhb | 5,41641 | 18,1255 | 23,1669 | 19,1786 | 83,6425 | 272,881 | 255,905 | 490,385 | 467,747 | 430,524 | Hadhb | 3,379503692 | 5,66E-11 |
| Vsig10l | 0,28293 | 0,186073 | 0,314884 | 0,285047 | 2,59238 | 2,93101 | 3,00677 | 1,28657 | 1,01032 | 1,0988 | Vsig10l | 3,439750058 | 5,73E-11 |
| Inpp4b | 0,467448 | 0,537993 | 0,65031 | 0,627951 | 2,47202 | 6,16734 | 5,25255 | 6,73376 | 8,2853 | 8,70172 | Inpp4b | 3,286193622 | 5,95E-11 |
| Pccb | 13,6446 | 25,9796 | 43,2281 | 26,7508 | 90,4086 | 116,106 | 114,143 | 160,723 | 149,674 | 145,231 | Pccb | 2,26758254 | 7,45E-11 |
| Eci2 | 20,7775 | 31,0832 | 54,2829 | 43,7999 | 138,438 | 252,943 | 245,298 | 223,668 | 223,832 | 219,534 | Eci2 | 2,699565334 | 7,45E-11 |
| Mpc1 | 1,46129 | 4,6802 | 9,57584 | 5,55355 | 65,6373 | 96,5881 | 98,4841 | 298,762 | 300,649 | 280,089 | Mpc1 | 3,270967704 | 7,82E-11 |
| Sorbs2 | 21,2535 | 30,8683 | 59,4508 | 49,6532 | 142,817 | 174,922 | 177,837 | 88,5722 | 96,9924 | 106,875 | Sorbs2 | 2,461403136 | 8,57E-11 |
| Mybpc3 | 217,208 | 283,398 | 493,708 | 445,827 | 978,291 | 1598,16 | 1670,32 | 1894,97 | 1500,67 | 1849,87 | Mybpc3 | 2,295577093 | 9,07E-11 |
| Retsat | 6,36938 | 15,01 | 6,34989 | 10,5147 | 29,0437 | 56,8968 | 55,3058 | 46,2524 | 53,5589 | 53,9369 | Retsat | 2,513150339 | 9,79E-11 |
| Tmem191c | 1,78898 | 0,756323 | 1,91966 | 1,54427 | 6,46261 | 12,1633 | 11,4363 | 0,463114 | 1,46953 | 1,16511 | Tmem191c | 2,790238152 | 1,04E-10 |
| D830005E20R | 0 | 0 | 0 | 0 | 4,24681 | 7,90495 | 6,7787 | 8,48717 | 8,78696 | 9,81102 | D830005E20R | 6,918598211 | 1,05E-10 |
| Acot13 | 12,9936 | 33,964 | 44,9901 | 56,8684 | 150,292 | 329,827 | 275,906 | 389,64 | 414,318 | 472,719 | Acot13 | 2,889750931 | 1,06E-10 |
| Irgq | 3,94875 | 3,3835 | 7,9661 | 5,67787 | 15,8567 | 17,3909 | 17,1191 | 8,7251 | 10,5812 | 10,6871 | Irgq | 2,057509895 | 1,09E-10 |
| Ndrg2 | 12,1849 | 44,024 | 97,7766 | 70,091 | 458,331 | 559,035 | 538,933 | 1080,26 | 1096,69 | 1096,32 | Ndrg2 | 3,430000588 | 1,09E-10 |
| Sdhb | 73,4492 | 132,275 | 217,872 | 171,717 | 505,485 | 633,737 | 592,03 | 1005,39 | 1008,45 | 973,942 | Sdhb | 2,226846509 | 1,10E-10 |
| Impa2 | 7,55107 | 16,6956 | 30,4236 | 19,1422 | 66,243 | 139,603 | 140,287 | 33,4966 | 40,7012 | 30,954 | Impa2 | 2,896711524 | 1,23E-10 |
| Gpam | 7,75123 | 8,64395 | 10,9006 | 9,67688 | 20,8159 | 44,6059 | 44,0021 | 93,8106 | 75,6178 | 94,1991 | Gpam | 2,288834437 | 1,51E-10 |
| Obscn | 7,25855 | 13,7912 | 25,38 | 23,0032 | 68,7151 | 81,2304 | 77,7784 | 102,297 | 98,9827 | 105,534 | Obscn | 2,58549512 | 1,55E-10 |
| Pla2g4e | 0,0714143 | 0 | 0,59611 | 0,0719526 | 1,96229 | 4,18551 | 4,05222 | 2,72931 | 4,39262 | 4,15657 | Pla2g4e | 4,702813908 | 1,59E-10 |
| Ndufc2 | 118,203 | 161,288 | 206,556 | 181,032 | 370,153 | 656,693 | 678,16 | 607,766 | 596,774 | 566,318 | Ndufc2 | 2,049286991 | 1,62E-10 |
| Fgf18 | 1,38241 | 2,27283 | 2,11525 | 0,696142 | 9,29194 | 9,73994 | 10,9069 | 1,87989 | 3,50121 | 4,04756 | Fgf18 | 2,909678754 | 1,64E-10 |
| Coq5 | 16,7166 | 21,5386 | 29,6135 | 25,7731 | 53,2864 | 96,0778 | 96,3632 | 136,739 | 137,919 | 143,591 | Coq5 | 2,096614776 | 1,68E-10 |
| Acacb | 1,47168 | 4,10139 | 12,5918 | 9,9354 | 51,2898 | 107,37 | 101,133 | 108,113 | 102,724 | 114,671 | Acacb | 3,805089872 | 1,81E-10 |
| Kbtbd12 | 0,333486 | 0,164489 | 1,1338 | 1,51164 | 8,08956 | 17,8407 | 17,1999 | 9,08343 | 9,94471 | 11,8779 | Kbtbd12 | 4,124197692 | 1,83E-10 |
| Hspa12a | 2,37148 | 1,79541 | 2,99234 | 2,94513 | 6,77846 | 13,9863 | 14,1567 | 7,53053 | 13,021 | 9,12493 | Hspa12a | 2,477044495 | 1,90E-10 |
| Pstpip2 | 0,148798 | 0,293572 | 0,124196 | 0,299781 | 3,16527 | 6,49937 | 6,75835 | 19,5973 | 24,1934 | 19,1578 | Pstpip2 | 4,619739439 | 1,93E-10 |
| Trak2 | 11,0612 | 14,7983 | 28,7112 | 17,6082 | 57,6632 | 63,3989 | 61,6677 | 61,8917 | 69,0965 | 70,0322 | Trak2 | 2,042774692 | 1,97E-10 |
| D5Ertd579e | 3,18166 | 9,65041 | 10,4186 | 8,30166 | 26,3398 | 40,6331 | 38,4977 | 35,5179 | 40,2125 | 41,383 | D5Ertd579e | 2,484131978 | 2,11E-10 |
| Spsb1 | 1,71593 | 3,4913 | 4,1178 | 4,97004 | 12,7822 | 13,6866 | 14,3708 | 21,7439 | 26,6979 | 23,162 | Spsb1 | 2,226520654 | 2,12E-10 |
| Fahd1 | 3,32518 | 8,07423 | 12,1681 | 14,9405 | 41,134 | 84,1522 | 84,9595 | 80,6492 | 85,6514 | 82,127 | Fahd1 | 3,084417338 | 2,12E-10 |
| Ampd3 | 1,05159 | 2,23438 | 3,24102 | 2,93398 | 8,11296 | 20,9045 | 19,5106 | 17,7722 | 22,4576 | 23,8634 | Ampd3 | 3,01456093 | 2,13E-10 |
| Lgr6 | 0 | 0 | 0,447361 | 0,629958 | 4,42938 | 6,51386 | 5,88675 | 17,7931 | 23,6263 | 24,0475 | Lgr6 | 4,431923918 | 2,19E-10 |
| Pygb | 39,1952 | 65,5623 | 113,544 | 79,1058 | 257,381 | 293,391 | 299,798 | 458,703 | 479,863 | 436,923 | Pygb | 2,117599023 | 2,26E-10 |
| Msrb2 | 3,44166 | 5,55548 | 10,4446 | 8,82138 | 26,2195 | 64,8326 | 59,5841 | 103,616 | 108,342 | 108,134 | Msrb2 | 3,063643348 | 2,28E-10 |
| Ptpn3 | 0,741725 | 1,36587 | 0,949354 | 0,448414 | 3,20655 | 7,52532 | 7,2469 | 19,7632 | 19,5773 | 17,4777 | Ptpn3 | 3,031128877 | 2,52E-10 |
| Iigp1 | 0 | 0 | 0,279798 | 0,112568 | 2,28045 | 7,46708 | 5,44821 | 12,4942 | 20,8211 | 23,2676 | Iigp1 | 5,284783899 | 2,76E-10 |
| Fam134b | 0,598007 | 2,09331 | 4,44115 | 10,6811 | 145,291 | 76,0248 | 74,5138 | 154,81 | 144,63 | 132,396 | Fam134b | 4,405598454 | 2,84E-10 |
| Tmem181b-ps | 4,26269 | 4,87301 | 6,96279 | 4,28242 | 7,83984 | 11,5062 | 10,3785 | 5,38172 | 5,74916 | 5,5713 | Tmem181b-ps | 2,185168232 | 2,91E-10 |
| Ttn | 17,4997 | 25,3098 | 27,3764 | 25,5073 | 314,707 | 93,225 | 88,5848 | 78,3367 | 83,9598 | 110,916 | Ttn | 2,839538778 | 2,95E-10 |
| Eif4e3 | 1,32525 | 2,87615 | 4,09278 | 4,80613 | 16,7787 | 25,0443 | 25,6548 | 38,3534 | 44,5257 | 42,8149 | Eif4e3 | 2,876994591 | 3,15E-10 |
| Mybpc2 | 0 | 0 | 0 | 0,0899677 | 2,25199 | 3,17765 | 3,09306 | 6,66031 | 6,24603 | 13,2049 | Mybpc2 | 4,891656645 | 4,04E-10 |
| Dixdc1 | 2,82087 | 1,90976 | 3,41657 | 2,17348 | 11,9047 | 7,78202 | 7,52255 | 2,36642 | 2,71523 | 2,68111 | Dixdc1 | 2,092849477 | 4,32E-10 |
| Me3 | 0,0732546 | 0,433594 | 0,183441 | 0,885677 | 3,86501 | 8,94805 | 9,60737 | 37,0826 | 43,0715 | 39,8762 | Me3 | 4,256320677 | 4,38E-10 |
| Tns1 | 1,97812 | 10,0669 | 25,214 | 17,6814 | 250,629 | 147,491 | 137,479 | 199,596 | 196,912 | 233,816 | Tns1 | 3,816628509 | 4,50E-10 |
| Cirbp | 32,5216 | 50,8671 | 53,5494 | 68,448 | 260,523 | 143,029 | 143,921 | 58,0852 | 69,9337 | 68,9831 | Cirbp | 2,049598181 | 4,61E-10 |
| Sucla2 | 27,5175 | 57,4658 | 97,1089 | 58,1932 | 189,681 | 358,628 | 373,359 | 603,492 | 661,474 | 638,878 | Sucla2 | 2,6246123 | 5,12E-10 |
| Fam73b | 6,78108 | 13,004 | 11,5641 | 13,9554 | 25,8229 | 43,4299 | 42,6545 | 36,3511 | 44,184 | 37,5725 | Fam73b | 2,067172595 | 5,28E-10 |
| Hdac11 | 2,40536 | 5,66849 | 9,59234 | 7,53859 | 23,7018 | 31,6865 | 27,3454 | 25,3163 | 21,7962 | 17,8448 | Hdac11 | 2,45419859 | 5,57E-10 |
| Lrrc14b | 0 | 0 | 3,74911 | 3,34637 | 17,1491 | 38,2548 | 38,0388 | 66,4321 | 79,1026 | 73,6886 | Lrrc14b | 3,40180669 | 5,62E-10 |
| Fndc5 | 1,34686 | 5,07308 | 12,3663 | 4,68728 | 34,0539 | 83,5702 | 86,3046 | 218,338 | 240,88 | 230,577 | Fndc5 | 3,704563167 | 5,65E-10 |
| 4631405J19Ri | 0 | 0 | 0 | 0 | 3,24375 | 12,9822 | 13,2152 | 13,5864 | 10,6419 | 12,0453 | 4631405J19Ri | 6,824655364 | 5,97E-10 |
| Trap1 | 29,2585 | 55,1284 | 54,3371 | 54,9683 | 105,285 | 185,075 | 181,954 | 158,932 | 167,656 | 168,394 | Trap1 | 2,001349404 | 6,22E-10 |
| Atcayos | 0 | 49,5006 | 20,9384 | 5,59888 | 91,7237 | 94,4474 | 109,759 | 9,66876 | 30,5691 | 34,1609 | Atcayos | 3,474077493 | 6,22E-10 |
| Idh3a | 25,8786 | 72,2632 | 94,5299 | 82,9734 | 244,43 | 388,372 | 383,635 | 424,084 | 483,224 | 459,517 | Idh3a | 2,542173043 | 6,39E-10 |
| Rabgap1l | 6,11407 | 9,08776 | 17,3874 | 14,3747 | 45,3298 | 75,329 | 74,0178 | 23,408 | 33,676 | 27,676 | Rabgap1l | 2,36610737 | 6,46E-10 |
| Nrtn | 2,26665 | 7,45297 | 5,35935 | 2,28201 | 20,0044 | 31,2189 | 30,2822 | 37,8854 | 32,8407 | 30,6519 | Nrtn | 2,930799806 | 6,98E-10 |
| Vwa8 | 4,14489 | 6,9759 | 16,6048 | 10,5237 | 37,2802 | 104,04 | 99,3598 | 111,211 | 106,367 | 112,04 | Vwa8 | 3,237041444 | 7,40E-10 |
| Tmem143 | 6,54737 | 7,22777 | 7,93703 | 7,8516 | 18,9705 | 54,2106 | 53,3982 | 115,381 | 119,089 | 126,577 | Tmem143 | 2,658633854 | 7,80E-10 |
| Angptl4 | 0,360994 | 0,534168 | 5,72473 | 1,45447 | 21,8202 | 65,2618 | 61,5805 | 28,3418 | 1,64047 | 4,21976 | Angptl4 | 4,583323606 | 8,14E-10 |
| Slc48a1 | 0 | 0 | 13,1542 | 9,95758 | 32,6441 | 45,899 | 45,7883 | 33,0586 | 38,2587 | 43,9429 | Slc48a1 | 2,648870283 | 8,30E-10 |
| Tmem38a | 6,63991 | 10,5441 | 30,8192 | 25,9382 | 90,8152 | 149,583 | 146,059 | 308,373 | 317,934 | 302,755 | Tmem38a | 3,081613506 | 8,63E-10 |
| Plekhf1 | 2,26794 | 3,35586 | 0,567855 | 3,19782 | 21,4478 | 13,8661 | 13,3843 | 11,1037 | 22,2815 | 20,1395 | Plekhf1 | 2,992752765 | 8,82E-10 |
| 2810013P06R | 2,18443 | 3,64674 | 4,34772 | 2,70814 | 10,4407 | 12,3661 | 11,2977 | 7,93295 | 8,76022 | 7,52577 | 2810013P06R | 2,044733918 | 9,12E-10 |
| Fgf13 | 6,39508 | 15,7721 | 14,2925 | 20,7669 | 47,5441 | 70,5928 | 70,4215 | 50,9095 | 61,7985 | 50,9866 | Fgf13 | 2,410186218 | 1,01E-09 |

| Gene | d9-11 (1) | d9-11 (2) | d9-11(3) | d9-11(4) | newborn(1) | newborn(2) | newborn(3) | adult(1) | adult(2) | adult(3) | Gene | log2(fold) | FDR(*p* -value) |
|---|---|---|---|---|---|---|---|---|---|---|---|---|---|
| Ppargc1a | 2,38612 | 6,34117 | 11,5603 | 8,14496 | 48,9109 | 32,2959 | 33,7962 | 39,9087 | 40,5133 | 28,9045 | Ppargc1a | 2,670445374 | 1,02E-09 |
| Tnni3 | 52,1871 | 182,646 | 532,044 | 374,238 | 2352,54 | 3111,87 | 3334,57 | 6173,26 | 5204,14 | 4813,36 | Tnni3 | 3,570294938 | 1,07E-09 |
| Ntn1 | 2,47953 | 5,59878 | 16,6499 | 21,9297 | 146,167 | 130,648 | 128,493 | 36,2142 | 25,4662 | 30,5016 | Ntn1 | 3,640196629 | 1,08E-09 |
| 2310015A10R | 0,197204 | 2,13996 | 1,31689 | 1,78824 | 8,75101 | 12,626 | 12,0033 | 5,1448 | 5,88699 | 5,80151 | 2310015A10R | 3,174951174 | 1,09E-09 |
| Tex2 | 3,8631 | 5,62116 | 6,55674 | 9,08199 | 17,2699 | 22,9751 | 20,6463 | 17,872 | 18,0498 | 17,7748 | Tex2 | 2,000773885 | 1,11E-09 |
| Ppif | 19,2227 | 26,4121 | 54,8152 | 50,4744 | 115,836 | 192,012 | 202,374 | 283,128 | 282,564 | 276,63 | Ppif | 2,463835625 | 1,16E-09 |
| Tmem53 | 1,7303 | 2,77261 | 6,48696 | 7,80093 | 20,9471 | 33,5672 | 28,6983 | 22,1572 | 22,7163 | 23,3235 | Tmem53 | 3,003463875 | 1,20E-09 |
| Synj2 | 2,97578 | 4,12431 | 3,76519 | 1,42571 | 7,88309 | 10,4929 | 11,0201 | 24,0435 | 19,2682 | 23,5485 | Synj2 | 2,267402986 | 1,42E-09 |
| Dtx3l | 0,207615 | 1,14869 | 1,09957 | 0,679228 | 4,30374 | 5,39512 | 4,88042 | 4,76564 | 4,9157 | 4,81121 | Dtx3l | 2,815886116 | 1,47E-09 |
| Rbl2 | 1,85958 | 2,90348 | 6,47863 | 4,92766 | 22,4557 | 17,9671 | 15,022 | 15,9316 | 20,2691 | 19,059 | Rbl2 | 2,41333165 | 1,55E-09 |
| Ptgr2 | 8,43101 | 18,6096 | 25,9502 | 22,9414 | 52,7704 | 138,914 | 137,874 | 159,255 | 168,103 | 168,68 | Ptgr2 | 2,797397042 | 1,55E-09 |
| Itga7 | 6,07985 | 5,761 | 4,74106 | 3,46577 | 12,6581 | 23,4388 | 22,3838 | 99,5989 | 118,988 | 113,117 | Itga7 | 2,239661901 | 1,59E-09 |
| Ank1 | 1,75005 | 4,30804 | 5,96085 | 2,71624 | 12,4456 | 29,9392 | 29,3058 | 61,2355 | 67,9624 | 57,7707 | Ank1 | 2,785523593 | 1,61E-09 |
| Ncam1 | 6,93947 | 14,4846 | 22,544 | 14,2294 | 104,119 | 65,6832 | 64,6225 | 1,18623 | 2,52617 | 1,45372 | Ncam1 | 2,254568969 | 1,62E-09 |
| Parp14 | 0,0864944 | 0,0426637 | 0,0722003 | 0,0871524 | 1,84902 | 0,974074 | 1,04628 | 1,10261 | 1,52971 | 1,28516 | Parp14 | 4,025055075 | 1,62E-09 |
| Gmpr | 21,3984 | 41,7684 | 63,0778 | 57,776 | 136,368 | 190,828 | 202,942 | 140,757 | 141,883 | 147,84 | Gmpr | 2,270706527 | 1,69E-09 |
| Wdfy1 | 1,01782 | 3,25872 | 3,10543 | 2,93505 | 11,2188 | 11,4936 | 10,7798 | 7,75024 | 8,65214 | 8,81425 | Wdfy1 | 2,789768882 | 1,74E-09 |
| Ap5s1 | 5,44495 | 5,37196 | 13,1865 | 12,5306 | 32,9386 | 43,529 | 44,4572 | 29,3331 | 36,3006 | 31,7219 | Ap5s1 | 2,361811 | 1,78E-09 |
| Selenbp1 | 4,56097 | 8,58949 | 16,2765 | 21,4029 | 66,4493 | 74,8171 | 70,2432 | 82,5258 | 98,4552 | 85,5202 | Selenbp1 | 2,664569725 | 1,91E-09 |
| Kcnb1 | 0,0556305 | 0,19208 | 0,905533 | 0,700698 | 4,30693 | 7,82766 | 7,63847 | 12,3895 | 13,399 | 16,5712 | Kcnb1 | 3,925996955 | 1,95E-09 |
| Il13ra1 | 1,40364 | 4,67331 | 6,07783 | 4,24249 | 15,2625 | 26,2185 | 25,2764 | 16,8771 | 21,3716 | 18,1945 | Il13ra1 | 2,682428373 | 2,03E-09 |
| Coro6 | 6,4887 | 20,0245 | 40,8912 | 21,8299 | 88,0548 | 137,52 | 134,909 | 126,281 | 129,5 | 138,912 | Coro6 | 3,004551463 | 2,03E-09 |
| Zfp46 | 3,23495 | 3,46288 | 8,96264 | 5,33979 | 26,733 | 20,5476 | 19,6971 | 10,2094 | 12,1465 | 10,8302 | Zfp46 | 2,333144596 | 2,19E-09 |
| Cs | 103,208 | 163,584 | 258,618 | 198,694 | 437,303 | 764,765 | 782,56 | 957,73 | 1025,2 | 1037,42 | Cs | 2,139434868 | 2,25E-09 |
| Fam212b | 0,420877 | 1,17639 | 2,07653 | 1,2988 | 6,69743 | 6,10838 | 4,71878 | 5,8346 | 7,50542 | 9,24573 | Fam212b | 2,489711598 | 2,42E-09 |
| Thra | 13,4974 | 31,2162 | 41,016 | 43,3578 | 112,469 | 127,391 | 129,762 | 49,0379 | 61,4692 | 48,6879 | Thra | 2,207358986 | 3,01E-09 |
| Ppargc1b | 3,2733 | 3,49091 | 9,37825 | 9,80435 | 59,3555 | 31,8569 | 28,8253 | 13,2774 | 11,5462 | 17,8619 | Ppargc1b | 2,838741875 | 3,04E-09 |
| Cpt2 | 9,41531 | 10,2732 | 23,6951 | 25,436 | 52,9186 | 164,97 | 169,538 | 226,531 | 241,543 | 248,63 | Cpt2 | 3,099727224 | 3,10E-09 |
| Lrg1 | 0 | 0 | 0,447256 | 0 | 4,41654 | 8,03232 | 8,40772 | 69,6567 | 56,3266 | 66,7816 | Lrg1 | 4,771756901 | 3,21E-09 |
| Lyrm9 | 0,361715 | 0,992925 | 2,20542 | 1,01412 | 6,39644 | 10,3019 | 10,3685 | 11,6225 | 12,7157 | 12,9836 | Lyrm9 | 3,156700544 | 3,26E-09 |
| Mir208b | 0 | 0 | 0 | 0 | 3620,63 | 0 | 0 | 0 | 0 | 0 | Mir208b | 6,461675427 | 3,58E-09 |
| Ndufa3 | 211,716 | 240,883 | 357,414 | 345,561 | 629,663 | 1190,72 | 1098,69 | 1530,97 | 1561,41 | 1499,65 | Ndufa3 | 2,016173208 | 3,61E-09 |
| Anxa4 | 2,21502 | 5,04246 | 4,69303 | 3,60419 | 11,7959 | 16,3278 | 18,8848 | 10,8279 | 10,3172 | 14,0747 | Anxa4 | 2,253508892 | 3,63E-09 |
| Pirt | 0,435119 | 1,28772 | 2,99633 | 1,86302 | 7,58496 | 14,7422 | 13,3427 | 1,18677 | 1,9683 | 3,09975 | Pirt | 3,113947286 | 3,66E-09 |
| Mapk10 | 0,08712 | 0,0429722 | 0,690864 | 0,395022 | 3,97417 | 4,9284 | 4,66177 | 3,88562 | 5,95582 | 6,6482 | Mapk10 | 3,828866445 | 3,88E-09 |
| Gja3 | 0 | 0,489855 | 3,10856 | 0,625303 | 14,7296 | 33,4919 | 33,3571 | 15,1183 | 18,1958 | 15,4051 | Gja3 | 4,611894375 | 3,92E-09 |
| Tcea3 | 9,75313 | 18,0758 | 31,5719 | 13,6902 | 53,61 | 105,942 | 106,222 | 117,179 | 147,829 | 112,583 | Tcea3 | 2,563522799 | 3,97E-09 |
| Mrm1 | 4,0391 | 2,52353 | 4,83226 | 6,23899 | 14,7177 | 21,0812 | 18,5881 | 12,2332 | 15,9516 | 15,3614 | Mrm1 | 2,281259823 | 4,12E-09 |
| Chmp4c | 0 | 1,33597 | 0,322955 | 0,19487 | 6,09548 | 7,34318 | 6,23988 | 9,83065 | 8,7992 | 9,49056 | Chmp4c | 3,942257659 | 4,50E-09 |
| Nol3 | 1,36096 | 1,26779 | 3,14053 | 2,80121 | 6,97812 | 15,3657 | 18,8515 | 27,116 | 28,1129 | 27,7395 | Nol3 | 3,166739868 | 4,53E-09 |
| Mrpl41 | 2,85395 | 6,66741 | 8,5196 | 6,45919 | 15,3401 | 26,4295 | 23,5306 | 33,0839 | 32,6218 | 33,3235 | Mrpl41 | 2,036343357 | 4,60E-09 |
| Fmo5 | 0,122316 | 0,414516 | 0,204244 | 0,123372 | 1,52221 | 1,99466 | 2,1344 | 3,92848 | 4,53782 | 4,23483 | Fmo5 | 3,714327806 | 4,71E-09 |
| D830031N03F | 2,29256 | 2,92899 | 3,54505 | 2,3479 | 29,9314 | 9,3005 | 8,23984 | 1,26635 | 1,36649 | 1,23797 | D830031N03F | 2,687943932 | 4,76E-09 |
| Atp5g1 | 112,158 | 193,666 | 289,828 | 280,496 | 570,403 | 843,025 | 841,449 | 1420,65 | 1089,07 | 1118,45 | Atp5g1 | 2,202258193 | 4,88E-09 |
| Cbx6 | 3,28846 | 5,34976 | 10,673 | 7,12002 | 31,5926 | 24,3258 | 22,0152 | 5,50424 | 7,41873 | 7,02815 | Cbx6 | 2,162044132 | 4,96E-09 |
| Actn2 | 110,766 | 236,143 | 449,553 | 389,686 | 1292,11 | 1439,25 | 1544,85 | 1239,54 | 1061,3 | 1245,16 | Actn2 | 2,535230709 | 4,97E-09 |
| Slc2a4 | 2,6864 | 6,33721 | 16,1836 | 16,3557 | 69,8803 | 77,5385 | 76,372 | 283,685 | 309,455 | 299,335 | Slc2a4 | 3,029213217 | 5,33E-09 |
| Fgl2 | 0,685219 | 3,21084 | 6,43454 | 4,14215 | 19,8639 | 37,8157 | 36,8718 | 12,2324 | 20,677 | 22,8616 | Fgl2 | 3,313654743 | 5,35E-09 |
| Eda | 0,323828 | 0,38335 | 1,13533 | 0,717816 | 2,87994 | 3,77232 | 3,33302 | 0,176826 | 0,302863 | 0,188434 | Eda | 2,660396999 | 6,21E-09 |
| Dio2 | 0,0543427 | 0,107219 | 0,181446 | 0 | 1,15422 | 1,6189 | 1,51476 | 0,148407 | 0,0181465 | 0,0702419 | Dio2 | 4,083186815 | 6,27E-09 |
| Fam195a | 16,2723 | 46,1468 | 61,525 | 57,8324 | 176,563 | 176,544 | 172,543 | 245,507 | 202,574 | 222,224 | Fam195a | 2,282942029 | 6,29E-09 |
| 2010111I01Ri | 0,970403 | 4,95579 | 5,43729 | 7,52756 | 37,5007 | 23,4632 | 23,295 | 33,658 | 37,4592 | 39,087 | 2010111I01Ri | 2,477011256 | 7,39E-09 |
| Htra1 | 1,00211 | 6,75514 | 12,128 | 4,5424 | 41,6993 | 62,4618 | 63,3758 | 77,7907 | 84,9609 | 82,8223 | Htra1 | 3,363938819 | 7,48E-09 |
| Adamts19 | 1,91464 | 1,88879 | 4,85165 | 3,16921 | 8,42563 | 14,1797 | 13,5974 | 0,124454 | 0,0457084 | 0,0442167 | Adamts19 | 2,337045268 | 8,06E-09 |
| Osbp2 | 0 | 0,154703 | 0,850851 | 0,631997 | 4,86487 | 6,59022 | 6,0554 | 15,4089 | 16,3108 | 13,9865 | Osbp2 | 3,831775849 | 8,69E-09 |
| Tef | 4,75947 | 7,00556 | 10,7203 | 11,189 | 51,7431 | 26,1911 | 23,6945 | 46,2225 | 40,7143 | 68,8521 | Tef | 2,253566609 | 8,78E-09 |
| Asb14 | 0 | 0,333919 | 0 | 0 | 2,64751 | 14,1497 | 15,5791 | 68,0952 | 71,0496 | 61,1389 | Asb14 | 5,371948611 | 8,81E-09 |
| Ccbl1 | 5,52654 | 5,80353 | 13,0941 | 10,9539 | 24,1055 | 52,0398 | 50,0354 | 25,8908 | 27,2408 | 28,9407 | Ccbl1 | 2,536595331 | 9,06E-09 |
| Hist2h2be | 0,887091 | 2,00024 | 2,53865 | 2,80863 | 7,40175 | 14,211 | 13,6837 | 5,87349 | 9,18269 | 8,70561 | Hist2h2be | 2,748364443 | 9,40E-09 |
| Pyurf | 3,33915 | 3,00513 | 3,56968 | 3,5415 | 7,18396 | 16,8232 | 16,0434 | 14,8236 | 15,3211 | 17,9298 | Pyurf | 2,250717925 | 9,46E-09 |
| Txnrd3 | 2,67067 | 8,13701 | 7,5705 | 5,38317 | 18,6403 | 23,1921 | 23,3731 | 12,8836 | 14,5271 | 13,923 | Txnrd3 | 2,07743863 | 9,49E-09 |
| Rnmtl1 | 3,10336 | 4,23878 | 7,17252 | 6,73175 | 17,7656 | 17,6646 | 20,2565 | 24,6669 | 25,2455 | 25,4032 | Rnmtl1 | 2,043391189 | 9,49E-09 |
| Dsp | 26,4444 | 66,5475 | 103,826 | 85,1046 | 271,18 | 290,584 | 287 | 266,719 | 308,661 | 281,096 | Dsp | 2,290679606 | 9,74E-09 |
| Ank2 | 4,05395 | 6,97165 | 14,8849 | 11,9228 | 30,16 | 49,8938 | 46,8736 | 52,8865 | 54,1504 | 57,7085 | Ank2 | 2,565511864 | 9,98E-09 |
| Urgcp | 1,71694 | 2,48489 | 7,8267 | 4,63014 | 25,2155 | 20,7653 | 20,7059 | 19,0587 | 24,5811 | 22,7882 | Urgcp | 2,579495542 | 1,04E-08 |
| Mut | 7,0385 | 13,4529 | 10,4283 | 10,0164 | 19,9816 | 41,2717 | 40,3976 | 75,7061 | 85,4554 | 78,5689 | Mut | 2,042222697 | 1,14E-08 |
| Camk2d | 13,0883 | 22,6619 | 42,2879 | 36,5156 | 95,4699 | 110,826 | 112,671 | 77,3075 | 85,9275 | 89,1784 | Camk2d | 2,15632371 | 1,17E-08 |
| Sdpr | 1,50091 | 11,3163 | 14,5862 | 6,58838 | 93,8953 | 55,238 | 54,7345 | 28,4619 | 39,8889 | 33,4289 | Sdpr | 3,198512152 | 1,20E-08 |
| Myh7b | 0 | 4,92166 | 14,21 | 11,9192 | 31,2694 | 60,2427 | 63,3196 | 20,9486 | 15,6913 | 17,7572 | Myh7b | 2,95077923 | 1,20E-08 |
| Slc37a1 | 2,71822 | 1,62806 | 1,21549 | 0,782439 | 7,12548 | 10,4844 | 9,9359 | 8,4564 | 7,28743 | 10,3136 | Slc37a1 | 2,692972357 | 1,42E-08 |
| Nwd1 | 0 | 0 | 0,101503 | 0,0408414 | 0,826278 | 1,12587 | 0,844027 | 0 | 0,0270463 | 0,0392731 | Nwd1 | 4,544922739 | 1,49E-08 |
| Ppcs | 3,73315 | 4,17362 | 6,23136 | 5,2635 | 12,7268 | 20,3014 | 18,6737 | 17,5913 | 11,6959 | 15,1512 | Ppcs | 2,182553161 | 1,53E-08 |
| Akap6 | 0,272517 | 1,10524 | 4,87826 | 3,69183 | 30,8238 | 36,7953 | 34,3936 | 36,2396 | 44,8439 | 38,013 | Akap6 | 3,825328124 | 1,60E-08 |
| Cpped1 | 5,10019 | 9,69442 | 17,6512 | 14,4114 | 37,1375 | 65,2387 | 67,8049 | 45,1082 | 50,3329 | 50,0586 | Cpped1 | 2,495301616 | 1,66E-08 |
| Phyhd1 | 0,562149 | 0,883216 | 3,23798 | 5,668 | 12,6469 | 31,573 | 29,818 | 86,7898 | 97,035 | 100,932 | Phyhd1 | 3,194206087 | 1,69E-08 |
| Pak6 | 0 | 0,765619 | 2,21313 | 2,13541 | 16,1769 | 18,3318 | 17,8378 | 9,39124 | 9,80792 | 10,6388 | Pak6 | 3,930874387 | 1,72E-08 |
| Cyp2u1 | 0,143939 | 0,283987 | 2,16255 | 1,30498 | 9,1442 | 12,2112 | 13,6041 | 11,3757 | 14,4535 | 12,6014 | Cyp2u1 | 3,649588973 | 1,78E-08 |
| Cox14 | 17,4614 | 46,9198 | 54,7598 | 61,1929 | 135,058 | 255,591 | 232,629 | 343,921 | 302,111 | 345,544 | Cox14 | 2,474500398 | 1,79E-08 |
| Mtus2 | 5,27346 | 9,91305 | 17,8158 | 11,966 | 31,8313 | 46,8534 | 47,1436 | 22,8376 | 35,4259 | 26,7424 | Mtus2 | 2,208440548 | 1,79E-08 |
| Cerk | 1,68662 | 3,3277 | 7,03928 | 5,94768 | 17,8465 | 20,1356 | 21,9422 | 12,8538 | 18,3227 | 14,8835 | Cerk | 2,383993457 | 1,82E-08 |
| Fbxo31 | 5,8081 | 5,72969 | 6,0755 | 5,33333 | 34,8853 | 14,1983 | 14,2832 | 59,1031 | 49,3693 | 66,2489 | Fbxo31 | 2,119175319 | 1,93E-08 |
| Acbd4 | 0,491076 | 1,29183 | 3,00579 | 1,484 | 7,01236 | 8,22191 | 8,34159 | 5,79661 | 8,69669 | 8,33937 | Acbd4 | 2,652733028 | 1,93E-08 |
| Zfp810 | 1,00712 | 1,68897 | 2,52192 | 1,31902 | 4,95214 | 6,63977 | 6,02114 | 1,92318 | 2,56343 | 2,60815 | Zfp810 | 2,166431764 | 1,98E-08 |
| Smpx | 27,2826 | 56,3235 | 122,877 | 100,113 | 328,44 | 439,788 | 427,167 | 317,555 | 354,687 | 336,843 | Smpx | 2,631319082 | 2,07E-08 |
| Ndufa5 | 99,0765 | 253,389 | 166,093 | 118,102 | 391,233 | 634,116 | 648,081 | 1042,49 | 945,518 | 917,975 | Ndufa5 | 2,099820388 | 2,16E-08 |
| Gm19277 | 0 | 0 | 0 | 0,323358 | 2,28353 | 10,0468 | 8,10944 | 2,90521 | 5,89028 | 5,0322 | Gm19277 | 5,753507429 | 2,18E-08 |
| Entpd5 | 3,11069 | 6,89693 | 10,5229 | 7,4837 | 17,2427 | 47,8757 | 47,8597 | 50,3369 | 71,1955 | 53,5226 | Entpd5 | 2,712985162 | 2,37E-08 |
| Fam210a | 2,45004 | 5,2474 | 8,98795 | 6,78904 | 18,371 | 24,3486 | 24,498 | 37,5825 | 46,1169 | 40,4163 | Fam210a | 2,189436328 | 2,41E-08 |
| Cep83os | 0,232948 | 0,919212 | 1,36112 | 0,782334 | 3,93769 | 4,9019 | 4,42603 | 0,777115 | 2,05143 | 2,46119 | Cep83os | 2,640615119 | 2,56E-08 |
| Wdr72 | 0 | 0 | 0,101555 | 0,0612911 | 0,671925 | 2,31517 | 2,75806 | 0,221468 | 0,406429 | 0,235944 | Wdr72 | 5,040331947 | 2,65E-08 |
| Sned1 | 0,142929 | 0,352499 | 1,13341 | 0,864039 | 3,13783 | 6,80178 | 5,14599 | 2,79626 | 2,36509 | 2,33395 | Sned1 | 3,429814871 | 2,65E-08 |
| Cox17 | 89,7701 | 106,402 | 106,146 | 137,617 | 225,873 | 476,341 | 450,046 | 638,135 | 593,77 | 606,042 | Cox17 | 2,033282991 | 2,86E-08 |
| Ndufs8 | 40,2939 | 77,1882 | 110,809 | 98,7617 | 210,559 | 338,103 | 339,98 | 487,557 | 492,673 | 472,518 | Ndufs8 | 2,081884644 | 2,90E-08 |
| Lrrc39 | 13,2256 | 14,8517 | 15,3759 | 18,8199 | 35,3005 | 91,6962 | 88,5303 | 62,413 | 78,4178 | 74,1052 | Lrrc39 | 2,418163454 | 3,24E-08 |
| Kat2b | 1,56699 | 6,02641 | 7,32246 | 4,65837 | 24,819 | 37,4534 | 35,941 | 31,2449 | 39,0096 | 36,6508 | Kat2b | 2,855057271 | 3,27E-08 |
| Eef1a2 | 2,16292 | 13,4586 | 45,2711 | 32,6806 | 206,657 | 361,942 | 378,308 | 1167,02 | 1020,03 | 1005,1 | Eef1a2 | 3,832494176 | 3,47E-08 |
| Kremen1 | 7,00548 | 12,0301 | 22,7408 | 22,4932 | 57,2862 | 63,6337 | 61,8403 | 61,9852 | 64,3957 | 69,9325 | Kremen1 | 2,202219356 | 3,77E-08 |
| Fbp2 | 0,847925 | 0 | 0 | 0 | 2,89229 | 18,591 | 18,5253 | 40,4362 | 47,8831 | 42,566 | Fbp2 | 5,369673527 | 3,82E-08 |
| Ybx2 | 1,12976 | 1,78315 | 3,01731 | 6,3719 | 11,871 | 25,4398 | 24,8283 | 12,7016 | 14,4933 | 17,1243 | Ybx2 | 3,285208611 | 3,83E-08 |
| Snx13 | 2,53126 | 3,99537 | 6,67687 | 6,17214 | 13,3324 | 17,6904 | 17,3821 | 18,9426 | 22,0222 | 21,2313 | Snx13 | 2,028043795 | 3,97E-08 |
| Dgke | 1,41106 | 2,82612 | 3,71417 | 1,41252 | 7,18158 | 7,93747 | 7,66525 | 4,65551 | 5,23312 | 5,85134 | Dgke | 2,039410727 | 4,01E-08 |
| Kcnip2 | 0,16991 | 0 | 0,850895 | 1,96822 | 32,6395 | 85,7455 | 85,1963 | 161,042 | 173,438 | 182,898 | Kcnip2 | 5,645819248 | 4,21E-08 |
| Slc12a7 | 8,35931 | 18,3866 | 25,7143 | 25,3923 | 50,316 | 83,4386 | 78,0779 | 97,5547 | 98,2008 | 97,4234 | Slc12a7 | 2,162191677 | 4,29E-08 |
| 4930481A15R | 0 | 0 | 0 | 0 | 0 | 0 | 0 | 0 | 0 | 0 | 4930481A15R | 2,46328487 | 4,51E-08 |
| Gm5105 | 0 | 0 | 0,701691 | 0,846774 | 2,745 | 3,69997 | 3,81035 | 3,62209 | 3,04805 | 3,27955 | Gm5105 | 3,497253598 | 4,99E-08 |
| Fitm1 | 7,01094 | 26,3662 | 38,0276 | 36,8308 | 82,3739 | 204,84 | 197,884 | 265,862 | 276,764 | 284,588 | Fitm1 | 2,784245898 | 5,05E-08 |
| Wif1 | 0 | 0 | 0,877963 | 0,908225 | 5,66868 | 9,3554 | 7,93214 | 2,18375 | 7,16888 | 2,97289 | Wif1 | 4,055150249 | 5,25E-08 |
| Hint3 | 6,99711 | 10,6665 | 7,96089 | 13,4404 | 25,4597 | 54,4226 | 41,7918 | 65,126 | 70,124 | 67,8887 | Hint3 | 2,319362746 | 5,67E-08 |
| Serac1 | 1,33647 | 1,69006 | 2,5775 | 2,88922 | 5,22932 | 8,26471 | 8,41068 | 3,45585 | 5,53066 | 3,99856 | Serac1 | 2,131785306 | 6,09E-08 |
| Antxr2 | 10,2446 | 22,3356 | 26,0849 | 25,5867 | 46,1384 | 104,95 | 102,82 | 67,0591 | 89,3442 | 71,8273 | Antxr2 | 2,23523559 | 7,29E-08 |
| Slc25a12 | 4,94549 | 21,76 | 37,6363 | 29,9502 | 111,583 | 136,952 | 133,06 | 167,794 | 178,284 | 182,993 | Slc25a12 | 2,685634916 | 7,60E-08 |
| 9430091E24R | 0 | 0 | 0 | 0 | 0 | 0 | 0 | 0 | 0 | 0 | 9430091E24R | 3,022402219 | 8,87E-08 |
| Ppm1e | 0,149659 | 1,18111 | 2,91493 | 1,20636 | 7,47292 | 12,5744 | 13,6444 | 3,86044 | 4,89617 | 5,82655 | Ppm1e | 3,196993928 | 8,93E-08 |

| Gene | d9-11 (1) | d9-11 (2) | d9-11(3) | d9-11(4) | newborn(1) | newborn(2) | newborn(3) | adult(1) | adult(2) | adult(3) | Gene | log2(fold) | FDR(*p* -value) |
|---|---|---|---|---|---|---|---|---|---|---|---|---|---|
| Il23a | 0 | 0 | 0 | 0 | 3,33279 | 5,84125 | 7,21537 | 0 | 0,180436 | 0,260497 | Il23a | 6,078641531 | 9,07E-08 |
| Zcchc5 | 0,334351 | 0,549723 | 0,372107 | 0 | 5,81284 | 2,65496 | 2,76837 | 0,709235 | 0,859079 | 0,360928 | Zcchc5 | 3,616268049 | 9,38E-08 |
| Rab20 | 0,603313 | 2,9755 | 13,5912 | 7,89191 | 49,6491 | 47,1184 | 43,9098 | 38,499 | 31,5152 | 32,8524 | Rab20 | 3,253509564 | 9,48E-08 |
| Ddx60 | 0 | 0,103986 | 0,131982 | 0 | 0,970136 | 1,10824 | 0,968599 | 0,239897 | 0,35195 | 0,25545 | Ddx60 | 4,13082712 | 9,56E-08 |
| Sorbs2os | 0 | 0 | 0 | 0 | 0 | 0 | 0 | 0 | 0 | 0 | Sorbs2os | 3,339646142 | 1,04E-07 |
| Lipe | 0,685623 | 2,09819 | 1,8569 | 3,81903 | 14,0693 | 26,8326 | 24,8665 | 26,9244 | 16,4746 | 19,0852 | Lipe | 3,332536232 | 1,04E-07 |
| Aamdc | 7,92247 | 12,6088 | 24,2292 | 9,54019 | 30,8064 | 43,1934 | 46,6935 | 67,7202 | 70,983 | 72,1501 | Aamdc | 2,006691519 | 1,06E-07 |
| Hrc | 25,162 | 41,5055 | 47,6276 | 47,3707 | 73,5961 | 168,331 | 183,155 | 697,026 | 737,659 | 653,364 | Hrc | 2,109826587 | 1,06E-07 |
| Emc9 | 0 | 0 | 0 | 0 | 22,1733 | 53,7449 | 53,3355 | 72,3095 | 65,2286 | 62,3689 | Emc9 | 2,93771181 | 1,09E-07 |
| Chrm2 | 0,334587 | 0,330063 | 4,46823 | 0,674058 | 37,6389 | 18,6142 | 17,6678 | 57,2618 | 64,5655 | 60,8671 | Chrm2 | 4,201103489 | 1,16E-07 |
| Tmem70 | 10,4666 | 22,2384 | 21,6713 | 17,6377 | 35,1115 | 82,4684 | 76,6251 | 108,191 | 114,623 | 114,37 | Tmem70 | 2,13731323 | 1,18E-07 |
| Lmod3 | 7,51339 | 10,8213 | 16,9321 | 20,8892 | 34,8219 | 73,5983 | 79,4136 | 75,3394 | 111,32 | 97,9348 | Lmod3 | 2,408993066 | 1,25E-07 |
| Ppm1j | 13,5666 | 9,98733 | 19,6041 | 12,2368 | 27,1314 | 60,3961 | 62,6746 | 23,8796 | 27,3024 | 27,5715 | Ppm1j | 2,068004091 | 1,26E-07 |
| 1700040L02Ri | 0 | 1,7647 | 6,22077 | 3,90307 | 22,6344 | 49,2139 | 43,9303 | 19,4311 | 28,2473 | 25,2525 | 1700040L02Ri | 3,595691409 | 1,29E-07 |
| Ahrr | 0,0673802 | 0,531765 | 0,112488 | 0 | 1,81315 | 2,86865 | 2,82831 | 1,04241 | 1,03586 | 0,522823 | Ahrr | 3,902436968 | 1,29E-07 |
| Tmco4 | 0,979864 | 1,15996 | 2,12652 | 2,6654 | 6,37297 | 7,39291 | 7,54015 | 6,95038 | 5,57786 | 3,96454 | Tmco4 | 2,276027822 | 1,31E-07 |
| Trim55 | 30,2391 | 49,5255 | 93,3505 | 72,4349 | 169,158 | 232,258 | 213,694 | 60,855 | 67,8523 | 65,3654 | Trim55 | 2,080552851 | 1,31E-07 |
| Rmdn1 | 0,55064 | 3,25916 | 5,20866 | 2,21855 | 9,37604 | 28,8036 | 27,9867 | 29,4849 | 29,1475 | 30,637 | Rmdn1 | 3,120088039 | 1,37E-07 |
| Rrm2b | 1,48748 | 1,88665 | 2,60151 | 2,21235 | 4,7152 | 10,946 | 10,331 | 9,02367 | 12,6438 | 11,5204 | Rrm2b | 2,245274778 | 1,40E-07 |
| Rdh14 | 2,42527 | 7,41654 | 13,1569 | 13,1896 | 35,6179 | 55,4931 | 53,2424 | 49,4566 | 56,0626 | 53,2029 | Rdh14 | 2,658838366 | 1,40E-07 |
| Ifngr2 | 8,21112 | 24,9483 | 36,9173 | 28,9489 | 67,6491 | 130,618 | 131,101 | 174,236 | 211,143 | 194,296 | Ifngr2 | 2,4107727 | 1,40E-07 |
| Mtl5 | 0 | 0,164941 | 0,125717 | 0,168422 | 1,61799 | 3,54791 | 2,95325 | 1,78131 | 1,31318 | 1,6147 | Mtl5 | 4,605274194 | 1,42E-07 |
| Asb8 | 9,83618 | 11,8595 | 28,9301 | 23,3858 | 55,1507 | 79,547 | 74,8458 | 76,3305 | 83,8987 | 83,1204 | Asb8 | 2,190900239 | 1,42E-07 |
| Pitpnc1 | 3,2241 | 5,38254 | 9,72998 | 4,59799 | 12,3907 | 30,0809 | 30,5056 | 17,8148 | 23,4603 | 17,548 | Pitpnc1 | 2,34355098 | 1,46E-07 |
| Pcdhgb4 | 0 | 0 | 0 | 0 | 0 | 0 | 0 | 0 | 0 | 0 | Pcdhgb4 | 2,085280366 | 1,50E-07 |
| Ankrd33b | 1,2163 | 0,618328 | 0,350101 | 1,17356 | 4,71649 | 3,72263 | 3,71088 | 8,3328 | 10,9726 | 9,42043 | Ankrd33b | 2,478837343 | 1,50E-07 |
| Tcaim | 1,07816 | 1,48903 | 2,96978 | 3,04132 | 6,17301 | 11,8403 | 12,4307 | 32,2498 | 37,335 | 41,6837 | Tcaim | 2,519133088 | 1,51E-07 |
| Endou | 0,139101 | 0 | 0 | 0 | 1,61733 | 2,61815 | 3,11566 | 0,252749 | 0,700501 | 0,631329 | Endou | 4,939483836 | 1,61E-07 |
| Ankrd23 | 0,35039 | 6,39455 | 4,09433 | 1,05882 | 16,5541 | 41,0643 | 39,0007 | 331,503 | 290,11 | 391,496 | Ankrd23 | 3,65169706 | 1,66E-07 |
| Pde7b | 0,311233 | 1,68867 | 1,3639 | 0,548755 | 3,69226 | 5,61366 | 4,95969 | 7,36228 | 11,0849 | 10,0411 | Pde7b | 2,612878976 | 1,75E-07 |
| Rpl3l | 0,370413 | 0 | 1,23646 | 0 | 4,5441 | 7,65168 | 11,4993 | 333,036 | 330,518 | 342,262 | Rpl3l | 4,43122923 | 1,79E-07 |
| Myl3 | 444,967 | 747,492 | 1697,57 | 1600,08 | 4020,85 | 4978,13 | 5010,1 | 7546,41 | 5902,97 | 6118,69 | Myl3 | 2,328187768 | 1,91E-07 |
| Oxld1 | 1,02733 | 5,57345 | 5,5719 | 3,10182 | 14,9494 | 25,9361 | 22,7084 | 42,5611 | 40,1029 | 48,8234 | Oxld1 | 2,742962992 | 1,93E-07 |
| Slc4a3 | 7,34635 | 10,3024 | 25,2801 | 16,6539 | 41,7638 | 56,061 | 57,9206 | 90,9647 | 96,2974 | 96,4999 | Slc4a3 | 2,20078197 | 1,94E-07 |
| Coq3 | 3,93675 | 6,38907 | 5,72388 | 5,75691 | 10,1177 | 27,7488 | 27,3468 | 33,3524 | 40,557 | 39,957 | Coq3 | 2,265073181 | 2,07E-07 |
| Clybl | 9,21576 | 19,648 | 26,5469 | 32,9292 | 74,1411 | 160,034 | 149,825 | 163,069 | 169,918 | 176,211 | Clybl | 2,423627818 | 2,11E-07 |
| Cav3 | 0,323873 | 7,0286 | 21,0823 | 21,1972 | 160,571 | 203,644 | 198,739 | 86,4284 | 105,407 | 109,76 | Cav3 | 3,900334493 | 2,22E-07 |
| Amigo1 | 1,59231 | 0,670754 | 1,79727 | 1,66647 | 3,97278 | 6,37856 | 5,87887 | 6,75652 | 8,31335 | 6,37038 | Amigo1 | 2,247705 | 2,25E-07 |
| Eppk1 | 0,175859 | 0,19827 | 0,272626 | 0,708818 | 6,61226 | 2,79266 | 2,54951 | 0,0915408 | 0,0586154 | 0,10542 | Eppk1 | 3,154046206 | 2,32E-07 |
| Pm20d1 | 0,732452 | 1,26448 | 0,84065 | 1,1069 | 2,8279 | 4,70375 | 4,35016 | 4,33018 | 8,2436 | 5,39226 | Pm20d1 | 2,261471122 | 2,38E-07 |
| 2010315B03R | 0,122738 | 0,874012 | 0,950804 | 0,510023 | 2,83476 | 3,73962 | 3,59723 | 4,60093 | 3,7815 | 3,99976 | 2010315B03R | 2,760534676 | 2,43E-07 |
| Spryd4 | 4,28038 | 8,00117 | 12,0342 | 7,82348 | 19,8328 | 38,4851 | 34,9678 | 32,2731 | 33,6906 | 36,9306 | Spryd4 | 2,219319734 | 2,45E-07 |
| Acot11 | 0,446467 | 1,26627 | 2,00316 | 1,34955 | 10,405 | 6,79724 | 7,02612 | 10,8715 | 9,48656 | 15,0142 | Acot11 | 2,481391258 | 2,61E-07 |
| Parp9 | 0,328356 | 0,269017 | 1,63862 | 0,549483 | 4,44883 | 6,10312 | 6,80491 | 3,43623 | 4,51496 | 5,05399 | Parp9 | 3,157394336 | 2,85E-07 |
| Abcd3 | 6,50179 | 12,9223 | 17,4783 | 21,2888 | 35,509 | 68,4121 | 68,6094 | 72,7816 | 80,7821 | 80,3044 | Abcd3 | 2,244455968 | 3,00E-07 |
| Lyrm5 | 3,38496 | 9,0165 | 16,4824 | 9,23796 | 27,8472 | 61,1835 | 61,5074 | 62,5113 | 72,7102 | 76,6358 | Lyrm5 | 2,610183265 | 3,03E-07 |
| Gm20300 | 1,13278 | 1,11748 | 2,36383 | 1,14125 | 3,39731 | 9,02599 | 9,54962 | 13,6504 | 18,2818 | 18,2693 | Gm20300 | 2,652816368 | 3,16E-07 |
| LOC10263651 | 8,15855 | 11,625 | 41,2199 | 10,941 | 22,6333 | 61,447 | 63,4552 | 35,2045 | 42,0669 | 42,6175 | LOC10263651 | 2,319876904 | 3,51E-07 |
| Gfm1 | 24,2805 | 45,6357 | 55,4661 | 65,6561 | 101,78 | 188,738 | 186,953 | 183,952 | 186,839 | 199,145 | Gfm1 | 2,010354267 | 3,57E-07 |
| Phyhip | 0 | 0,233401 | 0 | 1,43012 | 4,99695 | 13,0252 | 12,0458 | 10,3227 | 20,2682 | 18,2599 | Phyhip | 4,345701362 | 3,83E-07 |
| Trim72 | 1,88909 | 5,99003 | 7,32077 | 2,71862 | 9,75711 | 34,9765 | 35,6858 | 391,96 | 352,376 | 404,04 | Trim72 | 2,861478799 | 4,03E-07 |
| Dmd | 0,267708 | 0,924339 | 4,39491 | 1,93327 | 14,8138 | 21,6741 | 20,4053 | 18,1353 | 22,7619 | 25,6808 | Dmd | 3,364452711 | 4,06E-07 |
| Camta1 | 5,14489 | 18,1515 | 10,2422 | 9,72364 | 26,6054 | 35,2696 | 34,0591 | 42,4013 | 44,3154 | 44,2641 | Camta1 | 2,13909797 | 4,06E-07 |
| Parp10 | 0,494874 | 0,390552 | 0,330457 | 1,29628 | 4,30846 | 3,17451 | 3,69051 | 1,56716 | 2,45265 | 2,32935 | Parp10 | 2,823259704 | 4,18E-07 |
| Pnmal2 | 0,619298 | 0,840038 | 1,22773 | 1,4039 | 4,56164 | 3,37428 | 2,98911 | 1,76079 | 0,931961 | 1,90283 | Pnmal2 | 2,085232002 | 4,22E-07 |
| 3830406C13R | 0,812961 | 2,12877 | 1,57664 | 1,01163 | 3,40677 | 13,3278 | 13,3927 | 6,77843 | 9,0718 | 8,61229 | 3830406C13R | 3,12169268 | 4,27E-07 |
| B330016D10R | 0,94356 | 1,55134 | 1,57511 | 1,42571 | 5,88852 | 4,7779 | 4,68217 | 4,56999 | 3,43576 | 3,92833 | B330016D10R | 2,123505947 | 4,37E-07 |
| Lgals4 | 1,99264 | 0 | 0 | 0 | 17,6907 | 45,998 | 49,48 | 20,6798 | 27,1167 | 23,8403 | Lgals4 | 5,231096518 | 4,43E-07 |
| Parp12 | 0,909114 | 1,29543 | 4,46871 | 1,32297 | 10,5208 | 9,7996 | 9,92592 | 5,23562 | 8,72835 | 6,96503 | Parp12 | 2,573496513 | 4,48E-07 |
| Naprt1 | 1,4552 | 0,897201 | 0,910932 | 1,83221 | 4,74908 | 10,1926 | 9,71973 | 11,2264 | 14,8191 | 13,349 | Naprt1 | 2,791447081 | 4,49E-07 |
| Amy1 | 0 | 0 | 0 | 0,211986 | 4,09476 | 4,12856 | 4,6841 | 4,64531 | 6,27034 | 1,97113 | Amy1 | 5,100312539 | 4,59E-07 |
| Map2k3os | 0,579831 | 1,14395 | 0,483909 | 0,87583 | 5,19238 | 7,08861 | 5,97806 | 7,87401 | 7,1742 | 7,27893 | Map2k3os | 3,170763565 | 4,74E-07 |
| Acads | 18,042 | 36,3225 | 66,0736 | 56,3733 | 109,144 | 230,374 | 219,203 | 271,511 | 250,052 | 253,176 | Acads | 2,324046684 | 4,98E-07 |
| Adam22 | 0,309314 | 1,33949 | 4,12625 | 2,48623 | 6,71352 | 4,56595 | 4,26081 | 1,84673 | 1,97146 | 1,63132 | Adam22 | 2,554320975 | 5,03E-07 |
| Gpr155 | 0,448134 | 0,38575 | 1,06877 | 1,4446 | 3,51076 | 4,58681 | 4,96221 | 6,05621 | 8,32212 | 6,70875 | Gpr155 | 2,603493196 | 5,14E-07 |
| Ldhd | 0 | 0 | 0,380214 | 0,152957 | 1,76585 | 5,07004 | 3,35534 | 60,6805 | 63,6213 | 63,3352 | Ldhd | 4,433530289 | 5,25E-07 |
| Echdc3 | 0,1908 | 0,376439 | 2,07024 | 2,69055 | 7,90649 | 16,6885 | 17,9915 | 28,3942 | 31,3632 | 31,3607 | Echdc3 | 3,539500487 | 5,37E-07 |
| Hspb7 | 58,4935 | 123,979 | 289,807 | 235,445 | 637,119 | 766,263 | 835,264 | 1162,53 | 939,031 | 1181,32 | Hspb7 | 2,403252822 | 5,46E-07 |
| Kcna4 | 0 | 0,194584 | 0,548822 | 0,331227 | 2,21616 | 2,23102 | 2,29643 | 0,359017 | 0,856875 | 0,467622 | Kcna4 | 3,126637375 | 5,68E-07 |
| Tuba4a | 9,34947 | 8,21968 | 22,0388 | 24,3388 | 42,9059 | 109,365 | 114,735 | 570,043 | 606,384 | 493,361 | Tuba4a | 2,668258626 | 5,90E-07 |
| Klhl31 | 1,56133 | 5,81323 | 10,9309 | 7,15542 | 23,5756 | 36,0562 | 34,8478 | 44,1102 | 50,0892 | 50,8606 | Klhl31 | 2,560860571 | 6,24E-07 |
| Lyrm7 | 0,310032 | 1,37628 | 2,45833 | 0,624608 | 4,39718 | 11,9292 | 12,4656 | 17,3161 | 18,5853 | 18,1529 | Lyrm7 | 3,159017936 | 6,24E-07 |
| Fsd2 | 7,71949 | 21,649 | 39,673 | 38,5513 | 85,7271 | 153,82 | 152,769 | 143,167 | 162,698 | 141,901 | Fsd2 | 2,547244907 | 6,45E-07 |
| Lmcd1 | 0,805654 | 4,17246 | 7,90097 | 5,68026 | 23,6709 | 25,1674 | 26,9784 | 42,0258 | 42,3971 | 46,2744 | Lmcd1 | 2,659006155 | 6,46E-07 |
| Scin | 0,122689 | 0,484125 | 0,487217 | 0,123598 | 1,67519 | 4,19871 | 3,28899 | 2,12095 | 2,4572 | 2,09101 | Scin | 3,405317364 | 6,49E-07 |
| E330017L17Ri | 0 | 0 | 0 | 0 | 3,82686 | 9,95493 | 9,29037 | 8,05824 | 8,70601 | 6,69585 | E330017L17Ri | 5,210826627 | 6,69E-07 |
| Ttc30b | 0,444471 | 0,856052 | 1,97862 | 0,426438 | 4,29672 | 5,71861 | 5,79788 | 10,5116 | 10,97 | 10,1641 | Ttc30b | 2,991805083 | 7,09E-07 |
| Txlnb | 1,82248 | 9,48487 | 18,866 | 15,9765 | 53,7116 | 93,8943 | 90,8929 | 255,054 | 287,677 | 259,681 | Txlnb | 2,988968426 | 7,20E-07 |
| Etfa | 7,32251 | 38,2839 | 57,0395 | 42,035 | 181,699 | 407,655 | 408,32 | 540,417 | 564,471 | 531,484 | Etfa | 2,655625853 | 7,31E-07 |
| Gm12295 | 0,204745 | 0,504949 | 0,341801 | 1,34077 | 2,55324 | 3,28498 | 3,07602 | 0,651606 | 1,44015 | 2,07135 | Gm12295 | 2,730810879 | 7,39E-07 |
| Cox6a2 | 10,3409 | 53,7205 | 192,684 | 208,07 | 1017,62 | 1382,87 | 1426,47 | 3163,72 | 2412,99 | 2401,79 | Cox6a2 | 3,516420668 | 7,41E-07 |
| Mapk1ip1 | 1,6301 | 2,29719 | 7,19118 | 4,45613 | 17,839 | 20,1003 | 20,259 | 25,5436 | 16,8313 | 24,1692 | Mapk1ip1 | 2,463192666 | 7,42E-07 |
| Nudt6 | 4,2189 | 2,78291 | 5,00513 | 8,8495 | 16,365 | 30,2188 | 27,0839 | 42,2717 | 38,1896 | 44,4696 | Nudt6 | 2,40719277 | 7,64E-07 |
| 9330158H04R | 0 | 0,104 | 0 | 0 | 0 | 0 | 0 | 0 | 0 | 0 | 9330158H04R | 4,604602527 | 7,69E-07 |
| AI317395 | 0,205977 | 0,121916 | 0 | 0,249004 | 1,52417 | 2,99161 | 2,55249 | 3,59425 | 3,82036 | 3,40354 | AI317395 | 4,153092216 | 7,73E-07 |
| Efnb3 | 12,99 | 31,4357 | 40,8298 | 52,4501 | 89,8955 | 157,49 | 156,587 | 59,8904 | 77,4969 | 50,9878 | Efnb3 | 2,214057617 | 8,45E-07 |
| Thnsl2 | 0,506899 | 0,333363 | 0,987197 | 0,510596 | 3,59549 | 3,64989 | 4,04626 | 3,98842 | 4,48996 | 2,68714 | Thnsl2 | 2,871419021 | 8,48E-07 |
| March3 | 1,18803 | 0,781303 | 2,14839 | 1,39604 | 6,96946 | 6,42118 | 6,63 | 3,4132 | 4,20374 | 4,11747 | 03. Mrz | 2,288317748 | 8,54E-07 |
| Tmem88b | 1,02848 | 0,461179 | 0,546305 | 0,565188 | 1,65153 | 5,85047 | 6,06997 | 0,51021 | 0,319252 | 0,459929 | Tmem88b | 3,167139267 | 8,60E-07 |
| Cbr4 | 4,40745 | 9,69868 | 9,90277 | 12,9719 | 21,3177 | 45,359 | 50,0291 | 56,7183 | 62,1421 | 61,1376 | Cbr4 | 2,308179911 | 8,71E-07 |
| Acyp2 | 6,04046 | 15,8885 | 2,79984 | 7,42771 | 32,8289 | 80,915 | 75,6574 | 227,665 | 247,647 | 274,453 | Acyp2 | 2,893605196 | 8,82E-07 |
| Arhgap24 | 1,44488 | 1,54413 | 3,83433 | 3,14632 | 9,07305 | 10,1474 | 8,82639 | 11,1627 | 8,68079 | 8,65946 | Arhgap24 | 2,07657199 | 9,77E-07 |
| Dpyd | 0,144183 | 0,213354 | 0,541585 | 0,217904 | 1,36826 | 3,37661 | 3,75533 | 5,83827 | 7,10156 | 5,86287 | Dpyd | 3,258830455 | 9,79E-07 |
| Ccng1 | 7,16414 | 21,0186 | 38,5978 | 23,3405 | 58,8868 | 130,57 | 136,275 | 359,761 | 428,385 | 416,841 | Ccng1 | 2,519068411 | 9,90E-07 |
| BC037032 | 0 | 0 | 0 | 0 | 0 | 0 | 0 | 0 | 0 | 0 | BC037032 | 3,504355648 | 1,02E-06 |
| Pitpnm3 | 0,0474398 | 0,279205 | 0,315001 | 0,0956992 | 0,901633 | 1,51121 | 1,37245 | 0,649038 | 1,99872 | 1,4934 | Pitpnm3 | 3,036868134 | 1,02E-06 |
| Fbxo40 | 0,132237 | 1,69585 | 4,30463 | 4,52926 | 30,3595 | 27,8707 | 24,2601 | 65,1256 | 68,6501 | 62,4877 | Fbxo40 | 3,393829984 | 1,04E-06 |
| Adssl1 | 18,5614 | 43,2962 | 60,8383 | 47,7128 | 84,1299 | 230,944 | 226,541 | 164,033 | 156,981 | 148,705 | Adssl1 | 2,336223776 | 1,11E-06 |
| Aqp4 | 0,125064 | 1,17207 | 1,7225 | 2,20516 | 6,44682 | 12,8709 | 11,9924 | 2,56107 | 5,89151 | 4,00218 | Aqp4 | 3,113673534 | 1,17E-06 |
| Pln | 1,56701 | 15,7394 | 91,318 | 54,3881 | 1124,72 | 1922,74 | 2047,84 | 3155,73 | 3295,76 | 3041,02 | Pln | 4,881162313 | 1,20E-06 |
| Scn5a | 1,16854 | 4,23428 | 11,3013 | 20,0101 | 77,5806 | 90,6873 | 86,223 | 82,2855 | 87,6913 | 99,9912 | Scn5a | 3,310178548 | 1,24E-06 |
| Syne3 | 0,748075 | 0,922472 | 0,312219 | 1,00497 | 5,21715 | 2,63837 | 2,2351 | 1,30627 | 1,02871 | 1,25075 | Syne3 | 2,554387466 | 1,28E-06 |
| Kcnj2 | 1,04681 | 4,3602 | 2,76703 | 2,46104 | 6,88466 | 10,913 | 10,1368 | 20,2214 | 21,163 | 16,9723 | Kcnj2 | 2,112662217 | 1,36E-06 |
| Rsph3b | 18,2966 | 18,1242 | 27,2593 | 22,8565 | 41,0436 | 54,8109 | 59,6867 | 39,3071 | 43,0368 | 41,293 | Rsph3b | 2,158566053 | 1,39E-06 |
| Zadh2 | 2,35243 | 6,67189 | 13,4996 | 12,9363 | 30,1946 | 53,7817 | 53,6395 | 49,7379 | 54,7464 | 56,1243 | Zadh2 | 2,583629985 | 1,59E-06 |
| Optn | 1,93987 | 4,64744 | 8,09579 | 5,8625 | 17,9377 | 19,1251 | 18,7489 | 44,3122 | 46,5409 | 36,1601 | Optn | 2,062647254 | 1,64E-06 |
| Ablim2 | 0 | 0,557527 | 1,49385 | 0,284694 | 3,18195 | 8,43327 | 7,95448 | 6,97369 | 9,53776 | 10,1597 | Ablim2 | 3,661188302 | 1,65E-06 |
| Cyb5d2 | 1,1275 | 0,834197 | 1,41164 | 0,567902 | 3,55725 | 4,79284 | 4,4366 | 13,2519 | 14,3939 | 13,9442 | Cyb5d2 | 2,275809362 | 1,65E-06 |
| Fam13a | 1,11958 | 0,589049 | 2,30519 | 1,05281 | 4,01742 | 5,40319 | 5,14618 | 10,1869 | 11,0305 | 9,07577 | Fam13a | 2,156302441 | 1,85E-06 |
| Ldhb | 5,67392 | 32,4629 | 101,099 | 81,4202 | 542,146 | 693,182 | 703,054 | 1001,76 | 958,224 | 852,033 | Ldhb | 2,948935696 | 1,88E-06 |
| Plbd1 | 0,1732 | 0,341715 | 0,578244 | 2,268 | 4,22641 | 12,3131 | 10,689 | 41,1939 | 33,4496 | 36,4322 | Plbd1 | 3,360121969 | 1,92E-06 |
| Mgst3 | 10,9348 | 40,2673 | 40,1411 | 44,7341 | 138,37 | 214,387 | 187,612 | 401,799 | 421,888 | 418,177 | Mgst3 | 2,44089248 | 2,08E-06 |
| Scx | 0,66256 | 2,61431 | 3,04113 | 1,66782 | 24,2202 | 10,4508 | 10,3061 | 7,49067 | 5,51731 | 6,0587 | Scx | 2,829573635 | 2,15E-06 |

| Gene | d9-11 (1) | d9-11 (2) | d9-11(3) | d9-11(4) | newborn(1) | newborn(2) | newborn(3) | adult(1) | adult(2) | adult(3) | Gene | log2(fold) | FDR($p$-value) |
|---|---|---|---|---|---|---|---|---|---|---|---|---|---|
| Col6a2 | 10,6553 | 6,49484 | 24,5852 | 13,7899 | 195,836 | 40,3315 | 40,8816 | 1,73358 | 2,46387 | 1,90589 | Col6a2 | 2,832213393 | 2,21E-06 |
| 5730409E04R | 0,461793 | 0,341665 | 1,15745 | 1,28057 | 3,30226 | 4,0186 | 4,2705 | 8,19336 | 9,31645 | 7,88244 | 5730409E04R | 2,247284556 | 2,35E-06 |
| Gm14446 | 0,304599 | 0 | 0,381356 | 0,306833 | 2,59196 | 3,15444 | 2,61758 | 2,21317 | 2,76341 | 2,14884 | Gm14446 | 3,461701359 | 2,38E-06 |
| Trib2 | 2,59829 | 7,01114 | 17,2869 | 14,0132 | 40,6904 | 53,6111 | 52,1065 | 14,5319 | 13,135 | 14,6439 | Trib2 | 2,477268835 | 2,44E-06 |
| Echdc2 | 6,85089 | 5,5097 | 7,20903 | 15,7488 | 19,2792 | 45,7422 | 46,1886 | 46,2789 | 41,909 | 37,5923 | Echdc2 | 2,334012959 | 2,65E-06 |
| Rapsn | 0 | 0,427983 | 5,06945 | 3,05877 | 15,2142 | 26,2998 | 26,7656 | 17,7051 | 20,5668 | 18,4802 | Rapsn | 3,382552724 | 2,70E-06 |
| Efcab2 | 1,70426 | 3,49178 | 3,06387 | 3,8299 | 5,12979 | 13,9463 | 12,0252 | 32,7595 | 47,8177 | 41,5241 | Efcab2 | 2,231199679 | 2,83E-06 |
| Smpdl3a | 2,51352 | 7,62928 | 13,0714 | 5,84239 | 22,9395 | 31,7353 | 33,1987 | 47,5432 | 51,9796 | 48,1552 | Smpdl3a | 2,240705934 | 2,86E-06 |
| Clip4 | 1,20648 | 5,39381 | 16,6152 | 11,426 | 30,9137 | 49,4476 | 51,4278 | 92,454 | 101,076 | 98,9807 | Clip4 | 3,036355173 | 2,99E-06 |
| Rsad1 | 1,31467 | 1,14979 | 3,87309 | 3,23348 | 6,8574 | 11,8981 | 10,7143 | 8,05622 | 8,75407 | 9,31686 | Rsad1 | 2,310320558 | 3,00E-06 |
| Cdnf | 0 | 0,107349 | 0,544982 | 0,22769 | 1,38799 | 3,99043 | 4,10434 | 20,2071 | 23,5005 | 22,1406 | Cdnf | 3,785127993 | 3,11E-06 |
| Ppifos | 0 | 0 | 0 | 0 | 0 | 0 | 0 | 0 | 0 | 0 | Ppifos | 2,963358607 | 3,18E-06 |
| Deptor | 0,261144 | 0,110409 | 0,217989 | 0,263135 | 0,868243 | 1,15101 | 1,0732 | 2,7929 | 3,00788 | 2,69985 | Deptor | 2,461984075 | 3,21E-06 |
| Cdkn2c | 0,68494 | 2,02696 | 3,14383 | 0 | 13,1548 | 13,9199 | 14,2439 | 4,95468 | 7,80453 | 10,2942 | Cdkn2c | 2,788105655 | 3,23E-06 |
| Aldh5a1 | 1,40061 | 6,35584 | 10,3819 | 8,63665 | 22,0699 | 34,5737 | 32,1776 | 21,419 | 23,3501 | 24,125 | Aldh5a1 | 2,447187945 | 3,25E-06 |
| 2900076A07R | 2,39067 | 8,48945 | 4,78769 | 3,85004 | 3,8177 | 6,30192 | 8,12408 | 6,46467 | 8,20327 | 8,32303 | 2900076A07R | 2,335897261 | 3,30E-06 |
| Them6 | 1,43683 | 1,65361 | 0,599592 | 2,17059 | 6,93826 | 6,14796 | 5,2274 | 19,969 | 20,876 | 17,7686 | Them6 | 2,346209468 | 3,33E-06 |
| Metap1d | 3,79786 | 8,64554 | 17,7984 | 9,41392 | 21,7604 | 47,8691 | 42,7093 | 44,6954 | 45,9574 | 48,1094 | Metap1d | 2,225761279 | 3,40E-06 |
| Hibch | 7,98198 | 14,1328 | 18,6194 | 10,7193 | 21,5561 | 69,2669 | 70,9292 | 90,6008 | 95,7725 | 93,5894 | Hibch | 2,280870945 | 3,45E-06 |
| Cdh13 | 1,32108 | 4,63369 | 16,785 | 11,8293 | 62,1079 | 70,4062 | 68,7509 | 89,1114 | 94,2048 | 88,4896 | Cdh13 | 2,946199317 | 3,65E-06 |
| Cp | 0 | 0 | 0,409229 | 0 | 4,86237 | 0 | 1,19306 | 20,2503 | 16,6156 | 19,1884 | Cp | 3,240846969 | 4,07E-06 |
| Pygm | 13,3774 | 62,7269 | 131,671 | 87,2195 | 248,118 | 477,306 | 496,98 | 888,531 | 872,631 | 852,296 | Pygm | 2,695332742 | 4,29E-06 |
| Plekhb2 | 8,68578 | 10,166 | 18,8419 | 9,98664 | 19,8591 | 49,4724 | 46,984 | 63,0111 | 66,6486 | 66,998 | Plekhb2 | 2,009174929 | 4,34E-06 |
| Igip | 0,524354 | 0,258626 | 0,21881 | 0,792087 | 2,75648 | 4,55627 | 7,17723 | 3,56262 | 8,07312 | 6,14852 | Igip | 3,534190527 | 4,48E-06 |
| Cited4 | 8,77272 | 11,8269 | 24,6488 | 23,2629 | 39,7387 | 64,5754 | 73,3511 | 29,9137 | 42,4332 | 40,146 | Cited4 | 2,051210738 | 4,64E-06 |
| Zfp52 | 0,103241 | 0,101847 | 0,706638 | 0,83209 | 4,00871 | 2,56235 | 2,40577 | 2,81624 | 4 | 4,26155 | Zfp52 | 2,974449207 | 4,87E-06 |
| Fam120aos | 2,20205 | 2,17224 | 5,71771 | 3,69617 | 9,41799 | 12,7497 | 12,3381 | 7,5392 | 7,02946 | 8,04375 | Fam120aos | 2,025277808 | 4,97E-06 |
| Arhgap26 | 0,475185 | 3,04703 | 3,23937 | 2,19452 | 9,69792 | 8,81895 | 8,60132 | 16,912 | 19,2348 | 27,3278 | Arhgap26 | 2,247320168 | 5,21E-06 |
| Acox3 | 0,976411 | 5,05695 | 4,55055 | 3,77103 | 10,443 | 15,3844 | 14,4386 | 8,66073 | 10,7373 | 12,3443 | Acox3 | 2,161784871 | 5,25E-06 |
| Slc25a40 | 0,835567 | 1,17754 | 2,05906 | 1,44303 | 4,01103 | 6,43555 | 6,32932 | 4,55684 | 5,26912 | 4,63488 | Slc25a40 | 2,412443821 | 5,31E-06 |
| Ppil6 | 0,356938 | 0 | 0 | 0,718751 | 2,94593 | 6,04004 | 6,05001 | 1,29056 | 1,09368 | 0,583429 | Ppil6 | 4,201978345 | 5,34E-06 |
| Krt222 | 0 | 0,596021 | 0,605165 | 0,365198 | 1,6101 | 5,84964 | 5,07761 | 18,3414 | 21,1521 | 17,2637 | Krt222 | 3,39955696 | 5,63E-06 |
| Nudt7 | 0,690382 | 2,53465 | 10,398 | 2,83188 | 17,1642 | 41,8211 | 33,8644 | 74,3831 | 86,1647 | 87,6879 | Nudt7 | 3,173671551 | 5,70E-06 |
| Hcn2 | 0,612998 | 0,907077 | 0,255834 | 0,205857 | 1,96897 | 4,09792 | 3,79027 | 18,8586 | 17,3836 | 21,3331 | Hcn2 | 2,916884787 | 5,97E-06 |
| Gpihbp1 | 0,525193 | 0 | 0 | 0 | 19,7843 | 5,33213 | 2,76084 | 7,56606 | 20,9689 | 20,0458 | Gpihbp1 | 5,009885964 | 5,99E-06 |
| Grip2 | 3,08787 | 2,61972 | 5,61901 | 6,22243 | 8,25597 | 19,5461 | 17,9926 | 5,84572 | 8,6758 | 8,38409 | Grip2 | 2,085232578 | 6,18E-06 |
| Pex11a | 0 | 0,907608 | 1,53557 | 0,463078 | 4,45916 | 6,41003 | 8,24025 | 17,0091 | 18,2904 | 20,0823 | Pex11a | 3,439177371 | 6,19E-06 |
| Lysmd4 | 0,786679 | 0,544338 | 3,50039 | 1,78325 | 7,22692 | 8,09385 | 7,37756 | 7,72998 | 12,0648 | 9,28878 | Lysmd4 | 2,447707962 | 6,25E-06 |
| Camk2b | 0,324949 | 0,560982 | 1,28839 | 0,98217 | 2,09446 | 4,96611 | 4,22007 | 7,08377 | 7,321 | 7,71459 | Camk2b | 2,503117032 | 6,54E-06 |
| Dnajc4 | 14,1779 | 2,89349 | 15,0941 | 18,2067 | 27,6787 | 43,6329 | 43,6584 | 41,1929 | 41,8848 | 42,7299 | Dnajc4 | 2,098177143 | 6,69E-06 |
| Lingo3 | 0,0944859 | 0,0932099 | 0,0788678 | 0,285578 | 0,589059 | 3,38643 | 3,30204 | 10,0539 | 16,0057 | 7,81299 | Lingo3 | 3,962722125 | 6,83E-06 |
| Gca | 0,990715 | 3,12747 | 5,12709 | 1,79661 | 7,63694 | 16,6429 | 17,0802 | 8,9191 | 12,2145 | 10,6469 | Gca | 2,461821161 | 6,85E-06 |
| Rtn4ip1 | 5,7241 | 6,94988 | 13,3229 | 8,09555 | 14,2593 | 38,3391 | 37,9585 | 55,3419 | 67,6165 | 67,2656 | Rtn4ip1 | 2,067283047 | 7,06E-06 |
| Ttll11 | 0,204607 | 0,201844 | 0,59775 | 0,721469 | 2,87046 | 1,74397 | 2,03884 | 1,48839 | 1,5077 | 1,35789 | Ttll11 | 2,464335165 | 7,56E-06 |
| Dusp28 | 1,67235 | 4,94913 | 1,79454 | 4,81217 | 9,16127 | 16,4158 | 14,4699 | 19,056 | 17,355 | 19,8256 | Dusp28 | 2,274063169 | 8,22E-06 |
| Filip1l | 1,20257 | 4,64644 | 6,60824 | 5,85369 | 13,1996 | 14,1331 | 13,4856 | 4,09881 | 6,83477 | 6,75312 | Filip1l | 2,061679698 | 8,43E-06 |
| Pcp4l1 | 5,15476 | 17,2448 | 52,2816 | 32,73 | 89,3267 | 189,848 | 185,264 | 218,656 | 273,12 | 264,058 | Pcp4l1 | 2,706653572 | 8,52E-06 |
| Larp1b | 6,34038 | 3,6317 | 6,65751 | 8,6521 | 24,706 | 34,4558 | 32,4873 | 43,0437 | 50,6462 | 47,7933 | Larp1b | 2,287171629 | 8,67E-06 |
| 1110054M08I | 0 | 0 | 0 | 0 | 0 | 0 | 0 | 30,0235 | 35,228 | 42,249 | 1110054M08I | 2,848389796 | 8,87E-06 |
| Ndrg4 | 4,04183 | 38,3217 | 92,2487 | 66,8467 | 245,492 | 416,793 | 418,559 | 290,794 | 277,294 | 325,806 | Ndrg4 | 2,985171881 | 9,66E-06 |
| Myl2 | 177,996 | 828,003 | 1842,53 | 2059,54 | 5849,31 | 7818,63 | 7980,41 | 9925,4 | 7756,64 | 7499,64 | Myl2 | 2,769577002 | 1,04E-05 |
| Mcee | 7,63321 | 20,7055 | 16,9832 | 32,0009 | 40,099 | 87,8159 | 79,7066 | 123,397 | 131,685 | 138,895 | Mcee | 2,323986457 | 1,05E-05 |
| Zfp641 | 0,0880322 | 0,26053 | 1,10222 | 0,44346 | 2,41956 | 4,08103 | 4,45225 | 3,60332 | 3,06375 | 3,75983 | Zfp641 | 3,054609762 | 1,11E-05 |
| Ip6k3 | 0 | 0 | 0 | 0,293607 | 0,826641 | 3,62141 | 3,66712 | 10,3255 | 9,74047 | 14,2715 | Ip6k3 | 4,576926823 | 1,12E-05 |
| Slc16a8 | 0 | 0,190732 | 0,322749 | 0,389492 | 0,878068 | 1,26358 | 1,36596 | 1,05262 | 1,23761 | 1,06779 | Slc16a8 | 2,841802775 | 1,14E-05 |
| Klhdc1 | 0,878504 | 1,3866 | 2,63967 | 0,530939 | 2,8418 | 10,0862 | 10,1426 | 25,1994 | 27,2569 | 23,3864 | Klhdc1 | 2,883793561 | 1,25E-05 |
| Mov10l1 | 2,14192 | 7,0042 | 12,449 | 10,3905 | 20,9805 | 36,4788 | 37,8292 | 22,8068 | 14,0091 | 18,7827 | Mov10l1 | 2,285267724 | 1,25E-05 |
| Ppp2r3a | 6,60255 | 16,0979 | 32,0331 | 21,4697 | 40,0277 | 86,5327 | 89,2814 | 103,211 | 115,945 | 128,933 | Ppp2r3a | 2,084039637 | 1,26E-05 |
| Habp4 | 4,14331 | 12,5174 | 24,1003 | 14,0871 | 54,7201 | 53,9247 | 54,0395 | 37,191 | 41,2274 | 42,4216 | Habp4 | 2,122914585 | 1,26E-05 |
| Lrrc3b | 0,421821 | 0,62417 | 0,880155 | 1,27456 | 2,75455 | 6,15276 | 6,12427 | 19,1294 | 37,4876 | 22,0772 | Lrrc3b | 2,927954304 | 1,30E-05 |
| Dhrs4 | 12,9525 | 29,6509 | 27,1503 | 32,7613 | 35,0371 | 100,599 | 97,6036 | 128,847 | 135,667 | 128,97 | Dhrs4 | 2,006120636 | 1,31E-05 |
| Sirt5 | 1,8609 | 5,5071 | 14,1996 | 9,90559 | 17,2244 | 39,341 | 42,1968 | 49,0046 | 48,2729 | 39,7505 | Sirt5 | 2,409744245 | 1,34E-05 |
| C230035I16Ri | 1,52356 | 0 | 0,635551 | 0 | 6,98221 | 5,64459 | 5,44251 | 2,04667 | 4,25878 | 4,04527 | C230035I16Ri | 3,339514068 | 1,36E-05 |
| Mtm1 | 1,78445 | 2,24933 | 3,15417 | 2,75257 | 3,34519 | 6,65885 | 6,87761 | 11,2005 | 13,9816 | 14,3972 | Mtm1 | 2,227721114 | 1,38E-05 |
| D830046C22R | 0,284379 | 0,140268 | 0,356044 | 0,429708 | 2,82275 | 2,17691 | 1,83249 | 1,67923 | 1,9576 | 1,15254 | D830046C22R | 2,997635854 | 1,43E-05 |
| Col4a4 | 0 | 0 | 0 | 0,0403787 | 0,794218 | 0,411143 | 0,4295 | 0,0364917 | 0,133695 | 0,10354 | Col4a4 | 4,024651413 | 1,48E-05 |
| Ifi35 | 0,532618 | 2,62702 | 3,55612 | 1,07275 | 7,94625 | 9,92808 | 8,51975 | 6,27193 | 9,19404 | 9,36961 | Ifi35 | 2,210143464 | 1,50E-05 |
| 3110057O12R | 3,00934 | 2,19902 | 3,98172 | 6,08558 | 6,44109 | 27,5306 | 27,786 | 37,0254 | 45,9279 | 42,9507 | 3110057O12R | 2,547086092 | 1,54E-05 |
| Hfe2 | 0,506055 | 2,16325 | 11,8267 | 5,09747 | 14,8366 | 63,7664 | 60,5407 | 195,721 | 235,809 | 216,144 | Hfe2 | 3,391023041 | 1,55E-05 |
| Rab11fip5 | 0,783229 | 2,77835 | 14,1965 | 9,25467 | 53,5914 | 56,1819 | 53,75 | 29,5219 | 34,1369 | 26,5786 | Rab11fip5 | 3,05060064 | 1,67E-05 |
| Gm5512 | 2,028225 | 2,293585 | 4,08755 | 3,83662 | 4,99901 | 13,7045 | 14,2543 | 10,9062 | 13,21975 | 12,1193 | Gm5512 | 2,237572418 | 1,69E-05 |
| Krt80 | 0 | 0 | 0 | 0,787491 | 7,65633 | 1,96533 | 2,46168 | 0 | 0,263431 | 0,698527 | Krt80 | 4,004930691 | 1,74E-05 |
| N28178 | 0 | 0,472508 | 1,19941 | 1,54418 | 3,692 | 6,79494 | 6,69577 | 2,43934 | 2,85448 | 2,38735 | N28178 | 3,129959096 | 1,74E-05 |
| Gm14420 | 1,71282 | 2,35835 | 1,8186 | 1,02946 | 3,14879 | 3,45715 | 3,05256 | 17,2219 | 18,7432 | 27,1888 | Gm14420 | 2,034859261 | 1,90E-05 |
| Dram1 | 0,117477 | 0,115889 | 0,490281 | 0,118349 | 1,1656 | 2,35706 | 2,30591 | 1,28139 | 1,37851 | 0,684957 | Dram1 | 3,309301086 | 1,95E-05 |
| Clcnkb | 0 | 0 | 0 | 0,216575 | 0,65038 | 1,54688 | 1,62731 | 0,390608 | 0,385059 | 0,92946 | Clcnkb | 4,233266946 | 2,01E-05 |
| Bche | 0,863072 | 3,05511 | 3,94121 | 3,06925 | 6,05257 | 11,6915 | 10,9782 | 6,1472 | 8,84751 | 8,26732 | Bche | 2,062617572 | 2,04E-05 |
| Fam63b | 0,69628 | 1,67905 | 2,84149 | 2,84532 | 4,76951 | 7,99996 | 8,67057 | 9,19396 | 13,4204 | 10,8809 | Fam63b | 2,079716241 | 2,06E-05 |
| Il17rc | 0,887575 | 1,39485 | 1,48166 | 1,04311 | 5,05254 | 2,83324 | 2,95029 | 2,28425 | 1,51929 | 2,03293 | Il17rc | 2,051737987 | 2,06E-05 |
| Glt28d2 | 0,362223 | 1,90575 | 1,00781 | 0,486544 | 3,14918 | 5,48156 | 4,8884 | 6,03579 | 5,70893 | 5,63262 | Glt28d2 | 2,504155938 | 2,21E-05 |
| Csrp3 | 76,1365 | 224,423 | 398,388 | 394,967 | 982,141 | 975,173 | 895,277 | 1949,78 | 1819,72 | 1957,22 | Csrp3 | 2,063823086 | 2,24E-05 |
| Oplah | 1,1349 | 2,95887 | 12,5181 | 7,51392 | 24,6779 | 34,4244 | 36,2226 | 27,2866 | 29,2574 | 28,9753 | Oplah | 2,652324014 | 2,25E-05 |
| Sdsl | 3,37569 | 2,11905 | 8,96384 | 7,10733 | 10,9082 | 19,2322 | 20,0766 | 11,1087 | 10,7266 | 9,31196 | Sdsl | 2,370470326 | 2,27E-05 |
| Klhl41 | 0,917678 | 2,6253 | 16,5248 | 12,0631 | 36,0092 | 64,4155 | 67,0925 | 67,6454 | 81,9793 | 74,3738 | Klhl41 | 3,063474484 | 2,28E-05 |
| Gm16576 | 0,645143 | 0,462862 | 0,244781 | 0,0590932 | 1,59459 | 2,07047 | 1,86851 | 1,49475 | 1,25375 | 1,40269 | Gm16576 | 2,59348905 | 2,31E-05 |
| Dynlt1b | 14,9549 | 16,3734 | 10,7045 | 11,7734 | 21,1196 | 26,1217 | 21,9803 | 4,86502 | 6,0498 | 7,51379 | Dynlt1b | 3,60164715 | 2,35E-05 |
| Zbtb4 | 0,439882 | 0,788994 | 2,83736 | 1,41028 | 6,26112 | 5,45368 | 5,17988 | 4,91603 | 6,1637 | 5,59229 | Zbtb4 | 2,265754778 | 2,86E-05 |
| Acss1 | 0,270276 | 3,9994 | 9,92646 | 6,71675 | 22,1587 | 48,3039 | 47,1182 | 249,723 | 264,245 | 231,471 | Acss1 | 2,962129472 | 2,92E-05 |
| Rph3al | 0,795347 | 1,62306 | 8,36443 | 3,36491 | 11,8905 | 30,0106 | 27,8286 | 5,20573 | 4,2243 | 4,15443 | Rph3al | 2,789179237 | 3,15E-05 |
| 2310040G24R | 0 | 0,538503 | 0,455521 | 7,14224 | 15,897 | 41,4568 | 43,2348 | 34,5655 | 31,9402 | 41,7085 | 2310040G24R | 3,89869641 | 3,15E-05 |
| Btnl9 | 0 | 0 | 0 | 0 | 2,99652 | 0,7479 | 0,623478 | 0,369277 | 2,41006 | 2,2396 | Btnl9 | 5,129380607 | 3,25E-05 |
| Rxfp1 | 0 | 0 | 0,493167 | 0,892794 | 2,63941 | 4,33829 | 3,6718 | 0,134173 | 0,148857 | 0,191611 | Rxfp1 | 3,135692585 | 3,30E-05 |
| Gm15417 | 0,852788 | 4,18073 | 0,407425 | 1,96655 | 6,56395 | 9,92744 | 9,03606 | 13,4938 | 15,0558 | 16,684 | Gm15417 | 2,65146853 | 3,32E-05 |
| Ggact | 3,53208 | 4,18101 | 15,6223 | 5,33434 | 25,0285 | 31,0409 | 26,4487 | 16,6034 | 25,8475 | 28,7085 | Ggact | 2,168879637 | 3,33E-05 |
| Rab7l1 | 1,566 | 0 | 2,61383 | 3,46911 | 6,23507 | 9,32221 | 8,67926 | 5,38449 | 6,1671 | 5,62185 | Rab7l1 | 2,280912353 | 3,41E-05 |
| Adra1d | 0 | 0,137883 | 0,583319 | 0,844809 | 2,29899 | 3,50777 | 3,62394 | 5,46024 | 4,92738 | 4,48605 | Adra1d | 3,121870394 | 3,41E-05 |
| Snta1 | 1,92392 | 12,0201 | 32,6517 | 20,026 | 74,3863 | 90,2238 | 87,6341 | 76,142 | 67,428 | 88,9775 | Snta1 | 2,539510763 | 3,46E-05 |
| Fam213b | 5,36305 | 11,1092 | 25,5071 | 14,0329 | 27,4014 | 67,8118 | 62,2035 | 173,311 | 137,251 | 187,663 | Fam213b | 2,11700076 | 3,61E-05 |
| Pxmp4 | 3,52789 | 5,12018 | 11,356 | 6,98343 | 10,4714 | 30,005 | 30,1231 | 34,84 | 35,2764 | 33,577 | Pxmp4 | 2,158412897 | 3,70E-05 |
| E530011L22Ri | 0 | 0 | 0 | 0 | 0 | 4,3041 | 3,3872 | 0 | 2,74798 | 2,40045 | E530011L22Ri | 2,020621944 | 3,84E-05 |
| Spata2l | 0 | 0 | 0 | 0 | 2,19423 | 4,58975 | 3,3669 | 3,25895 | 3,73149 | 3,14699 | Spata2l | 2,610326712 | 3,88E-05 |
| 2010002M12I | 0 | 0 | 0,080672 | 0,0973699 | 0,849057 | 1,51404 | 0,93332 | 1,36234 | 1,52091 | 1,67355 | 2010002M12I | 4,120316179 | 3,89E-05 |
| Pkia | 1,93168 | 8,92163 | 11,4333 | 7,07693 | 13,4098 | 40,528 | 39,6699 | 98,7546 | 138,86 | 116,705 | Pkia | 2,341093329 | 3,91E-05 |
| 9030612E09R | 0,184437 | 0 | 0,461817 | 0,557327 | 1,7273 | 3,52241 | 3,39855 | 3,18034 | 3,91231 | 3,0547 | 9030612E09R | 3,298098411 | 4,05E-05 |
| Zfp3 | 0,37875 | 1,49451 | 2,52895 | 2,28896 | 7,90099 | 5,56864 | 5,23482 | 6,35836 | 4,65614 | 5,47451 | Zfp3 | 2,20235925 | 4,17E-05 |
| Sec31b | 0,246277 | 0 | 1,37047 | 1,24063 | 2,32586 | 4,38091 | 5,33385 | 5,00365 | 7,88132 | 7,40902 | Sec31b | 2,755760447 | 4,33E-05 |
| Doc2g | 2,06362 | 7,88824 | 29,0635 | 31,6928 | 59,4902 | 156,079 | 162,244 | 304,049 | 234,038 | 207,58 | Doc2g | 2,914589557 | 4,67E-05 |
| Corin | 1,83344 | 13,4537 | 18,6534 | 13,1316 | 35,0973 | 57,2998 | 56,8895 | 92,8578 | 97,5023 | 109,447 | Corin | 2,347785061 | 4,69E-05 |
| A530016L24R | 1,88676 | 2,97802 | 4,51455 | 2,28087 | 4,77882 | 10,6023 | 12,0575 | 23,7769 | 28,6494 | 21,5605 | A530016L24R | 2,067888652 | 4,73E-05 |
| Nlrx1 | 0,35091 | 0,0891395 | 2,56441 | 0,972236 | 6,8358 | 7,73061 | 7,97922 | 11,5706 | 11,6252 | 12,4608 | Nlrx1 | 3,01681574 | 4,80E-05 |
| Plce1 | 0,65474 | 1,97001 | 5,00086 | 3,19973 | 9,10086 | 12,0765 | 12,2134 | 5,63523 | 7,60355 | 6,89696 | Plce1 | 2,253607414 | 4,83E-05 |
| D830013O20I | 0 | 0 | 0 | 0 | 1,29207 | 4,11051 | 3,60169 | 4,6371 | 3,13494 | 5,12102 | D830013O20I | 5,033013837 | 5,01E-05 |
| Ctnna3 | 0,790459 | 5,37185 | 9,45718 | 8,224 | 30,2575 | 39,498 | 35,0241 | 16,324 | 19,0844 | 21,3321 | Ctnna3 | 2,521619913 | 5,18E-05 |

| Gene | d9-11 (1) | d9-11 (2) | d9-11(3) | d9-11(4) | newborn(1) | newborn(2) | newborn(3) | adult(1) | adult(2) | adult(3) | Gene | log2(fold) | FDR(*p* -value) |
|---|---|---|---|---|---|---|---|---|---|---|---|---|---|
| Fam131a | 1,03702 | 0 | 1,60748 | 1,3431 | 3,15136 | 5,35511 | 4,31906 | 30,815 | 33,3925 | 35,216 | Fam131a | 2,330160512 | 5,19E-05 |
| Slitrk4 | 0,490964 | 0,32289 | 1,16114 | 0,906861 | 2,08651 | 2,84965 | 2,83385 | 0,223322 | 0,739042 | 0,608802 | Slitrk4 | 2,138354094 | 5,39E-05 |
| D2hgdh | 3,51627 | 4,47243 | 9,14771 | 7,21079 | 10,9125 | 33,77 | 29,2594 | 29,9979 | 41,2279 | 34,4424 | D2hgdh | 2,15470011 | 5,41E-05 |
| Adamts6 | 0,85882 | 1,69445 | 4,35641 | 3,6609 | 6,35282 | 9,47966 | 8,48078 | 0,36073 | 0,485685 | 0,555292 | Adamts6 | 2,072459356 | 5,64E-05 |
| Btc | 0 | 0 | 0,0955112 | 0 | 0,616301 | 1,77915 | 1,43879 | 0,208037 | 0,997261 | 0,518855 | Btc | 4,110119837 | 5,71E-05 |
| Cox5b | 105,441 | 166,686 | 170,407 | 248,74 | 337,443 | 558,58 | 534,946 | 680,396 | 628,37 | 615,463 | Cox5b | 2,134369317 | 5,76E-05 |
| Gm16907 | 0 | 0 | 0 | 0 | 0 | 0 | 0 | 0 | 0 | 0 | Gm16907 | 3,547536871 | 5,94E-05 |
| Tha1 | 0,375014 | 0,369942 | 3,13002 | 2,45526 | 5,74737 | 13,0531 | 12,7271 | 13,1023 | 13,9553 | 13,3981 | Tha1 | 2,803312699 | 6,16E-05 |
| 4732491K20R | 0 | 0 | 0,429419 | 0,621957 | 1,80802 | 3,06954 | 3,34404 | 1,49693 | 2,06786 | 1,83207 | 4732491K20R | 3,353692714 | 6,21E-05 |
| Nt5c1a | 0 | 2,3728 | 0 | 0,345989 | 2,83548 | 8,24172 | 7,83443 | 20,196 | 19,6374 | 18,7504 | Nt5c1a | 3,555259165 | 6,21E-05 |
| Mamstr | 1,08566 | 0,305997 | 1,16507 | 0,312463 | 3,8715 | 5,10123 | 4,90312 | 3,38047 | 4,69106 | 4,02486 | Mamstr | 2,322935199 | 6,45E-05 |
| Chpt1 | 1,67801 | 14,9088 | 20,3231 | 10,9183 | 23,2235 | 57,023 | 55,7498 | 241,785 | 244,773 | 247,457 | Chpt1 | 2,3852162 | 6,89E-05 |
| 4632428C04R | 0,412865 | 0,407285 | 0,574346 | 0,138636 | 1,05372 | 3,97163 | 4,39143 | 5,93917 | 7,20707 | 6,915 | 4632428C04R | 3,034028886 | 6,97E-05 |
| Cpt1a | 0,148985 | 2,86599 | 6,77765 | 7,28018 | 46,384 | 83,0719 | 78,6733 | 17,2838 | 18,7424 | 19,6817 | Cpt1a | 3,820690951 | 7,00E-05 |
| Gm5860 | 0,117873 | 0 | 0,0737759 | 0,356036 | 1,53466 | 4,68839 | 3,86843 | 6,71296 | 5,3562 | 5,49032 | Gm5860 | 4,45444485 | 7,04E-05 |
| Ltbp3 | 0 | 0 | 8,01693 | 5,54726 | 26,6358 | 22,6083 | 22,3767 | 8,57446 | 8,37894 | 9,35262 | Ltbp3 | 2,816434006 | 7,11E-05 |
| Pxmp2 | 1,13898 | 2,99607 | 8,87123 | 9,55579 | 17,943 | 35,1187 | 38,0432 | 135,484 | 133,2 | 112,173 | Pxmp2 | 2,52030154 | 7,20E-05 |
| Rgs1 | 0 | 0 | 0,228252 | 0 | 6,76243 | 2,27389 | 1,178 | 0 | 0 | 0,0891205 | Rgs1 | 4,519084427 | 7,74E-05 |
| Slc7a4 | 0,201891 | 0 | 0,421296 | 0,406796 | 1,14436 | 2,25215 | 2,29026 | 0,734337 | 1,01425 | 1,20908 | Slc7a4 | 2,994514382 | 8,63E-05 |
| Tmem82 | 1,81036 | 0,174841 | 0 | 2,85142 | 3,65714 | 12,5614 | 11,0392 | 29,693 | 20,7883 | 27,0027 | Tmem82 | 3,147530781 | 8,93E-05 |
| Slc40a1 | 1,15427 | 0,9489 | 2,89042 | 2,71343 | 3,54365 | 10,4153 | 10,0555 | 8,22281 | 10,2091 | 9,24634 | Slc40a1 | 2,282128386 | 8,94E-05 |
| B430212C06R | 0 | 0,74853 | 0,633342 | 0,1274 | 1,2191 | 2,91875 | 3,14224 | 0,459711 | 0,551504 | 0,77859 | B430212C06R | 2,948136039 | 9,02E-05 |
| Ppp1r1b | 3,15055 | 3,7295 | 5,43435 | 10,1543 | 21,6466 | 31,111 | 31,3007 | 1,84061 | 1,70005 | 1,09242 | Ppp1r1b | 2,328149998 | 9,15E-05 |
| Gsn | 3,9876 | 17,501 | 62,6378 | 41,8734 | 253,496 | 143,823 | 138,596 | 9,75592 | 22,124 | 15,9282 | Gsn | 2,645393754 | 9,44E-05 |
| Lsmem1 | 0 | 3,91956 | 6,88717 | 0,923319 | 11,7333 | 27,9028 | 26,1587 | 26,5544 | 35,5741 | 27,6228 | Lsmem1 | 3,00433094 | 9,51E-05 |
| Lace1 | 3,44565 | 3,26835 | 7,41124 | 6,94219 | 8,39246 | 24,8281 | 24,557 | 41,1833 | 44,3403 | 45,2717 | Lace1 | 2,039086189 | 9,55E-05 |
| 2610507I01Ri | 1,24511 | 0,334989 | 4,06266 | 1,25437 | 6,46323 | 12,4989 | 10,6568 | 7,61451 | 10,9834 | 10,9978 | 2610507I01Ri | 2,526464296 | 9,61E-05 |
| Pcdhgb8 | 0 | 0 | 0 | 0 | 0 | 0 | 0 | 0 | 0 | 0 | Pcdhgb8 | 2,193396557 | 9,87E-05 |
| Asb18 | 0,769291 | 0,252958 | 0,428029 | 0 | 1,45719 | 3,35788 | 3,47011 | 9,99033 | 9,84488 | 12,9852 | Asb18 | 3,219892091 | 9,91E-05 |
| Tst | 2,42459 | 2,39171 | 1,15623 | 7,32363 | 13,6011 | 13,3804 | 14,3006 | 17,8504 | 21,9186 | 19,0151 | Tst | 2,244523024 | 0,000100481 |
| Klhdc8a | 0 | 0 | 0,250686 | 0,0756462 | 0,382855 | 2,59539 | 1,84061 | 12,4332 | 15,6401 | 14,3987 | Klhdc8a | 3,891864702 | 0,000103633 |
| Trmt2b | 2,19891 | 9,80598 | 13,7657 | 10,4725 | 16,0045 | 39,6369 | 39,7297 | 28,6302 | 37,1213 | 37,3694 | Trmt2b | 2,186624449 | 0,000105101 |
| Aox1 | 0,511305 | 0,144115 | 0,121942 | 0,147188 | 1,05528 | 1,24411 | 1,18626 | 8,77318 | 11,1234 | 10,5111 | Aox1 | 2,673474346 | 0,000107898 |
| 9330102E08R | 0,167623 | 0,248039 | 0,279834 | 0,337762 | 0,926108 | 1,42796 | 1,69537 | 1,21988 | 1,71039 | 1,4099 | 9330102E08R | 2,552180307 | 0,000111414 |
| Irgm2 | 0,138418 | 0,546195 | 0,847281 | 0,278908 | 1,84786 | 1,96576 | 2,89945 | 4,23837 | 6,36303 | 5,22591 | Irgm2 | 2,644715307 | 0,000113023 |
| Acad12 | 0,423009 | 1,80826 | 3,64843 | 2,98287 | 5,73824 | 17,8025 | 15,7487 | 60,3338 | 59,583 | 57,677 | Acad12 | 2,492180538 | 0,000117373 |
| Tceal7 | 0,403857 | 1,19513 | 1,01105 | 2,03284 | 4,25593 | 9,29478 | 7,59921 | 1,09396 | 0,275823 | 0,396788 | Tceal7 | 2,665829177 | 0,00011829 |
| Mypn | 0,0604923 | 0,298379 | 2,8277 | 2,74274 | 23,5233 | 35,6798 | 33,0698 | 61,1706 | 60,3002 | 59,6507 | Mypn | 3,968960995 | 0,000119935 |
| Gm3646 | 0 | 0 | 0 | 0 | 7,79976 | 18,9504 | 16,9865 | 26,3495 | 30,8027 | 28,919 | Gm3646 | 4,805811106 | 0,000122672 |
| D830032E09R | 0 | 0 | 0 | 0 | 0,646691 | 1,03029 | 1,35281 | 3,45119 | 3,88139 | 4,48624 | D830032E09R | 4,788812426 | 0,000127972 |
| BC029214 | 0,803447 | 8,32168 | 16,0913 | 8,49286 | 30,6628 | 19,2014 | 17,4718 | 8,83814 | 8,22984 | 12,1029 | BC029214 | 2,106881994 | 0,00012967 |
| 1700123M08R | 0,358622 | 0 | 0,299278 | 0 | 2,24544 | 4,52897 | 3,20516 | 3,2415 | 5,12847 | 3,04833 | 1700123M08R | 3,575841459 | 0,000139255 |
| Cacng6 | 6,99664 | 6,37057 | 3,5926 | 2,70818 | 6,14508 | 19,4054 | 23,9716 | 2,90668 | 2,22482 | 2,12614 | Cacng6 | 2,155947354 | 0,000142791 |
| P2rx5 | 0,146929 | 0,144943 | 0,245273 | 0,296017 | 1,62522 | 2,21289 | 2,38973 | 14,4139 | 12,7824 | 13,4841 | P2rx5 | 2,289079765 | 0,000144165 |
| Vstm4 | 0 | 2,2577 | 0,703789 | 0,84943 | 6,79559 | 5,19657 | 5,02454 | 3,17497 | 5,57408 | 5,307 | Vstm4 | 2,717817603 | 0,000147028 |
| Nhlrc1 | 0 | 0,289048 | 1,46739 | 0,295162 | 3,65654 | 3,86132 | 4,27107 | 7,31928 | 7,52895 | 7,26879 | Nhlrc1 | 2,974511978 | 0,00014795 |
| Ucp2 | 3,02987 | 11,1895 | 23,702 | 19,5683 | 26,5545 | 88,9313 | 88,7802 | 8,97672 | 12,4708 | 5,37718 | Ucp2 | 2,456093114 | 0,000151552 |
| Aifm2 | 0 | 0 | 6,61915 | 0 | 19,7888 | 35,496 | 34,7944 | 25,6066 | 24,4179 | 29,8776 | Aifm2 | 2,919442241 | 0,000156204 |
| Mansc1 | 0 | 1,25129 | 2,11746 | 1,56173 | 4,83627 | 6,8267 | 6,42196 | 1,79253 | 1,41967 | 1,69066 | Mansc1 | 2,542715603 | 0,000158891 |
| Gm572 | 0 | 0 | 1,73998 | 1,31223 | 2,36962 | 4,59437 | 4,96972 | 1,88874 | 3,17429 | 4,07418 | Gm572 | 3,295021479 | 0,0001644 |
| Nqo1 | 0,227984 | 2,24898 | 4,56667 | 1,14807 | 6,92792 | 12,4479 | 13,1745 | 37,6188 | 42,26 | 42,4204 | Nqo1 | 2,631776808 | 0,000166138 |
| Fgf9 | 0 | 0,0765229 | 0,841738 | 0,868374 | 1,82399 | 5,07178 | 4,30265 | 3,03472 | 3,29449 | 2,96044 | Fgf9 | 3,198129843 | 0,000167786 |
| 6430571L13Ri | 0,390932 | 0,257099 | 1,41398 | 1,18146 | 2,25389 | 3,95601 | 4,35741 | 2,84191 | 4,26314 | 3,92749 | 6430571L13Ri | 2,21542166 | 0,000172378 |
| D630045M09I | 2,79525 | 0 | 3,49927 | 2,53338 | 8,18349 | 20,6383 | 15,6536 | 1,26561 | 0,946912 | 1,45771 | D630045M09I | 2,79516917 | 0,000173643 |
| Atp8b1 | 0 | 0,192898 | 0,122416 | 0,443298 | 1,88258 | 0,929916 | 0,898323 | 0,0890112 | 0,195817 | 0,315866 | Atp8b1 | 2,90792981 | 0,000183216 |
| Tap2 | 0 | 0 | 1,22909 | 0,134855 | 5,16248 | 3,72848 | 4,02486 | 10,4605 | 12,2204 | 11,368 | Tap2 | 3,467978012 | 0,000187019 |
| Cystm1 | 1,61213 | 22,2627 | 18,1575 | 21,8914 | 37,1945 | 56,2906 | 52,3715 | 66,3485 | 53,9401 | 65,6406 | Cystm1 | 2,208515285 | 0,000199294 |
| Gdf15 | 1,05351 | 0,346406 | 1,17224 | 0,707144 | 3,6975 | 4,43388 | 5,30227 | 4,12709 | 3,46583 | 3,21368 | Gdf15 | 2,64003184 | 0,000199671 |
| Sema3b | 0,834305 | 1,55463 | 3,47303 | 6,26132 | 7,93304 | 14,1158 | 14,0343 | 17,6249 | 16,8324 | 16,9729 | Sema3b | 2,295547573 | 0,000208952 |
| Olfr608 | 0 | 0,409242 | 0 | 0 | 0,945474 | 3,25379 | 3,71212 | 0 | 0 | 0,135929 | Olfr608 | 4,220499466 | 0,000211212 |
| Itgb1bp2 | 6,5068 | 45,4887 | 59,4984 | 35,8959 | 78,8483 | 153,644 | 152,781 | 186,742 | 214,688 | 209,233 | Itgb1bp2 | 2,114381228 | 0,000217324 |
| Nmnat3 | 0 | 1,23596 | 4,05226 | 1,5776 | 8,30762 | 16,1226 | 13,8923 | 16,6404 | 22,1079 | 16,8168 | Nmnat3 | 2,876074344 | 0,00021747 |
| Fam19a3 | 0,296807 | 1,098 | 0,929064 | 0,672845 | 1,21918 | 4,55368 | 3,84269 | 1,35031 | 1,76145 | 2,06365 | Fam19a3 | 2,280147404 | 0,000222647 |
| Gm11346 | 0 | 0 | 0 | 0 | 2,31504 | 2,22516 | 1,77556 | 0 | 0,230583 | 0,110757 | Gm11346 | 3,534743633 | 0,000225371 |
| Nod1 | 0,631513 | 0,778733 | 1,12179 | 0,159065 | 1,9735 | 2,73084 | 2,31642 | 5,25981 | 7,21062 | 8,84369 | Nod1 | 2,107614918 | 0,00024654 |
| A930016O22R | 0 | 0,140722 | 0 | 0 | 0 | 0 | 0 | 0 | 0 | 0 | A930016O22R | 4,127426947 | 0,000255587 |
| Rnf150 | 0,0321284 | 0,412034 | 1,15322 | 1,1007 | 3,22428 | 3,84209 | 3,85605 | 4,91604 | 8,76359 | 6,47294 | Rnf150 | 2,588847094 | 0,000274127 |
| Dbp | 0 | 3,5512 | 7,32821 | 5,91309 | 26,0155 | 19,4017 | 18,4348 | 91,3435 | 56,5355 | 112,883 | Dbp | 2,24067788 | 0,000286932 |
| Slc41a3 | 0,15154 | 1,61861 | 5,63324 | 5,86006 | 16,8465 | 23,3479 | 20,521 | 55,5592 | 76,3413 | 38,1658 | Slc41a3 | 2,722425652 | 0,000290201 |
| 6820408C15R | 2,37772 | 1,49262 | 1,08245 | 1,08854 | 2,34891 | 6,77541 | 4,01574 | 2,44511 | 2,93807 | 2,83213 | 6820408C15R | 2,18423447 | 0,000290835 |
| Atp5s | 3,60058 | 3,55183 | 8,21381 | 1,93398 | 7,90933 | 17,6118 | 14,7601 | 19,5808 | 21,2911 | 22,7326 | Atp5s | 2,043715401 | 0,000292774 |
| Nppb | 34,5673 | 68,194 | 117,174 | 125,018 | 735,17 | 171,576 | 146,184 | 459,408 | 283,064 | 427,6 | Nppb | 2,217154243 | 0,000296764 |
| Phkg1 | 0 | 0 | 0 | 0 | 0 | 1,71725 | 1,45266 | 15,9536 | 22,7658 | 16,0815 | Phkg1 | 4,64898781 | 0,000298076 |
| Col4a3 | 0 | 0 | 0 | 0,0365246 | 0,441278 | 0,217671 | 0,199785 | 0,0990338 | 0,132994 | 0,17558 | Col4a3 | 4,094056068 | 0,000303264 |
| Steap3 | 0,599421 | 2,71847 | 5,67068 | 6,03345 | 10,8758 | 20,3869 | 21,4553 | 21,7931 | 25,9688 | 22,836 | Steap3 | 2,328514815 | 0,000305918 |
| Enpp5 | 0,136326 | 0,941388 | 1,42435 | 2,33464 | 3,28365 | 8,29964 | 6,40297 | 9,41093 | 8,62346 | 9,48058 | Enpp5 | 2,563279588 | 0,00030615 |
| Myo5c | 0,747086 | 0,921255 | 0,155905 | 0,329334 | 1,104 | 2,55935 | 2,61695 | 1,57297 | 1,74546 | 2,23245 | Myo5c | 2,302559479 | 0,000322666 |
| Ssh3 | 1,45409 | 0 | 3,25634 | 0,244144 | 5,49651 | 8,64704 | 7,76779 | 2,97362 | 3,29145 | 4,08266 | Ssh3 | 2,676496274 | 0,00032951 |
| 0610009L18Ri | 0,757994 | 4,48602 | 10,1183 | 6,8624 | 20,4072 | 25,5623 | 23,5439 | 19,0088 | 20,6535 | 20,3747 | 0610009L18Ri | 2,203904015 | 0,000364956 |
| Mt3 | 0,948635 | 9,3583 | 16,6217 | 12,4034 | 32,9588 | 49,9182 | 41,9735 | 0 | 2,01094 | 3,16689 | Mt3 | 2,212413944 | 0,000367048 |
| Ush1c | 0,333121 | 0,164308 | 2,93017 | 1,34252 | 2,80202 | 6,44678 | 6,67136 | 1,4688 | 2,12862 | 2,1422 | Ush1c | 2,596744179 | 0,000373223 |
| Cpt1b | 28,9148 | 54,1585 | 100,717 | 87,6707 | 300,468 | 642,683 | 662,522 | 537,835 | 519,386 | 524,706 | Cpt1b | 2,491498002 | 0,000373223 |
| Ddn | 0,172715 | 0,170383 | 0,144167 | 0,17401 | 2,39794 | 0,649169 | 0,767605 | 0,0785537 | 0,115583 | 0,16764 | Ddn | 2,929392182 | 0,000373247 |
| Rab3a | 2,39526 | 3,4031 | 15,4527 | 9,19091 | 21,318 | 32,6091 | 31,5284 | 51,3878 | 60,6172 | 57,4428 | Rab3a | 2,095539152 | 0,000373418 |
| 2310002L09Ri | 0 | 0 | 0 | 0,178943 | 0,806576 | 1,78533 | 1,88372 | 33,7018 | 41,0439 | 28,5142 | 2310002L09Ri | 4,047464295 | 0,000377383 |
| Tmem25 | 0,142389 | 0 | 1,18848 | 0,717186 | 1,3326 | 5,3606 | 4,4566 | 11,5124 | 13,8183 | 11,5186 | Tmem25 | 2,853308419 | 0,000379352 |
| S100a4 | 0 | 0 | 0 | 0 | 11,1658 | 6,66359 | 4,37736 | 12,4749 | 11,5146 | 12,6467 | S100a4 | 4,536446483 | 0,000380512 |
| Dok7 | 0 | 0,529862 | 1,45705 | 4,32868 | 7,50116 | 12,6353 | 10,1757 | 13,543 | 12,9345 | 16,7133 | Dok7 | 3,041759923 | 0,000390198 |
| Rab17 | 0,453588 | 0 | 0,189285 | 1,14208 | 2,18952 | 4,03999 | 3,40265 | 1,05328 | 1,6885 | 0,817068 | Rab17 | 3,024919487 | 0,000397115 |
| E330011O21R | 0 | 0 | 0 | 0 | 0,830712 | 3,58351 | 2,82776 | 6,76853 | 9,82374 | 10,3737 | E330011O21R | 4,119014528 | 0,000402259 |
| Kbtbd3 | 0,43387 | 0,281061 | 2,17283 | 1,01981 | 2,24783 | 6,61261 | 6,93183 | 7,93557 | 9,15707 | 9,28694 | Kbtbd3 | 2,607853892 | 0,000406172 |
| Rasd2 | 0 | 0,347266 | 0,195886 | 1,53675 | 4,2911 | 4,04159 | 3,23892 | 2,77315 | 2,01196 | 2,16651 | Rasd2 | 2,93313618 | 0,000408811 |
| Nudt13 | 1,3195 | 1,1715 | 4,18516 | 4,12066 | 5,051 | 11,6868 | 11,2536 | 15,1072 | 13,5052 | 17,7047 | Nudt13 | 2,125614967 | 0,000410357 |
| 5930403L14Ri | 0,167149 | 1,48404 | 4,04621 | 5,05247 | 17,633 | 13,5616 | 13,3407 | 5,07178 | 6,33561 | 5,59066 | 5930403L14Ri | 2,582558386 | 0,000411802 |
| Hnmt | 0 | 0,211131 | 1,25042 | 0,215564 | 2,00539 | 4,41404 | 4,53805 | 7,57015 | 10,3245 | 7,79343 | Hnmt | 3,117077121 | 0,000426469 |
| Rbm12b1 | 2,33198 | 1,81102 | 1,69801 | 2,79928 | 5,20976 | 4,7556 | 3,84096 | 1,62429 | 2,17109 | 1,25273 | Rbm12b1 | 2,040617752 | 0,000433146 |
| Myoz2 | 0 | 3,85018 | 33,116 | 21,6153 | 373,276 | 512,65 | 552,204 | 1101,71 | 992,645 | 1158,25 | Myoz2 | 4,171856395 | 0,000459248 |
| Lrrc48 | 0,370898 | 0,182941 | 0,309567 | 0,373589 | 1,42103 | 2,28356 | 2,33513 | 2,69281 | 3,0598 | 3,37272 | Lrrc48 | 2,568889101 | 0,000463592 |
| Zfp385b | 0,49616 | 0,30165 | 1,27617 | 0,821497 | 2,32642 | 3,91129 | 3,20986 | 3,93075 | 4,90188 | 4,2283 | Zfp385b | 2,107742035 | 0,000491876 |
| Il17d | 0,302908 | 0,597606 | 0,50559 | 1,52508 | 7,06271 | 2,36786 | 2,05145 | 2,74146 | 2,56917 | 2,07516 | Il17d | 2,374664605 | 0,000501996 |
| Znrd1as | 1,202 | 4,34749 | 2,34047 | 0,806715 | 5,02084 | 11,3483 | 12,6056 | 17,3663 | 17,7147 | 15,2193 | Znrd1as | 2,377698923 | 0,000516661 |
| Lrtm1 | 1,67073 | 0,253566 | 0,679431 | 0,43165 | 1,30999 | 5,55337 | 5,88607 | 68,9849 | 113,8 | 87,0747 | Lrtm1 | 2,580766628 | 0,000526074 |
| Gm6086 | 0 | 0 | 0 | 0 | 0,236804 | 0,93017 | 0,806488 | 1,16468 | 1,55593 | 1,60079 | Gm6086 | 4,476090075 | 0,000547392 |
| Prr33 | 0 | 0,296034 | 0,375713 | 0,151147 | 1,5321 | 1,31836 | 0,967021 | 5,17876 | 6,40203 | 7,10485 | Prr33 | 2,837281391 | 0,0005665 |
| Podxl2 | 3,24278 | 8,75498 | 20,6556 | 13,2376 | 26,7792 | 42,3355 | 42,9687 | 8,98675 | 9,76058 | 8,64193 | Podxl2 | 2,036087187 | 0,000569388 |
| Mir208a | 0 | 0 | 0 | 0 | 0 | 1215,04 | 0 | 0 | 0 | 0 | Mir208a | 3,91145051 | 0,000576324 |
| Aes | 21,6713 | 118,854 | 247,136 | 165,378 | 397,617 | 538,397 | 584,609 | 763,486 | 728,874 | 743,275 | Aes | 2,094280378 | 0,000576802 |
| Syt7 | 0,367784 | 0,979922 | 2,16168 | 2,60226 | 2,56873 | 7,13334 | 7,46101 | 9,45696 | 13,3668 | 12,7199 | Syt7 | 2,213841781 | 0,000609573 |
| Kcnk3 | 0,931422 | 7,50529 | 21,934 | 12,6259 | 32,1563 | 60,2202 | 61,862 | 172,185 | 167,886 | 170,298 | Kcnk3 | 2,463268817 | 0,000611768 |
| 2700046G09R | 0,427453 | 0,843337 | 2,31895 | 0,430523 | 1,88119 | 4,24654 | 4,26894 | 1,74452 | 1,65822 | 2,432 | 2700046G09R | 2,065508135 | 0,000616869 |
| Ces1d | 0 | 0 | 0 | 0 | 0,345589 | 1,34042 | 1,97667 | 55,8863 | 83,1699 | 65,5751 | Ces1d | 4,443036416 | 0,000643936 |
| Prdm9 | 0,187631 | 0,185097 | 0,234925 | 0,37807 | 0,664629 | 1,63023 | 1,29737 | 0,511937 | 0,376872 | 0,738797 | Prdm9 | 2,480042437 | 0,000690293 |
| Nsun3 | 1,89065 | 0,532869 | 1,12707 | 1,08798 | 3,07034 | 4,89904 | 3,73909 | 1,4678 | 2,83466 | 1,62802 | Nsun3 | 2,008852895 | 0,000772494 |

| Gene | d9-11 (1) | d9-11 (2) | d9-11(3) | d9-11(4) | newborn(1) | newborn(2) | newborn(3) | adult(1) | adult(2) | adult(3) | Gene | log2(fold) | FDR(*p*-value) |
|---|---|---|---|---|---|---|---|---|---|---|---|---|---|
| 1700012D14R | 0 | 0 | 0 | 0 | 0,211774 | 1,32204 | 1,60755 | 0,662687 | 0,258071 | 0,245283 | 1700012D14R | 4,360922368 | 0,000776994 |
| Colq | 0,11622 | 0,802547 | 3,68628 | 2,80999 | 6,45742 | 11,1633 | 11,0495 | 7,55351 | 9,62349 | 8,41977 | Colq | 2,549174635 | 0,000777142 |
| Sln | 0 | 0 | 3,35296 | 4,0415 | 7,21597 | 29,4332 | 20,05 | 30,4317 | 43,3179 | 8,39716 | Sln | 3,180461736 | 0,000780046 |
| Thrsp | 0 | 0 | 0 | 0 | 1,16128 | 3,4671 | 3,3154 | 29,8258 | 23,6321 | 32,6306 | Thrsp | 3,937995039 | 0,000793085 |
| Kcne1 | 0 | 2,04318 | 5,61856 | 4,90351 | 24,0234 | 83,2103 | 77,2633 | 2,07155 | 3,3995 | 2,01161 | Kcne1 | 3,829661924 | 0,000795343 |
| Clec2d | 0 | 1,50601 | 0 | 0 | 9,49295 | 4,13241 | 3,10204 | 7,09253 | 8,7031 | 5,54552 | Clec2d | 3,507613334 | 0,000797815 |
| S100a9 | 0 | 0 | 0 | 0 | 18,4119 | 2,0586 | 3,43468 | 0 | 0 | 0 | S100a9 | 4,411432965 | 0,000803059 |
| Mamld1 | 0 | 1,91843 | 1,19291 | 1,07757 | 8,65093 | 4,14327 | 2,96333 | 3,84308 | 3,39854 | 3,98081 | Mamld1 | 2,522768333 | 0,000808172 |
| Lancl3 | 0,246476 | 1,37784 | 0,205737 | 0,579431 | 2,11825 | 2,07902 | 2,11513 | 0,672676 | 0,96192 | 0,983357 | Lancl3 | 2,046921312 | 0,000824304 |
| Gm15328 | 0,110238 | 0 | 0,552092 | 0,333176 | 1,12491 | 1,71381 | 1,62264 | 0,501103 | 0,406365 | 0,392686 | Gm15328 | 2,477317211 | 0,000841793 |
| Sardh | 0,420489 | 0,345677 | 5,14787 | 1,97708 | 10,5711 | 13,7083 | 13,5283 | 1,59432 | 1,80376 | 2,13004 | Sardh | 2,584498391 | 0,000841885 |
| Ppara | 0 | 0 | 1,01717 | 0,263103 | 17,0919 | 13,8885 | 12,8431 | 20,0987 | 25,3321 | 22,8916 | Ppara | 4,216091881 | 0,000908147 |
| Armc2 | 0,0654659 | 0,0645822 | 0,27323 | 0,197881 | 0,50066 | 0,786915 | 0,762178 | 7,80485 | 9,34478 | 6,82181 | Armc2 | 2,529298175 | 0,000911405 |
| Rtn4rl1 | 0,295291 | 1,65072 | 5,75119 | 7,93324 | 13,9614 | 23,2171 | 23,8699 | 2,86433 | 4,0871 | 3,02675 | Rtn4rl1 | 2,515796795 | 0,000990406 |
| Lama2 | 0,096111 | 2,68641 | 7,60833 | 3,22816 | 40,0218 | 37,7678 | 36,6198 | 15,494 | 17,0392 | 21,8254 | Lama2 | 3,251797068 | 0,001016308 |
| Smco1 | 0 | 0,0285987 | 0 | 0,0243286 | 0,164838 | 2,30197 | 2,76597 | 20,0787 | 27,3553 | 25,8954 | Smco1 | 4,346939095 | 0,001043248 |
| Mkln1os | 0 | 0 | 0 | 0 | 0,928891 | 2,9902 | 4,5272 | 0,368043 | 2,36707 | 1,7358 | Mkln1os | 3,876997461 | 0,001056746 |
| Adprhl1 | 0,265148 | 9,67762 | 29,874 | 32,3093 | 200,311 | 280,706 | 285,466 | 583,685 | 432,777 | 559,313 | Adprhl1 | 3,485181854 | 0,001056746 |
| 6430710C18R | 0 | 0 | 0,964937 | 0 | 0,505427 | 2,79849 | 2,18117 | 0,403467 | 0,450551 | 0,868273 | 6430710C18R | 3,627887446 | 0,00111402 |
| Gm10033 | 0,946877 | 0,643132 | 2,01332 | 2,37713 | 1,77249 | 4,98205 | 5,04669 | 4,43644 | 5,00845 | 3,74907 | Gm10033 | 2,359366546 | 0,001147405 |
| Adra1b | 0 | 0,803107 | 4,16464 | 3,28171 | 20,6193 | 46,0489 | 43,7385 | 32,5914 | 41,7096 | 32,4689 | Adra1b | 3,610712905 | 0,001173491 |
| Tatdn3 | 0,894471 | 2,97676 | 4,97662 | 6,82619 | 7,37526 | 15,2466 | 12,8946 | 9,1758 | 10,5781 | 12,059 | Tatdn3 | 2,057943974 | 0,001196513 |
| Rufy4 | 0 | 0,287387 | 0 | 0,146733 | 0,950181 | 1,13343 | 1,41007 | 1,05852 | 0,733851 | 1,27532 | Rufy4 | 3,094998741 | 0,001241911 |
| Lmod2 | 2,26061 | 18,698 | 47,0252 | 45,0159 | 139,587 | 111,133 | 114,154 | 761,495 | 709,439 | 674,51 | Lmod2 | 2,273458635 | 0,001289441 |
| Aldh1l2 | 0,41866 | 0,464636 | 6,02835 | 3,74379 | 9,27719 | 15,1766 | 14,5226 | 4,00187 | 5,88839 | 4,85306 | Aldh1l2 | 2,439109705 | 0,001301241 |
| Dand5 | 1,20802 | 3,56946 | 0,756126 | 1,21644 | 3,69253 | 10,8137 | 8,83198 | 15,5801 | 14,8559 | 17,832 | Dand5 | 2,178559025 | 0,0013851 |
| Agt | 0 | 0,925425 | 0 | 0,188983 | 1,75723 | 2,68769 | 3,22697 | 5,10794 | 4,42299 | 4,26567 | Agt | 3,093712006 | 0,00139061 |
| Nme8 | 0 | 0 | 0 | 0 | 0,323158 | 0,858196 | 0,799261 | 0 | 0 | 0 | Nme8 | 4,204857376 | 0,001395603 |
| Slc36a2 | 0 | 0,813466 | 0,841233 | 0,669167 | 0,740499 | 3,74205 | 3,3664 | 22,0181 | 21,7531 | 24,6967 | Slc36a2 | 2,877890857 | 0,001415987 |
| Gal3st3 | 0,827977 | 1,36131 | 2,55704 | 3,19731 | 2,80454 | 8,93463 | 8,29874 | 19,7465 | 24,4833 | 21,3162 | Gal3st3 | 2,055694088 | 0,001457972 |
| Snap91 | 0,96709 | 0,66782 | 11,9841 | 7,39497 | 37,761 | 79,5638 | 78,9143 | 4,41578 | 6,20797 | 6,05158 | Snap91 | 3,272139695 | 0,001469069 |
| Slc1a1 | 0,174299 | 0,0859726 | 1,45489 | 0,439014 | 2,91382 | 2,48952 | 2,43082 | 0,158545 | 0,291621 | 0,592136 | Slc1a1 | 2,43287617 | 0,001522825 |
| 9330188P03R | 0 | 0 | 0,293738 | 0 | 1,50247 | 3,15542 | 2,16924 | 0,954592 | 0,598971 | 0,69026 | 9330188P03R | 4,165453539 | 0,001530882 |
| Sycp3 | 1,68228 | 0 | 0 | 0 | 0 | 0 | 0 | 0 | 0 | 0 | Sycp3 | 2,855870998 | 0,001608271 |
| Ppm1k | 1,51831 | 3,16791 | 9,39129 | 9,31713 | 15,3643 | 21,3142 | 20,4062 | 53,2265 | 61,9835 | 49,9446 | Ppm1k | 2,012310991 | 0,001629037 |
| Pbld2 | 0,152605 | 0,150542 | 0 | 0,307449 | 0,62765 | 1,97311 | 1,56902 | 0,485099 | 1,74345 | 1,16321 | Pbld2 | 3,300525687 | 0,001634208 |
| Mettl15 | 0,545375 | 3,04866 | 2,88289 | 3,47913 | 2,99219 | 13,3079 | 11,9656 | 9,73472 | 10,4038 | 7,73306 | Mettl15 | 2,150527277 | 0,001649023 |
| Gstm7 | 0,839345 | 3,3119 | 15,8779 | 5,35317 | 21,3524 | 31,1624 | 29,3562 | 55,5687 | 74,7813 | 57,7093 | Gstm7 | 2,270204406 | 0,001656978 |
| Osmr | 0,263508 | 0,454915 | 0,329935 | 0,265498 | 3,67593 | 0,676387 | 0,58532 | 0,119905 | 0,902581 | 0,468536 | Osmr | 2,399250613 | 0,001700344 |
| Fam71e1 | 0,481922 | 0,475372 | 0,402134 | 0,485064 | 1,78666 | 2,44329 | 2,23338 | 1,30306 | 1,15784 | 1,10814 | Fam71e1 | 2,297196288 | 0,00171247 |
| Slc22a4 | 0 | 0,073465 | 0,124318 | 0 | 0,50694 | 1,47065 | 1,48555 | 3,02138 | 4,06125 | 4,291 | Slc22a4 | 3,565784636 | 0,001718796 |
| Xkrx | 0,0419882 | 0,0282416 | 0,0955831 | 0,1269 | 0,519382 | 0,721529 | 0,702377 | 0,936868 | 1,07479 | 0,64906 | Xkrx | 3,691575318 | 0,001737202 |
| Cyp39a1 | 0,434652 | 3,63622 | 7,38408 | 2,54639 | 10,7607 | 13,2414 | 12,9329 | 3,87162 | 3,22823 | 3,37569 | Cyp39a1 | 2,050943385 | 0,001773906 |
| 4930520O04R | 0 | 0,684905 | 0,289716 | 0,34954 | 1,99097 | 2,72916 | 1,17412 | 0,627749 | 1,18117 | 0,226893 | 4930520O04R | 2,836418131 | 0,001785569 |
| Myot | 0 | 0 | 0,38289 | 0 | 0,693975 | 1,57399 | 1,78338 | 35,6915 | 23,5844 | 38,4636 | Myot | 3,398034459 | 0,00180976 |
| Mrap | 0 | 6,97625 | 9,5084 | 6,72435 | 17,5659 | 29,2053 | 26,9 | 10,6456 | 10,4579 | 12,219 | Mrap | 2,305977476 | 0,001812637 |
| Slc47a1 | 0,251776 | 10,3075 | 9,98227 | 8,37011 | 13,7951 | 50,6606 | 45,8952 | 33,6545 | 40,3777 | 37,786 | Slc47a1 | 2,518432766 | 0,001838672 |
| Mir1839 | 0 | 0 | 0 | 0 | 0 | 0 | 0 | 7907,27 | 0 | 0 | Mir1839 | 3,080328376 | 0,00184416 |
| Rab40b | 0,184661 | 0,364327 | 2,00364 | 0,372002 | 2,09625 | 4,08356 | 5,89799 | 4,69248 | 3,10882 | 2,99847 | Rab40b | 2,565588705 | 0,00184865 |
| C1qtnf4 | 0 | 2,87713 | 3,9832 | 3,20423 | 9,34306 | 9,95126 | 11,9896 | 5,7644 | 5,83298 | 5,87341 | C1qtnf4 | 2,183676511 | 0,001853933 |
| Ephx2 | 0 | 3,90491 | 13,904 | 5,15033 | 37,2004 | 147,943 | 157,68 | 229,574 | 258,289 | 232,343 | Ephx2 | 3,660799928 | 0,001886244 |
| Aldh1b1 | 1,02159 | 5,18289 | 35,204 | 29,697 | 92,5932 | 223,039 | 226,301 | 17,8968 | 17,3524 | 17,6522 | Aldh1b1 | 3,151955595 | 0,001921797 |
| Sync | 0 | 0 | 0 | 0 | 0 | 0 | 0 | 22,2657 | 28,5194 | 17,4683 | Sync | 2,272478959 | 0,001956244 |
| 5031414D18R | 0 | 0,369958 | 0,104342 | 0 | 0,637966 | 0,940775 | 0,920294 | 0,681655 | 0,419313 | 0,567064 | 5031414D18R | 2,805731111 | 0,001993566 |
| Abi3bp | 0 | 0,287402 | 0,138505 | 0,167177 | 4,15605 | 0,311804 | 0,61011 | 0,226423 | 1,02691 | 0,482965 | Abi3bp | 3,160097182 | 0,002023995 |
| Klhl32 | 0,587999 | 0 | 0,981568 | 0,88848 | 1,63116 | 2,88703 | 2,02314 | 1,78429 | 3,34425 | 2,0012 | Klhl32 | 2,32107945 | 0,00205489 |
| 4930599N23R | 0 | 0 | 0 | 0 | 0 | 0 | 0 | 0 | 0 | 0 | 4930599N23R | 4,070969055 | 0,002111021 |
| Abcc9 | 0,468328 | 3,48607 | 14,6887 | 18,71 | 57,1178 | 111,957 | 107,753 | 81,9169 | 98,236 | 87,0789 | Abcc9 | 3,091106701 | 0,002135792 |
| A330102I10Ri | 0 | 0 | 0 | 0 | 1,11165 | 1,93484 | 3,84131 | 0,654872 | 0,413164 | 0,197107 | A330102I10Ri | 4,072701614 | 0,002141814 |
| Igtp | 0,0842491 | 0,831099 | 0,281275 | 1,69728 | 2,29472 | 4,23225 | 5,40588 | 4,20688 | 9,03978 | 6,26162 | Igtp | 2,753082308 | 0,002278302 |
| Ifnlr1 | 0 | 0,474631 | 0,535471 | 0,969485 | 2,15824 | 2,00894 | 2,03973 | 1,89675 | 1,79699 | 1,27095 | Ifnlr1 | 2,211047676 | 0,002340827 |
| Park2 | 0 | 0,100584 | 0,510639 | 0,924494 | 1,67604 | 3,19504 | 2,03199 | 3,89395 | 3,75654 | 2,77265 | Park2 | 2,533948655 | 0,002341196 |
| Tigd4 | 0 | 0 | 0 | 0 | 0,398901 | 0,564483 | 0,230146 | 0,0853492 | 0 | 0 | Tigd4 | 4,038276236 | 0,002387953 |
| Gm9899 | 0 | 0 | 0 | 0 | 0 | 0 | 0 | 0 | 0 | 0 | Gm9899 | 2,415604841 | 0,002393684 |
| Mir568 | 0 | 0 | 0 | 0 | 0 | 0 | 1827,82 | 0 | 0 | 0 | Mir568 | 3,614783044 | 0,002396553 |
| Jade2 | 0,310258 | 0,204047 | 0,733785 | 0,625224 | 4,04155 | 1,10024 | 0,791918 | 2,73035 | 3,67731 | 3,62563 | Jade2 | 2,200687613 | 0,002433493 |
| Acot3 | 0,359791 | 0 | 0,0890752 | 0,0322521 | 0,5223 | 1,54191 | 1,96066 | 0,789411 | 1,14355 | 0,537406 | Acot3 | 3,475724599 | 0,002436706 |
| C1qa | 0 | 0 | 2,05617 | 0,354389 | 18,831 | 3,5554 | 2,16924 | 0,636395 | 0,598971 | 0,34513 | C1qa | 3,282005452 | 0,002472963 |
| Myh14 | 0,975018 | 0,242997 | 0,534593 | 1,14168 | 1,04916 | 3,854 | 3,62228 | 40,4786 | 40,2783 | 38,4863 | Myh14 | 2,07983351 | 0,002731189 |
| Enpp4 | 0,140195 | 1,72878 | 2,16494 | 2,9663 | 4,14998 | 7,00498 | 7,96848 | 11,1628 | 13,3204 | 11,6726 | Enpp4 | 2,051786524 | 0,002863083 |
| 1700120C14R | 0 | 0 | 0 | 0 | 0 | 0 | 0 | 10,7712 | 0 | 0 | 1700120C14R | 2,906511222 | 0,00286555 |
| Sstr3 | 0 | 0,0808747 | 0 | 0 | 0,510999 | 0,287558 | 0,376886 | 1,26787 | 0,795292 | 1,64422 | Sstr3 | 3,547521463 | 0,002888802 |
| Tnip3 | 0 | 0 | 0 | 0 | 0,194149 | 0,364851 | 0,393789 | 0,311654 | 0,888411 | 0,692775 | Tnip3 | 3,973194205 | 0,002917704 |
| Vdr | 0,36769 | 0 | 0,122767 | 0,815011 | 1,77073 | 1,78672 | 1,73512 | 0,401476 | 0,393379 | 0,261588 | Vdr | 2,468457935 | 0,003178825 |
| Klhl38 | 0 | 0,4514 | 3,94661 | 2,15106 | 8,13371 | 8,38626 | 7,44326 | 30,3382 | 27,8811 | 23,203 | Klhl38 | 2,515371857 | 0,003178992 |
| 1700020L24Ri | 0 | 5,04861 | 5,33867 | 0,858696 | 9,47871 | 7,98766 | 10,2667 | 1,53962 | 0,875165 | 0,41941 | 1700020L24Ri | 2,202271581 | 0,003217284 |
| Best3 | 0 | 0,119967 | 0,304518 | 0 | 0,724018 | 0,823596 | 0,932513 | 3,31588 | 3,71121 | 4,09742 | Best3 | 2,582209663 | 0,003382645 |
| Spata33 | 5,0559 | 0 | 0 | 0 | 4,35843 | 13,2562 | 13,5611 | 10,8099 | 16,4917 | 17,2461 | Spata33 | 2,491233159 | 0,003457945 |
| Tnni3k | 0,107701 | 6,79977 | 15,9119 | 10,8503 | 50,5272 | 109,744 | 104,632 | 84,7005 | 105,047 | 87,7307 | Tnni3k | 3,100438475 | 0,003495433 |
| D330023K18R | 4,12621 | 0 | 4,30399 | 2,07507 | 8,89457 | 16,7126 | 13,8385 | 7,34886 | 5,49224 | 10,3548 | D330023K18R | 2,396232013 | 0,003562682 |
| Gm19461 | 0 | 1,45147 | 0,350894 | 0,317639 | 1,01277 | 2,52948 | 2,90023 | 5,25564 | 6,5134 | 5,13806 | Gm19461 | 2,133487695 | 0,003648223 |
| 1500017E21R | 0 | 0 | 0 | 1,47005 | 0,795465 | 3,90781 | 4,84095 | 5,42877 | 5,07154 | 5,4839 | 1500017E21R | 2,617235998 | 0,00366036 |
| Trim12a | 0 | 1,6068 | 0,388415 | 0 | 3,20534 | 3,54266 | 3,60042 | 0,738301 | 1,45392 | 2,6876 | Trim12a | 2,66012452 | 0,003706723 |
| Clic3 | 0 | 0 | 0 | 1,58023 | 1,94169 | 1,59405 | 0,161838 | 0 | 0,180123 | 0,688807 | Clic3 | 2,296525309 | 0,00378017 |
| Acss3 | 0,636277 | 0 | 2,38984 | 1,4421 | 1,62448 | 8,74757 | 6,35977 | 0,433347 | 0,909026 | 0,928942 | Acss3 | 2,436070045 | 0,003879775 |
| Tuba3b | 0 | 0 | 0 | 0 | 2,04878 | 0,718471 | 0,727594 | 0,068137 | 0,158559 | 0,207768 | Tuba3b | 3,89009873 | 0,003918349 |
| T2 | 0,396267 | 0 | 0,177181 | 1,19739 | 2,02471 | 2,24141 | 2,40419 | 0,539276 | 1,06832 | 0,128755 | T2 | 2,433366922 | 0,003931925 |
| Lzts1 | 0,384495 | 0,189647 | 2,08594 | 1,35547 | 3,3831 | 3,28585 | 2,77537 | 0,872214 | 0,971433 | 0,99922 | Lzts1 | 2,079325769 | 0,00396891 |
| Itgad | 0 | 0 | 0 | 0 | 0 | 0 | 0 | 0 | 0 | 0 | Itgad | 2,84131953 | 0,004022095 |
| Acot6 | 0,306989 | 0,187046 | 0,47477 | 0,986753 | 2,44604 | 1,77446 | 1,57449 | 1,49693 | 2,36826 | 1,83739 | Acot6 | 2,388410547 | 0,004063954 |
| Gpr133 | 0,0615076 | 0 | 0 | 0,185918 | 0,888453 | 0,338823 | 0,433391 | 0,0559819 | 0,61645 | 0,457267 | Gpr133 | 2,774816824 | 0,004065706 |
| Catsper2 | 0,59959 | 0,147871 | 1,87671 | 1,20798 | 2,67853 | 4,96716 | 5,38236 | 3,40373 | 2,82013 | 2,96552 | Catsper2 | 2,109763551 | 0,004095898 |
| Scn10a | 0 | 0,153339 | 0,320957 | 1,20348 | 2,39887 | 3,24196 | 2,71875 | 0,841634 | 1,17975 | 1,0419 | Scn10a | 2,843010677 | 0,004157436 |
| Mir1b | 0 | 0 | 0 | 0 | 0 | 0 | 0 | 0 | 0 | 0 | Mir1b | 3,853034893 | 0,004282504 |
| Slc26a7 | 0 | 0 | 0 | 0 | 0,240373 | 0,496356 | 0,520155 | 0,128334 | 0,0474332 | 0,0916006 | Slc26a7 | 3,844702447 | 0,004391153 |
| Itk | 0 | 0 | 0 | 0 | 0,239675 | 0,485987 | 0,0913261 | 0,40483 | 0,284572 | 1,16124 | Itk | 3,86139984 | 0,004425825 |
| Atp5l | 35,2981 | 37,7846 | 52,7617 | 79,6227 | 96,7457 | 163,668 | 91,676 | 161,297 | 173,002 | 140,514 | Atp5l | 2,727785232 | 0,004425825 |
| C920021L13R | 0 | 0,48918 | 0,617251 | 0 | 0,469952 | 0,585069 | 0,865302 | 0,182962 | 0,228723 | 0,15974 | C920021L13R | 2,076887834 | 0,004430439 |
| C130083M11I | 0,492401 | 0,161915 | 0 | 0 | 0,535499 | 0,906862 | 0,856571 | 1,04321 | 1,21423 | 0,905441 | C130083M11I | 2,368247244 | 0,004480424 |
| Zfp14 | 0,0947803 | 0,374001 | 1,10759 | 0,286468 | 0,77891 | 2,70726 | 2,32542 | 1,89636 | 2,57126 | 1,4435 | Zfp14 | 2,208534303 | 0,004489199 |
| Tmem171 | 0 | 1,92098 | 1,08345 | 0,980418 | 2,03106 | 7,78005 | 4,99789 | 5,87196 | 5,62365 | 2,2194 | Tmem171 | 2,382175815 | 0,00451784 |
| Smim6 | 0 | 0 | 0 | 0 | 0,457654 | 1,32073 | 1,36305 | 0,725491 | 0 | 0,131553 | Smim6 | 3,830558064 | 0,004613933 |
| Kcnd2 | 0,0702758 | 0,0693271 | 0,586607 | 0,920474 | 1,19441 | 2,34094 | 2,56243 | 11,319 | 21,0711 | 12,1341 | Kcnd2 | 2,33032051 | 0,004838491 |
| Lrp2bp | 0,189975 | 0,187409 | 0,0792865 | 0 | 0,215342 | 1,04753 | 0,960938 | 0,604705 | 0,699616 | 0,522616 | Lrp2bp | 2,749555976 | 0,004974093 |
| Sox30 | 0 | 0 | 0 | 0 | 0,465667 | 0,274624 | 0,201512 | 0,298718 | 0,146805 | 0,106399 | Sox30 | 3,797482123 | 0,005041698 |
| 2900052N01R | 0 | 0 | 0 | 0 | 0,546492 | 0,225658 | 0,31534 | 0,583836 | 0,08622 | 0,0416395 | 2900052N01R | 3,804180544 | 0,005050538 |
| Tuba8 | 3,0062 | 0 | 0,836321 | 1,76605 | 2,49103 | 16,4624 | 16,101 | 195,982 | 240,602 | 196,781 | Tuba8 | 2,654273233 | 0,005230224 |
| Pcdhga1 | 0 | 0 | 0 | 0 | 0 | 0 | 0 | 0 | 0 | 0 | Pcdhga1 | 2,132450455 | 0,005235212 |
| B230119M05I | 0 | 0 | 0 | 0,402299 | 0,341425 | 0,505272 | 0,246515 | 0 | 0 | 0,130851 | B230119M05I | 2,838052553 | 0,005448477 |
| Toporsos | 0 | 0 | 0 | 0 | 0 | 42,0612 | 55,6475 | 29,7775 | 68,3418 | 57,4358 | Toporsos | 2,50994403 | 0,005910037 |
| Zfp536 | 0 | 0 | 0,242197 | 0,657764 | 1,84937 | 1,85323 | 1,80041 | 0,99007 | 1,40652 | 1,17283 | Zfp536 | 2,662637732 | 0,005925873 |
| Sh2d4a | 0 | 0,588539 | 4,48177 | 2,64451 | 6,241 | 11,911 | 11,9305 | 9,86929 | 12,6828 | 8,81289 | Sh2d4a | 2,459202645 | 0,006045252 |
| 4930552P12R | 0 | 0 | 0,222882 | 0 | 0,719325 | 0,60302 | 0,737307 | 0,242629 | 0,537702 | 0,692391 | 4930552P12R | 3,13939094 | 0,006060327 |

| Gene | d9-11 (1) | d9-11 (2) | d9-11(3) | d9-11(4) | newborn(1) | newborn(2) | newborn(3) | adult(1) | adult(2) | adult(3) | Gene | log2(fold) | FDR(*p* -value) |
|---|---|---|---|---|---|---|---|---|---|---|---|---|---|
| Tmem74b | 0,249899 | 0,246515 | 0,625695 | 1,51003 | 1,41993 | 4,40425 | 3,38129 | 3,39675 | 4,64719 | 3,01027 | Tmem74b | 2,249806221 | 0,006579284 |
| Asb5 | 0 | 1,10219 | 0 | 0 | 1,14224 | 3,48552 | 3,43405 | 12,7649 | 23,986 | 21,658 | Asb5 | 3,041451923 | 0,00681413 |
| Chrna2 | 0 | 0 | 0,076723 | 0 | 0,234415 | 0,391604 | 0,648049 | 2,25718 | 1,16912 | 1,16004 | Chrna2 | 3,346764695 | 0,00695534 |
| Epn3 | 0,0829656 | 2,61905 | 8,76043 | 7,02144 | 14,762 | 22,1797 | 24,4106 | 14,2646 | 13,1835 | 16,6402 | Epn3 | 2,287455669 | 0,00696589 |
| Clec4n | 0 | 0 | 0 | 0 | 1,26289 | 1,07858 | 0,708841 | 0 | 0 | 0 | Clec4n | 3,69344026 | 0,006970005 |
| Slc9a2 | 0 | 0 | 0,0775194 | 0,0935654 | 0,263169 | 0,512057 | 0,569381 | 2,78735 | 3,97923 | 2,22403 | Slc9a2 | 3,051688114 | 0,007118336 |
| Pbld1 | 0,0311936 | 0,0824838 | 0,294317 | 0,207401 | 0,775465 | 1,30671 | 1,21811 | 0,240064 | 0,415335 | 0,228959 | Pbld1 | 2,844209935 | 0,007430735 |
| Kcnn1 | 1,03399 | 0 | 2,39009 | 0,480814 | 2,63647 | 5,08116 | 5,48449 | 7,67034 | 7,04958 | 7,33213 | Kcnn1 | 2,336733872 | 0,007508539 |
| Abcc8 | 0 | 0,316909 | 1,93069 | 5,24348 | 14,2945 | 22,3902 | 23,5814 | 17,8343 | 21,5216 | 20,4718 | Abcc8 | 3,010511486 | 0,007592457 |
| Zfp939 | 0 | 0,681103 | 0,432207 | 0,347735 | 1,17544 | 1,08401 | 1,32494 | 1,41026 | 1,39378 | 1,1206 | Zfp939 | 2,343023324 | 0,007655788 |
| Scgb1c1 | 0 | 0 | 0 | 0 | 4,36254 | 9,25613 | 12,0664 | 24,3445 | 21,3611 | 16,6942 | Scgb1c1 | 3,65526831 | 0,007676766 |
| Dleu2 | 0,412105 | 0,203265 | 0 | 0 | 0,526478 | 1,19168 | 1,32947 | 3,36436 | 3,05685 | 4,28615 | Dleu2 | 2,010228958 | 0,008453112 |
| 6330410L21Ri | 0 | 0 | 0 | 0 | 0 | 0,748242 | 0,714907 | 0,573139 | 0 | 0 | 6330410L21Ri | 3,404836659 | 0,008726727 |
| Phospho1 | 0 | 0 | 5,68419 | 0 | 19,1643 | 32,3294 | 35,6598 | 80,3516 | 81,5593 | 95,337 | Phospho1 | 2,053141679 | 0,008986574 |
| Ifit1 | 0 | 0 | 0 | 0 | 0,677033 | 0,283754 | 0,289141 | 0,171326 | 0,548058 | 0,610848 | Ifit1 | 3,600303318 | 0,009181328 |
| 2200002J24Ri | 0 | 0 | 0 | 0 | 1,55484 | 2,97263 | 1,9272 | 0 | 0,271136 | 0 | 2200002J24Ri | 3,590768569 | 0,009208375 |
| Rbp7 | 0 | 0 | 0 | 0 | 5,32773 | 1,44725 | 1,25627 | 0,723056 | 5,09748 | 4,02974 | Rbp7 | 3,58937523 | 0,00967593 |
| Cdsn | 0 | 0,121966 | 0 | 0 | 0,210317 | 0,527234 | 0,72055 | 0,337098 | 0,414675 | 0,20029 | Cdsn | 3,224245081 | 0,009864421 |
| Tnfsf13b | 0 | 0 | 0 | 0 | 0,233356 | 0,568383 | 0,631365 | 0 | 0,346419 | 0,601099 | Tnfsf13b | 3,569920338 | 0,009943712 |
| Angptl3 | 0 | 0 | 0 | 0 | 0,124619 | 0,496779 | 0,741805 | 1,1939 | 0,740481 | 1,21305 | Angptl3 | 3,571595201 | 0,0099496 |
| BB014433 | 0 | 0 | 0 | 0 | 0,223944 | 0,396655 | 0,181764 | 0 | 0,0664632 | 0 | BB014433 | 3,53358247 | 0,010833175 |
| Itgb6 | 0,133109 | 0,393937 | 0,166664 | 0,469399 | 0,23951 | 1,66145 | 1,63413 | 3,84783 | 13,8019 | 7,98002 | Itgb6 | 2,193035841 | 0,011234479 |
| Hpdl | 0,439497 | 0,650323 | 0,366812 | 0,221324 | 1,43519 | 1,93473 | 1,48576 | 3,58684 | 4,22693 | 5,0022 | Hpdl | 2,036206816 | 0,011610823 |
| Gpr52 | 0 | 0 | 0 | 0 | 0,897307 | 0,70777 | 0,431834 | 0 | 0 | 0 | Gpr52 | 3,500867062 | 0,011783129 |
| Zfp759 | 0,0931833 | 0,275775 | 0,155561 | 0,0938806 | 0,422492 | 0,817389 | 0,77126 | 1,01699 | 1,62204 | 1,62843 | Zfp759 | 2,231260377 | 0,01184723 |
| Dct | 0 | 0 | 1,24136 | 0,149817 | 2,10915 | 2,53874 | 2,69288 | 2,02632 | 1,39893 | 2,074 | Dct | 2,615741732 | 0,011982304 |
| Tdrd6 | 0 | 0 | 0 | 0 | 0,214579 | 0,0892163 | 0,0955017 | 0 | 0 | 0 | Tdrd6 | 3,499656226 | 0,011998864 |
| Ly6c1 | 0 | 0,475372 | 0,402134 | 0,510088 | 10,1416 | 2,13788 | 1,53855 | 14,556 | 33,4634 | 31,8273 | Ly6c1 | 2,890481569 | 0,012011655 |
| Lcn2 | 0 | 0 | 0 | 0 | 3,14094 | 0,364153 | 1,03562 | 3,88484 | 6,7376 | 2,51664 | Lcn2 | 3,522946037 | 0,012099412 |
| Tmod4 | 0 | 0 | 0,496203 | 0 | 0,591664 | 1,34913 | 2,19614 | 21,5273 | 22,9538 | 17,453 | Tmod4 | 2,586909603 | 0,012154915 |
| Gm2011 | 1,84775 | 1,21507 | 3,23037 | 2,56799 | 1,5318 | 10,408 | 7,2976 | 10,14 | 10,965 | 11,9814 | Gm2011 | 3,484728231 | 0,012341805 |
| Grm1 | 0 | 0 | 0 | 0 | 0,0257584 | 0,273211 | 0,20895 | 10,1419 | 8,81054 | 9,19981 | Grm1 | 3,511946432 | 0,012394471 |
| Gm10560 | 0 | 0,479934 | 0 | 0,489714 | 1,66518 | 1,48024 | 1,50335 | 0 | 0,167044 | 0 | Gm10560 | 2,33416787 | 0,012868867 |
| Slc25a35 | 0 | 0 | 0 | 0 | 0 | 1,57063 | 1,97824 | 4,46315 | 5,73018 | 6,20378 | Slc25a35 | 2,26427692 | 0,01296781 |
| Ppp3cc | 0 | 6,937 | 7,63012 | 3,8958 | 12,3216 | 19,3449 | 20,3779 | 24,5756 | 25,853 | 26,3107 | Ppp3cc | 2,059416354 | 0,012978689 |
| Ctsk | 0 | 4,83541 | 1,30171 | 0,224402 | 16,0709 | 4,8205 | 3,69786 | 0,202028 | 1,20326 | 1,59419 | Ctsk | 2,239451571 | 0,013000805 |
| 4833418N02R | 0 | 0,228669 | 0,193468 | 0 | 0,921713 | 0,408322 | 0,855093 | 0,210144 | 0,391431 | 0,678743 | 4833418N02R | 2,46196133 | 0,013175377 |
| Thrb | 0 | 0,363447 | 0,219665 | 0,212122 | 0,372612 | 0,869556 | 0,905997 | 5,98912 | 6,37903 | 7,38076 | Thrb | 2,03656623 | 0,013175377 |
| Inpp5j | 0 | 0,460895 | 4,05577 | 2,3535 | 7,06985 | 19,9527 | 20,4377 | 12,4925 | 13,8883 | 17,1762 | Inpp5j | 2,85579622 | 0,013391046 |
| Irf7 | 0,590043 | 1,16414 | 0 | 0 | 5,18047 | 1,95614 | 1,36607 | 1,60707 | 2,43385 | 2,71168 | Irf7 | 2,581174088 | 0,013403471 |
| Ifit2 | 0 | 0,241259 | 1,42897 | 1,23199 | 4,75784 | 2,34876 | 2,1986 | 0,964105 | 1,7452 | 2,16234 | Ifit2 | 2,153871102 | 0,014113294 |
| Abca12 | 0,0375273 | 0,0740419 | 0,0313256 | 0,0756266 | 0,127495 | 0,33804 | 0,252804 | 0,820202 | 1,23928 | 0,884683 | Abca12 | 2,293136874 | 0,014120708 |
| Mthfs | 6,95045 | 7,22593 | 10,0336 | 9,55126 | 10,8558 | 12,7252 | 12,1268 | 8,12562 | 7,37137 | 8,31211 | Mthfs | 2,605753135 | 0,01424852 |
| Gpr135 | 0,263084 | 0 | 0,6587 | 0 | 1,42041 | 1,06021 | 1,86124 | 0 | 0,0890278 | 0,0856945 | Gpr135 | 2,045568393 | 0,014724141 |
| Zfp354b | 0 | 0,131825 | 0,334616 | 0,134618 | 0,530496 | 0,832677 | 0,498804 | 0,546395 | 0,747469 | 0,967317 | Zfp354b | 3,073319622 | 0,015235176 |
| 2810055G20R | 0 | 0,102298 | 1,03869 | 0,626833 | 0,940499 | 1,28687 | 1,71695 | 1,32006 | 1,05946 | 1,51077 | 2810055G20R | 2,124314166 | 0,015660955 |
| Mmaa | 0,231731 | 1,6002 | 3,86847 | 2,91815 | 3,27359 | 8,5866 | 8,43945 | 15,1664 | 17,1099 | 15,9874 | Mmaa | 2,045580378 | 0,01567092 |
| Tssk1 | 0 | 0 | 0 | 0 | 0,350357 | 0,496765 | 0,606763 | 0,447058 | 0,583666 | 0,883223 | Tssk1 | 3,382209417 | 0,015958491 |
| Cyp4b1 | 0,183991 | 0,181502 | 0,767833 | 0 | 2,06251 | 1,31769 | 1,1301 | 6,26186 | 10,9235 | 6,1941 | Cyp4b1 | 2,771126863 | 0,016010756 |
| Adc | 0 | 0,176589 | 1,34469 | 0,901556 | 5,02898 | 2,24883 | 1,20933 | 0,324962 | 0,843526 | 0,406851 | Adc | 2,065177193 | 0,01604056 |
| Abhd3 | 0 | 1,79763 | 0,152095 | 0 | 1,60312 | 3,47998 | 3,69344 | 3,96934 | 2,69928 | 3,90545 | Abhd3 | 2,265466422 | 0,016181609 |
| Tmie | 0 | 0,135211 | 0 | 0,276148 | 0,427548 | 1,16944 | 1,09339 | 1,12077 | 0,966214 | 0,799758 | Tmie | 2,57317183 | 0,016439669 |
| Ceacam15 | 0 | 0 | 0 | 0 | 0,515692 | 0,58766 | 0,797447 | 0 | 0,0877162 | 0,197028 | Ceacam15 | 3,357453757 | 0,01696621 |
| 2310007L24Ri | 0 | 0 | 0 | 0 | 0,759996 | 0,675998 | 0,823509 | 0,958842 | 0,733546 | 0,701116 | 2310007L24Ri | 2,746207588 | 0,017947186 |
| Lgi3 | 0 | 0 | 1,3709 | 0,620488 | 1,48372 | 3,0108 | 3,21023 | 1,40006 | 1,47844 | 1,22956 | Lgi3 | 2,407559604 | 0,018072993 |
| AI427809 | 0 | 0 | 0,139863 | 0 | 0,713139 | 0,561269 | 0,41156 | 0,304276 | 0,0563602 | 0,0543834 | AI427809 | 3,009720956 | 0,018148854 |
| Elfn2 | 0 | 0,0646239 | 0 | 0,0660029 | 0,222659 | 0,262475 | 0,301054 | 0,0596175 | 0,197013 | 0,0847101 | Elfn2 | 2,727993507 | 0,018471921 |
| Slc44a3 | 0 | 0,128111 | 0,433584 | 0,130825 | 0,368225 | 1,20554 | 0,916304 | 0,11801 | 0,0871479 | 0 | Slc44a3 | 2,170339859 | 0,018613355 |
| Pcdhb16 | 0 | 0 | 0 | 0 | 0,426283 | 0,0904458 | 0,0553306 | 0 | 0,14078 | 0,0437831 | Pcdhb16 | 3,339607468 | 0,018684747 |
| B430010I23Ri | 0 | 0 | 0 | 0 | 0 | 0 | 0 | 0 | 0 | 0 | B430010I23Ri | 3,011449998 | 0,018877313 |
| Sds | 0 | 0 | 0 | 0 | 0 | 0 | 0 | 0 | 0 | 0 | Sds | 2,060376322 | 0,019068794 |
| Pipox | 0 | 0,385631 | 1,46824 | 0,196873 | 1,49798 | 2,80059 | 2,34159 | 0,709376 | 0,526862 | 0,127005 | Pipox | 2,094978989 | 0,019111877 |
| Sspn | 0 | 5,58561 | 19,7532 | 12,4333 | 74,7407 | 125,365 | 126,531 | 105,088 | 126,872 | 127,273 | Sspn | 2,936134716 | 0,019635472 |
| Samd9l | 0 | 0 | 0 | 0,0599602 | 0,85957 | 0,1341 | 0,0364607 | 0,216665 | 0,51684 | 0,461617 | Samd9l | 3,126797865 | 0,019658261 |
| A730056A06R | 0 | 0 | 0 | 0 | 0 | 0 | 0 | 0 | 0 | 0 | A730056A06R | 3,300823413 | 0,020060388 |
| Pcdhb20 | 0 | 0,0943826 | 0,35937 | 0,19278 | 0,542257 | 0,823317 | 0,674599 | 1,43566 | 0,752756 | 1,16894 | Pcdhb20 | 2,152964083 | 0,020247757 |
| Blk | 0 | 0 | 0 | 0 | 0,0764244 | 0,101442 | 0,0826873 | 0 | 0,0452289 | 0,0436791 | Blk | 2,320529254 | 0,020251896 |
| Tnfrsf18 | 0 | 0 | 0,643291 | 0 | 0,296152 | 0,389713 | 0,4754 | 0 | 0 | 0 | Tnfrsf18 | 2,051590498 | 0,021078156 |
| Gm15408 | 0,261951 | 0 | 0,437243 | 0 | 1,11652 | 1,18756 | 1,12802 | 1,66113 | 0,975018 | 1,87708 | Gm15408 | 2,545120314 | 0,021122973 |
| Fuom | 0 | 1,32496 | 10,0491 | 4,04239 | 9,82174 | 17,4365 | 16,0654 | 10,286 | 13,5573 | 10,6061 | Fuom | 2,107165892 | 0,021329954 |
| Slc4a10 | 0 | 0 | 0 | 0 | 0,149572 | 0,0881509 | 0,16178 | 0 | 0 | 0 | Slc4a10 | 3,257985977 | 0,021501445 |
| Cilp | 0 | 0 | 0 | 0 | 1,24816 | 0,116177 | 0 | 0 | 0,0258481 | 0,0249989 | Cilp | 3,32546981 | 0,021505066 |
| 9330179D12R | 0 | 0 | 0,929848 | 0 | 0,942109 | 1,53655 | 1,98462 | 0,508564 | 0,951362 | 0,829563 | 9330179D12R | 2,451403874 | 0,022277328 |
| Cabyr | 0 | 0,499924 | 0,389309 | 0,282786 | 0,904864 | 0,240291 | 0,864994 | 0,254282 | 0,235436 | 0,364175 | Cabyr | 2,262499423 | 0,022383703 |
| Ly6a | 0 | 0 | 0 | 0 | 3,28316 | 0,625447 | 0,508519 | 5,58093 | 16,8908 | 13,2759 | Ly6a | 3,255011395 | 0,022497204 |
| A330093E20R | 0 | 0 | 0 | 0 | 0 | 0 | 0 | 0 | 0 | 0 | A330093E20R | 3,239328578 | 0,022596428 |
| C330013E15R | 0 | 0,454491 | 0,76895 | 0,463779 | 1,83893 | 1,28397 | 2,41894 | 1,66191 | 1,10545 | 2,57087 | C330013E15R | 2,321985184 | 0,023286694 |
| 2310061J03Ri | 0 | 0 | 0 | 0 | 5,33093 | 6,42969 | 5,81416 | 3,07873 | 4,29203 | 3,73937 | 2310061J03Ri | 2,026036362 | 0,026739485 |
| Crhr2 | 0 | 0,526607 | 0,850111 | 5,56191 | 10,476 | 15,2614 | 15,6361 | 18,493 | 24,8637 | 24,0882 | Crhr2 | 2,652275023 | 0,026750773 |
| Chrna10 | 0 | 0 | 0 | 0 | 0,112735 | 0,33252 | 0,365916 | 1,32461 | 2,35711 | 2,31942 | Chrna10 | 3,150905358 | 0,028014469 |
| Tmem117 | 0 | 0,23413 | 0,594305 | 1,31504 | 1,48022 | 2,35129 | 3,34844 | 9,38408 | 11,5652 | 10,6225 | Tmem117 | 2,110047955 | 0,028375525 |
| Kcne4 | 0 | 0,133645 | 0 | 0,136476 | 2,15135 | 0,101986 | 0,166261 | 0,369277 | 0,318309 | 0,0878276 | Kcne4 | 2,824220627 | 0,029220411 |
| Gm8234 | 0,597355 | 0,0841825 | 2,70659 | 0,257876 | 0,62953 | 1,07179 | 0,602609 | 0,929664 | 0,507167 | 0,747859 | Gm8234 | 3,123036838 | 0,029250215 |
| Ifi27 | 0 | 0 | 4,15183 | 0 | 3,6544 | 5,88829 | 5,62712 | 12,8444 | 11,0406 | 10,8376 | Ifi27 | 2,532239268 | 0,029256955 |
| Nipal2 | 0 | 0,249552 | 0 | 0,254866 | 0,406153 | 0,697217 | 0,723642 | 0,153413 | 0,197478 | 0,0545588 | Nipal2 | 2,071333655 | 0,029339545 |
| 4933432I09Ri | 0 | 0 | 0 | 0 | 0,45248 | 1,27007 | 0,816333 | 0,721174 | 0,359271 | 0,1729 | 4933432I09Ri | 2,853173157 | 0,029724336 |
| Timp4 | 0,507724 | 0 | 0 | 0 | 4,50792 | 0,894922 | 0,546506 | 3,91034 | 8,67147 | 8,59678 | Timp4 | 2,587475084 | 0,029868418 |
| Oas2 | 0 | 0 | 0 | 0 | 0,638158 | 0,0627038 | 0,0255675 | 0,0758874 | 0,362801 | 0,404801 | Oas2 | 3,141289673 | 0,030088191 |
| Gpr65 | 0 | 0 | 0 | 0 | 1,16287 | 0,237538 | 0,0483953 | 0 | 0 | 0 | Gpr65 | 3,136606003 | 0,030089466 |
| Cd28 | 0 | 0,0732 | 0,309688 | 0 | 0,483468 | 0,483248 | 0,545707 | 0,877703 | 1,43893 | 1,58373 | Cd28 | 2,292799112 | 0,030398176 |
| 1700073E17R | 0 | 0,0684885 | 0 | 0 | 0,196658 | 0,243428 | 0,212722 | 0 | 0 | 0 | 1700073E17R | 2,805520496 | 0,030705626 |
| Clec1a | 0,199442 | 0 | 0 | 0 | 1,35655 | 0,249972 | 0,18344 | 0,272044 | 0,834887 | 1,09752 | Clec1a | 2,768239707 | 0,03079166 |
| Mlana | 0 | 0 | 1,03817 | 0 | 10,0291 | 0,64229 | 1,55619 | 0 | 0,446311 | 0 | Mlana | 2,920425487 | 0,030986764 |
| Zpbp | 0,0880067 | 0,0868183 | 0,209991 | 0 | 0,166316 | 0,57382 | 0,591137 | 0,200127 | 0,47693 | 0,313228 | Zpbp | 2,059201306 | 0,031293098 |
| A730020M07l | 0 | 0,388573 | 0 | 0,542595 | 1,28628 | 0,425448 | 0,390116 | 2,55268 | 1,5653 | 1,54291 | A730020M07l | 2,550815364 | 0,031306127 |
| 4930506C21R | 0,879431 | 0 | 0 | 0 | 0 | 0 | 0 | 0 | 0 | 0 | 4930506C21R | 2,251749697 | 0,031599758 |
| Rd3 | 0 | 0 | 0,0740383 | 0 | 0,442957 | 0,32813 | 0,16313 | 0 | 0 | 0 | Rd3 | 2,59586673 | 0,031830407 |
| Ltf | 0 | 0 | 0,100581 | 0 | 1,40065 | 0,151123 | 0,0369598 | 0 | 0 | 0 | Ltf | 2,939264289 | 0,031871032 |
| Myh8 | 0 | 0,0885482 | 0,618669 | 0,0947472 | 0,559805 | 0,256905 | 0,164955 | 0,077824 | 0,0348191 | 0,0635278 | Myh8 | 2,713963203 | 0,032128963 |
| 4921504A21R | 0 | 0 | 0 | 0,331202 | 0,652971 | 0,619323 | 0,454217 | 1,79125 | 2,65358 | 2,45402 | 4921504A21R | 2,620420649 | 0,032645073 |
| Trim30d | 0 | 0,0979568 | 0 | 0 | 0,918334 | 0,0717825 | 0,0304433 | 0,120396 | 0,182891 | 0,107141 | Trim30d | 3,088863813 | 0,033439127 |
| Gpr115 | 0 | 0 | 0 | 0 | 0,187484 | 0,193495 | 0,20283 | 0 | 0,221654 | 0,107019 | Gpr115 | 3,057305762 | 0,033653747 |
| 0610038B21R | 0 | 0 | 0 | 0 | 0 | 0 | 0 | 0 | 0 | 0 | 0610038B21R | 2,388917204 | 0,034641852 |
| Cd33 | 0 | 0 | 0 | 0 | 0,78787 | 0,134478 | 0,164425 | 0 | 0,0449677 | 0,0434277 | Cd33 | 2,839982056 | 0,034712629 |
| Mcpt4 | 0 | 0 | 0 | 0 | 2,91659 | 0,49799 | 0,121487 | 0 | 0 | 0,128959 | Mcpt4 | 3,059087629 | 0,035322899 |
| Oas1b | 0 | 0 | 0,156937 | 0 | 0,800801 | 0,393793 | 0,288749 | 0,170632 | 0,379945 | 0,111964 | Oas1b | 2,780340422 | 0,035469666 |
| Mir7052 | 0 | 0 | 0 | 0 | 0 | 0 | 0 | 0 | 0 | 0 | Mir7052 | 3,038962501 | 0,035631315 |
| Phf11d | 0 | 0 | 0 | 0 | 0,486233 | 0,141684 | 0,0661508 | 0 | 0,764643 | 0,412584 | Phf11d | 3,032724716 | 0,036216848 |
| Agmat | 0 | 0 | 0,428029 | 0 | 0,437157 | 1,42064 | 0,63093 | 0,464666 | 0,346956 | 0,417529 | Agmat | 2,636033816 | 0,036525445 |
| 9430037G07R | 0,113067 | 0,152099 | 0,514781 | 1,19088 | 0,621481 | 1,23381 | 1,24171 | 0,607511 | 1,05583 | 1,21237 | 9430037G07R | 2,190616841 | 0,0380138 |
| A330069E16R | 0,902366 | 0 | 1,88247 | 0,0756931 | 2,37918 | 3,08526 | 3,27375 | 2,03469 | 1,77728 | 1,77639 | A330069E16R | 2,161030354 | 0,038180265 |
| Gm11837 | 0 | 0 | 1,49709 | 1,35445 | 5,36928 | 8,74611 | 6,64702 | 2,02282 | 2,74625 | 2,79573 | Gm11837 | 2,176953158 | 0,039018159 |
| Ccl3 | 0 | 0 | 1,33075 | 0 | 6,22098 | 1,61952 | 1,15088 | 0 | 0,549165 | 0 | Ccl3 | 2,596863949 | 0,039190325 |

| Gene | d9-11 (1) | d9-11 (2) | d9-11(3) | d9-11(4) | newborn(1) | newborn(2) | newborn(3) | adult(1) | adult(2) | adult(3) | Gene | log2(fold) | FDR(*p* -value) |
|---|---|---|---|---|---|---|---|---|---|---|---|---|---|
| AI429214 | 0 | 0 | 0,291612 | 0,175964 | 0,644396 | 0,78995 | 0,59003 | 0,158578 | 0,293911 | 0,510392 | AI429214 | 2,340247745 | 0,040064467 |
| E030011O05R | 0 | 0 | 0 | 0 | 0 | 0 | 0 | 0 | 0 | 0 | E030011O05R | 2,35388205 | 0,041383571 |
| Tnf | 0,219273 | 0 | 1,46407 | 0 | 5,72831 | 0,882522 | 1,61731 | 0 | 0 | 0,213912 | Tnf | 2,316809418 | 0,042203716 |
| Gm14858 | 0 | 0 | 0 | 0 | 0,448471 | 0,0992932 | 0,0606671 | 0 | 0 | 0 | Gm14858 | 2,950312449 | 0,04228926 |
| Aldh3b1 | 0 | 0 | 0,314553 | 0 | 1,07408 | 0,489366 | 0,578753 | 0 | 0,0634629 | 0,183613 | Aldh3b1 | 2,57479924 | 0,042402663 |
| Clec4a1 | 0 | 0 | 0 | 0 | 1,28912 | 0,342629 | 0,0697621 | 0,205776 | 0,0766267 | 0 | Clec4a1 | 2,953153046 | 0,043189307 |
| 1700067K01R | 0 | 0 | 0 | 0 | 0,499662 | 0,177355 | 0,324629 | 0 | 0,119511 | 0,114774 | 1700067K01R | 2,145090496 | 0,043438421 |
| Cd46 | 0 | 0 | 0 | 0 | 0,173827 | 0,462477 | 0,376385 | 0 | 0,207429 | 0,0997214 | Cd46 | 2,674886442 | 0,043516364 |
| Oscar | 0 | 0 | 0 | 0,19006 | 0,214457 | 0,617192 | 0,406137 | 0,171423 | 0,0636224 | 0,368144 | Oscar | 2,685979275 | 0,043518999 |
| Iyd | 0 | 0 | 0,384009 | 0 | 0,588031 | 0,752538 | 0,353581 | 0 | 0 | 0 | Iyd | 2,550155789 | 0,043587334 |
| Slfn10-ps | 0,231822 | 0,137215 | 0,554709 | 0,155705 | 0,249629 | 0,379582 | 0,487973 | 0 | 0,0310401 | 0,0250071 | Slfn10-ps | 2,441567035 | 0,043841394 |
| Ssu2 | 0 | 0 | 0 | 0 | 0,940394 | 0,138945 | 0,0848329 | 0 | 0,280142 | 0,359421 | Ssu2 | 2,942121683 | 0,043967286 |
| Slc9a3 | 0 | 0 | 0 | 0,404564 | 0,341635 | 0,974107 | 0,49284 | 0,121634 | 0,269567 | 1,34507 | Slc9a3 | 2,480817804 | 0,044352691 |
| 9330162012R | 0 | 0 | 0 | 0 | 0 | 0 | 0 | 0 | 0 | 0 | 9330162012R | 2,915355033 | 0,045171962 |
| AI507597 | 1,4122 | 0 | 2,35681 | 0 | 2,12615 | 8,60994 | 6,22864 | 10,5221 | 8,10607 | 11,2791 | AI507597 | 2,260826887 | 0,04584646 |
| Lrrc52 | 0 | 0 | 0 | 0 | 0,376555 | 0,167029 | 0,509642 | 2,09512 | 1,79957 | 1,18879 | Lrrc52 | 2,906034106 | 0,046262213 |
| Gpr113 | 0 | 0 | 0 | 0,106357 | 0,0897659 | 0,39702 | 0,194223 | 0 | 0 | 0,0683621 | Gpr113 | 2,65126002 | 0,046921006 |
| Enpp6 | 0 | 0 | 0 | 0 | 0,454199 | 0,0535702 | 0,0655157 | 0,0971306 | 0,107378 | 0 | Enpp6 | 2,906946276 | 0,04703976 |
| Pkib | 0,120867 | 0 | 0,151454 | 0 | 0,209835 | 0,272581 | 0,27792 | 0,0549746 | 0,0808152 | 0,0586451 | Pkib | 2,141823554 | 0,047138056 |
| Lilrb4 | 0 | 0 | 0,69492 | 0 | 2,06326 | 0,539473 | 0,401918 | 0 | 0,0803059 | 0,1547 | Lilrb4 | 2,539342707 | 0,047855431 |
| 9430083A17R | 0 | 0 | 0,366335 | 0,110538 | 0,342106 | 0,852874 | 0,706568 | 0,0997518 | 0,110304 | 0,213184 | 9430083A17R | 2,231500115 | 0,048025184 |
| Arl5c | 0 | 0 | 0,189003 | 0 | 0,643115 | 0,512788 | 0,765662 | 0 | 0,229359 | 0,14732 | Arl5c | 2,503701151 | 0,048165258 |
| Prnd | 0 | 0 | 0 | 0 | 0 | 0 | 0 | 0 | 0 | 0 | Prnd | 2,878530784 | 0,04862799 |
| Tm4sf4 | 0 | 0 | 0 | 0 | 0,366158 | 0,454337 | 0,554885 | 0 | 0,0871873 | 0 | Tm4sf4 | 2,865803073 | 0,049521831 |
| Otoa | 0 | 0 | 0 | 0,0893123 | 0,527492 | 0,199929 | 0,217379 | 0 | 0 | 0 | Otoa | 2,486519964 | 0,049613965 |
| Tslp | 0 | 0,327327 | 0,276922 | 0,668225 | 1,7906 | 1,58852 | 0,918361 | 0,299627 | 0,451321 | 0,649146 | Tslp | 2,102078719 | 0,050219888 |

**Genes up-regulated in cardiac progenitor cells**

| Gene | d9-11 (1) | d9-11 (2) | d9-11(3) | d9-11(4) | newborn(1) | newborn(2) | newborn(3) | adult(1) | adult(2) | adult(3) | Gene | log2(fold) | FDR(*p* -value) |
|---|---|---|---|---|---|---|---|---|---|---|---|---|---|
| Myl7 | 4021,69 | 5004,97 | 6775,69 | 3944,8 | 175,377 | 197,653 | 152,77 | 58,4526 | 70,1616 | 18,7405 | Myl7 | -4,45989084 | 2,20E-84 |
| Ddah1 | 26,0283 | 32,2873 | 34,4595 | 32,5623 | 2,61296 | 2,1164 | 2,00766 | 2,90073 | 2,97032 | 3,48554 | Ddah1 | -3,46833843 | 5,38E-76 |
| Slc2a3 | 132,035 | 100,176 | 71,1895 | 132,126 | 1,4051 | 1,16105 | 1,12127 | 0 | 0,0815863 | 0,157796 | Slc2a3 | -6,0764088 | 9,12E-74 |
| Sox11 | 32,5845 | 32,3635 | 41,3084 | 42,6976 | 2,60437 | 1,83953 | 1,75341 | 0,0672884 | 0,0123229 | 0,0596513 | Sox11 | -3,86119497 | 1,54E-63 |
| Slc16a3 | 281,912 | 182,569 | 193,359 | 268,061 | 6,28826 | 6,85245 | 6,8448 | 1,0196 | 0,721728 | 0,591263 | Slc16a3 | -4,7285371 | 5,50E-61 |
| Hba-x | 358,002 | 1743,67 | 2657,25 | 2097,95 | 0,683931 | 0 | 0 | 0 | 0 | 0 | Hba-x | -11,1167461 | 3,54E-49 |
| Dok4 | 73,3609 | 82,8465 | 72,0573 | 63,5964 | 9,27826 | 7,81879 | 7,19813 | 2,10336 | 4,39776 | 3,30674 | Dok4 | -2,8175231 | 3,55E-49 |
| Crb2 | 14,9755 | 10,8729 | 12,0062 | 15,7877 | 0 | 0 | 0 | 0 | 0 | 0 | Crb2 | -7,58568596 | 4,07E-48 |
| Prrx2 | 65,7636 | 58,9445 | 55,1609 | 56,8049 | 2,8491 | 1,6851 | 1,33678 | 0,301911 | 0 | 0 | Prrx2 | -4,60097989 | 1,12E-47 |
| Greb1 | 6,7549 | 6,87587 | 6,56678 | 7,98861 | 0,0902207 | 0,0199348 | 0,0121974 | 0,0725442 | 0,0531552 | 0,0854981 | Greb1 | -6,96307075 | 4,67E-46 |
| Cacna2d2 | 38,1688 | 40,5484 | 47,3198 | 59,2386 | 3,05898 | 2,51705 | 1,60136 | 0,993408 | 1,28565 | 0,723992 | Cacna2d2 | -3,94140141 | 6,75E-45 |
| Crabp1 | 112,471 | 118,132 | 101,233 | 135,727 | 0,59435 | 0,164776 | 0,200524 | 0 | 0 | 0,1713 | Crabp1 | -7,93348801 | 8,74E-45 |
| Igdcc3 | 16,6835 | 47,3748 | 38,1329 | 67,3362 | 0,206227 | 0,234538 | 0,318716 | 0,189029 | 0,03482 | 0 | Igdcc3 | -6,63408523 | 1,67E-43 |
| Sphkap | 12,9994 | 19,1645 | 23,6384 | 15,6301 | 1,3684 | 1,40309 | 1,34686 | 0 | 0,0806541 | 0,0156123 | Sphkap | -3,37862038 | 2,43E-43 |
| Rps6ka6 | 16,2806 | 22,512 | 13,3623 | 17,0211 | 0,443795 | 0,443558 | 0,751373 | 0,0619903 | 0 | 0,11012 | Rps6ka6 | -4,62716663 | 2,52E-43 |
| Dpysl5 | 13,8684 | 10,4952 | 13,0564 | 13,4415 | 0,125553 | 0,126858 | 0,310414 | 0,0576345 | 0,12695 | 0,0818815 | Dpysl5 | -5,72416668 | 3,88E-43 |
| Cadm1 | 39,1786 | 36,2001 | 26,476 | 35,8605 | 2,37855 | 1,68622 | 1,9154 | 0,468389 | 0,417965 | 0,35671 | Cadm1 | -3,69063609 | 6,40E-43 |
| Col2a1 | 378,476 | 270,869 | 264,837 | 282,2 | 0,841195 | 0,0646698 | 0,0593407 | 0,0587583 | 0,0215731 | 0 | Col2a1 | -8,95153735 | 7,69E-41 |
| Skp2 | 20,6358 | 28,5801 | 19,065 | 18,0246 | 1,26764 | 0,891913 | 1,12202 | 0,18487 | 0,069736 | 0,0673991 | Skp2 | -3,83565229 | 4,16E-40 |
| Rps6ka1 | 50,0025 | 41,1608 | 32,387 | 39,7206 | 2,09017 | 1,21271 | 1,01121 | 0,499658 | 0,706856 | 0,961068 | Rps6ka1 | -4,49511732 | 1,05E-38 |
| Krt8 | 108,948 | 65,3167 | 36,0691 | 60,4825 | 0,761511 | 0,457598 | 0,431667 | 0,260843 | 0,387334 | 0,217886 | Krt8 | -6,93099803 | 1,10E-38 |
| Cdc6 | 12,6202 | 12,3803 | 8,96388 | 9,58978 | 0,37946 | 0,423799 | 0,345648 | 0 | 0 | 0 | Cdc6 | -4,54630848 | 1,55E-38 |
| Hbb-bh1 | 278,347 | 1144,88 | 694,082 | 900,285 | 0 | 0 | 0 | 0 | 0 | 0 | Hbb-bh1 | -11,3629293 | 1,57E-38 |
| Lama1 | 27,9957 | 18,0034 | 8,27209 | 7,38191 | 0,0185174 | 0,0327293 | 0,0100136 | 0 | 0 | 0,0105598 | Lama1 | -8,55529355 | 1,62E-36 |
| Itga2b | 38,5503 | 29,9204 | 20,269 | 28,8434 | 0,669385 | 0,331538 | 0,376548 | 0,0859312 | 0,0316318 | 0 | Itga2b | -4,81866692 | 1,92E-36 |
| Gata5 | 53,0585 | 33,0785 | 34,1627 | 26,1313 | 0,782581 | 0,420255 | 0,151178 | 0,358736 | 0,165125 | 0,255382 | Gata5 | -5,92173077 | 2,51E-36 |
| Gata3 | 10,5544 | 13,6155 | 12,3675 | 9,71298 | 0,28762 | 0,0508812 | 0,0933453 | 0 | 0 | 0,032853 | Gata3 | -5,87800038 | 4,20E-36 |
| Hand1 | 32,4051 | 48,0396 | 55,6045 | 62,9105 | 3,49343 | 2,95671 | 2,11256 | 0 | 0 | 0 | Hand1 | -3,72597331 | 8,42E-35 |
| Kif5c | 3,72044 | 5,66394 | 5,40603 | 5,22969 | 0,234103 | 0,2644 | 0,309471 | 0,125461 | 0,30659 | 0,133539 | Kif5c | -3,73576433 | 1,46E-34 |
| Kif21b | 7,17598 | 8,19161 | 11,4383 | 9,19336 | 0,541742 | 0,350528 | 0,18833 | 0,0933641 | 0,0227923 | 0 | Kif21b | -4,184582 | 3,07E-34 |
| Rfc5 | 18,7737 | 21,0331 | 18,2154 | 25,1222 | 4,39673 | 4,82956 | 5,19078 | 0,54744 | 1,15258 | 1,53607 | Rfc5 | -2,36813676 | 7,71E-34 |
| Dusp9 | 50,3642 | 38,3206 | 18,4109 | 28,7636 | 0,312673 | 0,0614758 | 0,0751735 | 0,111373 | 0 | 0 | Dusp9 | -7,10297598 | 1,41E-33 |
| Cpne5 | 14,9584 | 10,995 | 17,0765 | 14,7016 | 0,872414 | 1,13872 | 0,988627 | 0 | 0,0980589 | 0,0948468 | Cpne5 | -3,48114567 | 1,52E-33 |
| Hmga2 | 169,706 | 121,191 | 78,3944 | 70,339 | 0,576907 | 0,0288751 | 0,023549 | 0 | 0,0128486 | 0,0248533 | Hmga2 | -8,11912726 | 1,53E-33 |
| Slc2a1 | 446,01 | 444,827 | 541,986 | 878,897 | 35,572 | 22,0749 | 18,827 | 15,2633 | 15,7794 | 11,9331 | Slc2a1 | -4,12283828 | 1,71E-33 |
| Marcksl1 | 505,53 | 476,489 | 392,846 | 390,725 | 76,5887 | 60,2138 | 60,7916 | 0,199412 | 1,00185 | 0,822389 | Marcksl1 | -2,53118017 | 1,16E-32 |
| Rnd2 | 31,9707 | 64,3578 | 53,8928 | 39,3875 | 4,47792 | 3,82294 | 2,42718 | 0,585757 | 0 | 0,215717 | Rnd2 | -3,91082661 | 1,34E-32 |
| Fndc3c1 | 27,4461 | 31,703 | 25,8361 | 22,8804 | 0,100016 | 0 | 0 | 0 | 0 | 0 | Fndc3c1 | -9,46324952 | 1,41E-32 |
| Krt18 | 117,842 | 93,6572 | 52,611 | 96,3921 | 3,50914 | 2,59179 | 3,36321 | 0 | 1,04445 | 0,586538 | Krt18 | -4,72683978 | 2,16E-32 |
| Cnn1 | 88,2845 | 116,601 | 181,263 | 116,665 | 12,1956 | 8,03297 | 6,49282 | 0,312156 | 0,636259 | 0,279034 | Cnn1 | -3,43249354 | 4,58E-32 |
| Edil3 | 4,99981 | 12,3895 | 7,9495 | 5,81728 | 0,134073 | 0,0745146 | 0,0364292 | 0 | 0 | 0 | Edil3 | -6,13519077 | 5,40E-32 |
| Col22a1 | 3,96278 | 8,73247 | 14,3484 | 7,46714 | 0,0291477 | 0,0515305 | 0,0315265 | 0,0468522 | 0,171823 | 0,166287 | Col22a1 | -7,06290384 | 4,04E-31 |
| Igf2bp1 | 83,911 | 67,0097 | 51,6933 | 56,5047 | 4,28293 | 1,91126 | 1,98524 | 0,067865 | 0,0248577 | 0,0120327 | Igf2bp1 | -4,2144256 | 4,78E-31 |
| Lef1 | 35,3826 | 24,1737 | 21,4364 | 22,7283 | 0,576383 | 0,310276 | 0,140667 | 0 | 0,0489471 | 0,140948 | Lef1 | -5,59342947 | 1,44E-30 |
| Msi1 | 64,5678 | 54,4662 | 52,7465 | 80,3101 | 3,69335 | 5,1666 | 4,55846 | 0,413702 | 0,539245 | 0,29687 | Msi1 | -3,39963623 | 2,16E-30 |
| Nnat | 105,456 | 126,87 | 192,594 | 212,741 | 7,30104 | 3,16475 | 3,05926 | 0,289914 | 0,217689 | 0,313842 | Nnat | -3,97770806 | 2,68E-30 |
| Dach1 | 12,6482 | 15,3853 | 11,156 | 11,9958 | 1,66378 | 0,915539 | 1,04672 | 1,41858 | 2,00239 | 1,08498 | Dach1 | -3,13632704 | 4,71E-30 |
| Hn1l | 65,9746 | 72,0249 | 51,1875 | 42,3411 | 6,78901 | 6,26264 | 5,66515 | 2,02551 | 3,97369 | 2,31121 | Hn1l | -3,05016413 | 4,96E-30 |
| Isl1 | 17,7901 | 23,3996 | 16,9389 | 10,6202 | 0,18683 | 0,033064 | 0 | 0 | 0 | 0 | Isl1 | -7,14911371 | 5,13E-30 |
| Mcm10 | 15,1332 | 21,1569 | 10,8494 | 18,3332 | 0,999737 | 0,814386 | 0,739969 | 0,0844393 | 0,0310782 | 0 | Mcm10 | -3,77664231 | 1,16E-29 |
| Smc2 | 29,8612 | 31,5749 | 26,2444 | 22,4603 | 5,50313 | 4,21959 | 4,25434 | 0,596761 | 0,477833 | 0,500871 | Smc2 | -2,33833809 | 3,32E-29 |
| Rimklb | 5,04385 | 9,63558 | 9,69008 | 7,34032 | 0,249506 | 0,0802303 | 0,0525775 | 0 | 0 | 0 | Rimklb | -5,41433596 | 4,19E-29 |
| Igfbp5 | 145,926 | 253,787 | 341,326 | 259,943 | 25,8206 | 17,725 | 14,5302 | 10,2676 | 17,2636 | 12,3292 | Igfbp5 | -3,34272398 | 4,94E-29 |
| Grip1 | 5,6481 | 12,2146 | 11,8926 | 7,65349 | 0,41686 | 0,213196 | 0,289673 | 0 | 0,0287479 | 0 | Grip1 | -4,26788844 | 5,65E-29 |
| Nptx2 | 47,0623 | 36,1379 | 15,5604 | 18,7802 | 0,0739297 | 0 | 0,0399934 | 0 | 0,0874868 | 0,0422493 | Nptx2 | -8,38425387 | 1,38E-28 |
| Frmd4b | 23,5181 | 26,6932 | 27,3886 | 23,0801 | 3,13361 | 1,8522 | 1,41391 | 1,72616 | 1,2885 | 0,940068 | Frmd4b | -3,18295479 | 3,21E-28 |
| Atp7b | 7,92033 | 6,21066 | 8,24997 | 11,7314 | 0,191751 | 0,152581 | 0,373342 | 0 | 0,0226243 | 0 | Atp7b | -4,49470804 | 3,29E-28 |
| Sdf2l1 | 68,3401 | 76,5142 | 44,7702 | 52,2951 | 4,47226 | 3,88182 | 2,84234 | 12,6681 | 11,2732 | 14,2898 | Sdf2l1 | -3,66953461 | 3,86E-28 |
| Lig1 | 74,3451 | 87,9464 | 55,9353 | 66,6953 | 6,5939 | 8,53641 | 7,8473 | 1,05701 | 1,13373 | 0 | Lig1 | -2,80833187 | 8,21E-28 |
| Mthfd1l | 36,1034 | 37,7618 | 36,0905 | 37,0101 | 4,84083 | 2,25515 | 2,38145 | 0 | 0,0896824 | 0,0867096 | Mthfd1l | -3,20814396 | 1,47E-27 |
| Praf2 | 19,7 | 24,2913 | 31,3402 | 19,217 | 2,88856 | 1,39735 | 1,7058 | 0,278554 | 1,67142 | 1,00436 | Praf2 | -3,6734826 | 2,03E-27 |
| Rgs4 | 16,6427 | 9,36589 | 12,2133 | 11,3652 | 0,126645 | 0 | 0,0342529 | 0,304626 | 0,748683 | 0,3979 | Rgs4 | -7,09487393 | 2,08E-27 |
| Melk | 19,9681 | 28,1008 | 20,2455 | 18,5254 | 2,81414 | 2,99875 | 2,35257 | 0,102556 | 0 | 0 | Melk | -2,65204528 | 2,21E-27 |
| Rad54l | 9,09362 | 14,1945 | 13,5312 | 12,4566 | 0 | 0 | 0 | 0 | 0 | 0 | Rad54l | -3,0960122 | 3,29E-27 |
| Arid3a | 29,831 | 22,3018 | 21,1285 | 23,7903 | 1,80893 | 0,763523 | 0,707171 | 0,166467 | 0,256969 | 0,363168 | Arid3a | -4,09939792 | 4,56E-27 |
| Rab11fip1 | 21,3522 | 17,8371 | 11,3108 | 10,5186 | 0,646509 | 0,460456 | 0,45527 | 0,179115 | 0,0985118 | 0,111234 | Rab11fip1 | -4,55054656 | 4,60E-27 |
| Crmp1 | 28,424 | 27,8218 | 30,0031 | 23,4286 | 0,83077 | 0,138664 | 0,135663 | 0 | 0 | 0 | Crmp1 | -5,60805695 | 4,86E-27 |
| Wnt5a | 25,0397 | 20,8205 | 23,6327 | 23,5612 | 2,96717 | 1,36001 | 1,75307 | 4,57211 | 6,19997 | 4,93191 | Wnt5a | -3,23636395 | 6,03E-27 |
| Vsnl1 | 9,3826 | 19,21 | 24,8233 | 17,4752 | 0,150703 | 0,40029 | 0,434877 | 0 | 0,059578 | 0 | Vsnl1 | -5,03669034 | 7,22E-27 |
| Pole | 14,0834 | 16,5852 | 12,1968 | 16,8512 | 2,36768 | 1,89447 | 2,05541 | 0,881611 | 0,837142 | 0,895618 | Pole | -2,45278321 | 7,87E-27 |
| Gldc | 13,2166 | 23,2162 | 7,04625 | 17,3534 | 0,072479 | 0 | 0 | 0 | 0,0285292 | 0,110345 | Gldc | -7,58721098 | 1,31E-26 |
| Clspn | 13,12 | 13,0689 | 9,59265 | 13,187 | 1,68174 | 1,32694 | 1,50611 | 0 | 0,0426643 | 0,0412767 | Clspn | -2,76131018 | 3,28E-26 |
| Ap3b2 | 6,27291 | 10,7039 | 30,3197 | 17,4227 | 0,0720542 | 0 | 0,0519629 | 0 | 0 | 0 | Ap3b2 | -7,63898355 | 3,74E-26 |
| Elovl6 | 9,02226 | 10,2356 | 11,7904 | 9,06393 | 1,55555 | 1,30199 | 1,01601 | 0,0966275 | 0,070883 | 0,0342982 | Elovl6 | -2,58217886 | 6,32E-26 |
| Cd24a | 157,413 | 196,573 | 128,749 | 109,03 | 22,8497 | 17,6032 | 18,4139 | 0,345408 | 1,15391 | 0,525482 | Cd24a | -2,71080312 | 8,09E-26 |
| Myl4 | 1253,08 | 1640,33 | 2330,79 | 1902,94 | 219,913 | 123,481 | 95,4587 | 33,8157 | 47,7934 | 16,2076 | Myl4 | -3,25038184 | 1,02E-25 |
| Mdk | 878,955 | 1032,98 | 822,377 | 779,208 | 56,6092 | 12,1956 | 15,0093 | 3,86208 | 1,29149 | 0,880803 | Mdk | -4,96844436 | 1,98E-25 |
| Tead2 | 155,581 | 168,44 | 140,056 | 118,864 | 26,8346 | 26,8135 | 25,2961 | 0,308166 | 0,661517 | 0,468166 | Tead2 | -2,13334306 | 2,24E-25 |
| Mcm4 | 66,9475 | 62,4831 | 49,2541 | 49,7231 | 6,36603 | 8,52426 | 8,32472 | 2,46195 | 2,56678 | 2,59842 | Mcm4 | -2,66244954 | 3,13E-25 |
| Crabp2 | 33,8878 | 73,0126 | 50,9749 | 99,6421 | 0,381254 | 0 | 0 | 0 | 0 | 0 | Crabp2 | -7,31722016 | 3,24E-25 |
| Map1b | 65,0726 | 44,8642 | 48,4925 | 38,9102 | 6,57531 | 4,83273 | 4,54609 | 0,14289 | 0,479305 | 0,312252 | Map1b | -2,90033409 | 1,18E-24 |
| Kcnmb1 | 9,72982 | 5,29049 | 9,78434 | 6,94703 | 0,130225 | 0,0383835 | 0,0234777 | 0 | 0,0256192 | 0,0991117 | Kcnmb1 | -6,26379629 | 1,67E-24 |
| Akap5 | 13,1425 | 11,5564 | 10,1591 | 6,92126 | 0,681104 | 0,803802 | 0,746824 | 0,182161 | 0,363017 | 0,170546 | Akap5 | -3,44979569 | 3,81E-24 |
| 3632451O06R | 107,112 | 118,206 | 76,6618 | 72,7856 | 7,22519 | 10,5434 | 9,40235 | 5,57256 | 9,07008 | 8,25626 | 3632451O06R | -3,04347933 | 5,86E-24 |
| Tonsl | 10,6747 | 11,5836 | 8,59213 | 11,216 | 1,25284 | 1,33701 | 1,28512 | 0,138748 | 0,0764897 | 0,123298 | Tonsl | -2,68509783 | 8,22E-24 |
| Mcm7 | 73,5223 | 69,9801 | 50,2359 | 68,0854 | 13,6229 | 11,565 | 10,6261 | 3,29852 | 2,29731 | 2,35413 | Mcm7 | -2,70775024 | 8,31E-24 |
| Brca1 | 6,95351 | 6,67302 | 6,83097 | 5,48115 | 0,964395 | 0,734042 | 0,898211 | 0,0861351 | 0,0315772 | 0,030563 | Brca1 | -2,57890047 | 8,78E-24 |
| Sema3a | 11,649 | 12,1478 | 10,4347 | 5,40665 | 0,655798 | 0,740351 | 0,51797 | 0,0446346 | 0,278214 | 0,20591 | Sema3a | -3,6503468 | 9,41E-24 |
| Lmtk3 | 4,1451 | 3,58024 | 6,97996 | 7,03168 | 0,111888 | 0,0164869 | 0,121026 | 0 | 0 | 0 | Lmtk3 | -5,44323981 | 1,86E-23 |
| Itga6 | 28,7313 | 60,0734 | 50,2445 | 51,6243 | 7,08043 | 4,43704 | 3,99957 | 2,14829 | 4,69716 | 2,97233 | Itga6 | -2,89795757 | 2,05E-23 |
| Trim71 | 30,2076 | 13,5125 | 5,86991 | 9,61138 | 0 | 0 | 0 | 0 | 0 | 0 | Trim71 | -9,54810822 | 2,81E-23 |
| Lpcat4 | 21,1307 | 21,5576 | 23,9677 | 19,8289 | 3,12634 | 1,81517 | 1,88568 | 0 | 0,303973 | 0,293214 | Lpcat4 | -2,91747267 | 3,40E-23 |
| Atad2 | 27,3243 | 39,5352 | 20,4063 | 23,328 | 3,60153 | 2,99566 | 3,03458 | 0,557285 | 0,408899 | 0,35971 | Atad2 | -2,71318276 | 3,93E-23 |
| Trp53 | 113,295 | 80,531 | 71,3828 | 73,9664 | 17,1101 | 12,2992 | 13,2184 | 3,97926 | 5,49636 | 5,34112 | Trp53 | -2,82707576 | 4,69E-23 |
| Uhrf1 | 99,7942 | 78,7378 | 49,2541 | 59,8346 | 6,41165 | 7,31862 | 7,68792 | 0,608115 | 0,255853 | 0,556498 | Uhrf1 | -3,3150427 | 9,91E-23 |
| 4833424O15R | 6,51869 | 9,9597 | 8,56 | 5,92694 | 0,74323 | 0,438145 | 0,438529 | 0,289296 | 0,186118 | 0,102853 | 4833424O15R | -3,461091 | 1,19E-22 |
| Wdr90 | 11,8023 | 14,464 | 11,1972 | 11,5251 | 0 | 0 | 0 | 0 | 0 | 0 | Wdr90 | -2,00572879 | 2,48E-22 |
| Hells | 23,1287 | 24,1017 | 16,4957 | 24,9418 | 1,41579 | 2,66818 | 1,69815 | 0,394887 | 0,254692 | 0,28129 | Hells | -3,10882958 | 2,88E-22 |
| Afp | 10,7458 | 31,3198 | 31,2559 | 104,472 | 0 | 0 | 0 | 0 | 0 | 0 | Afp | -9,46344104 | 3,07E-22 |
| Hmgcr | 43,275 | 44,5914 | 44,927 | 48,9148 | 10,7853 | 7,20197 | 6,75192 | 1,84068 | 2,48766 | 2,59313 | Hmgcr | -2,14077206 | 3,69E-22 |
| Wnt11 | 19,3547 | 28,6675 | 24,6823 | 15,6234 | 1,26596 | 0,385864 | 0,327693 | 3,33751 | 2,51583 | 3,18496 | Wnt11 | -4,4072328 | 4,93E-22 |
| Kntc1 | 13,1262 | 11,3026 | 12,6513 | 10,0445 | 1,64761 | 2,09978 | 1,78203 | 0 | 0 | 0,0437129 | Kntc1 | -2,33646815 | 5,51E-22 |
| Mcm3 | 110,35 | 95,8144 | 60,0417 | 83,8371 | 7,72331 | 8,40487 | 9,1247 | 0,414518 | 0,687805 | 0,92304 | Mcm3 | -3,01770118 | 5,65E-22 |
| Bub1 | 26,3437 | 32,075 | 19,7351 | 24,8297 | 4,13284 | 3,34097 | 3,44185 | 0 | 0,0493943 | 0,0238879 | Bub1 | -2,48291943 | 6,11E-22 |
| Atad5 | 4,95862 | 5,82752 | 5,14696 | 5,73494 | 0,952278 | 0,809309 | 0,660233 | 0,117785 | 0,201434 | 0,33426 | Atad5 | -2,3936553 | 1,14E-21 |
| Timeless | 21,6431 | 22,5142 | 17,0433 | 19,4374 | 3,19453 | 2,46183 | 1,99976 | 0,463025 | 0,81572 | 0,799243 | Timeless | -2,55018354 | 1,16E-21 |
| Ctnnd2 | 2,86739 | 3,87636 | 5,40756 | 7,49036 | 0,0751865 | 0,0132926 | 0 | 0,048339 | 0,0531902 | 0,0171581 | Ctnnd2 | -6,48387842 | 1,49E-21 |
| Asns | 41,1093 | 54,4223 | 28,2218 | 41,0498 | 4,79747 | 2,86157 | 3,33346 | 0,807672 | 0,299477 | 0,635571 | Asns | -3,13069314 | 1,58E-21 |
| Chtf18 | 21,4842 | 17,7655 | 15,1198 | 20,0528 | 2,26855 | 2,00668 | 1,55001 | 0,478776 | 0,28225 | 0,204587 | Chtf18 | -2,90059889 | 1,81E-21 |
| 2810417H13R | 41,8565 | 55,4694 | 29,855 | 31,9003 | 3,06861 | 3,61885 | 4,47825 | 0,254782 | 0 | 0,0909213 | 2810417H13R | -3,30304193 | 1,90E-21 |
| Gpc2 | 22,1317 | 19,0872 | 24,2248 | 22,5621 | 1,61586 | 2,69341 | 2,1955 | 0 | 0 | 0,128857 | Gpc2 | -2,88150875 | 2,03E-21 |
| Cnn2 | 372,002 | 315,751 | 327,744 | 261,775 | 72,0078 | 42,9659 | 46,3525 | 12,3509 | 17,3947 | 13,33 | Cnn2 | -2,4888505 | 3,33E-21 |

**Fold Change > 2.0**
***p* < 0.05**
**n=1915**

| Gene | d9-11 (1) | d9-11 (2) | d9-11(3) | d9-11(4) | newborn(1) | newborn(2) | newborn(3) | adult(1) | adult(2) | adult(3) | Gene | log2(fold) | FDR(*p* -value) |
|---|---|---|---|---|---|---|---|---|---|---|---|---|---|
| Igsf9 | 5,66841 | 13,9414 | 20,2547 | 23,6265 | 0,285977 | 0,0778079 | 0,0951835 | 0,0706468 | 0,0259675 | 0,100457 | Igsf9 | -5,71699446 | 4,02E-21 |
| Fmnl2 | 6,42545 | 6,81814 | 6,8509 | 7,23583 | 1,08567 | 0,594753 | 0,496176 | 0,638992 | 0,775294 | 0,88985 | Fmnl2 | -2,81808654 | 4,78E-21 |
| Hs3st3b1 | 3,7323 | 3,68193 | 3,11548 | 2,95287 | 0,210891 | 0,120625 | 0,056355 | 0,0957102 | 0,070207 | 0,127393 | Hs3st3b1 | -4,38673817 | 4,92E-21 |
| Chaf1b | 29,2475 | 41,2423 | 25,0103 | 29,6408 | 4,63354 | 3,47141 | 2,92029 | 0,162863 | 0 | 0,0582596 | Chaf1b | -2,71703667 | 9,22E-21 |
| Tlx1 | 5,87466 | 6,95427 | 8,40566 | 6,93196 | 0,142865 | 0,126475 | 0,0515304 | 0 | 0 | 0 | Tlx1 | -5,11157896 | 1,03E-20 |
| Igfbp2 | 46,6195 | 61,678 | 96,3538 | 220,929 | 2,64997 | 2,21099 | 0,928086 | 0 | 0 | 0 | Igfbp2 | -5,25259138 | 1,38E-20 |
| Myrf | 14,8148 | 17,0693 | 29,0279 | 15,041 | 0,640553 | 0,0424693 | 0,173211 | 1,33825 | 1,05753 | 1,15124 | Myrf | -5,46898966 | 1,43E-20 |
| Acta1 | 400,769 | 348,358 | 614,416 | 529,264 | 58,0402 | 68,414 | 72,7089 | 346,272 | 211,189 | 395,075 | Acta1 | -2,54437269 | 1,66E-20 |
| Eme1 | 0 | 0 | 0 | 0 | 0 | 0 | 0 | 0 | 0 | 0 | Eme1 | -3,63345878 | 1,83E-20 |
| Tiparp | 27,2779 | 40,7456 | 24,8979 | 30,1302 | 5,24272 | 3,87158 | 2,91274 | 4,42931 | 7,13227 | 7,77269 | Tiparp | -2,56439914 | 3,35E-20 |
| Top2a | 78,072 | 86,1015 | 62,2908 | 70,2843 | 19,2054 | 14,686 | 13,7651 | 0,055378 | 0,0609765 | 0,196659 | Top2a | -2,03004636 | 4,12E-20 |
| Hmga2-ps1 | 33,5829 | 24,1109 | 13,0974 | 16,6741 | 0,568306 | 0,236963 | 0,144732 | 0 | 0,19879 | 0 | Hmga2-ps1 | -5,23144894 | 6,16E-20 |
| Wnt2 | 45,8535 | 22,3005 | 12,0437 | 13,0815 | 0,272894 | 0,0402605 | 0 | 0 | 0,107798 | 0 | Wnt2 | -6,90560762 | 7,76E-20 |
| Mcm5 | 88,3307 | 69,7664 | 47,0781 | 58,8653 | 7,23636 | 7,6136 | 7,77 | 1,03599 | 0,85808 | 0,739269 | Mcm5 | -2,80280264 | 7,88E-20 |
| Hs6st2 | 7,21455 | 14,4285 | 14,0582 | 12,6193 | 1,27662 | 0,487995 | 0,486445 | 0,270326 | 0,173834 | 0,0960778 | Hs6st2 | -3,6451132 | 1,10E-19 |
| Rps20 | 891,77 | 1019,67 | 630,371 | 824,502 | 248,895 | 209,319 | 194,735 | 168,12 | 157,913 | 149,781 | Rps20 | -2,42868195 | 1,21E-19 |
| B4galnt4 | 4,9515 | 6,6514 | 9,40917 | 12,9486 | 0,0298597 | 0 | 0 | 0 | 0 | 0 | B4galnt4 | -7,92593476 | 1,26E-19 |
| Msx2 | 8,37014 | 17,7487 | 21,9381 | 23,1668 | 1,15388 | 0,746373 | 0,528228 | 0,426241 | 0,630999 | 0,7613 | Msx2 | -4,0474556 | 1,85E-19 |
| Kif15 | 14,7289 | 17,7227 | 15,2993 | 15,4057 | 2,07661 | 0,793935 | 0,971334 | 0 | 0,0220718 | 0,0213527 | Kif15 | -3,15541381 | 3,33E-19 |
| Cgn | 2,51359 | 4,30569 | 3,66958 | 5,2326 | 0,140004 | 0,0737389 | 0,0773294 | 0,0574315 | 0,0421667 | 0,030597 | Cgn | -4,881547 | 3,35E-19 |
| Aplp1 | 11,192 | 15,043 | 43,5556 | 29,7364 | 0,91251 | 0,315991 | 0,343411 | 0,381297 | 0,98639 | 0,544267 | Aplp1 | -5,00106974 | 3,35E-19 |
| Gap43 | 48,8618 | 52,9091 | 28,3577 | 16,1188 | 1,03766 | 0,424594 | 0,432041 | 0 | 0 | 0,0915323 | Gap43 | -5,15834046 | 3,52E-19 |
| Tmem108 | 26,9656 | 30,5744 | 31,2049 | 24,9626 | 4,86223 | 2,63264 | 2,12 | 0,318547 | 0,292971 | 0,254945 | Tmem108 | -2,81895546 | 4,00E-19 |
| Cdh24 | 9,08803 | 12,0732 | 8,29365 | 10,4982 | 1,0993 | 1,09418 | 0,929162 | 0,330402 | 0,0406351 | 0,157025 | Cdh24 | -2,57427169 | 4,14E-19 |
| Slc39a8 | 23,2176 | 25,7145 | 11,2299 | 13,213 | 1,18206 | 1,10661 | 0,662858 | 0,262182 | 0,698964 | 0,497645 | Slc39a8 | -3,80354828 | 5,45E-19 |
| Hmgn1 | 297,279 | 344,92 | 231,727 | 226,503 | 71,1603 | 61,4327 | 63,5345 | 26,483 | 32,2702 | 22,5559 | Hmgn1 | -2,06313652 | 5,91E-19 |
| Cdh1 | 5,7278 | 5,57895 | 3,63122 | 12,4185 | 0,0205388 | 0 | 0 | 0 | 0,0242388 | 0 | Cdh1 | -7,93662819 | 7,11E-19 |
| Tuba1a | 474,439 | 555,417 | 591,595 | 529,852 | 126,753 | 82,6196 | 87,3452 | 17,091 | 21,4773 | 18,4244 | Tuba1a | -2,12071843 | 8,55E-19 |
| 9430015G10R | 16,1528 | 13,7199 | 16,3281 | 14,9776 | 2,93384 | 2,84313 | 2,52146 | 0,844033 | 0,935182 | 1,04922 | 9430015G10R | -2,04548957 | 1,25E-18 |
| Wnk3 | 2,96859 | 2,03205 | 2,75617 | 1,52616 | 0,162927 | 0,0909355 | 0,0649194 | 0 | 0,0100998 | 0 | Wnk3 | -3,92971809 | 1,25E-18 |
| Cap1 | 100,169 | 115,507 | 94,5411 | 95,6952 | 25,2741 | 14,3008 | 13,5799 | 7,87652 | 8,38043 | 6,34684 | Cap1 | -2,30092756 | 1,35E-18 |
| Ahcy | 59,8339 | 55,7964 | 40,7624 | 55,1981 | 11,3972 | 8,32739 | 9,37876 | 8,20907 | 7,90799 | 8,38033 | Ahcy | -2,51848241 | 1,54E-18 |
| Mapk8ip1 | 14,2416 | 16,3586 | 18,3104 | 22,3809 | 2,69806 | 1,21317 | 1,30471 | 0,610818 | 0,638032 | 0,906615 | Mapk8ip1 | -3,04193467 | 1,64E-18 |
| Pawr | 65,5499 | 57,5819 | 39,7847 | 27,2091 | 4,5036 | 1,66167 | 2,32078 | 0,514268 | 0,508979 | 0,490859 | Pawr | -3,73072958 | 1,75E-18 |
| Adcy1 | 2,87992 | 4,7102 | 8,22429 | 4,68387 | 0,4362 | 0,470757 | 0,348014 | 0,805437 | 1,55776 | 1,06007 | Adcy1 | -3,2054333 | 2,26E-18 |
| Mcm6 | 98,454 | 97,6831 | 71,4391 | 79,8854 | 13,1428 | 17,4546 | 14,914 | 0,412207 | 1,29187 | 1,35843 | Mcm6 | -2,26206636 | 2,31E-18 |
| Exo1 | 9,37545 | 7,09272 | 4,84922 | 4,92605 | 0,586428 | 0,489615 | 0,458106 | 0,0523557 | 0,0192107 | 0 | Exo1 | -3,36734866 | 2,47E-18 |
| Fdps | 35,2389 | 33,0454 | 32,5015 | 28,9568 | 9,56277 | 11,3561 | 9,20587 | 4,08686 | 6,0834 | 5,98792 | Fdps | -2,15155223 | 2,51E-18 |
| Npas4 | 18,0958 | 9,41615 | 4,56458 | 5,80987 | 0,14089 | 0,0747705 | 0,0914496 | 0 | 0 | 0 | Npas4 | -5,90463258 | 2,85E-18 |
| Tmem132c | 3,2932 | 6,92658 | 12,2923 | 12,5211 | 0,246383 | 0,0777933 | 0,0380716 | 0 | 0 | 0 | Tmem132c | -5,52428791 | 3,16E-18 |
| Zwilch | 13,6309 | 21,164 | 15,0381 | 20,2995 | 3,15859 | 2,76473 | 3,01734 | 0 | 0,23845 | 0 | Zwilch | -2,29235206 | 3,93E-18 |
| 2610318N02R | 19,4429 | 16,8894 | 10,1718 | 18,1177 | 1,07247 | 0,219446 | 0,178621 | 0 | 0 | 0 | 2610318N02R | -4,71935409 | 3,93E-18 |
| Tcf7 | 20,923 | 20,4906 | 14,0469 | 21,9929 | 1,84921 | 1,33222 | 0,837578 | 0,137705 | 0 | 0,0983515 | Tcf7 | -3,34500375 | 4,01E-18 |
| Mycn | 51,2877 | 28,6409 | 28,1834 | 43,9364 | 4,06156 | 2,88784 | 3,09954 | 0,464915 | 0,343272 | 0,248675 | Mycn | -3,19899443 | 4,08E-18 |
| Slit1 | 4,36238 | 1,91267 | 5,91728 | 2,86921 | 0,0686405 | 0,0303411 | 0 | 0,0551467 | 0 | 0 | Slit1 | -5,54719595 | 4,92E-18 |
| B3gnt5 | 9,53386 | 12,5514 | 10,0393 | 12,1368 | 2,32274 | 1,65373 | 2,02058 | 0 | 0,0469595 | 0,0227124 | B3gnt5 | -2,11918428 | 5,00E-18 |
| Lrp2 | 2,6728 | 2,24321 | 0,715962 | 3,31637 | 0,0338731 | 0,00498838 | 0,0183166 | 0,0545161 | 0,0265845 | 0,0128742 | Lrp2 | -5,93839365 | 5,25E-18 |
| Cadm3 | 5,32518 | 3,57832 | 3,47871 | 4,58767 | 0,131185 | 0,0193333 | 0,0709522 | 0,0702174 | 0 | 0 | Cadm3 | -5,17863411 | 5,98E-18 |
| Mms22l | 8,37908 | 9,8909 | 8,36911 | 11,3285 | 1,84612 | 1,34531 | 1,36055 | 0 | 0,0718221 | 0,046315 | Mms22l | -2,29839312 | 6,70E-18 |
| Ube2c | 186,029 | 203,316 | 195,07 | 239,579 | 58,9717 | 32,2385 | 29,8569 | 0,384905 | 0,583444 | 0,978624 | Ube2c | -2,50444803 | 7,37E-18 |
| Ncapg | 15,1989 | 19,3924 | 14,8199 | 19,1957 | 2,98504 | 2,36777 | 1,95391 | 0 | 0 | 0 | Ncapg | -2,32653905 | 9,99E-18 |
| Fndc4 | 18,0825 | 16,6684 | 19,4228 | 12,2431 | 2,64844 | 1,41385 | 1,59201 | 1,61554 | 1,89201 | 2,11496 | Fndc4 | -2,72641251 | 1,04E-17 |
| Phf21b | 2,96648 | 3,95927 | 7,28278 | 6,32905 | 0,123609 | 0,0436863 | 0,0802289 | 0,634906 | 0,321153 | 0,282294 | Phf21b | -5,04391249 | 1,27E-17 |
| Fsd1 | 7,55367 | 10,9811 | 8,62726 | 10,0108 | 0,112853 | 0 | 0 | 0 | 0 | 0 | Fsd1 | -6,79753777 | 1,29E-17 |
| E2f8 | 34,1108 | 25,6588 | 25,1465 | 28,0579 | 5,16141 | 3,33564 | 2,95276 | 0,238979 | 0,293056 | 0,283354 | E2f8 | -2,58532532 | 1,33E-17 |
| Flrt3 | 9,86882 | 9,51978 | 12,0134 | 13,9142 | 1,35547 | 0,490754 | 0,470278 | 0 | 0 | 0,0522484 | Flrt3 | -3,53198656 | 1,46E-17 |
| 5730559C18R | 6,03943 | 6,01153 | 6,08563 | 5,26218 | 0,0308442 | 0 | 0 | 0 | 0 | 0 | 5730559C18R | -7,39616084 | 1,74E-17 |
| Spdl1 | 14,7295 | 15,5777 | 15,728 | 11,8974 | 1,99426 | 2,49718 | 2,36122 | 0 | 0,0890649 | 0,0430088 | Spdl1 | -2,31071645 | 1,76E-17 |
| Nap1l1 | 105,602 | 154,818 | 93,8564 | 105,725 | 28,6399 | 23,0809 | 23,5197 | 17,439 | 19,6009 | 17,8388 | Nap1l1 | -2,26827579 | 1,98E-17 |
| Stra6 | 8,46992 | 5,17915 | 9,69246 | 8,2885 | 0 | 0 | 0 | 0 | 0 | 0,038653 | Stra6 | -8,29971295 | 3,06E-17 |
| Cdca7 | 18,1663 | 25,061 | 10,7799 | 22,5891 | 2,65647 | 1,92349 | 1,17583 | 0,515725 | 0,190626 | 0,276091 | Cdca7 | -3,24861197 | 3,11E-17 |
| Fancd2 | 8,66726 | 12,9569 | 8,06955 | 9,67313 | 1,01978 | 1,48599 | 1,30733 | 0 | 0,0222807 | 0,0215545 | Fancd2 | -2,51946459 | 3,76E-17 |
| Tagln | 259,645 | 439,971 | 498,945 | 451,851 | 33,0379 | 0 | 0 | 11,9452 | 7,80379 | 6,90089 | Tagln | -4,07279412 | 4,09E-17 |
| Etv4 | 15,2215 | 14,8741 | 14,3829 | 13,3083 | 1,54758 | 0,576676 | 0,705033 | 0 | 0,0964483 | 0,139684 | Etv4 | -3,47915373 | 4,67E-17 |
| Mgarp | 24,4211 | 15,7338 | 10,9851 | 21,2442 | 0,598125 | 0,379391 | 0,555037 | 0 | 0 | 0 | Mgarp | -4,37670117 | 4,69E-17 |
| Tyms | 15,5644 | 16,8973 | 10,2828 | 8,43121 | 1,26778 | 1,23468 | 1,3801 | 0,0771766 | 0,227268 | 0,151172 | Tyms | -3,00382066 | 5,14E-17 |
| Fgf8 | 49,938 | 20,8547 | 13,4103 | 23,4111 | 0 | 0 | 0,0954472 | 0 | 0 | 0,126139 | Fgf8 | -7,68203242 | 5,48E-17 |
| Wwc1 | 5,99068 | 8,3318 | 13,8536 | 13,456 | 0,862786 | 0,393869 | 0,511844 | 0,0893083 | 0,197313 | 0,0953648 | Wwc1 | -3,72721593 | 5,53E-17 |
| Haus5 | 8,30735 | 10,0921 | 10,7218 | 9,91794 | 1,65822 | 2,20158 | 1,84145 | 1,11774 | 1,13733 | 0,79842 | Haus5 | -2,46903919 | 5,89E-17 |
| Polr2h | 32,8669 | 22,931 | 16,4991 | 21,5134 | 6,17779 | 8,48014 | 6,2809 | 5,77302 | 7,54619 | 4,74951 | Polr2h | -2,20677745 | 6,76E-17 |
| Hspa1b | 77,1612 | 23,479 | 40,9074 | 142,37 | 3,56483 | 1,54503 | 1,7934 | 7,94614 | 9,12034 | 11,295 | Hspa1b | -4,81041316 | 7,41E-17 |
| Alcam | 14,1781 | 20,4421 | 28,9176 | 25,1445 | 4,42464 | 2,95595 | 2,67307 | 0,291935 | 0,278669 | 0,311081 | Alcam | -2,43807759 | 8,63E-17 |
| Atp2a1 | 2,54874 | 9,6848 | 15,6013 | 10,8419 | 0 | 0 | 0 | 0 | 0,948064 | 0,611013 | Atp2a1 | -4,81149578 | 8,75E-17 |
| **Hoxb2** | 15,3036 | 11,2692 | 6,6562 | 8,24947 | 0,183605 | 0,0542136 | 0,066237 | 0 | 0,145436 | 0,350407 | Hoxb2 | -5,77355872 | 9,81E-17 |
| Epcam | 5,61083 | 10,9071 | 6,61144 | 20,7787 | 0,0468184 | 0,0414457 | 0 | 0 | 0,055495 | 0 | Epcam | -7,05435286 | 1,32E-16 |
| Gstcd | 9,12056 | 11,4966 | 11,5249 | 12,2507 | 1,86762 | 1,81138 | 2,05947 | 0,575594 | 0,424946 | 0,738846 | Gstcd | -2,12922597 | 1,35E-16 |
| Rps21 | 330,774 | 405,808 | 255,257 | 334,274 | 104,889 | 97,995 | 111,387 | 80,2682 | 71,888 | 85,1247 | Rps21 | -2,11600969 | 1,37E-16 |
| Cdh6 | 30,8681 | 16,0334 | 9,46472 | 9,39244 | 0,535831 | 0,0948207 | 0,0772963 | 0 | 0 | 0 | Cdh6 | -5,32694727 | 1,55E-16 |
| Me2 | 50,1695 | 38,091 | 33,0236 | 34,1039 | 5,89885 | 6,37997 | 4,46343 | 5,80596 | 4,28986 | 5,02417 | Me2 | -2,49985104 | 1,69E-16 |
| Snrpa | 134,113 | 124,224 | 116,304 | 120,152 | 44,4779 | 39,989 | 47,9587 | 35,5332 | 28,3275 | 31,346 | Snrpa | -2,05847285 | 1,84E-16 |
| Tmem237 | 11,3696 | 14,6837 | 13,1344 | 12,4063 | 1,7528 | 0,463193 | 0,58201 | 0,343908 | 0,893522 | 0,430849 | Tmem237 | -2,81949882 | 1,89E-16 |
| Myo16 | 1,00084 | 3,70252 | 4,59492 | 3,57997 | 0,0425121 | 0 | 0 | 0 | 0 | 0 | Myo16 | -6,69499859 | 1,94E-16 |
| Limd2 | 46,0897 | 45,0278 | 29,0389 | 33,6468 | 5,94845 | 3,57327 | 3,92607 | 1,11329 | 1,34316 | 1,08159 | Limd2 | -2,55221466 | 2,54E-16 |
| Pax6 | 2,47823 | 5,07644 | 5,49484 | 6,50534 | 0,0367753 | 0,0325404 | 0 | 0 | 0 | 0 | Pax6 | -5,94722811 | 2,62E-16 |
| Cenpt | 28,077 | 24,2108 | 29,1696 | 28,2799 | 5,18309 | 4,63951 | 4,34255 | 0,712567 | 0,661591 | 0,956874 | Cenpt | -2,09054222 | 2,80E-16 |
| Uba6 | 8,05667 | 8,25598 | 11,782 | 9,69075 | 1,73354 | 2,00175 | 1,60723 | 1,36423 | 2,52479 | 2,05919 | Uba6 | -2,10139177 | 2,86E-16 |
| E2f2 | 4,84493 | 4,9123 | 4,38119 | 5,22049 | 0,686174 | 0,640374 | 0,721601 | 0 | 0,0449772 | 0,0217553 | E2f2 | -2,46250433 | 2,90E-16 |
| Rpl13a | 456,131 | 471,631 | 350,782 | 452,214 | 178,295 | 135,107 | 100,022 | 99,239 | 82,0247 | 84,2831 | Rpl13a | -2,12465112 | 3,19E-16 |
| Hsph1 | 106,429 | 80,5176 | 46,0247 | 60,7213 | 6,43769 | 7,55261 | 7,80788 | 29,3591 | 25,7973 | 25,6945 | Hsph1 | -2,97304309 | 3,19E-16 |
| Rassf5 | 23,3815 | 28,9341 | 42,8569 | 26,9215 | 5,11144 | 2,43981 | 2,13573 | 0 | 0,223659 | 0,185324 | Rassf5 | -2,90586165 | 3,83E-16 |
| Vcan | 84,9083 | 114,171 | 115,872 | 90,2485 | 23,1362 | 13,0073 | 11,0547 | 0,335128 | 0,270797 | 0,142862 | Vcan | -2,44261543 | 4,00E-16 |
| Rnf165 | 9,23382 | 4,61817 | 3,69259 | 5,97191 | 0,474255 | 0,204213 | 0,227975 | 0,0782133 | 0,0573245 | 0,194213 | Rnf165 | -3,95696343 | 7,36E-16 |
| Arid3b | 115,133 | 62,716 | 37,2033 | 36,8768 | 2,72851 | 3,80394 | 2,76872 | 1,55043 | 1,11201 | 0,992549 | Arid3b | -3,90346505 | 8,13E-16 |
| Cfl1 | 1058,46 | 931,818 | 800,291 | 736,189 | 254,998 | 172,568 | 168,382 | 42,0154 | 55,0487 | 51,8892 | Cfl1 | -2,09881351 | 8,69E-16 |
| Ran | 134,911 | 163,663 | 88,5408 | 116,902 | 41,9501 | 46,6359 | 40,3624 | 37,3027 | 38,2649 | 35,8463 | Ran | -2,1382583 | 8,73E-16 |
| Lmod1 | 10,0458 | 11,1993 | 8,86261 | 8,96928 | 0,416516 | 0,0204625 | 0,0250311 | 0,074301 | 0,191242 | 0,0792592 | Lmod1 | -5,2608935 | 1,07E-15 |
| Phgdh | 20,929 | 21,1753 | 17,8905 | 29,0406 | 3,05461 | 0,863049 | 1,07359 | 0 | 0,185844 | 0,209126 | Phgdh | -3,76889601 | 1,09E-15 |
| Wdhd1 | 10,4562 | 16,8378 | 9,04885 | 11,232 | 1,78605 | 1,73338 | 1,83775 | 0,209855 | 0,128551 | 0,0994634 | Wdhd1 | -2,43277416 | 1,14E-15 |
| Slc25a1 | 28,3302 | 28,1524 | 27,8178 | 27,4851 | 5,85494 | 5,9193 | 3,77629 | 4,91263 | 3,0908 | 2,70843 | Slc25a1 | -2,02614807 | 1,86E-15 |
| Cldn6 | 11,7971 | 14,4304 | 6,10448 | 32,7921 | 0 | 0 | 0 | 0 | 0 | 0 | Cldn6 | -7,44472651 | 2,03E-15 |
| Mybl2 | 28,5033 | 19,4202 | 13,0072 | 16,5793 | 1,55837 | 0,776583 | 0,48165 | 0 | 0,0876377 | 0,0141228 | Mybl2 | -4,08049216 | 2,26E-15 |
| Ercc6l | 4,73806 | 4,90026 | 4,1463 | 4,54273 | 0,736087 | 0,516872 | 0,51521 | 0,0695306 | 0,0255545 | 0,0247155 | Ercc6l | -2,6562819 | 2,36E-15 |
| **Hoxb9** | 3,6777 | 6,89322 | 8,28825 | 8,95364 | 0 | 0,00256253 | 0 | 0 | 0 | 0 | Hoxb9 | -7,94954631 | 2,49E-15 |
| Cotl1 | 79,19 | 91,4197 | 88,4227 | 84,0701 | 19,8716 | 8,98957 | 9,66803 | 1,94107 | 1,01079 | 1,3221 | Cotl1 | -2,43836851 | 2,65E-15 |
| Fjx1 | 7,98567 | 8,42104 | 7,35496 | 10,125 | 0,819932 | 0,276413 | 0,253467 | 0,250183 | 0,0924405 | 0,178533 | Fjx1 | -3,80773534 | 2,82E-15 |
| Trip13 | 29,9838 | 39,3891 | 17,2212 | 24,9706 | 2,69442 | 3,31624 | 2,96093 | 0 | 0,29918 | 0,144414 | Trip13 | -2,82702465 | 3,26E-15 |
| Dnah8 | 0,33878 | 1,71283 | 1,83819 | 1,21615 | 0,0179792 | 0,00529554 | 0,0194443 | 0 | 0,021167 | 0,0136671 | Dnah8 | -5,68628292 | 3,26E-15 |
| Cnksr2 | 2,74963 | 3,05879 | 1,12318 | 3,30095 | 0,0331375 | 0 | 0,0179214 | 0,266253 | 0,0390815 | 0,0945388 | Cnksr2 | -6,0016605 | 3,27E-15 |
| Hbb-y | 40,1737 | 1171,6 | 2251,31 | 1880,04 | 0 | 0 | 0 | 0 | 0 | 0 | Hbb-y | -8,80929942 | 3,44E-15 |
| Myo1b | 27,3997 | 31,5874 | 29,0622 | 21,8035 | 5,23274 | 2,11078 | 2,23559 | 0,296938 | 1,84004 | 1,26573 | Myo1b | -2,77022994 | 3,65E-15 |
| Sec61b | 85,0101 | 93,7247 | 43,4286 | 34,7493 | 18,2655 | 11,8915 | 12,7406 | 7,30493 | 7,37139 | 7,27618 | Sec61b | -3,0832064 | 3,73E-15 |
| Meis3 | 15,7782 | 27,5672 | 29,8271 | 23,8417 | 3,88443 | 3,89367 | 3,59722 | 2,74276 | 2,027 | 2,14725 | Meis3 | -2,21616303 | 4,03E-15 |
| Snrpe | 59,1525 | 103,104 | 58,1352 | 65,6168 | 21,3091 | 27,0545 | 27,6247 | 17,2061 | 17,1405 | 22,6104 | Snrpe | -2,03131771 | 4,04E-15 |

| Gene | d9-11 (1) | d9-11 (2) | d9-11(3) | d9-11(4) | newborn(1) | newborn(2) | newborn(3) | adult(1) | adult(2) | adult(3) | Gene | log2(fold) | FDR(*p*-value) |
|---|---|---|---|---|---|---|---|---|---|---|---|---|---|
| Lpcat1 | 35,0837 | 34,6089 | 24,1572 | 30,035 | 5,2302 | 2,59241 | 2,2238 | 0,795359 | 1,47988 | 0,641737 | Lpcat1 | -2,79283676 | 4,81E-15 |
| Cdca7l | 6,7113 | 16,6756 | 9,44168 | 12,2917 | 0,900904 | 0,478279 | 0,662798 | 0 | 0 | 0 | Cdca7l | -3,35868214 | 4,89E-15 |
| Barx1 | 10,643 | 8,66159 | 11,3252 | 13,1297 | 0,0756121 | 0 | 0 | 0 | 0 | 0 | Barx1 | -7,01443117 | 4,96E-15 |
| Dtl | 24,2391 | 25,0039 | 12,3627 | 16,7681 | 2,72511 | 1,54906 | 1,35692 | 0,0694609 | 0,0255287 | 0,0987624 | Dtl | -3,05269146 | 5,28E-15 |
| Mex3a | 27,7998 | 31,5815 | 28,2149 | 29,343 | 6,41602 | 2,71256 | 2,44734 | 0,249078 | 0,12792 | 0,141472 | Mex3a | -2,63024177 | 5,91E-15 |
| Sall2 | 6,88525 | 6,2576 | 6,64006 | 9,02512 | 1,34436 | 0,954216 | 0,839723 | 0,0608341 | 0,223397 | 0,043223 | Sall2 | -2,4674188 | 6,38E-15 |
| Chaf1a | 42,325 | 27,4796 | 22,5118 | 26,973 | 4,18424 | 3,30615 | 3,68157 | 1,08531 | 0,791052 | 0,764653 | Chaf1a | -2,66294493 | 6,55E-15 |
| Palm3 | 6,27101 | 4,84142 | 8,53411 | 7,82791 | 0,541219 | 0,581563 | 0,376441 | 0 | 0,183042 | 0,132572 | Palm3 | -3,3606297 | 6,67E-15 |
| Pabpc1 | 598,796 | 495,037 | 399,12 | 464,535 | 135,131 | 118,088 | 129,715 | 27,8377 | 28,0584 | 31,4698 | Pabpc1 | -2,08465674 | 7,92E-15 |
| Sall1 | 30,3068 | 21,3215 | 9,07098 | 12,2266 | 0,444568 | 0,0151163 | 0,0924742 | 0 | 0 | 0 | Sall1 | -5,77935512 | 9,93E-15 |
| BC055324 | 10,2609 | 23,0605 | 13,4632 | 8,51158 | 1,21032 | 0,814946 | 1,31314 | 0,308158 | 0 | 0 | BC055324 | -3,25609503 | 1,06E-14 |
| Susd1 | 7,11692 | 10,4337 | 6,60056 | 4,58088 | 0,504234 | 0,346882 | 0,181827 | 0 | 0 | 0,0639913 | Susd1 | -3,71771175 | 1,10E-14 |
| Mthfd2 | 46,4749 | 53,9467 | 28,2363 | 51,4463 | 8,19989 | 5,81532 | 5,14221 | 1,04377 | 1,43579 | 2,34477 | Mthfd2 | -2,5293049 | 1,30E-14 |
| Fanca | 13,3051 | 15,4125 | 10,6979 | 8,2035 | 0,880492 | 1,9019 | 1,48686 | 0 | 0 | 0 | Fanca | -2,63779961 | 1,60E-14 |
| Recql4 | 6,89762 | 6,23744 | 5,43528 | 7,19761 | 0,772958 | 0,83445 | 0,689625 | 0,235651 | 0 | 0 | Recql4 | -2,44468304 | 1,66E-14 |
| Tmsb10 | 1037,24 | 1115,51 | 809,274 | 700,689 | 137,236 | 171,278 | 171,029 | 4,30109 | 6,72057 | 7,6676 | Tmsb10 | -2,18990471 | 1,67E-14 |
| Trim6 | 4,12952 | 3,40866 | 5,83872 | 3,31143 | 0,286537 | 0,295625 | 0,12915 | 0 | 0,0845862 | 0,0272643 | Trim6 | -3,60736628 | 1,78E-14 |
| Lrrn4 | 8,49791 | 3,6849 | 6,54761 | 2,16389 | 0,132313 | 0,0234043 | 0,0572534 | 0,169862 | 0,656471 | 0,241769 | Lrrn4 | -5,55191421 | 1,87E-14 |
| Amot | 57,2439 | 36,0598 | 14,352 | 15,5054 | 1,6739 | 0,666355 | 0,414611 | 2,91731 | 3,29822 | 4,16915 | Amot | -4,61851804 | 1,87E-14 |
| Ubash3b | 14,9477 | 15,1756 | 16,6172 | 16,106 | 3,96085 | 2,31881 | 2,21951 | 0,182858 | 0,0336732 | 0,0325477 | Ubash3b | -2,1876118 | 2,04E-14 |
| P4ha1 | 44,8476 | 38,398 | 33,3861 | 31,3248 | 9,48429 | 6,0958 | 6,57966 | 19,8219 | 16,622 | 15,8178 | P4ha1 | -2,05116911 | 2,26E-14 |
| Cdca8 | 115,994 | 85,8712 | 78,0614 | 76,3613 | 19,013 | 14,6147 | 13,2094 | 0,195889 | 0 | 0 | Cdca8 | -2,22423571 | 2,42E-14 |
| Tes | 12,308 | 9,4435 | 7,76195 | 11,0211 | 1,08585 | 0,480448 | 0,545464 | 0,124251 | 0,459058 | 0,221653 | Tes | -3,39806225 | 2,46E-14 |
| Cdkn1c | 675,913 | 845,452 | 803,618 | 542,242 | 38,5938 | 3,85264 | 5,65124 | 2,61966 | 3,66234 | 3,23361 | Cdkn1c | -4,83451184 | 2,63E-14 |
| Zic2 | 2,87662 | 3,05606 | 6,2798 | 12,9297 | 0 | 0,027743 | 0 | 0 | 0 | 0,0358292 | Zic2 | -7,21774539 | 2,85E-14 |
| Plxna3 | 3,79914 | 7,13881 | 10,1934 | 8,52191 | 1,16538 | 0,792356 | 0,734115 | 0,0411804 | 0,0754703 | 0,01461 | Plxna3 | -2,65602271 | 3,46E-14 |
| Nefl | 3,07039 | 6,05774 | 17,5118 | 9,53605 | 0,145194 | 0,0428473 | 0 | 0 | 0,0573826 | 0,166097 | Nefl | -5,9204093 | 5,15E-14 |
| Slc39a10 | 25,3088 | 29,7962 | 16,4332 | 17,3805 | 3,57818 | 3,70944 | 4,01446 | 1,05052 | 1,99271 | 1,11942 | Slc39a10 | -2,21675774 | 5,34E-14 |
| Suv39h2 | 7,46966 | 8,0661 | 4,07937 | 8,56573 | 0,747783 | 0,429355 | 0,59496 | 0,211889 | 0,753119 | 0,666677 | Suv39h2 | -3,12291101 | 5,38E-14 |
| Orc1 | 10,7769 | 6,98016 | 7,35988 | 3,83838 | 0,26227 | 0,136474 | 0,0333806 | 0 | 0 | 0,0352497 | Orc1 | -3,96402963 | 6,53E-14 |
| Ddx11 | 9,56601 | 9,08075 | 7,10247 | 6,19584 | 1,40251 | 1,2983 | 1,52897 | 0,563023 | 0,310416 | 0,400289 | Ddx11 | -2,1596217 | 6,93E-14 |
| Spidr | 9,63828 | 13,5421 | 10,029 | 12,4049 | 1,78194 | 1,45143 | 1,01658 | 0 | 0,468248 | 0,840749 | Spidr | -2,48983291 | 6,93E-14 |
| Fam84b | 5,29214 | 4,60321 | 6,96342 | 8,42834 | 0,729428 | 0,285195 | 0,331196 | 0,702828 | 0,826077 | 0,877566 | Fam84b | -3,32023739 | 6,93E-14 |
| Ptbp1 | 196,487 | 166,485 | 104,654 | 117,392 | 28,6626 | 10,4511 | 10,2138 | 2,77259 | 6,34434 | 5,03249 | Ptbp1 | -3,28860314 | 7,53E-14 |
| Mns1 | 22,3902 | 27,3777 | 13,5498 | 12,8795 | 2,094 | 1,93684 | 1,05821 | 0 | 0,257894 | 0 | Mns1 | -3,12326947 | 7,60E-14 |
| Shisa2 | 18,1783 | 19,9822 | 16,3005 | 15,279 | 1,14819 | 0,130209 | 0,159249 | 0,755602 | 0,521941 | 0,538083 | Shisa2 | -4,63830856 | 8,08E-14 |
| Myl1 | 21,8294 | 63,0955 | 167,464 | 108,83 | 7,38666 | 12,0201 | 10,3571 | 19,5097 | 20,7553 | 37,3852 | Myl1 | -3,47352612 | 8,09E-14 |
| Epha2 | 43,4035 | 29,4067 | 17,9097 | 21,411 | 1,90258 | 0,656566 | 0,250986 | 1,11751 | 1,64365 | 1,00666 | Epha2 | -4,41027578 | 9,15E-14 |
| Clmn | 1,26179 | 0,95289 | 1,83052 | 1,84139 | 0,125624 | 0,0326501 | 0,0559439 | 0,0237557 | 0,13923 | 0,0337076 | Clmn | -3,88348903 | 9,54E-14 |
| Pak1 | 5,56304 | 5,59343 | 10,9835 | 15,7356 | 0,576129 | 0,268229 | 0,131216 | 0 | 0,0358434 | 0 | Pak1 | -4,32599166 | 9,77E-14 |
| Bcr | 21,7885 | 23,5522 | 25,6342 | 23,4737 | 6,70588 | 3,32515 | 3,6339 | 1,44617 | 1,33354 | 1,29069 | Bcr | -2,03853547 | 1,06E-13 |
| Cenph | 17,574 | 39,9607 | 22,87 | 20,996 | 3,72635 | 2,85379 | 3,02564 | 0 | 0 | 0 | Cenph | -2,57212212 | 1,14E-13 |
| Lpar2 | 4,4378 | 3,06454 | 3,47285 | 3,80071 | 0,361326 | 0,0694374 | 0,118945 | 0 | 0,0185251 | 0 | Lpar2 | -3,83386885 | 1,24E-13 |
| Npm1 | 397,969 | 527,691 | 220,281 | 291,801 | 81,6103 | 68,9266 | 68,4649 | 36,0487 | 41,2941 | 33,0901 | Npm1 | -2,44440008 | 1,52E-13 |
| Prim2 | 21,1063 | 35,3273 | 11,6961 | 20,3888 | 2,40533 | 1,69492 | 2,07163 | 1,09935 | 1,22236 | 0,336949 | Prim2 | -3,08128512 | 1,52E-13 |
| Rad54b | 5,4462 | 5,46308 | 8,14979 | 5,27121 | 0,732575 | 0,648052 | 0,521128 | 0 | 0,0376581 | 0,0727748 | Rad54b | -2,67766387 | 1,60E-13 |
| Mogat2 | 23,0939 | 24,9052 | 7,67732 | 11,8276 | 0,222209 | 0,245984 | 0,300592 | 0 | 0 | 0 | Mogat2 | -4,00657751 | 2,01E-13 |
| Nup160 | 17,7508 | 24,9768 | 12,6805 | 16,5957 | 3,29357 | 3,34338 | 3,27266 | 2,17808 | 2,37861 | 2,19383 | Nup160 | -2,11464161 | 2,03E-13 |
| Galnt10 | 27,6134 | 17,4019 | 11,6097 | 19,1403 | 2,53676 | 1,46125 | 1,1641 | 1,17309 | 1,85935 | 1,44773 | Galnt10 | -3,1356442 | 2,05E-13 |
| Efnb2 | 38,4919 | 59,8484 | 43,7685 | 45,5817 | 4,85333 | 0,891714 | 0,886341 | 1,21469 | 2,28132 | 2,61428 | Efnb2 | -3,95631858 | 2,15E-13 |
| Pop1 | 14,4927 | 11,671 | 11,1919 | 8,54206 | 1,872 | 1,62301 | 1,66242 | 1,25517 | 0,924399 | 0,861634 | Pop1 | -2,31514293 | 2,27E-13 |
| Cdt1 | 73,0165 | 68,2785 | 40,2721 | 70,1874 | 7,30225 | 10,7594 | 9,41524 | 0,559998 | 0,621634 | 0,50003 | Cdt1 | -2,45685681 | 2,38E-13 |
| Gprc5c | 17,8327 | 14,0687 | 11,5779 | 13,3492 | 1,88428 | 0,729137 | 0,61794 | 1,75977 | 1,68605 | 1,7434 | Gprc5c | -3,58680106 | 2,40E-13 |
| Ptprn | 9,83423 | 6,33193 | 5,28112 | 3,69524 | 0,467697 | 0,160859 | 0,309187 | 3,08572 | 1,8722 | 1,92872 | Ptprn | -3,48250689 | 2,41E-13 |
| Syce2 | 19,2864 | 28,0645 | 19,7596 | 36,663 | 3,05778 | 1,08249 | 1,87333 | 1,01501 | 0,150189 | 0,185907 | Syce2 | -2,97106882 | 2,58E-13 |
| Grb7 | 10,4685 | 4,68156 | 3,26209 | 8,57707 | 0,118749 | 0,0350293 | 0,0428276 | 0 | 0 | 0 | Grb7 | -5,45974703 | 2,60E-13 |
| Sema4g | 3,42498 | 5,06811 | 5,76783 | 10,9524 | 0,611682 | 0,279754 | 0,319422 | 0,0677466 | 0,0497891 | 0 | Sema4g | -3,64376936 | 2,67E-13 |
| Maoa | 9,95072 | 11,6046 | 8,49919 | 7,53862 | 2,01048 | 1,68114 | 1,70199 | 1,33346 | 1,1608 | 0,926197 | Maoa | -2,04426667 | 2,89E-13 |
| Ddx25 | 11,0632 | 8,2366 | 4,5297 | 10,3018 | 0,355578 | 0,209967 | 0,256549 | 0 | 0 | 0 | Ddx25 | -4,4129012 | 3,00E-13 |
| Bex2 | 67,4589 | 87,8706 | 57,8748 | 70,4259 | 9,92144 | 5,80333 | 5,78755 | 0,601578 | 0,126852 | 0,291662 | Bex2 | -4,35204495 | 3,07E-13 |
| Krt19 | 40,8133 | 50,3907 | 52,0824 | 79,2861 | 2,31952 | 0,0663209 | 0,485963 | 0,47706 | 1,06927 | 0,257308 | Krt19 | -5,111023 | 3,23E-13 |
| Cdc7 | 12,1289 | 10,4007 | 9,25612 | 12,4644 | 1,89514 | 2,00316 | 2,11146 | 0,104841 | 0,116169 | 0,074829 | Cdc7 | -2,13588231 | 3,38E-13 |
| Npl | 83,4101 | 21,2664 | 9,63879 | 20,677 | 0,437528 | 0,517036 | 0,473599 | 0,465056 | 0,434067 | 0,167154 | Npl | -5,36179615 | 3,47E-13 |
| Cks1b | 160,22 | 173,343 | 100,028 | 119,505 | 32,1401 | 24,8007 | 21,387 | 7,10506 | 4,08808 | 4,83359 | Cks1b | -2,60243331 | 3,48E-13 |
| Astn1 | 1,30689 | 1,93389 | 2,46427 | 2,54588 | 0,0863 | 0 | 0 | 0 | 0,0726308 | 0,0562855 | Astn1 | -5,38485431 | 3,90E-13 |
| Tspan6 | 33,4647 | 68,16 | 34,5027 | 29,5428 | 6,64935 | 3,81035 | 3,26538 | 0,893004 | 1,12765 | 1,08729 | Tspan6 | -2,73634909 | 4,19E-13 |
| Prkcz | 1,56052 | 1,0263 | 1,55071 | 1,72201 | 0,11067 | 0,0782887 | 0,136627 | 0,338072 | 0,198765 | 0,216281 | Prkcz | -4,14766359 | 4,24E-13 |
| Phlda2 | 83,1959 | 30,0514 | 9,28806 | 30,657 | 0,167367 | 0 | 0 | 0 | 0 | 0,386455 | Phlda2 | -6,81254963 | 4,58E-13 |
| St8sia1 | 1,0916 | 1,48712 | 1,14986 | 0,663451 | 0,0441482 | 0,00867036 | 0,0127329 | 0,0315607 | 0,0192625 | 0,0335696 | St8sia1 | -5,05254649 | 4,63E-13 |
| Prmt8 | 2,49412 | 4,10068 | 4,85747 | 4,88541 | 0,0392939 | 0,0347729 | 0,0425146 | 0 | 0,0465176 | 0,0224597 | Prmt8 | -6,0879062 | 4,72E-13 |
| Prkar2b | 29,3495 | 14,3332 | 13,0171 | 12,0032 | 1,61956 | 0,728363 | 0,831471 | 0,0880881 | 0,291892 | 0,0313514 | Prkar2b | -3,5509998 | 5,27E-13 |
| Kcnc3 | 1,83742 | 1,6125 | 1,53375 | 1,60448 | 0,0712253 | 0 | 0,0385207 | 0,858272 | 1,22767 | 1,24025 | Kcnc3 | -5,00637564 | 5,27E-13 |
| Esrp1 | 1,86572 | 3,8411 | 1,82818 | 4,57491 | 0 | 0 | 0 | 0 | 0 | 0 | Esrp1 | -6,75858324 | 5,45E-13 |
| Sox2 | 2,1744 | 4,29003 | 7,25967 | 16,839 | 0,0385373 | 0 | 0 | 0 | 0,0456162 | 0 | Sox2 | -7,03370285 | 5,55E-13 |
| Hmgcll1 | 2,77559 | 3,88635 | 1,71892 | 2,52577 | 0,0253704 | 0,0224373 | 0 | 0 | 0 | 0 | Hmgcll1 | -5,77927194 | 5,87E-13 |
| Diap3 | 17,9834 | 23,5631 | 12,6245 | 10,3133 | 3,03144 | 2,44993 | 2,09199 | 0 | 0,0617393 | 0,0596884 | Diap3 | -2,40399046 | 6,26E-13 |
| Hey1 | 25,8681 | 27,7834 | 31,5563 | 24,9785 | 2,43136 | 0,384289 | 0,375827 | 0,139017 | 0,771514 | 0,695081 | Hey1 | -4,14417307 | 6,31E-13 |
| Lsr | 52,3198 | 23,7719 | 17,4841 | 21,4429 | 1,13406 | 0,217671 | 0,392675 | 0,435617 | 0,215024 | 0,0562548 | Lsr | -4,18906925 | 6,81E-13 |
| Nes | 39,9501 | 41,2394 | 57,4131 | 53,5845 | 5,21435 | 0,889203 | 1,08803 | 0,374945 | 1,94226 | 0,914887 | Nes | -3,90156861 | 7,00E-13 |
| Igf2 | 1343,97 | 1778,8 | 1895,42 | 1464,66 | 464,465 | 202,618 | 224,866 | 0,636935 | 1,6379 | 1,0454 | Igf2 | -2,23654041 | 7,10E-13 |
| Dusp5 | 69,4566 | 48,0246 | 36,8374 | 66,6684 | 6,64712 | 7,94385 | 6,3973 | 5,57836 | 7,829 | 5,82836 | Dusp5 | -2,55379023 | 7,53E-13 |
| Mylpf | 26,0478 | 27,3689 | 47,7198 | 46,7261 | 0,485125 | 0,767462 | 2,27819 | 0 | 0,586266 | 0,559844 | Mylpf | -4,18426305 | 7,76E-13 |
| Igf2os | 0 | 0 | 0 | 0 | 0 | 0 | 0 | 0 | 0 | 0 | Igf2os | -2,04339599 | 8,04E-13 |
| Plk1 | 64,6588 | 49,2745 | 44,121 | 46,1487 | 13,9075 | 7,42429 | 6,16039 | 0 | 0,360406 | 0,248495 | Plk1 | -2,49639079 | 8,04E-13 |
| Ccne1 | 39,7235 | 56,449 | 29,382 | 24,6774 | 3,02894 | 4,48415 | 4,78214 | 1,74735 | 0,883273 | 1,07937 | Ccne1 | -2,75230382 | 8,09E-13 |
| Cdk5rap2 | 20,8419 | 16,1587 | 14,2856 | 13,3738 | 3,47912 | 2,3332 | 2,57796 | 0,822622 | 1,61282 | 1,53328 | Cdk5rap2 | -2,18225751 | 8,14E-13 |
| Serp2 | 12,4232 | 15,4259 | 8,813 | 16,7829 | 0,317453 | 0,141227 | 0,344014 | 0 | 0 | 0 | Serp2 | -5,13151932 | 9,99E-13 |
| Egr4 | 33,4522 | 9,56519 | 3,507 | 7,16261 | 0,0916914 | 0,0405829 | 0,148823 | 0,146743 | 0,0543332 | 0 | Egr4 | -5,99341054 | 1,03E-12 |
| Kcnf1 | 4,52472 | 3,01952 | 1,55519 | 3,9555 | 0,113085 | 0 | 0,0203868 | 0 | 0 | 0 | Kcnf1 | -5,43455011 | 1,03E-12 |
| Rspo3 | 7,54075 | 26,0359 | 39,2796 | 31,9326 | 1,30684 | 0,210272 | 0,128542 | 0 | 0 | 0,0452718 | Rspo3 | -4,89248676 | 1,04E-12 |
| E2f7 | 14,4037 | 10,443 | 8,20861 | 8,39292 | 1,50721 | 0,73862 | 0,567044 | 0 | 0,0386414 | 0,0560854 | E2f7 | -3,07363044 | 1,04E-12 |
| Olfm1 | 6,68056 | 19,2361 | 13,9927 | 13,2504 | 3,02071 | 1,2936 | 1,52415 | 0,683526 | 0,844151 | 0,971022 | Olfm1 | -2,65103135 | 1,04E-12 |
| Elovl2 | 2,10108 | 2,40434 | 4,0688 | 3,38695 | 0,142872 | 0,0210576 | 0,0257585 | 0 | 0 | 0 | Elovl2 | -4,7820403 | 1,06E-12 |
| Siva1 | 46,9067 | 42,5457 | 33,7403 | 47,8383 | 11,3887 | 15,0594 | 14,1753 | 16,9853 | 18,3885 | 18,4599 | Siva1 | -2,15067372 | 1,09E-12 |
| Arg1 | 3,62345 | 10,0083 | 6,65311 | 12,1641 | 0,137238 | 0 | 0 | 0 | 0 | 0,0786197 | Arg1 | -5,59025448 | 1,37E-12 |
| Miat | 0,178959 | 1,37705 | 6,96135 | 5,6261 | 0,0101071 | 0 | 0 | 0 | 0 | 0,0115282 | Miat | -7,30975951 | 1,51E-12 |
| Axin2 | 11,8951 | 9,06082 | 7,35252 | 20,5559 | 1,25833 | 0,735481 | 0,992024 | 1,918 | 1,91316 | 1,8991 | Axin2 | -3,23382552 | 1,62E-12 |
| F5 | 6,89729 | 1,79498 | 2,54317 | 3,06984 | 0,0838666 | 0 | 0 | 0 | 0 | 0 | F5 | -5,96353646 | 1,79E-12 |
| Chek1 | 8,80208 | 14,3462 | 9,82278 | 11,7596 | 1,03029 | 2,03831 | 1,84781 | 0,0870099 | 0,0960965 | 0,247723 | Chek1 | -2,4092726 | 1,81E-12 |
| Nipsnap1 | 15,0599 | 34,0606 | 20,0808 | 27,3786 | 4,22177 | 4,24585 | 3,49693 | 0 | 0,309173 | 0,119282 | Nipsnap1 | -2,2491202 | 1,83E-12 |
| Nid2 | 128,252 | 99,1233 | 143,674 | 133,194 | 31,1876 | 15,493 | 15,6323 | 1,7204 | 2,75275 | 3,06853 | Nid2 | -2,19324217 | 2,08E-12 |
| Taf10 | 57,3515 | 50,6396 | 46,0744 | 44,3341 | 0 | 0 | 0 | 22,3305 | 20,9018 | 19,8924 | Taf10 | -2,11661575 | 2,45E-12 |
| Gpx3 | 342,994 | 363,988 | 563,738 | 540,916 | 109,4 | 55,1486 | 53,2609 | 80,4371 | 77,5392 | 66,2743 | Gpx3 | -2,31918102 | 2,55E-12 |
| Pole2 | 12,3548 | 14,6252 | 15,983 | 9,74763 | 0,81841 | 2,2264 | 1,39183 | 0 | 0,138853 | 0,267702 | Pole2 | -2,71756092 | 2,56E-12 |
| Nrp2 | 47,7015 | 62,9016 | 37,2445 | 28,5328 | 4,3027 | 0,742531 | 0,892941 | 1,24413 | 3,16001 | 1,99411 | Nrp2 | -3,97321818 | 2,99E-12 |
| Prtg | 128,459 | 62,1692 | 14,3198 | 14,9187 | 0,110836 | 0,0356198 | 0 | 0 | 0 | 0 | Prtg | -7,51466803 | 2,99E-12 |
| BC030867 | 9,81112 | 8,00093 | 6,33293 | 11,465 | 1,03858 | 1,0831 | 0,762496 | 0 | 0,438952 | 0,0847907 | BC030867 | -2,74761151 | 3,10E-12 |
| Dpy19l3 | 2,36699 | 3,5831 | 4,22417 | 3,98872 | 0,739499 | 0,653595 | 0,624858 | 0,0743237 | 0,136159 | 0,144988 | Dpy19l3 | -2,04721654 | 3,11E-12 |
| Pcyox1l | 16,2298 | 24,349 | 19,896 | 13,6235 | 4,31699 | 3,12113 | 3,26995 | 1,53486 | 1,93347 | 1,15227 | Pcyox1l | -2,01053995 | 3,29E-12 |
| Donson | 12,1368 | 17,8166 | 9,40664 | 13,9727 | 2,08987 | 2,71993 | 1,90651 | 1,31251 | 1,16459 | 0,983857 | Donson | -2,10871967 | 3,62E-12 |
| Rcc2 | 99,4984 | 105,679 | 80,5486 | 91,7112 | 27,609 | 14,2515 | 13,8692 | 4,91131 | 5,62023 | 5,18489 | Rcc2 | -2,1042834 | 3,63E-12 |
| Ppp1r17 | 5,22731 | 7,10574 | 12,024 | 6,44905 | 0 | 0,301822 | 0,184405 | 0 | 0 | 0 | Ppp1r17 | -4,6214252 | 4,10E-12 |
| Fbxl19 | 36,653 | 24,6589 | 25,0535 | 25,0554 | 5,96749 | 3,24798 | 3,30081 | 0,779907 | 1,14843 | 1,20271 | Fbxl19 | -2,37668268 | 4,20E-12 |

| Gene | d9-11 (1) | d9-11 (2) | d9-11(3) | d9-11(4) | newborn(1) | newborn(2) | newborn(3) | adult(1) | adult(2) | adult(3) | Gene | log2(fold) | FDR(*p* -value) |
|---|---|---|---|---|---|---|---|---|---|---|---|---|---|
| Prim1 | 30,2974 | 50,1923 | 36,096 | 30,2808 | 4,72938 | 7,6239 | 7,18054 | 0,419334 | 1,01536 | 0,752424 | Prim1 | -2,09641313 | 4,30E-12 |
| Aox4 | 1,40753 | 0,946726 | 3,4713 | 3,86772 | 0 | 0 | 0 | 0 | 0 | 0 | Aox4 | -7,32616295 | 4,55E-12 |
| Rab3b | 7,15762 | 3,28866 | 4,74684 | 2,66712 | 0,194508 | 0,147462 | 0,270539 | 0,0891647 | 0,229825 | 0,0952105 | Rab3b | -4,03008625 | 5,26E-12 |
| Dscc1 | 9,19895 | 16,2238 | 8,37516 | 13,755 | 0,871551 | 0,772669 | 1,2008 | 0 | 0 | 0 | Dscc1 | -3,17800642 | 5,34E-12 |
| Pfkfb3 | 16,9292 | 21,2372 | 13,8227 | 21,2398 | 2,80006 | 0,78973 | 0,757107 | 0,499047 | 0,778743 | 0,750323 | Pfkfb3 | -3,22535401 | 5,81E-12 |
| Lypd6 | 2,59878 | 3,11305 | 4,02687 | 3,08577 | 0,289309 | 0,0697832 | 0,113807 | 0 | 0,155343 | 0,120144 | Lypd6 | -3,84372513 | 5,82E-12 |
| Vnn1 | 3,5977 | 12,6948 | 9,2397 | 8,92114 | 0,863284 | 0,381978 | 0,551933 | 0,377147 | 0,0929072 | 0,134573 | Vnn1 | -3,41773494 | 6,19E-12 |
| Tmem163 | 6,61471 | 9,78799 | 24,5876 | 18,2582 | 0,600203 | 1,41627 | 0,933487 | 0 | 0,399562 | 0,128632 | Tmem163 | -3,48144611 | 6,28E-12 |
| Fermt3 | 24,65 | 18,111 | 18,5398 | 25,2215 | 3,42179 | 3,51083 | 3,54457 | 0 | 0 | 1,20652 | Fermt3 | -2,11042158 | 6,30E-12 |
| Zdhhc15 | 7,35953 | 7,37364 | 5,42331 | 3,18624 | 0,863025 | 0,40306 | 0,44032 | 0 | 0,0960169 | 0,092909 | Zdhhc15 | -3,01052716 | 6,60E-12 |
| Cfc1 | 40,7952 | 16,0967 | 10,2132 | 4,10723 | 0 | 0 | 0 | 0 | 0 | 0 | Cfc1 | -7,40198985 | 6,75E-12 |
| Emb | 50,1005 | 44,8057 | 18,4802 | 21,667 | 1,43484 | 0,0952165 | 0,116428 | 0,114976 | 0,169765 | 0 | Emb | -5,21610555 | 7,24E-12 |
| Cnpy4 | 22,7664 | 24,5699 | 15,5911 | 13,5234 | 3,20342 | 2,00539 | 2,21152 | 0,882767 | 1,31118 | 0,821808 | Cnpy4 | -2,55610273 | 7,42E-12 |
| H2afx | 163,69 | 132,925 | 115,438 | 128,178 | 37,1587 | 24,0516 | 21,1898 | 11,2163 | 11,4957 | 12,69 | H2afx | -2,01324871 | 7,67E-12 |
| Susd4 | 6,36065 | 14,0354 | 6,14719 | 14,6688 | 0,569881 | 0,336333 | 0,0513881 | 0 | 0,0562975 | 0 | Susd4 | -3,98653192 | 7,76E-12 |
| Fam57a | 10,2506 | 16,0097 | 21,3847 | 24,5072 | 2,92065 | 1,51333 | 1,44992 | 0,38544 | 0,292143 | 0,700015 | Fam57a | -2,75066498 | 7,77E-12 |
| Snrpd1 | 57,7234 | 75,5951 | 45,7066 | 59,0872 | 17,515 | 18,8511 | 20,1903 | 10,1816 | 11,2195 | 7,82945 | Snrpd1 | -2,19534858 | 7,99E-12 |
| Slc1a2 | 2,54043 | 1,13648 | 2,6048 | 1,64553 | 0,00887444 | 0,0635224 | 0,0505783 | 0,0257004 | 0,0188125 | 0,0516895 | Slc1a2 | -4,29205921 | 8,49E-12 |
| Gyltl1b | 3,97202 | 4,33816 | 1,43558 | 9,09614 | 0,0406033 | 0 | 0,0439356 | 0 | 0 | 0 | Gyltl1b | -5,85150685 | 9,22E-12 |
| Apoa1 | 17,3775 | 18,6999 | 3,29578 | 67,99 | 0 | 0,0998597 | 0 | 0 | 0,134789 | 0 | Apoa1 | -6,39492793 | 9,67E-12 |
| Scube3 | 2,97105 | 8,25 | 8,81749 | 15,7418 | 0,249526 | 0,0275919 | 0 | 0,200079 | 0,110625 | 0,0712678 | Scube3 | -5,41871818 | 9,75E-12 |
| Mpp2 | 23,5052 | 14,6572 | 17,6516 | 29,0678 | 1,64809 | 3,13132 | 2,74644 | 1,0013 | 0,42066 | 1,06792 | Mpp2 | -2,72658895 | 1,06E-11 |
| **Hoxb8** | 2,75699 | 2,59611 | 4,60242 | 2,7774 | 0,0355303 | 0 | 0 | 0 | 0 | 0 | Hoxb8 | -6,36172328 | 1,15E-11 |
| Lin28b | 11,3215 | 12,0367 | 3,23174 | 5,85136 | 0,382112 | 0,660914 | 0,413308 | 0,0533943 | 0 | 0,0189589 | Lin28b | -3,52300043 | 1,20E-11 |
| Oit3 | 58,0775 | 24,8629 | 8,84162 | 3,92507 | 0,517778 | 0,152702 | 0,112036 | 0 | 0 | 0 | Oit3 | -5,6143572 | 1,23E-11 |
| Prdm6 | 15,7412 | 13,3461 | 6,39188 | 12,6004 | 1,15801 | 0,256156 | 0,391522 | 0,231971 | 0,556637 | 0 | Prdm6 | -3,88118775 | 1,24E-11 |
| Shc2 | 11,8765 | 7,38277 | 7,80855 | 6,80243 | 0,668358 | 0,0815207 | 0,124653 | 0 | 0,10884 | 0,263134 | Shc2 | -4,33570062 | 1,30E-11 |
| Rad51 | 21,6489 | 32,5031 | 18,2458 | 17,7119 | 2,58382 | 4,33559 | 4,03569 | 0,143849 | 0,212988 | 0,359741 | Rad51 | -2,44375179 | 1,42E-11 |
| Soat2 | 4,14447 | 5,18918 | 2,2618 | 4,12128 | 0,0452194 | 0 | 0 | 0,289502 | 0 | 0 | Soat2 | -5,03368345 | 1,53E-11 |
| Edn1 | 20,8866 | 25,8259 | 12,7765 | 12,3936 | 3,04291 | 1,7953 | 1,93152 | 3,63884 | 1,6334 | 1,20601 | Edn1 | -2,61783545 | 1,54E-11 |
| Ugdh | 54,7021 | 53,8954 | 42,4624 | 32,5123 | 11,2463 | 6,72627 | 7,4267 | 3,23052 | 3,25931 | 2,65984 | Ugdh | -2,12323952 | 1,69E-11 |
| Ect2 | 23,671 | 29,2682 | 20,1429 | 24,6419 | 6,88818 | 3,56259 | 3,05573 | 0,14687 | 0,107998 | 0,182772 | Ect2 | -2,12106878 | 1,78E-11 |
| Ccnf | 28,2929 | 22,8076 | 21,413 | 29,3546 | 4,18907 | 1,27047 | 1,26245 | 0,383949 | 0,176838 | 0,170905 | Ccnf | -3,0988371 | 1,79E-11 |
| Prom1 | 9,55631 | 18,2471 | 13,9959 | 12,954 | 0,59781 | 0,0314747 | 0,0384867 | 0 | 0,0920196 | 0,122093 | Prom1 | -5,13108613 | 1,82E-11 |
| Sema5b | 1,89307 | 2,35169 | 4,44794 | 4,02662 | 0,218471 | 0,245846 | 0,19335 | 0,382813 | 0,328108 | 0,317386 | Sema5b | -3,42946034 | 1,82E-11 |
| Kif2c | 22,124 | 23,6766 | 19,1956 | 18,2412 | 5,19196 | 2,48225 | 2,35234 | 0 | 0,0792345 | 0,0382759 | Kif2c | -2,25763148 | 1,85E-11 |
| **Hoxb3** | 5,94492 | 5,14148 | 8,1637 | 4,05922 | 0,501297 | 0,0492655 | 0,0602559 | 0 | 0 | 0,159044 | Hoxb3 | -4,35907666 | 1,92E-11 |
| Hsd11b2 | 37,8539 | 20,9703 | 13,1514 | 7,13734 | 0,582141 | 0 | 0,0572699 | 0 | 0 | 0 | Hsd11b2 | -5,64968412 | 1,97E-11 |
| Ttll12 | 36,9914 | 27,7068 | 23,0863 | 24,5871 | 4,41565 | 5,51413 | 5,77395 | 3,81669 | 3,95374 | 4,29399 | Ttll12 | -2,10557145 | 2,22E-11 |
| Fzd10 | 1,86661 | 5,41587 | 3,39105 | 9,29189 | 0,155615 | 0,027532 | 0 | 0,0998236 | 0 | 0 | Fzd10 | -5,43063306 | 2,25E-11 |
| Stil | 8,86872 | 10,4518 | 7,99918 | 6,29706 | 1,60147 | 0,923981 | 0,547018 | 0 | 0 | 0 | Stil | -2,68496341 | 2,29E-11 |
| Npnt | 3,59373 | 13,3281 | 16,3305 | 13,1674 | 0,941317 | 0,225011 | 0,113136 | 0,0671884 | 0,0740644 | 0,119396 | Npnt | -4,23842903 | 2,56E-11 |
| Pdia5 | 35,9381 | 53,5551 | 29,038 | 24,6457 | 6,34801 | 3,65153 | 2,75961 | 0,693842 | 1,80296 | 1,49029 | Pdia5 | -2,64607904 | 2,72E-11 |
| Grhl1 | 2,10769 | 6,45302 | 10,3113 | 8,39136 | 0,450834 | 0,304913 | 0,258196 | 0,383816 | 0,125313 | 0,242292 | Grhl1 | -3,78760177 | 2,79E-11 |
| Nkd1 | 13,5146 | 13,5519 | 13,5743 | 11,0727 | 2,20876 | 0,706804 | 0,728143 | 0 | 0,0785271 | 0 | Nkd1 | -2,94632734 | 2,80E-11 |
| Ina | 3,33165 | 0 | 7,94579 | 9,20451 | 0 | 0 | 0 | 0 | 0 | 0 | Ina | -6,09727668 | 2,86E-11 |
| Slc9a3r1 | 73,0002 | 42,4331 | 13,4074 | 28,4056 | 2,9376 | 0,896878 | 0,767309 | 1,62006 | 2,2227 | 1,0431 | Slc9a3r1 | -4,18155425 | 2,97E-11 |
| Rims3 | 18,4306 | 9,48544 | 6,68705 | 6,85708 | 0 | 0,101324 | 0 | 0 | 0 | 0 | Rims3 | -6,46222628 | 3,04E-11 |
| Tipin | 50,5334 | 70,6942 | 41,4062 | 48,726 | 6,48816 | 10,62 | 8,67439 | 3,32593 | 5,81994 | 6,69495 | Tipin | -2,20128153 | 3,07E-11 |
| Ebf4 | 2,23272 | 2,44728 | 5,38377 | 4,99834 | 0,457176 | 0,248922 | 0,266334 | 0 | 0,0416038 | 0,0803785 | Ebf4 | -3,27433758 | 3,18E-11 |
| Slc29a4 | 2,83918 | 3,38432 | 3,35725 | 3,81366 | 0,268293 | 0 | 0,0725652 | 0 | 0,0396638 | 0,0383208 | Slc29a4 | -4,23821859 | 3,52E-11 |
| Ephb2 | 28,9905 | 14,3158 | 9,64675 | 16,9113 | 0,948551 | 0,0492939 | 0,180926 | 0,238856 | 0,164445 | 0,0636358 | Ephb2 | -4,85824456 | 3,75E-11 |
| E2f5 | 21,653 | 16,8529 | 11,0534 | 17,4074 | 2,65031 | 2,497 | 2,25793 | 0,901131 | 1,60672 | 1,74277 | E2f5 | -2,37851929 | 3,81E-11 |
| Zfp503 | 25,2763 | 10,8903 | 13,6632 | 15,8015 | 1,38203 | 0,276535 | 0,139984 | 0,289106 | 0,152748 | 0,246224 | Zfp503 | -4,26534613 | 3,85E-11 |
| Ttk | 15,1269 | 22,9708 | 9,92415 | 13,1187 | 2,72834 | 1,41979 | 1,73629 | 0 | 0 | 0,0394694 | Ttk | -2,57345129 | 4,19E-11 |
| Rian | 58,233 | 87,0403 | 89,3151 | 49,324 | 11,8469 | 2,76674 | 2,63524 | 0,164617 | 0,151439 | 0,146414 | Rian | -3,56784816 | 4,39E-11 |
| F10 | 3,46072 | 2,89015 | 2,00017 | 3,8215 | 0,113229 | 0 | 0,0376933 | 0 | 0 | 0 | F10 | -4,90584585 | 4,56E-11 |
| Psrc1 | 28,9878 | 22,0922 | 25,1757 | 27,3904 | 4,42739 | 1,2041 | 1,71962 | 0 | 0,0673255 | 0,0649105 | Psrc1 | -2,9743911 | 4,92E-11 |
| Metrn | 30,2797 | 39,5641 | 32,1429 | 70,0811 | 5,43452 | 6,15522 | 6,6181 | 22,857 | 20,137 | 21,9776 | Metrn | -2,4811108 | 4,94E-11 |
| Usp29 | 2,88607 | 2,13095 | 2,60065 | 1,26677 | 0,224046 | 0,177222 | 0,178093 | 0 | 0 | 0,0222876 | Usp29 | -3,04264531 | 4,96E-11 |
| Tagln3 | 8,68905 | 5,39672 | 23,6343 | 22,0349 | 0,183071 | 0,162391 | 0,297319 | 0 | 0,327945 | 0,630373 | Tagln3 | -4,86643578 | 4,96E-11 |
| Capn6 | 76,3341 | 91,3844 | 106,537 | 89,0112 | 21,2729 | 8,31939 | 6,37396 | 0 | 0,0915756 | 0 | Capn6 | -2,58311296 | 5,00E-11 |
| Tbx1 | 3,69474 | 3,8268 | 7,95285 | 2,34942 | 0 | 0,0488805 | 0,0596661 | 0 | 0,19655 | 0,18929 | Tbx1 | -5,5050307 | 5,03E-11 |
| **Hoxb4** | 3,10881 | 3,70571 | 2,8111 | 2,10961 | 0,146913 | 0 | 0,0397373 | 0,117709 | 0,0434617 | 0,0839561 | Hoxb4 | -4,89367228 | 5,30E-11 |
| Hcn4 | 29,3957 | 19,6119 | 15,0341 | 26,9627 | 2,86455 | 4,00206 | 3,20287 | 4,17332 | 7,64633 | 8,65803 | Hcn4 | -2,42168083 | 5,43E-11 |
| Nefm | 2,22154 | 2,29115 | 24,2747 | 15,0565 | 0 | 0,0759406 | 0,0309598 | 0 | 0,135278 | 0,326888 | Nefm | -6,23294792 | 5,70E-11 |
| Gins2 | 51,3524 | 67,7024 | 42,6361 | 39,2306 | 0 | 0 | 0 | 0 | 0 | 0 | Gins2 | -2,2833199 | 5,76E-11 |
| Acat2 | 31,3743 | 32,0536 | 34,6576 | 41,2221 | 5,51082 | 6,52614 | 6,4195 | 2,52673 | 3,79792 | 3,50572 | Acat2 | -2,34957837 | 6,55E-11 |
| Tspan14 | 43,5069 | 36,8067 | 37,1012 | 24,7635 | 9,23052 | 6,15517 | 5,82687 | 3,05965 | 3,43586 | 2,4886 | Tspan14 | -2,03953155 | 6,67E-11 |
| Cck | 25,7127 | 10,145 | 11,7992 | 21,9901 | 0 | 0 | 0 | 0 | 0,22294 | 0 | Cck | -6,32284715 | 6,94E-11 |
| Sept9 | 122,866 | 125,314 | 94,3051 | 97,4924 | 22,5126 | 6,93166 | 7,36987 | 8,81849 | 11,7669 | 10,4862 | 09. Sep | -2,75960458 | 6,94E-11 |
| Foxred2 | 1,72853 | 2,22987 | 1,41259 | 1,87554 | 0,169479 | 0,033298 | 0,040738 | 0 | 0,0444342 | 0,0214931 | Foxred2 | -4,0342167 | 8,00E-11 |
| Nr6a1 | 22,8869 | 10,2222 | 3,65921 | 4,26287 | 0,490774 | 0,36367 | 0,263483 | 0,232026 | 0,153159 | 0,230573 | Nr6a1 | -4,23176278 | 8,18E-11 |
| Rbp4 | 14,9488 | 8,00486 | 5,34616 | 52,8839 | 0 | 0,108095 | 0 | 0 | 0 | 0 | Rbp4 | -6,67173987 | 9,05E-11 |
| Dpp6 | 1,74799 | 0,973955 | 2,53118 | 2,99688 | 0,0599286 | 0,0353248 | 0 | 0 | 0 | 0 | Dpp6 | -5,13105911 | 9,46E-11 |
| Flnb | 44,3967 | 31,8496 | 24,9696 | 19,536 | 3,91711 | 0,95468 | 0,863092 | 0,782691 | 1,67901 | 1,35084 | Flnb | -3,51939325 | 9,76E-11 |
| Emid1 | 6,28755 | 5,67087 | 8,09672 | 11,9427 | 1,07054 | 0,767325 | 0,717147 | 0,163055 | 0,241871 | 0,174987 | Emid1 | -2,86467521 | 9,77E-11 |
| 4930503L19Ri | 10,2701 | 12,3023 | 8,57215 | 7,53752 | 2,1222 | 0,773392 | 0,990525 | 0,13327 | 0,295684 | 0,380621 | 4930503L19Ri | -2,58818417 | 1,02E-10 |
| Depdc1b | 9,99784 | 9,21387 | 5,49012 | 5,69846 | 1,04444 | 0,858171 | 0,766794 | 0,119536 | 0 | 0 | Depdc1b | -2,72196713 | 1,03E-10 |
| Wfdc2 | 19,8572 | 34,889 | 5,17745 | 13,7363 | 0,177362 | 0 | 0 | 0,557501 | 0,214906 | 0,409842 | Wfdc2 | -6,36185212 | 1,08E-10 |
| Cmip | 33,3793 | 32,4414 | 33,8118 | 38,8924 | 9,84514 | 4,31263 | 4,47598 | 0,963238 | 1,29741 | 1,09527 | Cmip | -2,14082151 | 1,08E-10 |
| Ccne2 | 12,5682 | 20,6951 | 11,8605 | 11,4126 | 0 | 1,37195 | 1,445 | 1,82176 | 0 | 0,995141 | Ccne2 | -2,75878833 | 1,20E-10 |
| Dna2 | 14,8804 | 22,5957 | 13,5264 | 15,2278 | 1,16979 | 2,43961 | 2,5069 | 0,212645 | 0,156326 | 0,277178 | Dna2 | -2,48005331 | 1,22E-10 |
| Spire2 | 4,02648 | 3,26276 | 1,92045 | 3,76638 | 0,0407836 | 0 | 0,0441272 | 0 | 0,0482932 | 0,0466276 | Spire2 | -5,67764582 | 1,34E-10 |
| Tmem132e | 5,86865 | 1,85261 | 1,24099 | 1,4979 | 0,0221682 | 0,0196025 | 0 | 0 | 0 | 0 | Tmem132e | -5,81002195 | 1,34E-10 |
| Nrgn | 29,1392 | 7,53494 | 17,944 | 13,6746 | 0,723748 | 0,0712966 | 0,0870548 | 0,256162 | 0,0958569 | 0 | Nrgn | -4,30316631 | 1,42E-10 |
| Nkx2-6 | 7,78123 | 7,99556 | 8,38781 | 4,24395 | 0 | 0 | 0 | 0 | 0 | 0 | Nkx2-6 | -6,86975259 | 1,43E-10 |
| Unc13d | 2,19048 | 1,9208 | 2,57334 | 2,61563 | 0,229853 | 0,101628 | 0,0994556 | 0,0738065 | 0,0814113 | 0,052486 | Unc13d | -3,35042623 | 1,48E-10 |
| BC018242 | 7,38093 | 7,02632 | 11,2644 | 10,0147 | 1,70054 | 1,72404 | 1,72484 | 0,567813 | 1,17359 | 1,19587 | BC018242 | -2,04859034 | 1,51E-10 |
| Soga3 | 0,97903 | 1,44872 | 3,13264 | 2,30156 | 0,0231149 | 0 | 0 | 0 | 0 | 0 | Soga3 | -6,18322532 | 1,57E-10 |
| Adra2a | 2,97106 | 2,00979 | 1,48799 | 1,53944 | 0,14431 | 0,063809 | 0,0260178 | 0 | 0,113604 | 0,0274627 | Adra2a | -4,33445381 | 1,83E-10 |
| Nell2 | 1,34032 | 2,13588 | 6,71256 | 8,30962 | 0,029221 | 0 | 0 | 0 | 0,0345349 | 0 | Nell2 | -5,91868179 | 1,89E-10 |
| Map3k9 | 1,23589 | 0,941662 | 0,829968 | 1,71474 | 0,0910598 | 0,0365772 | 0,155171 | 0 | 0,0319849 | 0 | Map3k9 | -3,44854563 | 2,11E-10 |
| Adra2b | 1,71791 | 2,25962 | 2,45822 | 2,22533 | 0,139063 | 0 | 0 | 0 | 0,0820936 | 0,0264625 | Adra2b | -4,87210078 | 2,42E-10 |
| Alg13 | 12,075 | 11,9869 | 7,35208 | 7,57497 | 2,61748 | 2,27859 | 2,29674 | 2,42238 | 1,87686 | 2,4277 | Alg13 | -2,33657269 | 2,50E-10 |
| Cad | 64,2252 | 42,2824 | 34,3465 | 29,7656 | 6,48492 | 6,83932 | 7,10821 | 4,20283 | 3,96081 | 4,37351 | Cad | -2,25574929 | 2,90E-10 |
| Smad7 | 32,3384 | 31,4755 | 36,4322 | 31,8564 | 9,60954 | 4,05891 | 3,88418 | 4,98078 | 4,76732 | 4,93736 | Smad7 | -2,19436982 | 3,22E-10 |
| Fen1 | 46,846 | 38,6152 | 28,4638 | 30,6478 | 6,24196 | 7,92296 | 7,46595 | 4,22049 | 3,27611 | 3,70894 | Fen1 | -2,02340607 | 3,29E-10 |
| Car3 | 22,0384 | 12,1287 | 12,1978 | 3,50459 | 0,592983 | 0,0583765 | 0,142625 | 0 | 1,09685 | 0,075474 | Car3 | -4,72520649 | 3,44E-10 |
| Haus6 | 12,334 | 15,274 | 9,4191 | 11,1045 | 3,17259 | 1,62206 | 1,85005 | 0,477409 | 1,1416 | 1,10381 | Haus6 | -2,02788633 | 3,65E-10 |
| Jag1 | 31,021 | 32,8776 | 23,632 | 25,3881 | 4,27842 | 0,736235 | 0,989125 | 0,734783 | 0,731811 | 0,409964 | Jag1 | -3,41854166 | 3,72E-10 |
| Lrp8 | 1,58791 | 2,14001 | 2,28148 | 2,8495 | 0,293753 | 0,139804 | 0,259525 | 0,0782574 | 0,106009 | 0,274009 | Lrp8 | -2,69055742 | 4,04E-10 |
| Cldn7 | 3,06071 | 8,10697 | 4,58113 | 14,5704 | 0,0790863 | 0 | 0 | 0 | 0,282751 | 0,129129 | Cldn7 | -5,79767846 | 4,09E-10 |
| Eml4 | 60,0932 | 43,816 | 25,1771 | 22,9977 | 5,73721 | 2,63067 | 2,50901 | 0,953749 | 1,16659 | 1,14762 | Eml4 | -2,98066923 | 4,25E-10 |
| Cpa2 | 7,41765 | 9,75623 | 8,48344 | 7,19315 | 1,17123 | 0,346097 | 0,676198 | 0 | 0,0930375 | 0 | Cpa2 | -3,05609185 | 4,33E-10 |
| Cntfr | 29,3738 | 25,4087 | 32,682 | 42,7427 | 2,04424 | 0,0868696 | 0,159265 | 1,18655 | 0,290871 | 0,194109 | Cntfr | -4,7167126 | 4,57E-10 |
| Tmem56 | 5,87644 | 5,54726 | 2,74864 | 1,5313 | 0,358637 | 0,20289 | 0,201711 | 0,415076 | 1,25152 | 0,720184 | Tmem56 | -3,56691782 | 4,73E-10 |
| Tnnt3 | 18,7623 | 15,0375 | 24,1381 | 12,5926 | 2,00351 | 1,08703 | 0,843803 | 0,353045 | 0,133369 | 0,767697 | Tnnt3 | -3,2124069 | 4,89E-10 |
| Ston2 | 2,7028 | 1,86024 | 1,28548 | 3,64181 | 0,249113 | 0,157249 | 0,0769782 | 0,744241 | 1,10732 | 0,735649 | Ston2 | -3,42265194 | 4,93E-10 |
| Pde1b | 3,65983 | 6,01623 | 8,06941 | 5,94184 | 0,949139 | 0,330593 | 0,279382 | 0 | 0,102313 | 0,0328705 | Pde1b | -3,09507469 | 5,16E-10 |
| Kcnh7 | 1,35934 | 2,43053 | 4,32586 | 2,0543 | 0,0722154 | 0,0425751 | 0,0781188 | 0 | 0 | 0 | Kcnh7 | -4,01979252 | 5,43E-10 |
| Alx4 | 0,427098 | 1,20381 | 2,40645 | 4,54919 | 0,0172807 | 0 | 0,0186917 | 0,1666 | 0,0203828 | 0,15777 | Alx4 | -5,96142219 | 5,70E-10 |
| Tulp1 | 4,58757 | 6,03405 | 3,68721 | 4,96332 | 0,385667 | 0,0853578 | 0,0521658 | 0 | 0 | 0 | Tulp1 | -4,13831366 | 6,07E-10 |

| Gene | d9-11 (1) | d9-11 (2) | d9-11(3) | d9-11(4) | newborn(1) | newborn(2) | newborn(3) | adult(1) | adult(2) | adult(3) | Gene | log2(fold) | FDR(*p* -value) |
|---|---|---|---|---|---|---|---|---|---|---|---|---|---|
| Lhx1 | 7,80786 | 2,48846 | 3,0079 | 0,968095 | 0,0510812 | 0 | 0 | 0 | 0 | 0 | Lhx1 | -6,31397446 | 6,50E-10 |
| Hhex | 62,1783 | 35,1602 | 11,6868 | 8,87637 | 1,61206 | 0,344601 | 0,0601573 | 0,533055 | 0,263946 | 0,445382 | Hhex | -4,94199592 | 6,91E-10 |
| C330027C09R | 22,5118 | 31,794 | 15,0733 | 16,7251 | 4,65731 | 3,97639 | 3,87153 | 0,221008 | 0,243776 | 0,471491 | C330027C09R | -2,01224222 | 7,17E-10 |
| Pitx2 | 7,38984 | 13,7223 | 18,2968 | 14,4505 | 2,13695 | 2,50952 | 2,58753 | 0,867845 | 0,450432 | 0,2512 | Pitx2 | -2,0753972 | 7,25E-10 |
| Pou3f3 | 0,961818 | 1,15968 | 3,74652 | 9,58216 | 0 | 0,00297717 | 0,00655388 | 0 | 0 | 0 | Pou3f3 | -6,87086626 | 7,56E-10 |
| Fscn1 | 484,845 | 281,782 | 204,419 | 212,627 | 29,5064 | 4,08112 | 4,03801 | 2,82173 | 4,79035 | 4,78841 | Fscn1 | -4,07187277 | 7,69E-10 |
| BC005764 | 0 | 0 | 16,6179 | 0 | 0 | 1,84265 | 0 | 0 | 0 | 0 | BC005764 | -2,03369968 | 7,71E-10 |
| Frmpd1 | 2,31724 | 3,26566 | 4,91853 | 3,73558 | 0,487579 | 0,53056 | 0,608539 | 0,120505 | 0,0663745 | 0,042808 | Frmpd1 | -2,34595326 | 8,24E-10 |
| Stab2 | 14,18 | 8,30221 | 3,81723 | 0,542092 | 0,0870385 | 0,125004 | 0,0823701 | 0,209968 | 0,0640962 | 0,148923 | Stab2 | -4,8025469 | 8,47E-10 |
| Hspa1a | 68,7965 | 18,7039 | 35,8334 | 139,054 | 4,0326 | 1,62615 | 1,83792 | 13,8229 | 14,3013 | 13,238 | Hspa1a | -4,19691909 | 8,54E-10 |
| Ptpru | 1,90965 | 3,88193 | 4,10587 | 7,3466 | 0,622779 | 0,318742 | 0,248176 | 0,737436 | 1,04371 | 0,41145 | Ptpru | -3,09220614 | 9,57E-10 |
| Dll1 | 6,82206 | 5,70174 | 6,56438 | 12,2188 | 0,751817 | 0,0949916 | 0,116186 | 0,603201 | 0,570978 | 0,429314 | Dll1 | -4,03905966 | 9,77E-10 |
| Rem2 | 3,64454 | 2,8762 | 3,65027 | 6,05717 | 0,155141 | 0,183157 | 0,167884 | 0 | 0 | 0 | Rem2 | -4,03653921 | 9,79E-10 |
| Hes1 | 89,0431 | 62,5325 | 44,4656 | 55,3951 | 12,0873 | 9,74901 | 11,756 | 37,3293 | 33,2689 | 54,1175 | Hes1 | -2,09672131 | 1,07E-09 |
| Mob3a | 39,4425 | 24,9515 | 23,0977 | 21,8225 | 5,57366 | 2,35582 | 2,07408 | 0,227591 | 1,05002 | 0,608574 | Mob3a | -2,70334868 | 1,11E-09 |
| Vil1 | 2,41305 | 0 | 0 | 4,96776 | 0 | 0 | 0 | 0 | 0 | 0 | Vil1 | -3,75563961 | 1,26E-09 |
| Agpat4 | 24,446 | 23,4948 | 18,3034 | 16,5667 | 2,3719 | 0,316145 | 0,551979 | 0,489451 | 0,907559 | 0,233455 | Agpat4 | -3,72035883 | 1,36E-09 |
| Dok5 | 4,31676 | 11,0869 | 11,7459 | 10,0793 | 0,721249 | 0,0492294 | 0,18056 | 0 | 0,922009 | 0,573751 | Dok5 | -4,13096648 | 1,40E-09 |
| Tubb4a | 9,38149 | 7,46344 | 15,4082 | 6,70669 | 1,1159 | 0,202599 | 0,278635 | 0,137432 | 0,15252 | 0 | Tubb4a | -3,77924822 | 1,42E-09 |
| Acvr2b | 23,9944 | 19,6236 | 13,6932 | 34,2886 | 3,959 | 1,75324 | 2,3313 | 0,558093 | 2,21254 | 1,33305 | Acvr2b | -2,76108443 | 1,46E-09 |
| Dnmt3b | 31,517 | 22,6461 | 4,74663 | 10,1053 | 0,978977 | 0,599944 | 0,538404 | 0 | 0,181122 | 0,22522 | Dnmt3b | -3,8445669 | 1,47E-09 |
| Lor | 3,70184 | 5,38155 | 5,20368 | 5,1023 | 0,553024 | 0,0979499 | 0,119696 | 0 | 0,0656442 | 0,0632976 | Lor | -3,82627063 | 1,54E-09 |
| Tkt | 101,137 | 93,4015 | 71,3544 | 76,4521 | 21,4686 | 7,34581 | 8,25629 | 2,94027 | 4,26411 | 4,21966 | Tkt | -2,35775384 | 1,56E-09 |
| P2rx1 | 2,73634 | 10,1226 | 2,85494 | 9,37216 | 0,310383 | 0,137333 | 0,293844 | 0 | 0,0459267 | 0 | P2rx1 | -4,01669753 | 1,59E-09 |
| Rgs19 | 28,9247 | 21,4237 | 16,6524 | 28,8034 | 4,07923 | 3,53969 | 3,96778 | 2,54713 | 1,55597 | 1,74213 | Rgs19 | -2,00793116 | 1,60E-09 |
| Otx2 | 7,32552 | 1,86533 | 0,657313 | 10,9159 | 0 | 0 | 0 | 0 | 0 | 0 | Otx2 | -6,78771666 | 1,81E-09 |
| Rims2 | 0,859016 | 0,832966 | 1,10543 | 1,04183 | 0,0743637 | 0,0431264 | 0,0264691 | 0,0785537 | 0,014362 | 0 | Rims2 | -4,00376703 | 1,86E-09 |
| Sel1l3 | 1,48748 | 5,31058 | 8,39577 | 5,63793 | 0,200647 | 0,319333 | 0,412372 | 0,12891 | 1,23123 | 0,458067 | Sel1l3 | -3,4856028 | 1,95E-09 |
| Srm | 141,65 | 151,489 | 61,2816 | 80,1526 | 30,2913 | 13,9097 | 11,8856 | 8,6848 | 6,72667 | 10,2675 | Srm | -2,58586427 | 1,97E-09 |
| Syne4 | 5,35431 | 8,18677 | 10,543 | 8,21483 | 0,584995 | 0,173115 | 0,739413 | 0,242494 | 0 | 0 | Syne4 | -2,61039876 | 2,04E-09 |
| Plk4 | 11,6172 | 16,9784 | 11,1375 | 11,3355 | 3,98995 | 2,18534 | 1,69134 | 0,510994 | 0,642791 | 0,71009 | Plk4 | -2,22954104 | 2,12E-09 |
| Ryr1 | 0,281184 | 0,396274 | 1,14007 | 0,566679 | 0,017054 | 0,010046 | 0 | 0,0548939 | 0,00669226 | 0,0583358 | Ryr1 | -4,8039627 | 2,17E-09 |
| Etv2 | 22,7167 | 26,0227 | 3,05765 | 1,84445 | 0 | 0 | 0 | 0 | 0 | 0 | Etv2 | -6,77394024 | 2,28E-09 |
| Lrch2 | 8,41383 | 11,0375 | 7,91965 | 4,59915 | 1,42032 | 0,515905 | 0,466425 | 0,162828 | 0,14978 | 0,0289629 | Lrch2 | -2,8350834 | 2,36E-09 |
| Klf5 | 20,7869 | 10,9496 | 11,4539 | 26,73 | 1,94787 | 0,5074 | 0,620008 | 0,549248 | 0,272106 | 0,459081 | Klf5 | -3,57729604 | 2,42E-09 |
| Gcat | 9,20649 | 17,4778 | 14,2511 | 21,7525 | 2,63274 | 1,33295 | 1,20937 | 9,97914 | 10,7333 | 7,87031 | Gcat | -2,79463829 | 2,54E-09 |
| Gata1 | 2,88967 | 5,87933 | 3,16555 | 2,72875 | 0,153755 | 0 | 0 | 0 | 0 | 0 | Gata1 | -5,13094174 | 2,54E-09 |
| Fads2 | 51,6313 | 73,2588 | 69,0649 | 62,9216 | 10,9151 | 2,01733 | 2,10188 | 0 | 0,716772 | 0,230141 | Fads2 | -3,21397592 | 2,60E-09 |
| Pola1 | 12,5125 | 17,755 | 13,3156 | 12,3682 | 1,31842 | 2,74681 | 3,14277 | 0,809646 | 1,10927 | 1,07332 | Pola1 | -2,17827906 | 2,77E-09 |
| **Hoxb5** | 7,43125 | 2,86078 | 2,11791 | 4,01646 | 0,102871 | 0,0455424 | 0 | 0 | 0 | 0 | Hoxb5 | -5,30009037 | 3,19E-09 |
| Twist1 | 33,1702 | 33,3588 | 36,4933 | 32,3237 | 3,36369 | 0,32505 | 0,264759 | 0 | 0,072666 | 0 | Twist1 | -4,15051365 | 3,22E-09 |
| St8sia2 | 3,06691 | 4,65465 | 4,5293 | 1,84219 | 0,417596 | 0,0886012 | 0,0542027 | 0,0536837 | 0,0394009 | 0,0190621 | St8sia2 | -3,77860511 | 3,26E-09 |
| Insm1 | 1,19316 | 1,17705 | 2,62562 | 4,80818 | 0 | 0 | 0 | 0 | 0 | 0 | Insm1 | -6,57317165 | 3,26E-09 |
| Acp5 | 2,41156 | 11,0929 | 10,2554 | 11,5697 | 0,533031 | 0,202496 | 0,164857 | 0 | 0,181406 | 0,261888 | Acp5 | -4,2398859 | 3,30E-09 |
| Bmp2 | 40,3808 | 24,7348 | 15,1659 | 13,6373 | 2,90888 | 0,705761 | 0,640486 | 0,165255 | 0,243248 | 0,176382 | Bmp2 | -3,61250811 | 3,32E-09 |
| Tpx2 | 47,6202 | 46,8038 | 36,9667 | 35,5039 | 11,7981 | 4,87442 | 4,29688 | 0 | 0,154647 | 0,118857 | Tpx2 | -2,30863725 | 3,41E-09 |
| Rab11fip4 | 1,76649 | 1,64013 | 2,34183 | 4,50151 | 0,117804 | 0 | 0 | 0 | 0 | 0 | Rab11fip4 | -5,12787985 | 3,41E-09 |
| Fam64a | 30,9489 | 37,5013 | 26,2368 | 31,9052 | 9,69163 | 4,66215 | 3,8214 | 0 | 0,0823631 | 0,555233 | Fam64a | -2,18846674 | 3,46E-09 |
| Pitx1 | 1,0911 | 1,74908 | 4,32589 | 5,90792 | 0 | 0 | 0 | 0 | 0 | 0 | Pitx1 | -6,00551922 | 3,48E-09 |
| Sema3e | 1,55673 | 0,496851 | 0,64973 | 0,876549 | 0,0280899 | 0,0114593 | 0 | 0 | 0 | 0 | Sema3e | -5,44720456 | 3,57E-09 |
| Espl1 | 20,5396 | 16,355 | 12,5916 | 15,5815 | 4,31167 | 2,24394 | 1,97581 | 0,0863766 | 0,0474992 | 0,0459734 | Espl1 | -2,17538566 | 3,57E-09 |
| Elp6 | 8,23781 | 7,15125 | 9,62602 | 8,46394 | 1,2153 | 1,61374 | 1,71966 | 0,89756 | 1,4405 | 0,801967 | Elp6 | -2,08104445 | 3,58E-09 |
| Epor | 7,92906 | 6,89065 | 9,45415 | 4,75388 | 0,964456 | 0,14234 | 0,289917 | 0,342633 | 0,317912 | 0,183957 | Epor | -3,54463063 | 3,59E-09 |
| Tubb3 | 2,02083 | 7,97394 | 53,2135 | 37,4514 | 0,917868 | 0,355655 | 0,206945 | 0,916624 | 0,272472 | 0,197012 | Tubb3 | -5,08118817 | 3,63E-09 |
| Dkk1 | 3,46472 | 2,22905 | 2,38894 | 1,97269 | 0 | 0,00756344 | 0 | 0 | 0 | 0 | Dkk1 | -6,47944137 | 3,75E-09 |
| Snrpg | 119,006 | 136,25 | 75,8757 | 128,51 | 30,489 | 17,7037 | 19,9684 | 17,9848 | 11,8449 | 17,3009 | Snrpg | -2,43049431 | 3,84E-09 |
| Pacsin1 | 0,684686 | 1,88471 | 8,03745 | 3,75137 | 0,0439515 | 0,0971605 | 0,0950863 | 0,211725 | 0,0778227 | 0,173008 | Pacsin1 | -4,68414808 | 3,96E-09 |
| Fanci | 0 | 6,76261 | 6,5949 | 11,5742 | 0 | 0 | 0 | 0 | 0 | 0 | Fanci | -2,2560885 | 4,07E-09 |
| Sass6 | 4,83365 | 6,38075 | 7,49436 | 10,9026 | 0 | 0 | 0 | 0 | 0 | 0 | Sass6 | -2,23747869 | 4,08E-09 |
| Sox3 | 5,76334 | 4,06101 | 6,32225 | 23,3877 | 0,467167 | 0,124067 | 0,353855 | 0 | 0 | 0 | Sox3 | -4,33904197 | 4,12E-09 |
| Clstn2 | 0,588731 | 2,97651 | 9,02993 | 7,48871 | 0,187625 | 0,0552997 | 0,0676514 | 0,0669612 | 0,0984164 | 0,095192 | Clstn2 | -4,50474124 | 4,25E-09 |
| Ism1 | 2,98944 | 4,4782 | 3,7891 | 3,5694 | 0,37661 | 0,0277632 | 0,0679059 | 0,201314 | 0,259734 | 0,143421 | Ism1 | -3,72819931 | 4,38E-09 |
| Car13 | 3,94611 | 6,63212 | 1,82983 | 3,38621 | 0,165806 | 0,0733716 | 0 | 0 | 0,0490895 | 0 | Car13 | -4,54992324 | 4,40E-09 |
| Pou3f2 | 0,401925 | 0,644314 | 1,70546 | 2,58172 | 0,0142269 | 0 | 0 | 0,0457392 | 0 | 0 | Pou3f2 | -5,98879337 | 4,55E-09 |
| Prr5l | 6,1178 | 4,02347 | 3,53051 | 2,5112 | 0,530557 | 0,149459 | 0,162014 | 0 | 0,0258086 | 0 | Prr5l | -3,41011144 | 4,68E-09 |
| Colec12 | 38,2433 | 30,7791 | 34,163 | 28,5296 | 6,42157 | 1,37542 | 1,482 | 0,712859 | 0,951517 | 0,666043 | Colec12 | -3,00409756 | 4,71E-09 |
| Lck | 2,29416 | 3,23306 | 3,82967 | 1,81573 | 0,0464896 | 0 | 0 | 0 | 0 | 0,0531746 | Lck | -4,98156968 | 4,79E-09 |
| Col18a1 | 243,599 | 238,586 | 127,907 | 111,973 | 19,3233 | 2,11472 | 2,37492 | 0,688322 | 0,736103 | 0,692668 | Col18a1 | -3,94549773 | 4,86E-09 |
| Id2 | 145,713 | 142,031 | 192,164 | 168,525 | 35,3532 | 9,01838 | 9,41271 | 4,44092 | 4,49889 | 15,7111 | Id2 | -2,78678374 | 5,04E-09 |
| Ticrr | 8,82146 | 8,08409 | 5,56686 | 7,8674 | 1,33293 | 0,379363 | 0,307018 | 0 | 0 | 0 | Ticrr | -3,10725151 | 5,11E-09 |
| Sp8 | 0,856369 | 1,05601 | 1,01267 | 4,60172 | 0,0202155 | 0 | 0,00437339 | 0,0129876 | 0 | 0 | Sp8 | -6,02985642 | 5,21E-09 |
| Kif23 | 32,8619 | 30,8276 | 28,7361 | 32,1139 | 9,20495 | 4,0647 | 3,48886 | 0,086252 | 0,0952525 | 0,0920821 | Kif23 | -2,12508609 | 5,74E-09 |
| Gm8801 | 12,4557 | 10,9166 | 7,00422 | 8,1122 | 2,32924 | 1,71486 | 1,822 | 0,502158 | 1,6353 | 1,86676 | Gm8801 | -5,34878677 | 6,37E-09 |
| Sox1 | 0,955703 | 0,864232 | 1,32956 | 3,61082 | 0 | 0 | 0 | 0 | 0 | 0 | Sox1 | -6,49105722 | 6,51E-09 |
| Basp1 | 62,9825 | 49,8511 | 67,1441 | 49,2169 | 8,77911 | 1,21392 | 1,65472 | 1,51747 | 1,37634 | 0,482706 | Basp1 | -3,45876536 | 6,94E-09 |
| Pdgfc | 3,23171 | 7,4692 | 12,4857 | 14,6981 | 1,17742 | 0,485955 | 0,537787 | 0,503874 | 1,85444 | 1,16534 | Pdgfc | -3,25635527 | 7,31E-09 |
| Tmem151b | 1,18169 | 1,55432 | 4,10994 | 3,70414 | 0,14876 | 0,115081 | 0,100568 | 0 | 0 | 0,0848934 | Tmem151b | -3,90578142 | 7,31E-09 |
| Lhfpl1 | 5,86379 | 4,24194 | 2,28392 | 1,18124 | 0 | 0 | 0 | 0 | 0 | 0 | Lhfpl1 | -6,45109718 | 7,51E-09 |
| Sec14l2 | 4,73992 | 3,11724 | 4,0662 | 6,63185 | 0,522671 | 0,495529 | 0,323141 | 0,119639 | 0,0441826 | 0,085344 | Sec14l2 | -2,82238019 | 8,02E-09 |
| Iglon5 | 1,87744 | 4,47051 | 4,10678 | 4,95656 | 0,14685 | 0,0324847 | 0 | 0 | 0 | 0 | Iglon5 | -4,52382881 | 8,03E-09 |
| Actb | 3272,21 | 2284,07 | 1606,3 | 1737,46 | 506,913 | 149,951 | 157,263 | 158,846 | 193,914 | 157,158 | Actb | -2,77326176 | 8,06E-09 |
| Kif22 | 47,4498 | 54,6096 | 36,4414 | 49,456 | 14,2096 | 6,24809 | 6,62575 | 0,149593 | 0,277019 | 0,0534645 | Kif22 | -2,08762956 | 8,19E-09 |
| Capn5 | 8,55953 | 10,0183 | 6,47876 | 4,89669 | 1,20209 | 0,327041 | 0,222273 | 0,132009 | 0,291005 | 0,234565 | Capn5 | -3,24382639 | 8,69E-09 |
| Id1 | 181,781 | 130,295 | 85,4155 | 158,194 | 21,6607 | 4,94424 | 4,1942 | 8,04947 | 10,7472 | 17,401 | Id1 | -3,34743051 | 8,79E-09 |
| Lgi2 | 6,40665 | 3,51111 | 12,9226 | 10,9347 | 0,706996 | 0,894241 | 0,546469 | 0,323069 | 0,419267 | 0,231117 | Lgi2 | -3,05906832 | 8,82E-09 |
| Apitd1 | 20,5012 | 18,9589 | 29,2257 | 18,4879 | 2,55557 | 3,56713 | 4,34977 | 1,9272 | 1,0225 | 0,980021 | Apitd1 | -2,28268922 | 8,87E-09 |
| Gucy1a2 | 0,630618 | 0,977605 | 1,24083 | 0,862394 | 0,121122 | 0,0225319 | 0,0344717 | 0 | 0,157617 | 0,0290763 | Gucy1a2 | -3,45941514 | 9,06E-09 |
| Tspan13 | 17,3029 | 29,9145 | 29,7768 | 37,0135 | 5,82113 | 1,61343 | 1,75282 | 4,20971 | 8,94558 | 5,50199 | Tspan13 | -2,8234194 | 9,07E-09 |
| Wdr54 | 8,40755 | 6,8637 | 15,4851 | 8,46636 | 0,494513 | 0,876942 | 0,981473 | 1,04987 | 1,17932 | 0,945316 | Wdr54 | -3,09224793 | 9,21E-09 |
| Has2 | 18,9623 | 18,4082 | 8,76541 | 5,40422 | 0,87748 | 0,0756968 | 0,115753 | 0 | 0 | 0 | Has2 | -4,38378537 | 9,23E-09 |
| Sult4a1 | 1,5453 | 0,970083 | 5,15928 | 4,10398 | 0 | 0 | 0 | 0 | 0,141504 | 0,18218 | Sult4a1 | -6,44461051 | 9,51E-09 |
| Prss8 | 2,77835 | 2,92349 | 1,85514 | 5,03731 | 0 | 0 | 0 | 0 | 0 | 0 | Prss8 | -5,71955411 | 9,94E-09 |
| Lmo2 | 27,7541 | 27,9409 | 17,3806 | 11,8146 | 4,01984 | 2,65662 | 2,5861 | 9,95818 | 11,29 | 10,0068 | Lmo2 | -2,40477241 | 1,01E-08 |
| Mtfr2 | 6,40607 | 5,74498 | 6,44064 | 7,18649 | 1,44523 | 0,950163 | 0,938239 | 0,132252 | 0,146699 | 0,141633 | Mtfr2 | -2,07377525 | 1,04E-08 |
| Shh | 0,514509 | 2,6953 | 7,37473 | 2,62734 | 0 | 0 | 0 | 0 | 0 | 0 | Shh | -6,47409696 | 1,09E-08 |
| Foxg1 | 0,334351 | 0,955813 | 2,51172 | 14,7037 | 0 | 0 | 0 | 0 | 0 | 0 | Foxg1 | -6,6427333 | 1,10E-08 |
| Tfap2b | 0,252548 | 0,507359 | 2,23238 | 3,98414 | 0,00289565 | 0 | 0 | 0 | 0 | 0 | Tfap2b | -6,55294966 | 1,17E-08 |
| Smad6 | 43,6391 | 73,6747 | 64,9199 | 50,5305 | 17,9752 | 6,89793 | 6,87739 | 4,51973 | 5,66458 | 5,93896 | Smad6 | -2,18344569 | 1,19E-08 |
| Krt20 | 1,24355 | 2,27823 | 5,63447 | 5,72617 | 0 | 0,0446333 | 0,0545508 | 0 | 0 | 0 | Krt20 | -5,40940722 | 1,23E-08 |
| Six2 | 3,54026 | 2,06369 | 6,80524 | 3,72825 | 0,273907 | 0,0404103 | 0,049397 | 0 | 0 | 0,0522122 | Six2 | -4,35650059 | 1,24E-08 |
| Pvrl1 | 42,8944 | 23,605 | 16,5133 | 20,5336 | 4,37583 | 1,17945 | 1,17367 | 0,457273 | 0,482605 | 0,284213 | Pvrl1 | -3,15762116 | 1,24E-08 |
| Spint1 | 5,32545 | 1,02151 | 1,60513 | 5,81162 | 0,0419567 | 0 | 0 | 0 | 0 | 0 | Spint1 | -5,94675632 | 1,28E-08 |
| Pklr | 0,37349 | 4,0877 | 7,79146 | 8,94522 | 0,105892 | 0 | 0 | 0 | 0 | 0 | Pklr | -5,7127992 | 1,29E-08 |
| Cntnap2 | 0,473164 | 1,58703 | 1,89577 | 4,29033 | 0 | 0 | 0 | 0 | 0 | 0,0145192 | Cntnap2 | -6,42146768 | 1,29E-08 |
| Gypa | 2,83533 | 7,64507 | 8,67708 | 2,47509 | 0,268238 | 0,047506 | 0,0580558 | 0 | 0 | 0 | Gypa | -4,46183657 | 1,30E-08 |
| Lars2 | 7950,46 | 4096,62 | 4092,73 | 7802,95 | 824,959 | 190,694 | 221,754 | 328,584 | 270,537 | 327,934 | Lars2 | -3,15750196 | 1,39E-08 |
| Mpl | 0,734948 | 1,40643 | 3,88518 | 4,17421 | 0,0347274 | 0 | 0 | 0 | 0 | 0 | Mpl | -5,85124844 | 1,50E-08 |
| Dab1 | 0,657342 | 3,03216 | 4,12923 | 4,97617 | 0 | 0 | 0 | 0 | 0 | 0 | Dab1 | -5,75122058 | 1,53E-08 |
| Tpm3 | 179,139 | 160,703 | 84,4323 | 70,4147 | 26,9979 | 10,2541 | 9,32833 | 3,39873 | 4,67557 | 4,96477 | Tpm3 | -2,69007303 | 1,56E-08 |
| Sema3f | 11,687 | 15,8763 | 15,5923 | 8,29999 | 2,36179 | 0,552239 | 0,469879 | 0,958312 | 1,47535 | 1,33319 | Sema3f | -3,08270711 | 1,69E-08 |
| Plch1 | 0,403525 | 0,796158 | 1,26313 | 3,10023 | 0 | 0,0125906 | 0,030812 | 0,0457928 | 0 | 0,0162515 | Plch1 | -5,09320678 | 1,78E-08 |
| Lxn | 19,8334 | 32,6074 | 20,3265 | 21,3707 | 5,24767 | 1,93287 | 2,03703 | 0,629173 | 0,473572 | 0,909678 | Lxn | -2,44897055 | 1,79E-08 |
| Necab2 | 0,520488 | 5,64795 | 10,7158 | 11,8837 | 0,0984574 | 0 | 0 | 0 | 0 | 0 | Necab2 | -5,6472124 | 1,79E-08 |

| Gene | d9-11 (1) | d9-11 (2) | d9-11(3) | d9-11(4) | newborn(1) | newborn(2) | newborn(3) | adult(1) | adult(2) | adult(3) | Gene | log2(fold) | FDR(*p*-value) |
|---|---|---|---|---|---|---|---|---|---|---|---|---|---|
| Mcm8 | 9,02268 | 12,9958 | 9,77238 | 8,97496 | 1,03299 | 1,98162 | 2,38548 | 1,95521 | 1,47061 | 1,05872 | Mcm8 | -2,07860171 | 1,81E-08 |
| Prdm1 | 10,6223 | 7,55455 | 1,80427 | 1,36893 | 0,227389 | 0,0154637 | 0,0378394 | 0,112418 | 0,103161 | 0,0998106 | Prdm1 | -4,93690442 | 1,83E-08 |
| Slc4a1 | 1,8957 | 13,5063 | 28,1311 | 16,4825 | 0,815429 | 0,105509 | 0,0860526 | 0 | 0 | 0 | Slc4a1 | -4,61402064 | 1,89E-08 |
| H2-T10 | 3,05068 | 6,07572 | 4,10638 | 2,73895 | 0,664715 | 0,123661 | 0,160051 | 0,16569 | 0,338675 | 0,320664 | H2-T10 | -4,70158297 | 1,90E-08 |
| Ap1m2 | 2,41058 | 6,73757 | 1,84428 | 3,23729 | 0 | 0,0504044 | 0,0615914 | 0,181884 | 0 | 0 | Ap1m2 | -5,260977 | 2,04E-08 |
| Clstn3 | 2,88981 | 0,791886 | 1,5411 | 2,26452 | 0,0909693 | 0,0603319 | 0,024601 | 0,0730289 | 0,107397 | 0 | Clstn3 | -4,32554918 | 2,07E-08 |
| Epb4.2 | 0,280409 | 2,39739 | 4,29108 | 2,07172 | 0,026487 | 0 | 0 | 0 | 0 | 0 | Epb4.2 | -5,87888394 | 2,10E-08 |
| Snhg11 | 1,9214 | 0,584585 | 1,88865 | 1,03131 | 0,0457677 | 0,0809156 | 0,0495033 | 0 | 0,0179884 | 0,0174078 | Snhg11 | -3,87945604 | 2,19E-08 |
| Ephb1 | 12,374 | 6,03425 | 1,36957 | 2,12515 | 0,0796645 | 0,21131 | 0,021543 | 0,191936 | 0,754018 | 0,442067 | Ephb1 | -4,74810982 | 2,23E-08 |
| Alx1 | 1,06952 | 3,29708 | 1,78541 | 2,15483 | 0 | 0 | 0 | 0 | 0 | 0 | Alx1 | -6,25634978 | 2,24E-08 |
| Cnr1 | 0,816307 | 0,966349 | 1,31737 | 0,712828 | 0,0616464 | 0,0136237 | 0,0500087 | 0 | 0 | 0,0175857 | Cnr1 | -3,94571577 | 2,38E-08 |
| Apba2 | 0,797769 | 1,25783 | 4,04857 | 4,81926 | 0,0565228 | 0,04917 | 0 | 0 | 0 | 0 | Apba2 | -5,02558323 | 2,39E-08 |
| Dlx2 | 0,404626 | 3,19327 | 3,37732 | 5,97831 | 0,0382462 | 0 | 0 | 0 | 0 | 0 | Dlx2 | -5,86067435 | 2,47E-08 |
| Orc6 | 40,664 | 33,3518 | 21,2344 | 29,6507 | 7,92347 | 5,17265 | 5,32947 | 2,11298 | 2,59425 | 2,75406 | Orc6 | -2,02031683 | 2,70E-08 |
| Arc | 18,1117 | 2,95866 | 3,48737 | 4,64116 | 0,303693 | 0,0805682 | 0,197068 | 2,23989 | 2,90847 | 1,28394 | Arc | -4,39527043 | 2,78E-08 |
| Nr2f1 | 2,33745 | 16,0054 | 37,8726 | 28,9488 | 1,3257 | 0,517569 | 0,632805 | 0,124923 | 0,323099 | 0,267434 | Nr2f1 | -4,02584173 | 2,85E-08 |
| Ccdc88c | 3,30496 | 7,11352 | 7,14772 | 7,41692 | 1,36128 | 0,363495 | 0,475454 | 0,136766 | 0,468073 | 0,533902 | Ccdc88c | -2,69226252 | 2,92E-08 |
| Rbfox3 | 5,34358 | 1,38746 | 1,00942 | 2,08742 | 0,0332258 | 0,0546877 | 0 | 0 | 0 | 0 | Rbfox3 | -5,17812569 | 2,92E-08 |
| Apob | 1,22189 | 0,635577 | 0,352355 | 7,3427 | 0 | 0 | 0 | 0 | 0 | 0 | Apob | -6,67585508 | 2,92E-08 |
| Brsk1 | 30,2952 | 15,8768 | 22,5723 | 26,3615 | 6,73423 | 3,10213 | 3,08874 | 0,236513 | 0,089421 | 0,141814 | Brsk1 | -2,28427008 | 3,02E-08 |
| Vwa2 | 0,664087 | 2,53859 | 0,970062 | 1,42178 | 0 | 0 | 0 | 0 | 0 | 0 | Vwa2 | -5,62641678 | 3,02E-08 |
| Fgb | 2,24451 | 3,09978 | 3,18461 | 3,61692 | 0 | 0 | 0 | 0 | 0 | 0 | Fgb | -6,18785356 | 3,02E-08 |
| Zfp703 | 42,3316 | 30,1768 | 20,2892 | 31,1294 | 8,08597 | 3,76983 | 3,86851 | 10,6764 | 12,0576 | 13,3044 | Zfp703 | -2,26094424 | 3,03E-08 |
| T | 0,831892 | 8,69882 | 2,49963 | 3,35186 | 0,0472036 | 0 | 0,0510779 | 0 | 0 | 0 | T | -4,94883948 | 3,17E-08 |
| Tmem151a | 1,0609 | 1,28809 | 2,92912 | 1,72664 | 0,115606 | 0,0613378 | 0,0750327 | 0,371207 | 0,57326 | 0,343179 | Tmem151a | -3,73993194 | 3,22E-08 |
| Slc27a3 | 17,8295 | 18,3214 | 10,5412 | 12,273 | 2,94991 | 0,745955 | 1,25393 | 0,539826 | 0,3993 | 0,192741 | Slc27a3 | -2,70552841 | 3,23E-08 |
| Mme | 10,7171 | 11,5428 | 10,413 | 5,65252 | 1,2012 | 0,198726 | 0,0846364 | 2,51426 | 2,93564 | 2,57937 | Mme | -3,74901727 | 3,33E-08 |
| Oip5 | 11,305 | 13,2192 | 14,5543 | 11,729 | 2,80344 | 1,79012 | 1,32651 | 0 | 0,0769739 | 0,148032 | Oip5 | -2,3235466 | 3,33E-08 |
| Atp2c2 | 0,607286 | 2,09679 | 1,5207 | 4,3847 | 0 | 0,0253727 | 0,0310321 | 0 | 0,0338988 | 0 | Atp2c2 | -5,07370357 | 3,42E-08 |
| Arrb2 | 44,7425 | 28,5916 | 29,5414 | 20,6326 | 7,04632 | 4,56371 | 3,69615 | 0,661968 | 0,981917 | 1,0981 | Arrb2 | -2,17234616 | 3,52E-08 |
| Tal1 | 9,14869 | 7,28524 | 4,06719 | 2,76143 | 0,625478 | 0,0762317 | 0,116158 | 0,276878 | 0,101759 | 0,221602 | Tal1 | -3,90314214 | 3,61E-08 |
| Slc13a4 | 3,50797 | 2,15118 | 1,89932 | 2,86557 | 0,214939 | 0,0237627 | 0 | 0 | 0 | 0,0306844 | Slc13a4 | -4,36773243 | 3,62E-08 |
| Caskin1 | 0 | 0 | 0 | 0 | 0 | 0 | 0 | 0 | 0 | 0 | Caskin1 | -2,24887011 | 3,70E-08 |
| Lin28a | 186,388 | 76,9883 | 3,96924 | 12,8695 | 0 | 0 | 0 | 0 | 0 | 0 | Lin28a | -6,74392225 | 3,82E-08 |
| Rprm | 2,69356 | 3,14021 | 3,47426 | 5,42541 | 0,139123 | 0,0616425 | 0 | 0,443837 | 0,248302 | 0,318823 | Rprm | -4,78388042 | 3,82E-08 |
| Suv39h1 | 31,6436 | 30,8155 | 28,1173 | 28,7916 | 9,49993 | 3,95899 | 3,70935 | 2,11318 | 2,46111 | 2,38444 | Suv39h1 | -2,02425166 | 3,99E-08 |
| Cbln1 | 2,72967 | 2,12588 | 2,87795 | 3,47338 | 0 | 0,180297 | 0 | 0,130509 | 0,33773 | 0,0931667 | Cbln1 | -4,59722208 | 4,03E-08 |
| Fam19a2 | 3,73205 | 1,67794 | 0,918533 | 1,32422 | 0,0876123 | 0,0193677 | 0,0236929 | 0 | 0,0517095 | 0,0250053 | Fam19a2 | -4,65214677 | 4,03E-08 |
| Ccp110 | 8,25472 | 13,6557 | 7,42046 | 6,52572 | 2,2661 | 1,43103 | 1,16722 | 0,173372 | 0,487966 | 0,328417 | Ccp110 | -2,06916743 | 4,04E-08 |
| Fzd7 | 40,6776 | 21,4367 | 16,0372 | 24,2304 | 4,18467 | 1,01204 | 0,710427 | 2,65283 | 2,30249 | 1,97046 | Fzd7 | -3,32330607 | 4,05E-08 |
| Gfi1b | 2,84417 | 3,18958 | 0,658447 | 2,81024 | 0,0573668 | 0 | 0 | 0 | 0 | 0 | Gfi1b | -5,4964712 | 4,26E-08 |
| Wnt6 | 0,4932 | 2,27048 | 2,46991 | 14,0761 | 0 | 0 | 0 | 0 | 0 | 0,160044 | Wnt6 | -6,38479124 | 4,27E-08 |
| Sncb | 4,35343 | 3,37419 | 2,0761 | 6,26236 | 0 | 0 | 0 | 0 | 0 | 0 | Sncb | -6,16983869 | 4,33E-08 |
| Bcar3 | 3,84874 | 5,25706 | 5,93087 | 3,57922 | 0,727155 | 0,197896 | 0,121021 | 1,34613 | 2,54459 | 2,29994 | Bcar3 | -3,26146987 | 4,43E-08 |
| Foxa1 | 1,33174 | 1,51587 | 0,855078 | 2,27053 | 0 | 0 | 0 | 0 | 0 | 0 | Foxa1 | -6,1660961 | 4,43E-08 |
| Spred3 | 5,07185 | 4,22159 | 3,37361 | 2,55499 | 0,583742 | 0,138973 | 0,242864 | 0 | 0,0530094 | 0,0512653 | Spred3 | -3,11038734 | 4,91E-08 |
| H19 | 2133,97 | 2756,77 | 2196,51 | 1771,88 | 412,65 | 60,8148 | 69,2179 | 0 | 1,27698 | 0,411158 | H19 | -3,18675585 | 4,98E-08 |
| Upk1b | 1,56872 | 2,90156 | 13,0931 | 7,90028 | 0,22264 | 0,0492925 | 0,180705 | 0,711652 | 1,32145 | 0,382252 | Upk1b | -4,56905824 | 5,00E-08 |
| Kctd15 | 15,3944 | 16,9649 | 23,9622 | 26,6847 | 2,55644 | 0,382852 | 0,170214 | 0,126001 | 0,558725 | 0,449605 | Kctd15 | -3,81912507 | 5,05E-08 |
| Kcnq3 | 0,800387 | 1,69195 | 4,39021 | 0,806335 | 0 | 0 | 0 | 0,10394 | 0 | 0 | Kcnq3 | -6,20951063 | 5,15E-08 |
| Sspo | 0,296878 | 0,214774 | 0,512162 | 0,438758 | 0,028009 | 0,0296984 | 0,0121165 | 0,0180314 | 0,00659459 | 0,00638721 | Sspo | -3,49365973 | 5,31E-08 |
| 2610018G03R | 3,23733 | 9,91684 | 15,928 | 7,55165 | 1,11191 | 0,684784 | 0,4185 | 0,309369 | 0,229271 | 0,110607 | 2610018G03R | -3,07220186 | 5,52E-08 |
| Ccdc40 | 0,8305 | 3,46623 | 1,59978 | 0,96551 | 0,126662 | 0,0639894 | 0,0391445 | 0 | 0 | 0 | Ccdc40 | -3,68767456 | 5,58E-08 |
| Ppap2c | 22,3627 | 16,1774 | 10,4876 | 18,2332 | 2,96306 | 3,05316 | 2,87959 | 1,35212 | 0,86209 | 0,692404 | Ppap2c | -2,2131119 | 5,63E-08 |
| Hyou1 | 127,765 | 94,5925 | 46,8123 | 66,0911 | 14,8438 | 15,3966 | 14,254 | 19,7686 | 21,5472 | 20,0329 | Hyou1 | -2,18558054 | 5,66E-08 |
| Rimbp2 | 0,451043 | 4,82033 | 4,70616 | 1,13609 | 0,0851841 | 0,0376609 | 0,0230361 | 0 | 0,0251357 | 0 | Rimbp2 | -4,68863217 | 6,22E-08 |
| Sall4 | 32,4515 | 20,0177 | 6,87717 | 13,003 | 0,69058 | 1,12947 | 1,45854 | 1,33811 | 1,47643 | 1,26148 | Sall4 | -3,10113149 | 6,37E-08 |
| Gpc3 | 363,734 | 420,092 | 511,884 | 375,074 | 135,399 | 44,915 | 46,1224 | 0,404279 | 3,09 | 1,05852 | Gpc3 | -2,18396577 | 6,45E-08 |
| Katnal2 | 3,13539 | 1,85578 | 2,0935 | 2,94742 | 0,0593447 | 0,0525643 | 0,128451 | 0 | 0 | 0,0679454 | Katnal2 | -4,4238132 | 6,63E-08 |
| B3gnt7 | 2,43111 | 1,76714 | 1,3884 | 2,9647 | 0,108838 | 0 | 0 | 0,116278 | 0,171709 | 0 | B3gnt7 | -4,88062239 | 6,65E-08 |
| Dlgap5 | 23,2707 | 24,0381 | 23,0203 | 28,1192 | 7,86982 | 3,33281 | 2,95498 | 0 | 0 | 0,0358685 | Dlgap5 | -2,07065579 | 6,87E-08 |
| Grin3a | 2,61888 | 4,25421 | 3,68902 | 3,05113 | 0,22013 | 0,0102241 | 0 | 0 | 0,0136315 | 0 | Grin3a | -4,51726289 | 7,04E-08 |
| Stard4 | 5,16623 | 14,964 | 15,7809 | 12,0704 | 2,58602 | 1,90177 | 1,71906 | 0,199863 | 0,69986 | 0,177976 | Stard4 | -2,05617564 | 7,59E-08 |
| Gla | 7,89013 | 10,1293 | 6,58585 | 7,07777 | 1,84154 | 1,30081 | 0,629714 | 0,589601 | 0,941599 | 0,839956 | Gla | -2,31110015 | 7,59E-08 |
| Dbf4 | 33,8849 | 36,3064 | 21,5833 | 28,9434 | 8,60151 | 3,02132 | 3,90392 | 0,248446 | 0,27556 | 0,132962 | Dbf4 | -2,2292965 | 7,59E-08 |
| Ccdc120 | 2,84336 | 2,80497 | 2,00254 | 6,44551 | 0,453194 | 0,178131 | 0,217888 | 0,0808208 | 0,0594721 | 0,0575014 | Ccdc120 | -3,31771804 | 8,01E-08 |
| Neurl1a | 1,58138 | 4,68006 | 3,29997 | 2,56687 | 0,398295 | 0,220149 | 0,107715 | 1,03886 | 0,470388 | 0,596941 | Neurl1a | -3,0297172 | 8,12E-08 |
| Podxl | 178,733 | 97,7276 | 45,0957 | 43,7624 | 11,3407 | 2,30905 | 2,07786 | 3,52012 | 9,69878 | 6,09599 | Podxl | -3,64859263 | 8,25E-08 |
| Cdca4 | 21,4186 | 25,6276 | 18,2235 | 23,2469 | 6,83806 | 3,19035 | 2,45911 | 1,38099 | 1,85559 | 1,56787 | Cdca4 | -2,11112364 | 9,57E-08 |
| Sptbn2 | 1,0623 | 2,69477 | 4,30705 | 4,77854 | 0,236309 | 0,0759455 | 0,0232347 | 0,0690997 | 0,0126557 | 0,0612608 | Sptbn2 | -2,95920356 | 9,88E-08 |
| Hsd3b6 | 0,576015 | 2,84112 | 5,98283 | 3,86789 | 0 | 0 | 0,0147439 | 0 | 0,0323397 | 0 | Hsd3b6 | -6,12245325 | 1,00E-07 |
| Kbtbd8 | 2,35831 | 1,32487 | 0,944023 | 1,42566 | 0,0190858 | 0,118121 | 0,064975 | 0,192961 | 0,450372 | 0,217125 | Kbtbd8 | -3,89420839 | 1,05E-07 |
| Tbx2 | 13,0064 | 15,9712 | 21,9409 | 20,1598 | 3,99483 | 0,957339 | 0,864298 | 0,082726 | 0,547966 | 0,706374 | Tbx2 | -2,83341582 | 1,06E-07 |
| Micall2 | 4,95896 | 3,16541 | 4,70739 | 4,31029 | 0,744 | 0,170605 | 0,149048 | 0,0884253 | 0,195346 | 0,251777 | Micall2 | -3,14363301 | 1,06E-07 |
| Alg8 | 26,9512 | 32,981 | 19,7898 | 15,292 | 3,72666 | 4,15618 | 5,44662 | 2,63258 | 2,75437 | 1,7717 | Alg8 | -2,01542385 | 1,06E-07 |
| Dbn1 | 89,1759 | 72,6525 | 73,1007 | 67,5743 | 17,1276 | 4,14269 | 3,67154 | 0,308422 | 0,49293 | 0,512761 | Dbn1 | -2,75722823 | 1,07E-07 |
| Syt14 | 3,61825 | 2,37954 | 3,01991 | 0,404929 | 0,0570625 | 0 | 0 | 0 | 0 | 0 | Syt14 | -5,48651102 | 1,07E-07 |
| Gm5082 | 3,75471 | 2,61451 | 1,29037 | 0,75688 | 0,062718 | 0 | 0 | 0 | 0 | 0 | Gm5082 | -5,44165899 | 1,09E-07 |
| Cox4i2 | 31,6617 | 17,6777 | 28,4111 | 54,1007 | 3,41369 | 0,303895 | 0,924973 | 0,53699 | 1,65266 | 0,788428 | Cox4i2 | -3,7197997 | 1,12E-07 |
| Zc4h2 | 5,82236 | 8,52529 | 7,73452 | 4,73459 | 1,26037 | 1,33514 | 0,611336 | 0,139156 | 0,30892 | 0,347893 | Zc4h2 | -2,28776509 | 1,21E-07 |
| Ptpn7 | 7,38905 | 2,16708 | 1,33354 | 1,15687 | 0,088433 | 0 | 0,0765339 | 0 | 0,0167198 | 0,0161613 | Ptpn7 | -4,92907683 | 1,21E-07 |
| Syt13 | 1,89395 | 0,934189 | 3,1618 | 1,82142 | 0,170747 | 0,0431428 | 0,0791598 | 0 | 0,0288052 | 0,0278526 | Syt13 | -3,75889271 | 1,27E-07 |
| Lhx2 | 0,452592 | 2,74463 | 5,29436 | 15,1935 | 0,0855823 | 0,0367567 | 0 | 0,137024 | 0,0506873 | 0 | Lhx2 | -5,14994682 | 1,29E-07 |
| Treml1 | 2,31554 | 1,90343 | 12,2287 | 17,7662 | 0,109885 | 0,0975532 | 0 | 0 | 0 | 0 | Treml1 | -5,49947228 | 1,30E-07 |
| Slc12a5 | 0,943121 | 1,91247 | 2,01187 | 1,63658 | 0,19289 | 0,288552 | 0,256779 | 0,476986 | 0,227423 | 0,355531 | Slc12a5 | -2,40648758 | 1,30E-07 |
| Slc39a4 | 3,4837 | 4,03427 | 0,379268 | 3,96699 | 0 | 0 | 0 | 0 | 0 | 0 | Slc39a4 | -5,63622344 | 1,33E-07 |
| Mirg | 10,128 | 9,71331 | 9,39179 | 17,8484 | 0,959617 | 0,212693 | 0,173138 | 0 | 0 | 0,0917034 | Mirg | -3,62228969 | 1,40E-07 |
| Gpr98 | 0,159164 | 0,722278 | 0,837038 | 0,930234 | 0,0495507 | 0,00398006 | 0,00974319 | 0 | 0,00530199 | 0 | Gpr98 | -3,93937456 | 1,40E-07 |
| E130012A19R | 27,0733 | 21,7353 | 12,741 | 18,1333 | 3,55323 | 0,578322 | 0,839612 | 1,83144 | 2,03122 | 1,91446 | E130012A19R | -3,19090928 | 1,41E-07 |
| Tpbg | 10,6394 | 11,0317 | 12,2593 | 20,2784 | 1,96356 | 0,25478 | 0,226643 | 0,0840553 | 0,0618714 | 0,0897236 | Tpbg | -3,60862357 | 1,45E-07 |
| Pcbp3 | 9,32472 | 6,43906 | 19,198 | 12,0544 | 2,16014 | 1,40474 | 1,62183 | 0,42342 | 0,940149 | 0,453728 | Pcbp3 | -2,33408288 | 1,47E-07 |
| Nlgn3 | 1,99172 | 2,2923 | 3,74065 | 3,17723 | 0,564307 | 0,270304 | 0,305216 | 0 | 0,0277623 | 0,0536927 | Nlgn3 | -2,42658002 | 1,50E-07 |
| Gnaz | 4,52717 | 1,35333 | 2,74816 | 5,11335 | 0,272322 | 0,206559 | 0,0841832 | 0,124642 | 0 | 0 | Gnaz | -3,71247247 | 1,50E-07 |
| Tenm3 | 14,4527 | 16,6667 | 21,4504 | 19,6846 | 4,30018 | 0,887861 | 0,921452 | 0,103386 | 0,236651 | 0,137496 | Tenm3 | -2,78095571 | 1,50E-07 |
| Camsap3 | 7,3146 | 3,38454 | 3,79905 | 6,91338 | 0,782339 | 0,841017 | 0,480133 | 0,135775 | 0,324308 | 0,241288 | Camsap3 | -2,57951316 | 1,52E-07 |
| Arhgap19 | 19,1431 | 16,2156 | 10,7701 | 9,94611 | 3,73587 | 1,71343 | 1,40557 | 0,457273 | 0,461622 | 0,609027 | Arhgap19 | -2,29814036 | 1,55E-07 |
| H1fx | 90,8009 | 82,816 | 87,2016 | 98,6757 | 28,4069 | 9,37436 | 10,3277 | 1,62837 | 2,20857 | 2,23833 | H1fx | -2,18328878 | 1,56E-07 |
| Nhs | 3,10058 | 1,64701 | 0,983297 | 1,44187 | 0,168874 | 0,104505 | 0,0902153 | 0,0542723 | 0,139423 | 0,0192718 | Nhs | -3,25828396 | 1,56E-07 |
| Stk32a | 3,81583 | 3,97578 | 4,0798 | 1,20949 | 0,340145 | 0,322291 | 0,210258 | 0,0780012 | 0,114764 | 0,138712 | Stk32a | -2,81876747 | 1,58E-07 |
| Bmp5 | 32,9819 | 13,5081 | 17,0341 | 17,1102 | 2,05335 | 2,74193 | 3,43337 | 0 | 0,0441826 | 0,085344 | Bmp5 | -2,46307697 | 1,60E-07 |
| Ache | 2,29137 | 5,12231 | 9,42094 | 10,4638 | 0,865943 | 0,498192 | 0,281302 | 0,416215 | 0,461969 | 0,544619 | Ache | -3,1291853 | 1,65E-07 |
| Gramd1c | 7,67776 | 7,39157 | 4,81599 | 6,40873 | 0,58309 | 0,996871 | 1,34109 | 0,252396 | 0,299768 | 0,257562 | Gramd1c | -2,38311653 | 1,66E-07 |
| Mapk11 | 5,1967 | 5,70465 | 3,99512 | 2,75528 | 0,581707 | 0,343177 | 0,419588 | 0 | 0,229529 | 0,132992 | Mapk11 | -2,75732763 | 1,66E-07 |
| D430019H16F | 0,600883 | 1,61665 | 4,55978 | 4,34814 | 0,185637 | 0,184615 | 0,117127 | 0,149179 | 0,14593 | 0,229478 | D430019H16F | -3,62215293 | 1,75E-07 |
| Nmral1 | 24,6318 | 36,1322 | 18,9453 | 23,1245 | 3,17768 | 5,82217 | 5,7456 | 1,59427 | 2,8897 | 2,86418 | Nmral1 | -2,00522784 | 1,76E-07 |
| **Hoxc4** | 0,172317 | 3,05975 | 4,02707 | 4,86002 | 0,0488931 | 0 | 0 | 0 | 0 | 0 | Hoxc4 | -5,5864142 | 1,76E-07 |
| Elavl4 | 1,02957 | 2,43354 | 3,41633 | 1,72907 | 0,286206 | 0,107491 | 0,147567 | 0,0780451 | 0 | 0,0277581 | Elavl4 | -3,22856944 | 1,89E-07 |
| Stx1b | 2,69142 | 6,41625 | 3,93103 | 6,32494 | 0,382157 | 0,16928 | 0,275742 | 0 | 0 | 0 | Stx1b | -3,25965911 | 1,91E-07 |
| Sgsm1 | 2,21234 | 1,50228 | 1,79371 | 1,55819 | 0,158687 | 0 | 0,0443798 | 0,0670097 | 0,0246221 | 0 | Sgsm1 | -4,17446557 | 1,93E-07 |
| Tfap2c | 1,04518 | 1,14562 | 1,64787 | 6,08365 | 0,0329208 | 0,0595963 | 0 | 0 | 0 | 0,0376154 | Tfap2c | -5,06712942 | 1,94E-07 |
| Lamc3 | 0,585735 | 0,21012 | 2,80025 | 1,98514 | 0 | 0 | 0,0163102 | 0 | 0,0533397 | 0 | Lamc3 | -4,94653322 | 1,96E-07 |
| Bnc1 | 0,547409 | 0,540019 | 2,11332 | 1,58567 | 0,0387647 | 0 | 0 | 0,0622688 | 0,068609 | 0 | Bnc1 | -5,14557693 | 2,01E-07 |
| Trpv2 | 3,00184 | 6,39637 | 7,61701 | 4,7176 | 0,987179 | 0,511987 | 0,626079 | 0,327411 | 0,281844 | 0,23339 | Trpv2 | -2,41617219 | 2,05E-07 |

| Gene | d9-11 (1) | d9-11 (2) | d9-11(3) | d9-11(4) | newborn(1) | newborn(2) | newborn(3) | adult(1) | adult(2) | adult(3) | Gene | log2(fold) | FDR(*p*-value) |
|---|---|---|---|---|---|---|---|---|---|---|---|---|---|
| Map7d2 | 2,09125 | 1,89797 | 1,04736 | 2,52834 | 0,213305 | 0,0838355 | 0,111098 | 0 | 0,0279849 | 0,00169132 | Map7d2 | -3,27643569 | 2,10E-07 |
| 6330403K07R | 3,52616 | 8,69611 | 22,4403 | 17,7572 | 1,25162 | 0,443513 | 0,338657 | 0 | 0 | 0 | 6330403K07R | -3,62071002 | 2,10E-07 |
| Lrrn1 | 0,171934 | 1,78093 | 4,37721 | 6,14944 | 0,0730739 | 0,0215409 | 0 | 0 | 0,0287643 | 0,0834395 | Lrrn1 | -5,2178192 | 2,13E-07 |
| Foxd1 | 1,22037 | 2,67529 | 7,13031 | 3,96132 | 0,192256 | 0,0340259 | 0 | 0 | 0 | 0,0219766 | Foxd1 | -4,52642687 | 2,15E-07 |
| Fermt1 | 0,241537 | 0,893538 | 0,856874 | 1,27766 | 0,00285036 | 0 | 0 | 0 | 0 | 0 | Fermt1 | -5,9612017 | 2,19E-07 |
| Draxin | 0,183152 | 0,421587 | 2,08939 | 1,84538 | 0 | 0 | 0 | 0 | 0 | 0 | Draxin | -6,05154463 | 2,22E-07 |
| Enox1 | 2,31822 | 4,02497 | 7,81752 | 5,3251 | 0,656918 | 0,488034 | 0,419088 | 0,0843366 | 0,0620801 | 0,150043 | Enox1 | -2,79009815 | 2,25E-07 |
| Cacna1b | 0,161187 | 0,572444 | 0,861121 | 0,487252 | 0,0182523 | 0,00806513 | 0 | 0 | 0 | 0,0104085 | Cacna1b | -4,75894034 | 2,25E-07 |
| Tpm4 | 203,222 | 191,059 | 88,9301 | 92,8348 | 37,726 | 10,7973 | 9,71712 | 6,44471 | 9,73824 | 6,48576 | Tpm4 | -2,94365691 | 2,35E-07 |
| Tcfl5 | 3,66988 | 0,905068 | 2,5526 | 1,58439 | 0 | 0 | 0 | 0 | 0,0513821 | 0,0495985 | Tcfl5 | -5,95435802 | 2,45E-07 |
| Irf5 | 5,39805 | 15,0623 | 26,1621 | 16,8067 | 2,18741 | 0,425948 | 0,331358 | 0 | 0,0518293 | 0 | Irf5 | -3,45560407 | 2,46E-07 |
| Reln | 3,28535 | 4,10357 | 1,70297 | 1,60185 | 0,532484 | 0,192193 | 0,227111 | 0,193075 | 0,282615 | 0,40197 | Reln | -2,69975751 | 2,46E-07 |
| Pax3 | 0,0963109 | 2,46612 | 2,15623 | 3,68333 | 0,0272934 | 0 | 0 | 0 | 0 | 0 | Pax3 | -5,63833262 | 2,50E-07 |
| Fam83g | 1,59801 | 0,525479 | 1,72295 | 3,22014 | 0,0565797 | 0 | 0,0204002 | 0 | 0,0445026 | 0 | Fam83g | -5,08584549 | 2,60E-07 |
| BC048507 | 7,99729 | 15,3825 | 4,00327 | 1,60885 | 0,687556 | 1,22723 | 1,74009 | 0 | 0,280081 | 0,26576 | BC048507 | -3,27761073 | 2,68E-07 |
| Tvp23a | 2,36714 | 2,74835 | 5,16192 | 5,93267 | 0,455785 | 0 | 0 | 0 | 0 | 0 | Tvp23a | -3,41697625 | 2,71E-07 |
| Pcdh8 | 0,783446 | 0,686993 | 2,98218 | 5,47541 | 0,0986605 | 0,0218128 | 0,0266816 | 0 | 0,0291285 | 0,0281646 | Pcdh8 | -4,72504108 | 2,74E-07 |
| Hkdc1 | 0,652083 | 0,551381 | 1,16635 | 3,47252 | 0 | 0 | 0 | 0,0847232 | 0 | 0 | Hkdc1 | -5,98840489 | 2,74E-07 |
| Mir6236 | 1355240 | 664016 | 669384 | 1359750 | 175766 | 21496,4 | 27817,7 | 50842,4 | 38028,7 | 49403,4 | Mir6236 | -3,54651786 | 2,75E-07 |
| Ccdc92 | 2,0893 | 2,20827 | 2,6158 | 2,4053 | 0,126984 | 0,0374631 | 0,183197 | 0,406651 | 0,20054 | 0,241996 | Ccdc92 | -3,53038302 | 2,77E-07 |
| Zswim6 | 13,1182 | 12,1394 | 6,29874 | 6,31631 | 2,01378 | 0,728445 | 0,640118 | 1,32086 | 1,86118 | 1,70711 | Zswim6 | -2,65818514 | 2,88E-07 |
| Dusp26 | 4,77386 | 6,47526 | 15,7717 | 11,0191 | 1,58098 | 0,650075 | 0,733263 | 0,180461 | 0 | 0,258529 | Dusp26 | -2,79780733 | 2,92E-07 |
| Cenpu | 7,36876 | 10,2894 | 7,71919 | 6,16748 | 1,16171 | 1,1663 | 1,24118 | 0 | 0 | 0 | Cenpu | -2,19956126 | 2,98E-07 |
| Cenpw | 9,04867 | 16,925 | 8,82711 | 9,46826 | 3,30442 | 2,60254 | 2,0231 | 0,213051 | 0,238174 | 0,458846 | Cenpw | -2,13462451 | 3,14E-07 |
| Camkv | 1,18935 | 2,34656 | 0,451246 | 1,74284 | 0,0306465 | 0 | 0 | 0 | 0 | 0 | Camkv | -5,35059213 | 3,15E-07 |
| Sorl1 | 5,83874 | 6,56366 | 16,0949 | 10,991 | 2,26842 | 0,974191 | 0,928723 | 3,0084 | 2,76428 | 2,99729 | Sorl1 | -2,41947616 | 3,16E-07 |
| Rgs8 | 0,326698 | 0,752008 | 1,86349 | 1,04239 | 0,0308398 | 0 | 0,033357 | 0 | 0 | 0 | Rgs8 | -4,57558375 | 3,22E-07 |
| Nog | 12,3154 | 5,10605 | 7,44857 | 12,2252 | 0,810373 | 0 | 0,164423 | 0,162006 | 0,240292 | 0,405649 | Nog | -4,17376602 | 3,26E-07 |
| Tdgf1 | 38,4844 | 21,5934 | 4,90168 | 3,76439 | 0,151499 | 0,044712 | 0 | 0 | 0 | 0,0577792 | Tdgf1 | -6,02038598 | 3,27E-07 |
| Eps8l2 | 0,640987 | 1,68621 | 1,42674 | 8,17965 | 0,0605598 | 0,0803552 | 0 | 0,0971306 | 0,501098 | 0,3805 | Eps8l2 | -5,08282318 | 3,29E-07 |
| Six4 | 5,39686 | 3,13837 | 2,79779 | 5,72393 | 0,756167 | 0,796468 | 0,504659 | 2,06845 | 2,14395 | 2,06541 | Six4 | -2,34284167 | 3,40E-07 |
| Iqgap2 | 17,5274 | 18,7498 | 7,26964 | 6,89855 | 1,5977 | 0,219394 | 0,151 | 0,0997208 | 0,146326 | 0,230099 | Iqgap2 | -3,73648562 | 3,52E-07 |
| Nrg1 | 10,8103 | 7,79914 | 2,96274 | 0,650111 | 0,320408 | 0,121554 | 0,0495285 | 0,14651 | 0,0542459 | 0 | Nrg1 | -4,43945377 | 3,53E-07 |
| Gale | 30,6666 | 21,6082 | 15,0104 | 15,5589 | 2,42298 | 3,4812 | 3,96916 | 0,209035 | 0,700782 | 0,825134 | Gale | -2,1583763 | 3,60E-07 |
| Nrcam | 0,340602 | 0,714014 | 1,67035 | 3,13164 | 0,0723287 | 0,063928 | 0,0391145 | 0,0387676 | 0,0142064 | 0 | Nrcam | -4,02179051 | 3,64E-07 |
| Bmp4 | 16,6205 | 42,83 | 65,0594 | 35,2381 | 4,05951 | 0,258248 | 0,315649 | 0,155553 | 0,230576 | 0,389318 | Bmp4 | -3,93794842 | 3,70E-07 |
| Slc14a1 | 0,179599 | 1,4534 | 2,94607 | 4,80048 | 0,0508909 | 0 | 0 | 0,081678 | 0 | 0 | Slc14a1 | -5,37680306 | 3,74E-07 |
| Reep2 | 7,84874 | 5,28833 | 6,98527 | 5,5798 | 1,2677 | 0,807798 | 1,09745 | 0 | 0,180433 | 0 | Reep2 | -2,19457883 | 3,80E-07 |
| Onecut2 | 0,161628 | 0,0683348 | 0,520404 | 0,581663 | 0,00653555 | 0 | 0 | 0 | 0 | 0 | Onecut2 | -5,9220886 | 3,80E-07 |
| Apela | 10,8465 | 28,1352 | 5,36376 | 2,02211 | 0,228827 | 0 | 0 | 0 | 0 | 0 | Apela | -5,18544287 | 3,83E-07 |
| Ntng2 | 4,15315 | 1,52728 | 2,83361 | 6,46032 | 0,18841 | 0 | 0 | 1,80509 | 1,63339 | 2,6485 | Ntng2 | -4,24551874 | 3,88E-07 |
| Pcyt1b | 2,22058 | 1,0953 | 2,62301 | 1,31889 | 0,203874 | 0,0490658 | 0,0200337 | 0 | 0,0218503 | 0 | Pcyt1b | -3,74543259 | 4,13E-07 |
| BC068157 | 0,899045 | 1,22803 | 1,5009 | 3,48395 | 0,156718 | 0,0346407 | 0,0847594 | 0,125866 | 0,0693454 | 0,067081 | BC068157 | -3,59575937 | 4,16E-07 |
| **Hoxc8** | 1,27257 | 0,557941 | 4,60276 | 1,85169 | 0 | 0 | 0 | 0 | 0 | 0 | Hoxc8 | -5,907885 | 4,19E-07 |
| Prss23 | 14,7626 | 16,5581 | 7,68034 | 8,65879 | 3,0376 | 1,62223 | 1,51905 | 1,83889 | 2,13348 | 2,29127 | Prss23 | -2,23835319 | 4,22E-07 |
| Ric3 | 1,50597 | 1,0697 | 0,955407 | 2,24569 | 0,124796 | 0,120666 | 0,164916 | 0,0548304 | 0,281729 | 0,175237 | Ric3 | -3,05263068 | 4,22E-07 |
| Cdca5 | 29,3687 | 20,44 | 13,2337 | 20,1449 | 4,85733 | 1,76558 | 2,76646 | 0 | 0,060649 | 0 | Cdca5 | -2,35685809 | 4,24E-07 |
| Edem1 | 16,8796 | 12,9809 | 9,58836 | 8,04191 | 3,25296 | 1,37703 | 1,28846 | 0,736384 | 1,76475 | 1,09786 | Edem1 | -2,28026947 | 4,34E-07 |
| **Hoxb6** | 2,20379 | 1,63046 | 3,44857 | 2,77411 | 0 | 0 | 0 | 0 | 0 | 0 | Hoxb6 | -5,81514638 | 4,52E-07 |
| Mab21l2 | 1,95828 | 1,81109 | 3,67774 | 7,64459 | 0,312295 | 0,092102 | 0,112624 | 0 | 0 | 0,0793062 | Mab21l2 | -3,79456433 | 4,62E-07 |
| Gatm | 3,69692 | 6,59258 | 5,45934 | 4,29708 | 0,9678 | 0,35687 | 0,305415 | 0,258343 | 0,334225 | 0,230508 | Gatm | -2,76214643 | 4,83E-07 |
| Tle1 | 15,4214 | 14,5789 | 20,4015 | 13,6948 | 4,84481 | 1,42004 | 1,68284 | 3,56627 | 2,7019 | 3,40292 | Tle1 | -2,44086836 | 5,09E-07 |
| 6530402F18R | 1,13653 | 1,03111 | 0,872447 | 1,1283 | 0,105754 | 0,0187008 | 0,0686326 | 0,883082 | 0,706709 | 1,1391 | 6530402F18R | -3,72984495 | 5,17E-07 |
| Tuft1 | 7,649 | 11,5103 | 7,89942 | 6,13832 | 1,94826 | 0,748108 | 0,715882 | 0,235618 | 0,217495 | 0,126042 | Tuft1 | -2,40323011 | 5,40E-07 |
| Hes3 | 2,45423 | 1,6761 | 1,26055 | 2,47202 | 0 | 0 | 0 | 0 | 0 | 0 | Hes3 | -5,79527212 | 5,48E-07 |
| Disp2 | 1,51818 | 0,795647 | 0,594036 | 0,28682 | 0 | 0 | 0,0145303 | 0 | 0 | 0 | Disp2 | -4,7442996 | 5,65E-07 |
| Nkx6-1 | 1,15714 | 2,28298 | 2,41448 | 0,971263 | 0 | 0 | 0 | 0 | 0 | 0 | Nkx6-1 | -5,7818978 | 5,69E-07 |
| Cmtm8 | 21,2087 | 27,6583 | 15,2992 | 11,5815 | 1,73924 | 3,45024 | 3,102 | 6,49819 | 11,3834 | 7,28652 | Cmtm8 | -2,24915537 | 5,88E-07 |
| Ttr | 10,6123 | 5,08471 | 3,7957 | 42,4396 | 0,431085 | 0,535215 | 0,280024 | 0 | 0,205742 | 0,0989169 | Ttr | -4,13330596 | 5,93E-07 |
| Tmprss2 | 2,27362 | 1,52926 | 0,172526 | 4,37293 | 0,0292906 | 0 | 0 | 0 | 0 | 0 | Tmprss2 | -5,52914228 | 5,98E-07 |
| Pknox2 | 11,3144 | 6,97654 | 7,84792 | 9,19652 | 1,96582 | 0,549026 | 0,587605 | 4,56628 | 6,46752 | 6,35604 | Pknox2 | -2,68459584 | 6,09E-07 |
| Rad51ap1 | 11,0971 | 16,5133 | 6,1224 | 8,90415 | 2,72246 | 1,60717 | 2,1374 | 0,170684 | 0,126687 | 0,12218 | Rad51ap1 | -2,05460575 | 6,10E-07 |
| Leprel1 | 32,5866 | 26,0125 | 35,515 | 31,9958 | 6,92674 | 1,20357 | 1,19549 | 0,272165 | 0,755017 | 1,02041 | Leprel1 | -2,91952593 | 6,16E-07 |
| Gm1673 | 35,625 | 40,8259 | 95,2552 | 41,6005 | 9,86881 | 8,86256 | 8,04384 | 5,04221 | 5,67665 | 2,40526 | Gm1673 | -2,274048 | 6,28E-07 |
| D8Ertd82e | 19,068 | 12,2992 | 9,90606 | 13,7708 | 3,92802 | 1,50269 | 1,36862 | 0,788766 | 1,47053 | 1,03462 | D8Ertd82e | -2,30167432 | 6,37E-07 |
| 1110038B12R | 158,392 | 352,601 | 171,82 | 155,798 | 38,5836 | 15,3505 | 15,2283 | 3,46404 | 4,49418 | 5,21773 | 1110038B12R | -2,2819565 | 6,68E-07 |
| Zfp850 | 1,87455 | 1,51733 | 0,842558 | 1,23497 | 0,22117 | 0,170614 | 0,195053 | 0,0218805 | 0,128351 | 0,0951106 | Zfp850 | -2,45719479 | 6,89E-07 |
| Dll3 | 0 | 2,62236 | 6,52583 | 5,19804 | 0 | 0,0392625 | 0 | 0 | 0 | 0 | Dll3 | -5,23951629 | 6,93E-07 |
| Casq1 | 53,628 | 112,328 | 207,093 | 261,762 | 31,1681 | 16,8911 | 14,6082 | 10,1676 | 14,3456 | 18,8466 | Casq1 | -2,5079067 | 7,03E-07 |
| Eya1 | 1,29394 | 2,49995 | 4,74729 | 2,94521 | 0,442038 | 0,247866 | 0,139231 | 0,26921 | 0,407227 | 0,147697 | Eya1 | -2,9654769 | 7,12E-07 |
| Hunk | 23,3623 | 15,5943 | 25,3833 | 25,4578 | 7,02165 | 2,36865 | 2,27559 | 0 | 0,127269 | 0,102609 | Hunk | -2,20655586 | 7,36E-07 |
| Cdk5r1 | 3,49511 | 1,91551 | 3,8899 | 3,67791 | 0,660116 | 0,564252 | 0,499841 | 1,05997 | 0,805199 | 0,904341 | Cdk5r1 | -2,24143635 | 7,38E-07 |
| Peg10 | 112,963 | 107,43 | 80,1022 | 72,2007 | 4,98214 | 0,342383 | 0,31783 | 0 | 0,0629777 | 0 | Peg10 | -4,65370017 | 7,46E-07 |
| Rfx4 | 0,245152 | 2,15542 | 1,74319 | 2,88159 | 0,0463045 | 0 | 0,0610264 | 0 | 0 | 0 | Rfx4 | -4,66094792 | 7,50E-07 |
| Cdc25c | 22,2411 | 17,0647 | 13,9963 | 19,5581 | 4,75838 | 1,59649 | 1,46344 | 0 | 0 | 0 | Cdc25c | -2,33583017 | 7,82E-07 |
| Unc5b | 50,6927 | 54,5057 | 75,4611 | 67,2165 | 16,1131 | 3,56513 | 3,47988 | 1,12733 | 1,70799 | 1,77465 | Unc5b | -2,60043486 | 7,85E-07 |
| Tspan8 | 0,466457 | 6,18719 | 2,54246 | 3,42647 | 0 | 0 | 0 | 0 | 0,177121 | 0,0852477 | Tspan8 | -5,78421535 | 8,00E-07 |
| D030025E07R | 0,28558 | 3,23979 | 1,07264 | 1,58225 | 0 | 0 | 0 | 0 | 0 | 0 | D030025E07R | -5,76253041 | 8,17E-07 |
| Sfrp5 | 59,6702 | 29,0749 | 4,07483 | 3,82479 | 0,667065 | 0,863236 | 0,832908 | 0,656465 | 1,33909 | 0,821982 | Sfrp5 | -4,16156917 | 8,22E-07 |
| Klf1 | 1,38661 | 3,41961 | 4,822 | 3,72403 | 0,131272 | 0,116306 | 0,213121 | 0 | 0,156089 | 0,0751847 | Klf1 | -3,81205614 | 8,41E-07 |
| Dlx3 | 1,01977 | 1,25748 | 0,957577 | 1,92621 | 0 | 0 | 0 | 0,23168 | 0 | 0 | Dlx3 | -5,71196137 | 8,62E-07 |
| Ildr2 | 1,059 | 1,11933 | 2,77825 | 3,58229 | 0,235574 | 0,0567819 | 0,0115812 | 0,0344426 | 0,0630816 | 0,0610701 | Ildr2 | -3,79629985 | 8,65E-07 |
| Fgg | 0,23305 | 2,98863 | 1,16703 | 6,80666 | 0 | 0 | 0 | 0 | 0 | 0 | Fgg | -5,83893116 | 8,77E-07 |
| Slc1a3 | 4,88109 | 11,2856 | 7,19373 | 5,99359 | 1,20994 | 0,324784 | 0,163601 | 0,208174 | 0,306036 | 0,098663 | Slc1a3 | -3,09765338 | 8,82E-07 |
| Gmds | 11,7697 | 18,2283 | 8,64431 | 11,8536 | 3,87786 | 3,16938 | 3,03937 | 3,20068 | 4,13484 | 2,451 | Gmds | -2,0109343 | 9,26E-07 |
| Adamts18 | 0,196779 | 0,452953 | 3,01136 | 2,97397 | 0,0371579 | 0 | 0 | 0 | 0 | 0 | Adamts18 | -5,29082464 | 9,42E-07 |
| Trpc6 | 0,748725 | 1,51685 | 1,35663 | 2,83168 | 0,120198 | 0,027168 | 0 | 0,0460333 | 0,0363063 | 0 | Trpc6 | -4,38877997 | 9,62E-07 |
| **Hoxc6** | 3,32021 | 1,09176 | 5,72695 | 2,89813 | 0 | 0 | 0,0680284 | 0,200711 | 0,0747039 | 0 | Hoxc6 | -4,62035486 | 9,62E-07 |
| 1700010I14Ri | 1,09929 | 3,43401 | 1,68211 | 1,47636 | 0,0519947 | 0 | 0 | 0 | 0,0616838 | 0 | 1700010I14Ri | -5,11294325 | 1,01E-06 |
| Ier5l | 29,2564 | 16,1082 | 27,0678 | 31,293 | 6,37637 | 1,99717 | 2,68383 | 2,46749 | 2,14393 | 1,99182 | Ier5l | -2,48222566 | 1,01E-06 |
| Ggta1 | 17,5261 | 22,719 | 14,4006 | 9,05323 | 3,31085 | 0,777787 | 0,643567 | 0,581163 | 1,28322 | 1,2997 | Ggta1 | -2,89519828 | 1,02E-06 |
| Hapln3 | 22,7782 | 12,8399 | 0,626725 | 6,30215 | 0,284454 | 0,126044 | 0,0769746 | 0 | 0 | 0,0814938 | Hapln3 | -4,6973987 | 1,03E-06 |
| Alas2 | 3,54473 | 19,9418 | 21,3343 | 21,8286 | 1,8277 | 0,564008 | 0,761925 | 0 | 0 | 0,0575419 | Alas2 | -3,21898613 | 1,04E-06 |
| Baiap2l2 | 1,96854 | 6,14944 | 16,2936 | 12,0628 | 0,791178 | 0,494384 | 0,251796 | 0 | 0,110325 | 0 | Baiap2l2 | -3,3762975 | 1,06E-06 |
| Ascl1 | 0,447049 | 1,02901 | 1,24379 | 5,85432 | 0 | 0 | 0 | 0 | 0 | 0 | Ascl1 | -5,79614262 | 1,09E-06 |
| Klc3 | 0 | 0 | 0 | 0 | 0 | 0 | 0 | 0 | 0 | 0 | Klc3 | -3,30283775 | 1,12E-06 |
| Slc4a5 | 1,61191 | 1,34551 | 0,879752 | 0,374791 | 0 | 0,0155237 | 0 | 0 | 0,0207123 | 0 | Slc4a5 | -5,03338381 | 1,16E-06 |
| Itgb3 | 24,0656 | 18,9117 | 23,9233 | 30,9161 | 3,1753 | 0,136924 | 0,318318 | 0,0995776 | 0,0730574 | 0,0530232 | Itgb3 | -3,75087428 | 1,18E-06 |
| Clhc1 | 6,35995 | 5,98861 | 3,61897 | 10,1889 | 0,575279 | 0,582947 | 0,533813 | 0 | 0,195983 | 0 | Clhc1 | -2,88215049 | 1,20E-06 |
| Hps1 | 13,9121 | 8,16347 | 9,00947 | 8,21583 | 2,27802 | 1,41381 | 0,882829 | 0,436032 | 0,402154 | 0,310816 | Hps1 | -2,15904067 | 1,20E-06 |
| Mtcl1 | 6,25428 | 5,57004 | 5,40197 | 6,08304 | 1,35274 | 0,293476 | 0,239416 | 0,59321 | 1,15946 | 1,24859 | Mtcl1 | -2,81463582 | 1,22E-06 |
| Foxa2 | 2,29911 | 1,94402 | 1,09655 | 1,4888 | 0,0465903 | 0 | 0 | 0 | 0 | 0 | Foxa2 | -5,0643957 | 1,32E-06 |
| Tbx3 | 12,6053 | 13,6894 | 17,2518 | 18,5524 | 4,95106 | 2,02362 | 1,7153 | 2,50312 | 2,29643 | 2,78236 | Tbx3 | -2,04896046 | 1,32E-06 |
| Prkcq | 1,86715 | 0,872496 | 2,54284 | 2,57414 | 0,0278498 | 0,123164 | 0,180768 | 1,34049 | 1,48081 | 1,17693 | Prkcq | -3,67829227 | 1,33E-06 |
| Prr11 | 12,824 | 15,4637 | 7,3453 | 11,2221 | 3,32503 | 1,39788 | 1,2915 | 0,306655 | 0,0657893 | 0,0817928 | Prr11 | -2,1960074 | 1,34E-06 |
| Amn | 4,9756 | 3,88572 | 0,86516 | 22,1338 | 0,235431 | 0,0521318 | 0,127396 | 0 | 0 | 0 | Amn | -4,61442421 | 1,35E-06 |
| Slc30a10 | 0,362916 | 0,747858 | 0,858801 | 0,927514 | 0,0153347 | 0 | 0 | 0 | 0 | 0 | Slc30a10 | -5,06634076 | 1,38E-06 |
| Arhgef16 | 2,79926 | 2,16113 | 4,7746 | 4,04611 | 0,414056 | 0,457921 | 0,373304 | 0 | 0 | 0 | Arhgef16 | -2,37631743 | 1,38E-06 |
| Chrna7 | 1,46143 | 1,44167 | 1,62639 | 0,817842 | 0,0460666 | 0 | 0 | 0 | 0 | 0,052689 | Chrna7 | -5,00873152 | 1,38E-06 |
| Pkn3 | 10,7477 | 23,7914 | 11,839 | 10,5035 | 2,89191 | 0,843389 | 1,27204 | 0,580141 | 0,939353 | 0,762465 | Pkn3 | -2,52932072 | 1,44E-06 |
| Actn3 | 2,39307 | 3,03523 | 7,70453 | 2,18125 | 0,355333 | 0,0285773 | 0,104842 | 0,155386 | 0,038197 | 0 | Actn3 | -4,20070826 | 1,54E-06 |
| Matn3 | 0,652678 | 1,82427 | 0 | 2,52056 | 0 | 0 | 0 | 0 | 0 | 0 | Matn3 | -5,68874048 | 1,58E-06 |
| Cdo1 | 12,2857 | 16,4641 | 20,5076 | 17,0427 | 4,87191 | 1,80828 | 1,78144 | 0,210144 | 1,09601 | 0,226248 | Cdo1 | -2,15597427 | 1,58E-06 |

| Gene | d9-11 (1) | d9-11 (2) | d9-11(3) | d9-11(4) | newborn(1) | newborn(2) | newborn(3) | adult(1) | adult(2) | adult(3) | Gene | log2(fold) | FDR(*p* -value) |
|---|---|---|---|---|---|---|---|---|---|---|---|---|---|
| Slain1 | 1,32657 | 1,66555 | 4,1271 | 5,34563 | 0,262119 | 0,151243 | 0,258924 | 0 | 0 | 0 | Slain1 | -3,24403887 | 1,61E-06 |
| 2410076I21Ri | 0 | 0 | 0 | 0 | 0 | 0 | 0 | 0 | 0 | 0 | 2410076I21Ri | -2,80785972 | 1,64E-06 |
| Hnrnpa1 | 194,415 | 201,041 | 113,548 | 131,087 | 56,1284 | 36,9542 | 36,1312 | 16,8121 | 23,1189 | 19,2332 | Hnrnpa1 | -2,00683739 | 1,67E-06 |
| Slit3 | 39,5312 | 40,2536 | 37,3195 | 31,7485 | 8,42008 | 1,51433 | 0,897115 | 0,347613 | 0,935854 | 0,411555 | Slit3 | -2,9458421 | 1,68E-06 |
| Foxh1 | 92,1912 | 27,4396 | 4,64302 | 2,13429 | 0,150563 | 0,200185 | 0,0814903 | 0 | 0,0896586 | 0 | Foxh1 | -5,48054545 | 1,73E-06 |
| Nova1 | 1,47998 | 1,15031 | 1,53488 | 1,58161 | 0,304768 | 0,38162 | 0,274694 | 0,204173 | 0,449005 | 0,666403 | Nova1 | -2,08170112 | 1,74E-06 |
| Apoc2 | 8,96085 | 5,89258 | 3,98752 | 23,4436 | 0,682737 | 0,151948 | 0 | 0 | 0 | 0 | Apoc2 | -4,54846857 | 1,76E-06 |
| Ntf5 | 1,9172 | 2,75095 | 1,74567 | 2,98453 | 0,19782 | 0,08757 | 0,107031 | 0,474647 | 0,175938 | 0,226317 | Ntf5 | -3,45967828 | 1,78E-06 |
| Calca | 3,29205 | 16,2593 | 7,19469 | 3,47899 | 0,506302 | 0 | 0 | 0 | 0 | 0 | Calca | -4,57712234 | 1,79E-06 |
| Dgkk | 4,03322 | 5,18974 | 10,1367 | 8,34866 | 1,91159 | 0,822857 | 0,698132 | 0,0798349 | 0 | 0 | Dgkk | -2,23904021 | 1,82E-06 |
| Lgr5 | 0,682592 | 1,53597 | 0,452053 | 0,954883 | 0,0596558 | 0 | 0 | 0 | 0 | 0 | Lgr5 | -4,52850546 | 1,82E-06 |
| Fgf15 | 1,53121 | 0,944059 | 0,479251 | 7,90424 | 0 | 0 | 0 | 0 | 0 | 0 | Fgf15 | -5,7301034 | 1,83E-06 |
| Kcnab2 | 1,45527 | 1,3292 | 4,25152 | 3,29887 | 0,309277 | 0,18235 | 0,0836418 | 0,082726 | 0 | 0 | Kcnab2 | -3,22696152 | 1,84E-06 |
| Zfp711 | 7,39672 | 7,73468 | 3,14883 | 3,94978 | 1,13151 | 0,555827 | 0,793303 | 0,0673025 | 0 | 0,119599 | Zfp711 | -2,3586324 | 1,88E-06 |
| Ret | 3,53681 | 1,74508 | 1,34574 | 2,19285 | 0,265988 | 0,0390716 | 0,111552 | 0,142094 | 0,156339 | 0,0168111 | Ret | -3,388186 | 1,91E-06 |
| **Hoxd9** | 0,32321 | 1,27536 | 2,96746 | 7,48819 | 0,0458457 | 0 | 0 | 0 | 0 | 0 | Hoxd9 | -5,35166609 | 1,91E-06 |
| Gm13315 | 6,38294 | 8,5515 | 6,47942 | 8,25402 | 3,1164 | 2,77349 | 2,95735 | 2,34911 | 2,62151 | 2,28191 | Gm13315 | -2,47837213 | 1,97E-06 |
| Cenpe | 17,5807 | 18,1336 | 13,7226 | 14,8899 | 5,28374 | 1,20255 | 1,79045 | 0,0729354 | 0,0935249 | 0 | Cenpe | -2,2905803 | 1,98E-06 |
| Pcsk1n | 2,46393 | 0,607655 | 2,31362 | 1,55125 | 0 | 0,0386635 | 0 | 0 | 0 | 0,0499521 | Pcsk1n | -5,09113963 | 2,00E-06 |
| Add2 | 0,723144 | 2,07408 | 6,47263 | 4,34948 | 0,288974 | 0,0293125 | 0,0304832 | 0 | 0 | 0 | Add2 | -3,94020219 | 2,05E-06 |
| Dock11 | 9,52997 | 9,02904 | 10,2785 | 8,31889 | 2,20424 | 0,484133 | 0,404573 | 0,0429541 | 0,157469 | 0,152411 | Dock11 | -2,68119404 | 2,14E-06 |
| St6galnac3 | 2,15652 | 7,85501 | 2,49241 | 2,42341 | 0,422999 | 0,103906 | 0,0254208 | 0 | 0,249724 | 0,0536633 | St6galnac3 | -3,67763212 | 2,19E-06 |
| Cpn1 | 1,54678 | 0,381464 | 3,22749 | 5,0634 | 0 | 0,0485993 | 0 | 0 | 0 | 0 | Cpn1 | -5,18510093 | 2,20E-06 |
| Gm266 | 7,32082 | 6,31891 | 3,05483 | 3,07156 | 0,433691 | 0,0769235 | 0 | 0,276057 | 0 | 0 | Gm266 | -4,07759665 | 2,25E-06 |
| Dkkl1 | 2,66364 | 3,06536 | 4,81586 | 2,68125 | 0,126524 | 0 | 0 | 0 | 0 | 0 | Dkkl1 | -4,55515748 | 2,27E-06 |
| Foxl2os | 1,38487 | 0,512312 | 1,22821 | 0,26161 | 0 | 0 | 0 | 0 | 0 | 0 | Foxl2os | -5,60273582 | 2,29E-06 |
| Zic3 | 0,398418 | 0,393039 | 0,798157 | 3,45215 | 0,0225753 | 0 | 0 | 0 | 0 | 0 | Zic3 | -5,26507868 | 2,36E-06 |
| Tgfb1 | 59,0123 | 44,1406 | 32,4758 | 27,7622 | 11,7281 | 4,11204 | 4,32972 | 5,7412 | 6,75909 | 4,83959 | Tgfb1 | -2,27786541 | 2,38E-06 |
| Cdh4 | 1,24146 | 0,587857 | 0,82903 | 2,10148 | 0,126557 | 0,0372893 | 0,0456282 | 0,858982 | 0,381277 | 0,401097 | Cdh4 | -3,47464727 | 2,40E-06 |
| Elovl4 | 3,32461 | 3,2797 | 3,58069 | 5,72635 | 0,425566 | 0,0268893 | 0 | 0 | 0 | 0 | Elovl4 | -4,00384743 | 2,46E-06 |
| Jakmip2 | 0 | 0,820039 | 3,85478 | 4,4666 | 0 | 0 | 0 | 0 | 0 | 0 | Jakmip2 | -5,7721199 | 2,49E-06 |
| **Hoxa3** | 0,654075 | 2,83904 | 4,36754 | 3,03098 | 0,185462 | 0 | 0,0401314 | 0,11887 | 0,0877903 | 0,0847907 | Hoxa3 | -4,02423085 | 2,50E-06 |
| Tfap2a | 0,843256 | 1,70293 | 6,27057 | 9,03581 | 0,130386 | 0,0576748 | 0,176326 | 0 | 0,0385461 | 0,111732 | Tfap2a | -3,95056397 | 2,53E-06 |
| Rgs9bp | 1,52771 | 1,31871 | 1,35494 | 1,3469 | 0,135185 | 0,274831 | 0,27781 | 0 | 0,0159353 | 0,0154233 | Rgs9bp | -2,19711726 | 2,61E-06 |
| Gdf11 | 4,77301 | 11,1119 | 14,1821 | 13,4612 | 1,41306 | 1,87144 | 1,9938 | 0 | 0,0643096 | 0 | Gdf11 | -2,15353027 | 2,61E-06 |
| Prcp | 20,2691 | 23,601 | 14,2374 | 9,37308 | 4,9919 | 2,083 | 2,58137 | 0,604162 | 1,59609 | 1,54234 | Prcp | -2,06608359 | 2,67E-06 |
| Slco4a1 | 1,90822 | 1,5687 | 0,973366 | 0,747613 | 0,090142 | 0 | 0,0325062 | 0 | 0 | 0,0686491 | Slco4a1 | -3,97410666 | 2,68E-06 |
| D430041D05F | 0,337819 | 0,393854 | 0,281993 | 0,309454 | 0,0173876 | 0,0153657 | 0,00940257 | 0 | 0,0102398 | 0,00991507 | D430041D05F | -3,68182569 | 2,76E-06 |
| Msx1 | 10,9191 | 16,4637 | 17,0417 | 20,5664 | 2,01526 | 0,0892142 | 0,0545189 | 0,483476 | 0,358523 | 0,403503 | Msx1 | -3,97666261 | 2,80E-06 |
| Vtn | 0,398343 | 2,35772 | 14,7949 | 12,2372 | 0,226151 | 0 | 0,0611856 | 1,26487 | 3,35622 | 1,74737 | Vtn | -4,87919923 | 2,89E-06 |
| Cdh8 | 1,03171 | 0,934069 | 1,2897 | 0,828246 | 0,0584816 | 0,0258647 | 0 | 0 | 0 | 0,0334012 | Cdh8 | -4,24617268 | 2,95E-06 |
| Zfp423 | 12,065 | 13,9498 | 9,09634 | 17,1231 | 2,07612 | 0,129948 | 0,158985 | 0,0590335 | 0,151723 | 0,272601 | Zfp423 | -3,53374327 | 3,00E-06 |
| **Hoxa1** | 3,90966 | 2,26089 | 0,562637 | 0,679053 | 0,0382292 | 0 | 0 | 0 | 0 | 0 | Hoxa1 | -5,18459125 | 3,02E-06 |
| Rab27a | 7,95446 | 3,70553 | 3,31977 | 3,00512 | 0,852842 | 0,180067 | 0,304909 | 0,200877 | 0,407257 | 0,357769 | Rab27a | -2,92770105 | 3,05E-06 |
| **Hoxa5** | 0,549977 | 1,44677 | 5,96744 | 1,29259 | 0,0520261 | 0 | 0 | 0,166382 | 0,0617214 | 0,119065 | Hoxa5 | -5,15070174 | 3,05E-06 |
| Mcoln3 | 1,01785 | 1,40571 | 2,37868 | 1,6403 | 0,0577905 | 0,0511841 | 0,125085 | 0 | 0 | 0 | Mcoln3 | -3,99111759 | 3,06E-06 |
| Foxb1 | 0,769716 | 2,60338 | 0,183565 | 1,77244 | 0 | 0 | 0 | 0 | 0 | 0 | Foxb1 | -5,59275365 | 3,13E-06 |
| Fbxl16 | 1,20511 | 2,65201 | 3,36593 | 3,73577 | 0,341493 | 0,44142 | 0,170501 | 0,84311 | 1,14813 | 1,37997 | Fbxl16 | -2,67760531 | 3,31E-06 |
| Stmn2 | 0,90195 | 7,29596 | 10,991 | 13,4459 | 0,511906 | 0,226626 | 0 | 0,163729 | 0,485774 | 0,527155 | Stmn2 | -4,17305889 | 3,37E-06 |
| Scrt2 | 0 | 2,07772 | 4,47498 | 1,35031 | 0 | 0 | 0 | 0 | 0 | 0 | Scrt2 | -5,68380883 | 3,39E-06 |
| Cpm | 4,15562 | 3,43637 | 2,34655 | 3,20186 | 0,432713 | 0,0153019 | 0,0561654 | 0,166867 | 0,347067 | 0,25679 | Cpm | -3,66847851 | 3,41E-06 |
| Mpped1 | 0,400592 | 0,592772 | 0,585155 | 3,12776 | 0 | 0 | 0 | 0 | 0 | 0 | Mpped1 | -5,57543501 | 3,45E-06 |
| Mybph | 3,06826 | 2,55257 | 4,91575 | 1,93236 | 0 | 0 | 0 | 0 | 0 | 0 | Mybph | -3,91522488 | 3,46E-06 |
| Cit | 14,0738 | 13,7491 | 7,67986 | 9,0408 | 3,0178 | 0,638352 | 0,739286 | 0,0829398 | 0,106404 | 0,0882781 | Cit | -2,53336104 | 3,49E-06 |
| Foxf2 | 0,707968 | 0,41904 | 2,95461 | 1,99691 | 0 | 0 | 0 | 0 | 0 | 0 | Foxf2 | -5,5520934 | 3,52E-06 |
| Lrig3 | 3,33361 | 4,6197 | 11,6605 | 10,3159 | 0,584657 | 0,0596533 | 0 | 0 | 0,238918 | 0,0513458 | Lrig3 | -4,13992915 | 3,55E-06 |
| Fgfr3 | 12,3947 | 9,97406 | 12,9693 | 10,0187 | 2,30433 | 0,170275 | 0,448389 | 0,345832 | 0,609875 | 0,579067 | Fgfr3 | -3,12907109 | 3,65E-06 |
| Elf3 | 2,11943 | 1,74231 | 1,32673 | 6,45301 | 0,150352 | 0,0443728 | 0 | 0 | 0 | 0 | Elf3 | -4,10271983 | 3,67E-06 |
| Foxc2 | 11,1658 | 13,0503 | 9,42122 | 5,99108 | 1,51396 | 0,0913275 | 0,18613 | 0,110309 | 0,081401 | 0,0393193 | Foxc2 | -3,50619899 | 3,72E-06 |
| Zfp57 | 16,7144 | 16,1271 | 16,3955 | 13,2339 | 2,86344 | 0,431501 | 0,397931 | 0 | 0,20474 | 0,331257 | Zfp57 | -3,03676814 | 3,76E-06 |
| Tube1 | 5,60951 | 3,54159 | 7,02332 | 5,99029 | 0,413421 | 1,20972 | 0,86012 | 0,305971 | 0,37601 | 0,436002 | Tube1 | -2,31892781 | 3,79E-06 |
| Sh3bgrl2 | 9,08188 | 6,94798 | 3,71299 | 4,15478 | 1,28677 | 0,859345 | 0,511348 | 0,252856 | 0,279189 | 0,299902 | Sh3bgrl2 | -2,38233326 | 3,96E-06 |
| Mmp17 | 1,95266 | 2,9353 | 3,18218 | 2,99775 | 0,500634 | 0,0466077 | 0,114016 | 0,338273 | 0,217882 | 0,180548 | Mmp17 | -3,08441451 | 3,96E-06 |
| Efhd1 | 2,68299 | 5,86054 | 4,47864 | 2,70241 | 0,59833 | 0,144505 | 0,294318 | 0,173896 | 0,839224 | 0,311265 | Efhd1 | -3,00732473 | 3,98E-06 |
| Dpp7 | 5,07946 | 8,56 | 8,47888 | 4,90281 | 1,86281 | 0,638721 | 0,715369 | 0,959781 | 0,428309 | 0,481623 | Dpp7 | -2,20620676 | 4,03E-06 |
| Rn45s | 29538,3 | 12889,2 | 12765,9 | 24867,9 | 4781,02 | 906,562 | 1072,64 | 1961,67 | 1285,66 | 1567,67 | Rn45s | -2,8568463 | 4,05E-06 |
| Cth | 4,99528 | 5,30676 | 2,08462 | 8,1277 | 0,490784 | 0,193162 | 0,0590129 | 2,26633 | 1,35914 | 1,93476 | Cth | -3,54286731 | 4,25E-06 |
| Elavl3 | 0,189741 | 0,748719 | 6,65203 | 3,88722 | 0,0358276 | 0,0950244 | 0,0968832 | 0 | 0,0422635 | 0,102224 | Elavl3 | -4,24889741 | 4,35E-06 |
| Tnnc2 | 2,56585 | 6,96001 | 7,49277 | 10,3265 | 0,55021 | 0,163362 | 0,198813 | 0 | 0,667353 | 0,42399 | Tnnc2 | -3,49731266 | 4,35E-06 |
| Smad9 | 2,71041 | 4,06887 | 5,60696 | 4,03702 | 1,31832 | 0,560596 | 0,487351 | 0,375423 | 0,964385 | 0,647482 | Smad9 | -2,0527977 | 4,42E-06 |
| Prkcb | 1,34224 | 1,77712 | 2,56519 | 1,60755 | 0,289998 | 0,053026 | 0,0108154 | 0 | 0 | 0,0228117 | Prkcb | -3,33012601 | 4,43E-06 |
| Tub | 0,584653 | 0,314574 | 2,08506 | 1,87429 | 0,10062 | 0,0266084 | 0,0162788 | 0,0483812 | 0 | 0 | Tub | -3,995134 | 4,44E-06 |
| E130006D01R | 13,5933 | 13,0288 | 10,9967 | 3,60936 | 1,14011 | 0,431765 | 0,131625 | 0 | 0,145861 | 0,279607 | E130006D01R | -3,343636 | 4,59E-06 |
| Six3os1 | 0,33828 | 1,65606 | 1,67464 | 4,82077 | 0 | 0 | 0 | 0 | 0 | 0 | Six3os1 | -5,57064784 | 4,78E-06 |
| Ripk3 | 29,6944 | 29,4167 | 6,46544 | 5,60979 | 1,92653 | 0,313122 | 0,339855 | 0,334772 | 0,434701 | 0,119792 | Ripk3 | -3,63901882 | 4,85E-06 |
| Apom | 10,4007 | 3,62085 | 1,53138 | 33,8582 | 0 | 0 | 0 | 0 | 0 | 0,201972 | Apom | -5,09222755 | 4,97E-06 |
| **Hoxa10** | 0,469595 | 0,926404 | 1,30846 | 2,52365 | 0 | 0 | 0 | 0 | 0 | 0 | Hoxa10 | -5,47531979 | 4,97E-06 |
| Kcnk5 | 12,2781 | 6,29459 | 3,3086 | 3,79863 | 0,684942 | 0,0484647 | 0,0296386 | 0,0879205 | 0,129481 | 0,219039 | Kcnk5 | -3,97589291 | 5,00E-06 |
| Robo3 | 0,472289 | 0,133118 | 3,60439 | 0,543832 | 0 | 0 | 0,0206721 | 0 | 0 | 0 | Robo3 | -4,65180778 | 5,32E-06 |
| Sesn2 | 30,3272 | 15,6359 | 11,4588 | 29,7974 | 4,63534 | 3,94475 | 3,90481 | 2,49531 | 2,13496 | 3,43721 | Sesn2 | -2,08412819 | 5,39E-06 |
| Ninl | 4,49213 | 7,20119 | 7,4994 | 3,89747 | 1,60806 | 0,624904 | 0,57342 | 0,170354 | 0,396046 | 0,443681 | Ninl | -2,16642271 | 5,46E-06 |
| Tspan32 | 6,02461 | 4,51961 | 1,96462 | 3,86601 | 0,175254 | 0,0592408 | 0 | 0 | 0 | 0 | Tspan32 | -4,00258252 | 5,59E-06 |
| Tbc1d9 | 3,02535 | 5,59656 | 6,04283 | 3,70149 | 1,39223 | 0,494353 | 0,430249 | 0,280703 | 0,598065 | 0,453891 | Tbc1d9 | -2,18636693 | 5,66E-06 |
| Trp53i11 | 22,7321 | 18,9262 | 29,0844 | 26,5152 | 8,11502 | 2,46618 | 2,25218 | 1,69649 | 4,84197 | 2,94346 | Trp53i11 | -2,15207917 | 5,70E-06 |
| Bin1 | 43,9279 | 38,7642 | 21,9989 | 21,4006 | 4,21426 | 0,280783 | 0,490323 | 0,435161 | 0,429593 | 0,25913 | Bin1 | -3,47239655 | 5,79E-06 |
| Prss12 | 2,4189 | 0,753543 | 4,03803 | 3,71933 | 0,21658 | 0,0319388 | 0,0781072 | 0 | 0,0854196 | 0,0412542 | Prss12 | -3,57938935 | 5,87E-06 |
| Efna3 | 0,906016 | 1,48956 | 3,78063 | 5,77807 | 0 | 0 | 0,0929682 | 0,273336 | 0,204909 | 0 | Efna3 | -4,92709558 | 5,88E-06 |
| Pycr1 | 2,25629 | 6,9 | 6,87406 | 4,43247 | 0,895409 | 0,608238 | 0,31137 | 0,205112 | 0,302494 | 0,109609 | Pycr1 | -2,39726436 | 6,19E-06 |
| Nkx2-3 | 3,3722 | 2,53456 | 0,804182 | 0,323511 | 0 | 0 | 0 | 0 | 0 | 0 | Nkx2-3 | -5,4758897 | 6,19E-06 |
| Myt1 | 0 | 0,766788 | 3,14427 | 0,903646 | 0 | 0 | 0 | 0 | 0 | 0 | Myt1 | -5,57627614 | 6,42E-06 |
| Chac1 | 6,30323 | 4,66344 | 1,87884 | 19,045 | 0,0639293 | 0,623006 | 0,207575 | 3,87809 | 9,87921 | 12,0081 | Chac1 | -3,87409355 | 6,56E-06 |
| Six1 | 1,27161 | 2,28079 | 3,95614 | 4,19256 | 0,163852 | 0,318905 | 0,283626 | 1,47116 | 2,09264 | 1,75982 | Six1 | -2,8461002 | 6,58E-06 |
| Il2rg | 9,59957 | 15,4958 | 11,8359 | 2,85658 | 1,54876 | 0,98782 | 0,536385 | 0,395693 | 1,39873 | 1,41886 | Il2rg | -2,70196617 | 6,63E-06 |
| 5430416N02R | 49,1718 | 39,316 | 15,5394 | 37,473 | 8,96044 | 4,37634 | 5,32122 | 0,667466 | 0 | 0,741383 | 5430416N02R | -2,2017332 | 6,63E-06 |
| Dpysl2 | 57,3085 | 62,3378 | 73,0802 | 51,8499 | 22,9405 | 6,85163 | 6,65704 | 1,31353 | 2,65419 | 1,58707 | Dpysl2 | -2,00716186 | 6,89E-06 |
| Adcy3 | 3,16516 | 6,21897 | 6,54869 | 6,08983 | 2,05703 | 0 | 0 | 0 | 0 | 0 | Adcy3 | -2,33727249 | 7,01E-06 |
| Camk2n2 | 3,14412 | 3,94739 | 3,57817 | 2,01482 | 0,243755 | 0,0720397 | 0,26388 | 1,55275 | 0,48433 | 0,83866 | Camk2n2 | -3,33504626 | 7,07E-06 |
| Rhpn1 | 3,47332 | 0,435249 | 2,08684 | 3,34669 | 0,0856675 | 0 | 0,0463467 | 0 | 0 | 0 | Rhpn1 | -4,06842376 | 7,72E-06 |
| Pyy | 34,6354 | 27,768 | 5,41972 | 0 | 0 | 0 | 0 | 0 | 0 | 0 | Pyy | -5,55465812 | 7,72E-06 |
| Treml2 | 0,496172 | 0,909024 | 1,77499 | 1,21407 | 0,120472 | 0,0532594 | 0,021719 | 0 | 0 | 0,0114597 | Treml2 | -3,44611589 | 8,27E-06 |
| Klk8 | 17,9058 | 14,1306 | 8,27658 | 4,99339 | 1,25272 | 0,0694089 | 0,169511 | 0,249477 | 1,02624 | 0,359092 | Klk8 | -3,7320975 | 8,49E-06 |
| Atp1a3 | 0 | 4,25845 | 10,3593 | 6,43302 | 0,152899 | 0,0450741 | 0,0551336 | 0 | 0,21069 | 0,261904 | Atp1a3 | -4,55621779 | 8,51E-06 |
| Atp2a3 | 10,184 | 5,92842 | 7,94044 | 7,49484 | 1,68636 | 0,247907 | 0,216634 | 0,160838 | 0,378132 | 0,202304 | Atp2a3 | -2,98120867 | 8,61E-06 |
| Nsg2 | 1,46645 | 1,15731 | 4,1858 | 2,51129 | 0,222846 | 0,0394493 | 0,144675 | 0,142677 | 0,0528073 | 0 | Nsg2 | -3,49068885 | 8,69E-06 |
| Diras2 | 1,54654 | 2,54277 | 5,71696 | 2,00339 | 0,479835 | 0,0922333 | 0,0225669 | 0 | 0,0246221 | 0,0238153 | Diras2 | -3,49790829 | 8,76E-06 |
| Lmx1a | 0,292452 | 0,577004 | 2,03425 | 1,37497 | 0,0276263 | 0 | 0 | 0 | 0 | 0,0631061 | Lmx1a | -4,91058939 | 8,92E-06 |
| Grm2 | 0,391945 | 0,96663 | 1,06326 | 0,394874 | 0 | 0 | 0 | 0 | 0 | 0 | Grm2 | -5,33162547 | 8,99E-06 |
| Map4k1 | 4,90558 | 6,77502 | 3,78752 | 1,35903 | 0,382465 | 0,492204 | 0,451406 | 0,334387 | 0,246783 | 0,317867 | Map4k1 | -2,68581693 | 9,00E-06 |
| Spin2c | 12,0196 | 4,13607 | 3,73256 | 1,97041 | 0,238354 | 0,352189 | 0,172018 | 0 | 0 | 0 | Spin2c | -3,56497478 | 9,10E-06 |
| Apoa4 | 2,73048 | 4,35109 | 0 | 19,0393 | 0,0596325 | 0 | 0 | 0 | 0 | 0 | Apoa4 | -5,35170078 | 9,24E-06 |
| Myb | 0,949875 | 1,87409 | 1,7443 | 0,86101 | 0,188424 | 0,0856538 | 0,0582386 | 0,0863864 | 0 | 0,0614843 | Myb | -2,97875846 | 9,44E-06 |
| Neurog2 | 0,456441 | 1,05063 | 1,90487 | 2,45221 | 0 | 0 | 0,0466955 | 0 | 0 | 0 | Neurog2 | -4,82077189 | 9,73E-06 |
| Rubie | 2,5643 | 1,7707 | 1,07007 | 1,03298 | 0 | 0 | 0,0788662 | 0 | 0 | 0 | Rubie | -4,76789271 | 9,77E-06 |

| Gene | d9-11 (1) | d9-11 (2) | d9-11(3) | d9-11(4) | newborn(1) | newborn(2) | newborn(3) | adult(1) | adult(2) | adult(3) | Gene | log2(fold) | FDR(*p*-value) |
|---|---|---|---|---|---|---|---|---|---|---|---|---|---|
| Cacng4 | 0,397005 | 2,3497 | 3,97558 | 9,59222 | 0 | 0,100387 | 0 | 0 | 0 | 0 | Cacng4 | -5,00587391 | 9,78E-06 |
| Greb1l | 9,93716 | 8,55941 | 2,90943 | 3,17573 | 1,03926 | 0,417497 | 0,193013 | 0,0337672 | 0 | 0,011974 | Greb1l | -2,87308977 | 9,95E-06 |
| Kcnk1 | 1,16697 | 1,2951 | 1,09579 | 3,67361 | 0,0827436 | 0,0732302 | 0 | 0,265011 | 0,0489944 | 0,0473022 | Kcnk1 | -4,13009063 | 9,98E-06 |
| Man1a | 21,376 | 22,1499 | 13,2092 | 10,6013 | 5,23477 | 1,78524 | 1,72673 | 7,98939 | 7,9547 | 5,97692 | Man1a | -2,13457596 | 9,98E-06 |
| F2rl3 | 13,7885 | 2,91471 | 1,64405 | 1,98402 | 0,111826 | 0,198069 | 0,181528 | 0 | 0,531008 | 0,191999 | F2rl3 | -3,99910214 | 1,00E-05 |
| Rin1 | 4,76439 | 2,97545 | 3,33548 | 2,25633 | 0,718413 | 0,1155 | 0,117745 | 0 | 0,205578 | 0,14912 | Rin1 | -2,96972466 | 1,04E-05 |
| Trib3 | 3,65034 | 10,4756 | 4,15469 | 17,215 | 1,45931 | 0,541748 | 0,458443 | 0,45195 | 0,61381 | 0,26923 | Trib3 | -2,98002863 | 1,04E-05 |
| Gjb2 | 0,55653 | 1,23527 | 1,04517 | 4,34489 | 0,0394561 | 0 | 0,0426901 | 0 | 0 | 0 | Gjb2 | -4,6228818 | 1,05E-05 |
| Vat1l | 0,176986 | 0,611087 | 2,65918 | 3,56626 | 0,0501495 | 0 | 0 | 0 | 0 | 0 | Vat1l | -4,87406838 | 1,05E-05 |
| Lrfn1 | 7,92356 | 3,58054 | 4,29187 | 6,68715 | 0,765413 | 1,03201 | 0,670381 | 0,36665 | 0,172513 | 0,249943 | Lrfn1 | -2,09482864 | 1,05E-05 |
| Bcat1 | 30,4993 | 18,8789 | 15,4761 | 15,6457 | 0,667213 | 0,00998251 | 0,0366478 | 0,036327 | 0,0399269 | 0,0515356 | Bcat1 | -4,82242486 | 1,07E-05 |
| Gabra4 | 1,24621 | 3,91866 | 3,77083 | 0,941685 | 0,375121 | 0,175607 | 0,0716073 | 0,141727 | 0 | 0,0755747 | Gabra4 | -3,10573554 | 1,10E-05 |
| Gstt1 | 35,6465 | 27,471 | 11,4652 | 11,589 | 2,74823 | 3,18992 | 2,86162 | 13,7566 | 13,9186 | 11,1737 | Gstt1 | -2,44910204 | 1,12E-05 |
| Vgll2 | 0,243692 | 3,12511 | 2,23725 | 1,9634 | 0 | 0 | 0 | 0 | 0 | 0 | Vgll2 | -5,34226635 | 1,13E-05 |
| Pcdhac2 | 0 | 0 | 0 | 0 | 0 | 0 | 0 | 0 | 0 | 0 | Pcdhac2 | -4,30170384 | 1,15E-05 |
| Fgf10 | 6,32198 | 2,20522 | 0,903812 | 1,47804 | 0,178091 | 0 | 0,0214043 | 0 | 0,0233498 | 0 | Fgf10 | -4,29508944 | 1,16E-05 |
| Soat1 | 31,1664 | 30,9017 | 19,9294 | 18,2179 | 4,17838 | 0,233495 | 0,428499 | 0 | 0,133537 | 0,0215308 | Soat1 | -3,33766103 | 1,18E-05 |
| Hes5 | 2,01787 | 0,853084 | 2,16521 | 4,06398 | 0 | 0 | 0 | 0 | 0 | 0 | Hes5 | -5,29759497 | 1,18E-05 |
| Rpl38 | 661,572 | 972,063 | 731,743 | 754,54 | 95,2508 | 78,7575 | 61,1958 | 46,7931 | 48,0063 | 56,5485 | Rpl38 | -2,15896065 | 1,18E-05 |
| Lipg | 0,766983 | 0,420348 | 3,27218 | 1,37376 | 0,0482922 | 0 | 0,05224 | 0 | 0 | 0 | Lipg | -4,17091156 | 1,19E-05 |
| Fam171b | 1,43648 | 1,3226 | 3,75697 | 3,47332 | 0,298523 | 0,0240028 | 0,0293581 | 0 | 0 | 0 | Fam171b | -3,68569024 | 1,20E-05 |
| Dusp4 | 62,8527 | 42,3768 | 39,7627 | 35,6456 | 9,40969 | 1,38207 | 0,802644 | 0,875642 | 0,462203 | 0,357066 | Dusp4 | -3,07547618 | 1,21E-05 |
| Dab2 | 56,2461 | 33,9229 | 22,8548 | 19,8571 | 7,67645 | 1,81909 | 1,19232 | 0,515039 | 0,851394 | 0,526163 | Dab2 | -2,80418424 | 1,22E-05 |
| 2610524H06R | 13,8038 | 24,1382 | 13,8461 | 13,4749 | 3,69415 | 2,35489 | 1,03255 | 0 | 0 | 0 | 2610524H06R | -2,23965181 | 1,23E-05 |
| Trank1 | 1,06288 | 0,611647 | 0,739362 | 0,267749 | 0,0835776 | 0,0147716 | 0,0180782 | 0 | 0,019687 | 0,0190632 | Trank1 | -3,44631287 | 1,24E-05 |
| Rasl11a | 18,421 | 8,25967 | 10,6214 | 5,73297 | 1,23871 | 0,0845362 | 0 | 0 | 0 | 0,546989 | Rasl11a | -3,62109223 | 1,25E-05 |
| Bin2 | 2,41928 | 4,4322 | 12,8367 | 15,8399 | 1,22581 | 0,13023 | 0,106116 | 0 | 0,0581407 | 0 | Bin2 | -3,64737462 | 1,25E-05 |
| Mtmr7 | 4,1093 | 5,1367 | 6,62727 | 3,08642 | 0 | 0 | 0 | 0 | 0 | 0 | Mtmr7 | -2,01801838 | 1,26E-05 |
| **Hoxd3** | 0,113319 | 0,894303 | 2,55382 | 2,0549 | 0 | 0 | 0,0347517 | 0 | 0 | 0 | Hoxd3 | -4,94619395 | 1,26E-05 |
| Hmga1 | 395,5 | 263,295 | 174,075 | 232,367 | 36,4316 | 28,4714 | 35,0954 | 33,3369 | 23,5365 | 23,4741 | Hmga1 | -2,2020985 | 1,29E-05 |
| Hnf4a | 0,440802 | 0,14495 | 0,367945 | 2,73875 | 0 | 0 | 0 | 0 | 0 | 0 | Hnf4a | -5,37959054 | 1,30E-05 |
| **Hoxa9** | 0,10717 | 0,398593 | 1,48368 | 2,95123 | 0 | 0 | 0 | 0 | 0 | 0 | Hoxa9 | -5,37457527 | 1,34E-05 |
| **Hoxd11** | 0 | 1,04266 | 1,74005 | 1,72018 | 0 | 0 | 0 | 0 | 0,0323495 | 0 | Hoxd11 | -5,37181012 | 1,45E-05 |
| Tmem180 | 10,9149 | 7,27525 | 1,84669 | 5,05181 | 0,794852 | 0,555379 | 0,271588 | 1,74172 | 1,68457 | 1,67415 | Tmem180 | -2,84965578 | 1,45E-05 |
| Wnt7b | 0,0964025 | 0,570603 | 1,04608 | 6,00665 | 0,0273194 | 0 | 0 | 0 | 0,0645546 | 0 | Wnt7b | -5,13130626 | 1,48E-05 |
| Osbpl10 | 11,7683 | 8,34406 | 6,90632 | 4,25993 | 1,35678 | 1,57103 | 0,96 | 2,00259 | 0,804833 | 1,19428 | Osbpl10 | -2,19797916 | 1,48E-05 |
| Chst10 | 1,78233 | 2,27538 | 2,27531 | 1,16187 | 0,326869 | 0,10514 | 0,0321469 | 0 | 0 | 0 | Chst10 | -3,08723927 | 1,49E-05 |
| Nhlh1 | 0,134043 | 0,396696 | 4,36343 | 2,29556 | 0 | 0 | 0 | 0 | 0 | 0 | Nhlh1 | -5,35680769 | 1,53E-05 |
| Foxf1 | 9,64661 | 8,29272 | 1,61035 | 2,77649 | 0,547117 | 0 | 0,0845656 | 0 | 0,0462625 | 0 | Foxf1 | -3,84742911 | 1,54E-05 |
| Wasf3 | 3,69857 | 6,6891 | 2,77843 | 6,20995 | 0,489344 | 0 | 0,0378158 | 0,112048 | 0 | 0,0399438 | Wasf3 | -3,8495868 | 1,56E-05 |
| Tbx4 | 0,554407 | 1,42199 | 1,54522 | 1,09707 | 0,0942899 | 0 | 0 | 0 | 0 | 0 | Tbx4 | -4,22056902 | 1,57E-05 |
| Hapln1 | 59,2967 | 134,61 | 35,8233 | 20,5274 | 0,76583 | 0 | 0,0385287 | 0 | 0,0420179 | 0 | Hapln1 | -5,24905496 | 1,58E-05 |
| Olig3 | 0,493733 | 0,64941 | 1,78576 | 2,65248 | 0 | 0 | 0 | 0 | 0 | 0 | Olig3 | -5,25941814 | 1,58E-05 |
| Gal | 12,0106 | 1,24706 | 4,74678 | 10,1766 | 0,180711 | 0,160945 | 0 | 0 | 0 | 0 | Gal | -4,16137235 | 1,59E-05 |
| Amer2 | 0,716642 | 0,565565 | 1,79687 | 1,78712 | 0,0344908 | 0 | 0 | 0 | 0 | 0 | Amer2 | -4,75248168 | 1,60E-05 |
| Flt4 | 32,25 | 20,2846 | 4,38129 | 4,30715 | 2,03806 | 0,447018 | 0,381215 | 0,443322 | 1,31902 | 0,734385 | Flt4 | -3,41172125 | 1,63E-05 |
| Dbx1 | 0 | 1,77531 | 2,45796 | 5,10879 | 0 | 0 | 0 | 0 | 0 | 0 | Dbx1 | -5,36013351 | 1,66E-05 |
| Rai14 | 15,8827 | 25,6569 | 22,4276 | 12,8021 | 3,43031 | 0,265164 | 0,202758 | 0,542036 | 0,68557 | 0,534866 | Rai14 | -3,29250712 | 1,71E-05 |
| Ggt7 | 2,92325 | 1,12842 | 4,24345 | 4,60938 | 0,396398 | 0,255082 | 0,155953 | 0 | 0,0852757 | 0,247109 | Ggt7 | -2,84401637 | 1,71E-05 |
| Abcg5 | 1,04112 | 0,88033 | 0,124174 | 3,74543 | 0 | 0 | 0 | 0 | 0 | 0,0482326 | Abcg5 | -5,28464594 | 1,72E-05 |
| Vav3 | 9,124 | 10,2597 | 5,11286 | 3,47146 | 1,13852 | 0,244122 | 0,166153 | 0,295744 | 0,405055 | 0,139954 | Vav3 | -3,14930967 | 1,74E-05 |
| Slc6a4 | 1,44316 | 4,50827 | 3,71414 | 5,20968 | 0,44323 | 0,0301647 | 0,0368866 | 0 | 0,0403275 | 0,0389601 | Slc6a4 | -3,61166073 | 1,74E-05 |
| Aass | 0,611271 | 0,603017 | 3,20718 | 3,95906 | 0,0989725 | 0 | 0 | 0 | 0 | 0,0282538 | Aass | -4,43938606 | 1,74E-05 |
| Nup62cl | 6,34965 | 10,5491 | 1,11558 | 2,35545 | 0,19015 | 0,0843486 | 0 | 0 | 0 | 0,109154 | Nup62cl | -4,30922409 | 1,81E-05 |
| Tenm2 | 0,322271 | 0,921977 | 2,20592 | 2,11075 | 0,127726 | 0,161251 | 0,0592027 | 0,0293178 | 0,0967178 | 0,0103937 | Tenm2 | -2,96898527 | 1,84E-05 |
| Sorcs3 | 0,842381 | 0,332405 | 0,609407 | 0,0565842 | 0 | 0 | 0 | 0 | 0 | 0 | Sorcs3 | -5,23090059 | 1,86E-05 |
| Gdap1l1 | 0,515064 | 0,63513 | 1,71965 | 0,908027 | 0 | 0 | 0 | 0 | 0 | 0,041728 | Gdap1l1 | -5,19660076 | 1,88E-05 |
| Shox2 | 1,10674 | 5,8955 | 16,0724 | 10,9237 | 0,628554 | 0,0556828 | 0 | 0 | 0 | 0,0719844 | Shox2 | -4,2133696 | 1,95E-05 |
| Psd | 0 | 8,59187 | 17,031 | 9,6542 | 1,02979 | 0 | 0 | 0 | 0 | 0 | Psd | -2,12171285 | 1,99E-05 |
| Plvap | 201,293 | 152,016 | 32,7549 | 23,8543 | 6,4311 | 0,977226 | 0,88286 | 0,921425 | 1,70696 | 1,53722 | Plvap | -4,22679071 | 1,99E-05 |
| 2010204K13R | 38,5298 | 57,8925 | 18,3027 | 27,2747 | 4,43292 | 7,72146 | 4,8174 | 0 | 0,970591 | 0,61195 | 2010204K13R | -2,24253227 | 2,00E-05 |
| Mrvi1 | 5,02606 | 5,32358 | 11,7793 | 8,68795 | 1,51636 | 0,152473 | 0,210667 | 0,240813 | 0,635943 | 0,24601 | Mrvi1 | -3,07650711 | 2,00E-05 |
| Chrm4 | 0 | 0 | 0 | 0 | 0 | 0 | 0 | 0 | 0,0803291 | 0,039054 | Chrm4 | -3,73416007 | 2,00E-05 |
| Krt7 | 0,222318 | 10,0882 | 9,46307 | 22,3911 | 0,568189 | 0,223714 | 0,159432 | 0,201587 | 0,675326 | 0,144606 | Krt7 | -4,05920497 | 2,01E-05 |
| Icam1 | 30,6635 | 25,3368 | 8,20325 | 13,8609 | 2,93535 | 0,427415 | 0,361806 | 0,476297 | 0,879429 | 0,806907 | Icam1 | -3,18048398 | 2,02E-05 |
| Icam2 | 36,2042 | 59,4141 | 29,1153 | 15,2444 | 7,86465 | 1,66117 | 1,72335 | 2,67927 | 4,36221 | 2,90195 | Icam2 | -2,62707182 | 2,04E-05 |
| Gbx2 | 2,07224 | 0,786239 | 1,33047 | 1,60569 | 0,0904388 | 0,0400273 | 0,0489296 | 0 | 0,107171 | 0,413734 | Gbx2 | -3,87488907 | 2,05E-05 |
| Bhlha15 | 0,278526 | 0,549529 | 1,08494 | 2,24488 | 0,0263089 | 0 | 0,0284604 | 0 | 0 | 0 | Bhlha15 | -4,47745575 | 2,12E-05 |
| Nr4a3 | 11,501 | 11,3454 | 7,92317 | 12,5039 | 3,16022 | 0,688082 | 0,840936 | 2,48528 | 2,58128 | 2,19339 | Nr4a3 | -2,48968091 | 2,13E-05 |
| Gng3 | 3,3218 | 2,65979 | 5,96502 | 4,39783 | 0 | 0 | 0 | 0 | 0 | 0 | Gng3 | -2,81322709 | 2,17E-05 |
| Cdr2 | 26,8945 | 23,4097 | 17,0562 | 14,2106 | 6,28023 | 1,45545 | 1,37514 | 0,359384 | 1,4157 | 1,28183 | Cdr2 | -2,40105452 | 2,18E-05 |
| Amph | 1,21619 | 0,399613 | 1,18435 | 1,63369 | 0,0573332 | 0,0507123 | 0 | 0,368341 | 1,25594 | 0,68899 | Amph | -3,96593806 | 2,20E-05 |
| Kif5a | 0,49919 | 2,35886 | 3,46821 | 3,61553 | 0,275828 | 0,224964 | 0,20678 | 0,0681462 | 0,116652 | 0,0645132 | Kif5a | -2,64404644 | 2,22E-05 |
| Plac1 | 21,0594 | 8,80242 | 4,4681 | 1,4375 | 0,507919 | 0,270449 | 0 | 0 | 0 | 0 | Plac1 | -4,15804367 | 2,23E-05 |
| **Hoxa2** | 0,976671 | 0,770767 | 2,11942 | 2,16421 | 0 | 0 | 0,0600022 | 0,177231 | 0 | 0,0634614 | Hoxa2 | -4,63814563 | 2,25E-05 |
| Lcp2 | 6,41338 | 3,73854 | 4,54216 | 2,15377 | 0,647141 | 0,0487131 | 0,178743 | 0 | 0,0325367 | 0,0314519 | Lcp2 | -3,24690069 | 2,31E-05 |
| Ctnna2 | 0,242159 | 0,47778 | 2,93432 | 1,6086 | 0,0457385 | 0,0303344 | 0,0494767 | 0,0734351 | 0,0269996 | 0 | Ctnna2 | -4,44171037 | 2,34E-05 |
| Fgfr2 | 7,57334 | 12,7306 | 12,0679 | 16,1449 | 2,23746 | 0,229389 | 0,180828 | 0,0596816 | 0,0219141 | 0,0212004 | Fgfr2 | -3,17996844 | 2,40E-05 |
| Plac8 | 26,7162 | 23,2153 | 10,6145 | 12,1603 | 2,72783 | 0,161972 | 0,197131 | 0 | 0 | 0 | Plac8 | -3,57989357 | 2,49E-05 |
| Fez1 | 0 | 3,37397 | 6,42261 | 13,4901 | 0,0810221 | 0,0718355 | 0 | 0,774208 | 0,386355 | 0 | Fez1 | -4,43150518 | 2,57E-05 |
| Pnmt | 4,99264 | 8,9542 | 7,5748 | 5,4824 | 0,646887 | 0,45984 | 0,420443 | 2,8654 | 1,86595 | 5,36152 | Pnmt | -2,51818176 | 2,59E-05 |
| Chek2 | 6,18327 | 10,4667 | 5,37992 | 3,82025 | 1,31092 | 1,01601 | 0,851057 | 0,272299 | 0,453236 | 0,583382 | Chek2 | -2,07001669 | 2,59E-05 |
| Id4 | 7,59278 | 10,8189 | 12,1461 | 14,0202 | 2,99407 | 0,742575 | 0,259228 | 0,382588 | 0,426804 | 0,205688 | Id4 | -2,87703567 | 2,60E-05 |
| 4930412O13R | 0,398001 | 2,43429 | 3,85372 | 0,56139 | 0,0676549 | 0 | 0 | 0 | 0 | 0 | 4930412O13R | -4,39667729 | 2,62E-05 |
| Scn8a | 0,496678 | 0,136105 | 0,479397 | 0,305846 | 0,0468651 | 0,00689342 | 0,00843689 | 0 | 0 | 0 | Scn8a | -3,61961647 | 2,69E-05 |
| Plau | 22,7571 | 11,7189 | 3,82282 | 5,623 | 1,66424 | 0,718461 | 0,570947 | 0,13002 | 0,528702 | 0,324849 | Plau | -2,9554469 | 2,76E-05 |
| Fbln1 | 69,9612 | 51,7265 | 53,0219 | 69,8012 | 22,3647 | 5,26681 | 5,75907 | 14,8822 | 27,8019 | 19,9331 | Fbln1 | -2,10533141 | 2,77E-05 |
| Gprc5a | 11,3252 | 5,66701 | 1,91793 | 4,62933 | 0,838172 | 0,206105 | 0,0503866 | 0 | 0,22077 | 0,106522 | Gprc5a | -3,39914982 | 2,83E-05 |
| Alpl | 5,80328 | 4,03342 | 2,09167 | 3,94328 | 0,841847 | 0,198557 | 0,242777 | 11,0238 | 14,6493 | 17,1468 | Alpl | -2,7194696 | 2,85E-05 |
| Lat | 1,49665 | 2,3622 | 7,99393 | 8,13827 | 0,340434 | 0,150945 | 0 | 0 | 0 | 0 | Lat | -3,46834621 | 2,87E-05 |
| Dnah5 | 0,0987665 | 0,155695 | 0,0494675 | 0,298571 | 0 | 0 | 0 | 0 | 0 | 0 | Dnah5 | -5,13798065 | 2,88E-05 |
| **Hoxb1** | 2,61766 | 2,42088 | 0,40966 | 0,329599 | 0 | 0 | 0 | 0 | 0 | 0 | Hoxb1 | -5,18246254 | 2,90E-05 |
| Ajap1 | 1,49459 | 0,737199 | 1,51485 | 1,18307 | 0,121034 | 0 | 0 | 0 | 0,0357674 | 0 | Ajap1 | -4,00658361 | 2,93E-05 |
| Spag4 | 1,42116 | 2,33654 | 4,34908 | 2,38549 | 0,403666 | 0,23845 | 0,0728189 | 1,0735 | 0,320076 | 0,385376 | Spag4 | -2,78294627 | 2,96E-05 |
| Pnp | 24,6848 | 21,4864 | 15,5231 | 14,5159 | 5,17661 | 1,01765 | 1,21598 | 0,964649 | 2,54552 | 1,3203 | Pnp | -2,45287139 | 2,96E-05 |
| Trim10 | 0,600181 | 1,03612 | 1,87857 | 1,96491 | 0,0851168 | 0 | 0 | 0 | 0 | 0 | Trim10 | -4,33297238 | 2,97E-05 |
| Arnt2 | 2,1039 | 2,76734 | 5,94071 | 5,39125 | 0,838519 | 0,117034 | 0,175026 | 0,0945836 | 0,0520327 | 0 | Arnt2 | -2,87442714 | 2,99E-05 |
| Rcl1 | 24,3692 | 31,2326 | 8,64828 | 24,9322 | 5,55517 | 3,47251 | 3,2416 | 2,78583 | 2,77944 | 3,52177 | Rcl1 | -2,01865579 | 3,03E-05 |
| Dpf1 | 10,6302 | 4,80633 | 5,66867 | 3,12326 | 0,5025 | 0,0370614 | 0 | 0 | 0,0495946 | 0 | Dpf1 | -3,8684747 | 3,05E-05 |
| Celsr3 | 0,698046 | 0,158915 | 1,61362 | 1,70435 | 0,0531985 | 0,0738733 | 0,032877 | 0,122279 | 0,143194 | 0,112667 | Celsr3 | -3,00797822 | 3,10E-05 |
| Hist1h2ae | 39,135 | 32,7253 | 28,1543 | 30,2251 | 2,33202 | 0,311227 | 0,772232 | 1,36934 | 0,98075 | 0,562868 | Hist1h2ae | -3,39434214 | 3,18E-05 |
| Folr1 | 0,558536 | 3,30582 | 0,932282 | 9,063 | 0 | 0 | 0,0859295 | 0 | 0,0946027 | 0 | Folr1 | -4,81143705 | 3,20E-05 |
| Rgs10 | 19,9028 | 12,7838 | 22,4013 | 14,4427 | 3,69477 | 0,703591 | 0,428838 | 0 | 0,31733 | 0,45579 | Rgs10 | -2,90020871 | 3,24E-05 |
| Nr2e1 | 0,100879 | 0,199032 | 1,01044 | 4,26853 | 0 | 0 | 0 | 0 | 0 | 0 | Nr2e1 | -5,23237108 | 3,30E-05 |
| Igsf21 | 1,8774 | 1,85201 | 0,712262 | 1,0315 | 0,0484254 | 0 | 0 | 0 | 0,287076 | 0,055397 | Igsf21 | -4,59412808 | 3,34E-05 |
| Papolb | 1,14825 | 2,12387 | 0,599007 | 3,32551 | 0,122119 | 0,0720504 | 0 | 0 | 0,0482012 | 0,0232696 | Papolb | -3,57019544 | 3,37E-05 |
| Piezo1 | 49,811 | 54,5298 | 30,2919 | 31,0018 | 14,0446 | 3,75004 | 3,32615 | 1,52909 | 2,88931 | 2,21672 | Piezo1 | -2,17182186 | 3,43E-05 |
| Xylb | 1,41971 | 0,823847 | 2,85806 | 2,41887 | 0,354922 | 0,188318 | 0,153574 | 0,0759706 | 0,0279386 | 0,0270164 | Xylb | -2,63020437 | 3,46E-05 |
| Sowahb | 0,826051 | 0,570428 | 0,413709 | 1,33161 | 0 | 0 | 0,0506342 | 0 | 0,0276335 | 0 | Sowahb | -4,29999541 | 3,52E-05 |
| Rab39b | 0,573147 | 0,565407 | 0,318938 | 1,05863 | 0 | 0 | 0 | 0 | 0,0319816 | 0 | Rab39b | -5,06830351 | 3,58E-05 |
| Klhl4 | 14,9959 | 18,6931 | 3,46262 | 0,944987 | 0,987268 | 0,192288 | 0,287481 | 0,310259 | 0,627642 | 0,248277 | Klhl4 | -3,55427808 | 3,64E-05 |
| Pdpn | 11,3426 | 16,4726 | 25,2122 | 24,9457 | 5,1558 | 0,821461 | 1,47626 | 0,348885 | 0,841909 | 0,312256 | Pdpn | -2,51170759 | 3,71E-05 |
| St14 | 0,640489 | 2,68533 | 2,1385 | 4,27513 | 0,294871 | 0,280806 | 0,245362 | 0,072837 | 0,133891 | 0,0258966 | St14 | -2,69217124 | 3,76E-05 |

| Gene | d9-11 (1) | d9-11 (2) | d9-11(3) | d9-11(4) | newborn(1) | newborn(2) | newborn(3) | adult(1) | adult(2) | adult(3) | Gene | log2(fold) | FDR(*p* -value) |
|---|---|---|---|---|---|---|---|---|---|---|---|---|---|
| Dmrta2 | 1,44885 | 1,64917 | 1,11632 | 5,16484 | 0,22114 | 0,139735 | 0,205064 | 0,101319 | 0 | 0,0360928 | Dmrta2 | -3,04963192 | 3,85E-05 |
| Bai3 | 0,172034 | 0,226282 | 0,622274 | 0,404452 | 0,0162405 | 0 | 0 | 0 | 0 | 0 | Bai3 | -4,57941957 | 3,88E-05 |
| Alx3 | 0,173298 | 1,02573 | 1,44643 | 3,49121 | 0 | 0 | 0 | 0 | 0 | 0 | Alx3 | -5,11548325 | 3,88E-05 |
| Th | 2,76666 | 0,584835 | 3,62866 | 0,398092 | 0 | 0 | 0,0607066 | 0 | 0 | 0 | Th | -4,68284187 | 3,88E-05 |
| Sost | 10,3684 | 10,0659 | 2,19785 | 0,497341 | 0,420223 | 0,0413331 | 0,0505233 | 0 | 0 | 0,0534062 | Sost | -4,10053572 | 3,92E-05 |
| Trim36 | 2,74139 | 0,901357 | 2,82088 | 1,07106 | 0,278761 | 0,140841 | 0,258426 | 0,19187 | 0,599151 | 0,63639 | Trim36 | -2,71604982 | 4,05E-05 |
| Fam181b | 2,43309 | 2,03092 | 4,21772 | 11,1225 | 0,584297 | 0,32925 | 0,229928 | 1,52863 | 1,95387 | 1,64125 | Fam181b | -3,0982143 | 4,05E-05 |
| Sox8 | 2,29669 | 1,61834 | 1,4606 | 2,75451 | 0,247998 | 0,137114 | 0 | 0 | 0 | 0 | Sox8 | -3,23516227 | 4,11E-05 |
| Phyhipl | 0,687595 | 1,1407 | 0,908833 | 1,62259 | 0 | 0 | 0 | 0 | 0 | 0 | Phyhipl | -4,16801012 | 4,29E-05 |
| Car4 | 80,3873 | 57,6715 | 7,50649 | 13,5862 | 5,67247 | 0,991642 | 1,21084 | 3,81773 | 4,76128 | 5,49707 | Car4 | -3,31833599 | 4,34E-05 |
| Jph3 | 0,825378 | 0,814234 | 0,482267 | 0,582104 | 0,04677 | 0,0206795 | 0,0252964 | 0,150172 | 0,0276109 | 0,0267002 | Jph3 | -3,6850224 | 4,44E-05 |
| Hcn1 | 1,0675 | 1,1311 | 0,891087 | 0,478058 | 0,135571 | 0,148402 | 0,159409 | 0,288029 | 0,595438 | 0,84275 | Hcn1 | -2,16116816 | 4,48E-05 |
| Cdx2 | 0,789156 | 1,08988 | 0,395206 | 2,06683 | 0 | 0 | 0 | 0 | 0 | 0 | Cdx2 | -5,03804288 | 4,53E-05 |
| Kit | 10,9439 | 10,0077 | 6,82573 | 5,14165 | 2,01998 | 0,215527 | 0,301367 | 0,167609 | 0,24648 | 0,258351 | Kit = c-Kit | -2,81720624 | 4,59E-05 |
| Srcin1 | 0,495884 | 0,400249 | 0,413934 | 0,772193 | 0,0510584 | 0,0225648 | 0,0552241 | 0,0820925 | 0,0526565 | 0,0145623 | Srcin1 | -3,06208059 | 4,59E-05 |
| Oprl1 | 0,673638 | 4,20874 | 1,4606 | 1,5239 | 0,190951 | 0 | 0,103291 | 0 | 0 | 0 | Oprl1 | -3,56927962 | 4,63E-05 |
| Plxnd1 | 90,7834 | 96,9823 | 36,5965 | 29,277 | 17,2272 | 4,43834 | 4,2056 | 9,56381 | 12,7171 | 11,6262 | Plxnd1 | -2,44284979 | 4,70E-05 |
| Dock9 | 14,6003 | 8,07976 | 6,32729 | 4,71374 | 1,93983 | 0,435369 | 0,456934 | 0,82819 | 2,55888 | 1,73411 | Dock9 | -2,62978749 | 4,73E-05 |
| Sh3bgrl3 | 162,291 | 104,268 | 92,8657 | 73,827 | 33,2352 | 6,48898 | 6,9353 | 3,35174 | 6,46088 | 3,28559 | Sh3bgrl3 | -2,44575047 | 4,90E-05 |
| Irs4 | 0,201762 | 0,298559 | 0,210522 | 1,21976 | 0,0142836 | 0 | 0 | 0 | 0 | 0 | Irs4 | -4,65586412 | 4,95E-05 |
| Zcchc16 | 0,801334 | 0,263505 | 0,66888 | 0,448522 | 0 | 0 | 0 | 0 | 0 | 0 | Zcchc16 | -4,99745086 | 4,95E-05 |
| Tspan33 | 2,0963 | 7,0655 | 7,43612 | 3,34331 | 0,644396 | 0,702178 | 0,804586 | 1,26862 | 1,52834 | 1,1342 | Tspan33 | -2,30560715 | 4,96E-05 |
| Slc6a7 | 1,26526 | 0,28804 | 0,893638 | 0,19611 | 0 | 0 | 0 | 0 | 0,0325891 | 0 | Slc6a7 | -5,04998534 | 5,01E-05 |
| Col23a1 | 9,10178 | 8,04249 | 6,15261 | 4,66976 | 1,24929 | 0,111835 | 0,085523 | 0,254147 | 0,428902 | 0,216541 | Col23a1 | -3,15053002 | 5,02E-05 |
| Adam12 | 12,9724 | 8,00842 | 6,46992 | 2,95951 | 0,947101 | 0,0510415 | 0,0749528 | 0,0371469 | 0,0136104 | 0 | Adam12 | -3,64097065 | 5,04E-05 |
| 5930412G12R | 0,834413 | 3,15535 | 0,116077 | 0,420281 | 0 | 0 | 0 | 0 | 0 | 0 | 5930412G12R | -5,06036527 | 5,16E-05 |
| En2 | 0,549835 | 0,452008 | 0,994394 | 2,86208 | 0,051935 | 0 | 0,056182 | 0 | 0 | 0 | En2 | -4,03659664 | 5,29E-05 |
| Tmem67 | 3,19565 | 5,36413 | 4,23649 | 2,74609 | 1,19855 | 0,400462 | 0,403379 | 0 | 0,471969 | 0,577942 | Tmem67 | -2,10061754 | 5,30E-05 |
| Sgol1 | 4,48468 | 7,21131 | 3,59366 | 3,07243 | 1,55034 | 0,854128 | 0,742324 | 0 | 0,0600365 | 0,0290229 | Sgol1 | -2,01163799 | 5,37E-05 |
| Fignl2 | 29,1956 | 18,1983 | 10,0535 | 15,9749 | 1,16615 | 0,0763813 | 0,0233599 | 0,138713 | 0,178431 | 0,0246534 | Fignl2 | -4,29743381 | 5,41E-05 |
| L1cam | 0,0599912 | 0,82854 | 6,81038 | 2,84092 | 0,135926 | 0,0150207 | 0 | 0,163811 | 0,120239 | 0,0387798 | L1cam | -4,2982478 | 5,44E-05 |
| Klf8 | 2,08079 | 1,41565 | 1,5572 | 1,08438 | 0,345531 | 0,215656 | 0,219858 | 0,0652896 | 0,0479722 | 0,0928052 | Klf8 | -2,18360935 | 5,49E-05 |
| Wscd2 | 0,595795 | 2,93876 | 4,78671 | 5,5526 | 0,443051 | 0,130584 | 0,0228214 | 0 | 0,0249007 | 0 | Wscd2 | -3,43367114 | 5,56E-05 |
| Hhip | 1,20153 | 1,01599 | 0,401189 | 0,622645 | 0,07775 | 0 | 0 | 0 | 0 | 0 | Hhip | -3,91900542 | 5,66E-05 |
| Mmp25 | 1,25323 | 1,42651 | 0,885142 | 0,0971231 | 0,0546388 | 0 | 0 | 0 | 0 | 0 | Mmp25 | -4,34536945 | 5,67E-05 |
| Nup210l | 0,89081 | 1,26326 | 1,90544 | 1,00975 | 0,173441 | 0,0418152 | 0,0511633 | 0 | 0 | 0,0899606 | Nup210l | -2,6533228 | 5,74E-05 |
| Gata2 | 40,5197 | 28,1692 | 16,7011 | 10,9399 | 3,98934 | 0,302453 | 0,184959 | 1,18858 | 1,5153 | 1,39955 | Gata2 | -3,40644426 | 5,81E-05 |
| Slc35d3 | 2,90143 | 0,622223 | 1,47412 | 1,14374 | 0,1073 | 0 | 0,0386964 | 0 | 0 | 0 | Slc35d3 | -3,73818928 | 5,99E-05 |
| Zfp69 | 8,29095 | 4,1943 | 3,63755 | 3,99753 | 1,30628 | 0,604692 | 0,804529 | 0,128752 | 0,285536 | 0,091901 | Zfp69 | -2,12153665 | 6,00E-05 |
| Pax1 | 0 | 0,991854 | 1,88825 | 2,53221 | 0 | 0 | 0 | 0 | 0 | 0 | Pax1 | -5,06897539 | 6,07E-05 |
| Dmbx1 | 0,480131 | 0,236655 | 0,20023 | 1,60691 | 0 | 0 | 0 | 0 | 0 | 0 | Dmbx1 | -5,01692869 | 6,09E-05 |
| Efna2 | 9,69816 | 10,8217 | 20,5684 | 16,1751 | 0,99222 | 2,91432 | 3,8065 | 1,15499 | 1,65676 | 0,773711 | Efna2 | -2,08015694 | 6,24E-05 |
| Pla2g12b | 7,17243 | 1,41503 | 1,49639 | 2,16641 | 0,102066 | 0 | 0 | 0 | 0 | 0 | Pla2g12b | -4,60654342 | 6,28E-05 |
| Fam19a5 | 3,95325 | 5,48159 | 2,98038 | 6,93725 | 1,12088 | 0,255941 | 0,391192 | 4,51966 | 5,86113 | 5,28943 | Fam19a5 | -2,55478553 | 6,36E-05 |
| Grm4 | 0,564889 | 0,238826 | 0,40416 | 1,537 | 0,0288681 | 0 | 0 | 0 | 0,0539848 | 0,0989234 | Grm4 | -4,54329852 | 6,41E-05 |
| Prdm12 | 0,26999 | 0,665854 | 1,91551 | 0,883944 | 0 | 0 | 0 | 0 | 0 | 0 | Prdm12 | -4,96766219 | 6,51E-05 |
| Ptprz1 | 0,581194 | 0,191118 | 1,00264 | 2,10824 | 0,0767897 | 0,0290861 | 0 | 0 | 0,0387769 | 0,0125132 | Ptprz1 | -3,79010556 | 6,60E-05 |
| Pls1 | 0,699399 | 1,98362 | 1,02164 | 3,61129 | 0 | 0,131443 | 0,133984 | 0 | 0,0292548 | 0 | Pls1 | -3,49158527 | 6,62E-05 |
| Myo7a | 18,2693 | 12,9071 | 10,8232 | 6,23619 | 3,21838 | 0,625935 | 0,76087 | 0,912844 | 0,644278 | 0,76029 | Myo7a | -2,4897658 | 6,67E-05 |
| Cd248 | 10,8994 | 16,8962 | 30,7582 | 20,2606 | 5,00358 | 0,781303 | 0,597079 | 0,471638 | 0,60951 | 0,2523 | Cd248 | -2,73694239 | 6,67E-05 |
| Rnf208 | 6,54616 | 3,22875 | 12,7477 | 13,4591 | 1,24031 | 1,44309 | 1,34264 | 0,494063 | 0,277075 | 0,799904 | Rnf208 | -2,29732016 | 6,68E-05 |
| Mgat3 | 1,29951 | 1,41691 | 4,33892 | 2,48081 | 0,464967 | 0,222679 | 0,335297 | 0,871371 | 2,05734 | 0,818193 | Mgat3 | -2,36938862 | 6,68E-05 |
| Plg | 0,239769 | 1,30091 | 0,700461 | 1,57005 | 0 | 0,0300692 | 0 | 0 | 0 | 0 | Plg | -4,52151462 | 6,72E-05 |
| Ptges | 3,50364 | 3,41257 | 1,77692 | 3,66394 | 0,552933 | 0,0444547 | 0,135941 | 0,0806797 | 0 | 0,0287002 | Ptges | -3,01254069 | 6,79E-05 |
| Slc43a1 | 22,4052 | 23,1984 | 15,5492 | 15,8503 | 3,33986 | 0,26673 | 0,462589 | 0 | 0,184041 | 0,044431 | Slc43a1 | -2,93826896 | 6,86E-05 |
| Usp44 | 2,06319 | 0,648023 | 1,17018 | 0,132351 | 0,0372529 | 0 | 0 | 0 | 0 | 0 | Usp44 | -4,58269002 | 6,94E-05 |
| Tcerg1l | 1,16432 | 0,893342 | 1,29577 | 0,130325 | 0 | 0 | 0 | 0 | 0 | 0 | Tcerg1l | -4,95995911 | 7,09E-05 |
| Pvalb | 1,80473 | 6,67581 | 3,01195 | 0,908317 | 0 | 0 | 0 | 0 | 0,154546 | 0,296064 | Pvalb | -4,95100543 | 7,11E-05 |
| B3galnt1 | 7,71859 | 9,95711 | 6,44245 | 4,78479 | 1,93662 | 0,409873 | 0,318869 | 0,134825 | 0,0498633 | 0,0962759 | B3galnt1 | -2,58844005 | 7,29E-05 |
| Tram1l1 | 0,909696 | 4,48699 | 3,29026 | 3,66549 | 0,473055 | 0,0761267 | 0,0465321 | 0 | 0,0509424 | 0 | Tram1l1 | -3,31573674 | 7,41E-05 |
| Grrp1 | 20,516 | 13,6904 | 3,0215 | 7,29138 | 1,71574 | 0,532537 | 0,557253 | 0 | 1,33056 | 1,47631 | Grrp1 | -2,91146324 | 7,42E-05 |
| Lcp1 | 11,1785 | 13,6137 | 9,34324 | 5,08522 | 3,29798 | 1,0579 | 0,994741 | 0,438983 | 0,374788 | 0,473899 | Lcp1 | -2,13893747 | 7,44E-05 |
| Tlx3 | 0,231454 | 0,456641 | 3,86347 | 3,49659 | 0 | 0 | 0 | 0,209826 | 0 | 0 | Tlx3 | -5,00271618 | 7,46E-05 |
| Bai2 | 2,00107 | 5,207 | 14,7642 | 13,5872 | 1,47956 | 1,22219 | 0,865555 | 1,06215 | 1,30356 | 1,55766 | Bai2 | -2,40701446 | 7,51E-05 |
| Gngt2 | 29,4324 | 74,0746 | 0 | 0 | 0 | 0 | 0 | 0 | 0 | 0 | Gngt2 | -3,37587185 | 7,58E-05 |
| Lmx1b | 0,607631 | 0,299698 | 12,6776 | 4,28302 | 0,08639 | 0 | 0 | 0 | 0 | 0 | Lmx1b | -4,82218563 | 7,82E-05 |
| Col13a1 | 12,7012 | 20,3606 | 6,24734 | 7,1975 | 2,28237 | 0,284383 | 0,173888 | 0,20618 | 0,494137 | 0,036728 | Col13a1 | -2,93990788 | 7,96E-05 |
| March11 | 0,457683 | 2,70893 | 1,90994 | 0,460954 | 0 | 0 | 0 | 0,622411 | 0,0772702 | 0,372214 | 11. Mrz | -4,92269681 | 8,11E-05 |
| Six3 | 0,263563 | 0,173336 | 0,439996 | 1,85877 | 0,0248934 | 0 | 0 | 0 | 0 | 0 | Six3 | -4,9542722 | 8,23E-05 |
| Dmtn | 1,03357 | 4,98309 | 4,28436 | 3,24195 | 0,726214 | 0,226796 | 0,265419 | 0,519411 | 0,654802 | 0,527684 | Dmtn | -2,55891045 | 8,24E-05 |
| Col25a1 | 0,139628 | 0,550977 | 0,955859 | 0,686772 | 0,052716 | 0,0116489 | 0,0142543 | 0 | 0,0155342 | 0,0150355 | Col25a1 | -3,59528045 | 8,26E-05 |
| Traip | 4,78487 | 5,8095 | 3,0722 | 3,46075 | 1,35656 | 0,677055 | 0,48922 | 0,111507 | 0,102868 | 0,238497 | Traip | -2,21665489 | 8,50E-05 |
| Gem | 35,9481 | 36,7865 | 28,5107 | 19,834 | 9,77299 | 1,83479 | 2,10573 | 0,406452 | 0,501102 | 0,387003 | Gem | -2,28114451 | 8,52E-05 |
| 2810459M11R | 0,0977503 | 0,771441 | 1,0607 | 1,77324 | 0,0554306 | 0 | 0 | 0 | 0 | 0 | 2810459M11R | -4,29409246 | 8,63E-05 |
| Grid1 | 0,379003 | 1,58901 | 2,53084 | 0,572757 | 0,0805518 | 0 | 0 | 0 | 0 | 0 | Grid1 | -4,18759386 | 8,65E-05 |
| Rgs5 | 19,6895 | 31,6805 | 21,0605 | 9,24212 | 5,31713 | 1,08104 | 0,939441 | 8,67302 | 23,3097 | 14,7787 | Rgs5 | -2,56763216 | 8,75E-05 |
| Gpr97 | 5,75407 | 5,94661 | 2,05833 | 0,276024 | 0,271953 | 0,0687596 | 0 | 0 | 0,137968 | 0,0888218 | Gpr97 | -3,89429446 | 8,76E-05 |
| Rgs16 | 18,3894 | 7,22798 | 6,35547 | 7,52565 | 2,15948 | 1,18996 | 1,01397 | 0,522037 | 0,385977 | 0,885084 | Rgs16 | -2,14418743 | 8,85E-05 |
| Dlg4 | 23,6528 | 27,4966 | 23,2657 | 24,6205 | 5,70183 | 0,688825 | 0,571691 | 0 | 0,197042 | 0,127071 | Dlg4 | -2,76150662 | 8,97E-05 |
| Igsf11 | 7,14191 | 3,47576 | 1,27176 | 1,91875 | 0,485736 | 0,0954687 | 0,233539 | 0,259807 | 0,191285 | 0,184917 | Igsf11 | -3,07710735 | 9,03E-05 |
| Aadat | 4,86772 | 2,88113 | 1,62511 | 0,196117 | 0,110534 | 0,0489434 | 0 | 0 | 0,0656017 | 0 | Aadat | -4,2202872 | 9,11E-05 |
| Adcy8 | 0,450028 | 0,443953 | 0,160993 | 0,388652 | 0 | 0 | 0 | 0,760633 | 0,665928 | 0,436436 | Adcy8 | -4,86965567 | 9,17E-05 |
| Klhl34 | 1,76829 | 0,373799 | 0,94883 | 1,27241 | 0,0716225 | 0,0950578 | 0 | 3,44355 | 2,24562 | 3,60159 | Klhl34 | -3,67405462 | 9,34E-05 |
| Krt5 | 0,30846 | 0,152145 | 2,44587 | 7,63856 | 0 | 0 | 0 | 0 | 0 | 0 | Krt5 | -5,03721657 | 9,56E-05 |
| Tmem59l | 0,958641 | 1,65491 | 0,600064 | 2,1723 | 0 | 0 | 0 | 0 | 0 | 0 | Tmem59l | -4,86574448 | 9,77E-05 |
| Nxf7 | 3,8826 | 0,547155 | 1,23451 | 0,744908 | 0 | 0 | 0,0567807 | 0 | 0 | 0 | Nxf7 | -4,22624167 | 0,000100231 |
| Lrp11 | 16,3604 | 8,43151 | 3,993 | 2,12058 | 0,460918 | 0,911244 | 0,997233 | 1,91422 | 2,05006 | 2,50819 | Lrp11 | -2,7785111 | 0,000101478 |
| Serpinf2 | 0 | 0,303982 | 2,31479 | 8,22577 | 0 | 0 | 0 | 0 | 0 | 0 | Serpinf2 | -5,02724331 | 0,000101643 |
| Rhou | 21,9251 | 29,6326 | 20,3164 | 19,4616 | 7,52717 | 1,13783 | 1,27324 | 2,54727 | 1,55228 | 2,34461 | Rhou | -2,40674569 | 0,000102238 |
| Dyrk3 | 4,26518 | 2,75106 | 3,01227 | 3,13963 | 0,698099 | 0,288391 | 0,0503593 | 0,297901 | 0,0551623 | 0,212929 | Dyrk3 | -2,79157579 | 0,000102365 |
| Stc1 | 2,67481 | 4,68154 | 2,66482 | 3,12952 | 1,00236 | 0,237825 | 0,185126 | 0,392435 | 0,230967 | 0,223328 | Stc1 | -2,41829055 | 0,000106763 |
| Smpdl3b | 0,890855 | 2,63642 | 4,61 | 4,8456 | 0,404472 | 0,223823 | 0,382978 | 0,161723 | 0 | 0 | Smpdl3b | -2,74981569 | 0,000109481 |
| Hmha1 | 16,1786 | 15,2689 | 12,8136 | 9,76957 | 2,69317 | 0,308098 | 0,0857482 | 0,0848009 | 0,0624247 | 0,0905241 | Hmha1 | -3,07868208 | 0,00011046 |
| **Hoxc5** | 0,242155 | 0,238883 | 1,3138 | 1,4637 | 0 | 0 | 0 | 0 | 0 | 0 | Hoxc5 | -4,87513428 | 0,000110553 |
| Dpysl4 | 1,38191 | 4,21364 | 16,6727 | 13,162 | 1,24665 | 0,315152 | 0,385361 | 4,11014 | 5,01475 | 5,65818 | Dpysl4 | -3,0564296 | 0,000110629 |
| Maff | 45,7549 | 47,4118 | 26,9568 | 34,855 | 12,8503 | 2,60935 | 3,36587 | 6,27994 | 4,53335 | 5,37081 | Maff | -2,15120252 | 0,000111087 |
| Sla2 | 0 | 0 | 0 | 0 | 0 | 0 | 0 | 0 | 0 | 0 | Sla2 | -4,04822209 | 0,000111242 |
| F2rl2 | 0 | 0,534343 | 2,3736 | 7,09358 | 0,0383998 | 0 | 0 | 0 | 0 | 0 | F2rl2 | -4,79074814 | 0,000112397 |
| Lrr1 | 3,31863 | 7,48276 | 5,5395 | 3,10355 | 0,471312 | 0,835228 | 0,510131 | 0 | 0 | 0 | Lrr1 | -2,26146977 | 0,000112533 |
| Fga | 0 | 0,991445 | 0,838877 | 3,54078 | 0 | 0 | 0 | 0 | 0 | 0 | Fga | -4,93991986 | 0,000119011 |
| Aff3 | 8,24457 | 11,4488 | 15,2325 | 13,1749 | 3,23034 | 0,593425 | 0,27344 | 0,0478056 | 0,245469 | 0,124433 | Aff3 | -2,63270894 | 0,000119542 |
| Gck | 2,5441 | 0,557422 | 0,942214 | 1,84843 | 0,200311 | 0,141816 | 0,216731 | 16,2985 | 7,20985 | 20,2666 | Gck | -2,78778993 | 0,000122621 |
| Neurog1 | 0,210182 | 0,414676 | 4,03473 | 2,32863 | 0 | 0 | 0 | 0 | 0 | 0 | Neurog1 | -4,88224796 | 0,000123778 |
| Cklf | 14,6299 | 22,7351 | 10,1266 | 1,66066 | 1,72789 | 1,95593 | 1,09505 | 0 | 0,470705 | 0,29575 | Cklf | -2,08904738 | 0,00012703 |
| Dact1 | 47,696 | 30,1898 | 23,9835 | 24,5821 | 9,21575 | 1,0325 | 1,20612 | 1,24942 | 0,861704 | 0,831576 | Dact1 | -2,66105595 | 0,00012703 |
| Tagln2 | 180,511 | 327,147 | 156,857 | 103,688 | 59,6289 | 11,6035 | 10,4206 | 4,97064 | 11,2275 | 7,65898 | Tagln2 | -2,40298957 | 0,000128743 |
| Cadps | 0,463985 | 0,514969 | 2,60645 | 0,409034 | 0,0821008 | 0,0435067 | 0,053232 | 0 | 0 | 0 | Cadps | -3,29217666 | 0,000128803 |
| Stab1 | 60,0345 | 34,5676 | 16,9781 | 6,45467 | 5,52889 | 0 | 0 | 0 | 1,1873 | 0,871526 | Stab1 | -2,9690661 | 0,000131053 |
| Tdrp | 1,76119 | 2,13833 | 1,5831 | 2,45656 | 0,345752 | 0,271964 | 0,332522 | 0,861646 | 0,545673 | 0,966104 | Tdrp | -2,19867528 | 0,000131493 |
| Cacna2d3 | 3,93638 | 1,63959 | 0,730163 | 0,088131 | 0,0247859 | 0 | 0,0536246 | 0 | 0,263445 | 0,0849081 | Cacna2d3 | -4,02951493 | 0,000132357 |
| Btbd17 | 0,297886 | 0,29386 | 2,23772 | 2,70067 | 0,042245 | 0 | 0 | 0 | 0 | 0 | Btbd17 | -4,51198103 | 0,000135196 |
| Mybphl | 7,47369 | 23,2515 | 56,1359 | 37,0511 | 1,716 | 6,5931 | 3,71538 | 5,20499 | 4,9687 | 0,74975 | Mybphl | -2,37981407 | 0,000137169 |
| Trpc5 | 0,385981 | 0,456923 | 0,579932 | 0,933322 | 0,0218697 | 0,0193382 | 0 | 0 | 0 | 0 | Trpc5 | -3,75479839 | 0,000137936 |

| Gene | d9-11 (1) | d9-11 (2) | d9-11(3) | d9-11(4) | newborn(1) | newborn(2) | newborn(3) | adult(1) | adult(2) | adult(3) | Gene | log2(fold) | FDR(*p*-value) |
|---|---|---|---|---|---|---|---|---|---|---|---|---|---|
| Tle3 | 45,6007 | 38,9402 | 36,5929 | 44,886 | 10,7595 | 0,922382 | 0,768114 | 0,842279 | 1,18606 | 0,633527 | Tle3 | -2,8435013 | 0,000138517 |
| Tnni2 | 13,5426 | 30,7239 | 46,3257 | 67,4432 | 4,64897 | 4,48806 | 7,5604 | 1,21513 | 3,29494 | 3,13702 | Tnni2 | -2,36206869 | 0,000138948 |
| **Hoxd1** | 1,66498 | 0,912479 | 0,463221 | 0,931701 | 0 | 0 | 0 | 0 | 0 | 0 | Hoxd1 | -4,78493116 | 0,000140151 |
| Zfp185 | 2,36942 | 2,15762 | 3,42306 | 1,28539 | 0,542535 | 0,0913533 | 0,11174 | 0,0828868 | 0,152511 | 0,0294898 | Zfp185 | -2,66686669 | 0,000140757 |
| Emilin3 | 2,14552 | 1,97633 | 4,61939 | 2,20188 | 0,646634 | 0,0476596 | 0,167484 | 0 | 0 | 0,0615421 | Emilin3 | -2,86004864 | 0,000143499 |
| Zfp239 | 0,25188 | 1,11815 | 1,26144 | 1,29882 | 0,0714145 | 0,0315937 | 0 | 0 | 0 | 0 | Zfp239 | -3,87296963 | 0,000144945 |
| Adm2 | 3,04114 | 1,33329 | 0,845981 | 2,04137 | 0 | 0 | 0 | 0 | 0 | 0,110383 | Adm2 | -4,7753402 | 0,00014814 |
| **Hoxd4** | 0,259842 | 0,128165 | 2,27729 | 3,01027 | 0,00920957 | 0 | 0,0398564 | 0,11806 | 0,0435927 | 0 | Hoxd4 | -4,33101989 | 0,000150002 |
| Spp2 | 0 | 5,06073 | 2,72433 | 10,3282 | 0 | 0 | 0 | 0 | 0 | 0 | Spp2 | -4,86311725 | 0,000152709 |
| Kdf1 | 1,0011 | 1,18507 | 1,50399 | 1,20999 | 0,0568364 | 0,100674 | 0 | 0 | 0 | 0 | Kdf1 | -3,72103903 | 0,000157748 |
| Calcrl | 22,682 | 28,927 | 9,10141 | 4,23835 | 3,35537 | 0,438248 | 0,494922 | 0,490006 | 1,59705 | 1,20769 | Calcrl | -2,98720737 | 0,00016375 |
| Plcxd3 | 1,6371 | 0,17944 | 0,607288 | 2,19866 | 0 | 0 | 0 | 0 | 0 | 0 | Plcxd3 | -4,78633342 | 0,000164278 |
| Cd38 | 11,7154 | 17,8248 | 8,03115 | 3,08949 | 1,94028 | 0,439398 | 0,167927 | 0,697008 | 1,89011 | 1,06399 | Cd38 | -2,91554818 | 0,000166633 |
| Ecscr | 23,8387 | 47,0309 | 18,7574 | 6,51495 | 4,45715 | 1,37552 | 0,734411 | 0,307947 | 2,77966 | 1,44637 | Ecscr | -2,79871202 | 0,000167786 |
| Nr2f2 | 6,69471 | 13,302 | 26,1183 | 20,2353 | 4,93541 | 0,876623 | 0,722705 | 0,899836 | 1,39914 | 1,01052 | Nr2f2 | -2,48602992 | 0,000171944 |
| Ido2 | 1,19658 | 0,214621 | 0,453989 | 0,547947 | 0 | 0 | 0 | 0 | 0 | 0 | Ido2 | -4,75204902 | 0,000174407 |
| Arx | 0 | 0,238978 | 3,03305 | 2,44047 | 0 | 0 | 0 | 0 | 0 | 0 | Arx | -4,86711897 | 0,000175166 |
| C130021I20Ri | 0 | 0,250246 | 1,48219 | 1,70383 | 0 | 0 | 0 | 0 | 0 | 0 | C130021I20Ri | -4,84076262 | 0,000181951 |
| Chrng | 1,78589 | 0,176174 | 3,42836 | 1,25922 | 0 | 0,0448704 | 0,0548401 | 0 | 0 | 0 | Chrng | -4,16088465 | 0,000184679 |
| Ccdc18 | 1,32469 | 1,28428 | 1,14388 | 0,759391 | 0,252312 | 0,154441 | 0,209939 | 0 | 0,0458014 | 0,0443066 | Ccdc18 | -2,00591064 | 0,000185081 |
| Chst12 | 9,6982 | 12,8198 | 13,2749 | 15,2386 | 4,12911 | 0,682571 | 0,714955 | 0,351986 | 0,457433 | 1,07122 | Chst12 | -2,40315021 | 0,000185081 |
| Slc32a1 | 2,46891 | 0,927832 | 0,392528 | 0,23688 | 0 | 0 | 0,0360577 | 0 | 0 | 0 | Slc32a1 | -4,409887 | 0,000191094 |
| Ifitm10 | 0 | 0,597692 | 2,78148 | 2,33985 | 0,143092 | 0,0506271 | 0,0928795 | 0,0918226 | 0,135278 | 0,228821 | Ifitm10 | -3,39473646 | 0,000194145 |
| Clvs1 | 0 | 0 | 0 | 0 | 0 | 0 | 0 | 0 | 0 | 0 | Clvs1 | -3,63454547 | 0,000194145 |
| Ldlrad3 | 14,7463 | 14,8012 | 15,2094 | 12,118 | 6,02188 | 1,36244 | 1,28819 | 0,374871 | 0,60653 | 0,533423 | Ldlrad3 | -2,02329384 | 0,000197993 |
| Sema3c | 1,15575 | 19,0025 | 21,0633 | 13,2621 | 2,34604 | 1,01295 | 0,747518 | 3,33087 | 4,01194 | 3,98521 | Sema3c | -2,72616541 | 0,00019857 |
| Clic6 | 11,6483 | 5,10714 | 2,80886 | 3,99884 | 0,513406 | 0,972922 | 1,16365 | 0,156974 | 0,0288709 | 0,0279161 | Clic6 | -2,34429106 | 0,000199553 |
| Padi3 | 9,49797 | 2,52665 | 0,356311 | 0 | 0,030248 | 0 | 0,0327233 | 0 | 0 | 0 | Padi3 | -4,53947826 | 0,000199671 |
| Mfsd2a | 0,314846 | 0,310589 | 4,59881 | 5,39157 | 0,133969 | 0,0395279 | 0 | 0 | 0 | 0,0510705 | Mfsd2a | -4,04679756 | 0,00020028 |
| Stmn3 | 1,6564 | 0,653577 | 4,9764 | 13,009 | 0,188484 | 0 | 0 | 0 | 0 | 0 | Stmn3 | -4,22839361 | 0,000202471 |
| Car12 | 0,956451 | 0,771983 | 0,435468 | 0,350408 | 0,0492737 | 0 | 0 | 0 | 0,05819 | 0,0843967 | Car12 | -3,99539073 | 0,000208475 |
| Glipr2 | 31,4103 | 50,8096 | 50,8829 | 35,2116 | 10,1097 | 0,933173 | 0,622154 | 0,306632 | 0,056803 | 0,0548088 | Glipr2 | -2,8641583 | 0,000209549 |
| Uncx | 0 | 0,64006 | 6,49863 | 2,61431 | 0 | 0 | 0 | 0 | 0 | 0 | Uncx | -4,60760785 | 0,000209608 |
| Emx2 | 0,383376 | 0,504261 | 1,27999 | 3,0897 | 0 | 0,0641201 | 0 | 0 | 0 | 0 | Emx2 | -4,21197306 | 0,000210302 |
| Xylt1 | 2,23952 | 5,68098 | 3,11553 | 6,55374 | 1,08816 | 0,187166 | 0,327003 | 0 | 0,0357296 | 0,10359 | Xylt1 | -2,52963435 | 0,000210989 |
| Pou3f3os | 0 | 3,65271 | 12,3577 | 13,9656 | 0 | 0 | 0 | 0 | 0 | 0 | Pou3f3os | -4,78067429 | 0,00021117 |
| Nova2 | 4,2151 | 2,24471 | 3,36284 | 2,36792 | 0,902942 | 0,168003 | 0,102798 | 0,305729 | 0,497705 | 0,216828 | Nova2 | -2,48807216 | 0,000213783 |
| D430020J02Ri | 1,70614 | 1,79529 | 3,89253 | 1,94799 | 0,25794 | 0,484902 | 0,453465 | 0 | 0 | 0 | D430020J02Ri | -2,13868092 | 0,000217606 |
| Tac2 | 1,31337 | 3,23871 | 2,19158 | 5,2856 | 0 | 0 | 0 | 0 | 0 | 0 | Tac2 | -4,67836304 | 0,000219765 |
| Tert | 1,23253 | 1,12235 | 3,95691 | 2,19694 | 0,214939 | 0,285153 | 0,0871943 | 0 | 0 | 0,0306844 | Tert | -2,54649356 | 0,000221026 |
| 2310014L17Ri | 2,21492 | 1,5891 | 0,840362 | 1,11573 | 0,199732 | 0,0252379 | 0 | 0 | 0 | 0 | 2310014L17Ri | -3,46803978 | 0,000221026 |
| Peg12 | 4,64632 | 6,56561 | 6,39379 | 8,7289 | 1,45976 | 0,126008 | 0,0770403 | 0 | 0 | 0,122068 | Peg12 | -3,01891731 | 0,000222081 |
| Batf3 | 3,6931 | 4,24995 | 2,05423 | 1,23867 | 0,527741 | 0 | 0 | 0 | 0 | 0 | Batf3 | -3,67400304 | 0,000222344 |
| Gad2 | 0,337513 | 0,443944 | 0,140866 | 0,170035 | 0,00531028 | 0 | 0 | 0 | 0 | 0 | Gad2 | -4,67957068 | 0,000223588 |
| Tgif1 | 16,4903 | 19,4874 | 8,20414 | 16,9414 | 4,89504 | 1,2331 | 1,68227 | 0,237102 | 0,0885604 | 0,279328 | Tgif1 | -2,14779822 | 0,000225635 |
| Dscam | 0,251301 | 0,123955 | 0,174809 | 0,506427 | 0 | 0,0104813 | 0 | 0 | 0 | 0 | Dscam | -4,26712791 | 0,000225778 |
| 1600016N20Ri | 0,498862 | 3,28078 | 0 | 3,35018 | 0 | 0 | 0 | 0 | 0 | 0 | 1600016N20Ri | -3,69506045 | 0,000227624 |
| Spink3 | 20,6032 | 14,5422 | 0 | 11,8538 | 0 | 0 | 0 | 0 | 0 | 0 | Spink3 | -4,76131356 | 0,000231138 |
| Tdo2 | 0,429869 | 4,66457 | 1,6145 | 1,29887 | 0 | 0,108119 | 0 | 0 | 0,0725098 | 0,0698817 | Tdo2 | -4,12412474 | 0,000234529 |
| Atp10a | 1,56393 | 2,68565 | 1,88566 | 0,93379 | 0,492136 | 0,246535 | 0,159694 | 0,105448 | 0,0967326 | 0,0748804 | Atp10a | -2,11320464 | 0,000236133 |
| Snrpf | 46,678 | 70,6193 | 53,6614 | 51,1796 | 18,5675 | 18,9311 | 21,3025 | 13,8095 | 10,4568 | 16,723 | Snrpf | -2,53803674 | 0,000239292 |
| **Hoxc10** | 0 | 1,41604 | 1,94691 | 2,16884 | 0 | 0 | 0 | 0 | 0 | 0 | Hoxc10 | -4,73155797 | 0,000239292 |
| Nfasc | 0,165809 | 0,261715 | 2,57587 | 1,15399 | 0,110356 | 0,041483 | 0,0298386 | 0 | 0,0110587 | 0,0107074 | Nfasc | -3,43882893 | 0,000256138 |
| Slc44a5 | 0 | 1,31531 | 1,83307 | 1,02726 | 0,0222197 | 0,019648 | 0,0480711 | 0 | 0,0524599 | 0 | Slc44a5 | -3,91755268 | 0,000256138 |
| Gls2 | 3,34812 | 1,38727 | 0,938917 | 1,77414 | 0 | 0 | 0 | 0 | 0 | 0 | Gls2 | -3,15062034 | 0,00025619 |
| Zic5 | 0,561928 | 0,138583 | 0,820795 | 2,5473 | 0 | 0 | 0 | 0 | 0,047168 | 0,0455451 | Zic5 | -4,69717221 | 0,00025619 |
| Stac | 0 | 1,43682 | 1,43675 | 1,20049 | 0 | 0 | 0 | 0 | 0 | 0 | Stac | -4,71065879 | 0,000257202 |
| Hid1 | 8,76482 | 3,1803 | 3,36367 | 3,3494 | 1,19919 | 0,530348 | 0,494206 | 0,274831 | 0,472368 | 0,293514 | Hid1 | -2,20234817 | 0,00026138 |
| Rpph1 | 287,464 | 52,0683 | 85,8515 | 222,529 | 35,7891 | 7,5848 | 6,64448 | 6,87172 | 8,94086 | 14,3589 | Rpph1 | -2,86535211 | 0,000262041 |
| Prokr2 | 0,0704875 | 0,278144 | 0,470699 | 0,781209 | 0 | 0 | 0 | 0 | 0 | 0,0227916 | Prokr2 | -4,66541407 | 0,000262261 |
| Alox12 | 0,329046 | 0,757402 | 8,78888 | 16,5747 | 0,435251 | 0,275021 | 0,134536 | 0,698011 | 0,551318 | 0,355178 | Alox12 | -3,60579372 | 0,000265942 |
| Dll4 | 24,8363 | 11,6936 | 2,51284 | 4,17034 | 1,75951 | 0,330099 | 0,346075 | 1,62564 | 2,70848 | 1,97901 | Dll4 | -3,06553823 | 0,000267077 |
| Sprr1a | 4,25295 | 3,14632 | 0,887167 | 1,60505 | 0 | 0 | 0 | 0 | 0 | 0 | Sprr1a | -4,63739826 | 0,000267325 |
| Otx1 | 0,466662 | 0,805625 | 1,46069 | 4,58374 | 0,0992178 | 0 | 0,0357804 | 0 | 0,0391121 | 0,0755786 | Otx1 | -3,77594139 | 0,000271642 |
| Dach2 | 0,0554018 | 0,512183 | 1,27652 | 1,36997 | 0,0156898 | 0,0138697 | 0 | 0 | 0 | 0 | Dach2 | -4,03164296 | 0,000271882 |
| Dlgap3 | 0,92961 | 0,833689 | 6,31346 | 2,80976 | 0,383113 | 0,148224 | 0,181313 | 0 | 0 | 0,0273402 | Dlgap3 | -2,90576516 | 0,000276153 |
| Emcn | 17,6031 | 47,19 | 21,2372 | 14,2457 | 7,92362 | 2,03255 | 2,17508 | 3,66209 | 8,96925 | 5,25631 | Emcn | -2,20126511 | 0,000277286 |
| Glrb | 0,440634 | 1,95607 | 4,41353 | 1,66467 | 0,0624494 | 0,13811 | 0,270243 | 0,100148 | 0,147661 | 0 | Glrb | -3,05684186 | 0,0002854 |
| Cpxm1 | 17,5335 | 19,067 | 19,2441 | 13,9077 | 5,83349 | 0,692923 | 0,931916 | 0,501725 | 0,324431 | 0,268532 | Cpxm1 | -2,38522922 | 0,000286427 |
| Itih2 | 0 | 1,85975 | 2,53522 | 3,79854 | 0,0593692 | 0 | 0 | 0 | 0,0701711 | 0 | Itih2 | -4,29084537 | 0,000288462 |
| **Hoxa7** | 0,175498 | 0,34625 | 2,92958 | 1,23743 | 0 | 0,0440897 | 0 | 0 | 0 | 0,0569737 | Hoxa7 | -4,33103508 | 0,000291649 |
| Msx3 | 0,293999 | 1,01509 | 0,490784 | 1,03656 | 0 | 0 | 0 | 0 | 0 | 0 | Msx3 | -4,60959544 | 0,000295704 |
| Trh | 1,30414 | 2,83025 | 1,08842 | 0,788014 | 0,0741144 | 0 | 0 | 0 | 0 | 0 | Trh | -4,18144699 | 0,000305067 |
| Exoc3l | 18,2834 | 9,43504 | 5,34682 | 1,74773 | 1,62723 | 0,334863 | 0,245661 | 0,363785 | 0,582252 | 0,865108 | Exoc3l | -2,65893723 | 0,00030806 |
| **Hoxd10** | 0 | 0,919188 | 2,33313 | 2,62793 | 0 | 0 | 0 | 0 | 0 | 0 | Hoxd10 | -4,67806389 | 0,000308722 |
| Smoc1 | 4,12407 | 3,14375 | 5,65374 | 9,96238 | 1,48739 | 0,164437 | 0,545922 | 0,0851535 | 0,69029 | 0,394333 | Smoc1 | -2,53452097 | 0,000317478 |
| E330009J07Ri | 2,63247 | 3,80875 | 1,46479 | 1,94453 | 0,398387 | 0,352718 | 0,484988 | 0,637227 | 0,413396 | 0,398816 | E330009J07Ri | -2,07294598 | 0,000318812 |
| Prr5 | 15,7993 | 12,8952 | 21,5854 | 19,8922 | 5,84527 | 1,13447 | 0,92428 | 0,170684 | 0,570093 | 1,16071 | Prr5 | -2,27402717 | 0,000319579 |
| Cdhr2 | 0,233425 | 0,844336 | 0 | 2,11662 | 0 | 0 | 0 | 0 | 0 | 0 | Cdhr2 | -4,70719706 | 0,000319874 |
| Mis18a | 31,7614 | 25,7858 | 10,556 | 11,888 | 5,35289 | 3,11659 | 2,59465 | 0,509031 | 0,476128 | 0,82456 | Mis18a | -2,01702302 | 0,000323183 |
| Scrt1 | 0,72188 | 0,079126 | 2,67806 | 0,48487 | 0 | 0,0401893 | 0,0245815 | 0 | 0,0268278 | 0 | Scrt1 | -4,03145916 | 0,000330388 |
| Dusp14 | 1,98614 | 2,93887 | 2,69364 | 2,75032 | 0,423185 | 0,125008 | 0 | 0 | 0,167868 | 0,0808223 | Dusp14 | -3,08316209 | 0,000332211 |
| Wnt3 | 0,104987 | 0,517847 | 0,350531 | 1,48078 | 0 | 0 | 0 | 0 | 0 | 0 | Wnt3 | -4,61088741 | 0,000344199 |
| Map2 | 0,116781 | 1,38246 | 4,97149 | 4,58892 | 0,330739 | 0,073096 | 0,125209 | 0 | 0,0195031 | 0,0566141 | Map2 | -3,17683657 | 0,000352144 |
| Gpr126 | 1,79001 | 12,9098 | 5,65368 | 2,48605 | 1,06854 | 0,0726543 | 0,10372 | 0,044046 | 0,129191 | 0,0312594 | Gpr126 | -3,1159414 | 0,000354857 |
| Rasip1 | 80,4429 | 58,5969 | 12,0274 | 9,7126 | 7,22762 | 1,76722 | 2,09778 | 2,26218 | 4,02804 | 3,25543 | Rasip1 | -2,55940505 | 0,000361236 |
| Chga | 0,365549 | 0,360605 | 1,06786 | 2,02512 | 0 | 0 | 0 | 0 | 0 | 0 | Chga | -4,5828132 | 0,000362 |
| Gp5 | 0,153529 | 0 | 4,22879 | 7,42344 | 0,043549 | 0 | 0 | 0 | 0 | 0 | Gp5 | -4,5612563 | 0,000364212 |
| Kdr | 79,049 | 67,0795 | 22,1405 | 14,5932 | 11,2844 | 1,9369 | 1,19384 | 3,86503 | 12,5506 | 8,00839 | Kdr | -2,74196452 | 0,000367071 |
| Was | 10,2137 | 4,47803 | 6,49517 | 1,79638 | 0,597905 | 0,12214 | 0 | 0,294421 | 0,109018 | 0 | Was | -3,36303264 | 0,000375542 |
| Sowaha | 0,0894561 | 0,882481 | 1,49339 | 0,99139 | 0,025348 | 0,0473256 | 0,0274206 | 0,0813664 | 0 | 0,173675 | Sowaha | -3,50033097 | 0,000375597 |
| Cacna1e | 0,121869 | 0,14427 | 0,305191 | 0,270164 | 0,0275966 | 0 | 0 | 0 | 0 | 0,00786726 | Cacna1e | -3,62431746 | 0,000375706 |
| Abi3 | 10,2628 | 12,6364 | 4,09481 | 7,50509 | 3,08941 | 2,78299 | 2,39084 | 0 | 0 | 0 | Abi3 | -2,50253743 | 0,000378537 |
| Satb2 | 2,46061 | 1,00648 | 1,10211 | 1,45125 | 0,373944 | 0,240426 | 0,220623 | 0,0546251 | 0,0601432 | 0,116386 | Satb2 | -2,06385049 | 0,000381921 |
| Wnt3a | 0,360221 | 0,592257 | 0,100224 | 1,20964 | 0 | 0 | 0 | 0 | 0 | 0 | Wnt3a | -4,59852131 | 0,000382724 |
| Cyp26c1 | 3,40699 | 0,672175 | 1,32697 | 0,915027 | 0 | 0 | 0 | 0 | 0 | 0 | Cyp26c1 | -3,68197644 | 0,000384959 |
| Ptgs2 | 1,31668 | 1,91015 | 1,22835 | 1,87282 | 0,351074 | 0,058207 | 0,0237352 | 0 | 0,0259011 | 0 | Ptgs2 | -2,775648 | 0,00038638 |
| Zbed5 | 14,9864 | 16,0379 | 12,0993 | 12,536 | 13,4001 | 19,0203 | 17,7652 | 29,6049 | 20,4031 | 23,203 | Zbed5 | -2,45628263 | 0,000391998 |
| Fam196b | 0,11805 | 0,407598 | 0,837586 | 0,832603 | 0,0334336 | 0 | 0 | 0 | 0 | 0 | Fam196b | -4,02898595 | 0,000394974 |
| Fev | 0,964722 | 1,66541 | 0,805159 | 0,485793 | 0 | 0 | 0 | 0 | 0 | 0 | Fev | -4,52641592 | 0,000399363 |
| Lhx5 | 0 | 0,238314 | 4,03283 | 1,70358 | 0 | 0 | 0 | 0 | 0 | 0 | Lhx5 | -4,65559763 | 0,00040103 |
| Slamf1 | 0,492544 | 0,121472 | 1,74725 | 1,11643 | 0 | 0,0308878 | 0 | 0,223825 | 0,123895 | 0,119685 | Slamf1 | -4,22825039 | 0,000405839 |
| Derl3 | 7,6167 | 1,3417 | 1,58919 | 3,83516 | 0,386551 | 0,137066 | 0 | 0 | 0,0921062 | 0,265912 | Derl3 | -3,28019248 | 0,000410357 |
| 2810408I11Ri | 7,98556 | 11,5123 | 4,6127 | 8,65319 | 0,702221 | 1,87581 | 0,380572 | 0 | 0 | 0 | 2810408I11Ri | -2,61365108 | 0,000413288 |
| En1 | 0,260398 | 0,256879 | 0,326023 | 2,3609 | 0 | 0 | 0 | 0,118312 | 0 | 0 | En1 | -4,5823516 | 0,000418353 |
| Mybpc1 | 0 | 11,2598 | 35,1963 | 22,5029 | 0,186119 | 0,102815 | 0,176165 | 0 | 0,137275 | 0,0796885 | Mybpc1 | -4,4320991 | 0,000422589 |
| Car7 | 4,14723 | 3,63653 | 0,192296 | 16,4753 | 0,130872 | 0,347855 | 0,212472 | 0,835509 | 0,622438 | 1,64901 | Car7 | -3,58595924 | 0,000423053 |
| Cntn1 | 0,0557205 | 0,0549675 | 0,838069 | 0,562003 | 0 | 0 | 0 | 0 | 0 | 0 | Cntn1 | -4,57923725 | 0,000433382 |
| Btk | 1,31267 | 1,42442 | 1,20522 | 2,9092 | 0,297765 | 0,0329351 | 0,0402703 | 0 | 0 | 0,0425425 | Btk | -3,09436841 | 0,000444989 |
| Ankrd13d | 1,26528 | 0,468064 | 5,80842 | 3,02703 | 0,269195 | 0,0397137 | 0,0485467 | 0 | 0 | 0,0513109 | Ankrd13d | -3,45206179 | 0,000445967 |
| Grid2 | 0,650352 | 1,60392 | 0,542846 | 0,764393 | 0,0614462 | 0 | 0 | 0 | 0 | 0,0350984 | Grid2 | -3,72625488 | 0,000451502 |
| Fabp5 | 20,7384 | 75,077 | 37,2679 | 42,24 | 22,3084 | 4,56252 | 4,10177 | 1,12372 | 3,97526 | 4,27231 | Fabp5 | -2,400692 | 0,000465032 |
| Neurod1 | 0 | 0,395476 | 2,78847 | 1,34618 | 0 | 0 | 0 | 0 | 0 | 0 | Neurod1 | -4,58882643 | 0,000466074 |

| Gene | d9-11 (1) | d9-11 (2) | d9-11(3) | d9-11(4) | newborn(1) | newborn(2) | newborn(3) | adult(1) | adult(2) | adult(3) | Gene | log2(fold) | FDR(*p* -value) |
|---|---|---|---|---|---|---|---|---|---|---|---|---|---|
| Sall3 | 4,69688 | 3,86875 | 0,989683 | 3,76769 | 0,477746 | 0,0228257 | 0 | 0 | 0 | 0 | Sall3 | -3,49956337 | 0,000468899 |
| Arhgap28 | 14,9183 | 21,3691 | 14,7939 | 7,63586 | 3,46463 | 0,313761 | 0,146244 | 0,0543148 | 0,338865 | 0,173583 | Arhgap28 | -2,86311906 | 0,000469969 |
| Foxl2 | 0,528488 | 0,912355 | 0,330838 | 0 | 0 | 0 | 0 | 0 | 0 | 0,0107053 | Foxl2 | -4,50644958 | 0,000471469 |
| Hemgn | 0 | 1,43419 | 4,3685 | 2,34324 | 0,0824661 | 0 | 0,0446138 | 0 | 0 | 0 | Hemgn | -4,02475209 | 0,000472027 |
| Tmem88 | 46,2429 | 24,8014 | 16,8999 | 19,5981 | 8,52909 | 2,32004 | 2,47311 | 0 | 4,21049 | 3,45607 | Tmem88 | -2,18825412 | 0,000476569 |
| Tmem200c | 0,507761 | 0,400723 | 0,508594 | 0,920791 | 0,0287812 | 0 | 0,0311359 | 0,277028 | 0 | 0,06575 | Tmem200c | -3,72778248 | 0,00048718 |
| Slc2a2 | 0,77588 | 1,14809 | 0,107935 | 3,51727 | 0,0733313 | 0 | 0 | 0,11751 | 0,0433872 | 0,167625 | Slc2a2 | -4,11496287 | 0,000493711 |
| Igfbp4 | 411,198 | 289,231 | 157,078 | 125,33 | 75,0564 | 8,43195 | 8,53844 | 0,597728 | 2,21373 | 1,97603 | Igfbp4 | -2,57458144 | 0,000493947 |
| Bdkrb2 | 6,86484 | 3,6938 | 0,694494 | 1,67618 | 0,0590596 | 0,0523111 | 0,383502 | 0 | 0,140296 | 0 | Bdkrb2 | -3,42940018 | 0,000499107 |
| Scg3 | 0 | 2,88783 | 2,44339 | 1,86244 | 0,0873403 | 0 | 0 | 0 | 0 | 0 | Scg3 | -4,15102159 | 0,000499107 |
| AY358078 | 1,83504 | 1,56884 | 0,561561 | 1,60167 | 0,0347527 | 0,0307963 | 0,150465 | 0 | 0 | 0 | AY358078 | -2,80027191 | 0,000499675 |
| Dnah2 | 0,135323 | 0,133498 | 0,60246 | 0,227265 | 0,0255353 | 0,0564092 | 0,0276163 | 0 | 0,0225485 | 0 | Dnah2 | -2,23863739 | 0,000500267 |
| Slc6a9 | 2,39017 | 6,87718 | 9,89226 | 10,9364 | 2,2015 | 0,84879 | 0,671731 | 0,181126 | 0,266812 | 0,580275 | Slc6a9 | -2,08324857 | 0,000508735 |
| Prdx6b | 1,1387 | 2,24664 | 0,807894 | 0,573582 | 0,290509 | 0,371225 | 0,226988 | 0,414054 | 0,343514 | 0,405679 | Prdx6b | -2,2004225 | 0,000515813 |
| Eya4 | 0,0783696 | 3,3244 | 0,654163 | 1,65814 | 0,199823 | 0,0785313 | 0,0480339 | 0 | 0 | 0,0760434 | Eya4 | -2,91413986 | 0,000526074 |
| Upk3b | 0,50188 | 0,825157 | 28,7642 | 8,08869 | 0,996743 | 0,25211 | 0,205439 | 2,12643 | 3,43224 | 0,86869 | Upk3b | -3,4882595 | 0,000542836 |
| Slc35f2 | 1,6825 | 1,14907 | 0,864217 | 0,651901 | 0,110091 | 0,0324709 | 0,11911 | 0 | 0 | 0,0419421 | Slc35f2 | -3,01390286 | 0,000548141 |
| Galnt14 | 0,540941 | 2,13452 | 0,677266 | 1,6348 | 0,0766969 | 0,0339347 | 0 | 0 | 0 | 0 | Galnt14 | -3,43568197 | 0,000561768 |
| Ano1 | 0,486259 | 7,61439 | 5,34003 | 2,87251 | 0,689406 | 0,0521931 | 0,0638533 | 0,126426 | 0,674033 | 0,652017 | Ano1 | -3,07377157 | 0,00056287 |
| Hdx | 0,39034 | 1,23222 | 1,3815 | 0,47193 | 0,0663509 | 0,117342 | 0,071773 | 0 | 0 | 0 | Hdx | -2,63647901 | 0,00057082 |
| Eomes | 1,23782 | 0,872221 | 0,147603 | 0,534472 | 0,0501057 | 0 | 0,0271012 | 0 | 0 | 0 | Eomes | -3,47451338 | 0,000574343 |
| Ccdc42 | 0,267951 | 2,3789 | 2,68353 | 1,61904 | 0 | 0 | 0 | 0 | 0 | 0 | Ccdc42 | -3,81731252 | 0,000587214 |
| Baiap3 | 0 | 0,745679 | 1,16663 | 0,813511 | 0 | 0 | 0 | 0 | 0 | 0 | Baiap3 | -3,8428403 | 0,000592175 |
| Chrna4 | 0,0720914 | 0,640063 | 1,68493 | 1,23479 | 0,0816872 | 0 | 0,0220901 | 0 | 0 | 0 | Chrna4 | -3,6723491 | 0,000599452 |
| Zfhx4 | 0,178184 | 1,09863 | 4,03458 | 2,01993 | 0,346737 | 0,0835598 | 0,0681808 | 0,0405814 | 0,014845 | 0,0287545 | Zfhx4 | -2,86725158 | 0,000619694 |
| Kcnk10 | 0,26276 | 2,33288 | 0,32898 | 0,661753 | 0,0372529 | 0 | 0 | 0 | 0 | 0 | Kcnk10 | -4,12222883 | 0,000624644 |
| Aspg | 1,81472 | 2,14824 | 1,71668 | 0,365634 | 0,171492 | 0,0303453 | 0,185536 | 0 | 0 | 0 | Aspg | -2,85085535 | 0,000627714 |
| Nkx2-1 | 0 | 1,027 | 0,874737 | 1,94762 | 0 | 0 | 0 | 0 | 0 | 0 | Nkx2-1 | -4,47030607 | 0,000638779 |
| Ocln | 3,57943 | 4,43907 | 0,256092 | 1,03032 | 0,318832 | 0,0512758 | 0,0627124 | 0,278979 | 0,102763 | 0,198648 | Ocln | -3,2898958 | 0,000661786 |
| Lhx1os | 5,9895 | 6,6464 | 1,24926 | 0,753137 | 0 | 0 | 0 | 0 | 0 | 0 | Lhx1os | -4,44339454 | 0,000665545 |
| Steap1 | 2,74256 | 1,50299 | 2,79745 | 2,45479 | 0,433255 | 0,153691 | 0,0938116 | 0 | 0 | 0 | Steap1 | -2,90649851 | 0,000669629 |
| Spns2 | 22,3718 | 17,0481 | 6,41368 | 3,30003 | 2,75913 | 0,50176 | 0,485255 | 0,205492 | 1,00551 | 1,06104 | Spns2 | -2,70137895 | 0,000679667 |
| Tjp3 | 3,2751 | 0 | 0 | 4,68958 | 0 | 0 | 0 | 0 | 0 | 0 | Tjp3 | -2,4904711 | 0,000680969 |
| Elmo1 | 11,5804 | 9,02541 | 2,27447 | 2,60386 | 1,46297 | 0,28424 | 0,167934 | 0,103345 | 0,341258 | 0,385255 | Elmo1 | -2,79815614 | 0,000690293 |
| 2900092D14R | 0 | 0,552872 | 0,677498 | 2,21948 | 0 | 0 | 0 | 0 | 0 | 0 | 2900092D14R | -4,47783549 | 0,000695619 |
| Sash3 | 2,95661 | 2,53622 | 0,64378 | 0,517995 | 0,218689 | 0,0645009 | 0,0394342 | 0 | 0,0431283 | 0 | Sash3 | -3,15229924 | 0,000702192 |
| Zcchc18 | 0,807044 | 1,16024 | 1,9859 | 2,72517 | 0,300075 | 0,0294976 | 0 | 0 | 0 | 0,0380975 | Zcchc18 | -3,2211533 | 0,000707354 |
| Shank2 | 0,693404 | 0,526868 | 0,9509 | 0,699563 | 0,181494 | 0,0267394 | 0,0303394 | 0 | 0 | 0,0517731 | Shank2 | -2,61277227 | 0,000712542 |
| Slc13a3 | 0,202361 | 0,698699 | 0,760099 | 0,407743 | 0,0286757 | 0 | 0 | 0 | 0 | 0,0327543 | Slc13a3 | -3,97952334 | 0,000712548 |
| Fbln7 | 0,56512 | 1,78395 | 0,754721 | 1,5941 | 0,160192 | 0,113371 | 0,138644 | 0,102746 | 0,0378825 | 0 | Fbln7 | -2,61767251 | 0,00071687 |
| Acap1 | 2,95547 | 10,2043 | 3,3639 | 0,405993 | 0,380938 | 0,033709 | 0,0412155 | 0 | 0 | 0,0870868 | Acap1 | -3,43481176 | 0,000723924 |
| Anxa3 | 20,9518 | 31,3287 | 31,292 | 18,2143 | 7,51521 | 1,44247 | 1,35561 | 0,799949 | 3,34932 | 2,00819 | Anxa3 | -2,21419045 | 0,000726021 |
| Rmrp | 339,131 | 152,47 | 83,5687 | 318,094 | 52,962 | 4,39209 | 22,4125 | 0 | 9,76219 | 8,53899 | Rmrp | -2,65872112 | 0,000735993 |
| Klf6 | 130,666 | 96,8642 | 45,7101 | 54,7902 | 32,0398 | 7,25891 | 6,75909 | 9,13451 | 11,292 | 9,12575 | Klf6 | -2,07908352 | 0,000737779 |
| **Hoxc9** | 0,202621 | 0,999399 | 0,845568 | 1,22449 | 0 | 0 | 0 | 0 | 0 | 0 | Hoxc9 | -4,38614846 | 0,000743606 |
| Klk1 | 11,5538 | 6,83808 | 0,385643 | 0 | 0 | 0 | 0 | 0 | 0 | 0 | Klk1 | -4,49610131 | 0,000743629 |
| Epha1 | 0 | 0 | 0 | 0 | 0 | 0 | 0 | 0 | 0 | 0 | Epha1 | -2,92889189 | 0,000745706 |
| Arap3 | 42,1018 | 27,5061 | 10,6495 | 4,28622 | 4,96534 | 0,745604 | 0,669941 | 1,66047 | 3,55126 | 2,38035 | Arap3 | -2,74698004 | 0,000749944 |
| Ccdc160 | 1,84601 | 0,682883 | 0,385175 | 0,929596 | 0 | 0 | 0 | 0,209192 | 0 | 0 | Ccdc160 | -4,38466311 | 0,000754086 |
| Pou4f1 | 0 | 0,754513 | 1,27684 | 2,48298 | 0 | 0,0212935 | 0,0781407 | 0 | 0,142165 | 0,109974 | Pou4f1 | -3,86578558 | 0,000754113 |
| Pax7 | 0 | 0,147916 | 1,12642 | 0,906412 | 0 | 0 | 0 | 0 | 0 | 0 | Pax7 | -4,45914482 | 0,000755474 |
| Crispld1 | 1,12411 | 3,48522 | 4,95964 | 2,18422 | 0,591457 | 0,0201159 | 0,049215 | 0,0730481 | 0,134282 | 0,077916 | Crispld1 | -3,0580535 | 0,000759406 |
| Aqp3 | 0,787405 | 0,582564 | 4,76466 | 7,33612 | 0,391138 | 0,148455 | 0,06047 | 0 | 0 | 0 | Aqp3 | -3,29180544 | 0,000763332 |
| Gjb3 | 0,666467 | 0,657448 | 0,927074 | 2,23738 | 0 | 0,0558875 | 0 | 0 | 0 | 0,0722495 | Gjb3 | -4,0023341 | 0,000764175 |
| L1td1 | 0,893238 | 0,330439 | 0,186395 | 1,01237 | 0 | 0 | 0 | 0 | 0 | 0 | L1td1 | -4,02682408 | 0,00076489 |
| Drp2 | 0,224802 | 0,532245 | 1,12591 | 1,13256 | 0,127306 | 0,0450094 | 0,0137692 | 0 | 0 | 0 | Drp2 | -2,83830857 | 0,000765263 |
| H2-Ab1 | 3,43108 | 4,30764 | 1,30155 | 12,5632 | 0,887051 | 0,15735 | 0,192076 | 0 | 0,317653 | 0,101791 | H2-Ab1 | -3,03652008 | 0,000769295 |
| Hapln4 | 0,90575 | 0,625463 | 0,529225 | 0,182507 | 0,0256655 | 0,0226985 | 0,0277642 | 0 | 0,0303149 | 0,0586182 | Hapln4 | -3,46969125 | 0,000771826 |
| Actg2 | 13,5844 | 4,37562 | 11,3372 | 2,51262 | 1,65473 | 0,908119 | 0,959619 | 0,564881 | 1,71369 | 1,01646 | Actg2 | -2,42817907 | 0,000775838 |
| Itpr3 | 7,4202 | 8,89357 | 6,30976 | 6,21903 | 2,46389 | 0,338337 | 0,318503 | 0,347365 | 0,39318 | 0,324691 | Itpr3 | -2,28745958 | 0,000778039 |
| Evx1 | 1,02946 | 0,112839 | 1,24118 | 0,115234 | 0 | 0 | 0 | 0 | 0 | 0 | Evx1 | -4,41033527 | 0,000791939 |
| Wdr93 | 0,294284 | 2,90306 | 1,10533 | 2,07513 | 0 | 0,110806 | 0,135465 | 0 | 0,0494251 | 0,143149 | Wdr93 | -3,17637893 | 0,000798048 |
| Skap1 | 3,59247 | 1,32243 | 0,97904 | 0 | 0,0475426 | 0 | 0 | 0 | 0 | 0 | Skap1 | -4,2132151 | 0,000801903 |
| Egr3 | 8,24057 | 7,58047 | 5,76682 | 9,56038 | 1,81594 | 0,0851263 | 0,104129 | 0 | 0,113668 | 0,0274781 | Egr3 | -2,9200806 | 0,000803468 |
| Nrsn2 | 0 | 2,84 | 12,0138 | 3,1629 | 0,148741 | 0,0659183 | 0,161007 | 0 | 0 | 0 | Nrsn2 | -3,59219026 | 0,000821046 |
| Dnah11 | 0,223847 | 0 | 0 | 0 | 0 | 0 | 0 | 0 | 0,0146325 | 0,0141716 | Dnah11 | -3,55513012 | 0,000823942 |
| Zim1 | 5,37336 | 8,77372 | 22,2712 | 9,14707 | 3,85914 | 0,928738 | 0,908786 | 0,758337 | 0,744277 | 0,419734 | Zim1 | -2,17337468 | 0,000824304 |
| H2-Q10 | 0,411232 | 5,60125 | 1,14896 | 1,06026 | 0,426179 | 0,313881 | 0,381709 | 0,880807 | 0,645193 | 0,423233 | H2-Q10 | -4,06210129 | 0,000829159 |
| Nuak2 | 3,99919 | 4,56809 | 2,28397 | 4,4531 | 1,07383 | 0,369395 | 0,129078 | 1,62671 | 1,12821 | 0,954077 | Nuak2 | -2,35187688 | 0,000838128 |
| Fbxo2 | 0 | 2,95274 | 3,99696 | 7,23402 | 0 | 0,0754725 | 0,18428 | 0,541863 | 0,20306 | 0,390547 | Fbxo2 | -3,86349793 | 0,00084696 |
| Slc5a3 | 5,00438 | 3,26515 | 2,32163 | 3,10555 | 1,35022 | 0,36416 | 0,314686 | 0 | 0 | 0 | Slc5a3 | -2,15799041 | 0,000850173 |
| Piezo2 | 1,59231 | 1,19954 | 2,22335 | 1,10855 | 0,442551 | 0,0941496 | 0,0531809 | 0,0263702 | 0,019304 | 0,0186926 | Piezo2 | -2,41039481 | 0,000857097 |
| Snai3 | 0,887327 | 1,31298 | 2,03659 | 3,57473 | 0,125987 | 0,223222 | 0,136356 | 4,62641 | 5,91471 | 3,96788 | Snai3 | -2,85053286 | 0,000872372 |
| Kremen2 | 0 | 0,789965 | 0,668387 | 2,74261 | 0,0908685 | 0 | 0 | 0 | 0 | 0 | Kremen2 | -3,92838454 | 0,000873375 |
| D930028M14I | 0 | 0,874542 | 9,98761 | 2,71971 | 0 | 0 | 0 | 0 | 0 | 0 | D930028M14I | -4,42328821 | 0,000874509 |
| Nxpe3 | 2,00747 | 2,51171 | 2,32965 | 1,2827 | 0,63778 | 0,183832 | 0,104973 | 0,0891543 | 0,0653776 | 0,110731 | Nxpe3 | -2,19680171 | 0,000874747 |
| Ambp | 0 | 0,295565 | 7,00157 | 6,03428 | 0 | 0 | 0 | 0 | 0 | 0 | Ambp | -4,23175023 | 0,000878574 |
| Tead4 | 8,30076 | 8,3986 | 2,84251 | 1,49706 | 1,20653 | 0,503045 | 0,273419 | 0,202639 | 0,224108 | 0,0481238 | Tead4 | -2,49446327 | 0,000882674 |
| Zic4 | 0 | 0 | 0,852829 | 1,97958 | 0 | 0 | 0 | 0 | 0 | 0 | Zic4 | -4,43699019 | 0,000890468 |
| St18 | 0 | 0,0546243 | 1,61771 | 0,557911 | 0 | 0 | 0 | 0 | 0 | 0 | St18 | -4,43521896 | 0,000897582 |
| Lrrtm4 | 0,216091 | 0,2844 | 0,721919 | 0,774531 | 0,0306247 | 0 | 0 | 0 | 0 | 0 | Lrrtm4 | -3,95093705 | 0,00089803 |
| 4930539E08R | 0,41703 | 0,4114 | 0,522156 | 0,280117 | 0,0196883 | 0,0174076 | 0,0425931 | 0 | 0 | 0,022473 | 4930539E08R | -3,38594608 | 0,000904108 |
| Cd109 | 10,4454 | 6,00209 | 2,42698 | 1,73602 | 1,31131 | 0,229139 | 0,164923 | 0,0490142 | 0,0898943 | 0,104391 | Cd109 | -2,69091691 | 0,000909094 |
| Grem2 | 0,17247 | 0,595495 | 0,647832 | 0,347528 | 0 | 0 | 0 | 0 | 0 | 0 | Grem2 | -3,9190326 | 0,000909891 |
| Nkx3-1 | 0,312589 | 0,513944 | 0,695778 | 0,209946 | 0,0295318 | 0 | 0 | 0 | 0 | 0 | Nkx3-1 | -3,92203532 | 0,000911405 |
| Krt15 | 0,202486 | 0,798987 | 0,169001 | 2,24341 | 0 | 0 | 0 | 0 | 0 | 0 | Krt15 | -4,35283902 | 0,000911405 |
| Gad1 | 0,201891 | 0,89624 | 0,421296 | 0,915292 | 0,028609 | 0,0253051 | 0 | 0 | 0 | 0 | Gad1 | -3,6309019 | 0,000926506 |
| Qprt | 0,621283 | 3,06431 | 3,88871 | 9,38396 | 0,441705 | 0,705154 | 0 | 0 | 0 | 0 | Qprt | -2,77886081 | 0,000944602 |
| Atp2b4 | 5,13062 | 19,154 | 36,29 | 18,1821 | 5,74088 | 0,801394 | 0,614865 | 0,281093 | 0,746541 | 0,697801 | Atp2b4 | -2,52500349 | 0,000953998 |
| Tnc | 9,1044 | 70,0212 | 109,109 | 56,2887 | 7,14134 | 0,575162 | 0,419584 | 0,160968 | 0,265478 | 0,18559 | Tnc | -3,49391725 | 0,000960877 |
| Celf3 | 0,108122 | 0,942655 | 6,64527 | 6,53042 | 0,0919396 | 0,0342023 | 0 | 0 | 0 | 0,0441801 | Celf3 | -3,33927004 | 0,000969178 |
| Csn3 | 3,74926 | 0,870241 | 0,184071 | 0,222126 | 0 | 0 | 0 | 0 | 0 | 0 | Csn3 | -4,37787675 | 0,000971257 |
| F2rl1 | 1,31769 | 0,354514 | 1,09986 | 4,70645 | 0,1698 | 0,0300455 | 0,0734819 | 0 | 0 | 0,0388059 | F2rl1 | -3,24952824 | 0,000983371 |
| Chl1 | 0 | 0,292101 | 0,910012 | 0,0813684 | 0 | 0 | 0 | 0 | 0,0538838 | 0,0260811 | Chl1 | -4,38934604 | 0,000985552 |
| Spock3 | 0,211764 | 1,90534 | 2,82679 | 0,921714 | 0,0900362 | 0,106198 | 0,0649325 | 0 | 0 | 0 | Spock3 | -3,16119548 | 0,000989043 |
| B3gat1 | 0,28443 | 0,374118 | 1,26621 | 0,573114 | 0 | 0 | 0,0581295 | 0 | 0 | 0 | B3gat1 | -3,70997078 | 0,001013028 |
| Hist1h2ai | 38,2365 | 18,1087 | 20,1263 | 31,0149 | 2,86046 | 1,10769 | 0,94885 | 1,06522 | 0,527902 | 0,55846 | Hist1h2ai | -3,17172635 | 0,001015346 |
| Tmem229a | 0,319021 | 0,314715 | 0,159777 | 0,321432 | 0 | 0 | 0 | 0 | 0 | 0 | Tmem229a | -3,85844394 | 0,00104506 |
| Wdr86 | 11,2499 | 8,22763 | 5,18045 | 7,81467 | 1,54153 | 0,0975102 | 0,0595796 | 0 | 0,130695 | 0,0630129 | Wdr86 | -2,9931783 | 0,001055413 |
| Hoxa11os | 0,187623 | 0,555255 | 1,16756 | 2,07881 | 0 | 0 | 0 | 0 | 0 | 0 | Hoxa11os | -4,335954 | 0,001059345 |
| Lacc1 | 2,21095 | 7,36111 | 5,6516 | 3,06246 | 1,56746 | 0,381455 | 0,25439 | 0,376635 | 0,788625 | 0,985518 | Lacc1 | -2,31615887 | 0,001059949 |
| Lurap1l | 1,4375 | 2,8361 | 0,856983 | 4,24015 | 0,349797 | 0,206544 | 0,126187 | 1,11769 | 1,17702 | 1,80207 | Lurap1l | -2,85307311 | 0,001068889 |
| Tll1 | 1,74356 | 0,605194 | 0,431228 | 1,17115 | 0,164617 | 0,0808545 | 0,0989221 | 0,0587707 | 0,0647332 | 0,0417511 | Tll1 | -2,55715717 | 0,00107007 |
| Dcx | 0 | 0,13715 | 3,25135 | 1,74901 | 0,0786961 | 0,0173833 | 0,0212734 | 0 | 0 | 0 | Dcx | -3,738235 | 0,00109451 |
| Mpzl2 | 0,304352 | 0,300241 | 1,01617 | 2,14636 | 0,115009 | 0 | 0 | 0 | 0,0339784 | 0 | Mpzl2 | -3,71497491 | 0,001105405 |
| Wdr17 | 0,0774247 | 0,919212 | 0,0646277 | 1,4082 | 0 | 0,0193956 | 0 | 0 | 0 | 0 | Wdr17 | -4,08684649 | 0,001124039 |
| Sp5 | 5,50334 | 3,03381 | 0 | 4,72835 | 0,229596 | 0 | 0,0496871 | 0 | 0 | 0 | Sp5 | -3,79426085 | 0,001128396 |
| Ccbe1 | 3,63401 | 1,79248 | 1,19499 | 0,332868 | 0,311862 | 0,0275685 | 0,0168659 | 1,45353 | 2,05935 | 1,97503 | Ccbe1 | -3,17518191 | 0,001131074 |
| Tmem40 | 3,13042 | 0,441128 | 6,34395 | 8,41028 | 0,382384 | 0 | 0 | 0 | 0 | 0 | Tmem40 | -3,76841184 | 0,001156138 |
| Slc1a4 | 6,80815 | 8,49643 | 7,80536 | 14,7103 | 3,43949 | 0,534325 | 0,427368 | 0,373105 | 0,21951 | 0,185739 | Slc1a4 | -2,26979596 | 0,001183677 |
| Mmrn1 | 1,03301 | 2,64958 | 2,12744 | 1,87345 | 0,643969 | 0,103451 | 0,147707 | 0,12534 | 0,345269 | 0,111332 | Mmrn1 | -2,22279915 | 0,001199318 |
| Rims4 | 0,990524 | 2,93117 | 5,78565 | 3,98779 | 0 | 0,376786 | 0,30611 | 0,892478 | 0,170132 | 0,325543 | Rims4 | -2,95744931 | 0,001259261 |
| Stat4 | 0 | 0,760143 | 0,826939 | 2,66155 | 0,0312021 | 0,027602 | 0,0337559 | 0 | 0 | 0 | Stat4 | -3,70695359 | 0,001259716 |

| Gene | d9-11 (1) | d9-11 (2) | d9-11(3) | d9-11(4) | newborn(1) | newborn(2) | newborn(3) | adult(1) | adult(2) | adult(3) | Gene | log2(fold) | FDR(*p*-value) |
|---|---|---|---|---|---|---|---|---|---|---|---|---|---|
| Grid2ip | 0,0938446 | 0,925772 | 0,861657 | 2,26912 | 0,0265934 | 0 | 0 | 0 | 0 | 0 | Grid2ip | -3,48993935 | 0,001284316 |
| Plk3 | 20,9461 | 11,1692 | 10,1591 | 16,3955 | 3,29123 | 0,0710405 | 0,30399 | 0,51429 | 0,285134 | 0,230783 | Plk3 | -2,67437317 | 0,001287941 |
| Gp9 | 0 | 0 | 4,27885 | 20,1279 | 0 | 0 | 0 | 0 | 0 | 0 | Gp9 | -4,0368364 | 0,001305754 |
| 9330159F19Ri | 0,168166 | 0,608286 | 0,421119 | 0,677759 | 0,0635 | 0 | 0 | 0 | 0 | 0 | 9330159F19Ri | -3,43154645 | 0,001312701 |
| Rasef | 0,428819 | 0,423031 | 0,204541 | 1,66652 | 0,104102 | 0,0153389 | 0 | 0,278783 | 0,368374 | 0,118806 | Rasef | -3,31560878 | 0,001353486 |
| Kctd17 | 26,1469 | 22,7139 | 20,7845 | 20,3278 | 7,0715 | 0,795176 | 0,914404 | 0,623812 | 1,31765 | 1,10165 | Kctd17 | -2,27505878 | 0,001360287 |
| Pax8 | 0,517473 | 0,25524 | 1,18779 | 0,130325 | 0 | 0 | 0,0132288 | 0 | 0 | 0 | Pax8 | -4,23699527 | 0,001364418 |
| Arhgdig | 3,73598 | 4,79088 | 4,05304 | 7,14654 | 1,3826 | 0,660826 | 0,460692 | 0 | 0,381987 | 0,488841 | Arhgdig | -2,15122662 | 0,001364959 |
| Il3ra | 5,43082 | 15,8167 | 12,3025 | 11,4593 | 3,45364 | 0,260511 | 0,556782 | 0 | 0,174978 | 0,336892 | Il3ra | -2,27753323 | 0,001374313 |
| Lgi4 | 0,942809 | 0,279023 | 1,88872 | 0,759889 | 0,106869 | 0 | 0 | 0 | 0,126251 | 0,0305126 | Lgi4 | -3,53317938 | 0,001396477 |
| Isl2 | 0,372488 | 0,183725 | 5,2852 | 2,06354 | 0 | 0,0468041 | 0,114399 | 0,169019 | 0 | 0 | Isl2 | -3,73113737 | 0,001400063 |
| Cpne2 | 3,79448 | 8,63815 | 7,06515 | 4,70447 | 2,06958 | 0,293062 | 0,447855 | 0,397706 | 0,5392 | 0,567901 | Cpne2 | -2,21949821 | 0,001400925 |
| Muc13 | 0,884472 | 1,19972 | 0,461414 | 1,11382 | 0,0626768 | 0 | 0,0339035 | 0 | 0 | 0,035803 | Muc13 | -3,1960775 | 0,001406063 |
| Redrum | 0,605257 | 0,447807 | 0,252593 | 0,609699 | 0 | 0 | 0 | 0 | 0 | 0 | Redrum | -4,19727604 | 0,001414226 |
| Ifitm1 | 39,578 | 24,1671 | 16,773 | 0,667894 | 3,4693 | 0,169168 | 0,19466 | 0,595822 | 0 | 0,658713 | Ifitm1 | -3,56518967 | 0,001424907 |
| Epha6 | 0 | 0,225317 | 0,635498 | 0,920476 | 0 | 0 | 0 | 0 | 0,12729 | 0,123112 | Epha6 | -4,24828655 | 0,001427746 |
| Kif26a | 11,2584 | 6,94153 | 8,79148 | 11,0238 | 3,67202 | 0,511275 | 0,347574 | 0,330671 | 0,969635 | 0,659908 | Kif26a | -2,26782207 | 0,001429057 |
| Pmaip1 | 2,24738 | 2,95602 | 4,27277 | 3,0187 | 1,06196 | 0,250563 | 0,114895 | 0 | 0,125638 | 0,0404545 | Pmaip1 | -2,23861395 | 0,001450383 |
| Slc44a4 | 0,285983 | 1,26953 | 0 | 2,3047 | 0 | 0 | 0 | 0 | 0 | 0 | Slc44a4 | -4,26736203 | 0,001465683 |
| Brsk2 | 0,212794 | 0,470291 | 3,28917 | 3,12117 | 0,0602784 | 0 | 0,02435 | 0 | 0,00530274 | 0 | Brsk2 | -3,42821772 | 0,001471507 |
| Itga2 | 1,29265 | 3,82559 | 2,22146 | 0,383052 | 0,323118 | 0,038095 | 0,0233014 | 0 | 0 | 0 | Itga2 | -2,98427676 | 0,001503171 |
| Gria2 | 0,464635 | 1,00839 | 0,851725 | 0 | 0 | 0,0114939 | 0 | 0 | 0 | 0 | Gria2 | -3,98657422 | 0,001506148 |
| Rac3 | 0,707237 | 7,32527 | 7,67269 | 16,0216 | 0,704434 | 0,625118 | 1,19867 | 0,63933 | 0,240728 | 0,231171 | Rac3 | -2,51337905 | 0,001540432 |
| Slc4a8 | 0,131387 | 0,36292 | 0,833532 | 1,5887 | 0,111573 | 0,00657292 | 0,016089 | 0,0239364 | 0,0525548 | 0,0254469 | Slc4a8 | -3,09724949 | 0,001554397 |
| Olfm2 | 0,465386 | 6,65675 | 2,71891 | 3,5153 | 0,264358 | 0,0585563 | 0,0715316 | 0 | 0 | 0 | Olfm2 | -2,9301192 | 0,00160278 |
| Phox2b | 0 | 0,228862 | 1,83962 | 0,818017 | 0 | 0 | 0 | 0 | 0 | 0 | Phox2b | -4,23622689 | 0,001603201 |
| Tenm4 | 8,64439 | 4,75366 | 6,67958 | 7,53336 | 2,52145 | 0,314141 | 0,169879 | 0,106402 | 0,0973649 | 0 | Tenm4 | -2,34209879 | 0,001634208 |
| B4galnt3 | 0,443526 | 1,09384 | 1,48083 | 3,12778 | 0,3143 | 0,139019 | 0,0680051 | 0 | 0,0743186 | 0 | B4galnt3 | -2,63341608 | 0,001637877 |
| Fam89a | 0,831695 | 3,82867 | 2,77646 | 4,46687 | 0,551576 | 0,139711 | 0,255898 | 0 | 0,469503 | 0 | Fam89a | -2,51492968 | 0,001639217 |
| Osr2 | 1,4269 | 0 | 3,0624 | 4,9275 | 0 | 0,0512531 | 0 | 0,369829 | 0,0687193 | 0,0662474 | Osr2 | -3,67646042 | 0,00164733 |
| Gsg1 | 0,605816 | 1,79282 | 1,51677 | 1,22007 | 0 | 0 | 0 | 0 | 0 | 0 | Gsg1 | -3,76306326 | 0,001678243 |
| Ppfia3 | 0,588939 | 0,58099 | 1,63867 | 1,05491 | 0,11121 | 0,180257 | 0,160389 | 0,0595534 | 0,0218667 | 0,0634641 | Ppfia3 | -2,1324377 | 0,001706591 |
| Crym | 0 | 0,867052 | 5,86842 | 7,37586 | 0,0832963 | 0,147719 | 0,0901758 | 1,06092 | 0,794701 | 0,764356 | Crym | -3,62850937 | 0,001709251 |
| Rspo1 | 0 | 3,71315 | 8,6395 | 3,79134 | 0,213658 | 0 | 0 | 0,170789 | 0,316917 | 0,30564 | Rspo1 | -3,82076792 | 0,00171247 |
| Fhdc1 | 2,23644 | 1,10313 | 0,533381 | 1,1267 | 0,331748 | 0,159958 | 0,179413 | 0,0484743 | 0,0355598 | 0,0172062 | Fhdc1 | -2,09460151 | 0,001719477 |
| Nrxn3 | 0,0789816 | 0,19057 | 0,42854 | 0,243305 | 0,0273506 | 0 | 0,0189159 | 0 | 0 | 0 | Nrxn3 | -3,42152721 | 0,00173797 |
| Fgfr4 | 1,35039 | 1,33215 | 2,42774 | 2,09301 | 0,382731 | 0 | 0,0955495 | 0 | 0,0347961 | 0,0336302 | Fgfr4 | -2,81833819 | 0,001751656 |
| Rab38 | 1,34274 | 2,8699 | 1,68101 | 0,450785 | 0,127103 | 0 | 0,137565 | 0 | 0 | 0 | Rab38 | -3,24068805 | 0,001751825 |
| Tmem27 | 0,582607 | 2,87357 | 0,972451 | 0,29334 | 0 | 0 | 0 | 0 | 0 | 0 | Tmem27 | -4,16119099 | 0,001753812 |
| Angpt2 | 1,0921 | 3,41162 | 3,9502 | 2,01718 | 0,747868 | 0,0456149 | 0,0836919 | 0,165551 | 0,152305 | 0,176699 | Angpt2 | -2,45983943 | 0,001764189 |
| Mak | 0,796203 | 0,0981811 | 0,913812 | 0,300806 | 0 | 0 | 0 | 0 | 0 | 0 | Mak | -3,16624853 | 0,001770175 |
| Cxcr4 | 5,33572 | 14,0988 | 4,29434 | 9,40512 | 2,27161 | 0,143685 | 0,175591 | 0,518758 | 0,770247 | 0,928492 | Cxcr4 | -2,72740126 | 0,001777265 |
| Notch1 | 32,742 | 32,4621 | 15,3362 | 18,3141 | 8,8976 | 0,738714 | 0,713192 | 0,747008 | 1,80517 | 1,3347 | Notch1 | -2,39885077 | 0,001782302 |
| Ildr1 | 0,338946 | 0,445823 | 0,0943051 | 0,796753 | 0 | 0 | 0 | 0 | 0 | 0 | Ildr1 | -4,14817318 | 0,001788956 |
| Bai1 | 0 | 0,349536 | 0,676026 | 2,55005 | 0,0430007 | 0 | 0,0310066 | 0,0921626 | 0,0168983 | 0,0654167 | Bai1 | -3,71036867 | 0,001804432 |
| St6gal2 | 0,147642 | 0,728231 | 0,616159 | 0,44618 | 0 | 0 | 0 | 0 | 0 | 0 | St6gal2 | -4,13294934 | 0,001819472 |
| Fst | 1,29178 | 3,53978 | 3,23464 | 3,32551 | 0,814129 | 0,108076 | 0,0440437 | 0 | 0,0964024 | 0,139617 | Fst | -2,64413175 | 0,001820136 |
| Sp6 | 0,481707 | 0,570245 | 0,562915 | 1,74711 | 0,0819062 | 0,144886 | 0,0590703 | 0,525689 | 0,161284 | 0,187092 | Sp6 | -2,6872577 | 0,001822463 |
| Cldn3 | 1,1641 | 0,861246 | 1,94304 | 8,49873 | 0,330943 | 0,0733605 | 0,0895683 | 0 | 0,0986599 | 0,284692 | Cldn3 | -3,28031448 | 0,001826826 |
| Sst | 0 | 0,786782 | 11,3129 | 9,62791 | 0 | 0 | 0 | 0 | 0 | 0 | Sst | -4,20458763 | 0,001838853 |
| Abcg8 | 0,216214 | 0,213292 | 0,721965 | 0 | 0 | 0 | 0 | 0 | 0 | 0 | Abcg8 | -4,1257434 | 0,001856306 |
| Ece1 | 69,6386 | 109,313 | 52,5936 | 32,1358 | 27,2807 | 4,73774 | 4,80327 | 6,8327 | 10,338 | 7,93395 | Ece1 | -2,03301684 | 0,001859968 |
| Kcna1 | 0,139012 | 0,274274 | 0,17406 | 0,665346 | 0,0491937 | 0,00869515 | 0,010641 | 0 | 0,127496 | 0,123441 | Kcna1 | -3,01805149 | 0,001877217 |
| 2010015L04Ri | 0,512044 | 2,15109 | 0,309536 | 1,79255 | 0,246228 | 0 | 0 | 0 | 0 | 0 | 2010015L04Ri | -3,12971226 | 0,001887058 |
| Plxnc1 | 7,30762 | 9,4948 | 4,46338 | 2,55916 | 1,75373 | 0,13382 | 0,136461 | 0 | 0,0446103 | 0,0431805 | Plxnc1 | -2,58768801 | 0,001904533 |
| Ubxn11 | 1,2283 | 3,3865 | 2,05028 | 2,96887 | 0,488511 | 0,123686 | 0,206348 | 0,667894 | 0,453241 | 0,159931 | Ubxn11 | -2,56367524 | 0,001906531 |
| Hfm1 | 0,24466 | 0,625103 | 1,66191 | 0,446911 | 0,0358949 | 0,062072 | 0,058239 | 0 | 0 | 0 | Hfm1 | -2,75422649 | 0,001906531 |
| Rhag | 0,144968 | 1,00106 | 2,78301 | 2,77465 | 0,205574 | 0 | 0,0444858 | 0 | 0 | 0 | Rhag | -3,34919308 | 0,001955129 |
| Sim2 | 0 | 0,434594 | 0,514815 | 0,710153 | 0 | 0 | 0 | 0 | 0,0294849 | 0 | Sim2 | -4,14706963 | 0,00196805 |
| Pkd1l3 | 0 | 0 | 0 | 1,64365 | 0 | 0 | 0 | 0 | 0 | 0 | Pkd1l3 | -2,45455092 | 0,001978955 |
| Myh3 | 0 | 0,935388 | 9,81143 | 7,62973 | 0,0950985 | 0,0282568 | 0,0268915 | 0 | 0,021106 | 0 | Myh3 | -4,07237221 | 0,001980841 |
| Ccdc60 | 0,771942 | 1,3961 | 0,751712 | 0,388825 | 0,0729583 | 0 | 0 | 0 | 0 | 0 | Ccdc60 | -3,39887117 | 0,001980925 |
| Pcp4 | 5,74939 | 4,2533 | 1,79878 | 2,16894 | 0 | 0 | 0,223067 | 0 | 1,25221 | 0,714506 | Pcp4 | -3,7304783 | 0,001983896 |
| Lrrc34 | 1,50006 | 0,924854 | 1,56501 | 0,377732 | 0,106433 | 0,0471232 | 0 | 0 | 0 | 0 | Lrrc34 | -3,11699012 | 0,002036768 |
| Esx1 | 0,660199 | 0,434176 | 0 | 5,31945 | 0 | 0 | 0 | 0 | 0 | 0,0715642 | Esx1 | -4,18549893 | 0,002042738 |
| A2m | 0,409819 | 1,54977 | 1,76744 | 3,78502 | 0,328907 | 0 | 0,0418556 | 0,0621571 | 0,0456568 | 0 | A2m | -3,11912049 | 0,002071074 |
| Kcnmb4 | 5,02337 | 9,9104 | 14,1884 | 7,39144 | 1,98046 | 0,976786 | 0,476617 | 0 | 0 | 0 | Kcnmb4 | -2,28699118 | 0,002079977 |
| Unc5c | 5,22679 | 5,12322 | 5,50974 | 2,46454 | 0,37941 | 0,0335304 | 0 | 0 | 0 | 0 | Unc5c | -3,60442982 | 0,002088283 |
| Xkr7 | 0 | 0,72211 | 0,61099 | 0,491617 | 0 | 0 | 0 | 0 | 0 | 0 | Xkr7 | -4,12729805 | 0,002089742 |
| Palm | 22,867 | 49,6798 | 30,2296 | 31,8671 | 13,6226 | 1,95673 | 2,10901 | 1,59053 | 3,81575 | 2,12628 | Palm | -2,02566703 | 0,002112584 |
| Foxj1 | 0,902125 | 1,1442 | 2,36653 | 1,29828 | 0,32887 | 0,0969983 | 0,118604 | 0 | 0 | 0,0417636 | Foxj1 | -2,32667482 | 0,002118007 |
| Gp6 | 0 | 2,56953 | 0,271733 | 14,0978 | 0,0926192 | 0 | 0 | 0 | 0 | 0 | Gp6 | -4,05576923 | 0,002127686 |
| F2 | 0 | 1,01704 | 0,717089 | 1,21157 | 0 | 0 | 0 | 0 | 0 | 0 | F2 | -3,74255653 | 0,002145233 |
| Gm4961 | 1,34775 | 1,32953 | 1,93736 | 1,28222 | 0,169898 | 0,0375928 | 0,0229788 | 0 | 0 | 0 | Gm4961 | -3,09770464 | 0,00217999 |
| Fam181a | 0,255999 | 0,757596 | 1,70924 | 1,80468 | 0 | 0 | 0 | 0,231944 | 0 | 0,0833639 | Fam181a | -3,72411646 | 0,002206158 |
| Fmn2 | 0,146605 | 0,482089 | 1,95803 | 1,67414 | 0,166056 | 0,061159 | 0,0898034 | 0 | 0 | 0,0157882 | Fmn2 | -2,64683836 | 0,002217579 |
| G6b | 0,598116 | 0,442321 | 0,998431 | 1,13266 | 0,0848232 | 0 | 0 | 0 | 0 | 0 | G6b | -3,4729879 | 0,002238026 |
| Cacnb3 | 11,3859 | 17,4702 | 21,0264 | 26,1688 | 7,97101 | 1,0066 | 1,19072 | 0,470437 | 0,48509 | 0,125826 | Cacnb3 | -2,05048654 | 0,00226399 |
| Cldn9 | 0,248267 | 0,734718 | 1,03602 | 1,25015 | 0 | 0 | 0 | 0 | 0 | 0 | Cldn9 | -4,06647671 | 0,002269812 |
| Grem1 | 0,214027 | 0 | 2,14358 | 1,72452 | 0,0607694 | 0 | 0 | 0 | 0 | 0,0695842 | Grem1 | -3,83867186 | 0,002300824 |
| St6galnac5 | 0,170769 | 1,51614 | 2,13798 | 0,860068 | 0,0484527 | 0,0428956 | 0,104861 | 0 | 0 | 0 | St6galnac5 | -3,16226162 | 0,002322753 |
| Tram2 | 16,3481 | 16,8946 | 11,9114 | 7,05585 | 3,83414 | 0,653517 | 0,638505 | 1,17546 | 1,84364 | 1,43672 | Tram2 | -2,2881036 | 0,002342354 |
| Tox2 | 6,8884 | 9,88392 | 6,09767 | 6,72773 | 2,19273 | 0,472426 | 0,320686 | 0 | 0 | 0,271405 | Tox2 | -2,20773792 | 0,002356384 |
| Tceal6 | 1,77993 | 2,18977 | 1,98055 | 4,15327 | 1,84882 | 1,61181 | 1,54056 | 0,153293 | 0 | 0 | Tceal6 | -3,65821064 | 0,002398173 |
| Syt1 | 0,067338 | 0,332175 | 0,787209 | 0,474995 | 0,0190738 | 0 | 0 | 0 | 0 | 0 | Syt1 | -3,57286634 | 0,002416748 |
| Ldb2 | 12,7003 | 8,99535 | 2,58528 | 2,2569 | 1,93583 | 0,581061 | 0,56648 | 0,239693 | 0,752418 | 0,341971 | Ldb2 | -2,28300875 | 0,002425954 |
| Lhx9 | 0,0701947 | 0,161392 | 5,25069 | 2,29591 | 0,0959549 | 0 | 0 | 0 | 0 | 0 | Lhx9 | -3,78258327 | 0,002427453 |
| Mfng | 12,045 | 5,56973 | 6,28327 | 6,25571 | 2,83097 | 0,472991 | 0,809248 | 0,853945 | 1,33105 | 0,794664 | Mfng | -2,04766016 | 0,002440457 |
| Rpl34 | 70,773 | 50,8244 | 76,044 | 95,9799 | 19,615 | 12,792 | 11,9778 | 9,45413 | 7,5342 | 8,76305 | Rpl34 | -2,16297603 | 0,002445516 |
| 6720416L17Ri | 1,22419 | 0,134183 | 0,227068 | 0,411075 | 0 | 0 | 0 | 0 | 0 | 0 | 6720416L17Ri | -4,06172898 | 0,002481681 |
| Rab25 | 1,44471 | 0 | 0,904228 | 4,36367 | 0 | 0 | 0 | 0 | 0 | 0 | Rab25 | -4,08452873 | 0,002557158 |
| Prr7 | 3,72832 | 11,0335 | 4,66741 | 6,43649 | 1,89196 | 0,536653 | 0,409604 | 0,723684 | 0,180275 | 0,433778 | Prr7 | -2,13248734 | 0,002579566 |
| 2900011O08R | 0 | 0,982792 | 0,356377 | 2,52513 | 0 | 0 | 0 | 0 | 0 | 0 | 2900011O08R | -4,07069171 | 0,002595544 |
| St8sia3 | 0,301046 | 0,0494972 | 0,209411 | 0,10111 | 0 | 0 | 0 | 0 | 0 | 0 | St8sia3 | -4,03133446 | 0,002622409 |
| Kcnip1 | 0,150045 | 3,01651 | 2,41491 | 0,171436 | 0,0425584 | 0 | 0 | 0 | 0 | 0 | Kcnip1 | -3,71951565 | 0,002636145 |
| Pbx4 | 0,942812 | 0,265726 | 2,24816 | 5,69673 | 0,229661 | 0,06786 | 0 | 0 | 0 | 0 | Pbx4 | -3,12734954 | 0,002660397 |
| Dusp2 | 23,2444 | 11,3578 | 9,06552 | 2,84435 | 3,33094 | 0,710349 | 0,667596 | 0,197003 | 0,146595 | 0,282546 | Dusp2 | -2,36653767 | 0,002669154 |
| Psd2 | 0,0705366 | 0,139169 | 0,289234 | 0,42641 | 0 | 0 | 0 | 0 | 0,0235782 | 0 | Psd2 | -4,01665319 | 0,002726543 |
| Neurod4 | 0 | 0,191904 | 0,974254 | 0,78394 | 0 | 0 | 0 | 0 | 0 | 0 | Neurod4 | -4,05517661 | 0,002760032 |
| Tunar | 0,0992655 | 0,881323 | 0,165714 | 0,30002 | 0 | 0 | 0 | 0 | 0 | 0 | Tunar | -4,0212199 | 0,002763149 |
| Nts | 5,05809 | 12,4738 | 1,84679 | 1,59161 | 0,629369 | 0,637981 | 0,0973441 | 0 | 0,107344 | 0,206366 | Nts | -2,7660526 | 0,002834366 |
| Lyl1 | 13,4488 | 10,4258 | 6,25487 | 6,58143 | 0 | 0 | 0 | 0 | 0 | 0 | Lyl1 | -2,42576644 | 0,002845482 |
| Dkk2 | 0 | 1,54884 | 2,18421 | 0,878783 | 0,123574 | 0,0218567 | 0 | 0 | 0 | 0 | Dkk2 | -3,46001008 | 0,002846596 |
| Nfe2 | 3,77545 | 2,68982 | 4,02638 | 9,084 | 1,3101 | 0,263733 | 0,128896 | 0,190244 | 0 | 0,0681815 | Nfe2 | -2,49723338 | 0,002849722 |
| Areg | 5,58015 | 3,36387 | 0 | 2,49729 | 0 | 0 | 0 | 0 | 0 | 0 | Areg | -3,5637733 | 0,002858464 |
| Bsn | 0,0773996 | 0,190889 | 0,549183 | 0,662941 | 0,0876267 | 0,0241957 | 0,0118458 | 0 | 0,0257886 | 0 | Bsn | -2,71568873 | 0,002894591 |
| Ntrk2 | 0,344165 | 0,255432 | 0,50434 | 3,0593 | 0,290782 | 0,0223492 | 0,0136742 | 0 | 0,059603 | 0,0288462 | Ntrk2 | -3,12625022 | 0,002901613 |
| Adcyap1 | 1,29836 | 0 | 1,3546 | 0,326962 | 0 | 0 | 0 | 0 | 0 | 0 | Adcyap1 | -4,03316202 | 0,002971724 |
| Rinl | 1,8329 | 2,03416 | 0,669334 | 1,26948 | 0,259789 | 0,172371 | 0,351321 | 0,208271 | 0,345599 | 0,148413 | Rinl | -2,04845698 | 0,002989063 |
| Ptprn2 | 0,136087 | 0,402749 | 1,02235 | 1,23402 | 0,0963692 | 0,0681633 | 0,0625443 | 0,0619211 | 0,0454825 | 0,0219993 | Ptprn2 | -2,5797057 | 0,003007281 |
| Tmem30b | 1,48898 | 0,489624 | 0,579999 | 1,60011 | 0,225056 | 0,0497654 | 0,121733 | 0,45134 | 0,332419 | 0,353451 | Tmem30b | -2,50013313 | 0,003015399 |
| Vwa5b2 | 0,141171 | 1,50957 | 0,176758 | 0,142236 | 0,0399896 | 0 | 0 | 0 | 0,0235947 | 0,0228234 | Vwa5b2 | -2,95832139 | 0,003024636 |
| Nmnat2 | 0,0706513 | 0,766672 | 0,884611 | 1,06776 | 0,100067 | 0 | 0 | 0 | 0 | 0 | Nmnat2 | -3,28540762 | 0,003034534 |

| Gene | d9-11 (1) | d9-11 (2) | d9-11(3) | d9-11(4) | newborn(1) | newborn(2) | newborn(3) | adult(1) | adult(2) | adult(3) | Gene | log2(fold) | FDR(*p*-value) |
|---|---|---|---|---|---|---|---|---|---|---|---|---|---|
| Ugt8a | 0,269877 | 0,177488 | 0,0750893 | 0,634429 | 0 | 0 | 0 | 0 | 0 | 0 | Ugt8a | -3,99337886 | 0,00304198 |
| Klk13 | 0,74628 | 0 | 3,42529 | 3,00538 | 0 | 0 | 0 | 0 | 0 | 0 | Klk13 | -4,02386653 | 0,003044398 |
| Vgf | 1,30255 | 0 | 1,52209 | 1,44339 | 0,0738662 | 0,0653605 | 0,0399591 | 0 | 0,0437056 | 0,0844258 | Vgf | -3,22337806 | 0,0030567 |
| Zfp940 | 1,43353 | 1,06062 | 0,997121 | 0,481387 | 0,203199 | 0,179775 | 0,0366397 | 0 | 0,0400562 | 0,154795 | Zfp940 | -2,44557048 | 0,003098554 |
| Gdf2 | 0,439203 | 0,86654 | 0,45825 | 0,110618 | 0,0311231 | 0 | 0 | 0 | 0 | 0 | Gdf2 | -3,65007801 | 0,003099365 |
| Kcnk6 | 7,62149 | 4,1081 | 1,63963 | 2,84986 | 1,35786 | 0,0984183 | 0,337107 | 0,151905 | 0,238887 | 0,177894 | Kcnk6 | -2,39777948 | 0,003140236 |
| Pvrl4 | 0,189443 | 0,114207 | 0,55345 | 1,71774 | 0,0536844 | 0 | 0 | 0 | 0,031711 | 0 | Pvrl4 | -3,60500269 | 0,003141778 |
| Fgf17 | 0,207034 | 0,408466 | 1,38237 | 6,6728 | 0,117554 | 0,0520604 | 0,0636111 | 0 | 0 | 0 | Fgf17 | -3,33598149 | 0,003211384 |
| AI504432 | 0,285539 | 1,01407 | 0,667373 | 0,460321 | 0,129387 | 0,0142975 | 0 | 0 | 0 | 0 | AI504432 | -2,90906966 | 0,00321357 |
| Fosl1 | 5,25971 | 0 | 0 | 0 | 0 | 0 | 0 | 0 | 0 | 0 | Fosl1 | -3,84748083 | 0,00321596 |
| Ppbp | 0 | 4,17616 | 31,7973 | 43,6909 | 0,200797 | 0,178187 | 0 | 0 | 0 | 0 | Ppbp | -3,98420909 | 0,00321596 |
| Fibcd1 | 0 | 0,33096 | 0,784113 | 0,405625 | 0,0190057 | 0 | 0 | 0 | 0 | 0 | Fibcd1 | -3,75611214 | 0,003221575 |
| Cxcl14 | 0 | 0,933486 | 1,57961 | 7,81575 | 0,161144 | 0,0475654 | 0 | 0 | 0,637426 | 0,368837 | Cxcl14 | -3,60821349 | 0,003225695 |
| Cyp26b1 | 0,882196 | 9,43927 | 3,51201 | 3,41867 | 1,23022 | 0,560839 | 0,311883 | 0,617555 | 1,79173 | 0,943429 | Cyp26b1 | -2,13060108 | 0,003241412 |
| Ntng1 | 0,99402 | 0,478609 | 1,2873 | 0,190502 | 0,130744 | 0 | 0,0193083 | 0 | 0 | 0 | Ntng1 | -3,05935759 | 0,003268817 |
| Fgf3 | 0,934789 | 6,45484 | 0,260044 | 0 | 0 | 0 | 0 | 0 | 0 | 0 | Fgf3 | -4,01654868 | 0,003304622 |
| Cecr6 | 0,130791 | 0,258051 | 0,218349 | 0,131779 | 0 | 0 | 0 | 0 | 0 | 0 | Cecr6 | -3,94246975 | 0,003313965 |
| Hmx1 | 1,1788 | 0 | 1,37736 | 0,47488 | 0 | 0 | 0 | 0 | 0 | 0 | Hmx1 | -3,98629278 | 0,003315076 |
| Gm11110 | 0 | 6,68277 | 1,25634 | 2,27354 | 0 | 0 | 0 | 0 | 0 | 0 | Gm11110 | -3,57294404 | 0,003317869 |
| Hist1h2bk | 25,8817 | 9,61493 | 7,97407 | 17,9858 | 5,26239 | 3,36159 | 3,77936 | 5,33416 | 4,03257 | 6,63762 | Hist1h2bk | -3,35048682 | 0,003344024 |
| Krt79 | 0,809314 | 0,319349 | 2,9722 | 0,163046 | 0,0918382 | 0 | 0 | 0 | 0,0544209 | 0 | Krt79 | -3,57750487 | 0,003505397 |
| Mecom | 4,95547 | 8,44181 | 6,91746 | 5,812115 | 2,2582705 | 0,1899565 | 0,276652 | 0,065723 | 0,54329 | 0,362018 | Mecom | -2,10983599 | 0,003552696 |
| Wnt4 | 0,178113 | 4,78574 | 3,06345 | 4,73181 | 0,501998 | 0,0105698 | 0,0258587 | 0,0767488 | 0 | 0,0307064 | Wnt4 | -3,16764927 | 0,003593775 |
| Hist1h2bj | 31,7379 | 17,7164 | 6,5878 | 13,0432 | 5,50966 | 4,19713 | 4,25736 | 7,23838 | 6,92822 | 9,47094 | Hist1h2bj | -3,3968023 | 0,0036418 |
| Adcy4 | 14,8389 | 20,8717 | 11,1075 | 5,30479 | 4,04236 | 0,527895 | 0,440233 | 0,696517 | 2,17957 | 1,30137 | Adcy4 | -2,30800091 | 0,003665584 |
| AF357359 | 0 | 0 | 0 | 0 | 0 | 0 | 0 | 0 | 0 | 0 | AF357359 | -2,4482958 | 0,003665745 |
| Rsph4a | 0,177244 | 0,524552 | 0,591789 | 0,297621 | 0,0125557 | 0 | 0 | 0 | 0 | 0,028675 | Rsph4a | -3,2942742 | 0,003738092 |
| Prkch | 16,2832 | 21,515 | 3,54204 | 4,07634 | 3,3561 | 0,667898 | 0,877403 | 1,16665 | 1,75147 | 1,53329 | Prkch | -2,17829279 | 0,003785446 |
| Olig2 | 0,137063 | 0,405633 | 0,228807 | 0,828445 | 0 | 0 | 0 | 0 | 0 | 0 | Olig2 | -3,91318502 | 0,003837888 |
| Nol4 | 0,0632338 | 0,0902657 | 0,68609 | 0,254847 | 0 | 0 | 0 | 0 | 0 | 0 | Nol4 | -3,91838305 | 0,003858303 |
| Phox2a | 0 | 0 | 0 | 0 | 0 | 0 | 0 | 0 | 0 | 0 | Phox2a | -3,64214768 | 0,003861604 |
| Arhgef26 | 1,52474 | 4,33198 | 1,78184 | 2,51949 | 0,86369 | 0,13744 | 0,093421 | 0,111022 | 0,101873 | 0,23656 | Arhgef26 | -2,20988897 | 0,003912568 |
| Irf6 | 0,545076 | 1,53633 | 0,779972 | 3,29506 | 0,286784 | 0,136548 | 0,286356 | 0 | 0,026041 | 0,10074 | Irf6 | -2,21533203 | 0,003921321 |
| Edn3 | 0,110198 | 0,652259 | 1,65567 | 0,888148 | 0,0624721 | 0,0552641 | 0,0337927 | 2,30424 | 0,62779 | 1,17763 | Edn3 | -2,91720265 | 0,003930253 |
| Ppapdc1a | 1,04866 | 0 | 0 | 1,05584 | 0,0994704 | 0 | 0 | 0 | 0 | 0 | Ppapdc1a | -3,53055308 | 0,004013975 |
| Espn | 4,34584 | 2,91287 | 5,04962 | 14,6478 | 0,727837 | 0,632771 | 0,694289 | 1,4045 | 0,732934 | 1,13336 | Espn | -2,11344239 | 0,004078631 |
| Bmpr1b | 0,431163 | 0,546871 | 1,73895 | 1,42739 | 0 | 0,192944 | 0,0938544 | 0 | 0,102888 | 0,0198676 | Bmpr1b | -2,74519997 | 0,004081524 |
| Glis1 | 2,15145 | 1,67557 | 0,189031 | 0,798532 | 0,192589 | 0 | 0,0694515 | 0 | 0,151814 | 0,256712 | Glis1 | -2,9987103 | 0,004082772 |
| Zscan10 | 1,75879 | 0,960325 | 0 | 0,159816 | 0 | 0 | 0 | 0 | 0 | 0,0514725 | Zscan10 | -3,94139739 | 0,004119936 |
| Tubb1 | 0 | 1,3439 | 11,0864 | 24,186 | 0,0483166 | 0,0427749 | 0 | 0 | 0 | 0 | Tubb1 | -3,92507211 | 0,004229657 |
| Epha8 | 0,067665 | 0 | 1,12963 | 0,340878 | 0 | 0 | 0 | 0 | 0 | 0,0437532 | Epha8 | -3,92829876 | 0,004268299 |
| Dio3 | 0,368651 | 0,909162 | 0,461537 | 0,928315 | 0,0523107 | 0 | 0 | 0 | 0 | 0 | Dio3 | -3,51868008 | 0,004284177 |
| Wscd1 | 1,93109 | 1,78593 | 2,41778 | 5,10669 | 0,855415 | 0,21191 | 0,0740373 | 0,219396 | 0,566617 | 0,430098 | Wscd1 | -2,34471393 | 0,004319668 |
| Esrp2 | 2,47878 | 0,855849 | 0,724168 | 3,37116 | 0,281116 | 0,217635 | 0,190091 | 0,22528 | 0,457274 | 0,963752 | Esrp2 | -2,28269649 | 0,00432397 |
| Kcna3 | 0,868967 | 1,02866 | 0,580226 | 0,875296 | 0,14794 | 0 | 0 | 0 | 0 | 0 | Kcna3 | -3,17965081 | 0,004334271 |
| Mkx | 0,611765 | 0,402335 | 1,02128 | 0,513608 | 0,115589 | 0,0511207 | 0,0312614 | 0 | 0,170752 | 0,165039 | Mkx | -2,63108796 | 0,004399043 |
| Prph | 1,56872 | 2,51468 | 2,31497 | 0,197507 | 0,16698 | 0,147877 | 0,12047 | 0 | 0 | 0,254835 | Prph | -2,67355553 | 0,004428333 |
| Stk33 | 0,559087 | 0 | 0,699982 | 0,985611 | 0 | 0 | 0 | 0 | 0 | 0 | Stk33 | -3,88808811 | 0,004450218 |
| Lrmp | 1,88728 | 2,00499 | 2,3023 | 4,98296 | 0,741127 | 0,109317 | 0,255584 | 0 | 0 | 0 | Lrmp | -2,21929946 | 0,004465373 |
| Ermap | 0 | 1,45419 | 4,62885 | 1,7681 | 0,218723 | 0,0175807 | 0,0215082 | 0 | 0 | 0 | Ermap | -3,24040733 | 0,004489199 |
| Kng1 | 0 | 0,585216 | 0,365966 | 2,49615 | 0 | 0 | 0 | 0 | 0 | 0 | Kng1 | -3,89790188 | 0,004500635 |
| Grin2c | 3,38166 | 0,962312 | 0,922824 | 0,786278 | 0,294719 | 0,227993 | 0,358636 | 0,118369 | 0,195575 | 0,126139 | Grin2c | -2,03221161 | 0,004560651 |
| Mrc1 | 25,6445 | 10,6605 | 4,03232 | 0,660985 | 2,31403 | 0,47783 | 0,548092 | 0 | 0,119529 | 0,115654 | Mrc1 | -2,6178604 | 0,004580378 |
| Sim1 | 0 | 0,168168 | 0,533612 | 0,0858826 | 0 | 0 | 0 | 0 | 0 | 0 | Sim1 | -3,88775292 | 0,004608248 |
| Cldn2 | 0,320381 | 0,316053 | 0,0891402 | 0,968303 | 0,0302693 | 0 | 0 | 0 | 0 | 0,0345787 | Cldn2 | -3,57285819 | 0,004624662 |
| Notch4 | 12,6692 | 8,31652 | 4,92992 | 2,68747 | 2,83221 | 0,536448 | 0,481377 | 0,910643 | 1,90773 | 1,95415 | Notch4 | -2,06515171 | 0,004638793 |
| Gm5424 | 1,82661 | 7,99587 | 3,43015 | 10,6931 | 1,71838 | 0,517051 | 0,491281 | 0,931525 | 0,424015 | 0,817012 | Gm5424 | -2,15238006 | 0,004653503 |
| Lrrc73 | 1,8258 | 1,80107 | 0,653053 | 3,15206 | 0,444687 | 0 | 0,0802259 | 0 | 0,0882516 | 0 | Lrrc73 | -2,85555953 | 0,004668342 |
| Gjb5 | 0 | 0,807588 | 1,36656 | 2,06141 | 0,0581026 | 0 | 0 | 0 | 0,138001 | 0,066517 | Gjb5 | -3,63998278 | 0,004730415 |
| Tal2 | 0,985256 | 1,61983 | 0,548159 | 3,30684 | 0,0934235 | 0,0828774 | 0,101153 | 0 | 0 | 1,17969 | Tal2 | -3,09555563 | 0,004824874 |
| Cbfa2t3 | 7,81676 | 5,91974 | 1,96282 | 3,0103 | 1,75495 | 0,307239 | 0,202126 | 1,30101 | 2,54635 | 2,1484 | Cbfa2t3 | -2,15372869 | 0,004885668 |
| Col9a3 | 1,15733 | 3,42509 | 9,37033 | 3,9308 | 1,11547 | 0,0580491 | 0,177469 | 0 | 0 | 0 | Col9a3 | -2,61130313 | 0,004885668 |
| Nrxn1 | 0 | 0,375667 | 0,722443 | 0,313944 | 0,0489753 | 0,00864389 | 0,0211566 | 0 | 0,0345978 | 0,0223114 | Nrxn1 | -3,1135966 | 0,004941371 |
| Rasal1 | 1,00812 | 0,795602 | 0 | 0,101426 | 0 | 0 | 0 | 0 | 0 | 0 | Rasal1 | -3,87289344 | 0,004965375 |
| Il17b | 7,05609 | 0 | 10,1688 | 14,1989 | 0,733613 | 0 | 0,198813 | 0 | 0,222451 | 0 | Il17b | -3,31580419 | 0,004980005 |
| Bzrap1 | 0,246681 | 0,0405588 | 0,652064 | 0,414265 | 0,0349219 | 0 | 0 | 0 | 0 | 0 | Bzrap1 | -3,29776856 | 0,00498173 |
| Plcb1 | 0,628054 | 1,9915 | 1,28823 | 2,79902 | 0,642977 | 0,134729 | 0,0686938 | 0,326773 | 0,598851 | 0,413047 | Plcb1 | -2,0911702 | 0,005012377 |
| Cdk5r2 | 0 | 0,241381 | 0,102118 | 0,739501 | 0 | 0 | 0 | 0 | 0 | 0 | Cdk5r2 | -3,85206438 | 0,005014749 |
| Nxph4 | 0,417479 | 0,205915 | 1,21953 | 6,51749 | 0,118526 | 0 | 0,128274 | 0 | 0,0703919 | 0,0678514 | Nxph4 | -3,35541403 | 0,005016579 |
| Asphd2 | 1,55623 | 1,31586 | 3,89656 | 5,59776 | 0,568189 | 0 | 0 | 0,201587 | 0 | 0 | Asphd2 | -3,09388197 | 0,005146303 |
| Tlx2 | 0,500209 | 0 | 1,46116 | 0,503757 | 0 | 0 | 0 | 0 | 0 | 0 | Tlx2 | -3,81538736 | 0,005155211 |
| Dock6 | 34,2908 | 35,6732 | 25,9015 | 12,8057 | 11,3803 | 1,29642 | 1,4101 | 1,65925 | 2,80146 | 2,26211 | Dock6 | -2,00723682 | 0,005162995 |
| Mapk8ip2 | 0,091584 | 0,361389 | 4,73964 | 3,22944 | 0,259519 | 0,0229521 | 0 | 0,0832977 | 0,0306546 | 0,0296368 | Mapk8ip2 | -3,23783685 | 0,00517529 |
| Aim1l | 0 | 0,660463 | 0,223539 | 0,539643 | 0 | 0,0167665 | 0 | 0 | 0 | 0 | Aim1l | -3,61512122 | 0,005212433 |
| Ciart | 8,3331 | 10,9603 | 6,25387 | 11,7098 | 5,21274 | 0,390432 | 0,317889 | 8,42756 | 3,14688 | 15,652 | Ciart | -2,09009502 | 0,005252878 |
| Spock1 | 0,136684 | 0,205697 | 0,741355 | 0,413032 | 0,0295473 | 0,0171099 | 0,0209326 | 0 | 0,0228336 | 0 | Spock1 | -3,08657208 | 0,005343153 |
| Coro2a | 0,248693 | 2,94402 | 2,76783 | 0,668162 | 0,320887 | 0,0311549 | 0,0254069 | 0 | 0 | 0,026817 | Coro2a | -2,87194899 | 0,005354582 |
| Csmd2 | 0 | 0,0674945 | 0,0190372 | 0,183843 | 0,0129103 | 0,00570398 | 0 | 0 | 0 | 0 | Csmd2 | -2,92089029 | 0,005354822 |
| Casr | 0,422827 | 0,695199 | 0,235295 | 0,213008 | 0,0399246 | 0 | 0,043186 | 0 | 0 | 0 | Casr | -2,94449036 | 0,005363153 |
| Tnfrsf22 | 2,43964 | 1,89616 | 1,23412 | 2,68098 | 0,608076 | 0,16702 | 0,0680626 | 0,134303 | 0,273172 | 0,167824 | Tnfrsf22 | -2,54520188 | 0,005475902 |
| Snap25 | 0 | 1,26261 | 4,60702 | 1,12812 | 0,136159 | 0,0803508 | 0 | 0 | 0 | 0 | Snap25 | -3,24483646 | 0,005521198 |
| Gpr182 | 49,6587 | 27,8473 | 4,29943 | 3,11305 | 5,44027 | 0,77718 | 0,759699 | 0,560727 | 1,04211 | 0,60274 | Gpr182 | -2,59370532 | 0,005717367 |
| 2610035F20R | 0 | 0 | 0,272608 | 0,65782 | 0 | 0 | 0 | 0 | 0 | 0 | 2610035F20R | -3,80353409 | 0,005743011 |
| She | 10,8961 | 9,88046 | 2,61835 | 1,49709 | 2,05717 | 0,358196 | 0,337135 | 0,450852 | 0,845346 | 0,693551 | She | -2,2823515 | 0,005759332 |
| Cdx1 | 4,56599 | 0,979179 | 0,165692 | 0 | 0,0563526 | 0 | 0 | 0 | 0 | 0 | Cdx1 | -3,70207835 | 0,005792725 |
| 2700086A05R | 1,47856 | 0 | 0 | 0 | 0 | 0 | 0 | 0 | 0 | 0 | 2700086A05R | -3,7678223 | 0,005811415 |
| Ednrb | 2,67249 | 24,5427 | 9,51346 | 6,29349 | 3,59387 | 0,523925 | 0,760481 | 0,835835 | 1,3969 | 0,93524 | Ednrb | -2,19610884 | 0,005862708 |
| Esm1 | 1,61348 | 5,09332 | 0,13467 | 0,325055 | 0,18309 | 0 | 0,0495285 | 0,14651 | 0,108492 | 0,104703 | Esm1 | -3,28381105 | 0,005877653 |
| Foxc1 | 3,87044 | 7,23864 | 7,33641 | 4,63069 | 1,87329 | 0,0808059 | 0,0741362 | 0,146715 | 0,458504 | 0,156495 | Foxc1 | -2,49362309 | 0,005939794 |
| Wnt8b | 0,385244 | 0,19002 | 0,080391 | 1,35843 | 0 | 0 | 0,0295258 | 0 | 0,0322465 | 0 | Wnt8b | -3,37103493 | 0,005992569 |
| Gm6213 | 2,17138 | 4,28361 | 0,603865 | 0 | 0 | 0 | 0 | 0 | 0 | 0 | Gm6213 | -3,76521414 | 0,006045262 |
| Sult5a1 | 0,968524 | 2,54772 | 4,31082 | 5,20117 | 0,6428 | 0,24437 | 0 | 0 | 0 | 0 | Sult5a1 | -2,6267548 | 0,006046892 |
| Plek | 3,27633 | 4,23251 | 9,24624 | 11,7892 | 2,65199 | 0,351756 | 0,262965 | 0 | 0,052176 | 0,0756913 | Plek | -2,18121542 | 0,006121574 |
| Id3 | 265,625 | 194,677 | 149,154 | 202,009 | 91,8134 | 7,4317 | 11,7441 | 12,3254 | 21,2068 | 28,0594 | Id3 | -2,01751701 | 0,006172701 |
| Rltpr | 0 | 2,75622 | 0 | 0 | 0 | 0 | 0 | 0 | 0 | 0 | Rltpr | -2,50508495 | 0,006236654 |
| 2610528J11Ri | 4,36298 | 0,430372 | 0 | 2,63515 | 0 | 0 | 0 | 0 | 0 | 0 | 2610528J11Ri | -3,78997375 | 0,00625081 |
| Vstm2b | 0,737578 | 0,485061 | 1,23116 | 1,98085 | 0,139688 | 0 | 0,075599 | 0 | 0 | 0 | Vstm2b | -3,07035754 | 0,006411645 |
| Rell1 | 12,178 | 10,6236 | 6,88869 | 5,17121 | 3,90774 | 0,655968 | 0,401143 | 0,549117 | 0,53931 | 0,619023 | Rell1 | -2,01121725 | 0,00643425 |
| Fezf1 | 0 | 0,143556 | 0,850243 | 3,66482 | 0 | 0 | 0,0446566 | 0 | 0 | 0 | Fezf1 | -3,64433853 | 0,006444109 |
| Dcaf12l2 | 0,228425 | 0,450677 | 1,23931 | 0,345183 | 0,0647518 | 0,0572837 | 0,0350263 | 0 | 0 | 0 | Dcaf12l2 | -2,7624825 | 0,006490187 |
| Scn2b | 0,420102 | 0,690718 | 1,63645 | 0,775994 | 0,257835 | 0,0175361 | 0,0643609 | 0,0637141 | 0,0468075 | 0 | Scn2b | -2,4215409 | 0,006513765 |
| Cacng5 | 0 | 0,43359 | 1,10063 | 0,0885639 | 0 | 0 | 0 | 0 | 0 | 0 | Cacng5 | -3,76671629 | 0,006600485 |
| Fabp7 | 0 | 1,6541 | 5,59678 | 21,9379 | 0,15961 | 0 | 0,518898 | 0 | 0 | 0 | Fabp7 | -3,40267173 | 0,006625021 |
| Kcne3 | 5,29363 | 4,89611 | 0,42431 | 1,53537 | 0,290161 | 0 | 0,157191 | 0 | 0,174834 | 0,167211 | Kcne3 | -3,13483886 | 0,006880768 |
| Crlf1 | 0 | 2,73125 | 0,888781 | 6,86427 | 0 | 0,160693 | 0 | 0 | 0,0718408 | 0,0692404 | Crlf1 | -3,42954842 | 0,007086596 |
| Spag16 | 0,451128 | 0,623051 | 0,602496 | 0,363606 | 0,0895915 | 0,0396516 | 0,0553138 | 0 | 0 | 0 | Spag16 | -2,46474419 | 0,007122616 |
| Stk10 | 3,38298 | 2,19398 | 3,32065 | 5,90184 | 1,47274 | 0,219619 | 0,0767713 | 0,513163 | 0,920953 | 0,830275 | Stk10 | -2,20302786 | 0,007152339 |
| Kel | 0 | 0,78338 | 4,1979 | 1,99994 | 0,112587 | 0 | 0 | 0 | 0 | 0 | Kel | -3,22851621 | 0,007213616 |
| Snord16a | 2611,14 | 2689,47 | 15874,1 | 0 | 0 | 0 | 0 | 0 | 0 | 0 | Snord16a | -3,0632031 | 0,00727665 |
| Prdm8 | 0,104843 | 0,620559 | 1,31268 | 1,79561 | 0,104004 | 0,078855 | 0,160734 | 0 | 0 | 0 | Prdm8 | -2,57445947 | 0,007289468 |
| Efcab1 | 0,334192 | 0,988982 | 1,95227 | 1,34597 | 0,0950748 | 0 | 0 | 0 | 0 | 0 | Efcab1 | -3,25427048 | 0,00729316 |
| Prex2 | 4,49204 | 4,03955 | 1,8458 | 1,50417 | 0,891508 | 0,0821468 | 0,0630451 | 0,350792 | 1,0124 | 0,607013 | Prex2 | -2,47924637 | 0,007358919 |
| Mcf2 | 0,227603 | 0 | 0,367619 | 0,458631 | 0 | 0 | 0 | 0 | 0 | 0 | Mcf2 | -3,70878289 | 0,007408942 |

| Gene | d9-11 (1) | d9-11 (2) | d9-11(3) | d9-11(4) | newborn(1) | newborn(2) | newborn(3) | adult(1) | adult(2) | adult(3) | Gene | log2(fold) | FDR(*p*-value) |
|---|---|---|---|---|---|---|---|---|---|---|---|---|---|
| Ooep | 0 | 3,03334 | 0,513255 | 1,23855 | 0 | 0 | 0 | 0 | 0 | 0 | Ooep | -3,71526969 | 0,00742508 |
| Slc22a18 | 0 | 2,26255 | 0,98689 | 2,1436 | 0,0671727 | 0,0595191 | 0,0727051 | 0 | 0,159785 | 0 | Slc22a18 | -3,1202338 | 0,007451063 |
| Lmo1 | 0,35652 | 3,51684 | 4,16534 | 9,33282 | 0,710248 | 0,270129 | 0 | 0,966798 | 0,849621 | 0,81583 | Lmo1 | -2,69624885 | 0,007489412 |
| Ppp1r16b | 5,82939 | 6,51121 | 1,57119 | 0,597788 | 1,02301 | 0,22894 | 0,17414 | 0,224901 | 0,550918 | 0,372711 | Ppp1r16b | -2,32792025 | 0,007509396 |
| Nkx6-2 | 0 | 0 | 0 | 0 | 0 | 0 | 0 | 0 | 0 | 0 | Nkx6-2 | -3,69950833 | 0,007510716 |
| Gm3716 | 1,48979 | 1,24357 | 0 | 2,42447 | 0,0974576 | 0,0287393 | 0,140582 | 0 | 0 | 0 | Gm3716 | -2,95128317 | 0,007546678 |
| Magee2 | 0,579872 | 0,429025 | 0,242001 | 1,02224 | 0,0411148 | 0 | 0 | 0 | 0 | 0 | Magee2 | -3,12659839 | 0,007742137 |
| Mir703 | 8122,88 | 14089,7 | 11640,5 | 12290,3 | 3705,52 | 2645,95 | 3540,93 | 0 | 0 | 0 | Mir703 | -2,07572274 | 0,007784945 |
| Sox2ot | 0 | 0,218052 | 0,276748 | 0,779388 | 0 | 0 | 0 | 0 | 0 | 0 | Sox2ot | -3,67900581 | 0,007844301 |
| 5730457N03R | 0,338192 | 0,111208 | 0,282286 | 0,227138 | 0 | 0 | 0 | 0 | 0 | 0 | 5730457N03R | -3,66205919 | 0,007860657 |
| Tex16 | 0,162042 | 0,719342 | 0,135258 | 0,163259 | 0 | 0,0202986 | 0 | 0 | 0 | 0 | Tex16 | -3,38267299 | 0,008222413 |
| Nap1l2 | 1,401 | 0,502572 | 0,425234 | 0,513225 | 0,108335 | 0,0639045 | 0,03907 | 0 | 0 | 0 | Nap1l2 | -2,63768822 | 0,008244054 |
| C1ql1 | 1,47918 | 0,583657 | 1,48137 | 1,48951 | 0,168221 | 0,074584 | 0,182116 | 0 | 0 | 0 | C1ql1 | -2,75363809 | 0,008326311 |
| Tceal3 | 1,61041 | 3,73455 | 3,67817 | 5,86009 | 2,68481 | 2,18237 | 2,01057 | 0 | 0 | 0 | Tceal3 | -2,9151393 | 0,008338451 |
| Cnr2 | 0,668098 | 2,88346 | 0,209126 | 0,588975 | 0,16563 | 0 | 0 | 0,0759706 | 0,111754 | 0,108066 | Cnr2 | -3,11330922 | 0,008363849 |
| Cdx4 | 1,10433 | 0,435752 | 0,184338 | 0,444896 | 0 | 0 | 0 | 0 | 0 | 0 | Cdx4 | -3,64279222 | 0,008415474 |
| Syt5 | 0 | 0,391671 | 0,497077 | 2,19951 | 0,0563526 | 0,0499073 | 0 | 0 | 0,0669025 | 0,0645048 | Syt5 | -3,21935403 | 0,008481944 |
| BC024139 | 0 | 1,11538 | 0,734017 | 1,01246 | 0 | 0 | 0 | 0 | 0 | 0 | BC024139 | -3,14813975 | 0,008503522 |
| Tmem145 | 0,584889 | 0,144246 | 2,56299 | 1,17838 | 0,165886 | 0 | 0 | 0 | 0 | 0 | Tmem145 | -3,1870205 | 0,008511844 |
| Gm10857 | 0,594268 | 3,86918 | 0,0990276 | 1,07762 | 0,202165 | 0,029757 | 0,0363845 | 0,107842 | 0,0597113 | 0 | Gm10857 | -2,82591613 | 0,008519198 |
| Zfp473 | 1,44748 | 0,624719 | 0,424192 | 0,18229 | 0,10254 | 0,0848887 | 0 | 0,0770474 | 0 | 0 | Zfp473 | -2,6969946 | 0,00854707 |
| Runx1 | 2,7658 | 2,47763 | 1,36086 | 1,46134 | 0,848364 | 0,0567151 | 0,0944562 | 0,257499 | 0,102926 | 0,0732028 | Runx1 | -2,16566652 | 0,008556862 |
| Cdh15 | 0,115866 | 0,2286 | 1,93423 | 1,05053 | 0,0656908 | 0 | 0 | 0 | 0 | 0 | Cdh15 | -3,24712529 | 0,008565094 |
| Rnf32 | 2,44385 | 3,52959 | 3,16533 | 0,273451 | 0 | 0,0684096 | 0,206195 | 0,245936 | 0 | 0 | Rnf32 | -2,37083065 | 0,008655096 |
| Flt1 | 34,7457 | 25,3921 | 6,07293 | 4,70163 | 3,76516 | 0,427011 | 0,414925 | 2,46666 | 5,24278 | 3,95547 | Flt1 | -2,73807402 | 0,008713548 |
| Frem1 | 6,02362 | 4,96546 | 5,53337 | 5,8622 | 2,02635 | 0,133959 | 0,0922161 | 0,0609563 | 0,0334798 | 0,0324158 | Frem1 | -2,25811708 | 0,008775974 |
| Asic4 | 0,875419 | 0,37011 | 0,835083 | 0,503943 | 0,141829 | 0,0627442 | 0 | 0 | 0 | 0 | Asic4 | -2,66016014 | 0,008986943 |
| Gareml | 0 | 0 | 0 | 0 | 0 | 0 | 0 | 0 | 0 | 0 | Gareml | -2,52726556 | 0,009064572 |
| Fezf2 | 0 | 0 | 0,97357 | 4,25934 | 0 | 0 | 0,0223712 | 0 | 0 | 0 | Fezf2 | -3,67927225 | 0,009092185 |
| Stx11 | 0,315499 | 1,29251 | 7,54236 | 5,99855 | 0,671237 | 0,474453 | 0,365495 | 0,143255 | 0,300067 | 0,153531 | Stx11 | -2,33414786 | 0,009202491 |
| Sema3g | 11,6644 | 8,82483 | 1,75694 | 3,00423 | 2,26138 | 0,219767 | 0,215056 | 0,55842 | 0,968487 | 0,851292 | Sema3g | -2,27574962 | 0,009237203 |
| Cend1 | 1,2681 | 4,16981 | 13,5826 | 17,4553 | 0,900101 | 1,27566 | 2,87817 | 0,575072 | 0,49899 | 0,412241 | Cend1 | -2,1042576 | 0,009294239 |
| Pou3f4 | 0,70072 | 1,03682 | 2,63146 | 1,76379 | 0,199402 | 0 | 0 | 0 | 0 | 0 | Pou3f4 | -3,05190849 | 0,00934519 |
| Rapgef5 | 8,47333 | 11,2248 | 3,30064 | 2,27653 | 2,33204 | 0,262829 | 0,247318 | 1,32148 | 2,05142 | 1,85334 | Rapgef5 | -2,10270911 | 0,009433327 |
| Amigo2 | 3,94401 | 6,03481 | 1,83972 | 2,56832 | 1,47998 | 0,200197 | 0,105591 | 0,418265 | 0,642166 | 0,413314 | Amigo2 | -2,22055204 | 0,009451893 |
| Trap1a | 0,636228 | 0,627604 | 0 | 7,68761 | 0,090477 | 0 | 0 | 0 | 0 | 0 | Trap1a | -3,52183354 | 0,009451893 |
| Tmem184a | 1,67018 | 0,494279 | 0,464068 | 1,51413 | 0,157795 | 0,0465747 | 0 | 0,168199 | 0 | 0,12038 | Tmem184a | -2,85705951 | 0,009521848 |
| Klhl29 | 2,27216 | 1,40643 | 1,59914 | 2,64856 | 0,655976 | 0,01115 | 0,0409323 | 0,0405658 | 0,0594717 | 0,0575655 | Klhl29 | -2,48370808 | 0,009522842 |
| Cpne7 | 3,05951 | 2,19503 | 0,116077 | 0,140094 | 0,236628 | 0,0349006 | 0 | 0 | 0 | 0 | Cpne7 | -2,94435953 | 0,009526458 |
| Camk4 | 0,100441 | 0,473136 | 0,345646 | 0,455443 | 0,0995078 | 0,00628077 | 0,015374 | 0 | 0 | 0 | Camk4 | -2,43384765 | 0,009576784 |
| Iqcg | 0 | 2,82465 | 0 | 0 | 0 | 0 | 0 | 0 | 0 | 0 | Iqcg | -2,38262911 | 0,009613536 |
| Kcnh1 | 0,0444415 | 0,306893 | 0,556455 | 0,313456 | 0,0377507 | 0 | 0 | 0 | 0 | 0 | Kcnh1 | -2,8768557 | 0,009615122 |
| Sigirr | 2,05963 | 2,37041 | 1,00279 | 2,93908 | 0,633092 | 0,0431146 | 0,0526975 | 0,779074 | 0,63517 | 0,66854 | Sigirr | -2,39266958 | 0,009808247 |
| Samsn1 | 14,5806 | 7,92931 | 0,31204 | 0,188286 | 0,689681 | 0,0939552 | 0,287056 | 0 | 0 | 0 | Samsn1 | -2,95486666 | 0,009831163 |
| **Hoxa4** | 0 | 1,14509 | 1,5501 | 2,80581 | 0,0659375 | 0 | 0 | 0,210463 | 0,0784075 | 0 | Hoxa4 | -3,3010647 | 0,009870662 |
| Tmem121 | 0,660199 | 3,6905 | 1,65304 | 1,77315 | 0,437432 | 0 | 0,0676329 | 0 | 0 | 0 | Tmem121 | -2,71649164 | 0,010029169 |
| Nrg2 | 0,815095 | 0,459476 | 1,3607 | 1,05576 | 0,132037 | 0,175218 | 0,142847 | 0,529226 | 0,35133 | 0,301733 | Nrg2 | -2,00752384 | 0,010151635 |
| Tmem169 | 0,234229 | 0,693195 | 0,782034 | 0 | 0 | 0,0293722 | 0 | 0 | 0 | 0 | Tmem169 | -3,37669511 | 0,01017954 |
| Wnt10b | 0 | 0 | 0,12636 | 2,28751 | 0 | 0 | 0 | 0 | 0,101731 | 0 | Wnt10b | -3,60811557 | 0,010339264 |
| Kcnb2 | 0,0597437 | 0,294687 | 0,0498698 | 0,42137 | 0,0169206 | 0 | 0 | 0 | 0 | 0,0193098 | Kcnb2 | -3,30391055 | 0,010350669 |
| 4933402E13R | 0,887735 | 0,175146 | 0 | 1,60954 | 0 | 0 | 0 | 0 | 0 | 0 | 4933402E13R | -3,59736586 | 0,010356735 |
| Fsip1 | 0,209026 | 0,412396 | 0,697834 | 1,47371 | 0,0593447 | 0 | 0 | 0 | 0 | 0 | Fsip1 | -3,29389318 | 0,010542173 |
| Usp51 | 0,899385 | 0,443614 | 1,00091 | 1,96297 | 0,170065 | 0,22578 | 0,184012 | 0 | 0 | 0 | Usp51 | -2,10680114 | 0,010691891 |
| Pou3f1 | 2,83131 | 1,39653 | 2,45417 | 2,85239 | 0,92599 | 0,0546091 | 0,233748 | 0 | 0 | 0,0705248 | Pou3f1 | -2,11813158 | 0,010809704 |
| Crb3 | 0,972653 | 1,59911 | 1,89402 | 2,93812 | 0,276667 | 0,0818072 | 0,199701 | 0 | 0,330443 | 0 | Crb3 | -2,32300111 | 0,010809704 |
| Kcns3 | 0,344058 | 0,226273 | 0,0957272 | 0,462152 | 0,0325104 | 0 | 0 | 0 | 0 | 0 | Kcns3 | -3,2364911 | 0,010830274 |
| Rnd1 | 3,52406 | 6,04595 | 2,42985 | 3,39556 | 1,34729 | 0,234654 | 0,0470259 | 0,417469 | 0,257433 | 0,447291 | Rnd1 | -2,24528514 | 0,01084403 |
| Tfec | 5,45534 | 1,92198 | 1,46353 | 0 | 0,387117 | 0 | 0,179543 | 0 | 0 | 0 | Tfec | -2,90005311 | 0,010914652 |
| Mira | 0 | 1,03431 | 2,18735 | 0 | 0 | 0 | 0 | 0 | 0 | 0 | Mira | -3,54672382 | 0,010996698 |
| Cyp17a1 | 2,06131 | 1,47886 | 0,156404 | 0 | 0 | 0,047094 | 0 | 0 | 0 | 0 | Cyp17a1 | -3,43089869 | 0,011043243 |
| Pnliprp2 | 0,675846 | 0,222233 | 0,940118 | 0,226894 | 0 | 0 | 0 | 0 | 0 | 0 | Pnliprp2 | -3,54175427 | 0,011043679 |
| Aard | 8,61237 | 2,65491 | 1,12309 | 1,89723 | 0,917828 | 0,271198 | 0,248386 | 0 | 0,182223 | 0,0876864 | Aard | -2,37854607 | 0,011094534 |
| A730017C20R | 0 | 1,88456 | 2,23229 | 1,15454 | 0,150265 | 0,0480161 | 0,0586781 | 0 | 0 | 0 | A730017C20R | -2,85617993 | 0,011176364 |
| **Hoxa6** | 2,52855 | 0 | 1,58226 | 0,636035 | 0 | 0 | 0 | 0 | 0 | 0 | Hoxa6 | -3,54487623 | 0,011455321 |
| Apoa2 | 11,0142 | 14,8177 | 0 | 44,3114 | 0,575373 | 3,85988 | 1,24897 | 3,56876 | 3,90293 | 1,33904 | Apoa2 | -2,45575273 | 0,011485139 |
| Sh2b2 | 1,61903 | 0,912663 | 4,05417 | 2,33008 | 0,655657 | 0,0580049 | 0,0354668 | 0 | 0 | 0 | Sh2b2 | -2,43410854 | 0,011528149 |
| Ksr2 | 0,411231 | 0 | 0,30036 | 0,155381 | 0,0291132 | 0 | 0 | 0 | 0 | 0 | Ksr2 | -3,12166079 | 0,011619221 |
| Rmst | 0 | 0 | 0,102821 | 1,48918 | 0 | 0 | 0 | 0 | 0,0413152 | 0,119734 | Rmst | -3,55751338 | 0,011715737 |
| Chn2 | 1,24906 | 6,03765 | 2,23725 | 1,40485 | 1,01644 | 0,223542 | 0,311246 | 0,113501 | 0,0837885 | 0 | Chn2 | -2,09841498 | 0,011772428 |
| Ovol2 | 0,242527 | 0,301505 | 0,510161 | 1,84664 | 0 | 0 | 0 | 0 | 0 | 0 | Ovol2 | -3,52035833 | 0,011831997 |
| Fam167a | 0,0757164 | 2,09144 | 1,70645 | 1,83087 | 0,343199 | 0,0569001 | 0,0464052 | 0 | 0,050636 | 0 | Fam167a | -2,49468841 | 0,011911894 |
| Snph | 0,421947 | 0,13875 | 0,234805 | 0,212564 | 0,0398415 | 0 | 0 | 0,127987 | 0 | 0 | Snph | -3,06792797 | 0,012043767 |
| Fhad1 | 0,141896 | 0,909872 | 1,95431 | 0,78631 | 0,140683 | 0,142158 | 0,0869571 | 0 | 0 | 0 | Fhad1 | -2,11012142 | 0,0120456 |
| Ackr1 | 1,61764 | 0,957426 | 0,539998 | 0,977292 | 0,18405 | 0 | 0 | 0 | 0 | 0 | Ackr1 | -3,02387675 | 0,012053167 |
| Serpina1b | 0 | 0,360408 | 1,14609 | 4,98448 | 0 | 0 | 0 | 0 | 0 | 0 | Serpina1b | -3,53991512 | 0,012055574 |
| Hist1h3f | 7,6097 | 2,76543 | 1,12723 | 5,02149 | 0,85675 | 0,156571 | 0,291779 | 0,657919 | 0,574198 | 0,456606 | Hist1h3f | -2,76293883 | 0,012265814 |
| Sez6 | 0,0773729 | 0 | 0,581259 | 0,389774 | 0 | 0 | 0 | 0 | 0 | 0 | Sez6 | -3,52552096 | 0,012290268 |
| Susd5 | 0,166385 | 0,328277 | 0,277767 | 0,167634 | 0,0235708 | 0 | 0 | 0 | 0,027831 | 0 | Susd5 | -3,19256538 | 0,012339333 |
| Gm2694 | 1,31054 | 0 | 1,64014 | 1,31856 | 0 | 0,166944 | 0 | 1,17648 | 0,909758 | 0,650004 | Gm2694 | -3,33791908 | 0,012400023 |
| Ptpre | 4,77348 | 3,00455 | 0,596462 | 0,590136 | 0,663517 | 0,190636 | 0,0717687 | 1,27949 | 1,09555 | 2,00654 | Ptpre | -2,34182948 | 0,012658075 |
| Habp2 | 0 | 0 | 0 | 1,66038 | 0 | 0 | 0 | 0 | 0 | 0 | Habp2 | -3,18296528 | 0,012666888 |
| Zfp804a | 0 | 0,151348 | 1,21658 | 0,772866 | 0,0651948 | 0 | 0 | 0 | 0 | 0,0248093 | Zfp804a | -3,27822126 | 0,012679702 |
| Tmeff2 | 0,0974526 | 1,53818 | 0,244032 | 0,392723 | 0,11047 | 0 | 0 | 0 | 0,0652622 | 0,0315429 | Tmeff2 | -3,13001091 | 0,012713137 |
| Clec1b | 13,0975 | 6,45674 | 7,86122 | 6,75489 | 2,01363 | 0 | 0 | 0 | 0 | 0 | Clec1b | -2,77907941 | 0,012737973 |
| Nodal | 2,91819 | 3,03866 | 0,135316 | 0,326614 | 0,137978 | 0 | 0,0497668 | 0 | 0,109018 | 0,0526043 | Nodal | -2,97845615 | 0,012880868 |
| Ngfr | 0,573327 | 4,14762 | 12,4425 | 9,33809 | 1,92259 | 0,239499 | 0,0878804 | 0 | 0,095975 | 0,0309263 | Ngfr | -2,43930038 | 0,013132336 |
| Adrb2 | 4,89356 | 2,04799 | 1,48527 | 0,746895 | 0,630888 | 0,297797 | 0,0910164 | 0,269388 | 0,0996285 | 0,240454 | Adrb2 | -2,21349309 | 0,013158882 |
| A830082K12R | 0,72866 | 0 | 0 | 0 | 0 | 0,0731598 | 0 | 0 | 0 | 0 | A830082K12R | -2,74971302 | 0,013297363 |
| Plekhg4 | 1,23232 | 0,303921 | 0,12858 | 0,155199 | 0,0218198 | 0,077176 | 0,0236028 | 0 | 0 | 0 | Plekhg4 | -2,65582415 | 0,013390567 |
| Dmrt2 | 0,143531 | 0,283183 | 0,119801 | 0,867525 | 0 | 0,0360252 | 0 | 0 | 0,0482012 | 0 | Dmrt2 | -3,21235738 | 0,013668818 |
| Cass4 | 1,6945 | 0,0879477 | 0,595384 | 0,628793 | 0,151686 | 0,0223408 | 0,0273271 | 0 | 0,029836 | 0,028847 | Cass4 | -2,62784016 | 0,013724408 |
| Kcnab1 | 1,69581 | 0,590439 | 0,999175 | 0,232743 | 0,196473 | 0,0579386 | 0 | 0 | 0 | 0 | Kcnab1 | -2,57957826 | 0,014005567 |
| Grik4 | 0 | 0,0917563 | 0,621103 | 0,281125 | 0 | 0 | 0 | 0 | 0,0311355 | 0 | Grik4 | -3,46276837 | 0,014113294 |
| Crtac1 | 0 | 1,12607 | 1,48211 | 0,383314 | 0,0359609 | 0,0636371 | 0 | 0 | 0 | 0 | Crtac1 | -3,0320697 | 0,014271518 |
| Tmem130 | 0,120885 | 2,26578 | 2,92613 | 2,07028 | 0,411253 | 0,212248 | 0,14831 | 0,109872 | 0,0405375 | 0 | Tmem130 | -2,14743873 | 0,014284832 |
| Zbtbd6 | 1,04537 | 1,1398 | 1,05626 | 1,16401 | 0,748578 | 0,607023 | 0,320565 | 0,350139 | 0,276563 | 0,427607 | Zbtbd6 | -2,03528166 | 0,014305933 |
| Tmem54 | 0 | 1,72945 | 0,566861 | 5,30347 | 0 | 0 | 0 | 0 | 0 | 0 | Tmem54 | -3,33753304 | 0,014307076 |
| Myct1 | 3,80332 | 6,21414 | 3,17458 | 0,957884 | 1,38146 | 0,327908 | 0,145813 | 1,08031 | 2,11211 | 1,38605 | Myct1 | -2,04585695 | 0,014312305 |
| Tgm1 | 1,55973 | 0,355074 | 0,600868 | 2,05477 | 0,238096 | 0 | 0,0735981 | 0,109051 | 0,120694 | 0,0388674 | Tgm1 | -2,44988731 | 0,014330506 |
| Gabrb1 | 0 | 1,27903 | 0,61838 | 0,186567 | 0 | 0 | 0 | 0 | 0 | 0 | Gabrb1 | -3,45979599 | 0,014369367 |
| Gpr45 | 0,417561 | 0 | 0,278835 | 0,841392 | 0,0236615 | 0 | 0 | 0 | 0 | 0 | Gpr45 | -3,18130681 | 0,014472865 |
| Dcc | 0,0902819 | 0,267192 | 1,93431 | 2,42591 | 0,221496 | 0,158097 | 0,110563 | 0 | 0 | 0 | Dcc | -2,24851025 | 0,014508927 |
| Tnk1 | 0 | 0 | 0 | 0 | 0 | 0 | 0 | 0 | 0 | 0 | Tnk1 | -2,73616713 | 0,014512107 |
| Prokr1 | 0,871724 | 0,625421 | 0,330746 | 0,558904 | 0,179613 | 0 | 0,0242864 | 0 | 0,0265047 | 0,0512653 | Prokr1 | -2,49653705 | 0,014518438 |
| Kcnn4 | 0,350794 | 0,865125 | 2,7815 | 1,01053 | 0,248848 | 0,0440642 | 0,0538563 | 0 | 0 | 0,0569406 | Kcnn4 | -2,46953206 | 0,014699971 |
| Phldb3 | 0,147356 | 0,290728 | 2,09088 | 1,48439 | 0,167175 | 0 | 0,0904415 | 0 | 0 | 0,0477862 | Phldb3 | -2,73277937 | 0,014700847 |
| S100g | 2,31061 | 2,28038 | 0 | 20,9192 | 0 | 0 | 0 | 0 | 0 | 0 | S100g | -3,33332517 | 0,014737712 |
| Igfbp1 | 1,16793 | 0 | 0,58486 | 0,705759 | 0 | 0 | 0 | 0 | 0 | 0 | Igfbp1 | -3,43824162 | 0,014920867 |
| Ctla2b | 11,3767 | 12,3445 | 1,77577 | 4,60166 | 1,19222 | 0,650462 | 0,352885 | 0,592213 | 0,664883 | 0,318151 | Ctla2b | -2,09329176 | 0,015047165 |
| Vwa5b1 | 0 | 0,164905 | 0,139532 | 0,58946 | 0 | 0 | 0 | 0 | 0 | 0 | Vwa5b1 | -3,43445416 | 0,015141532 |
| Frmd7 | 0 | 0,338731 | 0,343939 | 0,415148 | 0,0389049 | 0,0515966 | 0,105207 | 0 | 0 | 0,0222034 | Frmd7 | -2,25075561 | 0,015245827 |
| Fendrr | 0,140677 | 0,416327 | 0,352257 | 0 | 0,039895 | 0 | 0 | 0 | 0 | 0,0456086 | Fendrr | -3,1736553 | 0,01543224 |
| Snai1 | 14,8953 | 12,9393 | 8,34976 | 21,7193 | 5,87128 | 0,111857 | 0,273312 | 0,403174 | 0,300145 | 0 | Snai1 | -2,34010166 | 0,015586253 |

| Gene | d9-11 (1) | d9-11 (2) | d9-11(3) | d9-11(4) | newborn(1) | newborn(2) | newborn(3) | adult(1) | adult(2) | adult(3) | Gene | log2(fold) | FDR(*p*-value) |
|---|---|---|---|---|---|---|---|---|---|---|---|---|---|
| Creb3l4 | 3,18938 | 3,37094 | 0,380271 | 3,90051 | 0,452888 | 0,401241 | 0,280098 | 0,206543 | 0 | 0,222316 | Creb3l4 | -2,08621362 | 0,015688429 |
| Serpini1 | 0,939053 | 0 | 2,61275 | 1,36652 | 0,236579 | 0,0523168 | 0,0959764 | 0,189741 | 0,559237 | 0,202685 | Serpini1 | -2,5960844 | 0,015958491 |
| Tdh | 0,182885 | 0 | 0 | 3,13161 | 0 | 0 | 0 | 0 | 0 | 0 | Tdh | -3,43834053 | 0,016167646 |
| Cdhr1 | 0,0744379 | 0,146866 | 1,30483 | 0,524988 | 0,0421746 | 0,0932286 | 0,0228102 | 0 | 0 | 0 | Cdhr1 | -2,54550621 | 0,016204386 |
| Faim2 | 0,0688059 | 0 | 0,402625 | 0,0693245 | 0 | 0 | 0 | 0 | 0 | 0 | Faim2 | -3,40606414 | 0,016327567 |
| Slc30a3 | 1,87635 | 0,336541 | 0,711862 | 0,343641 | 0,0967963 | 0,0428473 | 0,157114 | 0,61943 | 0 | 0,38756 | Slc30a3 | -2,59459036 | 0,016381925 |
| BC021767 | 2,03013 | 1,46781 | 1,43136 | 1,44041 | 0,17883 | 0,0506497 | 0,139318 | 0 | 0 | 0,0490773 | BC021767 | -2,82208812 | 0,016464867 |
| Otp | 0 | 0,311198 | 1,05312 | 2,54129 | 0 | 0 | 0 | 0 | 0 | 0 | Otp | -3,40353035 | 0,01649758 |
| Slc7a3 | 0,28253 | 2,22969 | 0,707459 | 2,8461 | 0,480746 | 0,141816 | 0,173385 | 0,128334 | 0 | 0,183201 | Slc7a3 | -2,11680204 | 0,016571125 |
| Rab36 | 0,506487 | 4,87152 | 2,8536 | 2,80629 | 0,82572 | 0,0635307 | 0,0388417 | 0 | 0 | 0 | Rab36 | -2,41794014 | 0,01676588 |
| Msx1os | 0 | 0 | 0 | 0 | 0 | 0 | 0 | 0 | 0 | 0 | Msx1os | -3,36816634 | 0,01682721 |
| Kcnip3 | 6,8361 | 5,24293 | 2,43565 | 0,952388 | 0,763521 | 0 | 0 | 0 | 0,0805001 | 0 | Kcnip3 | -2,78990135 | 0,017037721 |
| Pf4 | 1,64249 | 18,6318 | 93,1826 | 142,907 | 16,2441 | 2,10157 | 4,34124 | 0 | 0,287991 | 0,546202 | Pf4 | -2,4206283 | 0,017599128 |
| Ntrk1 | 1,02665 | 0,506389 | 2,46366 | 0 | 0,145546 | 0 | 0 | 0 | 0,0861099 | 0 | Ntrk1 | -2,9035099 | 0,01793686 |
| Atg9b | 0 | 0 | 0 | 0,914982 | 0 | 0 | 0,207371 | 0,150212 | 0 | 0,293782 | Atg9b | -2,08276126 | 0,018266083 |
| Egr2 | 21,7011 | 4,58742 | 7,39337 | 7,36188 | 4,45655 | 0,305396 | 0,373482 | 0,100657 | 0,111314 | 0,0717107 | Egr2 | -2,13628826 | 0,018361949 |
| Krt13 | 0,832454 | 2,18981 | 0,463165 | 0 | 0 | 0 | 0 | 0 | 0 | 0 | Krt13 | -3,1729181 | 0,018387525 |
| Vwce | 1,4649 | 1,17416 | 0,84065 | 0,276726 | 0,155664 | 0,0458903 | 0,0561311 | 0 | 0 | 0,0296278 | Vwce | -2,43261842 | 0,018466371 |
| Lipf | 0 | 1,58593 | 0,223628 | 1,3492 | 0 | 0 | 0 | 0 | 0 | 0 | Lipf | -3,35402725 | 0,018484098 |
| Foxd2 | 0,126674 | 0,124963 | 1,48026 | 1,14851 | 0,0718317 | 0,0635573 | 0 | 0,0575594 | 0,0849893 | 0,0205235 | Foxd2 | -2,82568221 | 0,018491675 |
| 1300017J02Ri | 0 | 2,04202 | 2,71505 | 1,63838 | 0,209681 | 0,0742313 | 0 | 0 | 0 | 0 | 1300017J02Ri | -2,72729982 | 0,018525628 |
| Fam189a1 | 0,0670534 | 0,0661483 | 0,391797 | 1,41875 | 0,0569793 | 0,0167924 | 0,0410887 | 0 | 0,0448179 | 0 | Fam189a1 | -2,70605436 | 0,018622016 |
| Atoh1 | 0,162468 | 0,160271 | 0,54242 | 1,63656 | 0,0460913 | 0 | 0,0498735 | 0 | 0 | 0 | Atoh1 | -2,96948289 | 0,018677584 |
| Doc2b | 0,0755284 | 0,372544 | 1,00872 | 1,06536 | 0,0855865 | 0 | 0 | 0 | 0,0757645 | 0 | Doc2b | -2,86413415 | 0,018725611 |
| Gna15 | 2,26839 | 1,03279 | 0,291278 | 0,351524 | 0,148536 | 0 | 0,0535773 | 0 | 0,0587141 | 0 | Gna15 | -2,72879548 | 0,01880942 |
| Slc39a5 | 1,76517 | 2,13667 | 0 | 5,81787 | 0,0512211 | 0 | 0 | 0 | 0 | 0 | Slc39a5 | -2,32321529 | 0,018982601 |
| Cyp26a1 | 12,2588 | 5,56661 | 3,08573 | 3,52785 | 1,9331 | 0,146736 | 0 | 0 | 0 | 0 | Cyp26a1 | -2,46474723 | 0,019053465 |
| Ccdc151 | 2,61066 | 0 | 0 | 0,618778 | 0 | 0 | 0,0471331 | 0 | 0,0516048 | 0 | Ccdc151 | -3,11317801 | 0,019111877 |
| Hmx3 | 0,264687 | 0 | 0,883616 | 1,86588 | 0 | 0 | 0 | 0 | 0 | 0 | Hmx3 | -3,33868386 | 0,019161877 |
| Nmur2 | 0,397841 | 0 | 0,0830195 | 0,300609 | 0 | 0 | 0 | 0 | 0 | 0 | Nmur2 | -3,33137882 | 0,019186657 |
| Plet1 | 4,10995 | 0,368578 | 0,467773 | 0,376341 | 0 | 0 | 0,0573758 | 0 | 0 | 0 | Plet1 | -3,18090999 | 0,019207495 |
| Ly6g6f | 0 | 0 | 3,58289 | 2,52154 | 0 | 0 | 0 | 0 | 0 | 0 | Ly6g6f | -3,35422906 | 0,019322537 |
| Gpr50 | 0,768505 | 0,189527 | 0,962133 | 1,35462 | 0,109063 | 0 | 0 | 0 | 0 | 0 | Gpr50 | -2,87184916 | 0,019348671 |
| Got1l1 | 0 | 0,790876 | 0,446077 | 1,07652 | 0,0759457 | 0 | 0 | 0 | 0 | 0 | Got1l1 | -3,15978202 | 0,019429388 |
| Rspo4 | 0 | 0,139356 | 0,471639 | 0,996136 | 0 | 0 | 0 | 0 | 0 | 0 | Rspo4 | -3,33022284 | 0,019513747 |
| Ephb6 | 2,06205 | 0,932343 | 3,44355 | 4,59043 | 1,16852 | 0,107643 | 0,131672 | 0,15631 | 0,28748 | 0,0277974 | Ephb6 | -2,08661414 | 0,019576512 |
| Fam167b | 3,35099 | 2,47912 | 2,09725 | 0,421669 | 0,238663 | 0,21197 | 0 | 0 | 0 | 0 | Fam167b | -2,59796991 | 0,01967238 |
| Ppp4r4 | 0,175556 | 0,60615 | 0,659422 | 1,32654 | 0,0994874 | 0,0879833 | 0 | 0,159686 | 0,117495 | 0,170406 | Ppp4r4 | -2,40490618 | 0,019782435 |
| Zic1 | 0,0485552 | 2,07564 | 8,51112 | 15,9473 | 0,0550406 | 0 | 0 | 0 | 0 | 0 | Zic1 | -3,34601372 | 0,019866378 |
| Sema4f | 0,483347 | 0,238408 | 0,857482 | 0 | 0,0342574 | 0,030309 | 0 | 0 | 0,0528571 | 0,0511183 | Sema4f | -3,02512994 | 0,019943563 |
| Dmrta1 | 0,631185 | 0 | 0,543324 | 1,03339 | 0,0894099 | 0,0494138 | 0,0402986 | 0 | 0 | 0 | Dmrta1 | -2,61687364 | 0,020051062 |
| Fam65c | 5,70906 | 1,87731 | 3,64406 | 1,24052 | 1,39618 | 0,0561416 | 0,102986 | 0 | 0 | 0,0362529 | Fam65c | -2,17497963 | 0,020219662 |
| Tex26 | 3,47161 | 0,342453 | 0,579432 | 1,7477 | 0,0987853 | 0 | 0,106966 | 0 | 0 | 0 | Tex26 | -2,82591484 | 0,020219662 |
| 2700069I18Ri | 0 | 0,531822 | 3,59897 | 1,62778 | 0 | 0 | 0 | 0 | 0 | 0 | 2700069I18Ri | -3,31416333 | 0,020219662 |
| Kcnmb2 | 0 | 0,111084 | 0,845918 | 0,340328 | 0 | 0 | 0 | 0 | 0 | 0 | Kcnmb2 | -3,31416333 | 0,020219662 |
| Amer3 | 0,220507 | 0,290041 | 0,184062 | 0 | 0 | 0 | 0 | 0 | 0 | 0 | Amer3 | -3,30532202 | 0,020256502 |
| 4933430I17Ri | 6,02441 | 0,191707 | 0,810997 | 1,56593 | 0 | 0,293094 | 0,298471 | 0,176329 | 0,196423 | 0 | 4933430I17Ri | -2,56353475 | 0,02027978 |
| Unc5d | 0,236516 | 0 | 0,253839 | 0,170229 | 0 | 0,0169069 | 0 | 0 | 0 | 0 | Unc5d | -3,04097023 | 0,020333821 |
| 6030468B19R | 0,353618 | 3,1394 | 1,18041 | 0 | 0 | 0 | 0 | 0 | 0 | 0 | 6030468B19R | -3,17538008 | 0,020396624 |
| Sh3gl3 | 0,450005 | 5,11136 | 2,25628 | 2,12182 | 0,383861 | 0,510112 | 0,161566 | 0 | 0,0888714 | 0,146337 | Sh3gl3 | -2,05046172 | 0,020496256 |
| Ly6h | 7,59129 | 2,34413 | 2,95007 | 6,27972 | 1,4 | 0 | 0 | 0 | 0 | 0 | Ly6h | -2,419798 | 0,020632395 |
| Hist1h3c | 43,4927 | 61,2988 | 14,1717 | 22,2972 | 9,96195 | 0,660645 | 0,238559 | 0,513214 | 0,275208 | 0,357979 | Hist1h3c | -2,32456353 | 0,020711219 |
| Cd226 | 0 | 0,155456 | 0,78318 | 0,63496 | 0 | 0 | 0 | 0 | 0 | 0 | Cd226 | -3,2987446 | 0,020832078 |
| Syndig1l | 0,533353 | 0 | 0,111294 | 0,805936 | 0 | 0 | 0 | 0 | 0 | 0 | Syndig1l | -3,30066879 | 0,020839617 |
| Gprin1 | 0,697776 | 0,458903 | 0,841309 | 1,17171 | 0,153752 | 0 | 0 | 0 | 0,0518552 | 0,0501513 | Gprin1 | -2,60285714 | 0,021276092 |
| Aatk | 0,429251 | 1,21602 | 2,6102 | 1,12234 | 0,52141 | 0,0614176 | 0,0563581 | 0 | 0,215406 | 0,158567 | Aatk | -2,06508701 | 0,021544151 |
| Trim46 | 0,607563 | 0,119871 | 1,13648 | 1,58273 | 0,207792 | 0,085086 | 0,0963407 | 0,110441 | 0,0350856 | 0,0393666 | Trim46 | -2,23073418 | 0,02155832 |
| Cd79a | 1,09693 | 0,270518 | 0 | 0,552322 | 0,0779403 | 0 | 0 | 0 | 0 | 0,178728 | Cd79a | -2,98438349 | 0,021597642 |
| Prdm13 | 0 | 0 | 0,772954 | 1,34757 | 0 | 0 | 0 | 0 | 0 | 0 | Prdm13 | -3,28185193 | 0,022596428 |
| 5330434G04R | 0,18827 | 0,417887 | 0,3143 | 0 | 0 | 0 | 0 | 0 | 0 | 0 | 5330434G04R | -3,25712868 | 0,022630554 |
| Gli1 | 8,83133 | 6,2727 | 4,57038 | 7,20699 | 2,91193 | 0,0885218 | 0,13535 | 0 | 0 | 0 | Gli1 | -2,13480238 | 0,022938594 |
| Mapk13 | 0,259937 | 5,64116 | 0,21694 | 3,14129 | 0,29544 | 0 | 0,0799502 | 0 | 0 | 0,253977 | Mapk13 | -2,80602837 | 0,023018787 |
| Actl6b | 0 | 0 | 2,2902 | 2,07273 | 0 | 0 | 0 | 0 | 0 | 0 | Actl6b | -3,26090378 | 0,023636658 |
| Lbx1 | 0 | 0 | 3,53522 | 0,473816 | 0 | 0 | 0 | 0 | 0 | 0 | Lbx1 | -3,24585501 | 0,02369394 |
| Fli1 | 33,4998 | 24,7332 | 6,10566 | 5,3401 | 8,17287 | 0,877109 | 0,812654 | 1,15665 | 2,88867 | 2,29975 | Fli1 | -2,00281542 | 0,023988229 |
| AU015836 | 0,109327 | 0,754952 | 0 | 0,330422 | 0 | 0 | 0 | 0 | 0 | 0 | AU015836 | -3,23333349 | 0,024287685 |
| Tmod2 | 2,01222 | 1,51982 | 2,04713 | 1,20388 | 0,676444 | 0,00784661 | 0,038505 | 0 | 0,08387 | 0,0101273 | Tmod2 | -2,22581075 | 0,024416353 |
| Cntn2 | 0,329424 | 0,429403 | 6,49251 | 2,8454 | 0,656168 | 0,318527 | 0,109949 | 0,148664 | 0,725752 | 0,474323 | Cntn2 | -2,10324067 | 0,024515092 |
| Gm3002 | 1,9071 | 5,10235 | 4,90187 | 5,36243 | 0,507978 | 0,21282 | 0,259339 | 0,00395715 | 0,0617928 | 0,0343338 | Gm3002 | -2,55862181 | 0,024522757 |
| Tbxas1 | 1,20964 | 0 | 2,01926 | 4,87382 | 0,196129 | 0,17364 | 0,212232 | 0 | 0 | 0 | Tbxas1 | -2,43981822 | 0,02464135 |
| Gfra3 | 2,22875 | 0,169124 | 1,71713 | 0,690768 | 0,243222 | 0 | 0 | 0 | 0,0576769 | 0,0556484 | Gfra3 | -2,77931778 | 0,024645691 |
| Barx2 | 0,386695 | 0 | 0,806874 | 1,75272 | 0 | 0 | 0,0593895 | 0,175436 | 0 | 0 | Barx2 | -3,06534752 | 0,02466443 |
| Grap | 17,0675 | 11,514 | 2,75706 | 0 | 1,68846 | 0,332393 | 0,203046 | 0,599098 | 1,59788 | 1,14586 | Grap | -2,55168749 | 0,02471159 |
| Prss50 | 0,491718 | 0,727591 | 0,820776 | 1,98085 | 0 | 0 | 0 | 0 | 0 | 0 | Prss50 | -2,74993196 | 0,024816879 |
| Gli3 | 11,154 | 10,8207 | 10,1149 | 12,3965 | 3,25227 | 0,101993 | 0,0794276 | 0,0337466 | 0,0247211 | 0,0119667 | Gli3 | -2,51291067 | 0,024851645 |
| P2rx2 | 0,212532 | 0,209656 | 0 | 0 | 0 | 0 | 0 | 0 | 0 | 0 | P2rx2 | -2,98791831 | 0,025536708 |
| Gulo | 0 | 2,052 | 1,48817 | 0,149671 | 0 | 0,0372977 | 0,0455974 | 0 | 0 | 0 | Gulo | -2,89011015 | 0,0257567 |
| AF529169 | 0 | 1,04312 | 0,315224 | 0,152194 | 0,0427933 | 0 | 0 | 0 | 0,0252548 | 0 | AF529169 | -2,97426118 | 0,025952315 |
| Pla2g4c | 0,578232 | 1,14084 | 0,160656 | 0 | 0 | 0,0482436 | 0 | 0 | 0 | 0 | Pla2g4c | -2,82577463 | 0,02606007 |
| Mat1a | 1,2099 | 0,183625 | 0,543798 | 0,562594 | 0,105493 | 0,0933012 | 0,0855905 | 0,169291 | 0 | 0 | Mat1a | -2,06339196 | 0,026064934 |
| Gng13 | 0 | 0 | 0 | 0 | 0 | 0 | 0 | 0 | 0 | 0 | Gng13 | -3,20341344 | 0,026329065 |
| Klb | 0,188854 | 0,745216 | 0 | 0,761068 | 0,0267588 | 0,0236666 | 0,0289472 | 0 | 0,031612 | 0 | Klb | -2,75001118 | 0,026407712 |
| Aldob | 0,701183 | 0,172925 | 0 | 1,76572 | 0 | 0,0880773 | 0 | 2,86423 | 7,37347 | 1,42269 | Aldob | -2,79106023 | 0,026444369 |
| Nkx3-2 | 0,249489 | 0,246111 | 0 | 0,502519 | 0 | 0 | 0 | 0 | 0 | 0,0812239 | Nkx3-2 | -3,17698908 | 0,026444369 |
| Zfp365 | 0,527128 | 1,63432 | 0,942864 | 1,06219 | 0,42666 | 0,0188632 | 0,0461521 | 0 | 0,0755383 | 0,0243534 | Zfp365 | -2,13859157 | 0,026576403 |
| Gm5741 | 15,3307 | 2,54422 | 0 | 0 | 0 | 0,673636 | 0,811553 | 0 | 0 | 0 | Gm5741 | -2,73733352 | 0,0266152 |
| Gp1bb | 0 | 0 | 0 | 10,5058 | 0,652624 | 0,365489 | 0 | 0,508254 | 0 | 0 | Gp1bb | -2,33275753 | 0,026644053 |
| Slc7a11 | 1,46076 | 1,88125 | 1,53031 | 2,0559 | 0,284348 | 0,191462 | 0,151211 | 0,0778964 | 0,0947653 | 0,0639545 | Slc7a11 | -2,15133078 | 0,026946013 |
| **Hoxb7** | 0,289364 | 0 | 0,482989 | 0,874165 | 0 | 0 | 0 | 0 | 0 | 0 | Hoxb7 | -3,16691381 | 0,027029276 |
| 9330182L06Ri | 0 | 1,22496 | 0,518248 | 1,52911 | 0,117245 | 0,0172771 | 0,0845477 | 0 | 0 | 0 | 9330182L06Ri | -3,31558422 | 0,027053996 |
| Npas3 | 0,161185 | 0,636039 | 1,03152 | 1,94888 | 0,380386 | 0,0403504 | 0,0658293 | 0 | 0 | 0 | Npas3 | -2,12504892 | 0,027738585 |
| Pax9 | 0,258188 | 0,509398 | 0,53876 | 0,130048 | 0,0366033 | 0 | 0,039602 | 0,117311 | 0,0433129 | 0 | Pax9 | -2,72395003 | 0,027770949 |
| **Hoxc11** | 0 | 1,29112 | 1,09223 | 1,31757 | 0 | 0 | 0 | 0 | 0 | 0 | Hoxc11 | -3,16345157 | 0,027805729 |
| Thsd7b | 0,100798 | 0,149156 | 2,56625 | 1,06641 | 0,228348 | 0,0378462 | 0,108055 | 0 | 0,0504758 | 0,0162836 | Thsd7b | -2,29322861 | 0,028077787 |
| Galnt9 | 0,226456 | 0,112248 | 0,380877 | 0,229261 | 0 | 0 | 0 | 0 | 0 | 0 | Galnt9 | -3,14792929 | 0,028152909 |
| Sox5 | 0,436444 | 1,94046 | 3,0697 | 2,4121 | 0,75717 | 0,0887577 | 0,0271528 | 0,040365 | 0,014794 | 0 | Sox5 | -2,06084252 | 0,028529104 |
| 1700007K13R | 1,03784 | 0,511863 | 0,865987 | 4,7003 | 0,296152 | 0 | 0 | 0 | 0 | 0,170703 | 1700007K13R | -2,84497812 | 0,028536975 |
| Tmem28 | 1,01545 | 1,0017 | 0 | 0,766981 | 0 | 0,063923 | 0 | 0 | 0 | 0,0826614 | Tmem28 | -2,79188489 | 0,028580027 |
| Itga4 | 0,284734 | 2,77772 | 8,72818 | 2,26339 | 0,510501 | 0,00791483 | 0,00968637 | 0 | 0,0210986 | 0,0153218 | Itga4 | -2,84243545 | 0,028848258 |
| Ctla2a | 31,447 | 14,6376 | 5,74872 | 7,2659 | 5,57134 | 0,773743 | 0,945172 | 1,47955 | 2,39694 | 1,58811 | Ctla2a | -2,00297978 | 0,029383348 |
| Rab37 | 0,461987 | 2,43062 | 1,02828 | 2,17175 | 0,436817 | 0,0386635 | 0 | 0 | 0,155249 | 0,0499521 | Rab37 | -2,39689245 | 0,029612919 |
| Sapcd1 | 4,50081 | 8,43282 | 3,33208 | 1,6962 | 0 | 0 | 0 | 0 | 0 | 0 | Sapcd1 | -2,46161271 | 0,029620475 |
| Mamdc2 | 0,584903 | 3,26969 | 1,95288 | 1,47318 | 0,635406 | 0,122174 | 0 | 0,0886515 | 0,163208 | 0,126212 | Mamdc2 | -2,22369907 | 0,0300882 |
| Mtnr1a | 0 | 0,466945 | 0,395063 | 0,238363 | 0 | 0 | 0 | 0 | 0 | 0 | Mtnr1a | -3,12668134 | 0,030139753 |
| Gm5523 | 32,6486 | 39,2613 | 46,9587 | 49,2879 | 8,61545 | 10,5583 | 10,1955 | 10,9033 | 11,7264 | 10,6851 | Gm5523 | -2,44041774 | 0,030140792 |
| A630089N07R | 1,82834 | 4,31301 | 1,19429 | 0,16015 | 0,180404 | 0,159689 | 0,0976026 | 0,288748 | 0,293941 | 0,309484 | A630089N07R | -2,00981897 | 0,030845127 |
| Stmn4 | 0 | 1,18343 | 21,2757 | 7,24827 | 0,341109 | 0,453736 | 1,01555 | 0 | 0,305202 | 0,0978316 | Stmn4 | -2,48311804 | 0,030951303 |
| Cmah | 0,0336236 | 0,359387 | 0,279768 | 0,429472 | 0,0713911 | 0,0336491 | 0,0205899 | 0,0612458 | 0,0448524 | 0,0325701 | Cmah | -2,28890313 | 0,031213921 |
| Atp8b3 | 0,290153 | 0,357796 | 0,121098 | 0,0730849 | 0,0410971 | 0 | 0 | 0 | 0 | 0 | Atp8b3 | -2,77606833 | 0,031448112 |
| Fut9 | 0,0258376 | 0,0764672 | 0,021568 | 0,156212 | 0 | 0 | 0,0079098 | 0 | 0 | 0 | Fut9 | -2,89211591 | 0,031545438 |
| Nr5a2 | 1,86923 | 0,0878091 | 0,148597 | 0,896781 | 0,126109 | 0,0669168 | 0 | 0,0809624 | 0,0893657 | 0,0288013 | Nr5a2 | -2,49330241 | 0,031690602 |
| Pou2f2 | 0 | 0 | 2,45877 | 0,593841 | 0,113793 | 0 | 0 | 0 | 0 | 0 | Pou2f2 | -2,78265407 | 0,031830407 |
| Optc | 0 | 2,56607 | 1,08971 | 0,915559 | 0 | 0 | 0 | 0 | 0 | 0 | Optc | -2,82622044 | 0,0318612 |

| Gene | d9-11 (1) | d9-11 (2) | d9-11(3) | d9-11(4) | newborn(1) | newborn(2) | newborn(3) | adult(1) | adult(2) | adult(3) | Gene | log2(fold) | FDR(*p* -value) |
|---|---|---|---|---|---|---|---|---|---|---|---|---|---|
| Mog | 1,22552 | 0,20149 | 0,170476 | 0,205726 | 0 | 0 | 0,0627533 | 0 | 0 | 0,132764 | Mog | -2,6643392 | 0,032310897 |
| Olig1 | 0 | 0,153625 | 0,779892 | 0,470614 | 0 | 0 | 0 | 0 | 0,0523385 | 0,0505181 | Olig1 | -3,09720515 | 0,032355314 |
| Tek | 42,4227 | 58,858 | 13,6546 | 6,70043 | 9,62573 | 0,862007 | 0,989833 | 1,53372 | 3,76574 | 1,96631 | Tek | -2,30077173 | 0,032357318 |
| Enc1 | 54,9158 | 26,0595 | 22,7795 | 20,1139 | 11,119 | 0,538666 | 0,741397 | 0,611696 | 1,37028 | 0,73886 | Enc1 | -2,28586647 | 0,032548733 |
| Procr | 4,49336 | 9,09835 | 0,592134 | 2,85813 | 1,41063 | 0,178557 | 0,0727051 | 0 | 0,0798927 | 0,307817 | Procr | -2,18109803 | 0,032644161 |
| Pla2g2f | 0,41119 | 0 | 0,114403 | 0,690371 | 0 | 0 | 0 | 0 | 0 | 0 | Pla2g2f | -3,0877075 | 0,033069151 |
| Trpm1 | 0 | 0 | 1,87065 | 0,339198 | 0 | 0 | 0,0344175 | 0 | 0 | 0 | Trpm1 | -3,03110255 | 0,033153538 |
| Pdzk1ip1 | 1,34358 | 0 | 2,24234 | 2,51693 | 0 | 0,126853 | 0,15458 | 0 | 0 | 0 | Pdzk1ip1 | -2,76205583 | 0,033185192 |
| Arhgap20os | 3,12296 | 0 | 0 | 0 | 0 | 0 | 0 | 0 | 0 | 0 | Arhgap20os | -2,98743821 | 0,033419692 |
| 2410003L11Ri | 1,54283 | 3,80456 | 0 | 0,775977 | 0 | 0 | 0 | 0 | 0 | 0 | 2410003L11Ri | -3,06997695 | 0,034272671 |
| Cyp4x1 | 0,465742 | 1,14859 | 0 | 0 | 0 | 0 | 0 | 0 | 0 | 0 | Cyp4x1 | -3,06997695 | 0,034272671 |
| Plek2 | 0,684465 | 0,6752 | 1,33295 | 1,83829 | 0,0647957 | 0,287034 | 0,0701301 | 0 | 0 | 0,0742176 | Plek2 | -2,07354363 | 0,034276315 |
| Sstr2 | 0 | 0,533817 | 0,903262 | 0,272479 | 0 | 0 | 0 | 0 | 0 | 0 | Sstr2 | -3,06431151 | 0,034462782 |
| Barhl2 | 0 | 0 | 1,21116 | 1,16939 | 0 | 0 | 0 | 0 | 0 | 0,0235263 | Barhl2 | -3,0790055 | 0,034501386 |
| Syt9 | 0,760802 | 0,667137 | 0,493927 | 0,255504 | 0,19161 | 0 | 0,0259091 | 0 | 0,0282821 | 0 | Syt9 | -2,31287678 | 0,034694314 |
| Pcsk1 | 0,132352 | 0,130563 | 0,441885 | 0,133329 | 0 | 0 | 0 | 0 | 0,0444152 | 0 | Pcsk1 | -3,05276554 | 0,034694314 |
| Ncan | 0 | 0 | 1,97326 | 1,27917 | 0,0743705 | 0,0547781 | 0,0268126 | 0 | 0 | 0 | Ncan | -2,72810427 | 0,034788197 |
| Nkx2-2 | 0 | 0,16386 | 0,223431 | 2,67025 | 0 | 0 | 0 | 0 | 0 | 0 | Nkx2-2 | -3,06222376 | 0,0351477 |
| Upk3bl | 0 | 0,363617 | 0 | 2,22674 | 0 | 0 | 0 | 0 | 0 | 0 | Upk3bl | -3,06222376 | 0,0351477 |
| Foxd3 | 0,141134 | 0,278453 | 1,76701 | 3,12781 | 0,24015 | 0 | 0 | 0 | 0 | 0 | Foxd3 | -2,7042683 | 0,035389411 |
| Mycl | 3,89699 | 6,31573 | 7,3576 | 8,03926 | 0,815076 | 0 | 0 | 0 | 0 | 0 | Mycl | -2,85258061 | 0,035401372 |
| Gsc | 0,318114 | 0,313802 | 1,06193 | 1,28127 | 0,090477 | 0 | 0 | 0 | 0 | 0 | Gsc | -2,83645637 | 0,035631315 |
| Spry3 | 0,0736805 | 0,0726858 | 0,24601 | 0,296942 | 0,0208725 | 0 | 0 | 0 | 0 | 0 | Spry3 | -2,83645637 | 0,035631315 |
| Cbln4 | 0,117841 | 0 | 0,393441 | 0,474861 | 0 | 0 | 0 | 0 | 0 | 0 | Cbln4 | -3,04850234 | 0,035660888 |
| Sez6l | 0,1074 | 0 | 0,403425 | 0,703391 | 0,0608294 | 0,013443 | 0,0164486 | 0 | 0 | 0 | Sez6l | -2,55317278 | 0,035724937 |
| Rtn1 | 1,2311 | 4,63819 | 30,6687 | 11,8597 | 3,25238 | 0,317134 | 0,21039 | 0,225897 | 0,710337 | 0,168177 | Rtn1 | -2,1850605 | 0,035757794 |
| Nppc | 3,69065 | 1,09217 | 0,923971 | 0,371574 | 0,210112 | 0 | 0 | 0 | 0 | 0 | Nppc | -2,70449633 | 0,035982674 |
| Gpr123 | 0 | 0,19827 | 0,33553 | 0,3375 | 0,0189763 | 0,0167776 | 0 | 0 | 0 | 0,0216592 | Gpr123 | -2,71997444 | 0,036267403 |
| A930011O12R | 0 | 0,0785255 | 0,930208 | 0,240596 | 0 | 0,0199418 | 0 | 0 | 0 | 0 | A930011O12R | -2,92354284 | 0,036388931 |
| Pou5f1 | 0,803853 | 3,08088 | 0 | 0 | 0 | 0,0994677 | 0 | 0 | 0 | 0 | Pou5f1 | -2,9207028 | 0,036418272 |
| 5730460C07R | 1,31742 | 0,185658 | 0 | 0,189567 | 0 | 0 | 0 | 0 | 0 | 0 | 5730460C07R | -3,03637772 | 0,036995063 |
| Hnf1b | 0,269631 | 0 | 0,112527 | 0,814863 | 0 | 0 | 0 | 0 | 0 | 0 | Hnf1b | -3,03247431 | 0,037010332 |
| Gstt3 | 3,30783 | 0,543849 | 1,22705 | 1,48082 | 0,573675 | 0,0923572 | 0 | 0,500346 | 0,309362 | 0,23871 | Gstt3 | -2,15030875 | 0,037456555 |
| Rnf112 | 0 | 0,106738 | 0,54188 | 0,21801 | 0 | 0 | 0 | 0 | 0 | 0 | Rnf112 | -3,02436277 | 0,037522988 |
| Slc18a3 | 0 | 0,136627 | 0,80921 | 0,279039 | 0 | 0 | 0 | 0 | 0 | 0 | Slc18a3 | -3,02436277 | 0,037522988 |
| Hist1h4d | 37,8377 | 33,4353 | 11,1984 | 26,6802 | 9,82077 | 1,75171 | 0,617537 | 2,72523 | 1,59614 | 3,22137 | Hist1h4d | -2,05858981 | 0,037606602 |
| Fgd3 | 0 | 2,07497 | 0,614489 | 2,54287 | 0,178848 | 0 | 0 | 0 | 0 | 0 | Fgd3 | -2,73690297 | 0,037623741 |
| Gckr | 0,338685 | 0 | 0,706711 | 1,19404 | 0 | 0,0425356 | 0 | 0 | 0 | 0,109925 | Gckr | -2,43674886 | 0,037662453 |
| Dmrt3 | 0,139965 | 0,552292 | 0,467299 | 0 | 0 | 0 | 0 | 0 | 0 | 0 | Dmrt3 | -3,01917449 | 0,037765794 |
| Brinp1 | 0,581919 | 1,2438 | 0,890504 | 0,390846 | 0,302337 | 0 | 0,0297333 | 0 | 0,129896 | 0 | Brinp1 | -2,25718503 | 0,037852677 |
| Vsx2 | 0,102684 | 0 | 0,428549 | 0,310349 | 0 | 0 | 0 | 0 | 0 | 0 | Vsx2 | -3,01777586 | 0,037889634 |
| Calcr | 0,186997 | 0 | 0,234131 | 0,94198 | 0,0264951 | 0 | 0,028662 | 0 | 0 | 0 | Calcr | -2,58065051 | 0,037934301 |
| Lhx3 | 0 | 0,155778 | 0,659019 | 0,159069 | 0 | 0 | 0 | 0 | 0,0530796 | 0 | Lhx3 | -3,01740729 | 0,038003204 |
| Tie1 | 80,0881 | 65,8935 | 21,7749 | 12,388 | 16,3122 | 2,0191 | 1,68053 | 4,68578 | 8,94929 | 7,31021 | Tie1 | -2,13434909 | 0,038103728 |
| 2610528A11R | 3,95643 | 3,34507 | 0 | 2,27509 | 0,161405 | 0,574509 | 0,524748 | 0 | 0 | 0 | 2610528A11R | -2,09182843 | 0,038119733 |
| Epo | 0 | 1,20695 | 0,510462 | 0 | 0 | 0 | 0 | 0 | 0 | 0 | Epo | -2,99977907 | 0,038421915 |
| Wdr52 | 0,430073 | 0,141423 | 0,119666 | 0 | 0,0135313 | 0 | 0 | 0 | 0 | 0 | Wdr52 | -2,795105 | 0,038504002 |
| Pof1b | 0,16566 | 0,49027 | 0 | 0,083452 | 0 | 0 | 0 | 0 | 0 | 0 | Pof1b | -3,00757722 | 0,038821472 |
| Ptprr | 1,59377 | 0,917136 | 1,10857 | 0 | 0,188302 | 0,0333249 | 0 | 0,120681 | 0,178287 | 0,301326 | Ptprr | -2,56054734 | 0,03902758 |
| Ccdc177 | 0,145493 | 0,143529 | 0,242892 | 0,146589 | 0 | 0,0182212 | 0,0222912 | 0 | 0,0486407 | 0 | Ccdc177 | -2,57406283 | 0,039047852 |
| Gucy1b2 | 0,123939 | 0,733585 | 0 | 0,249714 | 0 | 0 | 0 | 0 | 0,0415697 | 0 | Gucy1b2 | -3,00183417 | 0,039236575 |
| Serpina6 | 0 | 0 | 3,67272 | 0,73864 | 0 | 0 | 0 | 0 | 0 | 0 | Serpina6 | -3,01224597 | 0,039318856 |
| Hpse | 0,849577 | 0,314288 | 1,77285 | 0,64193 | 0,331099 | 0,0532512 | 0 | 0,0965556 | 0,0355789 | 0,0343847 | Hpse | -2,27504278 | 0,039388206 |
| Dleu7 | 0 | 0,538608 | 0,455684 | 0,824768 | 0 | 0 | 0 | 0 | 0 | 0 | Dleu7 | -2,98999617 | 0,03989866 |
| Scn3a | 0,0325583 | 0,192714 | 0,448435 | 0,492102 | 0,0952429 | 0,016291 | 0,00996857 | 0 | 0,0488569 | 0,0126147 | Scn3a | -2,1813821 | 0,039961512 |
| Sct | 0 | 0 | 0,863702 | 21,3577 | 1,45281 | 0 | 0 | 0 | 0 | 0 | Sct | -2,58027958 | 0,039963715 |
| Creb3l3 | 0,294569 | 0,145294 | 0,122934 | 3,1157 | 0 | 0,0369712 | 0,135596 | 0 | 0 | 0 | Creb3l3 | -2,67137639 | 0,040026508 |
| Kcng1 | 0,144968 | 0,143008 | 0,726002 | 2,62861 | 0,123345 | 0,145549 | 0,0889717 | 0 | 0 | 0 | Kcng1 | -2,23139885 | 0,040146775 |
| Ngef | 0,480294 | 0,118451 | 0,601343 | 1,20964 | 0,170203 | 0,0301169 | 0,0368282 | 0 | 0 | 0 | Ngef | -2,27092203 | 0,040368305 |
| Sdk2 | 5,23916 | 3,71189 | 11,3706 | 6,81465 | 1,86117 | 0,0596054 | 0,0291744 | 0 | 0,0476932 | 0 | Sdk2 | -2,41160991 | 0,040624272 |
| 1700003M02R | 1,21816 | 0 | 0,84726 | 2,04491 | 0 | 0 | 0,0623748 | 0 | 0 | 0 | 1700003M02R | -2,58600906 | 0,040917942 |
| Lhx8 | 0 | 0 | 0,432944 | 2,61246 | 0 | 0 | 0 | 0 | 0 | 0 | Lhx8 | -2,98912962 | 0,041165092 |
| Gata5os | 0 | 0 | 0 | 0 | 0 | 0 | 0 | 0 | 0 | 0 | Gata5os | -2,59091025 | 0,041236972 |
| L3mbtl1 | 0 | 0,119299 | 0,807528 | 0,852807 | 0,0342847 | 0 | 0,0370924 | 0 | 0 | 0 | L3mbtl1 | -2,40223477 | 0,041315491 |
| Col9a1 | 2,00723 | 9,17108 | 24,5142 | 9,47237 | 1,85645 | 0 | 0,097179 | 0 | 0 | 0 | Col9a1 | -2,68340704 | 0,041871619 |
| Pcdh10 | 0,379298 | 0,821841 | 1,6664 | 2,51807 | 0,38783 | 0 | 0,0696252 | 0 | 0,0253243 | 0,0244933 | Pcdh10 | -2,19029152 | 0,042051056 |
| Muc2 | 0 | 0 | 0,506536 | 0,470332 | 0,0264349 | 0,0116829 | 0,0142959 | 0 | 0,0155796 | 0 | Muc2 | -2,53284727 | 0,042102241 |
| Cnrip1 | 12,8927 | 10,9013 | 2,88225 | 2,08684 | 3,46548 | 0,231727 | 0,21231 | 0 | 0,388725 | 0,449386 | Cnrip1 | -2,02385492 | 0,042120422 |
| Cd40 | 5,07622 | 5,87839 | 2,57886 | 1,25818 | 1,60035 | 0,16656 | 0,0678301 | 0 | 0,210444 | 0,0717737 | Cd40 | -2,03008488 | 0,042507966 |
| Bmx | 0,654786 | 6,13644 | 1,36636 | 0,65966 | 0,52586 | 0 | 0,0334646 | 0 | 0,0731372 | 0,141354 | Bmx | -2,38089538 | 0,042871332 |
| **Hoxd13** | 0 | 0,264931 | 0,448322 | 0,270542 | 0 | 0 | 0 | 0 | 0 | 0 | Hoxd13 | -2,95276577 | 0,042932411 |
| Gja6 | 0,67624 | 0,500322 | 0,141106 | 0,170293 | 0,0479666 | 0 | 0 | 0,306972 | 0,341201 | 0,0548702 | Gja6 | -2,75762788 | 0,043239662 |
| Tcte2 | 1,53341 | 0 | 1,91966 | 0,257378 | 0,290455 | 0,128712 | 0,0785998 | 0 | 0,172885 | 0,166445 | Tcte2 | -2,16195831 | 0,043253156 |
| Aqp5 | 0 | 0 | 2,28505 | 1,14893 | 0 | 0 | 0 | 0 | 0 | 0 | Aqp5 | -2,95963518 | 0,043379086 |
| Dennd1c | 0,264129 | 0 | 1,10231 | 0,13304 | 0,112342 | 0 | 0,0405157 | 0 | 0 | 0,0428024 | Dennd1c | -2,12540789 | 0,043422979 |
| Mir3091 | 0 | 14334,5 | 0 | 0 | 0 | 0 | 0 | 0 | 0 | 0 | Mir3091 | -2,43834534 | 0,043762889 |
| Ticam2 | 0,199704 | 0,197007 | 0,0833466 | 1,40837 | 0,0848956 | 0,0250302 | 0 | 0 | 0 | 0 | Ticam2 | -2,54509547 | 0,044057987 |
| Pnma2 | 0,2122 | 0,139557 | 0,413298 | 0 | 0 | 0,0177159 | 0 | 0 | 0 | 0 | Pnma2 | -2,80187009 | 0,044152705 |
| Ubash3a | 0 | 0,109703 | 0,556936 | 0,224066 | 0 | 0 | 0 | 0 | 0 | 0 | Ubash3a | -2,93652011 | 0,044450829 |
| **Hoxd8** | 0 | 0,205633 | 1,38509 | 1,34074 | 0,118363 | 0 | 0 | 0,189076 | 0 | 0,0539498 | Hoxd8 | -2,76170377 | 0,045031214 |
| Lyve1 | 59,4541 | 27,5043 | 2,6566 | 0,769516 | 3,86234 | 0,159694 | 0,234322 | 0 | 0,726066 | 0,330033 | Lyve1 | -2,65959151 | 0,045215313 |
| Clca5 | 0,821366 | 0,32411 | 0 | 0 | 0 | 0 | 0 | 0 | 0 | 0,02657 | Clca5 | -2,9375981 | 0,045280716 |
| Ppp2r2c | 0,159534 | 0,15738 | 0,266331 | 0,482201 | 0 | 0,0399675 | 0 | 0,14514 | 0,0533587 | 0,103205 | Ppp2r2c | -2,34293839 | 0,045463768 |
| Gstm3 | 0,721148 | 0,609749 | 0 | 2,43782 | 0,60058 | 0,506667 | 0,3489 | 0,699194 | 1,04897 | 0,722753 | Gstm3 | -2,92404059 | 0,045619419 |
| Ncf2 | 3,32539 | 1,51406 | 0,960798 | 0,128846 | 0,417035 | 0,0962614 | 0,235408 | 0 | 0,042909 | 0,0828918 | Ncf2 | -2,05083747 | 0,045663156 |
| Spink10 | 0 | 0 | 0,276768 | 0 | 0 | 0 | 0 | 0 | 0 | 0 | Spink10 | -2,913233 | 0,045726131 |
| Apoc3 | 7,8565 | 0 | 0 | 5,04288 | 0 | 0 | 0 | 0 | 0 | 0 | Apoc3 | -2,93143514 | 0,045750237 |
| Fgf23 | 0,244884 | 0,241575 | 0,306602 | 0,616747 | 0,0694269 | 0,0307131 | 0 | 0 | 0 | 0 | Fgf23 | -2,24957982 | 0,04613531 |
| Slc6a15 | 0 | 1,17169 | 1,06767 | 0,276144 | 0,129447 | 0,0457935 | 0,0280064 | 0 | 0 | 0 | Slc6a15 | -2,30889559 | 0,046627204 |
| 4930426D05R | 0 | 1,92455 | 0,723691 | 0 | 0 | 0 | 0 | 0 | 0 | 0 | 4930426D05R | -2,91966693 | 0,046742641 |
| 4930562C15R | 0,252688 | 0 | 0 | 0 | 0,0238649 | 0 | 0 | 0 | 0 | 0 | 4930562C15R | -2,40961603 | 0,046921006 |
| Gucy2e | 0,0749805 | 0,147938 | 0,187769 | 0,528864 | 0,0636848 | 0,0187615 | 0 | 0 | 0 | 0 | Gucy2e | -2,1989375 | 0,046991442 |
| 4930556M19R | 0,341154 | 1,3049 | 1,28135 | 0,171821 | 0,145194 | 0 | 0,0523713 | 0,309715 | 0,172148 | 0,166097 | 4930556M19R | -2,38059294 | 0,047007285 |
| Creg2 | 0,108997 | 0,107526 | 0 | 0,164735 | 0,00347259 | 0,0136432 | 0 | 0 | 0 | 0,00587031 | Creg2 | -2,89686634 | 0,047894991 |
| Sytl5 | 0,0479308 | 0 | 0,120157 | 0,0482937 | 0 | 0 | 0 | 0,0436366 | 0 | 0 | Sytl5 | -2,89923635 | 0,048017333 |
| Dnajc6 | 0,0618905 | 0,427856 | 0,36203 | 0,749125 | 0,105291 | 0 | 0 | 0 | 0,0413662 | 0 | Dnajc6 | -2,46174508 | 0,048160818 |
| Barhl1 | 0 | 0,85105 | 1,52103 | 0,464297 | 0,0997468 | 0 | 0,0587719 | 0 | 0,0294505 | 0 | Barhl1 | -2,4334072 | 0,048354414 |
| Foxq1 | 0,352702 | 0 | 0,196264 | 1,53972 | 0,0666575 | 0,0294861 | 0 | 0,267163 | 0,0394167 | 0 | Foxq1 | -2,62036005 | 0,048428318 |
| Clgn | 0 | 0,145646 | 0,246464 | 0,743634 | 0 | 0 | 0 | 0 | 0 | 0 | Clgn | -2,89144471 | 0,048468159 |
| Btbd18 | 1,38166 | 0,272601 | 0,345985 | 0 | 0,130507 | 0,046169 | 0 | 0 | 0 | 0 | Btbd18 | -2,35073574 | 0,048684206 |
| Mapk15 | 0 | 0,876241 | 0,444827 | 0,178943 | 0 | 0 | 0 | 0 | 0 | 0 | Mapk15 | -2,59891468 | 0,048692447 |
| Dnajc22 | 0,733428 | 2,41166 | 0,20404 | 2,46213 | 0,0694494 | 0,184628 | 0,225515 | 0,221564 | 0 | 0 | Dnajc22 | -2,03013145 | 0,048795284 |
| Tmem200a | 0 | 0,0769941 | 0,781773 | 0,235905 | 0,0221113 | 0 | 0 | 0 | 0 | 0 | Tmem200a | -2,7824521 | 0,049224979 |
| Chrnb4 | 0,372277 | 0,459062 | 0,310742 | 0 | 0 | 0,0466506 | 0 | 0 | 0 | 0 | Chrnb4 | -2,66103166 | 0,049347154 |
| Kif19a | 0 | 0,0933258 | 1,10552 | 0,762488 | 0,107235 | 0,0237109 | 0,0290014 | 0 | 0 | 0 | Kif19a | -2,30073089 | 0,049444286 |
| Myod1 | 0 | 0 | 2,79291 | 0,374502 | 0 | 0 | 0 | 0,16871 | 0,0625994 | 0,0603754 | Myod1 | -2,88808707 | 0,049640824 |
| Gimap1 | 20,8926 | 4,21563 | 0,593979 | 1,19553 | 2,21497 | 0,238635 | 0,218787 | 0,64503 | 1,73787 | 1,07652 | Gimap1 | -2,20010348 | 0,050087136 |
| Hist1h4f | 30,0706 | 25,0035 | 8,79675 | 24,6637 | 5,15432 | 0,594497 | 0,159688 | 1,29519 | 0,155065 | 0,414115 | Hist1h4f | -2,22878027 | 0,050365767 |

**Overlapping CHD genes with CPC (*p* =0.0078318)**

PRKCZ
ARHGEF16
AJAP1
KCNAB2
HES3
ESPN
APITD1
SRM
FBXO2
DRAXIN
PDPN
FHAD1
EPHA2
PADI3
RCC2
IGSF21
PAX7
PLA2G2F
VWA5B1
ECE1
ALPL
WNT4
EPHA8
EPHB2
E2F2
ID3
GALE
CNR2
SH3BGRL3
UBXN11
AIM1L
LIN28A
RPS6KA1
KDF1
SMPDL3B
SESN2
PTPRU
FAM167B
LCK
MARCKSL1
TMEM54
CSMD2
GJB5
GJB3
DLGAP3
CLSPN
RSPO1
CDCA8
POU3F1

MYCL
MFSD2A
CAP1
ZFP69
RIMS3
ERMAP
SLC2A1
TIE1
MPL
SLC6A9
KIF2C
PLK3
RAD54L
DMBX1
CYP4X1
PDZK1IP1
TAL1
STIL
FOXD2
ELAVL4
DMRTA2
RAB3B
ORC1
LRP8
GLIS1
DAB1
L1TD1
FOXD3
DNAJC6
CTH
LHX8
SLC44A5
ST6GALNAC3
ST6GALNAC5
MCOLN3
DDAH1
BARHL2
HFM1
CDC7
CCDC18
BCAR3
TMEM56
SASS6
NTNG1
VAV3
PSRC1
MYBPHL
GSTM3
ALX3
KCNA3

PHGDH
CIART
CGN
TUFT1
CELF3
SPRR1A
LOR
SLC27A3
CREB3L4
NUP210L
TPM3
SHE
CKS1B
EFNA3
TRIM46
PKLR
FDPS
RAB25
MEX3A
NES
CRABP2
NTRK1
CADM3
ACKR1
TAGLN2
IGSF9
CASQ1
NHLH1
SLAMF1
PVRL4
APOA2
RGS4
RGS5
LMX1A
ILDR2
F5
ASTN1
SOAT1
CACNA1E
RGS16
RGS8
NPL
NMNAT2
NCF2
PTGS2
LHX9
NR5A2
KIF21B
LMOD1
ELF3

PTPN7
MYBPH
OPTC
ATP2B4
SNRPE
NFASC
CNTN2
NUAK2
RASSF5
DYRK3
IRF6
SYT14
KCNH1
DTL
BATF3
SLC30A10
SUSD4
WNT3A
RHOU
ACTA1
FAM89A
KCNK1
FMN2
GREM2
EXO1
SOX11
ID2
GRHL1
KCNF1
GREB1
MYCN
VSNL1
KCNS3
MATN3
APOB
KLHL29
ADCY3
GAREML
DPYSL5
CAD
SLC30A3
FNDC4
GCKR
GALNT14
EML4
ABCG5
ABCG8
SIX3
SIX2
EPCAM

NRXN1
CLHC1
OTX1
SLC1A4
CNRIP1
PLEK
PROKR1
SNRPG
ADD2
CYP26B1
EGR4
ACTG2
MTHFD2
SLC4A5
WDR54
TLX2
SEMA4F
LRRTM4
CTNNA2
TMSB10
KCNIP3
ADRA2B
DUSP2
AFF3
CHST10
CREG2
POU3F3
GPR45
ST6GAL2
BUB1
FBLN7
PAX8
EN1
BIN1
AMER3
TMEM163
MCM6
CXCR4
THSD7B
LYPD6
FMNL2
RPRM
KCNH7
SCN3A
LRP2
SP5
GAD1
DLX2
ITGA6
CDCA7

HOXD13
HOXD11
HOXD10
HOXD9
HOXD8
HOXD3
HOXD4
HOXD1
ITGA4
NEUROD1
FAM171B
CALCRL
STAT4
MYO1B
TMEFF2
SLC39A10
SATB2
TMEM237
FZD7
NRP2
MAP2
MYL1
SPAG16
TMEM169
IGFBP2
IGFBP5
VIL1
WNT6
CDK5R2
FEV
PTPRN
ASIC4
PAX3
SPHKAP
B3GNT7
NPPC
CHRNG
EFHD1
NGEF
SPP2
GBX2
CHL1
LRRN1
EDEM1
FANCD2
TMEM40
SGOL1
EOMES
OSBPL10
CMTM8

TRIM71
SUSD5
STAC
TRANK1
XYLB
ACVR2B
CCK
KIF15
LARS2
TDGF1
PRSS50
ELP6
CELSR3
BSN
TRAIP
CAMKV
SEMA3F
CACNA2D2
GRM2
SEMA3G
STAB1
TKT
CACNA2D3
WNT5A
FLNB
FEZF2
CADPS
KBTBD8
FRMD4B
EPHA6
NXPE3
ALCAM
TAGLN3
GRAMD1C
GAP43
IGSF11
UPK1B
ILDR1
CASR
SEMA5B
PDIA5
MUC13
GATA2
GP9
H1FX
PLXND1
TRH
TMEM108
EPHB1
FOXL2

CLSTN2
PLS1
ZIC4
ZIC1
ARHGEF26
MME
PLCH1
KCNAB1
TIPARP
SHOX2
LXN
B3GALNT1
SERPINI1
MECOM
LRRC34
SLC2A2
ECT2
KCNMB2
SOX2
B3GNT5
VWA5B2
CAMK2N2
POLR2H
KNG1
SST
HES1
GP5
IQCG
FGFR3
MSX1
CRMP1
PPP2R2C
NKX3-2
CD38
PROM1
LDB2
NCAPG
LGI2
SEL1L3
RELL1
KLB
UGDH
PHOX2B
GABRA4
GABRB1
KIT
KDR
UBA6
CSN3
AFP

PF4
PPBP
AREG
SOWAHB
ANXA3
PRDM8
PLAC8
HPSE
NKX6-1
MMRN1
GRID2
ATOH1
BMPR1B
UNC5C
EMCN
SLC39A8
CENPE
GSTCD
NPNT
DKK2
LEF1
RPL34
COL25A1
ELOVL6
PITX2
NEUROG2
TRAM1L1
PRSS12
PLK4
PCDH10
SLC7A11
MGARP
CLGN
TBC1D9
GYPA
HHIP
MAB21L2
FHDC1
FGB
FGA
FGG
PDGFC
GLRB
GRIA2
APELA
TLL1
SPOCK3
AADAT
WDR17
TENM3

CENPU
MTNR1A
TRIP13
TERT
LPCAT1
CTNND2
DNAH5
MARCH11
BASP1
CDH6
RAI14
SKP2
SLC1A3
DAB2
PLCXD3
FGF10
HCN1
EMB
ISL1
ITGA2
FST
ESM1
DEPDC1B
ZSWIM6
CENPH
OCLN
MAP1B
FOXD1
ENC1
HMGCR
IQGAP2
F2RL2
F2RL1
OTP
VCAN
HAPLN1
EDIL3
NR2F1
PCSK1
CAMK4
STARD4
TRIM36
TICAM2
CDO1
PRDM6
SOWAHA
TCF7
PITX1
NEUROG1
CXCL14

SPOCK1
GFRA3
CDC25C
REEP2
PSD2
NRG2
PCDHAC2
ARAP3
STK32A
JAKMIP2
ADRB2
PCYOX1L
IL17B
CDX1
SLC6A7
GPX3
NMUR2
GALNT10
HAND1
TENM2
WWC1
SLIT3
SPDL1
FAM196B
LCP2
KCNIP1
KCNMB1
TLX3
NPM1
STK10
MSX2
CDHR2
GPRIN1
SNCB
FGFR4
PRR7
DBN1
COL23A1
FLT4
FOXQ1
FOXF2
FOXC1
GMDS
TFAP2A
MAK
ELOVL2
EDN1
ID4
HIST1H3C
HIST1H4D

HIST1H2AE
HIST1H4F
HIST1H3F
HIST1H2BJ
HIST1H2BK
HIST1H2AI
MOG
ZFP57
TRIM10
POU5F1
APOM
LY6G6F
SAPCD1
HSPA1A
HSPA1B
NOTCH4
ITPR3
GRM4
HMGA1
PACSIN1
SCUBE3
TULP1
MAPK13
CPNE5
DNAH8
KCNK5
TREML1
TREML2
LRRC73
TMEM151B
RHAG
TFAP2B
MCM3
TRAM2
HMGCLL1
BMP5
COL9A1
OOEP
CD109
SH3BGRL2
ELOVL4
TTK
TPBG
CNR1
FUT9
MMS22L
POU3F2
PRDM13
SIM1
LIN28B

PRDM1
NR2E1
TUBE1
RSPH4A
VGLL2
FABP7
CENPW
RSPO3
SOGA3
TMEM200A
ARG1
VNN1
EYA4
MYB
MTFR2
SLC35D3
OLIG3
STX11
LRP11
MTHFD1L
MYCT1
ACAT2
PLG
AGPAT4
T
DLL1
CHST12
PAPOLB
ACTB
FSCN1
GRID2IP
TSPAN13
TWIST1
SP8
DNAH11
CDCA7L
RAPGEF5
HOXA1
HOXA2
HOXA3
HOXA4
HOXA5
HOXA6
HOXA7
HOXA9
HOXA10
EVX1
CHN2
PPP1R17
ELMO1

AMPH
GLI3
MYL7
GCK
ADCY1
IGFBP1
CLDN3
UPK3B
SEMA3C
SEMA3E
SEMA3A
DBF4
STEAP1
CALCR
PEG10
ASNS
BHLHA15
NPTX2
TMEM130
MCM7
CNPY4
GPC2
ACTL6B
EPO
ACHE
VGF
SH2B2
RELN
PRKAR2B
NRCAM
TFEC
TES
WNT2
PTPRZ1
AASS
FEZF1
TMEM229A
IRF5
TSPAN33
CPA2
PODXL
SLC13A4
TBXAS1
EPHB6
KEL
EPHA1
CNTNAP2
SSPO
GIMAP1
ATG9B

WDR86
EN2
SHH
RNF32
PTPRN2
TDRP
ANGPT2
FAM167A
MTMR7
FGF17
DMTN
EGR3
NKX3-1
NKX2-6
STC1
NEFM
NEFL
PNMA2
DPYSL2
STMN4
DUSP4
NRG1
DUSP26
UNC5D
RAB11FIP1
GOT1L1
IDO2
SPIDR
MCM4
EFCAB1
ST18
RPS20
CLVS1
PREX2
EYA1
KCNB2
CRISPLD1
ZFHX4
STMN2
HEY1
FABP5
E2F5
TMEM67
GEM
RAD54B
ESRP1
CCNE2
POP1
OSR2
PABPC1

RIMS2
AARD
DSCC1
HAS2
ATAD2
FAM84B
ADCY8
KCNQ3
COL22A1
ARC
LY6H
RHPN1
MAPK15
SCRT1
TONSL
FOXH1
RECQL4
DMRT3
DMRT2
RCL1
GLDC
LURAP1L
FREM1
HAUS6
DMRTA1
TEK
AQP3
CNTFR
GLIPR2
MELK
FRMPD1
MAMDC2
TLE1
RASEF
NTRK2
DIRAS2
FGD3
BARX1
HEMGN
CORO2A
SEC61B
NR4A3
ALDOB
GRIN3A
SMC2
TAL2
SUSD1
AMBP
TNC
BRINP1

CDK5RAP2
CRB2
LHX2
NR6A1
LMX1B
PKN3
IER5L
PRRX2
PTGES
PRDM12
FIBCD1
LAMC3
NTNG2
BARHL1
GFI1B
OLFM1
LHX3
NOTCH1
CACNA1B
KLF6
PFKFB3
PRKCQ
ITIH2
GATA3
MCM10
SUV39H2
GAD2
MKX
RET
GDF2
SLC18A3
DKK1
PHYHIPL
EGR2
DNA2
HKDC1
COL13A1
NODAL
UNC5B
OIT3
PLA2G12B
P4HA1
PLAU
MAT1A
TSPAN14
CDHR1
GRID1
LIPF
HHEX
CYP26C1

CYP26A1
RBP4
HELLS
SLIT1
ARHGAP19
SFRP5
CRTAC1
HPS1
NKX2-3
CPN1
WNT8B
SEMA4G
TLX1
LBX1
FGF8
PSD
TMEM180
CYP17A1
INA
SORCS3
DUSP5
ADRA2A
HABP2
VWA2
PNLIPRP2
EMX2
RGS10
PPAPDC1A
FGFR2
HMX3
ADAM12
PTPRE
TCERG1L
DPYSL4
NKX6-2
IFITM1
SCT
EPS8L2
CEND1
MUC2
BRSK2
IFITM10
TNNT3
IGF2
TH
TSPAN32
SLC22A18
PHLDA2
TRIM6
TAF10

SYT9
TUB
RIC3
LMO1
STK33
LYVE1
MRVI1
ZBED5
CALCA
MYOD1
E2F8
DBX1
PAX6
LMO2
SLC1A2
FJX1
LDLRAD3
PRR5L
ALX4
SYT13
MAPK8IP1
GYLTL1B
MDK
CHRM4
F2
NUP160
SLC43A1
BTBD18
VWCE
MYRF
FEN1
FADS2
GNG3
FERMT3
CDCA5
CFL1
FOSL1
TMEM151A
CD248
RIN1
NPAS4
ACTN3
SPTBN2
ANKRD13D
GAL
FGF3
ANO1
SHANK2
FOLR1
PHOX2A

KCNE3
MOGAT2
WNT11
MYO7A
PAK1
ALG8
TENM4
FAM181B
PRCP
PRSS23
RAB38
TRPC6
GUCY1A2
SLC35F2
PLET1
CADM1
APOA4
APOC3
APOA1
TAGLN
SCN2B
MPZL2
HYOU1
H2AFX
PVRL1
GRIK4
SORL1
UBASH3B
NRGN
ROBO3
PKNOX2
FEZ1
CHEK1
DDX25
FLI1
BARX2
ST14
B3GAT1
B4GALNT3
TEAD4
PRMT8
FGF23
RAD51AP1
KCNA1
CLEC1B
GPRC5A
GSG1
ST8SIA1
SOX5
BCAT1

LRMP
DDX11
CNTN1
NELL2
AMIGO2
COL2A1
CACNB3
RND1
WNT10B
TUBA1A
PRPH
DNAJC22
FAIM2
AQP5
BIN2
SLC4A8
SCN8A
FIGNL2
KRT7
KRT5
KRT79
KRT8
KRT18
SOAT2
ESPL1
HOXC11
HOXC10
HOXC4
HOXC9
HOXC8
HOXC6
HOXC5
HNRNPA1
NFE2
PDE1B
NEUROD4
GDF11
SLC39A5
TIMELESS
GLS2
PRIM1
GPR182
NXPH4
GLI1
KIF5A
LRIG3
FAM19A2
HMGA2
GRIP1
CPM

KCNMB4
PTPRR
TSPAN8
LGR5
NAP1L1
E2F7
SYT1
PAWR
SLC6A15
ALX1
NTS
PLXNC1
USP44
SNRPF
MYBPC1
ASCL1
STAB2
RFX4
WSCD2
RASAL1
LHX5
TBX3
KSR2
RFC5
CCDC60
CIT
MSI1
KNTC1
CCDC92
TMEM132C
FZD10
RIMBP2
RAN
MMP17
P2RX2
POLE
GJB2
AMER2
SHISA2
WASF3
RASL11A
CDX2
FLT1
TEX26
HSPH1
SMAD9
ENOX1
LACC1
SERP2
LCP1

DLEU7
ATP7B
PCDH8
DACH1
KLF5
SLAIN1
EDNRB
POU4F1
DOCK9
ZIC5
ZIC2
EFNB2
MYO16
SOX1
F10
PNP
SALL2
REM2
CDH24
TGM1
ADCY4
RIPK3
NOVA1
FOXG1
NPAS3
NKX2-1
PAX9
FOXA1
LRR1
POLE2
NID2
BMP4
WDHD1
DLGAP5
OTX2
DACT1
RTN1
SIX1
SIX4
TMEM30B
PRKCH
AKAP5
PLEK2
CCDC177
SMOC1
MAP3K9
VSX2
SYNDIG1L
NRXN3
STON2

KCNK10
CCDC88C
CHGA
FAM181A
PPP4R4
SERPINA6
GSC
CLMN
BDKRB2
DIO3
AMN
ASPG
KIF26A
SIVA1
CDCA4
ATP10A
APBA2
FAM189A1
TRPM1
CHRNA7
GREM1
LPCAT4
FSIP1
DISP2
RAD51
SPINT1
DLL4
CHAC1
OIP5
GATM
SCG3
TMOD2
RAB27A
PRTG
MNS1
FOXB1
IGDCC3
TIPIN
ZWILCH
SMAD6
KIF23
TLE3
HCN4
STRA6
ARID3B
ISL2
CRABP1
CHRNB4
ARNT2
AP3B2

BNC1
SH3GL3
HAPLN3
FANCI
TICRR
WDR93
ST8SIA2
NR2F2
ARHGDIG
SOX8
BAIAP3
HN1L
CCNF
KREMEN2
CLDN9
CLDN6
MMP25
ZSCAN10
NMRAL1
TVP23A
XYLT1
CCP110
CRYM
CDR2
PLK1
PRKCB
ATP2A1
LAT
QPRT
KIF22
MYLPF
FBXL19
STX1B
PRSS8
ORC6
CBLN1
NKD1
SALL1
CPNE2
DOK4
CDH8
CKLF
PLEKHG4
HSD11B2
RLTPR
CENPT
ESRP2
CDH1
PKD1L3
ADAMTS18

VAT1L
CMIP
NECAB2
ATP2C2
COTL1
GINS2
FOXF1
FOXC2
JPH3
CDH15
CPNE7
FANCA
SPIRE2
TUBB3
FAM57A
SERPINF2
ATP2A3
SPNS2
ARRB2
WSCD1
FAM64A
ALOX12
DLG4
CLDN7
ACAP1
TNK1
DNAH2
TMEM88
CCDC42
MYH3
HS3ST3B1
TRPV2
FAM83G
GRAP
RNF112
SEZ6
SLC6A4
ATAD5
RAB11FIP4
CDK5R1
TMEM132E
LHX1
DUSP14
HNF1B
SRCIN1
PNMT
GRB7
CDC6
TOP2A
IGFBP4

KRT13
KRT15
KRT19
RND2
BRCA1
ETV4
SOST
MPP2
PYY
SLC4A1
ITGA2B
C1QL1
WNT3
MYL4
ITGB3
SP6
SKAP1
HOXB1
HOXB2
HOXB3
HOXB4
HOXB5
HOXB6
HOXB7
HOXB8
HOXB9
IGF2BP1
GNGT2
ABI3
NGFR
DLX3
EME1
NOG
BZRAP1
PRR11
TBX2
TBX4
LIMD2
ICAM2
AXIN2
CACNG5
CACNG4
SSTR2
SDK2
RPL38
BTBD17
GPRC5C
RAB37
SLC9A3R1
GRIN2C

HID1
UNC13D
FOXJ1
13. Sep 09
RBFOX3
CCDC40
AATK
SLC16A3
COLEC12
TYMS
ADCYAP1
TGIF1
TMEM200C
ARHGAP28
LAMA1
MTCL1
PIEZO2
GREB1L
SNRPD1
TTR
NOL4
SLC14A1
RNF165
KATNAL2
SMAD7
LIPG
ME2
DCC
ST8SIA3
ONECUT2
CCBE1
PMAIP1
CD226
PPAP2C
SHC2
GNA15
TJP3
CREB3L3
FSD1
CHAF1A
UHRF1
CRB3
DENND1C
TUBB4A
CAMSAP3
OLFM2
ICAM1
AP1M2
DOCK6
EPOR

CCDC151
ELAVL3
CNN1
ACP5
KLF1
SYCE2
LYL1
PALM3
TPM4
F2RL3
PLVAP
CRLF1
TMEM59L
NCAN
HAPLN4
PBX4
LPAR2
VSTM2B
CCNE1
DPY19L3
RGS9BP
KCTD15
LGI4
LSR
HAUS5
ETV2
APLP1
SYNE4
DPF1
KCNK6
SPRED3
RYR1
MAP4K1
RINL
LRFN1
DLL3
SNRPA
TGFB1
CD79A
ATP1A3
POU2F2
TMEM145
PHLDB3
KCNN4
APOC2
KLC3
NOVA2
MEIS3
PLA2G4C
LIG1

LMTK3
RASIP1
PPFIA3
TEAD2
DKKL1
RPL13A
KCNC3
KLK1
KLK8
KLK13
IGLON5
SYT5
BRSK1
USP29
NRSN2
TRIB3
SCRT2
RSPO4
SNPH
EBF4
CPXM1
PROKR2
MCM8
LRRN4
FERMT1
BMP2
PLCB1
SNAP25
JAG1
ISM1
FLRT3
OVOL2
INSM1
NKX2-2
PAX1
FOXA2
NINL
ID1
COX4I2
TPX2
XKR7
DNMT3B
AHCY
GGT7
PROCR
SPAG4
SLA2
NNAT
SLC32A1
PPP1R16B

EMILIN3
L3MBTL1
MYBL2
TOX2
GDAP1L1
HNF4A
RIMS4
WFDC2
UBE2C
TNNC2
SLC12A5
CD40
SLC13A3
SNAI1
FAM65C
KCNG1
SALL4
DOK5
CBLN4
CASS4
TFAP2C
TUBB1
EDN3
CDH4
SLCO4A1
COL9A3
TCFL5
CHRNA4
STMN3
RGS19
OPRL1
MYT1
SAMSN1
HUNK
MIS18A
OLIG2
OLIG1
DONSON
SLC5A3
CLIC6
RUNX1
CHAF1B
SIM2
HMGN1
PCP4
DSCAM
TMPRSS2
UBASH3A
COL18A1
PCBP3

CECR6
SLC25A1
GP1BB
TBX1
SDF2L1
GNAZ
RAB36
BCR
DERL3
SGSM1
SEZ6L
ASPHD2
CHEK2
EMID1
NIPSNAP1
SEC14L2
MCM5
FOXRED2
PVALB
KCTD17
MFNG
GCAT
BAIAP2L2
MAFF
MGAT3
TTLL12
MPPED1
SULT4A1
PRR5
PHF21B
FBLN1
WNT7B
FAM19A5
ADM2
MAPK8IP2
BMX
TMEM27
S100G
NHS
MAP7D2
CNKSR2
KLHL34
PCYT1B
POLA1
ARX
SYTL5
MAOA
GATA1
PCSK1N
CCDC120

PRAF2
DGKK
WNK3
ALAS2
USP51
KLF8
ZC4H2
SLC7A3
IL2RG
NLGN3
ERCC6L
NAP1L2
CDX4
ZDHHC15
MAGEE2
POU3F4
RPS6KA6
HDX
POF1B
DACH2
KLHL4
TSPAN6
DRP2
BTK
GLA
BEX2
TCEAL3
ZCCHC18
ESX1
CLDN2
NUP62CL
IRS4
CAPN6
DCX
ALG13
TRPC5
ZCCHC16
LHFPL1
AMOT
LRCH2
DOCK11
DCAF12L2
SASH3
FRMD7
HS6ST2
GPC3
CCDC160
PLAC1
ZIC3
MCF2
